# Seasonal population structure and adaptive signatures across long-term marine microbial time series

**DOI:** 10.64898/2026.09.27.754832

**Authors:** Ting-Chun Huang, Francisco Latorre, Anders K. Krabberød, Sergio González-Motos, Vanessa Balagué, Josep M. Gasol, Pierre E. Galand, Ramiro Logares

## Abstract

Long-term microbial population dynamics can reveal how ocean microbiomes respond to environmental variation, but resolving within-species change requires suitable genomic references. We integrated 30 PacBio HiFi long-read metagenome-assembled genomes (MAGs) from the northwestern Mediterranean Sea with monthly Illumina metagenomes from neighboring time series spanning 12 and 7 years. Population differentiation varied widely among genomes and often differed in magnitude between sites, while a subset showed increasing differentiation with temporal distance. Genetic diversity and population structure were frequently seasonal, with populations associated with warm or cold waters and others spanning both thermal regimes. Candidate adaptive genes accounted for 1.6–3.0% of coding sequences, and many were detected in both time series. Their pN/pS trajectories included regular seasonal fluctuations consistent with changing strain or ecotype contributions, alongside irregular temporal patterns. These results show that combining long-read MAG references with dense short-read time series can resolve seasonal population structure, site-specific differentiation, and candidate adaptive variation within marine microbial species, providing a framework for tracking contemporary population-genomic change in the ocean.

## INTRODUCTION

Marine microbes play a crucial role in the functioning of the biosphere, underpinning global biogeochemical cycles^1^, with phytoplankton fixing atmospheric carbon on a scale comparable to terrestrial plants^2^. Part of this fixed carbon is transferred to higher trophic levels through microbial food webs, sustaining animal life and fisheries worldwide^3^. During the past two decades, major advances in marine microbial genomics have led to the discovery of thousands of species and millions of genes^4–6^. These studies have substantially improved our understanding of the structure of the ocean microbiome across spatiotemporal scales^7–14^, as well as the potential ecological interactions occurring within it^9,15,16^. Despite these advances, many questions remain unanswered regarding the structure and function of the ocean microbiome. In particular, we still lack a thorough understanding of the genetic composition of microbial populations and their dynamics^17^.

The genetic composition of populations is shaped by four key processes: mutation introduces new variants, selection alters allele or gene frequencies according to fitness, gene flow moves alleles and genes among populations, and genetic drift causes random changes in allele frequencies across generations^18,19^. Selection can favor genetic variants or accessory genes that enhance fitness, driving adaptation to specific environmental conditions or ecological niches and improving competitive abilities for particular resources^20^. Marine microbes experience substantial environmental variation in temperature and nutrient availability, while currents and geographic barriers influence connectivity across the ocean. Environmental gradients can be even steeper vertically, particularly those related to light and pressure^21^. Such heterogeneity has promoted the adaptive diversification of microbial populations^17^. Understanding population structure in relation to the environment can therefore provide insights into ecosystem function and how microbiomes respond to changing conditions^20^. This is particularly important under global change, as ocean warming can alter the genetic composition and structure of microbial populations, with potentially unpredictable consequences.

Culture-based and molecular studies have revealed extensive genomic diversity within microbial populations and associations between population differentiation and niche adaptation^17,20^. For example, hundreds of *Prochlorococcus* strains can coexist within small seawater samples^22,23^. These strains show substantial allelic variation in their core genomes and carry different sets of flexible genes, potentially reflecting distinct metabolic capabilities and ecological adaptations^23^. *Prochlorococcus* populations also differ between the Pacific and Atlantic Oceans, with distinct genomic backbones associated with each region^24^. Similarly, analyses of single amino-acid variants in SAR11 revealed distinct population structures associated with warm and cold oceanic currents^25^. Together, these studies demonstrate that substantial genomic and ecological variation can occur below the species level.

Tracking single nucleotide variants (SNVs) through time provides a complementary view of microbial population dynamics, revealing seasonal changes in ecotypes as well as processes such as immigration, local extinction and strain replacement^17^. A study of 30 bacterial MAGs from a nine-year freshwater metagenomic time series revealed substantial SNV heterogeneity within and between populations and patterns consistent with genome-wide and gene-specific selective sweeps^26^. More recent analyses of the same time series spanning 20 years identified both gradual and abrupt changes in strain composition, together with signatures of disturbance and resilience^27^. Long-term SNV analyses can therefore reveal contemporary population-genomic change occurring over years to decades^28^. A recent short-read metagenomic analysis spanning BBMO, SOLA and the global ocean further showed that marine microbial population differentiation varies strongly among taxa and across temporal and spatial scales, with geographic differentiation generally exceeding long-term temporal differentiation^29^.

However, the population-genomic resolution obtained from metagenomic time series also depends on the quality of the genomes used as references. Short-read MAGs can be fragmented and may provide incomplete representations of highly microdiverse populations and flexible genomic regions. Long-read sequencing offers an opportunity to improve this resolution by generating more contiguous genomes and recovering genomic regions that are difficult to reconstruct using short reads alone^30^. PacBio HiFi sequencing is particularly useful because it combines long reads with high sequence accuracy (>99.9%). Integrating long-read MAGs with dense short-read metagenomic time series therefore provides a means to link well-resolved genomic references with population variation occurring over annual-to-decadal timescales.

Here, we integrate 30 highly contiguous MAGs reconstructed from PacBio HiFi long reads from the LTER Blanes Bay Microbial Observatory (BBMO) in the Northwestern Mediterranean Sea with monthly Illumina metagenomes collected over 12 years at BBMO and seven years at the neighboring SOLA station in Banyuls Bay, France. We investigate the long-term population dynamics of these MAGs by tracking genome-wide SNV variation, nucleotide diversity, population differentiation and gene-level signatures of selection. We hypothesized that microbial populations would exhibit substantial genetic diversity and seasonal population structure, with some populations restricted to particular locations or periods, and that a subset of genes would show signatures of positive selection consistent with population differentiation. The investigated MAGs displayed substantial genetic diversity and varying degrees of population differentiation, frequently associated with seasonality. Some MAGs exhibited populations associated with cold or warm waters, whereas others included populations spanning both thermal regimes. Across the MAGs, ∼1.6%–3.0% of genes exhibited signatures of positive selection, and within individual MAGs, 0–100% of these genes were shared between the two time series. Temporal pN/pS patterns included regular seasonal fluctuations, consistent with recurrent strain or ecotype dynamics, and irregular changes suggesting intermittent strain occurrence or replacement. Together, this integrative approach provides a high-resolution view of long-term population structure and candidate adaptive variation in marine microbes.

## METHODS

### Sampling and DNA sequencing

Surface water samples (3m depth) were collected at the Long Term Ecological Research (LTER) Blanes Bay Microbial Observatory (BBMO) (http://bbmo.icm.csic.es) located in the Northwestern Mediterranean Sea at (41°40’N, 2°48’E; **Figure 1**)^31^. The LTER-BBMO, an oligotrophic coastal site approximately 1 kilometer offshore, is characterized by a depth of approximately 20 meters and minimal riverine or human influence. Similarly, surface water samples (3 m depth) were taken from the Banyuls Bay Microbial Observatory (SOLA; 42°31’N 3°11’E; **Figure 1**) in the Northwestern Mediterranean Sea ^32^. SOLA is also an oligotrophic coastal site with a depth of ∼26 m, located ∼1 km off the coast of Banyuls-sur-Mer, France, and with limited human impact^33^. Unlike BBMO, SOLA is affected by sporadic winter storms that bring nutrients from the sediments to the water column. Additionally, nutrient enrichment in SOLA is further enhanced by freshwater inputs from nearby rivers during flash floods^34^. Aside from these differences, both BBMO and SOLA exhibit similar environmental conditions (**Figure 1**). Approximately 150 kilometers separate BBMO and SOLA. The Northern Current, also known as the Liguro-Provençal-Catalan Current, mediates the connectivity between these two locations (**Figure 1**). This current flows along the Italian coast (west of Genoa), as well as the French and Catalan coasts, and can reach speeds of up to 1 m s^−1^ at the surface^35^.

**Figure 1.**
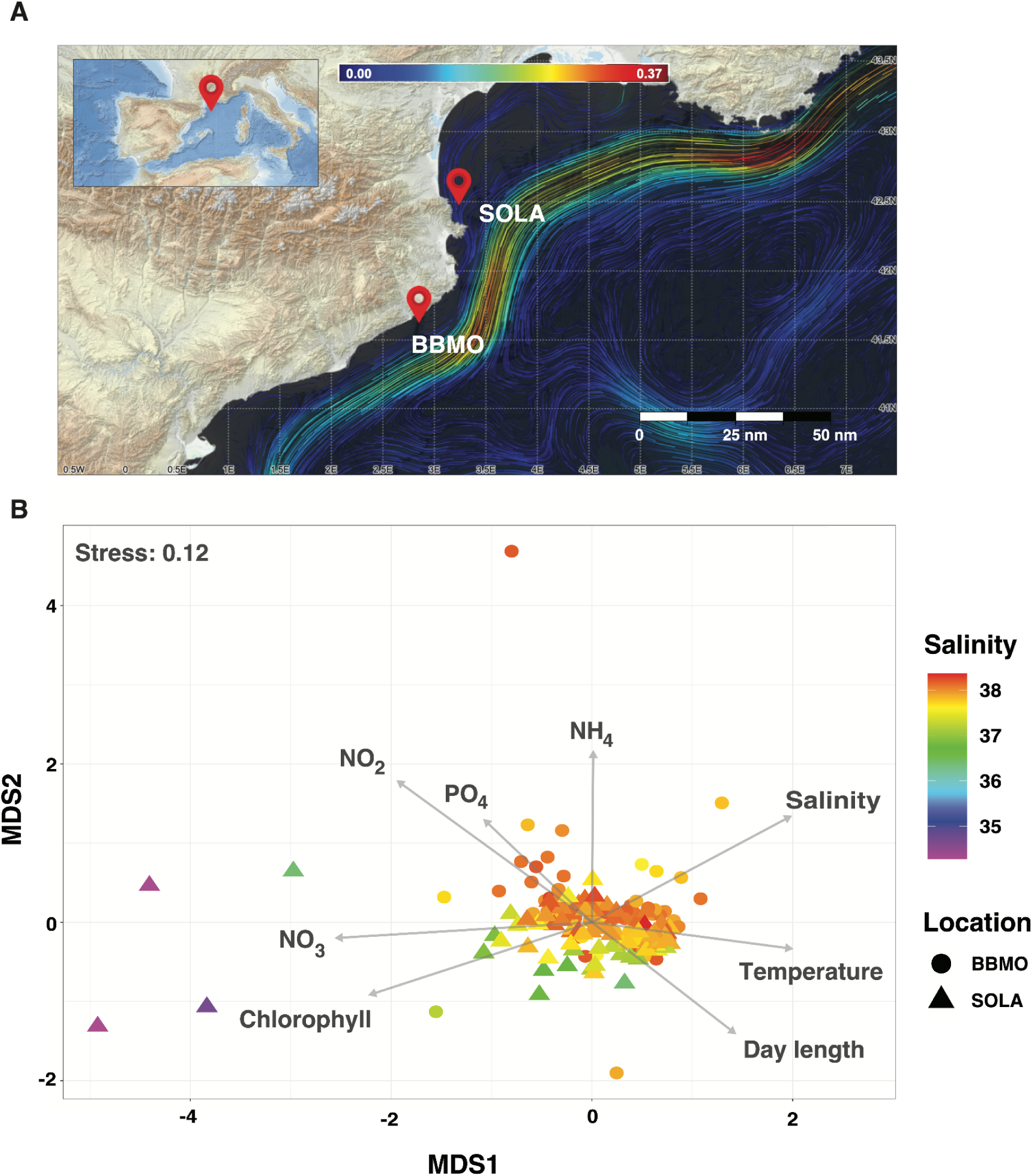
Location and environmental context of the BBMO and SOLA time series. **Panel A.** Location of the Blanes Bay Microbial Observatory (BBMO; 41°40′13″N, 2°48′00″E) in Spain and the Banyuls Bay Microbial Observatory (SOLA; 42°31’N 3°11’E) in France, in the NW Mediterranean Sea. Climatological January surface circulation for 1987–2019 (33 years), derived from MEDSEA products, shows the Northern Current flowing southwestward along the continental margin and potentially providing physical connectivity between the two sites^36^ (https://cosmo.icm.csic.es/currents). Streamline colors indicate estimated current speed (m s⁻¹; upper color bar). The inset shows the location of the study area within the western Mediterranean, and the scale bar indicates distance in nautical miles (nm). **Panel B.** Non-metric multidimensional scaling (NMDS) of environmental conditions at BBMO and SOLA during 2009–2015, based on temperature, salinity, day length, chlorophyll concentration, NH₄, NO₃, NO₂ and PO₄. Environmental variables were standardized as z-scores before analysis. Points represent individual samples, with shape indicating location and color indicating salinity; arrows show the direction of association of environmental variables with the ordination. BBMO and SOLA samples show substantial overlap in environmental space, although several low-salinity SOLA samples are clearly separated from the main cluster. These low-salinity observations likely reflect episodic freshwater inputs to the system^34^. NMDS stress = 0.12.

At BBMO, sampling occurred every month over 12 years, from January 2009 through December 2020 (140 samples). Surface seawater was pre-filtered through a 200-µm nylon mesh, transported to the laboratory in 20-L plastic carboys under dim light, and processed within two hours of collection. About 6 L of 200-μm pre-filtered seawater were sequentially filtered through a 20-μm mesh, a 3-μm pore-size polycarbonate filter (Poretics), and a 0.22-μm Sterivex (Merck-Millipore) filter using a peristaltic pump. Only the size fractions corresponding to the picoplankton (0.2-3μm) were used here. At SOLA, sampling occurred for 7 years, from January 2009 through December 2015 (90 samples). Samples were obtained using a 10-L Niskin bottle. The water was stored in high-density polyethylene carboys and kept in the dark until processed in the laboratory, which occurred within 1.5 hours. A total of 5 L was sequentially filtered through 3 μm pore-size polycarbonate filters (Millipore, Billerica, MA, USA) and 0.22-μm Sterivex filters (Merck-Millipore). Sterivex cartridges containing the pico-fraction of microbial biomass were stored at –80 °C until nucleic acid extraction was performed for both sampling stations. Both locations were sampled as part of the routine time-series sampling.

DNA extractions for BBMO samples followed the protocol of Schauer *et al.*^37^, where Sterivex cartridges were treated with lysozyme (20 mg/mL) for cell lysis, followed by proteinase K (20 mg/mL). The lysates were extracted twice with phenol-chloroform-isoamyl alcohol (25:24:1, pH 8) and once with chloroform-isoamyl alcohol (24:1), then purified using Amicon units (Millipore). For SOLA samples, DNA extraction followed the protocol of Hugoni *et al.*^38^, including lysozyme (20 mg/mL) and proteinase K (20 mg/mL) treatment. Pooled lysates were extracted using the AllPrep DNA/RNA kit (Qiagen, Hilden, Germany).

At BBMO, water temperature and salinity were sampled *in situ* with a SAIV A/S SD204 CTD. Inorganic nutrients (NO_3_^−^, NO_2_^−^, NH_4_^+^, PO_4_^3−^, SiO_2_) were measured on prefiltered seawater (200μm nylon mesh) using an Alliance Evolution II autoanalyzer^39^. Samples for chlorophyll *a* concentration were filtered through GF/F filters, extracted with acetone, and analyzed using fluorometry^40^. See further details in Gasol *et al*.^31^ and Krabberød *et al.*^9^. At SOLA, *in situ* temperature and salinity were measured using a Seabird CTD SBE9/11 instrument. Inorganic nutrients (NO_3_^−^, NO_2_^−^, PO_4_^3−^, SiO_2_) were measured using a Skalar auto-analyzer, following a previously established protocol^41^. Other physicochemical parameters were provided by the Service d’Observation en Milieu Littoral (SOMLIT; https://www.somlit.fr/en/). Chlorophyll *a* concentrations were determined from 1 L of seawater filtered through a GF/F filter under low pressure (< 0.2 bar), following the method outlined in Galand et al. ^42^. See further details in Galand *et al.*^32^.

The extracted DNA from BBMO was used for *Illumina* (short-read) and PacBio (long-read; Sequel II, HiFi) shotgun sequencing. For the short reads, the first 3 years were sequenced using an Illumina Hiseq4000, and for the following 9 years, using Illumina NovaSeq6000 (2 x 150 bp, Centre Nacional d’Anàlisi Genómica, CNAG, Spain). All metagenomes from BBMO had a sequencing coverage of >30Gb. SOLA metagenomes from January 2009 to December 2011 and from March to December 2015 had a sequencing coverage of >30Gb. SOLA metagenomes from January 2012 to February 2015 had a mean sequencing coverage of 14.9 Gb (SD = 3.8 Gb) and ranged in coverage from 7.0 to 22.9 Gb^32^. SOLA metagenomes with >30 Gb of sequencing coverage were produced with *Illumina* NovaSeq6000 (2 x 150 bp, Centre Nacional d’Anàlisi Genómica CNAG), while the rest were sequenced with HiSeq2500 (2x100bp).

Three BBMO samples from February 2009, August 2009, and January 2010 were shotgun sequenced on the PacBio Sequel II platform. The samples were fragmented to 12-16 kbp and multiplexed before all three were pooled, cleaned, and size-selected for fragments >3 kb. Sequencing was conducted on an 8M SMRT cell on the Sequel II using the Sequel II Binding Kit 2.0 and Sequencing Chemistry v2.0 (PacBio). HiFi Circular Consensus Sequences (CCS) were generated using the CCS pipeline in SMRT Link (v10.1.0.119588) with default settings. CCS reads were subsequently demultiplexed and assigned to samples (**Supp. Figure 1, Supp. Table 1**). Sequencing was conducted at the Norwegian Sequencing Centre (www.sequencing.uio.no).

**Table 1.**
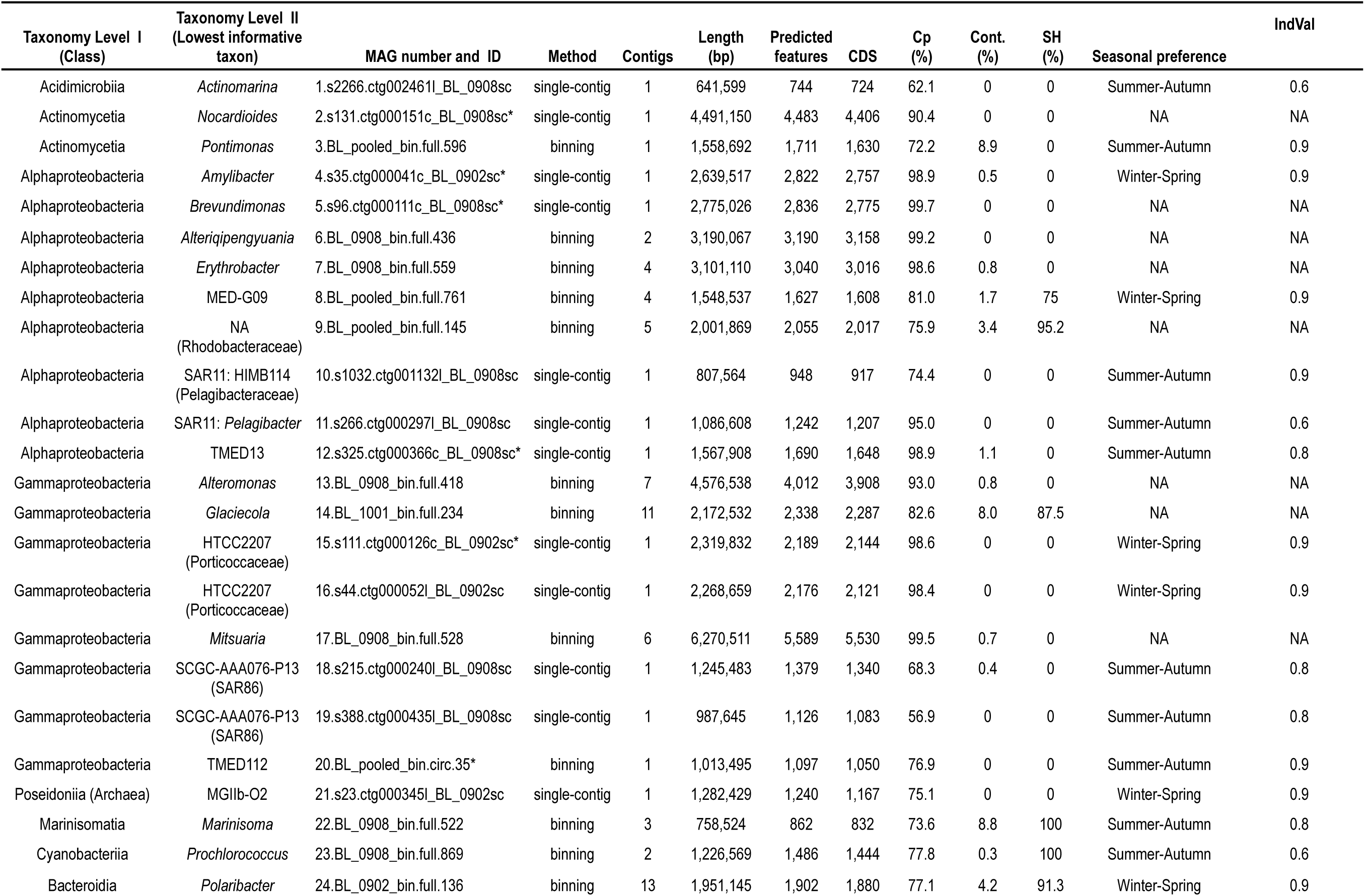
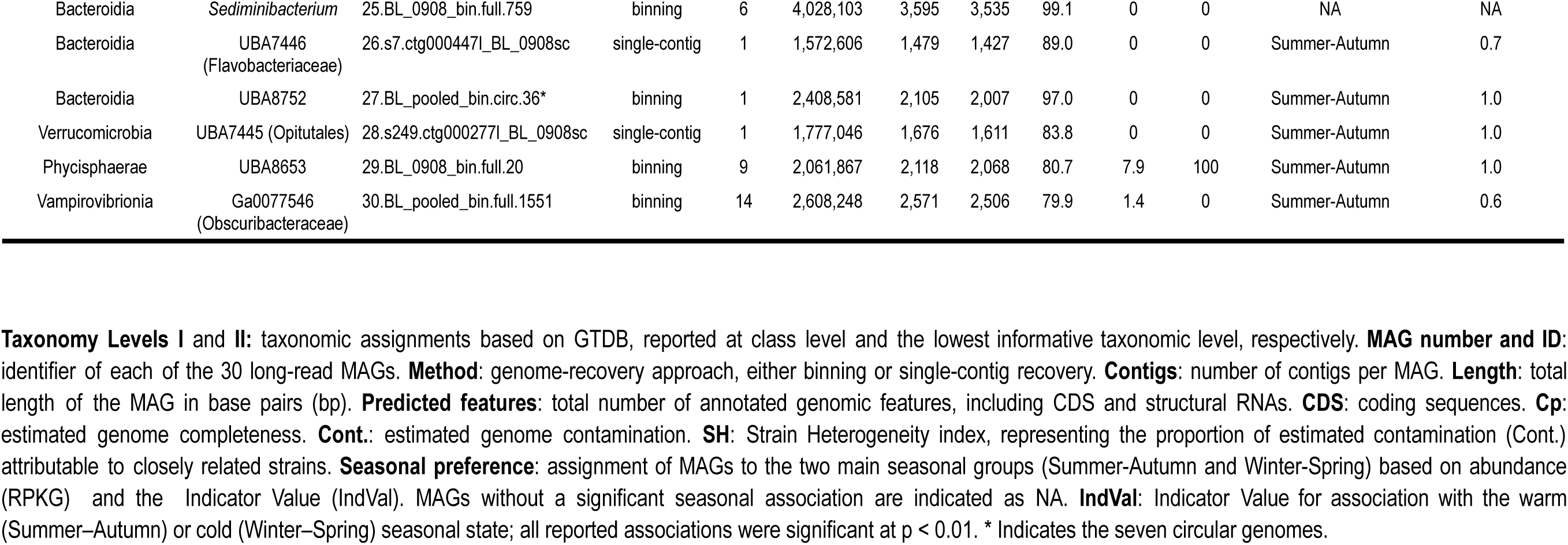
The 30 PacBio HiFi long-read BBMO MAGs investigated in this study.

Samples were assembled individually and co-assembled using hifiasm-meta^43^ (**Supp. Figure 2, Supp. Table 2**). A total of 20 single-contig MAGs were recovered from the single-sample assemblies featuring > 50% genome completeness and >500 kb contig length. No binning was used for these MAGs. In order to recover additional MAGs, contigs from individual samples, and also from the coassembly were binned using the HiFi MAG pipeline v1.5 (https://github.com/PacificBiosciences/pb-metagenomics-tools/blob/master/docs/Tutorial-HiFi-MAG-Pipeline.md) with MetaBat2^44^. This generated a total of 37 medium- to high-quality PacBio MAGs (>70% completeness, <10% contamination, <20 contigs)^45^.

**Table 2.**
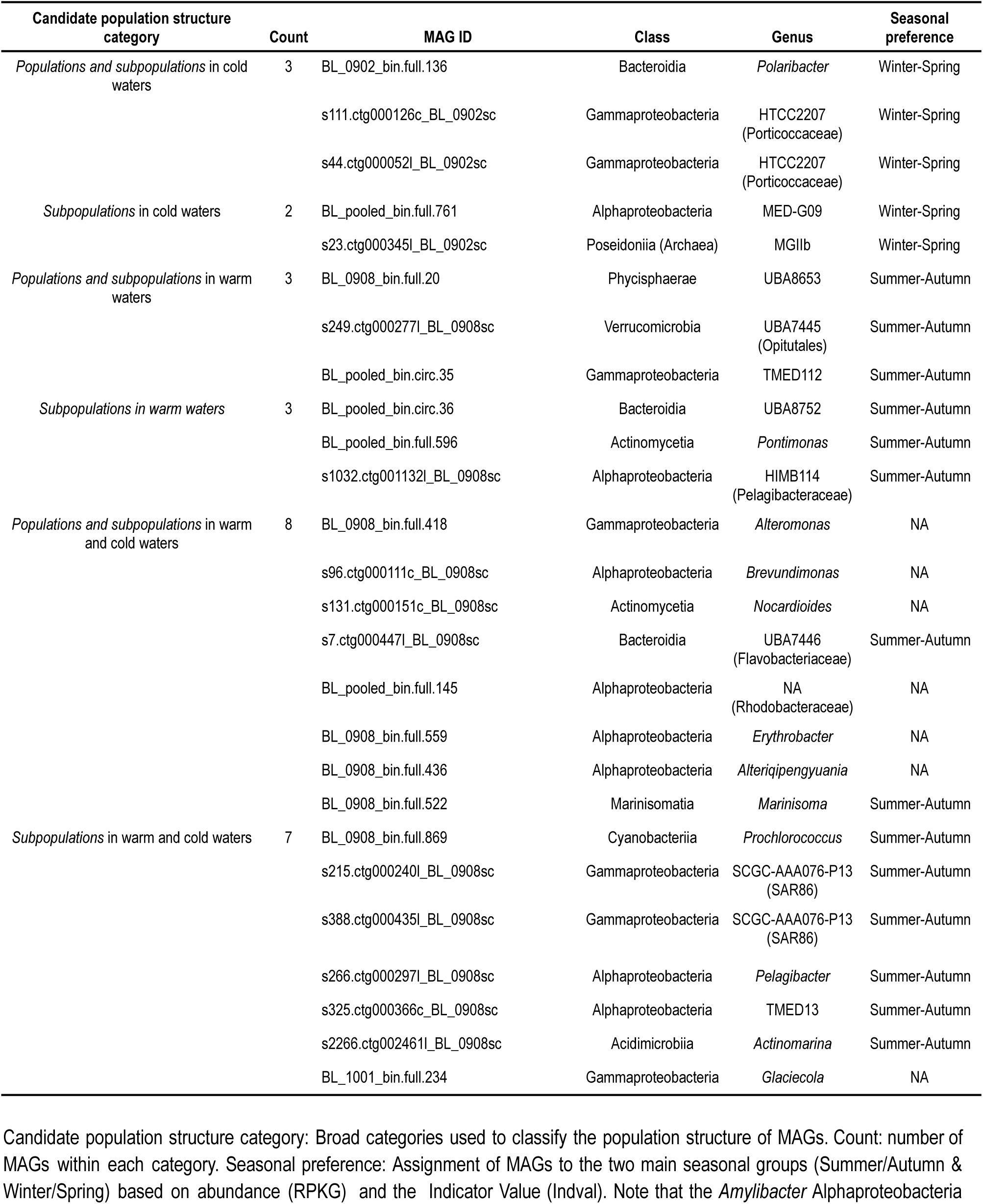
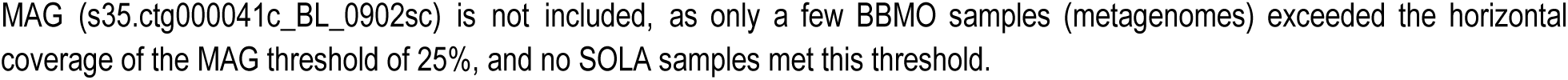
Main candidate population structure patterns identified in the 26 MAGs associated with temperature.

| Candidate population structure category | Count | MAG ID | Class | Genus | Seasonal preference |
| --- | --- | --- | --- | --- | --- |
| Populations and subpopulations in cold waters | 3 | BL_0902_bin.full.136 | Bacteroidia | *Polaribacter* | Winter-Spring |
|  |  | s111.ctg000126c_BL_0902sc | Gammaproteobacteria | HTCC2207 (Porticoccaceae) | Winter-Spring |
|  |  | s44.ctg000052l_BL_0902sc | Gammaproteobacteria | HTCC2207 (Porticoccaceae) | Winter-Spring |
| Subpopulations in cold waters | 2 | BL_pooled_bin.full.761 | Alphaproteobacteria | MED-G09 | Winter-Spring |
|  |  | s23.ctg000345l_BL_0902sc | Poseidoniiia (Archaea) | MGIIb | Winter-Spring |
| Populations and subpopulations in warm waters | 3 | BL_0908_bin.full.20 | Phycisphaerae | UBA8653 | Summer-Autumn |
|  |  | s249.ctg000277l_BL_0908sc | Verrucomicrobia | UBA7445 (Opitutales) | Summer-Autumn |
|  |  | BL_pooled_bin.circ.35 | Gammaproteobacteria | TMED112 | Summer-Autumn |
| Subpopulations in warm waters | 3 | BL_pooled_bin.circ.36 | Bacteroidia | UBA8752 | Summer-Autumn |
|  |  | BL_pooled_bin.full.596 | Actinomycetia | *Pontimonas* | Summer-Autumn |
|  |  | s1032.ctg001132l_BL_0908sc | Alphaproteobacteria | HIMB114 (Pelagibacteraceae) | Summer-Autumn |
| Populations and subpopulations in warm and cold waters | 8 | BL_0908_bin.full.418 | Gammaproteobacteria | *Alteromonas* | NA |
|  |  | s96.ctg000111c_BL_0908sc | Alphaproteobacteria | *Brevundimonas* | NA |
|  |  | s131.ctg000151c_BL_0908sc | Actinomycetia | *Nocardioides* | NA |
|  |  | s7.ctg000447l_BL_0908sc | Bacteroidia | UBA7446 (Flavobacteriaceae) | Summer-Autumn |
|  |  | BL_pooled_bin.full.145 | Alphaproteobacteria | NA (Rhodobacteraceae) | NA |
|  |  | BL_0908_bin.full.559 | Alphaproteobacteria | *Erythrobacter* | NA |
|  |  | BL_0908_bin.full.436 | Alphaproteobacteria | *Alteriqipengyuania* | NA |
|  |  | BL_0908_bin.full.522 | Marinisomatia | *Marinisoma* | Summer-Autumn |
| Subpopulations in warm and cold waters | 7 | BL_0908_bin.full.869 | Cyanobacteriia | *Prochlorococcus* | Summer-Autumn |
|  |  | s215.ctg000240l_BL_0908sc | Gammaproteobacteria | SCGC-AAA076-P13 (SAR86) | Summer-Autumn |
|  |  | s388.ctg000435l_BL_0908sc | Gammaproteobacteria | SCGC-AAA076-P13 (SAR86) | Summer-Autumn |
|  |  | s266.ctg000297l_BL_0908sc | Alphaproteobacteria | *Pelagibacter* | Summer-Autumn |
|  |  | s325.ctg000366c_BL_0908sc | Alphaproteobacteria | TMED13 | Summer-Autumn |
|  |  | s2266.ctg002461l_BL_0908sc | Acidimicrobiia | *Actinomarina* | Summer-Autumn |
|  |  | BL_1001_bin.full.234 | Gammaproteobacteria | *Glaciecola* | NA |

Candidate population structure category: Broad categories used to classify the population structure of MAGs. Count: number of MAGs within each category. Seasonal preference: Assignment of MAGs to the two main seasonal groups (Summer/Autumn & Winter/Spring) based on abundance (RPKG) and the Indicator Value (Indval). Note that the *Amylibacter* Alphaproteobacteria
MAG (s35.ctg000041c\_BL\_0902sc) is not included, as only a few BBMO samples (metagenomes) exceeded the horizontal coverage of the MAG threshold of 25%, and no SOLA samples met this threshold.

The 57 unbinned and binned MAGs were analyzed with CheckM2^46^ and compareM (https://github.com/dparks1134/CompareM) to assess completeness, contamination, strain heterogeneity, and redundancy. MAGs featuring >99% similarity in genome-wide Average Amino Acid Identity (AAI) were considered redundant, and the best was selected based on quality (quality = completeness – 5X contamination)^47^. We applied the following selection procedure. MAGs derived from binning were selected when they were clearly superior in terms of quality to single-contig MAGs, or if single-contig MAGs did not reach 70% completeness. Otherwise, single-contig MAGs were selected. Then, bins with the best values of quality and strain heterogeneity were selected. In addition, when a bin from a single-sample metagenome had an overall quality similar to that of another from the co-assembly, the former was selected. After performing this selection process, 30 non-redundant (<90% AAI) MAGs with completeness above 50% and a percentage of contamination below 10% were selected for downstream analyses (**Table 1; Supp. Figure 3**). The genome-wide Average Nucleotide Identity (ANI) among the 30 MAGs was calculated using FastANI^48^. All MAGs exhibited an ANI below 95%, a widely accepted threshold for species delimitation, beyond which homologous recombination approaches zero^48,49^. Thus, the 30 MAGs are considered different species.

While 11 MAGs were recovered by the three approaches (i.e., no-binning (single-contig MAGs), binning of single-sample contigs, and binning of the coassembly), there were MAGs that were recovered with only one or two methods (**Supp. Figure 4**). In particular, there were six MAGs that were recovered only as single-contig (no binning), four that were recovered only from the single-sample assemblies, and four that were recovered from the co-assembly only (**Supp. Figure 4**). This supports the use of different approaches for long-read MAG generation. The maximum number of contigs within a single MAG was 14 (**Table 1**). A total of 7 of 30 PacBio MAGs represented complete circular genomes, retrieved as a single contig (**Table 1**).

Gene prediction and annotation of the MAGs were conducted using prokka-v1.14.6^50^ and eggNOG-mapper v2.1.9^51^. As expected, the number of annotated features increased linearly with genome size, supporting MAG construction (**Supp. Figure 5**). Taxonomy assignment was performed using GTDB-Tk^52^, which indicated 29 Bacterial and 1 Archaeal MAGs (**Table 1**). For each of the 24 MAGs for which relatively close relatives could be identified in GTDB, the Average Nucleotide Identity (ANI) and the Alignment Fraction (AF) with the closest reference genome were calculated (**Supp. Figure 6**). The AF shows the amount of genome overlap with the reference genome, which was used to calculate the ANI. There were 15 MAGs with an AF > 0.8 and an ANI >90%, and 5 MAGs with an AF > 0.9 and an ANI > 93%. Such similarity between some MAGs (including several single-contig MAGs) and reference genomes supports our MAG construction strategy. Two pairs of single-contig MAGs had the same taxonomic classification: HTCC2207, Porticoccaceae (s111.ctg000126c_BL_0902sc & s44.ctg000052l_BL_0902sc) and SCGC-AAA076-P13, SAR86 (s215.ctg000240l_BL_0908sc & s388.ctg000435l_BL_0908sc). The Average Amino Acid Identity (AAI) between each pair of MAGs was 87.5% and 88.0%, respectively. In agreement, the Average Nucleotide Identity (ANI) was 82% and 89% for each pair, respectively. This indicates that both pairs are distinct species (using the ANI < 95% criterion) from the same lineage.

### Mapping

140 short-read monthly Illumina metagenomes from the BBMO covering 12 years and 90 from SOLA covering 7 years were mapped individually to the 30 PacBio long-read MAGs with the Burrows-Wheeler Aligner v0.7.17-r1188 (BWA)^53^, using the BWA-MEM algorithm. In addition, we implemented competitive mapping that involved combining the 30 PacBio MAGs to create a combined reference genome pool. Subsequently, the 230 short-read metagenomes (140 from BBMO plus 90 from SOLA) were mapped onto the 30 PacBio long-read MAGs using identical BWA parameters. This approach can resolve read-mapping ambiguity by ensuring that each short read is uniquely assigned to its best-matching MAG within the pool. This yielded the third dataset, named as “MIX”. BAM files containing the total number of reads mapped per metagenome and their corresponding genomic locations were generated for each MAG. Hits were filtered by >95% identity and >80% coverage of the read. The latter thresholds reduce the likelihood that mapped reads originate from taxa other than the query MAG species. BAM files were analyzed to calculate the percentage of MAGs covered by hits (horizontal coverage) and the average number of hits per nucleotide within the mapped regions of each MAG (vertical coverage). MAG abundances were estimated using RPKG (mapped Reads Per Kilobase of genome per Gigabase of metagenome).

### Single Nucleotide Variant (SNV) calling and Nucleotide Diversity

SNVs involve the substitution of a single nucleotide at a specific position in the genome. SNVs were called using the program Freebayes^54^, considering ploidy = 1. SNVs were included in the analysis only if they had a minimum coverage of 10 reads in a given sample and the position (locus) was present in at least four distinct samples. Variant Call Files (VCFs) were produced with information on the genomic position of variants.

Genome-wide nucleotide diversity (π) refers to the average number of nucleotide differences per site between any two sequence reads taken at random from the sample populations (0<= π <1)^55^. To calculate the genome-wide π with POGENOM, π is averaged over all loci by summing all π_i_ and dividing by the genome size.

### Fixation Index (Fst)

The Fixation Index (Fst) measures population differentiation based on genetic structure^56,57^. It ranges between 0 (no population differentiation) and 1 (complete population differentiation). Here, for each MAG, genome-wide Fst pairwise distances among all samples (metagenomes) were calculated with the program POGENOM^55^ from the VCF files.

UPGMA hierarchical clustering dendrograms based on the Fst distances were generated. We used the threshold of Fst=0.15 in the UPGMA to delineate candidate populations^58^. MAGs were considered to be present in a sample when more than 25% of horizontal coverage was detected; all samples featuring >25% of horizontal coverage displayed RPKG values >0, as expected. To identify the main patterns of population structure, we focused on samples in which MAGs displayed high abundance relative to the other samples.

### Detecting candidate positively selected genes

For each coding gene in each genome, the ratio of non-synonymous to synonymous polymorphisms (pN/pS) was calculated using POGENOM in each sample. pN/pS values were considered as follows: pN/pS = 1: signal coherent with neutral evolution; pN/pS <1: signal coherent with purifying or negative selection; pN/pS > 1: signal coherent with positive or diversifying selection. Genes showing signatures of positive selection (pN/pS > 1) in specific populations may represent part of the genetic basis of adaptation to particular conditions (e.g., summer or winter). Differences in pN/pS values for a coding gene across samples may reflect changes in the variants or strains contributing to that gene in each sample.

To identify positively selected genes, the criteria were:

1. Genes had a mean pN/pS > 0.8 over the 12 years for BBMO or 7 years for SOLA. This was also applied to the combined BBMO and SOLA data (MIX dataset).
2. Genes had a pN/pS > 1 in at least one sample

Genes matching the two criteria were considered candidate positively selected genes and used in downstream analyses. The reason for setting the mean pN/pS cutoff to 0.8 rather than 1 was to allow more genes to be considered. This is because we expect multiple genes to have low pN/pS across several samples, given that they represent the population of the template genome (i.e., the population of the MAG used for read mapping), thereby lowering the overall mean pN/pS.

### Statistical Analyses

Most statistical analyses were carried out in R 4.1.1^59^ and R-Studio v 2021.9.0.351^60^, using packages such as *tidyverse*^61^ and *dplyr*^62^. Plots were generated in *ggplot2*^63^. Mantel tests were carried out using *Vegan*^64^, and 999 permutations were run in parallel with six cores and 250 GB of RAM. We used the Indicator Value (IndVal) implemented in the R package *labdsv*^65^ to assign MAGs to seasons based on their RPKGs. Seasons were classified as follows: Winter (December to February), Spring (March to May), Summer (June to August), and Autumn (September to November). The Summer and Autumn seasons were categorized as the warm-water group, while Winter and Spring were considered the cold-water group. The group Others include MAGs that IndVal could not assign to any season. Only MAGs with an IndVal p-value below 0.01 were considered to belong to a seasonal group; the remaining were assigned to the group Others.

## RESULTS

### Abundance and seasonality of the 30 recovered MAGs

We mapped short reads from BBMO (12 years) and SOLA (7 years) back to the 30 selected MAGs from BBMO (**Table 1**) to determine their seasonal abundance at both sites (**Supp. Figure 7)**. The analysis of seasonal dynamics indicated that, of the 30 MAGs, 15 were associated with Summer–Autumn, 6 with Winter–Spring, and 9 showed no significant seasonal preference **(Table 1)**. Some MAGs showed clear rhythmic synchronous seasonality in both BBMO and SOLA, with peaks of abundance in the same months (e.g., Bacteroidia UBA8752, *Pontimonas*, Alphaproteobacteria MED-G09, Poseidoniia, and *Prochlorococcus*; **Supp. Figure 7**). Other MAGs displayed comparable seasonality patterns in both time series, with small shifts in their peaks of abundance (e.g., TMED112 & SCGC-AAA076-P13 [Gammaproteobacteria], Rhodobacteraceae, HIMB114 [Pelagibacteriaceae], *Actinomarina*). A number of MAGs displayed non-synchronous dynamics, including conditionally rare taxa (CRT; e.g., *Mitsuaria* [Gammaproteobacteria], *Sediminibacterium* [Bacteroidota], *Amylibacter* [Alphaproteobacteria], HTCC2207 [Gammaproteobacteria; two MAGs] (**Supp. Figure 7**). CRTs are taxa that are typically present at low relative abundances, but can become temporarily abundant under specific environmental conditions or at particular times ^66^.

CRTs were recovered during peaks in abundance in BBMO. For example, *Mitsuaria* and *Sediminibacterium* had a very low abundance in BBMO and SOLA during 12 and 7 years, respectively, except in August 2009 in BBMO, when the sample for long-read metagenomes was taken [**Supp. Figure 7**]. This supports the capability of long-read PacBio sequencing to capture highly contiguous MAGs from rare taxa during their occasional peaks of abundance (blooms). Other MAGs were also recovered from BBMO samples, in which they displayed peaks of abundance. For example, August 2009 (*Nocardioides*, *Alteromonas*, *Brevundimonas*) and February 2009 (*Amylibacter*, HTCC2207 [Porticoccaceae, Gammaproteobacteria; 111.ctg000126c_BL_0902sc]).

We analyzed the horizontal and vertical short-read coverage of the 30 long-read MAGs **(Supp. Figure 8)**. All MAGs had nearly 100% horizontal coverage in at least one sample. Three MAGs (BL_0908_bin.full.528 [*Mitsuaria*], BL_0908_bin.full.759 [*Sediminibacterium*], and BL_pooled_bin.full.1551 [Obscuribacteraceae (Cyanobacteria)]) were abundant in just a single sample **(Supp. Figure 8)**. Two of these three MAGs (*Mitsuaria* & *Sediminibacterium*) were conditionally rare taxa, which is consistent with the low sample abundance (**Supp. Figure 7**). These three MAGs were excluded from the downstream population differentiation analyses, as population-level information was available for only a single sample. For the 27 remaining long-read MAGs, we included only samples (short-read metagenomes) with ≥25% horizontal MAG coverage to ensure robust estimates of population diversity and differentiation (**Supp. Figure 8**). The abundance in these samples showed substantial variation in both BBMO and SOLA **(Supp. Figure 9).**

### Genetic diversity

The genome-wide nucleotide diversity (π) of the 27 MAGs across 12 years at BBMO and 7 years at SOLA frequently displayed rhythmic patterns that broadly coincided with seasonal abundance dynamics (**Supp. Figure 7**). In several MAGs (e.g., MED-G09, Alphaproteobacteria [BL_pooled_bin.full.761] and HIMB114, Pelagibacteraceae [s1032.ctg001132l_BL_0908sc]), similar levels of nucleotide diversity recurred across years, suggesting that comparable levels of within-population genetic diversity were maintained over time. In other MAGs, π peaked during particular periods or individual samples without necessarily coinciding with increases in MAG abundance (e.g., *Alteriqipengyuania* and *Brevundimonas*; **Supp. Figure 7**). In some cases, π showed no clear rhythmicity and remained relatively constant through time (e.g., the *Pelagibacter* s266.ctg000297l_BL_0908sc and SAR86 s215.ctg000240l_BL_0908sc and s388.ctg000435l_BL_0908sc MAGs at BBMO). In other MAGs, π initially increased with MAG abundance and then reached a plateau. Thus, further increases in abundance were not necessarily accompanied by increases in nucleotide diversity. For individual MAGs, π patterns often differed between the two time series, indicating location-specific differences in nucleotide-diversity dynamics (**Supp. Figure 7**).

### Population differentiation and structure

For a given MAG, pairwise Fst quantifies genome-wide population differentiation between the populations represented in two samples. We used this metric to assess population differentiation for 27 MAGs across 12 years at BBMO and 7 years at SOLA, considering only samples with at least 25% horizontal coverage of the corresponding MAG. Some MAGs showed consistently low averages with small standard deviations across both time series, indicating minimal population-level differentiation within and between years (e.g., the *Pelagibacter* MAG [s266.ctg000297l_BL_0908sc]; **Figure 2**). In turn, other MAGs had high mean Fst values and large standard deviations in both time series over the years, indicating typically high and more variable levels of population differentiation (e.g., Verrucomicrobia [s249.ctg000277l_BL_0908sc], *Glaciecola* [BL_1001_bin.full.234]; **Figure 2**). The Fst distribution of most MAGs (17 of the 23 for which Fst could be calculated in both time series) differed significantly between BBMO and SOLA (**Figure 2**), indicating site-specific differences in population differentiation and suggesting differences in the populations represented at each location during the sampling periods. In turn, comparable Fst values in both time series suggest similar levels of population divergence at the two locations, potentially reflecting the recurrent presence of the same or closely related populations at both sites.

**Figure 2.**
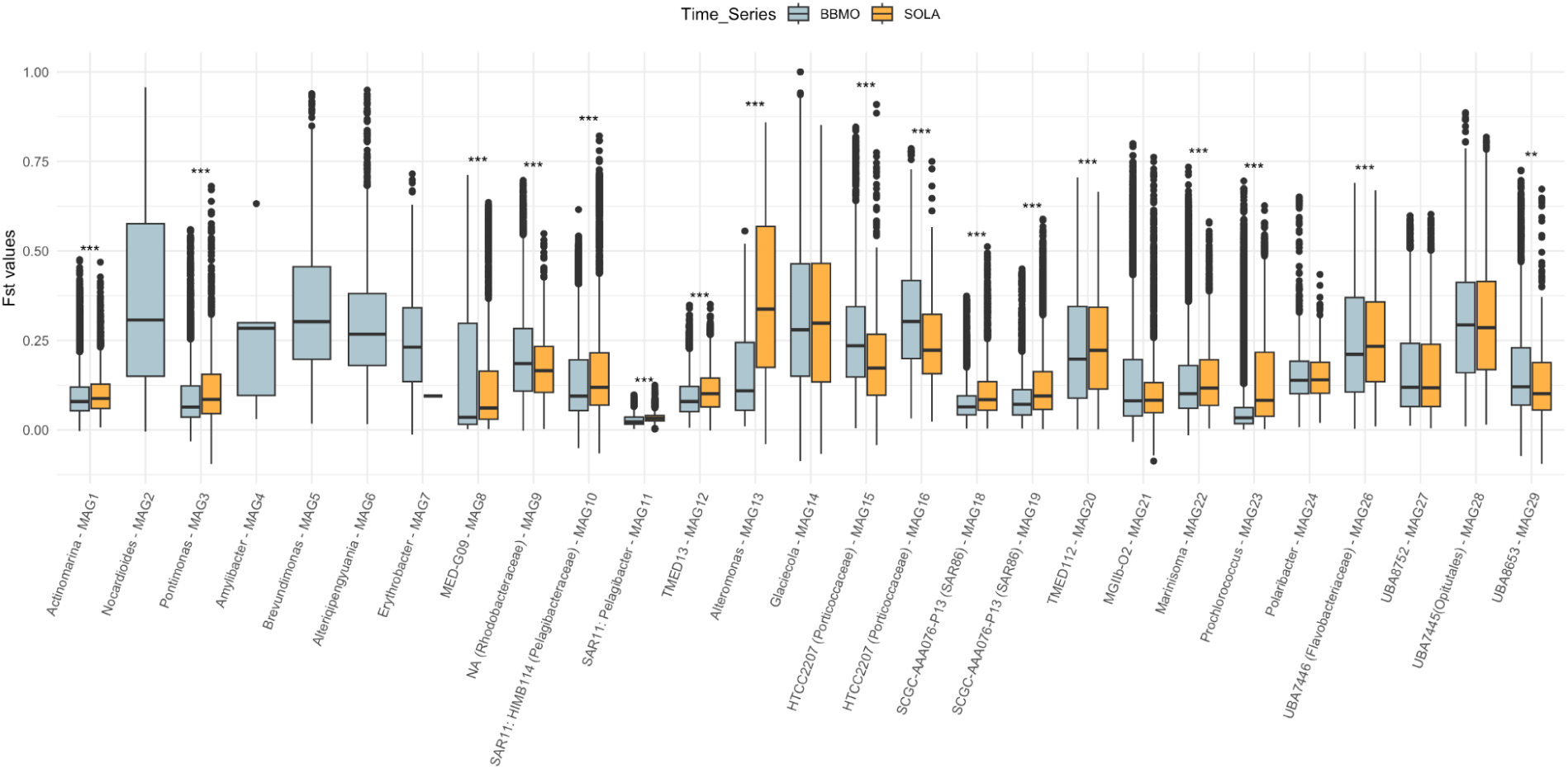
Distribution of pairwise genome-wide Fst values across the BBMO and SOLA time series for 27 MAGs. Pairwise Fst values were calculated between samples collected over 12 years at BBMO and 7 years at SOLA. Only sample pairs in which both samples had ≥25% horizontal coverage of the corresponding MAG were included. Boxplots show the distribution of pairwise Fst values for BBMO and SOLA. Asterisks indicate differences between the two time series based on Wilcoxon tests (*** P-value <= 0.001; ** P-value <= 0.01; * P-value <= 0.05).

We investigated whether genetic differentiation (Fst) was associated with temporal distance in both BBMO and SOLA. A positive association between Fst and temporal distance indicates that populations sampled farther apart in time tend to be more genetically differentiated. Such divergence could reflect processes including strain replacement, migration, demographic stochasticity, or ongoing evolutionary change^28^. Conversely, a negative association would indicate lower differentiation between populations sampled farther apart in time, potentially reflecting recurrent population or ecotype states. For each of the 27 MAGs, we tested the association between pairwise Fst and temporal distance (days between samples) using Mantel tests and characterized the direction and magnitude of the relationship using linear regression, considering only samples with >25% horizontal coverage of the corresponding MAG (**Supp. Figure 10; Supp. Table 5**).

In the linear regressions, the slope represents the fitted rate at which pairwise Fst changes with temporal distance. Positive slopes indicate greater genetic differentiation between samples collected farther apart in time, whereas negative slopes indicate lower differentiation. Among the 49 MAG × time-series combinations for which Fst could be analyzed, 35 combinations, representing 24 MAGs, had regression slopes with p < 0.05: 23 at BBMO and 12 at SOLA (**Supp. Table 5; Supp. Figure 10**). Of these, 31 slopes were positive (21 at BBMO and 10 at SOLA), whereas four were negative (two at each site). Slopes ranged from −6.73 × 10⁻⁶ to 2.13 × 10⁻⁴ Fst units day⁻¹. Adjusted R² values quantify the proportion of variation in pairwise Fst captured by the linear relationship with temporal distance. Values ranged from 0.0002 (0.02%) to 0.4859 (48.59%) and were generally low, indicating that temporal distance accounted for little of the variation in Fst in most MAGs. However, five MAGs at BBMO had adjusted R² values above 0.1, indicating stronger linear associations between genetic differentiation and temporal distance: *Nocardioides* (0.4859; 48.59%), *Erythrobacter* (0.1797; 17.97%), *Brevundimonas* (0.1780; 17.80%), *Pelagibacter* (0.1644; 16.44%), and *Alteriqipengyuania* (0.1264; 12.64%) (**Supp. Table 5; Supp. Figure 10**). Mantel tests supported significant positive associations between Fst and temporal distance in 21 MAG × time-series combinations, representing 18 MAGs: 15 at BBMO and 6 at SOLA. In contrast, none of the four negative regression slopes was supported by a significant Mantel test. Significant Mantel Pearson correlations ranged from 0.0634 to 0.6975, indicating substantial variation in the strength of temporal associations among MAGs. Eight MAG × time-series combinations, representing seven MAGs, had Mantel correlations >0.2, and the five BBMO MAGs with adjusted R² >0.1 also showed the strongest Mantel correlations (r > 0.35) (**Supp. Table 5**). Overall, these analyses support increasing population-level genetic differentiation with temporal distance in a subset of MAGs, with supported temporal associations being more frequent and generally stronger at BBMO than at SOLA within the analyzed time series.

For each MAG, we used Fst to identify candidate populations. We used an Fst = 0.15 threshold in UPGMA dendrograms to define sample groups that may represent different populations, while, in specific cases, distinct clusters with an Fst < 0.15 were regarded as candidate subpopulations **(Supp. Figure 9)**. We analyzed metagenomes from BBMO and SOLA together to detect candidate populations present in both time series. To identify main population structure patterns, we focused on samples with moderate or high MAG abundances. We classified MAGs based on whether their population structure was consistent with a seasonal preference (Summer—Autumn vs. Winter—Spring; Table 1,^9,14^).

The analysis revealed diverse population and subpopulation structures and seasonal preferences among the investigated MAGs (**Table 2, Supp. Figure 9**). In cold waters, we identified candidate populations and subpopulations in MAGs from *Polaribacter* and HTCC2207 (Gammaproteobacteria), as well as MED-G09 (Alphaproteobacteria) and MGIIb (Archaea). In warm waters, candidate populations and subpopulations were observed in UBA8653 (Phycisphaerae), UBA7445 (Verrucomicrobia), TMED112 (Gammaproteobacteria), UBA8752 (Bacteroidia), *Pontimonas*, HIMB114 (Pelagibacteraceae), and SAR86 (Gammaproteobacteria) (**Table 2, Supp. Figure 9**). Several MAGs showed candidate populations and subpopulations in both cold and warm waters, consistent with thermal niche differentiation (**Figure 3**). These MAGs include those from *Alteromonas*, *Brevundimonas*, *Marinisoma*, *Nocardioides*, UBA7446 (Flavobacteriaceae), Rhodobacteraceae, *Erythrobacter*, and *Alteriqipengyuania*. Additionally, MAGs from *Prochlorococcus*, SAR86, *Pelagibacter*, TMED13 (Alphaproteobacteria), *Actinomarina*, and *Glaciecola* exhibited subpopulation-level structure across both thermal environments (**Figure 3**, **Table 2, Supp. Figure 9**). Two pairs of MAGs, classified as HTCC2207 (Porticoccaceae, Gammaproteobacteria) and SCGC-AAA076-P13 (SAR86, Gammaproteobacteria), exhibited distinct population structures. This aligns with their measured AAI (∼88%) and ANI (82-89%), indicating they represent different species^49^ (**Supp. Figure 9**). These findings highlight the broad diversity in population-level structures observed among prokaryotic genomes, both within and across lineages.

**Figure 3.**
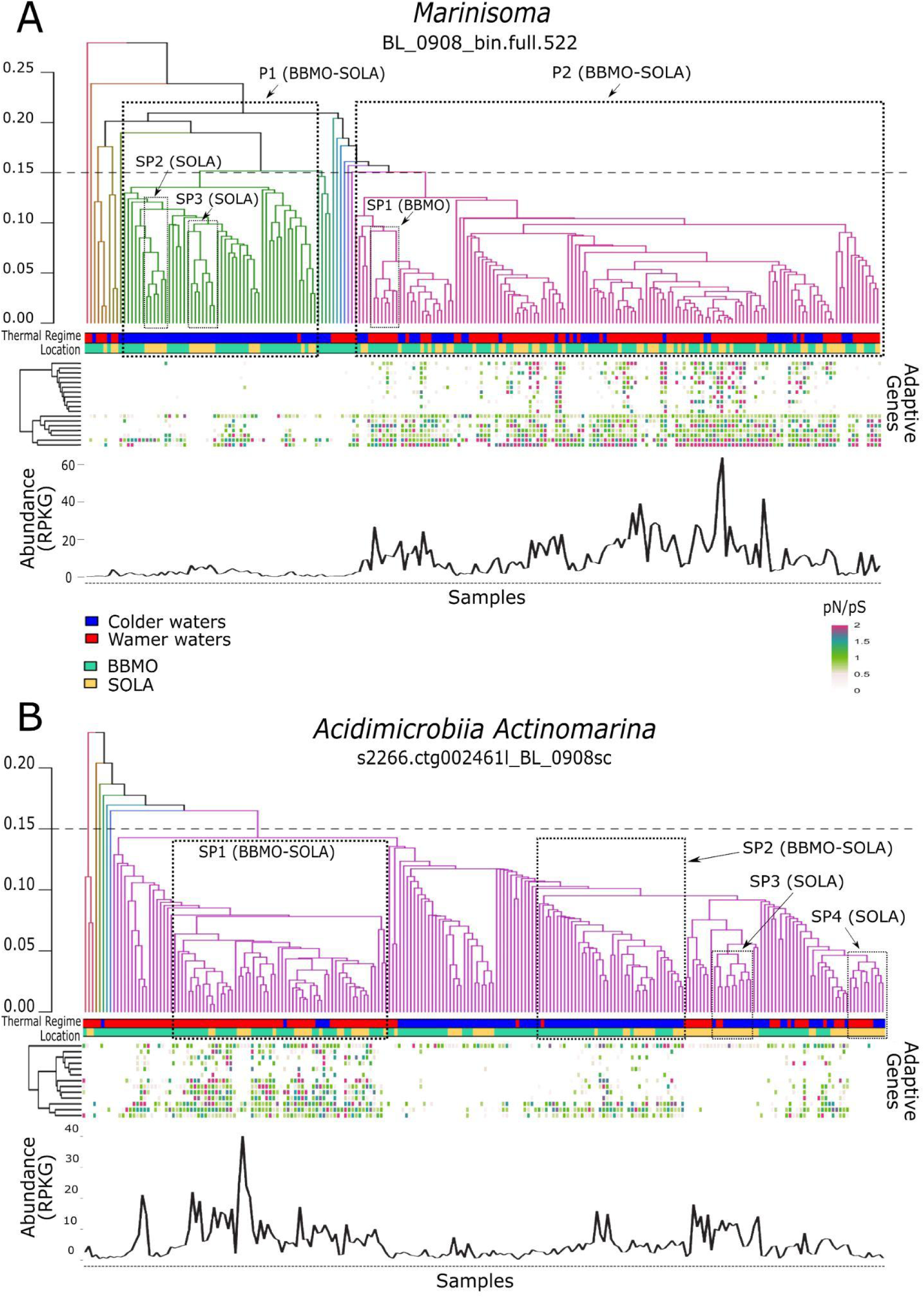
Examples of population and subpopulation structure in *Marinisoma* (A) and *Actinomarina* (B). For each MAG, the upper dendrogram shows UPGMA clustering based on pairwise genome-wide Fst among samples. A heuristic Fst cut-off of 0.15 (dashed horizontal line) was used to delineate candidate populations, with specific nested clusters below this threshold interpreted as candidate subpopulations. Branch colors indicate the resulting candidate population clusters. Bars below the dendrogram show the thermal regime of each sample (Winter–Spring, colder waters; Summer–Autumn, warmer waters) and sampling location (BBMO or SOLA). (A) *Marinisoma* separates into two candidate populations, P1 and P2, both represented at BBMO and SOLA. P1 is mainly associated with colder waters and lower MAG abundance, whereas P2 is predominantly associated with warmer waters and higher abundance. Three nested subpopulations are highlighted: SP1, composed of BBMO samples and mainly associated with warmer waters, and SP2 and SP3, composed of SOLA samples and mainly associated with colder waters. (B) *Actinomarina* shows two major candidate subpopulation clusters, SP1 and SP2, containing samples from both time series. SP1 is predominantly associated with warmer waters, whereas SP2 is more frequently associated with colder waters. Two additional smaller SOLA-specific clusters, SP3 and SP4, are also indicated. Heatmaps show pN/pS values for candidate adaptive genes across the corresponding samples, with values >2 capped at 2 for visualization. White cells indicate unavailable or undefined pN/pS values. The lower line plots show MAG abundance (RPKG) across samples with >25% horizontal coverage. Samples are ordered according to the Fst-based dendrogram and therefore are not arranged chronologically.

In several MAGs, within the candidate populations and subpopulations, we observed samples from both time series, suggesting that members of the same or closely related populations were present in both locations during the same or different months (**Figure 3, Supp. Figure 9**). Examples were observed in *Polaribacter*, UBA8653, *Marinisoma*, *Prochlorococcus*, TMED112, UBA8752, Rhodobacteraceae, *Pontimonas*, MED-G09, UBA7446, Poseidoniia, HTCC2207 (both MAGs), SAR86 (both MAGs), UBA7445, *Pelagibacter*, TMED13, HIMB114, and *Actinomarina* (**Supp. Figure 9**). In turn, we also observed population-level structure that was only detected in one time series (**Figure 3, Supp. Figure 9**).

### Candidate adaptive genes in the two time series

Genes showing elevated pN/pS may contain nonsynonymous variation associated with adaptation to particular environmental conditions, such as colder or warmer waters. Here, we used the long-read MAGs as species-level genomic references and mapped short-read metagenomes from the BBMO and SOLA time series to characterize gene-level polymorphism through SNV analysis. Variation in pN/pS among samples may therefore reflect changes in the relative contribution of strains or ecotypes carrying different genetic variants through time. Candidate adaptive genes were defined as genes exhibiting a mean pN/pS ratio >0.8 across the complete time series (12 years for BBMO, 7 years for SOLA, or their combination in the MIX dataset), together with pN/pS >1 in at least one individual sample. Mean pN/pS was calculated after excluding NA values; consequently, genes represented in only a small number of samples could still satisfy this criterion if their available pN/pS values were sufficiently high.

Candidate adaptive genes were not detected in three MAGs with limited metagenomic coverage. In the independent analyses, 664 candidate adaptive genes were detected at BBMO and 480 at SOLA, yielding 1,144 candidate adaptive-gene detections across 27 MAGs. These corresponded to 827 distinct genes out of the 52,232 CDS encoded collectively by these MAGs (∼1.6%), of which 317 were detected in both time series, 347 only at BBMO, and 163 only at SOLA (**Table 3**). A total of 22 MAGs contained one or more candidate adaptive genes detected in both time series, while four MAGs displayed adaptive genes only in BBMO (**Table 3; Supp. Table 3**).

**Table 3.**
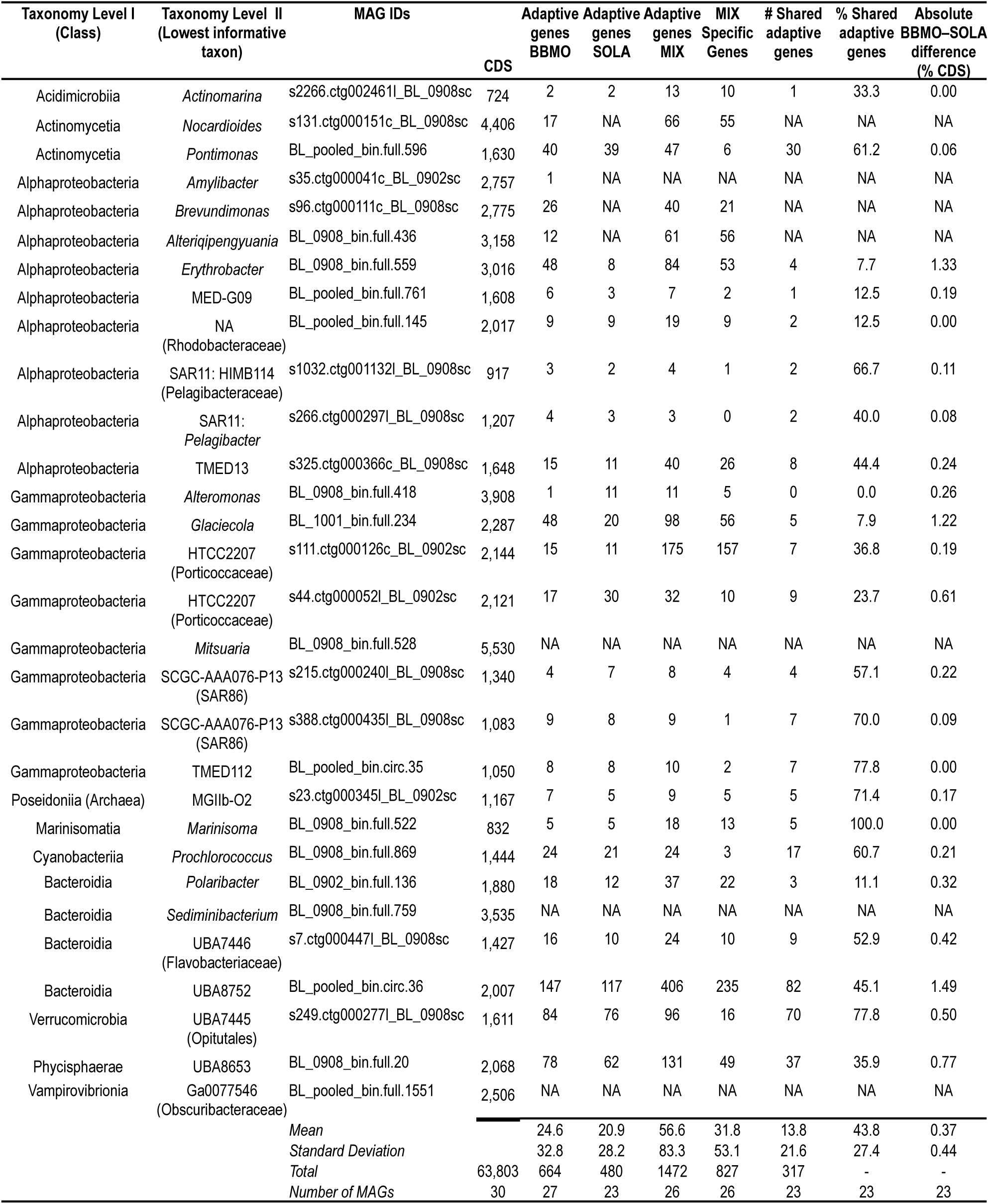
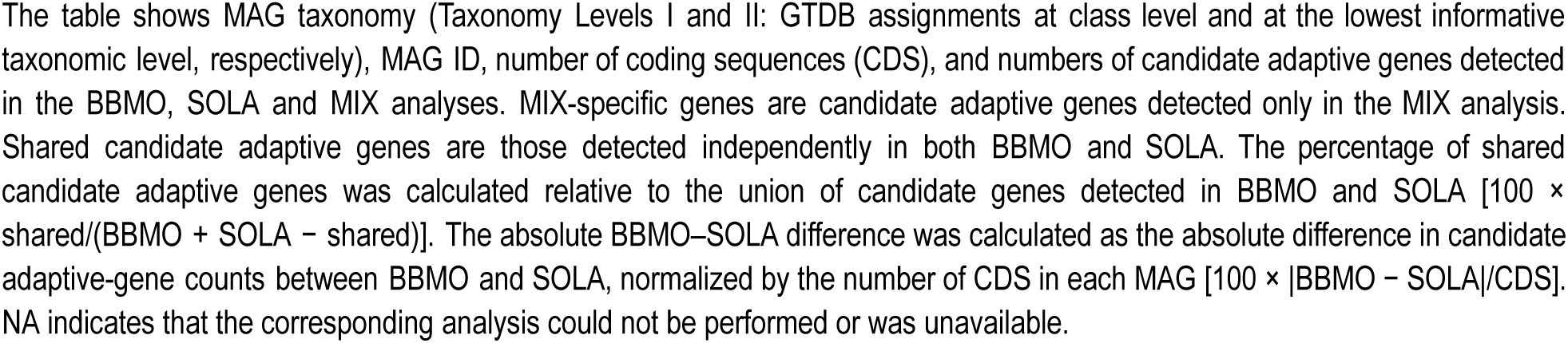
Detected candidate adaptive genes in each of the 30 long-read MAGs.

In the MIX dataset, 1,472 candidate adaptive genes were identified across the 26 MAGs that could be analyzed, which collectively encoded 49,475 CDS; these candidate genes represented ∼3.0% of the total. Of the 1,472 candidate adaptive genes, 827 were not detected in either independent BBMO or SOLA analysis (**Table 3**). The larger number of candidate adaptive genes detected in the MIX dataset, as well as differences in gene identity among datasets, may reflect the broader environmental and population variation captured by the combined dataset and the greater statistical power provided by the larger number of samples. Together, these factors may increase sensitivity for detecting signatures of positive selection relative to the individual datasets. Overall, 1,117 of the 1,472 candidate adaptive genes identified in the MIX analysis (75.9%) were represented in both time series, whereas 355 were detected only at BBMO. In several MAGs, pN/pS >1 occurred in the same samples for multiple candidate adaptive genes, indicating coordinated temporal patterns across genes (**Supp. Figure 9**). Similarly, different candidate adaptive genes within the same MAG sometimes exhibited comparable pN/pS dynamics, potentially reflecting shared population-genetic or selective processes (**Supp. Figure 9**).

Analysis of the 27 MAGs with candidate adaptive genes shows that BBMO had slightly more positively selected genes than SOLA. The mean absolute BBMO–SOLA difference in candidate adaptive-gene counts, normalized by the number of CDS per MAG, was 0.37% (**Table 3**). Across the three analyses, the number of candidate adaptive genes per MAG varied substantially, ranging from 1 to 406 (BBMO mean = 24.6, SD = 32.8; SOLA mean = 20.9, SD = 28.2; MIX mean = 56.6, SD = 83.3; **Table 3**). In most cases, the number of predicted adaptive genes per MAG was consistent across time series (**Table 3**). However, differences occurred in some cases, such as in *Brevundimonas* and *Alteriqipengyuania* (26 and 12 candidate adaptive genes detected only in BBMO, respectively). Furthermore, *Erythrobacter* displayed 48 adaptive genes in BBMO and only 8 in SOLA, with four being shared. Similarly, *Glaciecola* had 48 adaptive genes in BBMO and 20 in SOLA, with five shared genes. One Porticoccaceae MAG had 17 adaptive genes in BBMO and 30 in SOLA, with nine shared (**Table 3**). Two pairs of MAGs, each sharing the same taxonomic classification (HTCC2207 Porticoccaceae and SCGC-AAA076-P13 SAR86), displayed different amounts of adaptive genes in each time series as well as different numbers of shared adaptive genes, supporting their genomic differentiation (**Table 3**).

Functionally annotated shared candidate adaptive genes exhibited diverse functions (**Supp. Table 4**). Adaptive genes encoding methyltransferase domains (PF08241.11 and PF13649.5), involved in methylation processes that affect genome defense and gene regulation^67^, were detected in *Prochlorococcus* (BL_0908_bin.full.869) and TMED13 Alphaproteobacteria (s325.ctg000366c_BL_0908sc). Similarly, adaptive genes annotated as PF02321.17 (outer membrane efflux protein), key components in the efflux of toxic substances and antibiotic resistance^68^, were present in two MAGs: UBA8752 Bacteroidia (BL_pooled_bin.circ.36) and HTCC2207 Porticoccaceae (s44.ctg000052l_BL_0902sc). Furthermore, an open reading frame annotated with both PF13185 (GAF domain) and K08968 (L-methionine (R)-S-oxide reductase), which jointly play a role in oxidative stress response^69^, was detected in UBA8752 Bacteroidia (BL_pooled_bin.circ.36) and HTCC2207 Porticoccaceae (s44.ctg000052l_BL_0902sc). Additionally, the AMP-binding enzyme (PF00501), which participates in various metabolic pathways, including fatty acid metabolism, amino acid activation, and secondary metabolite biosynthesis, was detected in UBA8752 Bacteroidia (BL_pooled_bin.circ.36) and HTCC2207 Porticoccaceae (s111.ctg000126c_BL_0902sc). Genes annotated as PF01370 (NAD-dependent epimerase/dehydratase family) and PF16363 (NAD(P)-binding domain common to GDP-mannose 4,6-dehydratase and other dehydrogenases) were found in both *Marinisoma* sp. (BL_0908_bin.full.522) and HTCC2207 Porticoccaceae (s111.ctg000126c_BL_0902sc). The PF01370 family includes enzymes that use NAD as a cofactor to act on nucleotide-sugar substrates, playing crucial roles in carbohydrate metabolism and the modification of sugars^70,71^. PF16363 represents NAD- and NADP-binding domains with a core Rossmann-type fold, found in various enzymes involved in redox reactions in carbohydrate metabolism, including GDP-mannose 4,6-dehydratase. Altogether, these patterns suggest that specific genomic functions related to methylation, detoxification, oxidative stress response, nucleotide-sugar metabolism, NAD(P)-dependent redox processes, and other metabolic pathways have been or are under positive selection across different MAGs, reflecting specific selective pressures. Positive selection on the same functional genes across taxa suggests that these MAG populations may have developed comparable adaptations to environmental heterogeneity.

### pN/pS dynamics

We analyzed pN/pS ratios of candidate adaptive genes across 12 and 7 years for BBMO and SOLA, respectively, for all analyzable MAGs **(Supp. Figure 11)**. In most MAGs, candidate adaptive genes showed temporal fluctuations in pN/pS that were often regular, alternating between pN/pS >1 and NA values (pS = 0 in pN/pS calculations), consistent with temporal changes in the variants or strains contributing to these genes, although an undefined pN/pS does not by itself indicate gene or strain absence (**Figure 4A; Supp. Figure 11**). In contrast, other candidate adaptive genes showed irregular fluctuations (**Figure 4B**; **Supp. Figure 11)**. Some candidate adaptive genes exhibited specific pN/pS values only during certain periods of the analyzed time series. For example, in one SAR11 MAG (s266.ctg000297l_BL_0908sc), the candidate adaptive gene 1_994 (**Figure 4C**) showed fluctuating pN/pS values above one only in the early years of the studied period in SOLA. In BBMO, pN/pS values during the corresponding period generally remained just below one, whereas defined pN/pS estimates were unavailable for most of the remaining study period (**Supp. Figure 11**). This pattern is consistent with a temporal change in the variants or strains contributing to this gene. Similarly, in BBMO, several candidate adaptive genes in the *Brevundimonas* MAG (s96.ctg000111c_BL_0908sc), including 1_1097, 1_1275, and 1_1925, showed fluctuating pN/pS values at or above 1 during the initial years of the time series, whereas defined pN/pS estimates were unavailable in later years (**Supp. Figure 11**). In contrast, gene 1_2120 from the same *Brevundimonas* MAG displayed pN/pS values near or above one only in the final years of the studied period in BBMO (**Figure 4D**). This contrast indicates different temporal pN/pS profiles among genes from the same MAG, consistent with shifts in the variants or strains contributing to these genes.

**Figure 4.**
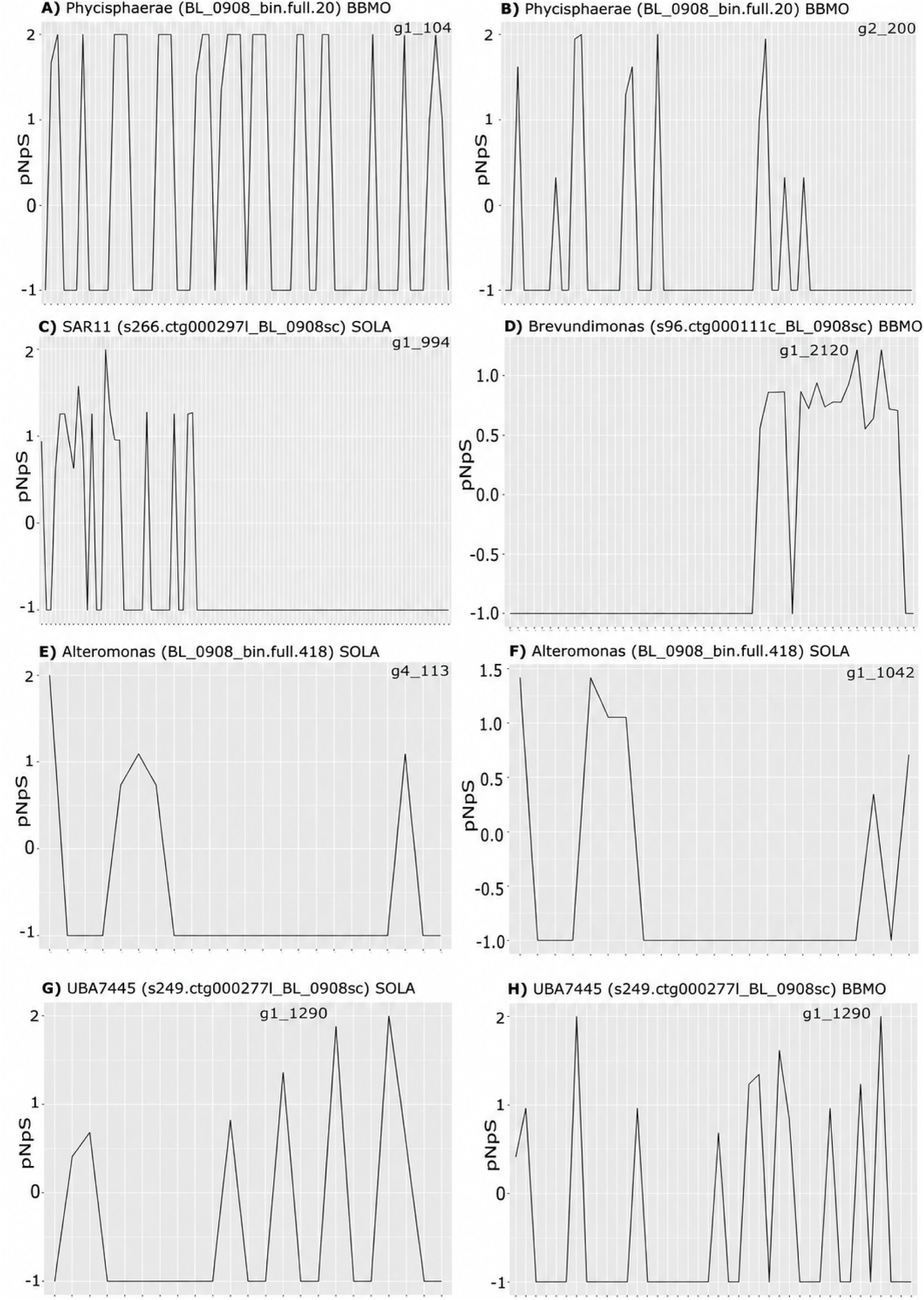
Representative temporal pN/pS patterns of candidate adaptive genes. The x-axes show samples from the BBMO or SOLA time series in chronological order, and the y-axes show pN/pS values. Values >2 were capped at 2 for visualization, whereas undefined pN/pS values resulting from pS = 0 were plotted as −1. Each panel shows the temporal pN/pS profile of a specific gene from the indicated MAG and time series; gene identifiers are shown in the upper-right corner. **(A)** A gene with a regular temporal pN/pS pattern. **(B)** A gene with an irregular pattern. **(C)** A SAR11 gene showing pN/pS >1 primarily during the early part of the SOLA time series. **(D)** A *Brevundimonas* gene showing elevated pN/pS primarily during the later part of the BBMO time series. **(E, F)** Two candidate adaptive genes from the same *Alteromonas* MAG showing broadly synchronized, irregular pN/pS fluctuations in SOLA. **(G, H)** Gene 1_1290 from the UBA7445 Opitutales MAG, showing a slightly increasing pN/pS trend over time in SOLA (G) but no corresponding trend in BBMO (H).

Within particular MAGs, multiple candidate adaptive genes exhibited regular, largely synchronized fluctuations in pN/pS, consistent with coordinated temporal changes in the variants or strains contributing to these genes at particular seasons or time points. For example, the UBA8653 Phycisphaerae MAG (BL_0908_bin.full.20) in BBMO exhibited several candidate adaptive genes with comparable and predominantly regular pN/pS fluctuations over time (**Supp. Figure 11**). We also observed irregular but synchronized pN/pS fluctuations for multiple candidate adaptive genes within the same MAG (**Supp. Figure 11**). This was the case, for example, for the *Alteromonas* MAG (BL_0908_bin.full.418) in SOLA (**Figure 4E, F; Supp. Figure 11**). These patterns are consistent with coordinated temporal shifts in the variants or strains contributing to these genes.

We investigated whether pN/pS ratios showed upward or downward trends over the study period, but most candidate adaptive genes showed no clear long-term trends. One possible exception was gene 1_1290 in the UBA7445 Opitutales (Verrucomicrobia; s249.ctg000277l_BL_0908sc) MAG, which showed an increasing pN/pS trend over time at SOLA but no corresponding trend at BBMO (**Figure 4G, H**). This increase suggests a rising ratio of nonsynonymous to synonymous polymorphism over the study period, consistent with changing selective dynamics.

## DISCUSSION

We leveraged the power of long-read sequencing to generate highly contiguous MAGs with the microbial community coverage offered by short-read shotgun metagenomics. By mapping short-read metagenomes from two neighboring long-term marine time series in the Mediterranean Sea, we reconstructed the temporal abundance variation of the analyzed MAGs across both locations over 12 (BBMO) and 7 (SOLA) years. We found that specific MAGs (e.g., Bacteroidia UBA8752, *Pontimonas*, *Prochlorococcus*) exhibited rhythmic and predominantly synchronous dynamics in both locations. Other MAGs displayed rhythmic dynamics that were slightly asynchronous (e.g., TMED112, Rhodobacteraceae). This suggests that members of the same or closely related populations were present in both locations at comparable time points. This may be explained by the geographic proximity, hydrodynamic connectivity, and comparable environmental conditions in both time series. The rhythmic and synchronous presence of MAGs in both locations aligns with the idea that the distribution of certain microbes follows large-scale patches across the ocean^72^. Therefore, patches of some microbial species could cover both BBMO and SOLA during specific periods of the year. Other MAGs, possibly conditionally rare or blooming taxa^66^ (e.g., *Mitsuaria*, *Sediminibacterium*), displayed arrhythmic, non-synchronous dynamics, typically appearing sporadically during specific time points. This may reflect local abiotic and biotic conditions that drive blooms or promote copiotroph growth, and that can vary depending on the specific location^73–75^. At SOLA, for instance, pronounced freshwater influxes from winter flash floods are known to impact microbial community structure^76,77^.

Genome-wide nucleotide diversity (π), measuring the genetic variation within a population, displayed rhythmic patterns in specific MAGs. These appeared to follow, in multiple cases, seasonal fluctuations in abundance and tended to remain consistent over time (no clear trends were detected). This suggests that interannual population diversity remained stable, with no clear evidence of increased diversity from factors such as immigration or a decline due to local extinctions. The latter is relevant in the context of global change, which may reduce the genetic diversity of populations^78^. We also observed contrasting examples where π did not display rhythmic patterns, remaining relatively constant during most of the studied period and across seasons (e.g., *Pelagibacter* [s266.ctg000297l_BL_0908sc], SAR86 [s215.ctg000240l_BL_0908sc, and s388.ctg000435l_BL_0908sc at BBMO]). In these MAGs, subpopulation structure was detected and seemed to be related to cold and warm waters. This suggests that closely related strains with similar π replaced each other during different periods of the year. Similarly, multiple genotypes can coexist within the same SAR11 population and remain stable over time, which could be attributed to widespread homologous recombination^79^. Multiple MAGs exhibited non-synchronous nucleotide diversity (π) dynamics between the two time series, with most showing different average Fst values. This highlights the inherent complexity and context dependency of the temporal dynamics of taxa that constitute ocean microbiomes.

### Population differentiation

The variability in long-term genome-wide population differentiation (i.e., Fst variance) among MAGs suggests that distinct eco-evolutionary processes shape these populations. The comparatively low average Fst values and small standard deviations observed in some MAGs, such as *Pelagibacter* (MAG 11; **Table 1**), in both time series indicate limited genome-wide population differentiation over time. Although this MAG peaks in summer, its year-round detection in both time series indicates that closely related strains persist across seasons, without constituting significantly different populations. SAR11 populations are known to undergo widespread recombination among distantly related members, which limits genetic divergence^79^. Freshwater relatives of SAR11 (LD12; *Ca.* Fonsibacter sp.) also displayed comparatively low Fst values (Fst < 0.04) over 7 years in Lake Erken, Sweden^80^. In contrast, MAGs like Verrucomicrobia (UBA7445) and *Glaciecola* demonstrated higher mean Fst values and greater variability, indicating more pronounced and fluctuating population differentiation over time. Differences in seasonal selective pressures, the introduction of strains via currents, and stochastic events may contribute to the dominance of distinct ecotypes or strains over time. Such effects may manifest both across different seasons or environmental states (i.e., warm vs. cold waters) [e.g., *Glaciecola*] and within the same season or environmental states (e.g., UBA7445 Verrucomicrobia). This is consistent with Fst clustering analyses that revealed potential population structure in both *Glaciecola* and UBA7445. These findings align with the expanding body of evidence supporting diverse population structures in marine microbes across spatiotemporal scales^17^, highlighting the differences in microdiversity between lineages^79,80^.

Marine microbial populations may show directional change over contemporary timescales^28,81^. Our analysis of 27 MAGs across 12 and 7 years in the two time series indicates minimal or no significant increase in genome-wide population differentiation for most cases. Nevertheless, five MAGs, affiliated with *Nocardioides*, *Erythrobacter*, *Brevundimonas*, *Pelagibacter*, and *Alteriqipengyuania*, showed a marked increase in genome-wide differentiation over time. Such population divergence could reflect strain replacement due to selection or drift, or the slow evolutionary differentiation of populations due to specific selective pressures^28,81^. Hoetzinger and colleagues^80^ found comparable patterns of increasing Fst over time for four microbial species in five continental waterbodies when considering different timescales (up to 7 years). At the higher end, *Polynucleobacter paneuropaeus* exhibited an Fst change of ∼ 0.4 per 1,000 days^80^. In our dataset, *Nocardioides* had the highest rate of Fst change, with an Fst of ∼0.2 per 1,000 days. In turn, Hoetzinger *et al.*,^80^ report a very low change in Fst (0.043) over ∼2,500 days in LD12, pointing to a slow and steady divergence. In our work, a marine relative of LD12, SAR11, also displayed a very low estimated change in Fst (0.03) over ∼4,300 days, with R^2^ = 0.164 and a Mantel Pearson correlation of 0.4. This suggests that populations of the marine SAR11 MAG and its freshwater relative LD12 may diverge slowly over contemporary timescales. Similarly, a recent study in Lake Mendota (Madison, USA) provided evidence of gradual changes in strain composition over a 20-year period, aligning with the patterns we observed in specific MAGs across both time series^27^. Overall, these findings suggest that population differentiation in environmental microbes may exhibit trends on contemporary timescales, emphasizing the importance of monitoring such changes to better understand microbial responses to ongoing global change^82^.

Most MAGs showed significant differences in average Fst values between BBMO and SOLA, which may partly reflect the different time spans analyzed (12 vs. 7 years) as well as slight differences in sampling days at each location. These Fst differences also suggest that local environmental conditions play an important role in shaping microbial population structures, even in relatively close marine locations (∼150 km apart) that are connected by currents and experience similar environmental variability. Specifically, the Northern Current (NC) may promote connectivity between BBMO and SOLA and facilitate microbial dispersal, with seasonally variable intensity (**Supp. Figure 12**). Nevertheless, unlike BBMO, SOLA experiences sporadic winter storms that transport nutrients from the sediments to the water column. Additionally, freshwater inputs from nearby rivers contribute to nutrient enrichment during flash floods^34^. These differences may drive differential environmental selection in both locations, potentially resulting in distinct resident populations developing at each site. Stochastic biotic and abiotic factors, including the recruitment of functionally redundant taxa from the rare biosphere or seed bank^83,84^, may also play a role in amplifying the differences between resident populations at BBMO and SOLA.

We also identified several instances where the same resident population appeared in both BBMO and SOLA. This was evident in UPGMA analyses, where the population most similar (based on Fst values) to a given resident population in BBMO was found in SOLA, and vice versa. This pattern may reflect the connectivity between the two locations via ocean currents (**Supp. Figure 12**) and is consistent with previous estimates suggesting that populations of abundant marine microbes can span large geographic areas. For example, *Prochlorococcus* is expected to be well-mixed within large water parcels (∼10 km² in area and 3 m in depth) over ecologically relevant time scales of approximately one week^23^. These abundant microbes were also predicted to disperse across large ocean provinces through turbulence and currents over weeks to months^85^. Lastly, similar environmental selection pressures at both sites could account for the presence of the same populations in both locations.

The observed patterns of seasonal population structure across the studied MAGs point to complex ecological adaptations in the sea primarily linked to temperature. Temperature is a well-established factor influencing microbial distribution and diversity, with numerous studies demonstrating its role in shaping community composition^4,11,86^. The detection of distinct candidate populations in cold and warm waters in 8 MAGs, such as *Alteromonas* and *Brevundimonas*, suggests the evolution of thermally adapted ecotypes, similar to what has been observed in *Prochlorococcus*^87,88^. Particularly interesting was the detection of population structure in ubiquitous marine taxa like SAR11 and SAR86, which aligns with previous observations of fine-scale niche partitioning in these groups^25,89^. The presence of distinct candidate populations within the same thermal regime in BBMO and SOLA (i.e., warm or cold waters) suggests that other environmental factors, such as resource availability or biotic interactions, may have driven population differentiation in these locations.

### Candidate adaptive genes

Analyzing candidate adaptive genes in marine microbes over extended periods can provide insights into microbial evolutionary dynamics. When we analyzed BBMO and SOLA independently, we identified 1,144 candidate adaptive-gene detections across 27 MAGs, corresponding to 827 distinct genes, of which 317 were detected in both time series. These 827 distinct candidate adaptive genes represented ∼1.6% of the total coding sequences. When BBMO and SOLA were analyzed jointly, we detected 1,472 candidate adaptive genes among the 26 MAGs that could be analyzed, representing ∼3.0% of the coding sequences in those MAGs (**Table 3**). This higher percentage may partly reflect increased detection sensitivity resulting from the larger number of samples included in the joint analysis. Overall, the small proportion of genes showing signatures of positive selection is consistent with previous observations of generally low pN/pS ratios across 66 species in the human gut microbiome, suggesting that purifying selection predominates across much of the genome^90^. Similarly, Zhao *et al*.^91^ reported evidence of parallel adaptive evolution in 16 genes of *Bacteroides fragilis*, an important species in the human gut microbiome. Together, these observations are consistent with adaptive evolution being concentrated in a relatively limited subset of genes within microbial genomes.

The overlap in detected adaptive genes between sites (38.3%) indicates similar adaptations across MAGs in both time series. Furthermore, this is consistent with both the potential connectivity between populations at the two locations via currents and the influence of similar selective pressures. The detection of multiple genes featuring elevated pN/pS in specific samples, which in several cases clustered by Fst similarity, points to the presence of adaptations in several genes in ecotypes or strains. Alternatively, selection could have acted on specific adaptive genes in a given strain, while other adaptive genes could be hitchhiking (a.k.a. codrivers)^92,93^.

While the generally consistent pattern of adaptive gene numbers between time series for most MAGs indicates similar adaptations in both sites, the marked differences in adaptive gene counts observed in some taxa (e.g., *Brevundimonas*, *Erythrobacter*) suggest locally specific selective pressures or different evolutionary trajectories. This variation may also reflect the presence of locally adapted populations at each time series, such as ecotypes differing in genomic content^23,89^. Given that long-read MAGs are effective at capturing the flexible genome^30^, it is plausible that some MAGs retrieved from BBMO represent ecotypes containing genes that were absent or underrepresented in the SOLA samples (e.g., in *Brevundimonas*, *Alteriqipengyuania*, *Erythrobacter*, and *Glaciecola*, which displayed more adaptive genes in BBMO than in SOLA).

Annotation of the shared candidate adaptive genes revealed diverse functions. Adaptive methyltransferase domains (PF08241 and PF13649) suggest that methylation may play a role in seasonal population adaptation, potentially contributing to genome defense and gene regulation in response to environmental fluctuations^67^. Outer membrane efflux proteins containing the PF02321 domain point to selection for tolerance to diverse stressors, including toxins, antibiotics, osmotic stress, and potentially phage infection, which may vary temporally^68^. An adaptive gene annotated with both the GAF domain (PF13185) and methionine-(R)-sulfoxide reductase activity (K08968) underscores the potential for diverse oxidative stress responses across populations. This aligns with the importance of oxidative stress mitigation in marine surface waters, where high levels of reactive oxygen species, driven by solar radiation and photosynthetic activity, may challenge microbial communities^94,95^.

Positively selected AMP-binding enzymes (PF00501) likely reflect adaptive responses to environmental fluctuations in marine ecosystems. These enzymes, such as acyl-CoA synthetases, may play a role in fatty acid metabolism^96^, enabling modifications in fatty acid composition in response to changes in temperature or nutrients. Marine bacteria may adjust their fatty acid profiles to maintain membrane fluidity and functionality under varying environmental conditions^97,98^. Positively selected genes in the NAD-dependent epimerase/dehydratase family, including the PF01370 protein domain, which contributes to the activity of enzymes like GDP-mannose 4,6-dehydratase (containing both PF01370 and PF16363 domains), are involved in nucleotide-sugar metabolism^70,71^. These genes indicate potential adaptations for carbohydrate metabolism and modifications of structural sugars critical for extracellular polymeric substances (EPS) and cell wall biosynthesis^71,99^, which may help modify cell surface structures in response to seasonal fluctuations in temperature, nutrient availability, or biotic pressures such as viral infection^100^. Collectively, these positively selected genes detected in both time series illustrate the spectrum of possible population-level adaptations that may enable microbes to thrive in seasonally changing environmental conditions, emphasizing the importance of genome regulation, stress tolerance, and metabolic flexibility.

### Changes in pN/pS over time

The cyclical pN/pS fluctuations observed in adaptive genes from most MAGs indicate ecotypes that thrive during particular seasons. Long-term analyses at BBMO using 16S rRNA gene metabarcoding indicated seasonal differentiation among closely related Amplicon Sequence Variants (ASVs)^10^. Our work goes further by identifying adaptive genes linked to potential ecotypes that exhibit seasonal abundance but likely share the same 16S rRNA sequence. Conversely, the irregular fluctuations in pN/pS ratios for some adaptive genes point to intermittent occurrences of specific strains. This may result from episodic environmental disturbances or immigration of strains^27^. Changes in the contribution of specific gene variants or strains, as observed for the SAR11 and *Brevundimonas* MAGs, could reflect local competitive exclusion or subtle environmental shifts favoring different strains. Furthermore, the synchronized fluctuations of multiple adaptive genes within specific MAGs support the concept of linked adaptive genes within ecotypes undergoing coordinated changes in abundance.

While most candidate adaptive genes did not exhibit significant long-term trends in pN/pS, the increasing trend observed for gene 1_1290 in the UBA7445 Opitutales MAG at SOLA is consistent with either ongoing positive selection or relaxed purifying selection. Such an upward trend may reflect an increasing relative contribution of nonsynonymous to synonymous polymorphism and could be consistent with selection favoring nonsynonymous variants under changing environmental conditions, including warming. A study analyzing ∼30 years of data (since 1992) from Mediterranean Sea surface waters revealed that the region encompassing BBMO and SOLA is warming at a rate of 3.9 ± 0.6 °C per century, alongside a marked increase in salinity throughout the water column^101^. Another study using Argo float data reveals warming trends in the Mediterranean Sea, particularly in the Western Mediterranean (5-700 m depth), where temperatures increase at a rate of 0.070 ± 0.015°C per year^102^. Furthermore, the Mediterranean Sea is experiencing more frequent marine heatwaves that can trigger mass mortality events in multicellular organisms^103^, while their impact on microbes remains unclear. Altogether, these and other impacts of global change may be exerting selective pressures on the microbiota of the Mediterranean Sea, driving shifts in allele frequencies or altering the relative abundance of different ecotypes, with potential cascading effects on ecosystem function.

### Technical considerations

The metagenomic approach used in this study, combining long- and short-read DNA sequencing, allowed us to explore the long-term population structure of 30 prokaryotic genomes from two neighboring coastal locations in the Mediterranean Sea, offering insights into their potential adaptations. Utilizing the PacBio Sequel II platform, we generated 30 highly contiguous long-read MAGs. Unlike other sequencing methods, PacBio can assemble MAGs from relatively few DNA molecules^104^, offering greater accuracy than short-read MAGs^30,105,106^ and allowing for more accurate metagenome-based population genomics analyses^17^. The main reason for this is related to a potential reduction of chimerism in long-read MAGs compared to short-read MAGs, as the former is assembled from fewer DNA strands than the latter. Our results indicate that combining assembly and binning strategies enhances the recovery of MAGs from marine metagenomic samples. Specifically, a number of MAGs were only recovered as single-contig (no-binning), while others were only recovered after binning contigs from individual samples or from their co-assembly. Furthermore, the retrieval of conditionally rare taxa from individual samples where they were abundant supports the use of PacBio to retrieve MAGs that are usually rare and that could be missed in binning approaches using short reads.

## CONCLUSIONS

Our results show that combining long-read MAGs as genomic references with dense short-read metagenomic time series can uncover population-genomic dynamics that are not apparent from species-level patterns alone. We identified recurrent seasonal population structure, site-specific differences in genome-wide differentiation, and increasing differentiation with temporal distance in a subset of marine microbes. We also show that long-read metagenomics can recover highly contiguous genomes from taxa that remain rare for most of the time but occasionally reach high abundance. Candidate adaptive genes represented only a small fraction of coding sequences, yet their temporal dynamics revealed coordinated and lineage-specific genomic changes through time. Together, these findings provide a framework for resolving contemporary population-genomic change in marine microbes and for understanding how within-species variation contributes to long-term ocean microbiome dynamics. These approaches are particularly valuable for assessing how ocean warming and increasingly frequent marine heatwaves may reshape microbial populations over time.

## ACKNOWLEDGMENTS

We thank all members of the BBMO sampling team (http://bbmo.icm.csic.es) as well as members of the Ecology of Marine Microbes (http://emm.icm.csic.es) group. Bioinformatics analyses were performed at the MARBITS platform of the Institut de Ciencies del Mar (ICM; http://marbits.icm.csic.es), the supercomputer Finisterrae III of the Centro de Supercomputación de Galicia (CESGA; https://www.cesga.es/), as well as the supercomputer DRAGO of the CSIC. This work acknowledges the Severo Ochoa Centre of Excellence accreditation (CEX2019-000928-S) funded by AEI 10.13039/501100011033.

## DATA AVAILABILITY

Short-read metagenomic sequences from the BBMO time series are available in the European Nucleotide Archive (ENA) under project PRJEB51979, while long-read sequences are available under PRJEB106312. Metagenomic sequences for the SOLA time series (Banyuls Bay Microbial Observatory) are available in the ENA under accession numbers PRJEB66489 and PRJEB26919. The 30 long-read MAGs are available at Zenodo (10.5281/zenodo.21517772).

## Supplementary Materials

### Supplementary Tables

**Supplementary Table 1.** Obtained HiFi reads per metagenome.

|  | PacBio HiFi<br>reads | Sum length (bp) | Minimum length<br>(bp) | Mean length<br>(bp) | Maximum length<br>(bp) |
| --- | --- | --- | --- | --- | --- |
| BL_0902 | 578,435 | 5,330,280,894 | 51 | 9,215 | 32,746 |
| BL_0908 | 1,195,188 | 10,320,701,278 | 51 | 8,635.2 | 37,636 |
| BL_1001 | 512,905 | 5,183,848,219 | 58 | 10,106.8 | 36,106 |

**Supplementary Table 2.**
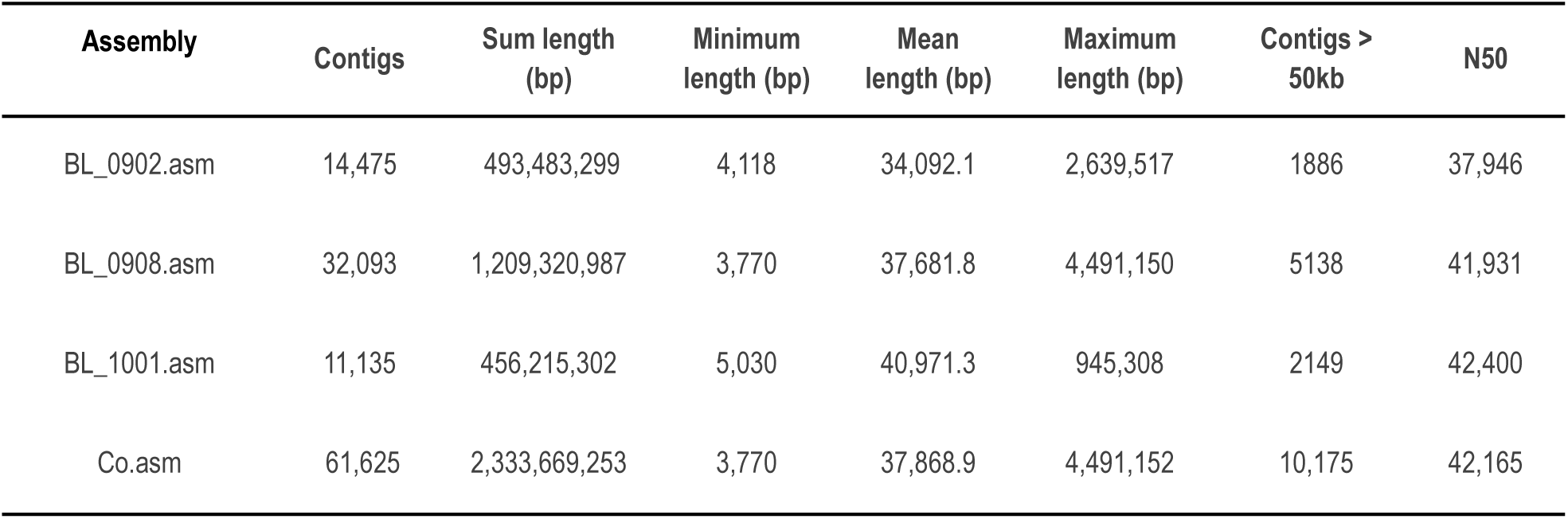
Assemblies using hifiasm-meta. The first three rows represent single-sample assemblies, while the fourth row represents the coassembly.

| Assembly | Contigs | Sum length<br>(bp) | Minimum<br>length (bp) | Mean<br>length (bp) | Maximum<br>length (bp) | Contigs ><br>50kb | N50 |
| --- | --- | --- | --- | --- | --- | --- | --- |
| BL_0902.asm | 14,475 | 493,483,299 | 4,118 | 34,092.1 | 2,639,517 | 1886 | 37,946 |
| BL_0908.asm | 32,093 | 1,209,320,987 | 3,770 | 37,681.8 | 4,491,150 | 5138 | 41,931 |
| BL_1001.asm | 11,135 | 456,215,302 | 5,030 | 40,971.3 | 945,308 | 2149 | 42,400 |
| Co.asm | 61,625 | 2,333,669,253 | 3,770 | 37,868.9 | 4,491,152 | 10,175 | 42,165 |

**Supplementary Table 3.** The gene IDs and the number of candidate adaptive genes shared between both time series for each of the 30 MAGs.

| MAG ID | Shared Candidate Adaptive Gene | Quantity |
| --- | --- | --- |
| BL_0902_bin.full.136 | 1_141, 1_322, 5_191 | 3 |
| BL_0908_bin.full.20 | 1_82, 1_104, 1_109, 1_117, 1_266, 1_369, 1_480, 1_623, 1_705, 1_709, 1_714, 1_898, 1_900, 1_908, 1_935, 1_936, 1_937, 1_1095, 1_1122, 1_1127, 2_63, 2_77, 2_115, 2_159, 2_161, 2_165, 2_209, 2_446, 2_492, 2_568, 2_575, 2_584, 2_593, 3_9, 3_21, 3_22, 3_52 | 37 |
| BL_0908_bin.full.418 | 0 | 0 |
| BL_0908_bin.full.436 | NA | NA |
| BL_0908_bin.full.522 | 1_101, 1_456, 2_86, 3_1, 3_7 | 5 |
| BL_0908_bin.full.528 | NA | NA |
| BL_0908_bin.full.559 | 1_161, 1_165, 1_263, 1_292 | 4 |
| BL_0908_bin.full.759 | NA | NA |
| BL_0908_bin.full.869 | 1_130, 1_170, 1_323, 1_640, 2_29, 2_33, 2_150, 2_166, 2_199, 2_234, 2_236, 2_525, 2_527, 2_540, 2_541, 2_543, 2_574 | 17 |
| BL_1001_bin.full.234 | 1_91, 2_194, 2_259, 5_246, 8_62 | 5 |
| BL_pooled_bin.circ.35 | 1_360, 1_411, 1_520, 1_521, 1_555, 1_878, 1_922 | 7 |
| BL_pooled_bin.circ.36 | 1_51, 1_83, 1_118, 1_129, 1_135, 1_140, 1_156, 1_159, 1_178, 1_185, 1_194, 1_201, 1_206, 1_232, 1_274, 1_316, 1_326, 1_331, 1_415, 1_471, 1_560, 1_564, 1_581, 1_590, 1_599, 1_663, 1_673, 1_689, 1_716, 1_763, 1_780, 1_795, 1_889, 1_912, 1_929, 1_935, 1_944, 1_958, 1_966, 1_982, 1_983, 1_1020, 1_1035, 1_1042, 1_1064, 1_1126, 1_1211, 1_1230, 1_1232, 1_1243, 1_1267, 1_1286, 1_1368, 1_1379, 1_1386, 1_1397, 1_1404, 1_1410, 1_1412, 1_1467, 1_1480, 1_1484, 1_1502, 1_1525, 1_1598, 1_1651, 1_1671, 1_1672, 1_1677, 1_1691, 1_1728, 1_1735, 1_1748, 1_1767, 1_1815, 1_1816, 1_1866, 1_1881, 1_1885, 1_1930, 1_1950, 1_1982 | 82 |
| BL_pooled_bin.full.145 | 1_79, 3_127 | 2 |
| BL_pooled_bin.full.1551 | NA | NA |
| BL_pooled_bin.full.596 | 1_397, 1_521, 1_554, 1_555, 1_646, 1_663, 1_843, 1_855, 1_1130, 1_1171, 1_1194, 1_1245, 1_1247, 1_1318, 1_1326, 1_1328, 1_1329, 1_1342, 1_1343, 1_1362, 1_1375, 1_1392, 1_1401, 1_1406, 1_1480, 1_1493, 1_1498, 1_1508, 1_1577, 1_1626 | 30 |
| BL_pooled_bin.full.761 | 1_216 | 1 |
| s1032.ctg001132l_BL_0908sc | 1_676, 1_700 | 2 |
| s111.ctg000126c_BL_0902sc | 1_3, 1_444, 1_591, 1_955, 1_959, 1_1915, 1_1951 | 7 |
| s131.ctg000151c_BL_0908sc | NA | NA |
| s215.ctg000240l_BL_0908sc | 1_58, 1_971, 1_1088, 1_1259 | 4 |
| s2266.ctg002461l_BL_0908sc | 1_157 | 1 |
| s23.ctg000345l_BL_0902sc | 1_158, 1_342, 1_805, 1_821, 1_1152 | 5 |
| s249.ctg000277l_BL_0908sc | 1_17, 1_250, 1_254, 1_331, 1_332, 1_402, 1_406, 1_436, 1_437, 1_477, 1_504, 1_505, 1_578, 1_582, 1_599, 1_647, 1_695, 1_696, 1_725, 1_727, 1_728, 1_732, 1_752, 1_792, 1_795, 1_845, 1_849, 1_856, 1_857, 1_858, 1_883, 1_885, 1_891, 1_892, 1_930, 1_931, 1_932, 1_935, 1_937, 1_938, 1_939, 1_940, 1_941, 1_947, 1_948, 1_955, 1_971, 1_1041, 1_1067, 1_1126, 1_1133, 1_1136, 1_1200, 1_1206, 1_1207, 1_1208, 1_1215, 1_1290, 1_1308, 1_1318, 1_1319, 1_1374, 1_1398, 1_1399, 1_1424, 1_1437, 1_1451, 1_1487, 1_1513, 1_1586 | 70 |
| s266.ctg000297l_BL_0908sc | 1_69, 1_994 | 2 |
| s325.ctg000366c_BL_0908sc | 1_67, 1_83, 1_512, 1_536, 1_812, 1_1241, 1_1303, 1_1403 | 8 |
| s35.ctg000041c_BL_0902sc | NA | NA |
| s388.ctg000435l_BL_0908sc | 1_124, 1_129, 1_272, 1_439, 1_772, 1_831, 1_1031 | 7 |
| s44.ctg000052l_BL_0902sc | 1_507, 1_807, 1_1104, 1_1513, 1_1515, 1_1520, 1_1673, 1_1708, 1_1936 | 9 |
| s7.ctg000447l_BL_0908sc | 1_110, 1_147, 1_309, 1_581, 1_736, 1_739, 1_829, 1_842, 1_1293 | 9 |
| s96.ctg000111c_BL_0908sc | NA | NA |

**Supplementary Table 4.**
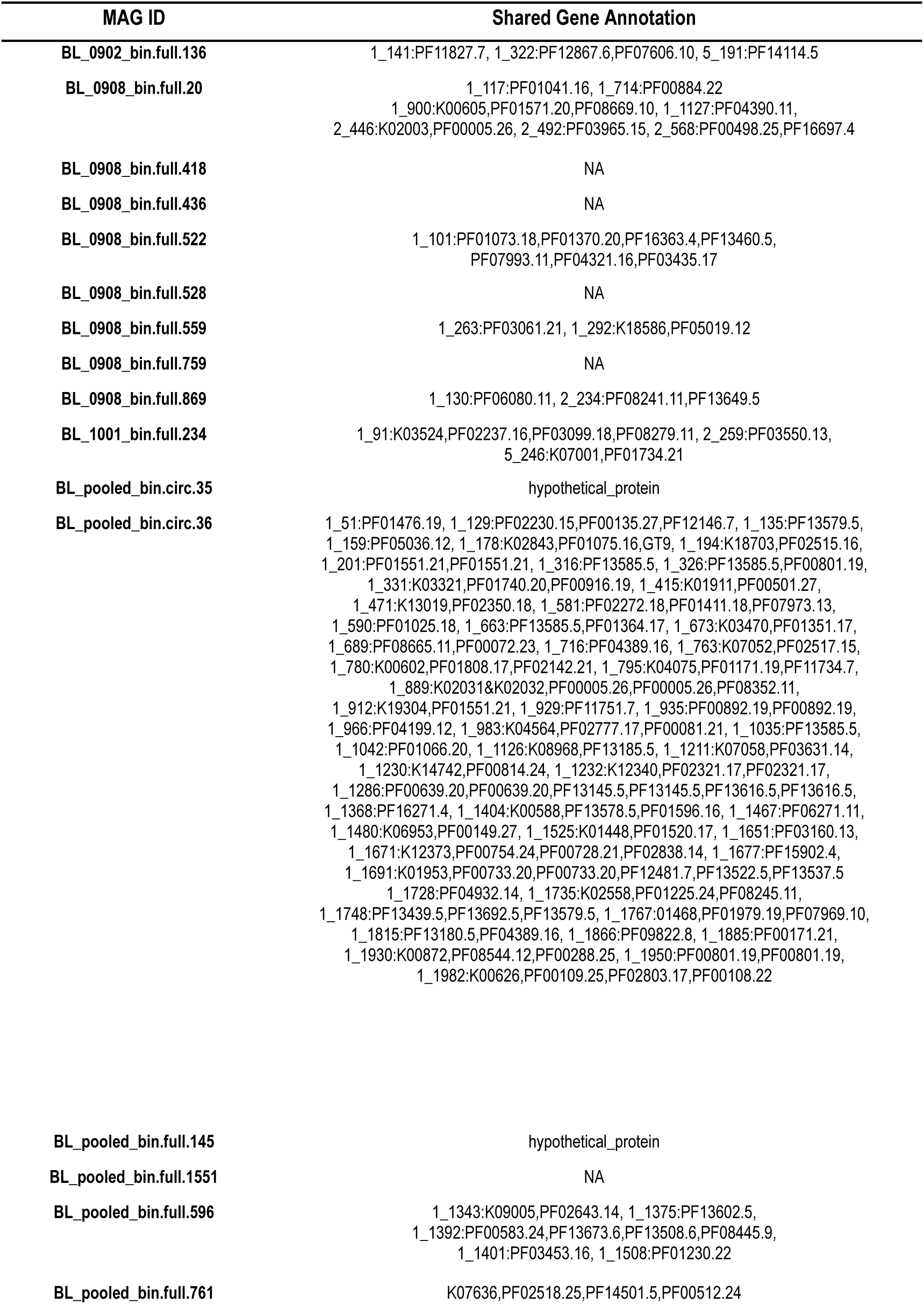
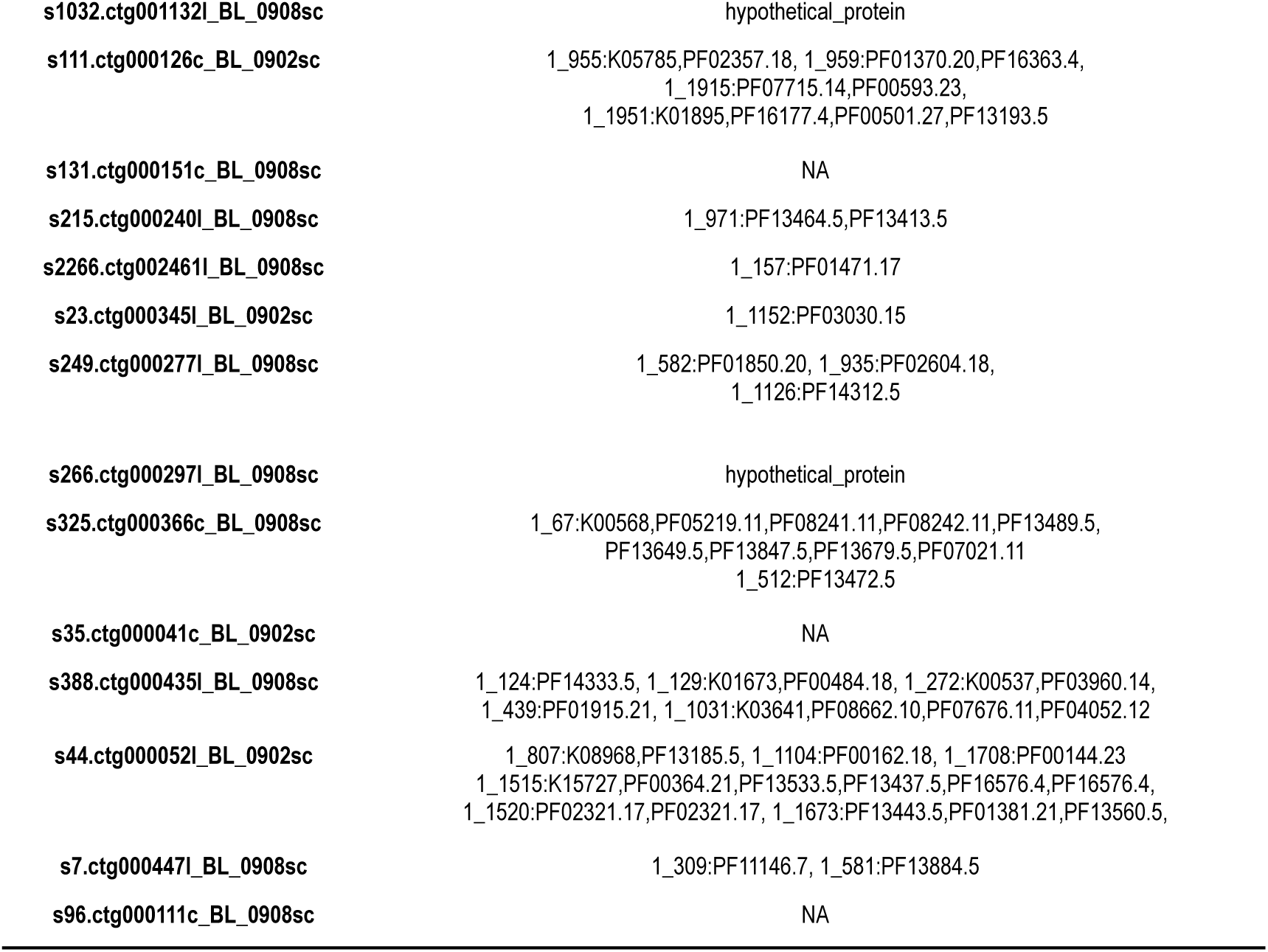
Annotation of the candidate adaptive genes present in both time series for each MAG (using KEGG and PFAM).

| MAG ID | Shared Gene Annotation |
| --- | --- |
| BL_0902_bin.full.136 | 1_141:PF11827.7, 1_322:PF12867.6,PF07606.10, 5_191:PF14114.5 |
| BL_0908_bin.full.20 | 1_117:PF01041.16, 1_714:PF00884.22<br>1_900:K00605,PF01571.20,PF08669.10, 1_1127:PF04390.11,<br>2_446:K02003,PF00005.26, 2_492:PF03965.15, 2_568:PF00498.25,PF16697.4 |
| BL_0908_bin.full.418 | NA |
| BL_0908_bin.full.436 | NA |
| BL_0908_bin.full.522 | 1_101:PF01073.18,PF01370.20,PF16363.4,PF13460.5,<br>PF07993.11,PF04321.16,PF03435.17 |
| BL_0908_bin.full.528 | NA |
| BL_0908_bin.full.559 | 1_263:PF03061.21, 1_292:K18586,PF05019.12 |
| BL_0908_bin.full.759 | NA |
| BL_0908_bin.full.869 | 1_130:PF06080.11, 2_234:PF08241.11,PF13649.5 |
| BL_1001_bin.full.234 | 1_91:K03524,PF02237.16,PF03099.18,PF08279.11, 2_259:PF03550.13,<br>5_246:K07001,PF01734.21 |
| BL_pooled_bin.circ.35 | hypothetical_protein |
| BL_pooled_bin.circ.36 | 1_51:PF01476.19, 1_129:PF02230.15,PF00135.27,PF12146.7, 1_135:PF13579.5,<br>1_159:PF05036.12, 1_178:K02843,PF01075.16,GT9, 1_194:K18703,PF02515.16,<br>1_201:PF01551.21,PF01551.21, 1_316:PF13585.5, 1_326:PF13585.5,PF00801.19,<br>1_331:K03321,PF01740.20,PF00916.19, 1_415:K01911,PF00501.27,<br>1_471:K13019,PF02350.18, 1_581:PF02272.18,PF01411.18,PF07973.13,<br>1_590:PF01025.18, 1_663:PF13585.5,PF01364.17, 1_673:K03470,PF01351.17,<br>1_689:PF08665.11,PF00072.23, 1_716:PF04389.16, 1_763:K07052,PF02517.15,<br>1_780:K00602,PF01808.17,PF02142.21, 1_795:K04075,PF01171.19,PF11734.7,<br>1_889:K02031&K02032,PF00005.26,PF00005.26,PF08352.11,<br>1_912:K19304,PF01551.21, 1_929:PF11751.7, 1_935:PF00892.19,PF00892.19,<br>1_966:PF04199.12, 1_983:K04564,PF02777.17,PF00081.21, 1_1035:PF13585.5,<br>1_1042:PF01066.20, 1_1126:K08968,PF13185.5, 1_1211:K07058,PF03631.14,<br>1_1230:K14742,PF00814.24, 1_1232:K12340,PF02321.17,PF02321.17,<br>1_1286:PF00639.20,PF00639.20,PF13145.5,PF13145.5,PF13616.5,PF13616.5,<br>1_1368:PF16271.4, 1_1404:K00588,PF13578.5,PF01596.16, 1_1467:PF06271.11,<br>1_1480:K06953,PF00149.27, 1_1525:K01448,PF01520.17, 1_1651:PF03160.13,<br>1_1671:K12373,PF00754.24,PF00728.21,PF02838.14, 1_1677:PF15902.4,<br>1_1691:K01953,PF00733.20,PF00733.20,PF12481.7,PF13522.5,PF13537.5<br>1_1728:PF04932.14, 1_1735:K02558,PF01225.24,PF08245.11,<br>1_1748:PF13439.5,PF13692.5,PF13579.5, 1_1767:01468,PF01979.19,PF07969.10,<br>1_1815:PF13180.5,PF04389.16, 1_1866:PF09822.8, 1_1885:PF00171.21,<br>1_1930:K00872,PF08544.12,PF00288.25, 1_1950:PF00801.19,PF00801.19,<br>1_1982:K00626,PF00109.25,PF02803.17,PF00108.22 |
| BL_pooled_bin.full.145 | hypothetical_protein |
| BL_pooled_bin.full.1551 | NA |
| BL_pooled_bin.full.596 | 1_1343:K09005,PF02643.14, 1_1375:PF13602.5,<br>1_1392:PF00583.24,PF13673.6,PF13508.6,PF08445.9,<br>1_1401:PF03453.16, 1_1508:PF01230.22 |
| BL_pooled_bin.full.761 | K07636,PF02518.25,PF14501.5,PF00512.24 |
| s1032.ctg001132l_BL_0908sc | hypothetical_protein |
| s111.ctg000126c_BL_0902sc | 1_955:K05785,PF02357.18, 1_959:PF01370.20,PF16363.4,<br>1_1915:PF07715.14,PF00593.23,<br>1_1951:K01895,PF16177.4,PF00501.27,PF13193.5 |
| s131.ctg000151c_BL_0908sc | NA |
| s215.ctg000240l_BL_0908sc | 1_971:PF13464.5,PF13413.5 |
| s2266.ctg002461l_BL_0908sc | 1_157:PF01471.17 |
| s23.ctg000345l_BL_0902sc | 1_1152:PF03030.15 |
| s249.ctg000277l_BL_0908sc | 1_582:PF01850.20, 1_935:PF02604.18,<br>1_1126:PF14312.5 |
| s266.ctg000297l_BL_0908sc | hypothetical_protein |
| s325.ctg000366c_BL_0908sc | 1_67:K00568,PF05219.11,PF08241.11,PF08242.11,PF13489.5,<br>PF13649.5,PF13847.5,PF13679.5,PF07021.11<br>1_512:PF13472.5 |
| s35.ctg000041c_BL_0902sc | NA |
| s388.ctg000435l_BL_0908sc | 1_124:PF14333.5, 1_129:K01673,PF00484.18, 1_272:K00537,PF03960.14,<br>1_439:PF01915.21, 1_1031:K03641,PF08662.10,PF07676.11,PF04052.12 |
| s44.ctg000052l_BL_0902sc | 1_807:K08968,PF13185.5, 1_1104:PF00162.18, 1_1708:PF00144.23<br>1_1515:K15727,PF00364.21,PF13533.5,PF13437.5,PF16576.4,PF16576.4,<br>1_1520:PF02321.17,PF02321.17, 1_1673:PF13443.5,PF01381.21,PF13560.5, |
| s7.ctg000447l_BL_0908sc | 1_309:PF11146.7, 1_581:PF13884.5 |
| s96.ctg000111c_BL_0908sc | NA |

**Supplementary Table 5.**
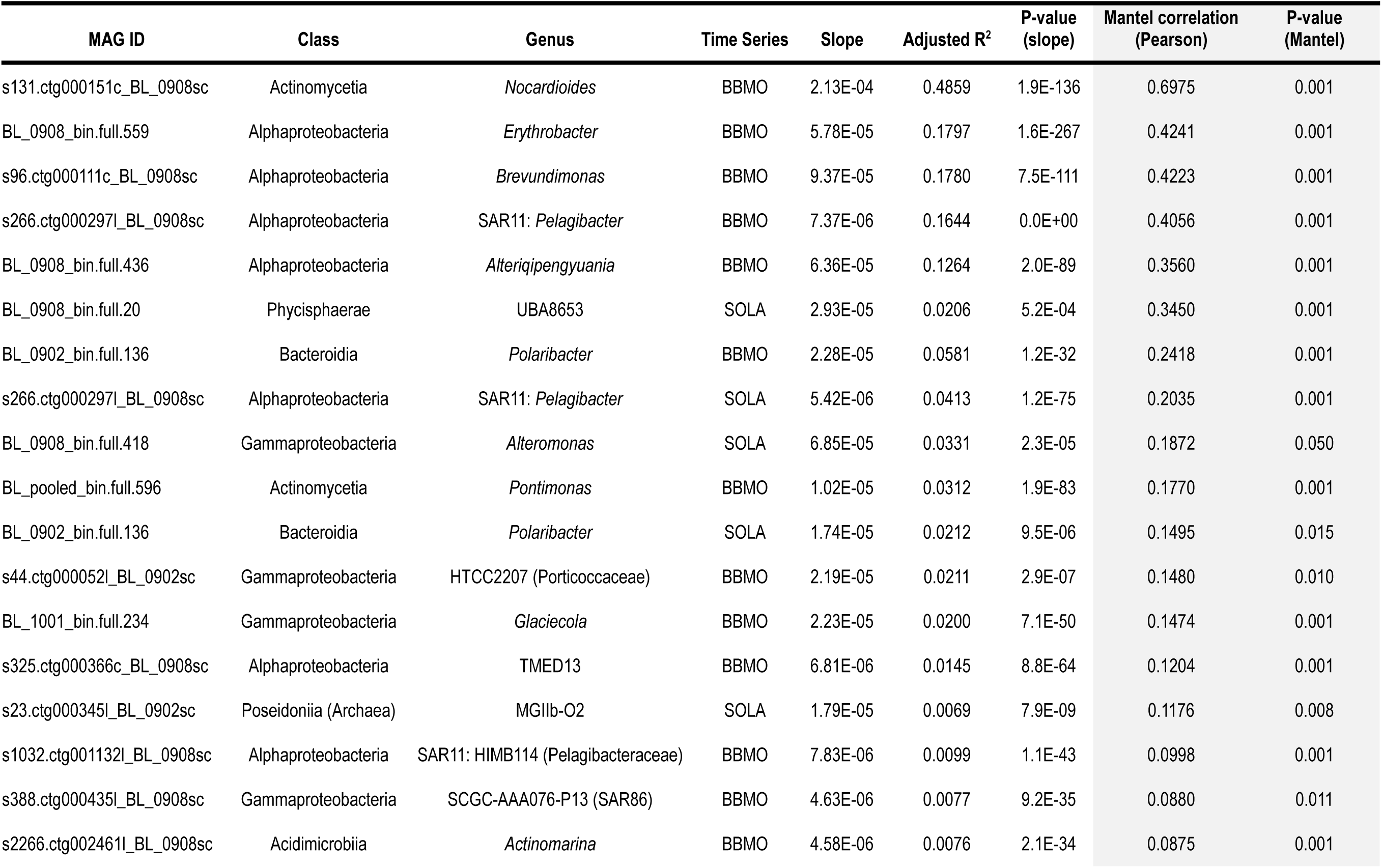
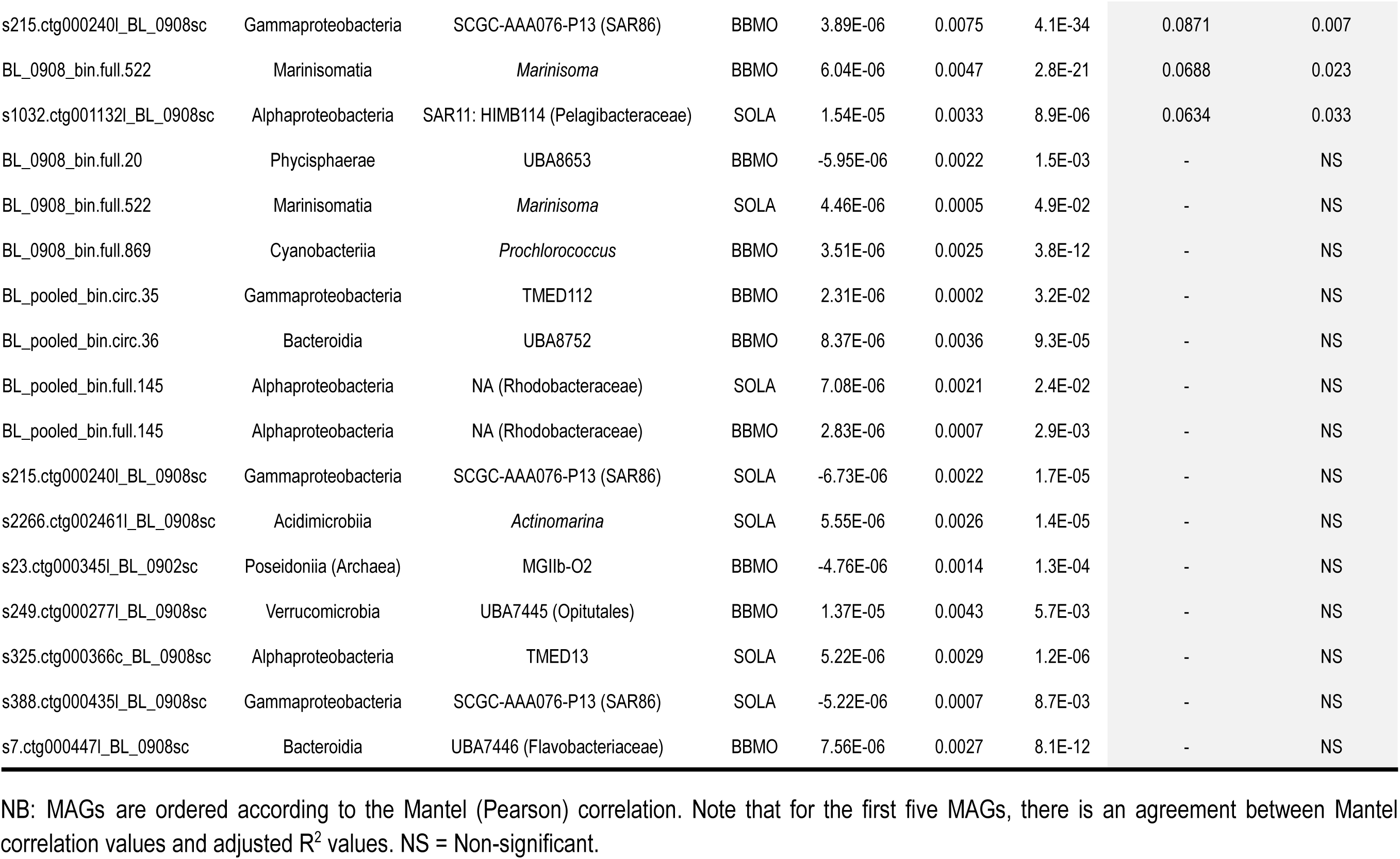
Regression and Mantel test results of the analysis of Fst vs. time between samples (in days). Results are presented for the 24 MAGs that yielded significant values in at least one of the tests.

### Supplementary Figures & Figure Legends

**Supplementary Figure 1.**
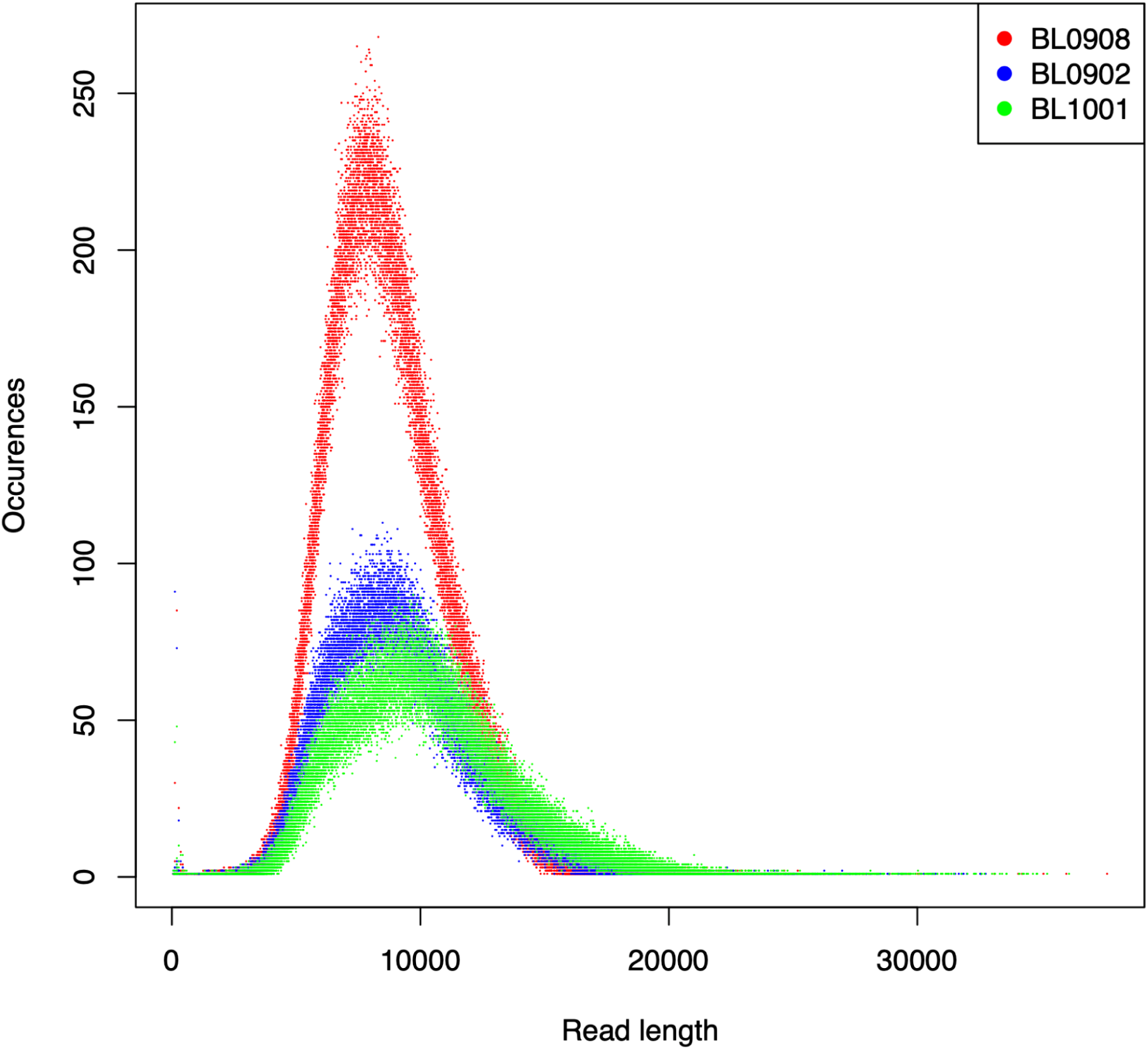
Read lengths. Distribution of PacBio HiFi read lengths for the three samples: February (BL0902) and August (BL0908) 2009, and January 2010 (BL1001).

**Supplementary Figure 2.**
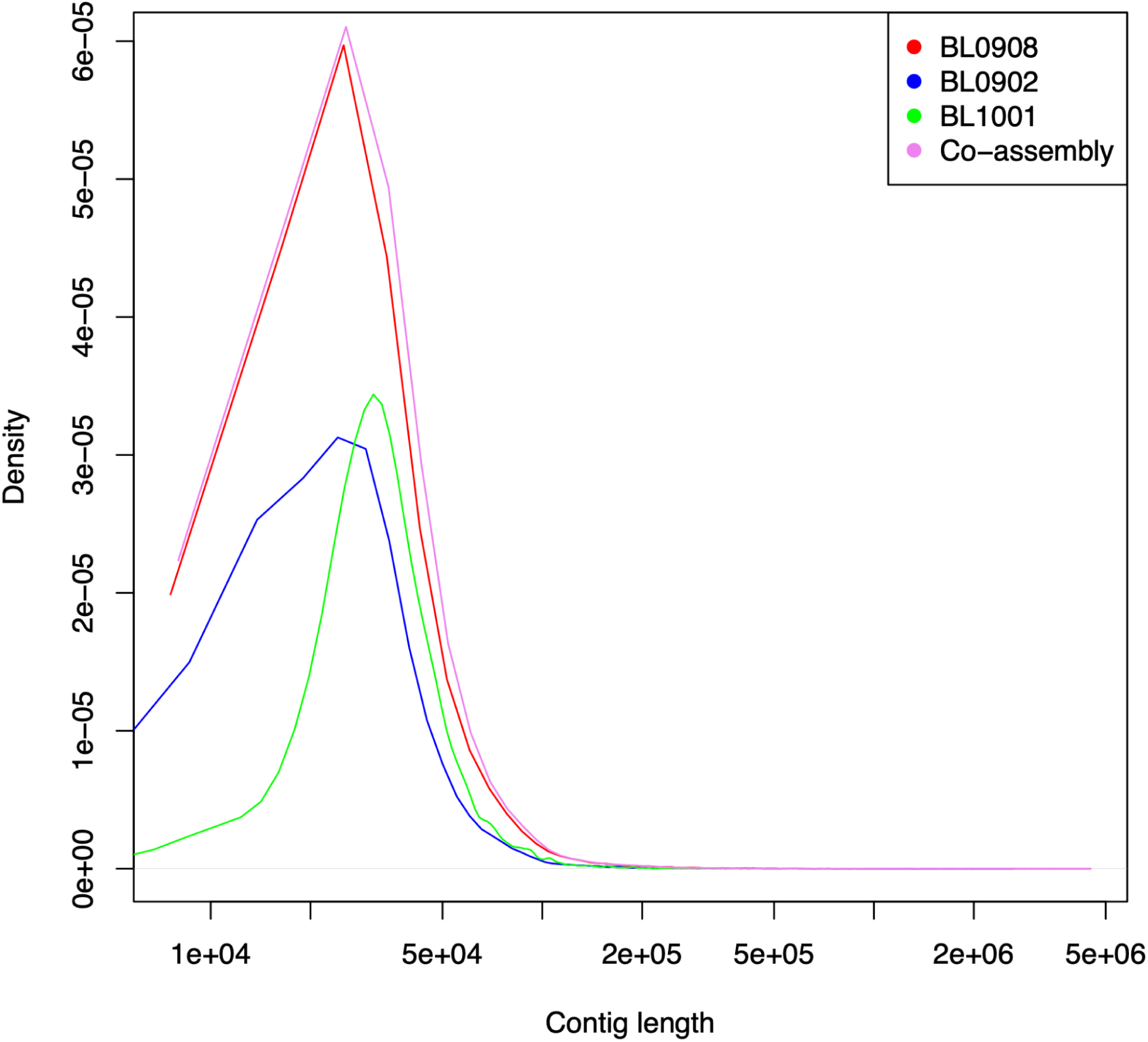
Contig lengths. Distribution of contig lengths for the three samples, February (BL0902) and August (BL0908) 2009, and January 2010 (BL1001), as well as their co-assembly.

**Supplementary Figure 3.**
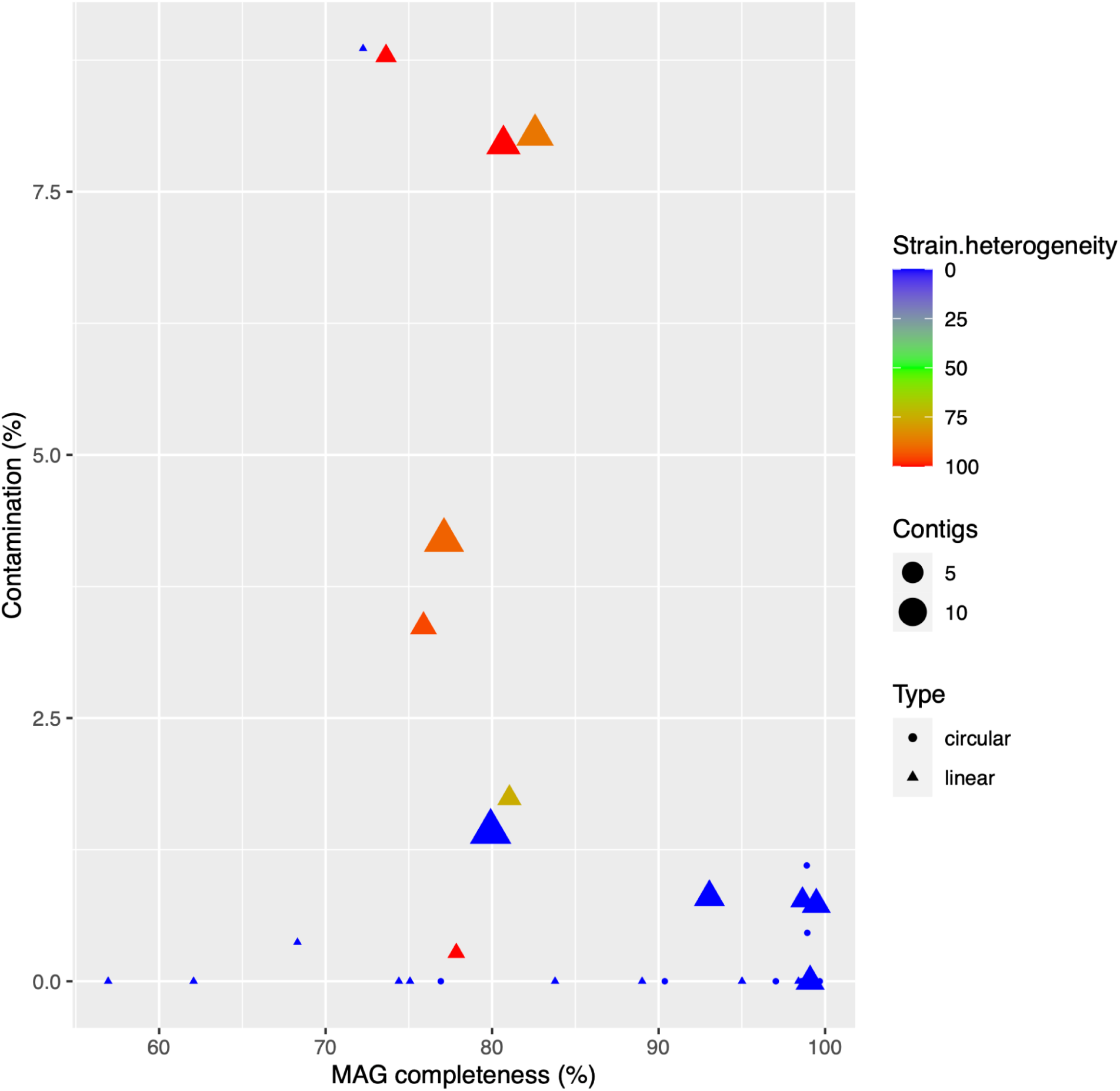
Main features of the 30 selected PacBio HiFi BBMO MAGs. The figure shows each MAG in terms of its completeness, contamination, strain heterogeneity, and number of contigs. The shapes indicate whether a MAG is circular or linear.

**Supplementary Figure 4.**
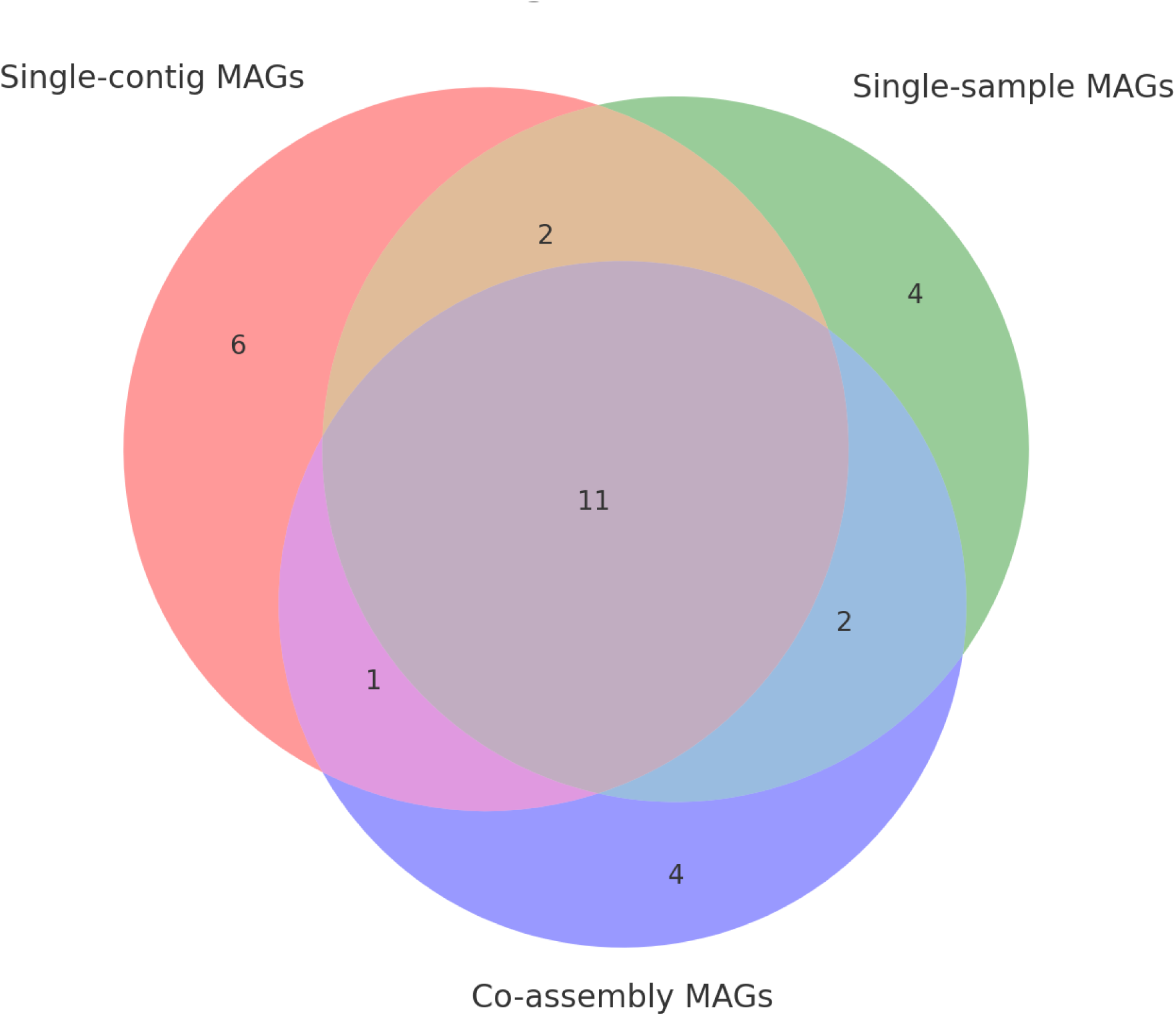
MAGs recovered by each of the approaches. No binning (single-contig MAGs) and Binning (Single-sample MAGs & Co-assembly MAGs).

**Supplementary Figure 5.**
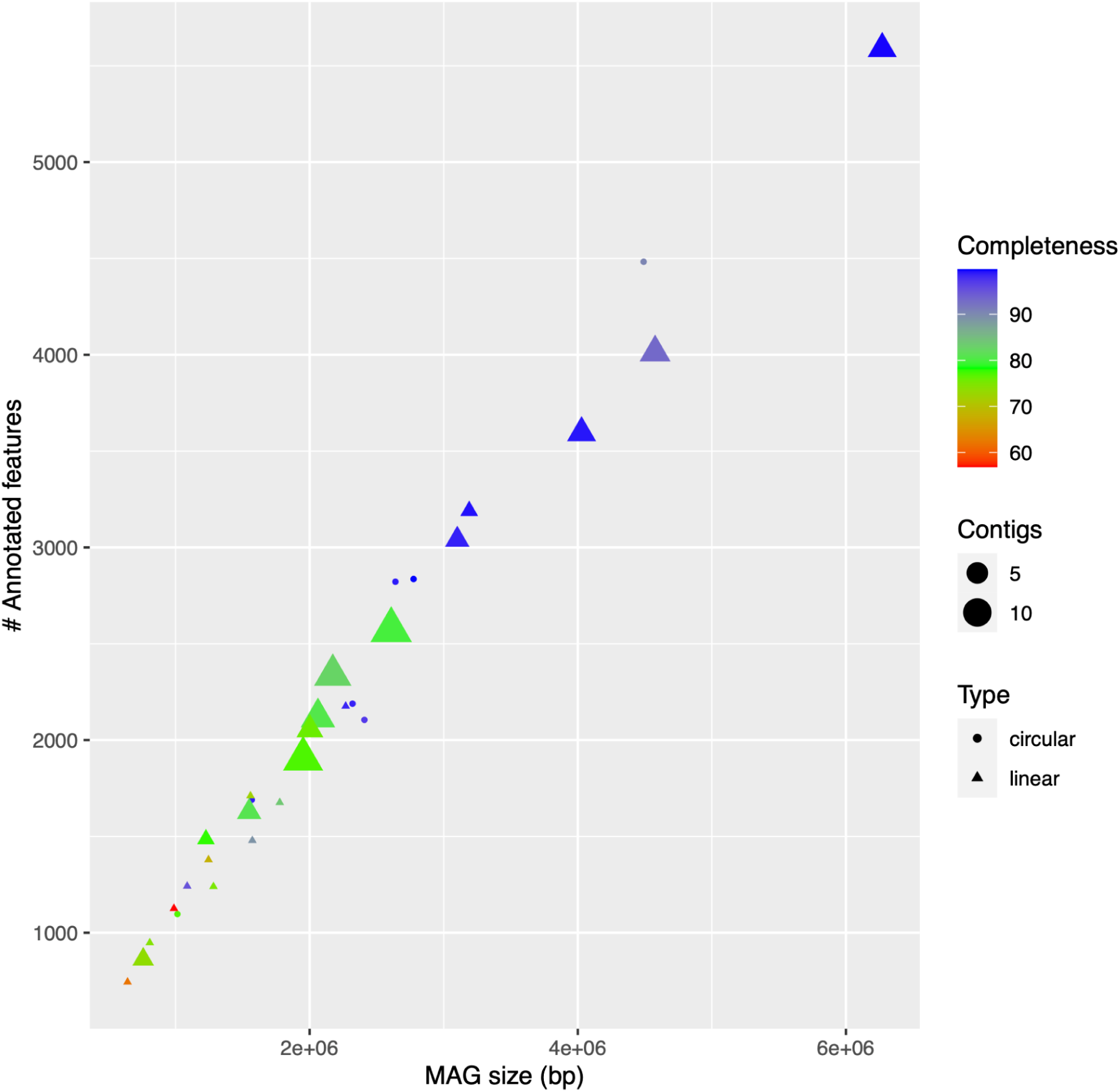
Number of annotated features for each MAG. MAG completeness and number of contigs are shown.

**Supplementary Figure 6.**
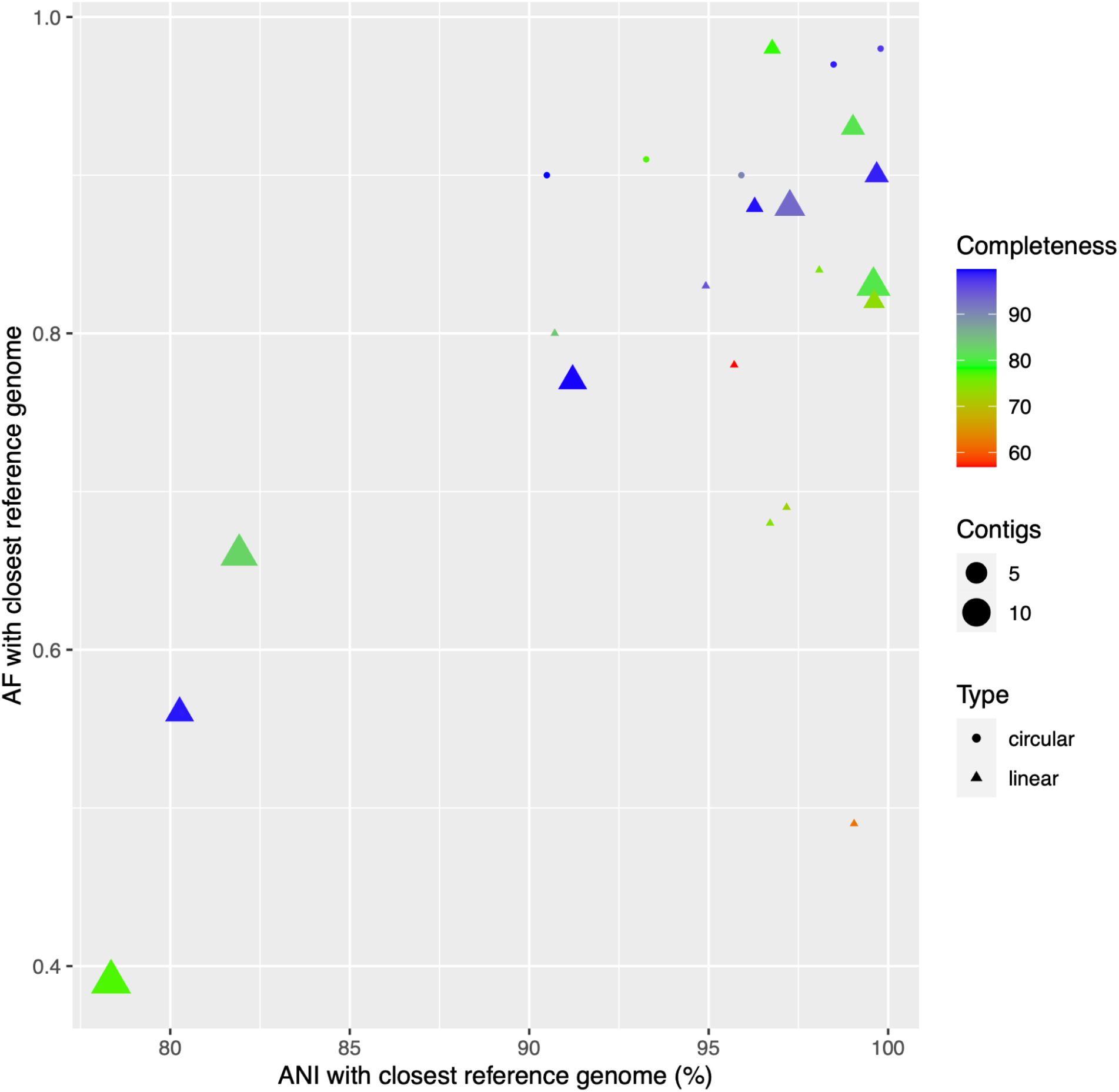
Similarity of the MAGs to reference genomes. For each of the 24 PacBio MAGs for which relatively close relatives could be identified in GTDB, the Average Nucleotide Identity (ANI) and the Alignment Fraction (AF) with the closest reference genome are shown. The AF shows the amount of genome overlap with the reference genome, which was used to calculate the ANI. Note that there are 15 MAGs with an AF > 0.8 and an ANI >90%, and 5 MAGs with an AF > 0.9 and ANI > 93%. Such similarity between some MAGs (including several Single-Contig MAGs) and reference genomes supports our MAG construction method.

**Supplementary Figure 7. Temporal abundance and nucleotide diversity patterns of the 30 long-read MAGs at BBMO and SOLA.** For each MAG, BBMO is shown in the upper row and SOLA in the lower row. From left to right, panels show MAG abundance (RPKG) by month, abundance through time, genome-wide nucleotide diversity (π; blue) and normalized π (red) across samples, and the relationship between nucleotide diversity and MAG abundance (RPKG). Red curves in the rightmost panels show smoothed relationships between π and RPKG, with grey shaded areas indicating confidence intervals. MAGs are presented in the same order as in Table 1. (See file Supp.Figure.7.pdf)

**Supplementary Figure 8. Horizontal and vertical coverage of the 30 long-read MAGs across the BBMO and SOLA time series.** Each point represents an individual metagenomic sample. Horizontal coverage (x-axis) indicates the percentage of the MAG covered by mapped reads, whereas vertical coverage (y-axis) indicates the mean sequencing depth across the mapped regions of the MAG. Point shapes indicate sampling location (BBMO or SOLA), and colors indicate seasonal state (Summer–Autumn or Winter–Spring). Each panel corresponds to one MAG, presented in the same order as in Table 1. (See file Supp.Figure.8.pdf)

**Supplementary Figure 9. Population structure, candidate adaptive-gene pN/pS, and abundance patterns across BBMO and SOLA.** For each MAG, the upper panel shows a UPGMA dendrogram based on pairwise genome-wide Fst among samples meeting the horizontal-coverage criterion described in the Methods. The dashed horizontal line indicates the Fst=0.15 threshold used to delineate candidate populations, and branch colors distinguish the resulting sample clusters. Bars below the dendrogram indicate the seasonal state of each sample (Winter–Spring, cold waters; Summer–Autumn, warm waters) and sampling location (BBMO or SOLA). The middle panel shows pN/pS values for candidate adaptive genes across the same samples; values >2 were capped at 2 for visualization. White cells indicate zero or unavailable pN/pS values. The lower panel shows MAG abundance (RPKG) across the corresponding samples. Samples with RPKG <1 were not considered when interpreting population structure. Samples are shown in the order defined by the Fst dendrogram and therefore are not arranged chronologically. MAGs are presented in the same order as in Table 1. Note that *Amylibacter* (s35.ctg000041c_BL_0902sc) is not included, as only a few BBMO samples (metagenomes) pass the threshold of horizontal coverage > 25%, while no sample from SOLA passes this threshold. (See file Supp.Figure.9.pdf)

**Supplementary Figure 10. Genome-wide population differentiation versus temporal distance for the 27 MAGs.** Pairwise genome-wide Fst values are plotted against the temporal distance (days) between samples for BBMO and SOLA, with each point representing one pair of samples. Only sample pairs for which both samples had >25% horizontal coverage of the corresponding MAG were included. Blue lines show linear regression fits and red dashed lines show LOESS fits; shaded areas indicate 95% confidence intervals around the fitted relationships. The slope, adjusted R^2^, and p-value of the linear regression are shown in each panel. Analyses are shown only for time series with sufficient data for the corresponding MAG. (See file Supp.Figure.10.pdf)

**Supplementary Figure 11. Temporal dynamics of pN/pS for candidate adaptive genes across the BBMO and SOLA time series.** For each MAG, temporal profiles of pN/pS are shown separately for BBMO and SOLA. Each panel represents one candidate adaptive gene, identified by its gene ID (top-left corner), with samples arranged in chronological sampling order. The x-axis indicates successive sampling months (1–140 for BBMO and 1–90 for SOLA), and the y-axis shows pN/pS. Samples with <25% horizontal coverage of the corresponding MAG were excluded. For visualization, pN/pS values >2 were capped at 2, whereas undefined values resulting from pS = 0 were plotted as −1. (See file Supp.Figure.11.pdf)

**Supplementary Figure 12.**
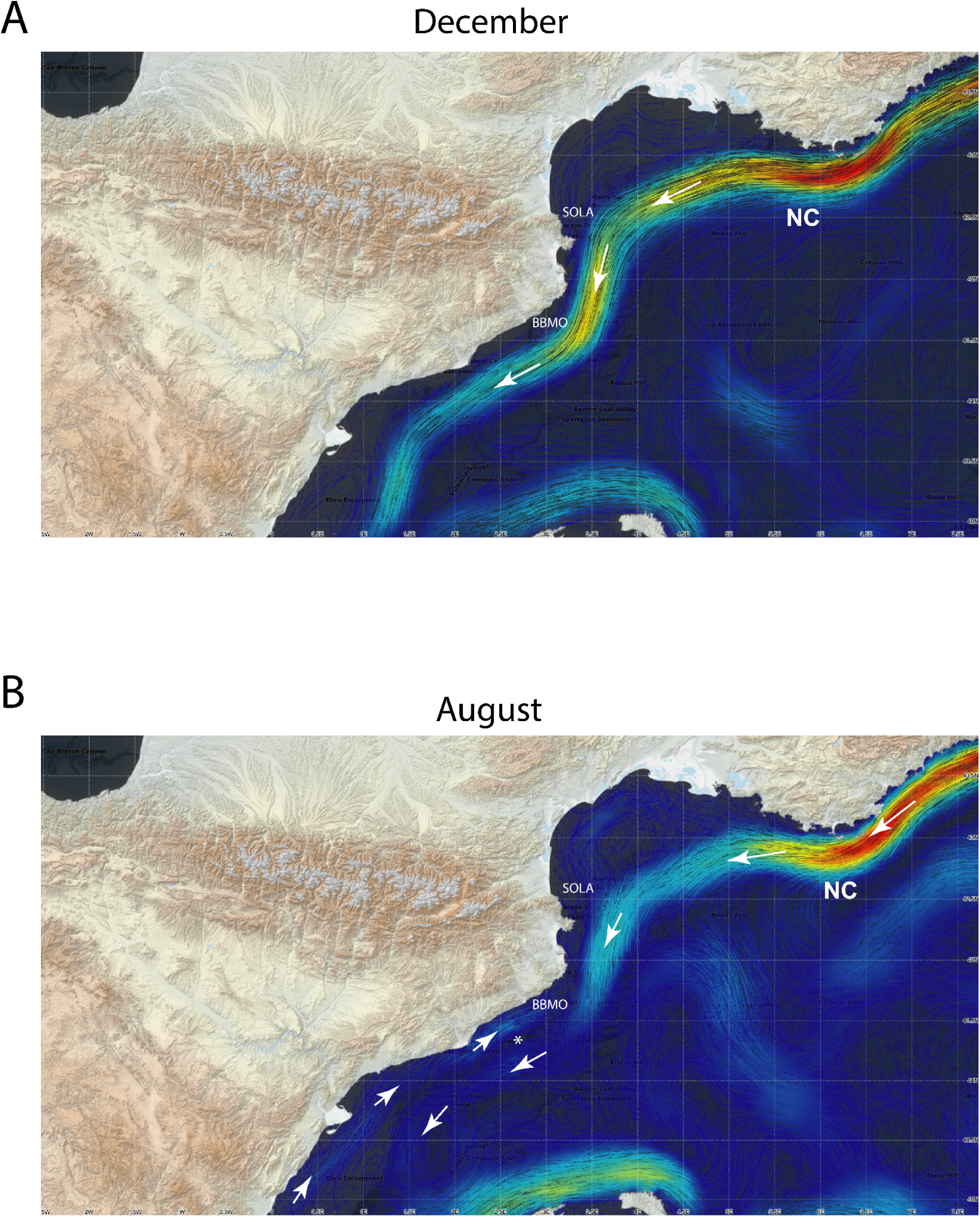
Seasonal surface circulation patterns around BBMO and SOLA in the NW Mediterranean Sea. The Blanes Bay Microbial Observatory (BBMO; 41°40′13″N, 2°48′00″E) and the Banyuls Bay Microbial Observatory (SOLA; 42°31’N 3°11’E) are shown together with climatological surface currents derived from MEDSEA products for 1987–2019 (33 years;^36^; https://cosmo.icm.csic.es/currents). (A) December circulation, representative of the dominant pattern during most of the year, with the Northern Current (NC) flowing southwestward along the continental margin and potentially providing physical connectivity between SOLA and BBMO. (B) August circulation, showing a weaker northeastward flow near BBMO and a mesoscale eddy (asterisk), while the influence of the NC on BBMO appears reduced. Analysis of the MEDSEA product indicates that this circulation pattern occurs predominantly during August and September. These seasonal changes in circulation could modulate dispersal pathways and connectivity between BBMO and SOLA, potentially contributing to differences in the microbial populations reaching each site.

