## Supplementary Figure 7 for "Seasonal population structure and adaptive signatures across long-term marine microbial time series"

**Supplementary Figure 7. Temporal abundance and nucleotide diversity patterns of the 30 long-read MAGs at BBMO and SOLA.** For each MAG, BBMO is shown in the upper row and SOLA in the lower row. From left to right, panels show MAG abundance (RPKG) by month, abundance through time, genome-wide nucleotide diversity ($\pi$; blue) and normalized $\pi$ (red) across samples, and the relationship between nucleotide diversity and MAG abundance (RPKG). Red curves in the rightmost panels show smoothed relationships between $\pi$ and RPKG, with grey shaded areas indicating confidence intervals. MAGs are presented in the same order as in Table 1.

### *Actinomarina* **Acidimicrobiia** (s2266.ctg002461l_BL_0908sc)

#### BBMO

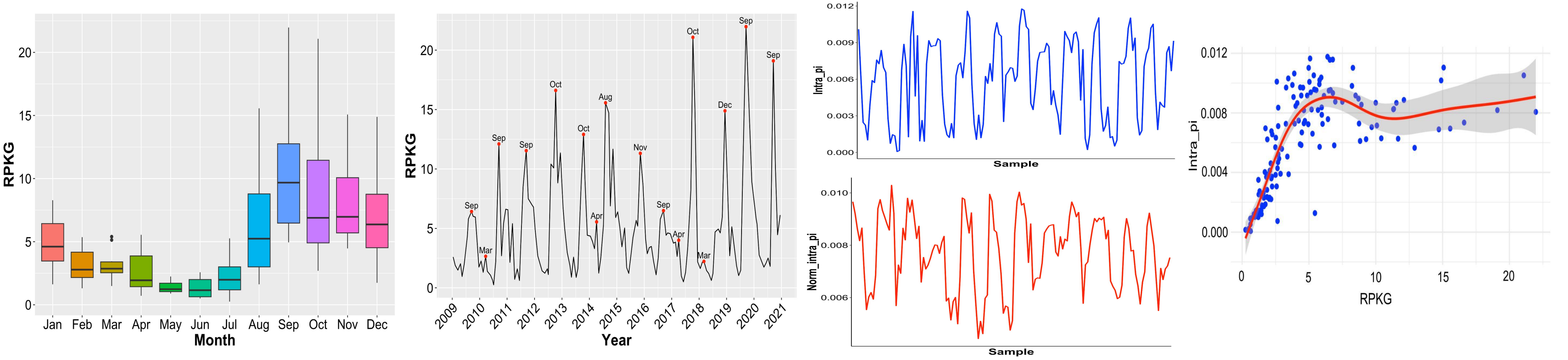


#### SOLA

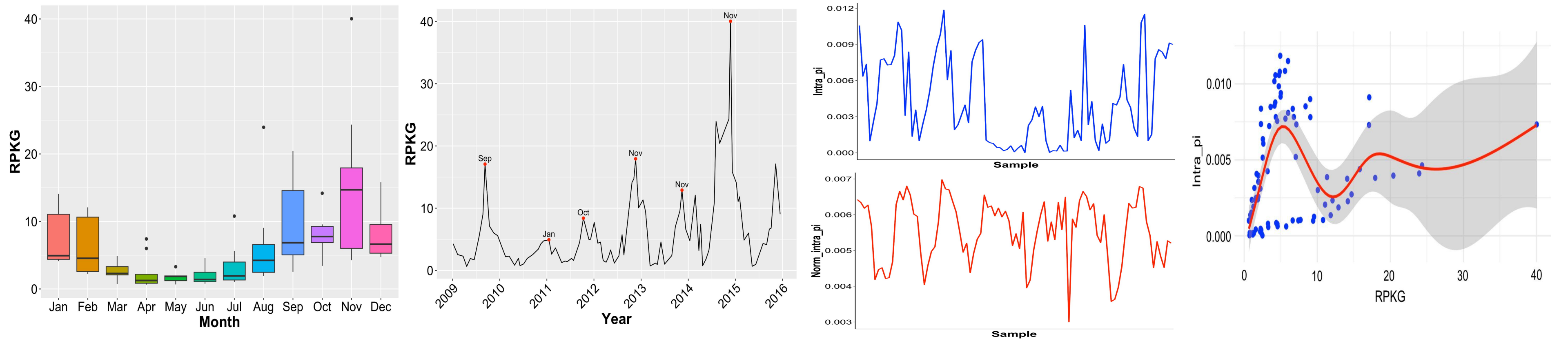

### *Nocardioides* Acidimicrobiia (s131.ctg000151c_BL_0908sc)

#### BBMO

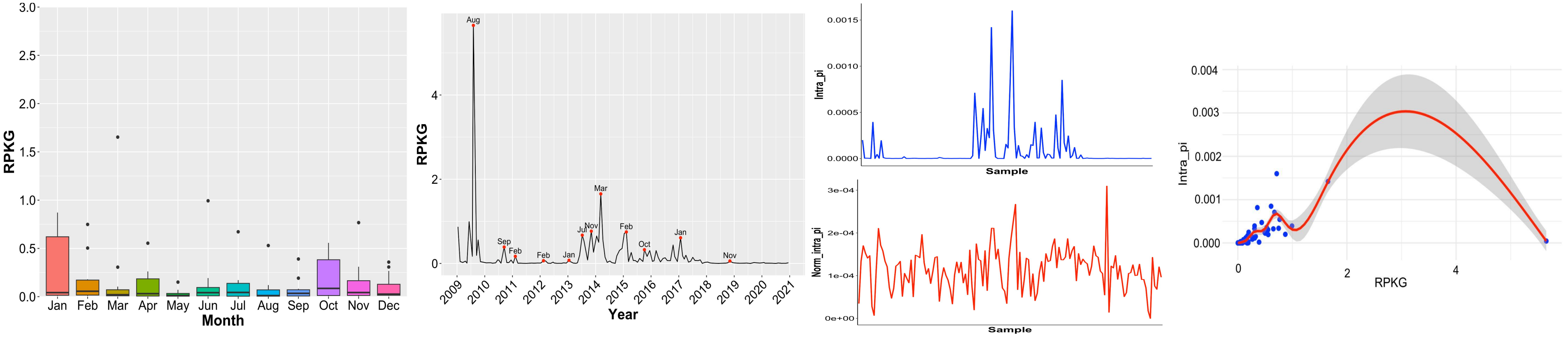


#### SOLA

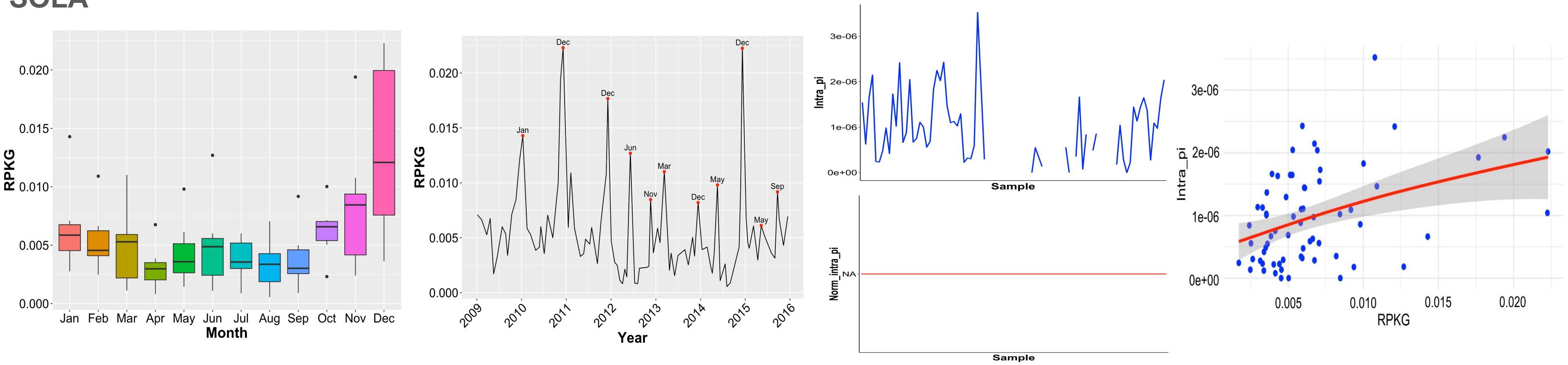

### *Pontimonas* Actinomycetia (BL_pooled_bin.full.596)

#### BBMO

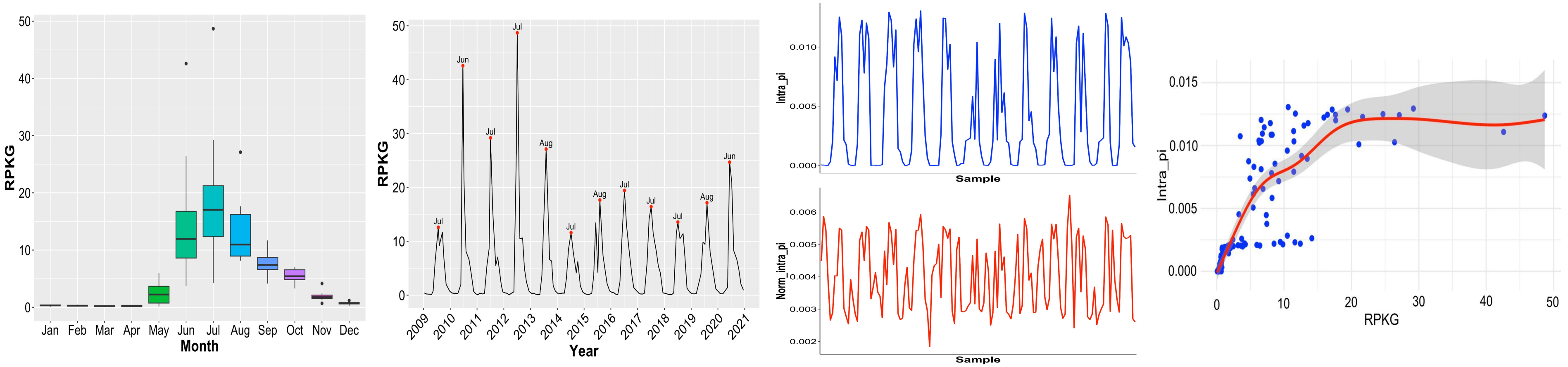


#### SOLA

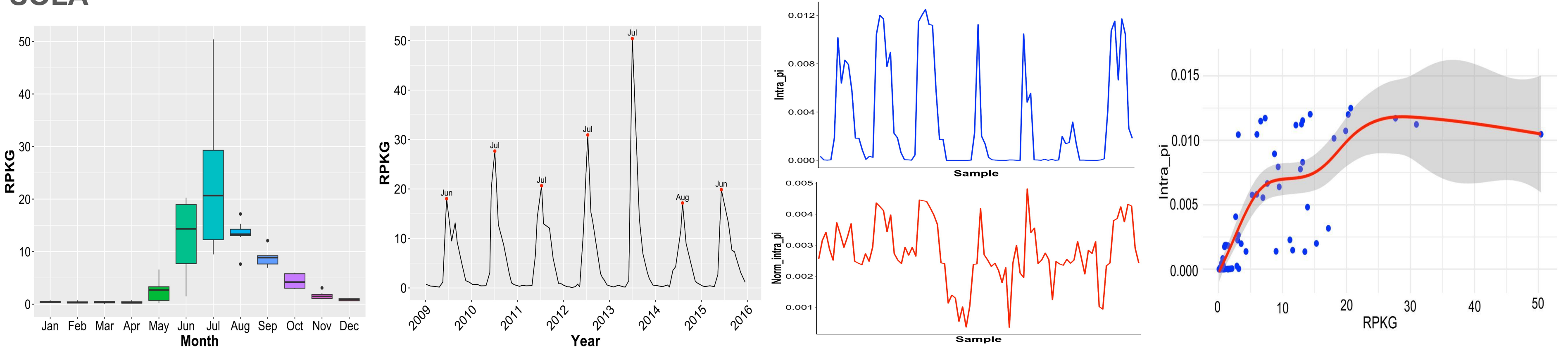

### *Amylibacter* **Alphaproteobacteria** (s35.ctg000041c_BL_0902sc)

#### BBMO

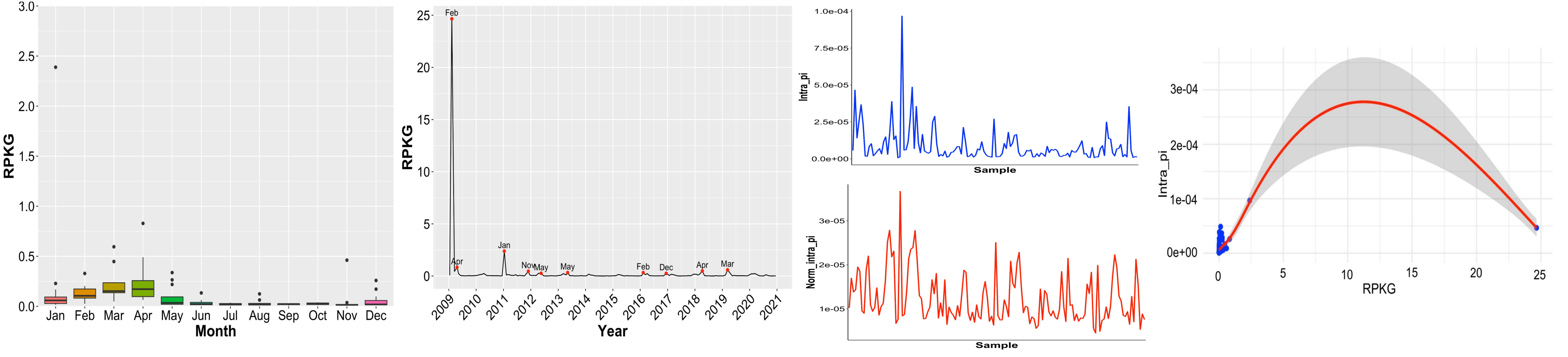


#### SOLA

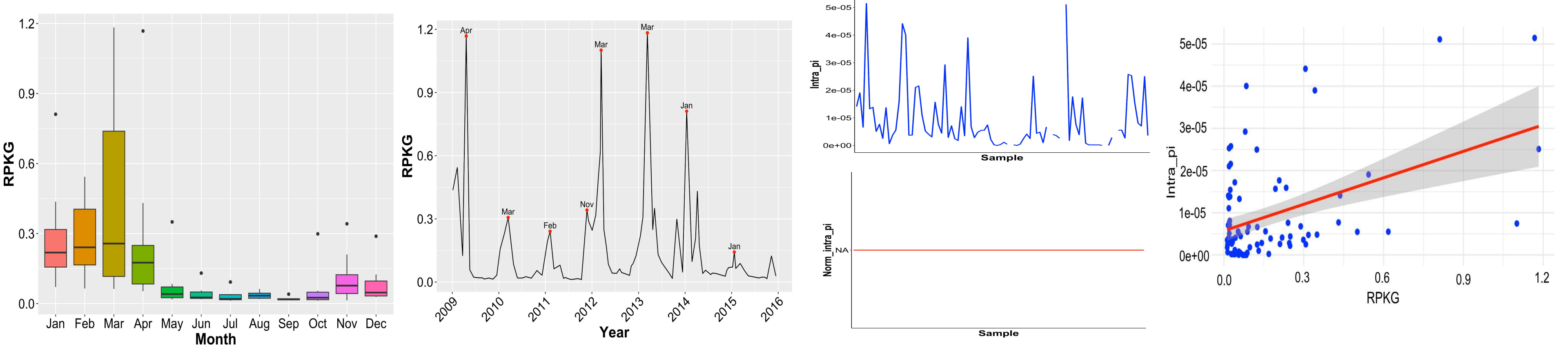

### *Brevundimonas* Alphaproteobacteria (s96.ctg000111c_BL_0908sc)

#### BBMO

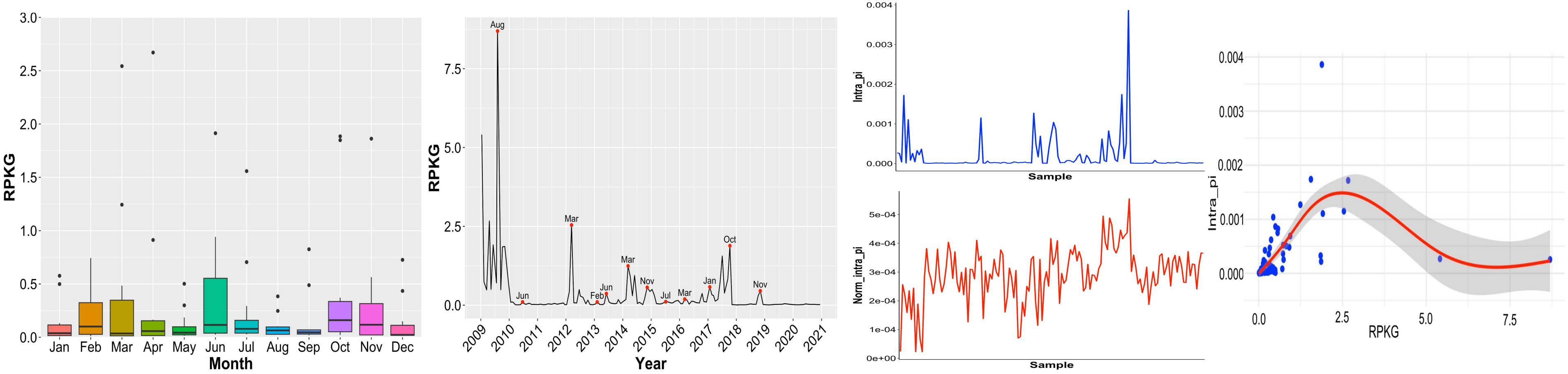


#### SOLA

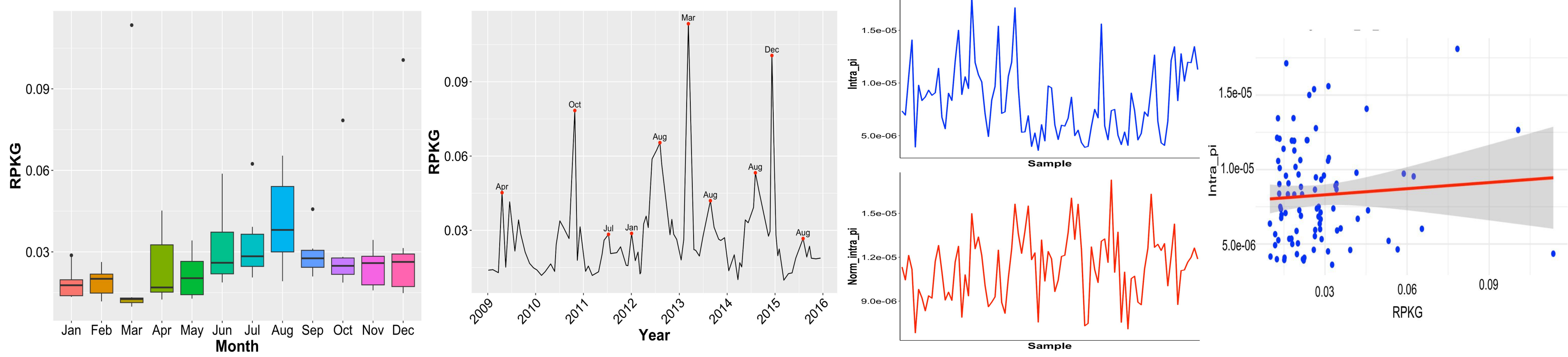

### *Alteriqipengyuania* Alphaproteobacteria (BL_0908_bin.full.436)

#### BBMO

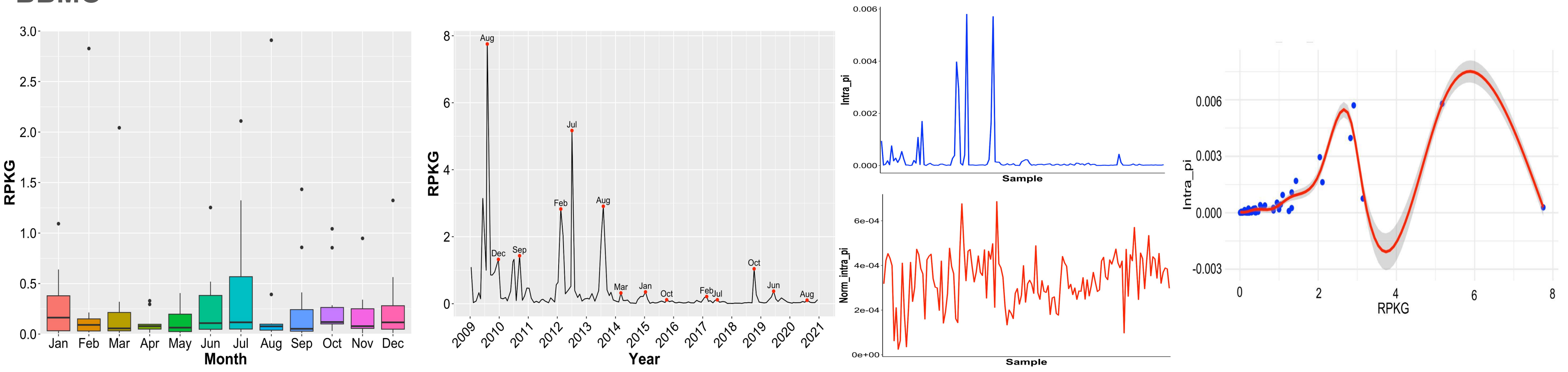


#### SOLA

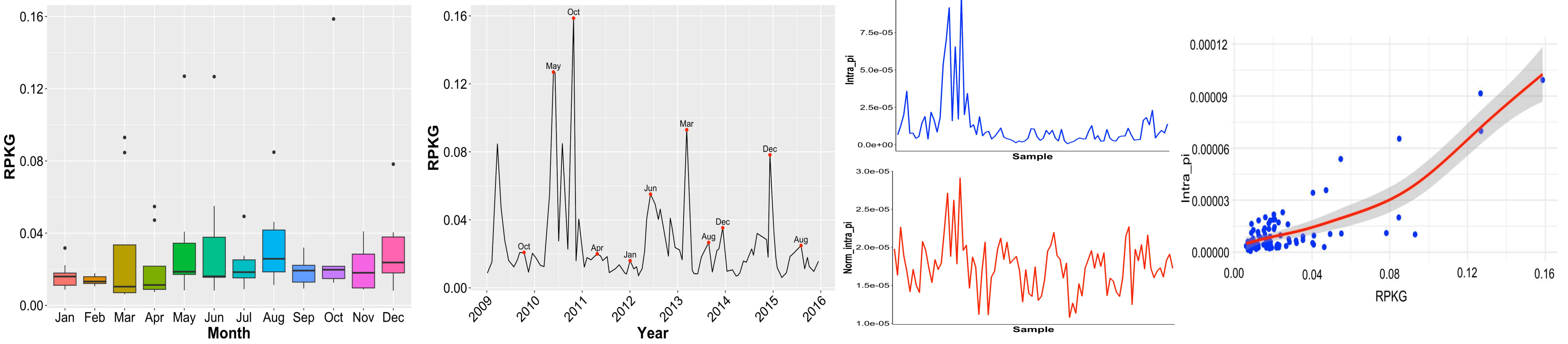

### *Erythrobacter* **Alphaproteobacteria** (BL_0908_bin.full.559)

#### BBMO

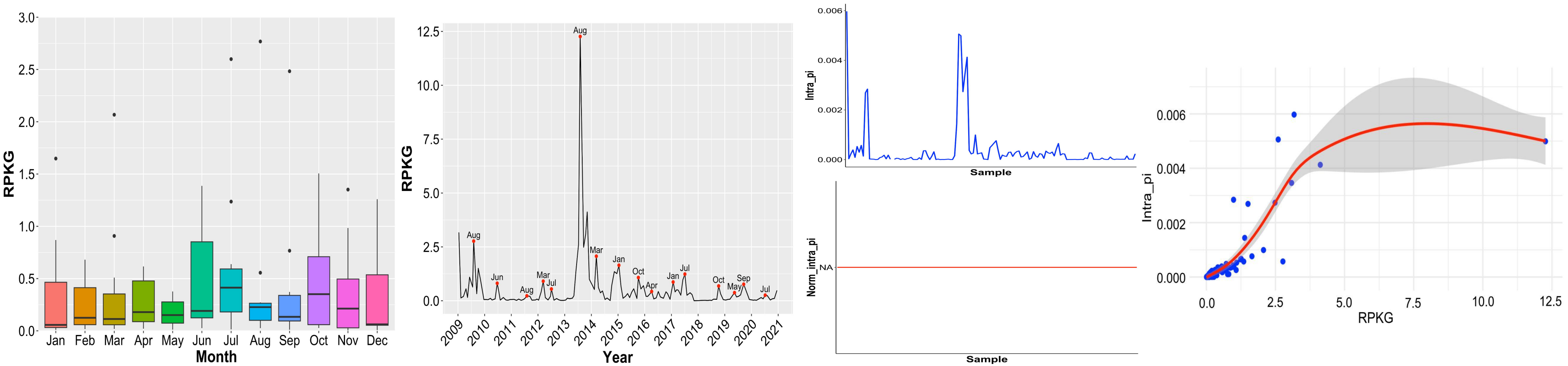


#### SOLA

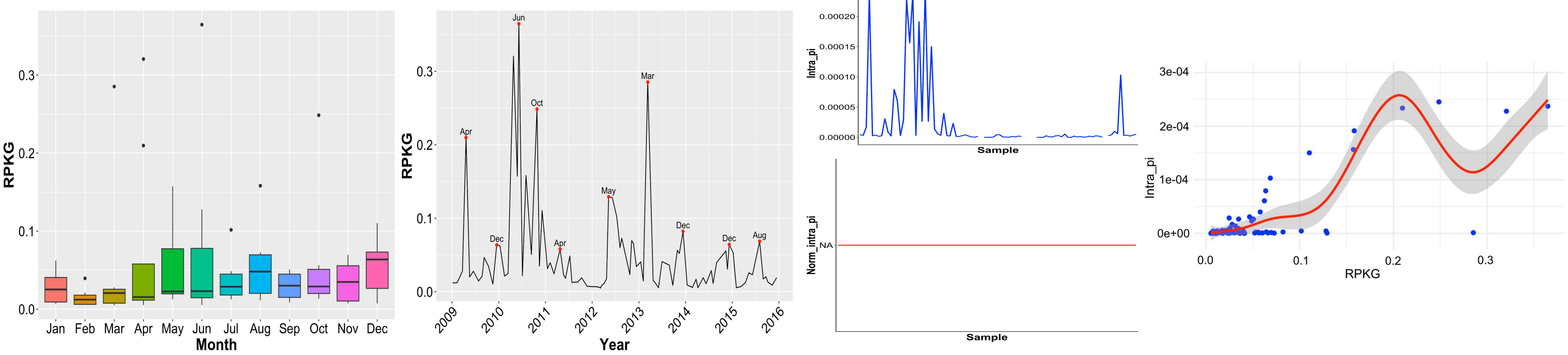

### **MED-G09 Alphaproteobacteria** (BL_pooled_bin.full.761)

#### BBMO

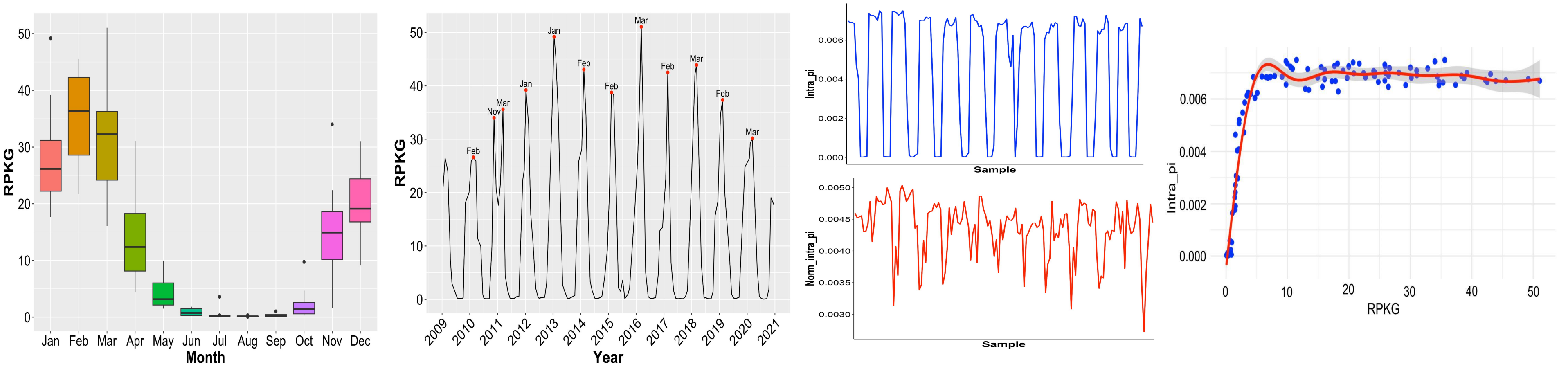


#### SOLA

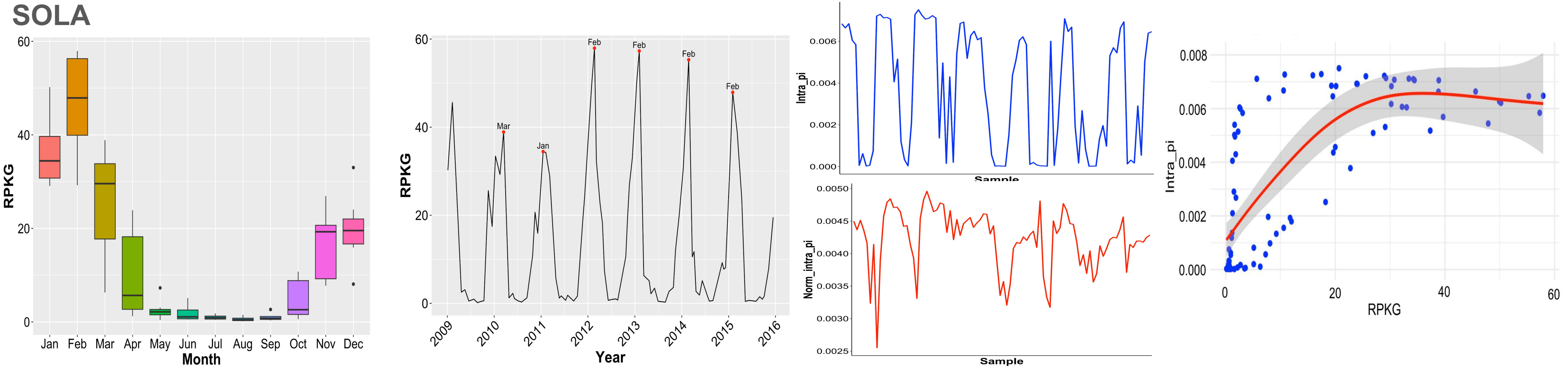

### NA (Rhodobacteraceae) Alphaproteobacteria (BL_pooled_bin.full.145)

#### BBMO

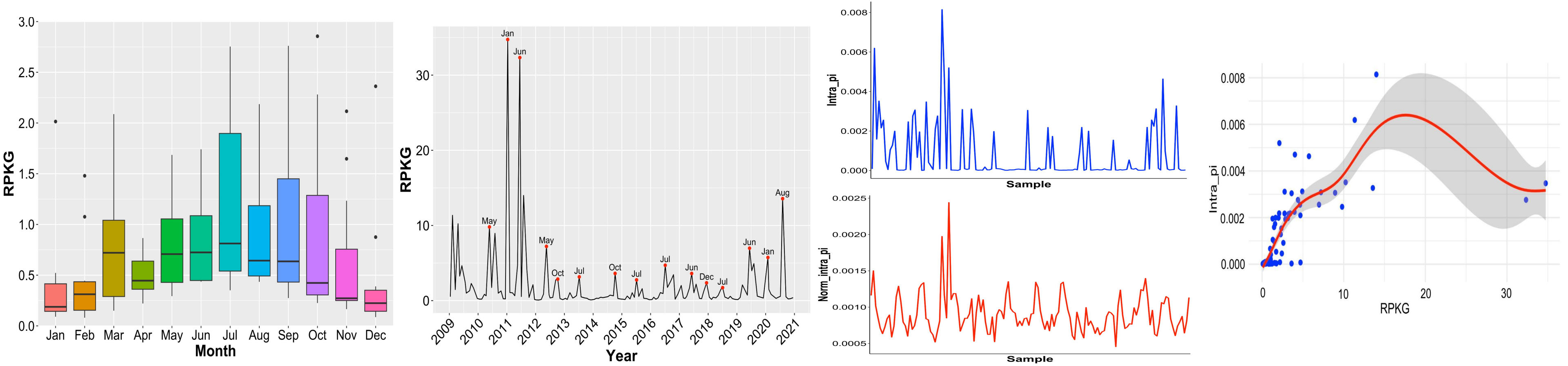


#### SOLA

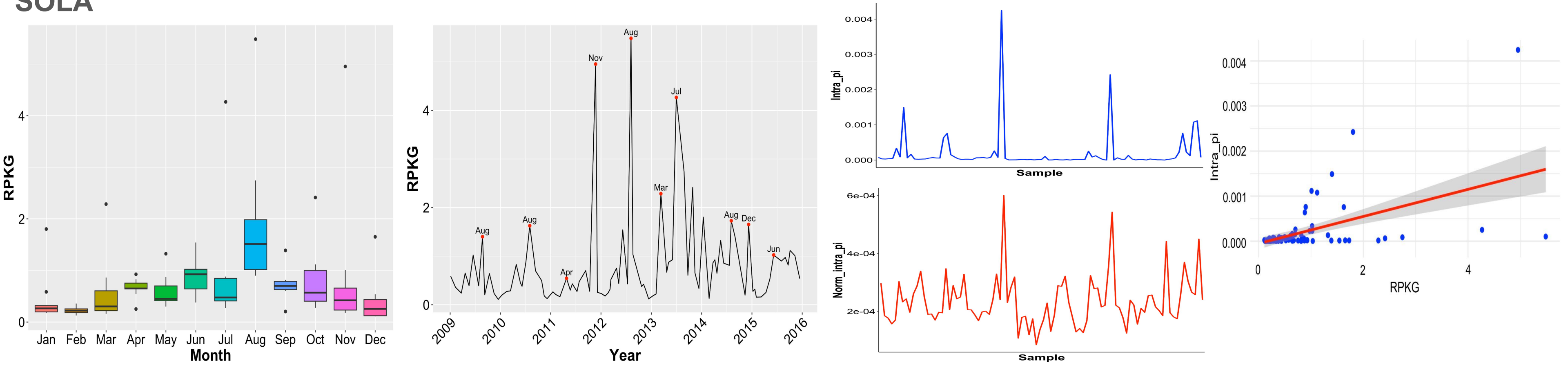

### SAR11: HIMB114 (Pelagibacteraceae) Alphaproteobacteria (s1032.ctg001132l_BL_0908sc)

#### BBMO

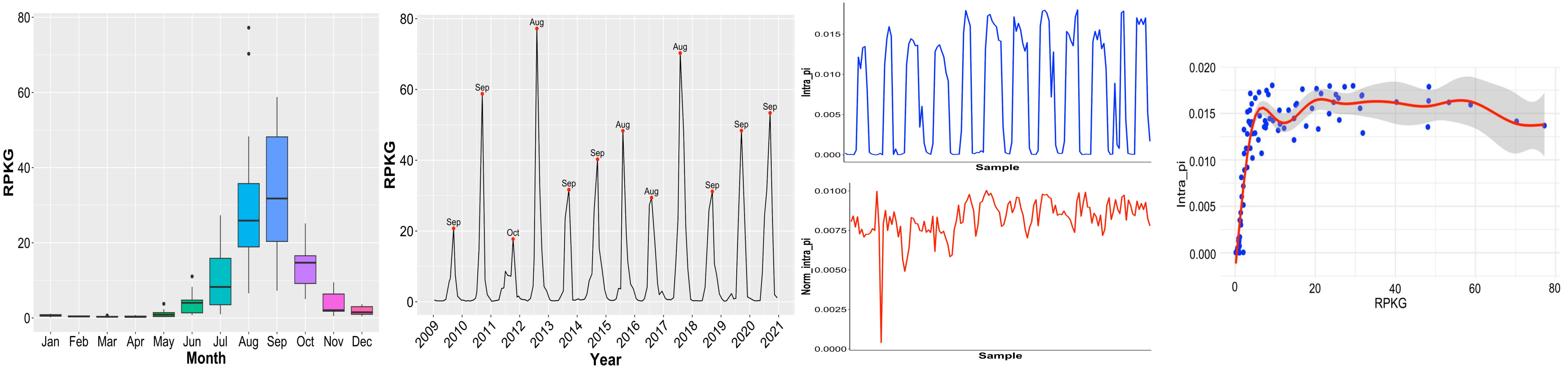


#### SOLA

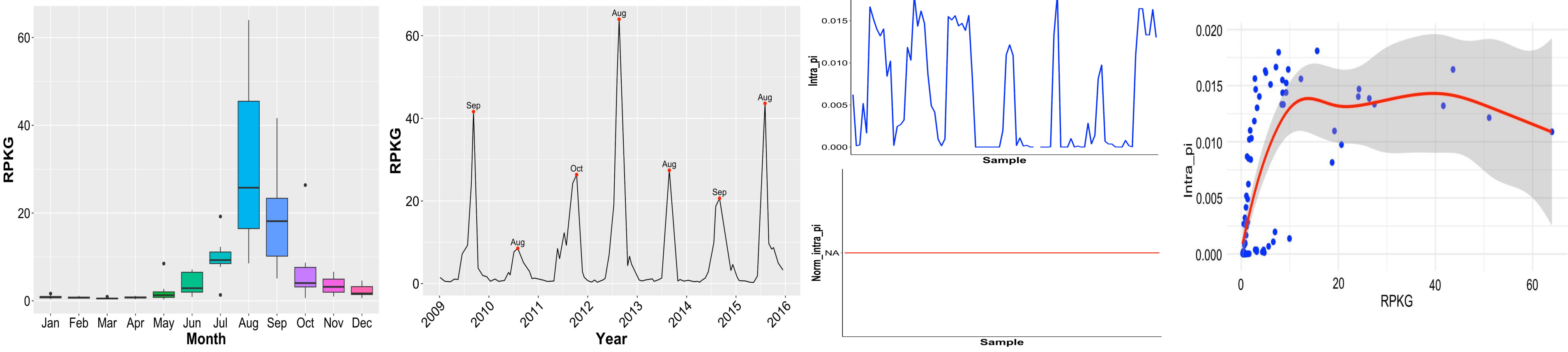

### SAR11: *Pelagibacter* Alphaproteobacteria (s266.ctg000297l_BL_0908sc)

#### BBMO

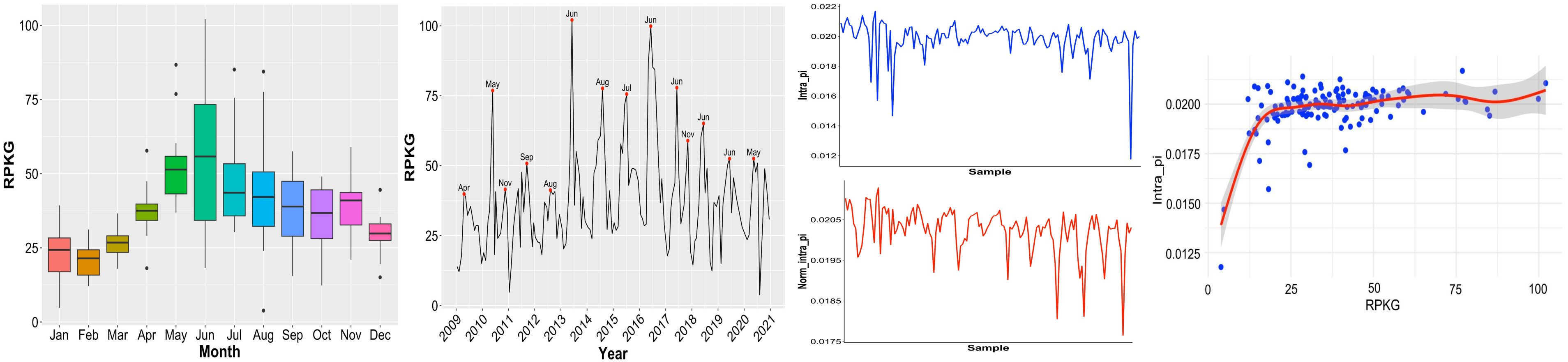


#### SOLA

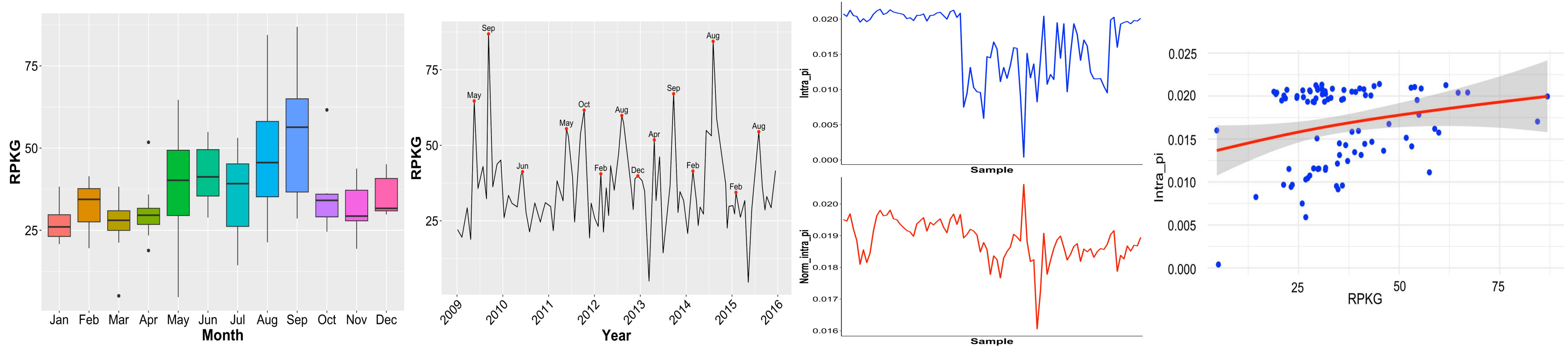

### **TMED13 Alphaproteobacteria** (s325.ctg000366c_BL_0908sc)

#### BBMO

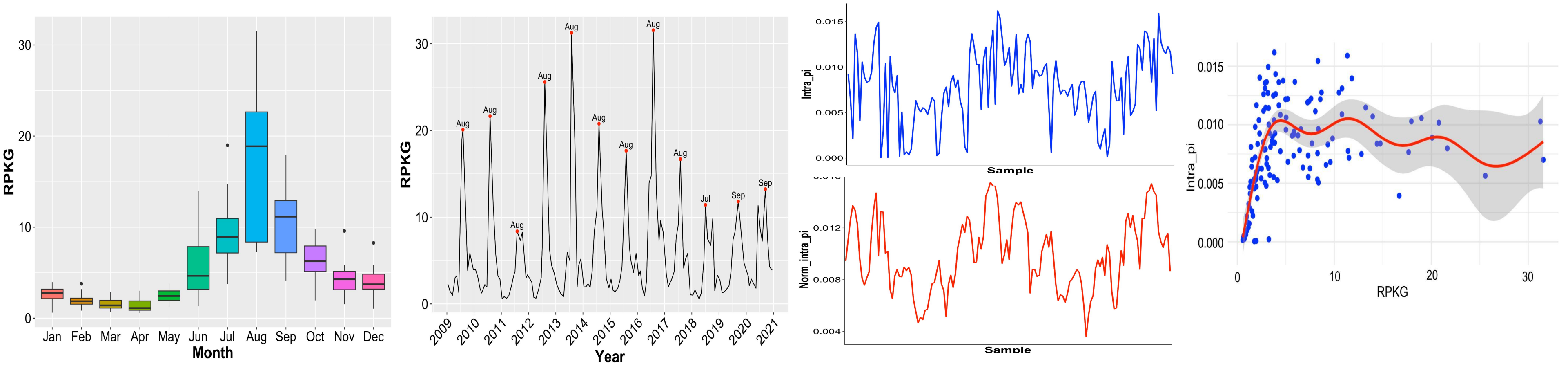


#### SOLA

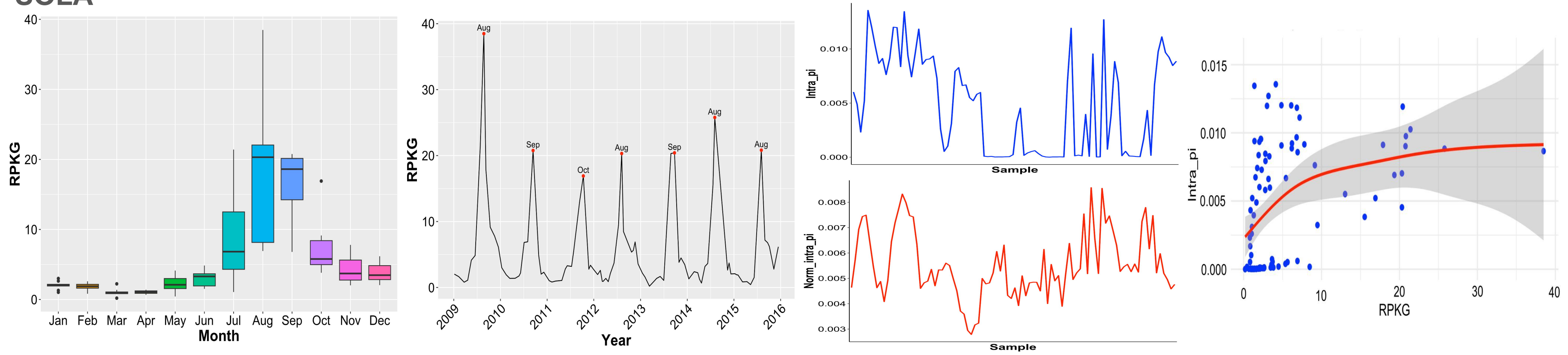

### *Alteromonas* **Gammaproteobacteria** (BL_0908_bin.full.418)

#### BBMO

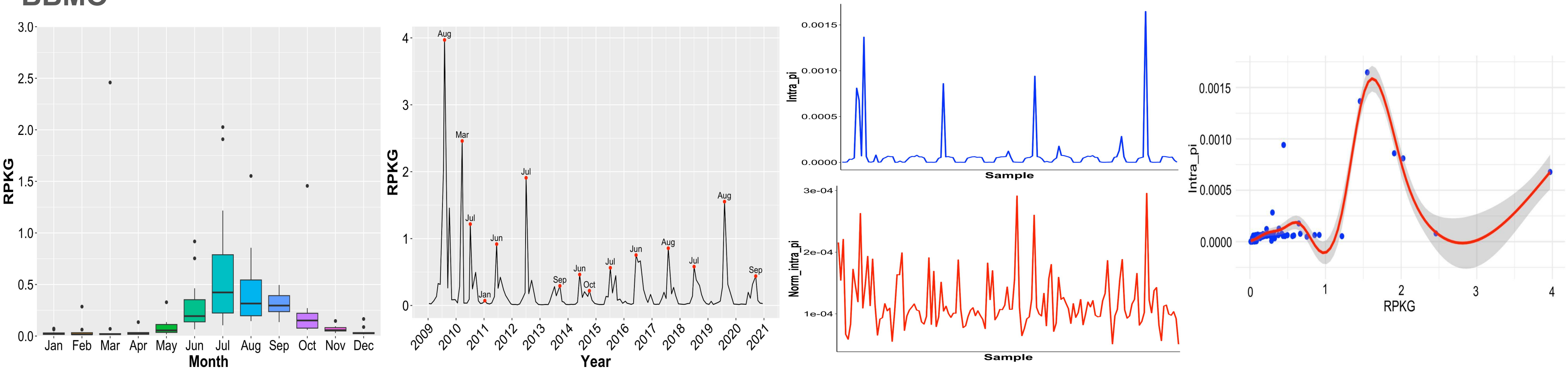


#### SOLA

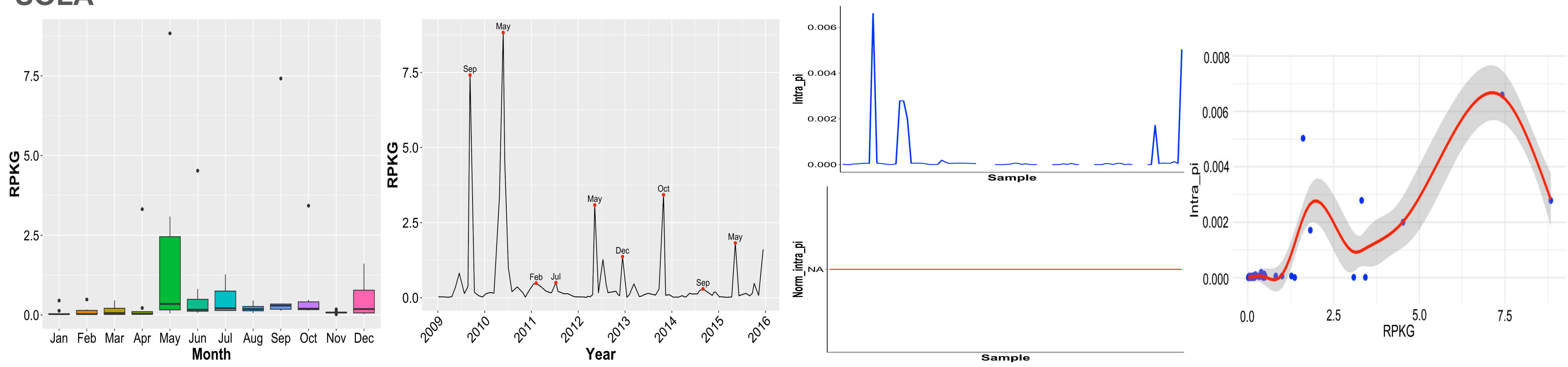

### *Glaciecola* **Gammaproteobacteria** (BL_1001_bin.full.234)

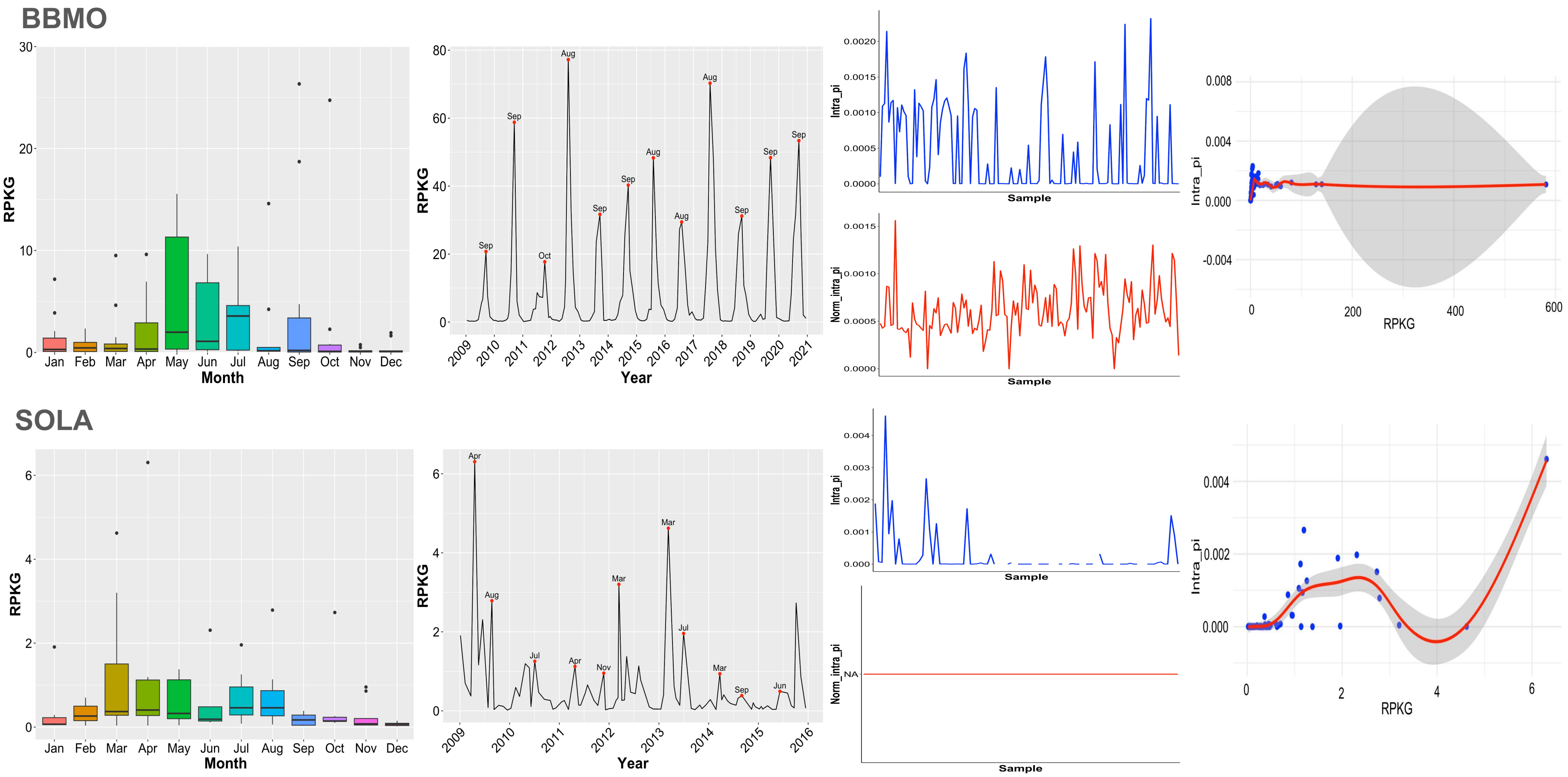

### HTCC2207 (Porticoccaceae) Gammaproteobacteria (s111.ctg000126c_BL_0902sc)

#### BBMO

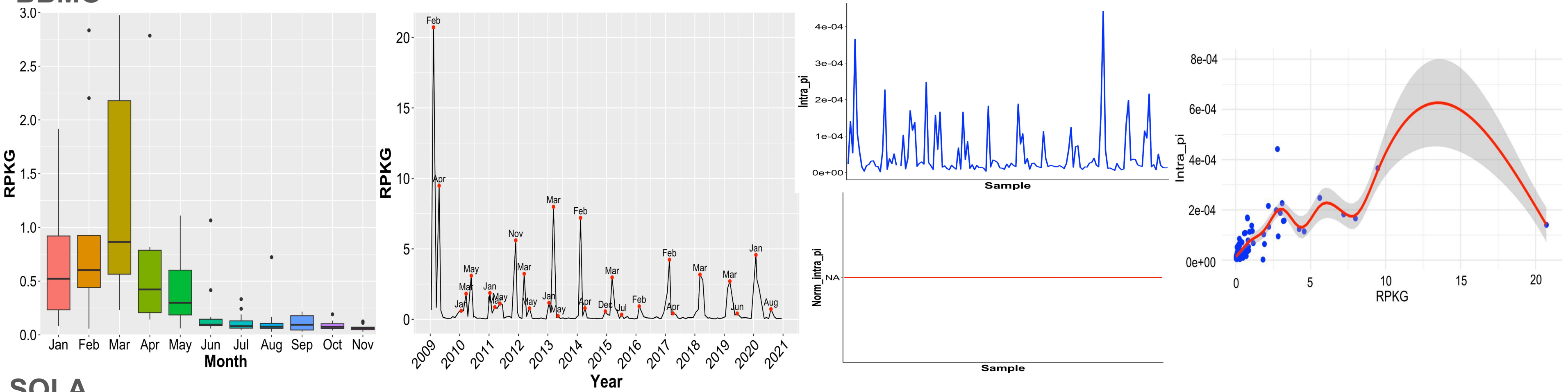


#### SOLA

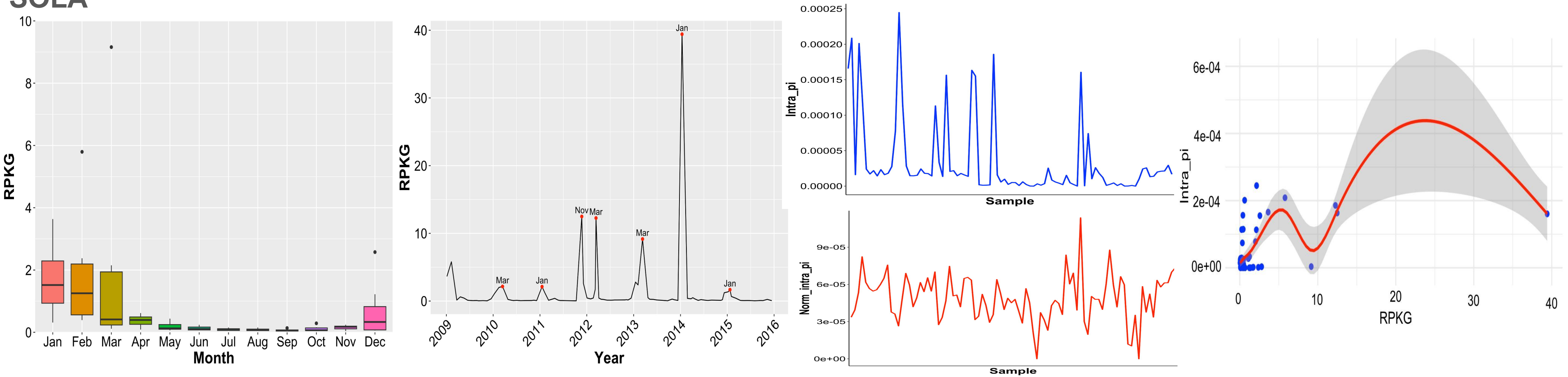

### HTCC2207 (Porticoccaceae) Gammaproteobacteria (s44.ctg000052l_BL_0902sc)

#### BBMO

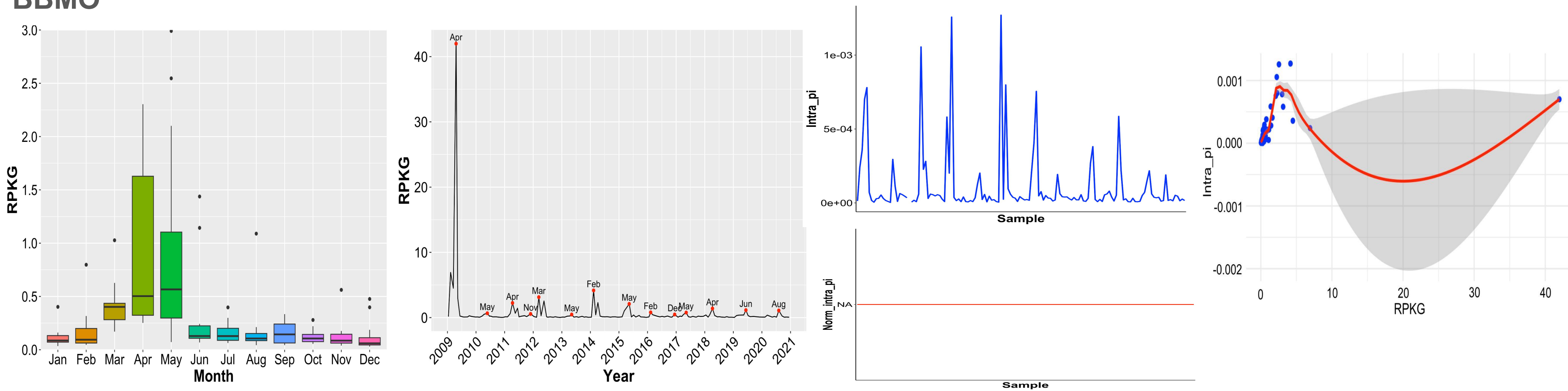


#### SOLA

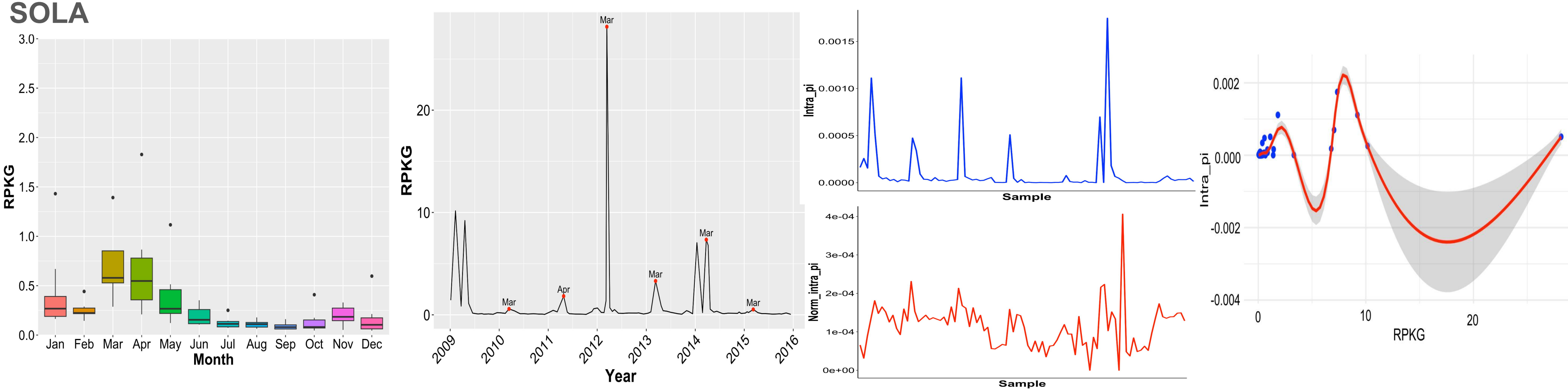

### *Mitsuaria* Gammaproteobacteria (BL_0908_bin.full.528)

#### BBMO

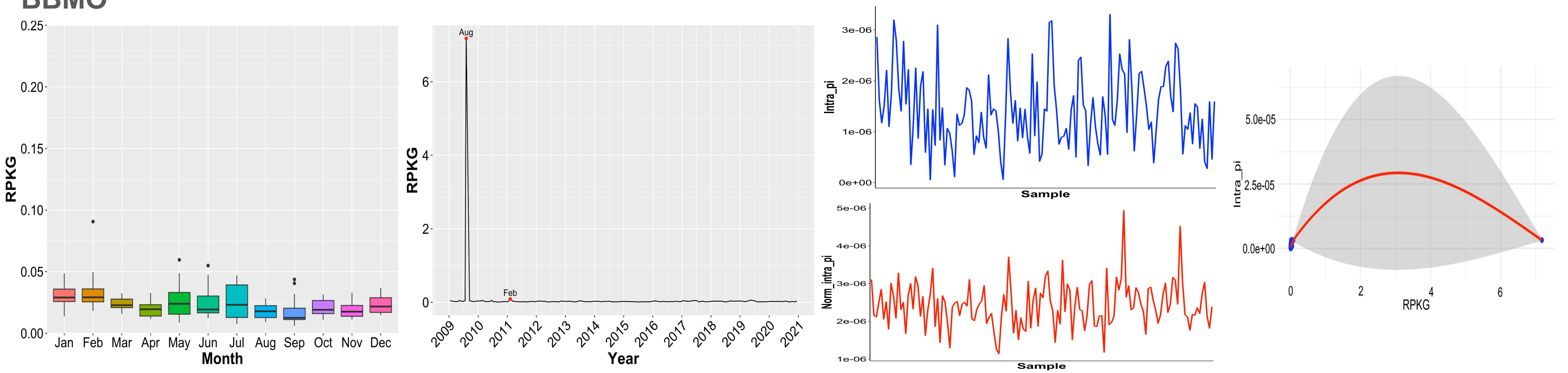


#### SOLA

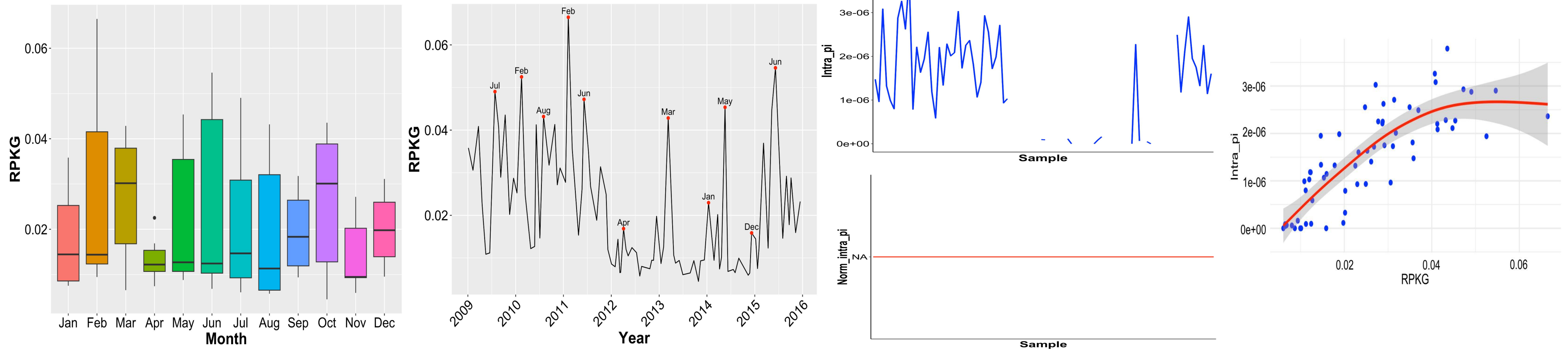

### SCGC-AAA076-P13 (SAR86) Gammaproteobacteria (s215.ctg000240l_BL_0908sc)

#### BBMO

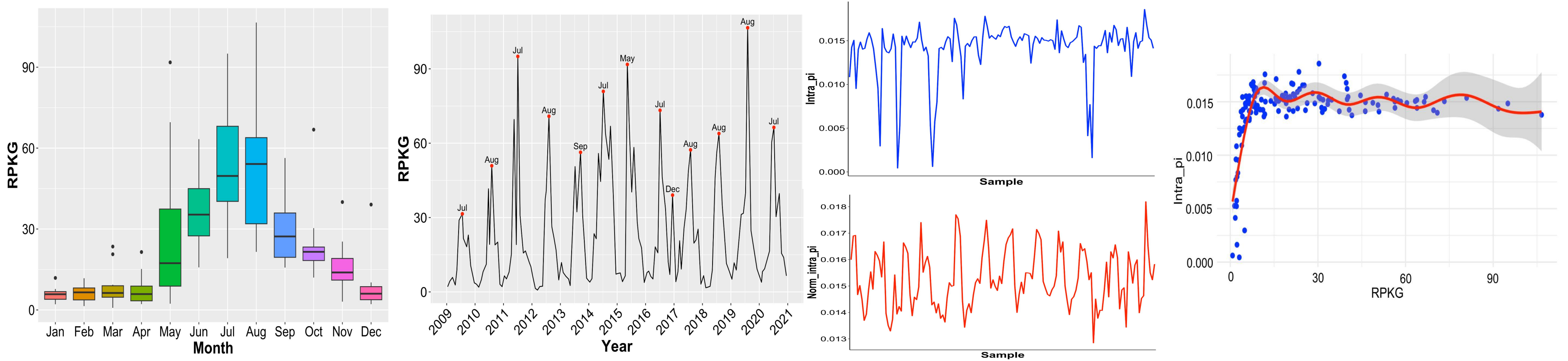


#### SOLA

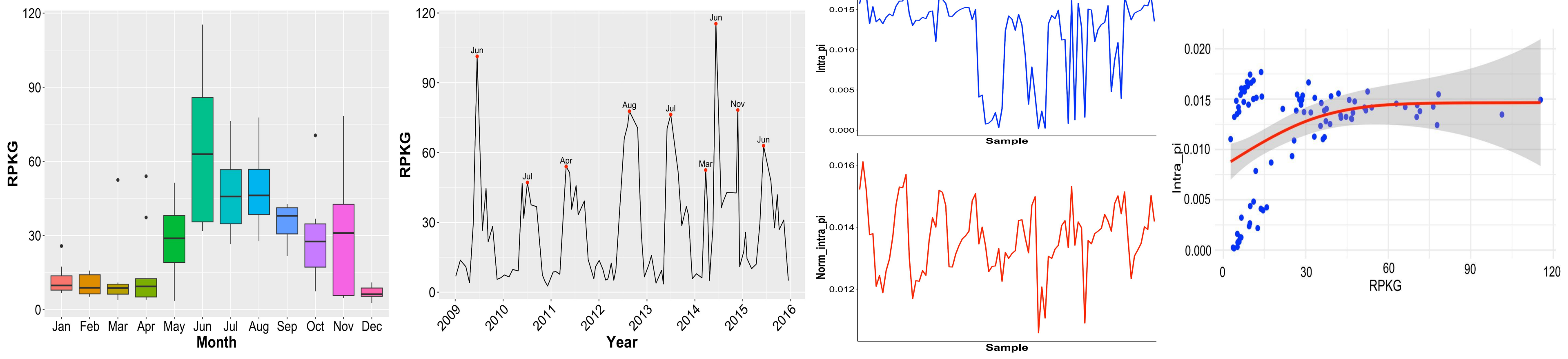

### **SCGC-AAA076-P13 (SAR86) Gammaproteobacteria** (s388.ctg000435l_BL_0908sc)

#### BBMO

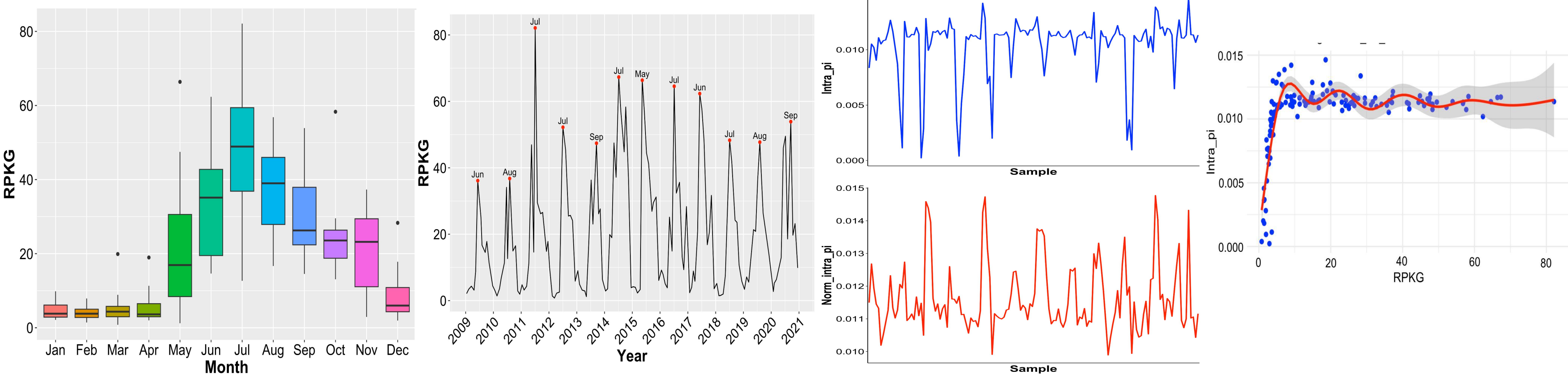


#### SOLA

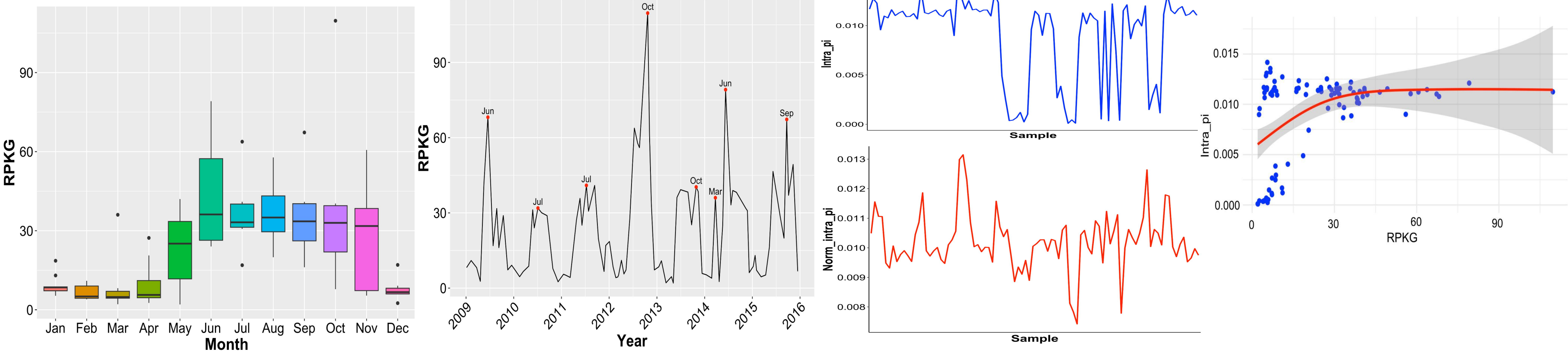

### TMED112 Gammaproteobacteria (BL_pooled_bin.circ.35)

#### BBMO

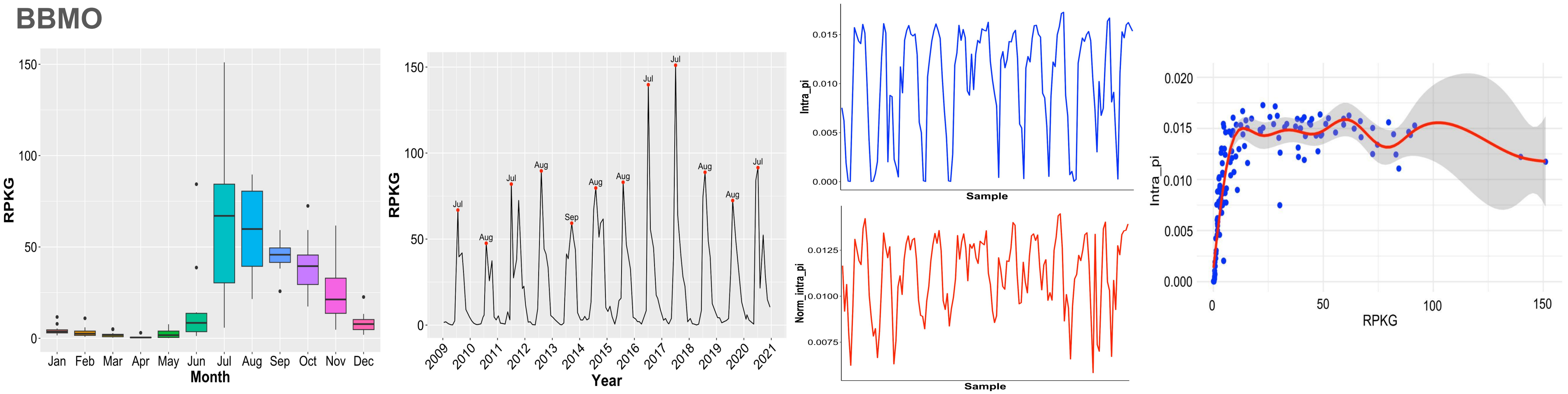


#### SOLA

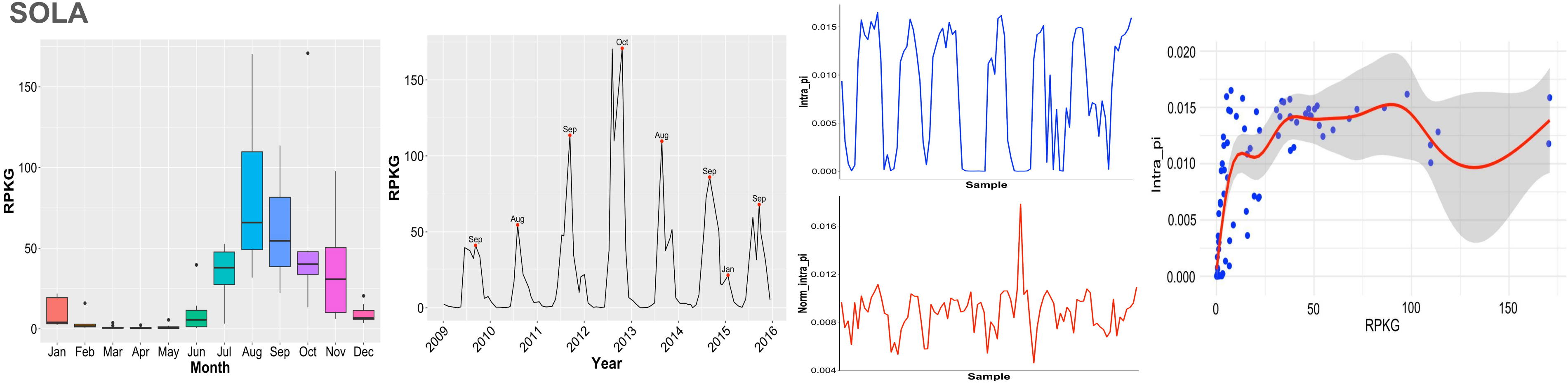

# MGIIb Poseidoniia (Archaea)   (s23.ctg000345l_BL_0902sc)

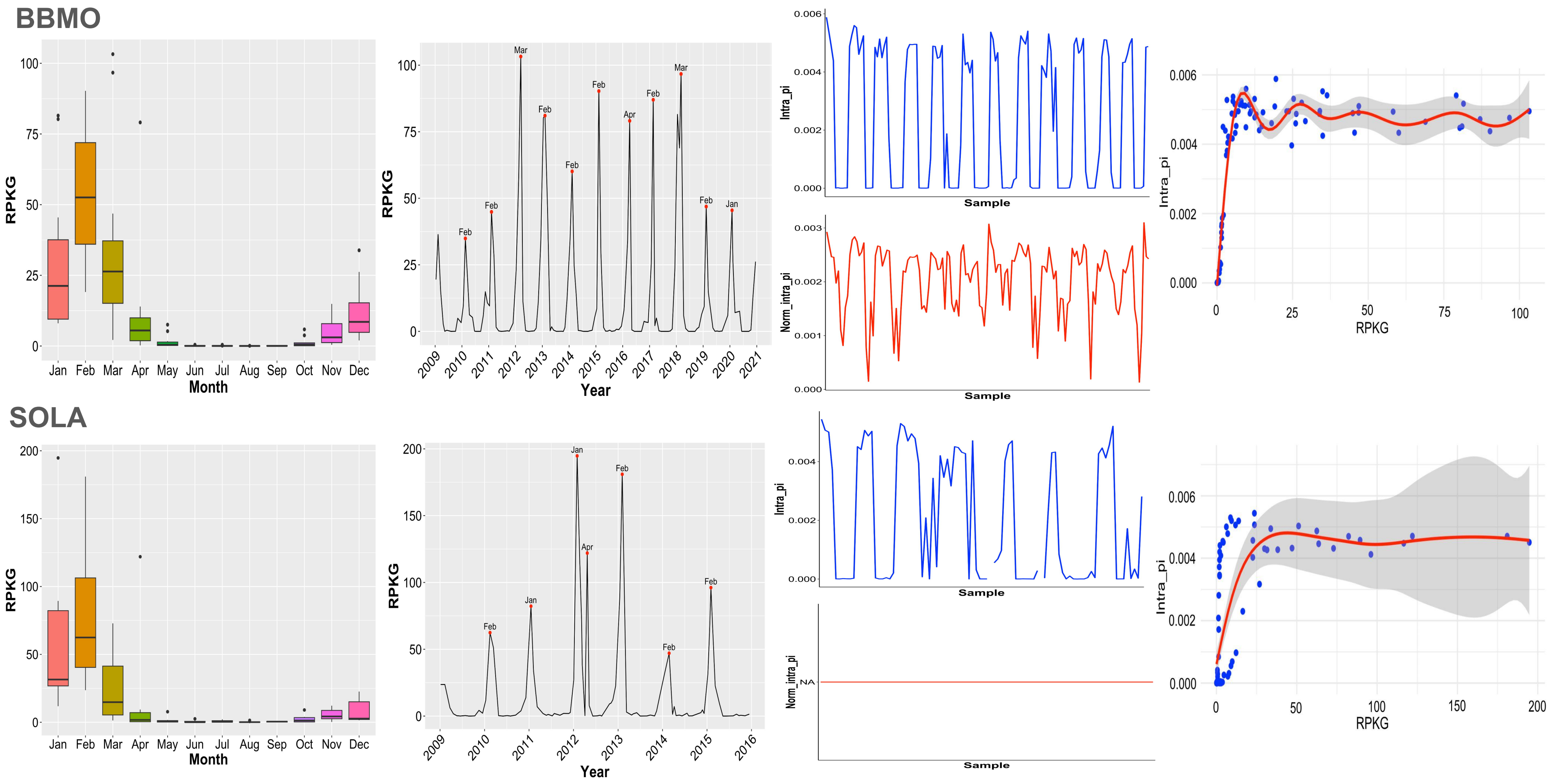

### *Marinisoma* **Marinisomatia** (BL_0908_bin.full.522)

#### BBMO

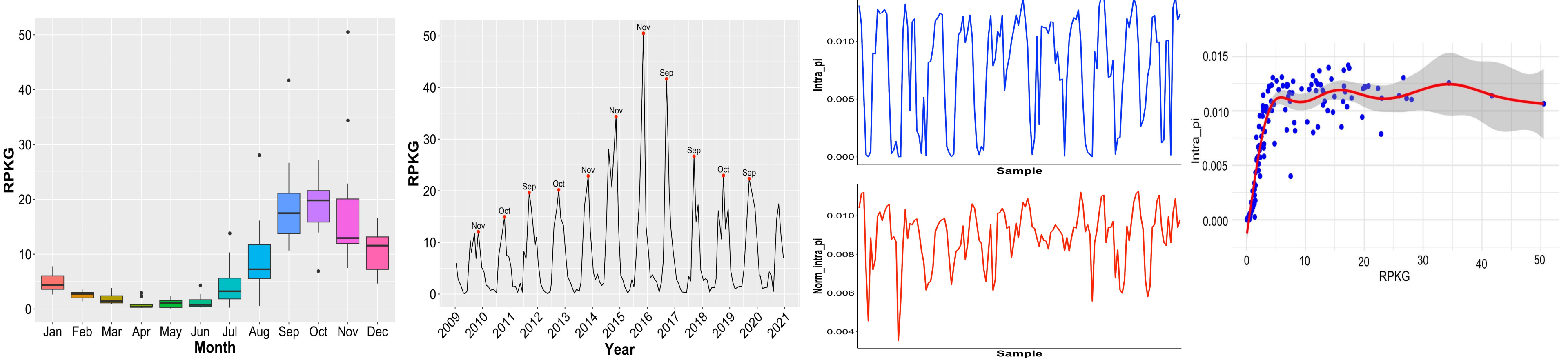


#### SOLA

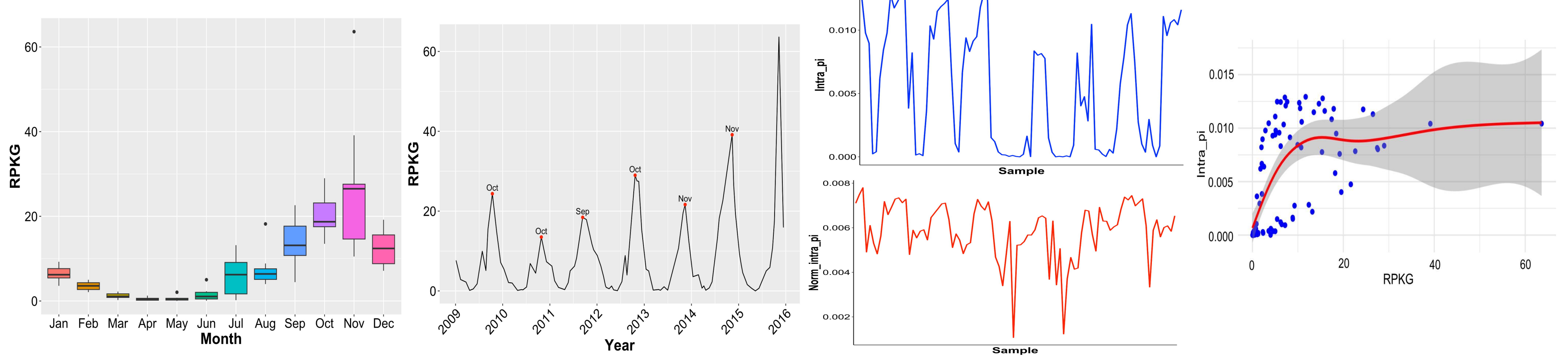

### *Prochlorococcus* Cyanobacteriia (BL_0908_bin.full.869)

#### BBMO

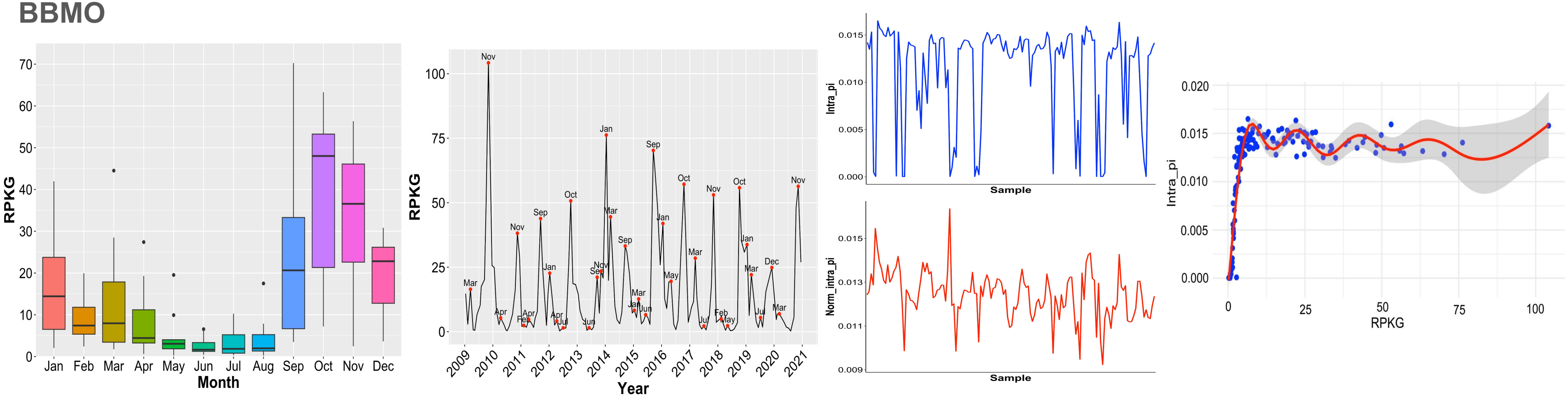


#### SOLA

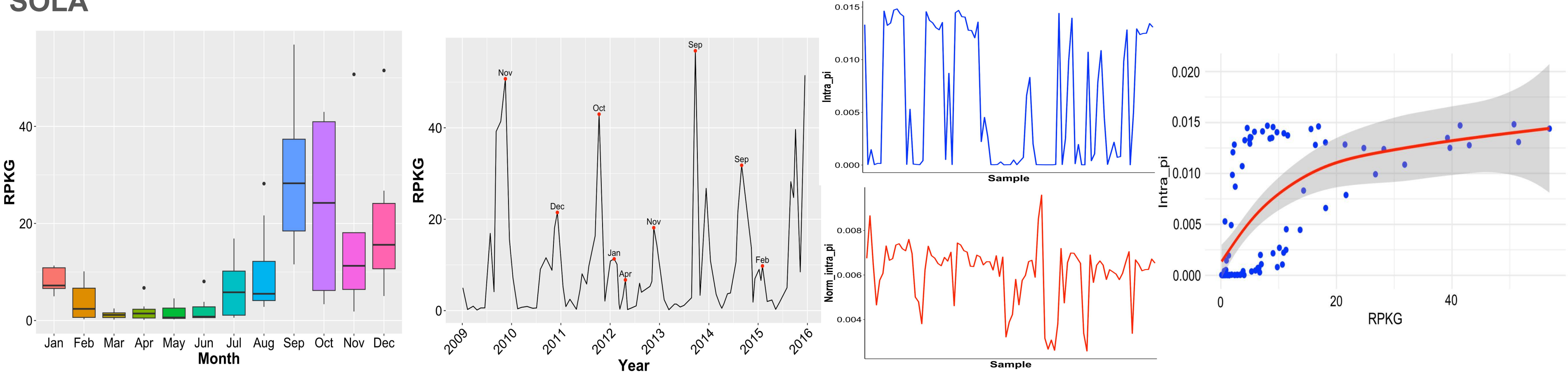

### *Polaribacter* **Bacteroidia** (BL_0902_bin.full.136)

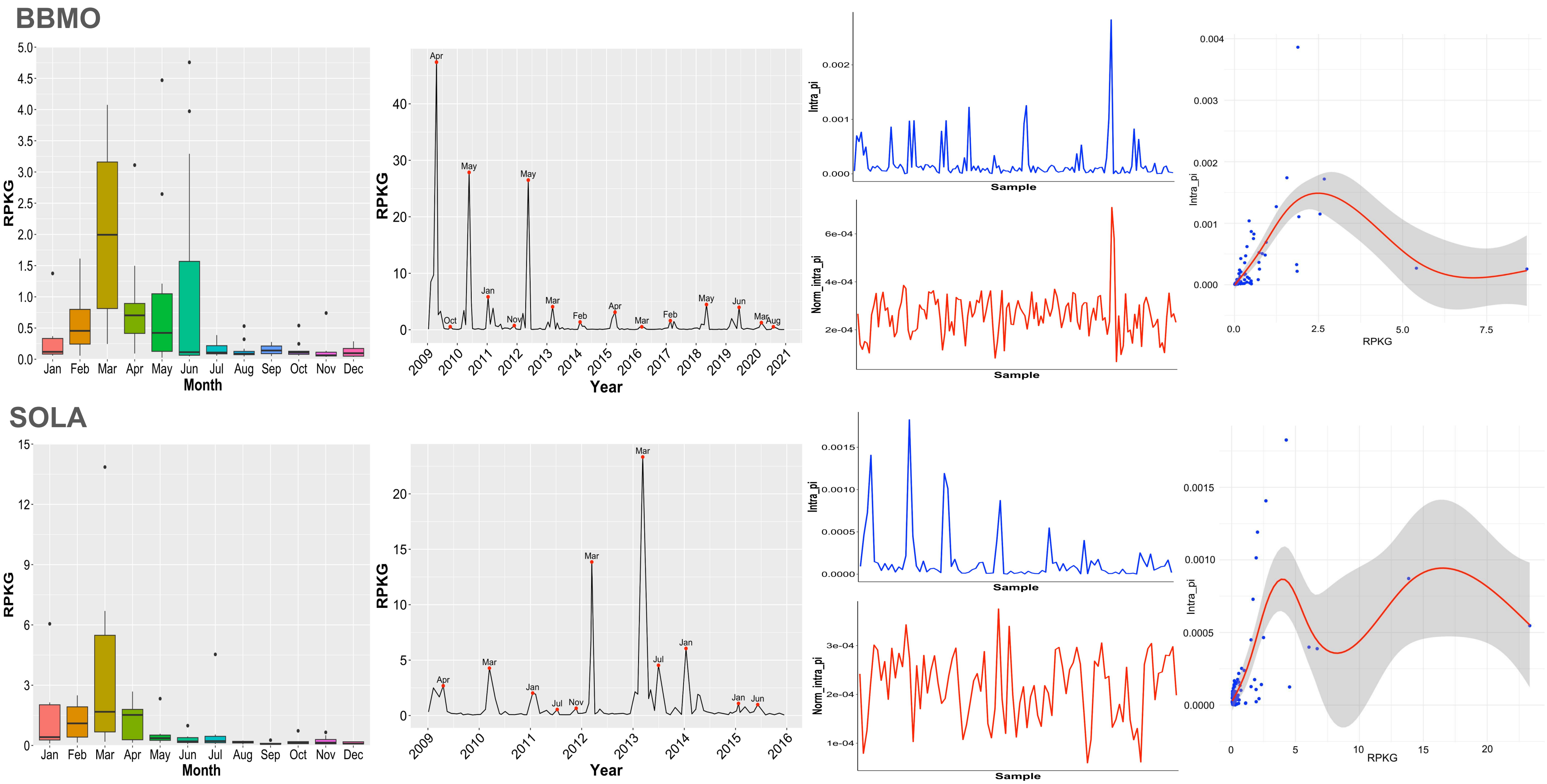

### *Sediminibacterium* **Bacteroidia** (BL_0908_bin.full.759)

#### BBMO

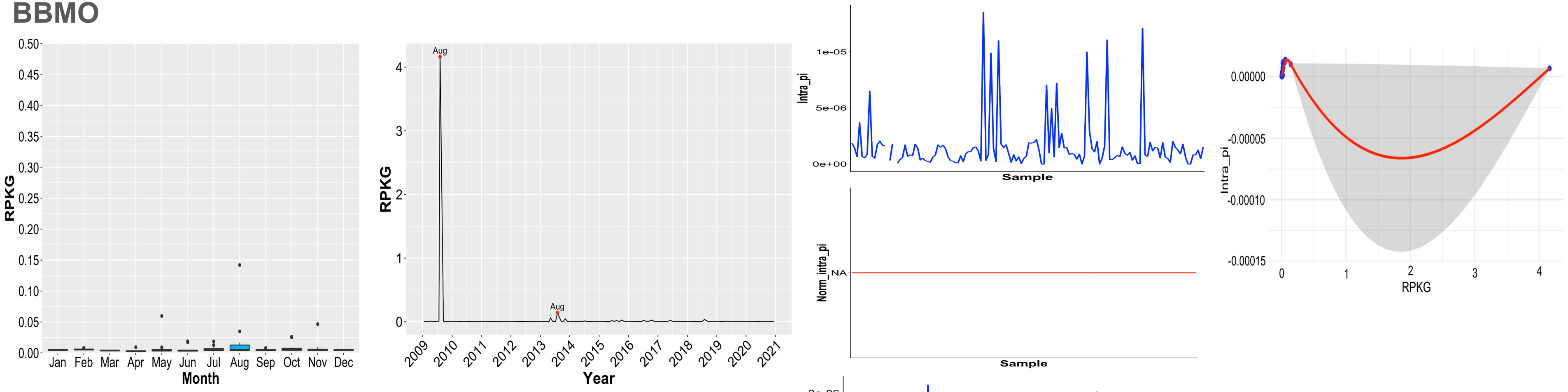


#### SOLA

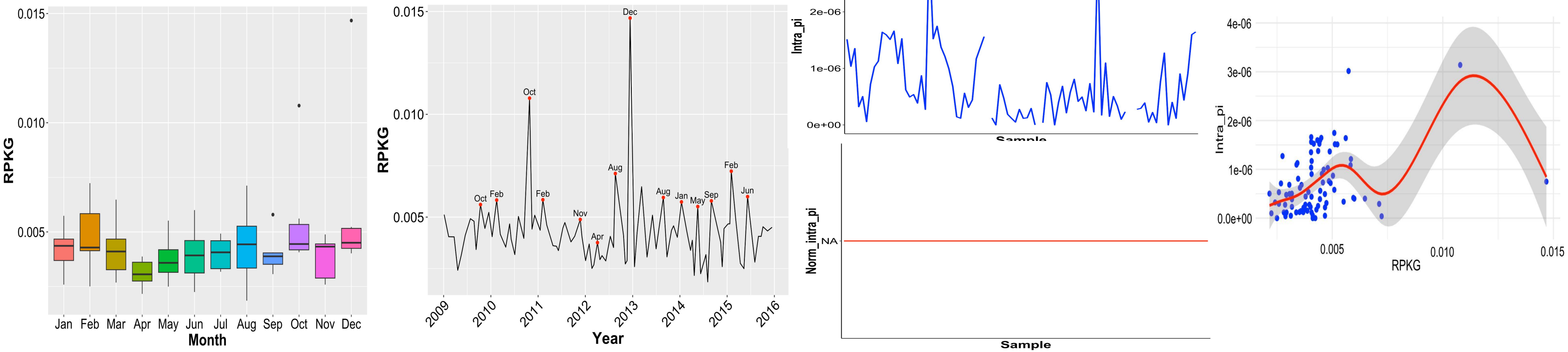

### **UBA7446 (Flavobacteriaceae) Bacteroidia** (S7.ctg000447l_BL_0908sc)

#### BBMO

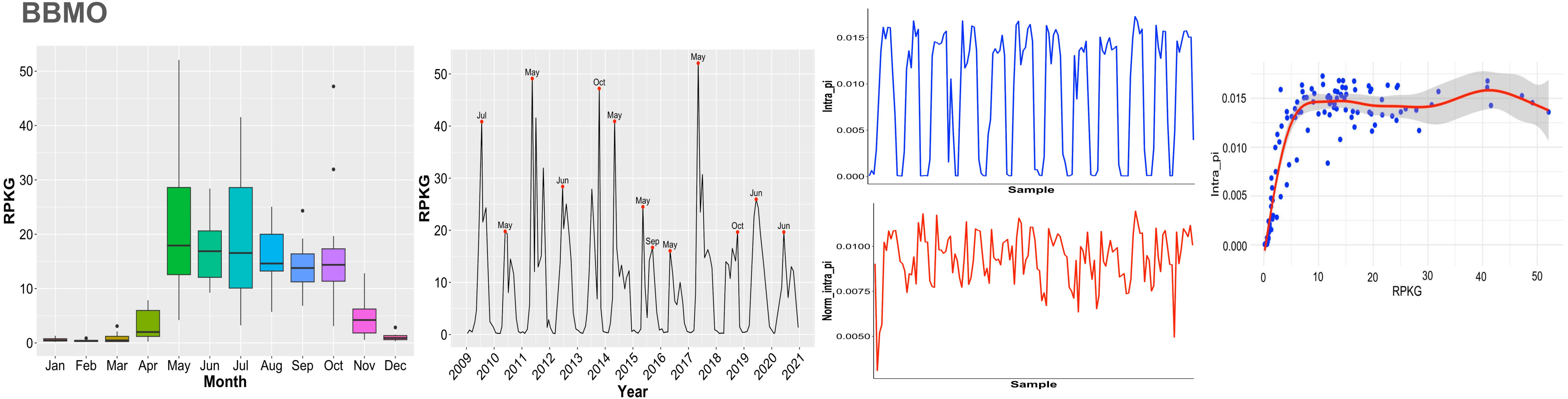


#### SOLA

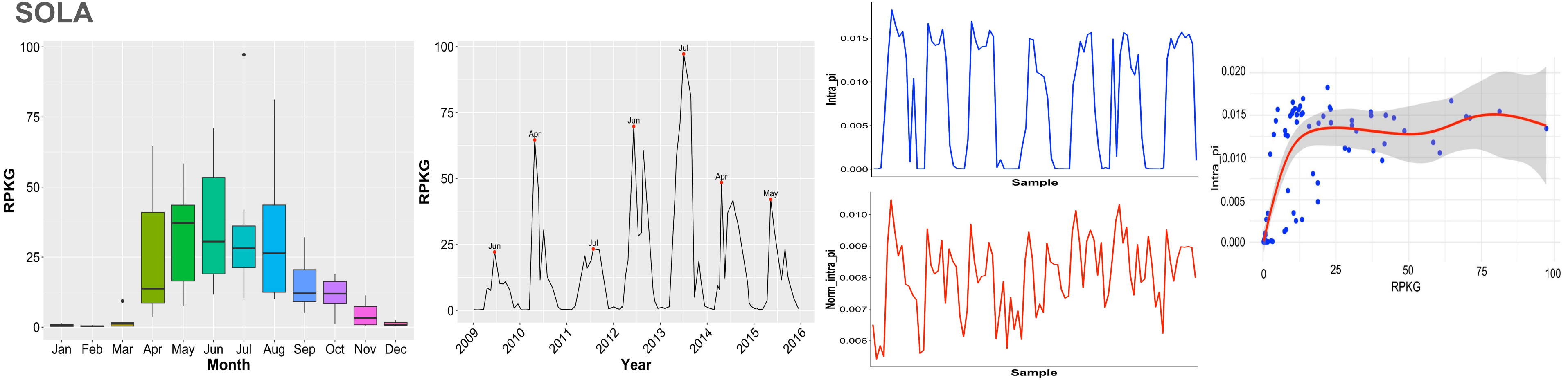

### **UBA8752 Bacteroidia** (BL_pooled_bin.circ.36)

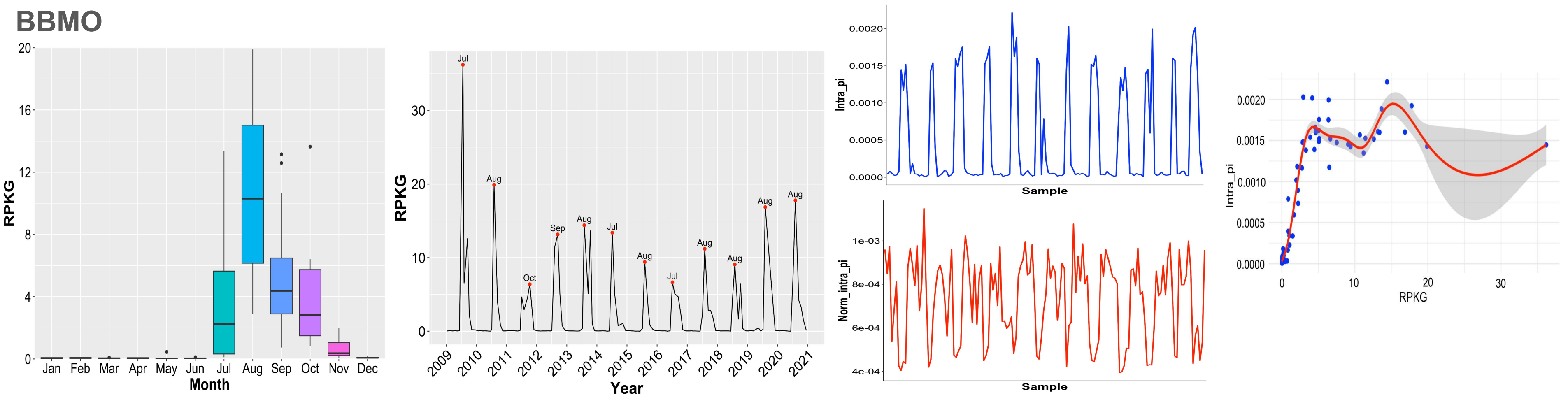


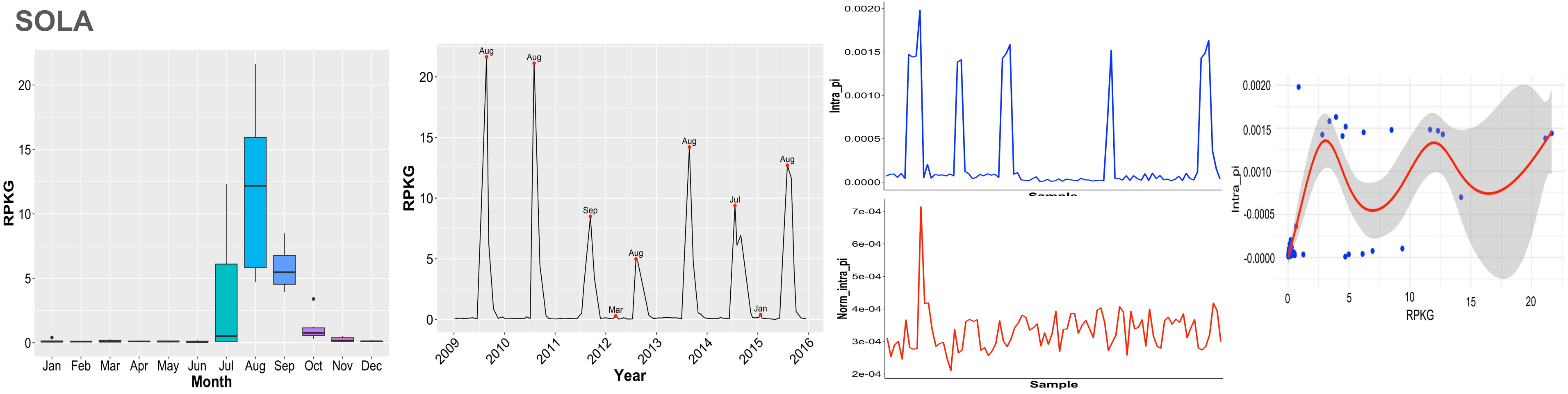

### **UBA7445 (Opitutales) Verrucomicrobia** (s249.ctg000277l_BL_0908sc)

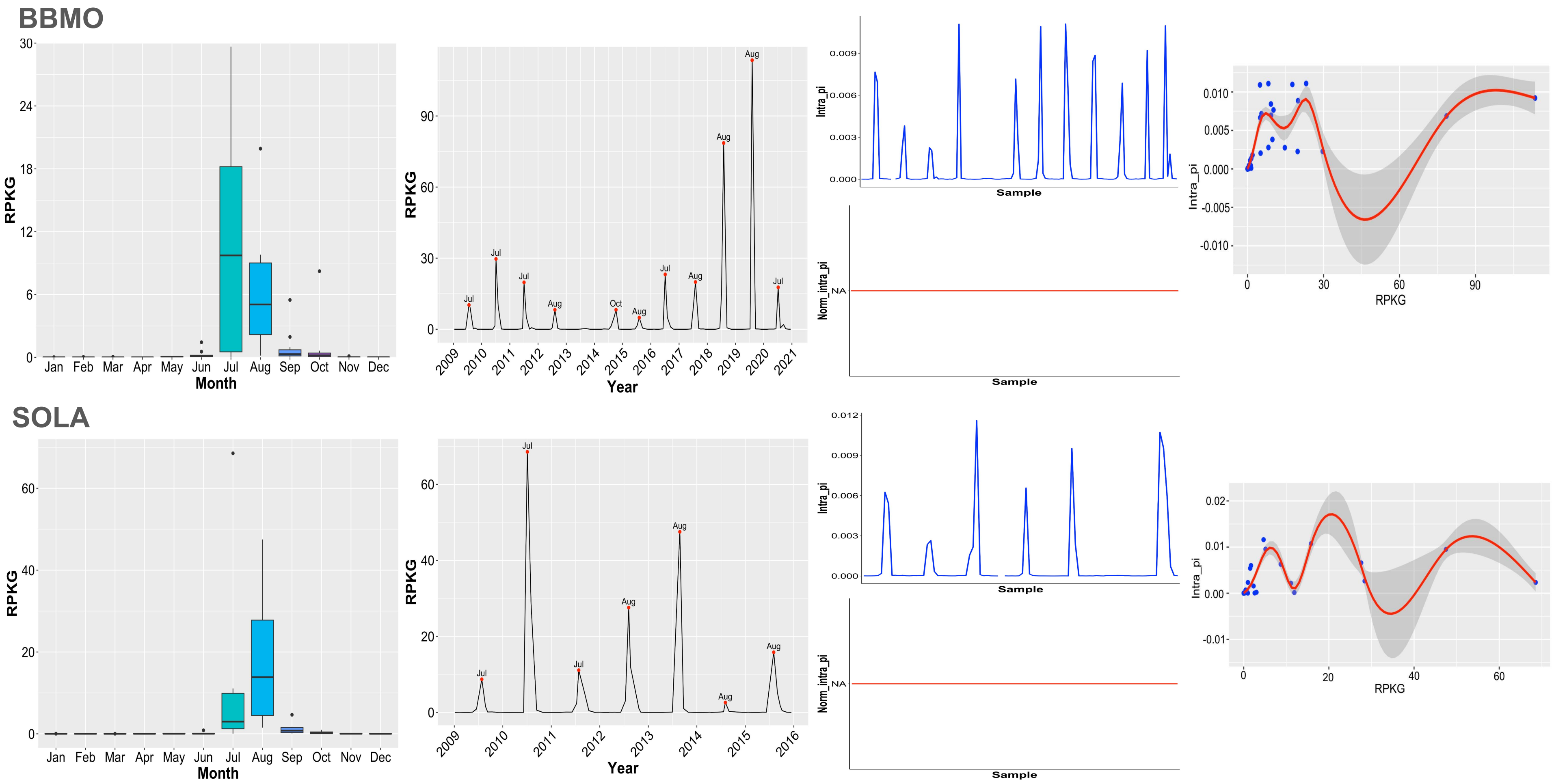

### UBA8653 Phycisphaerae (BL_0908_bin.full.20)

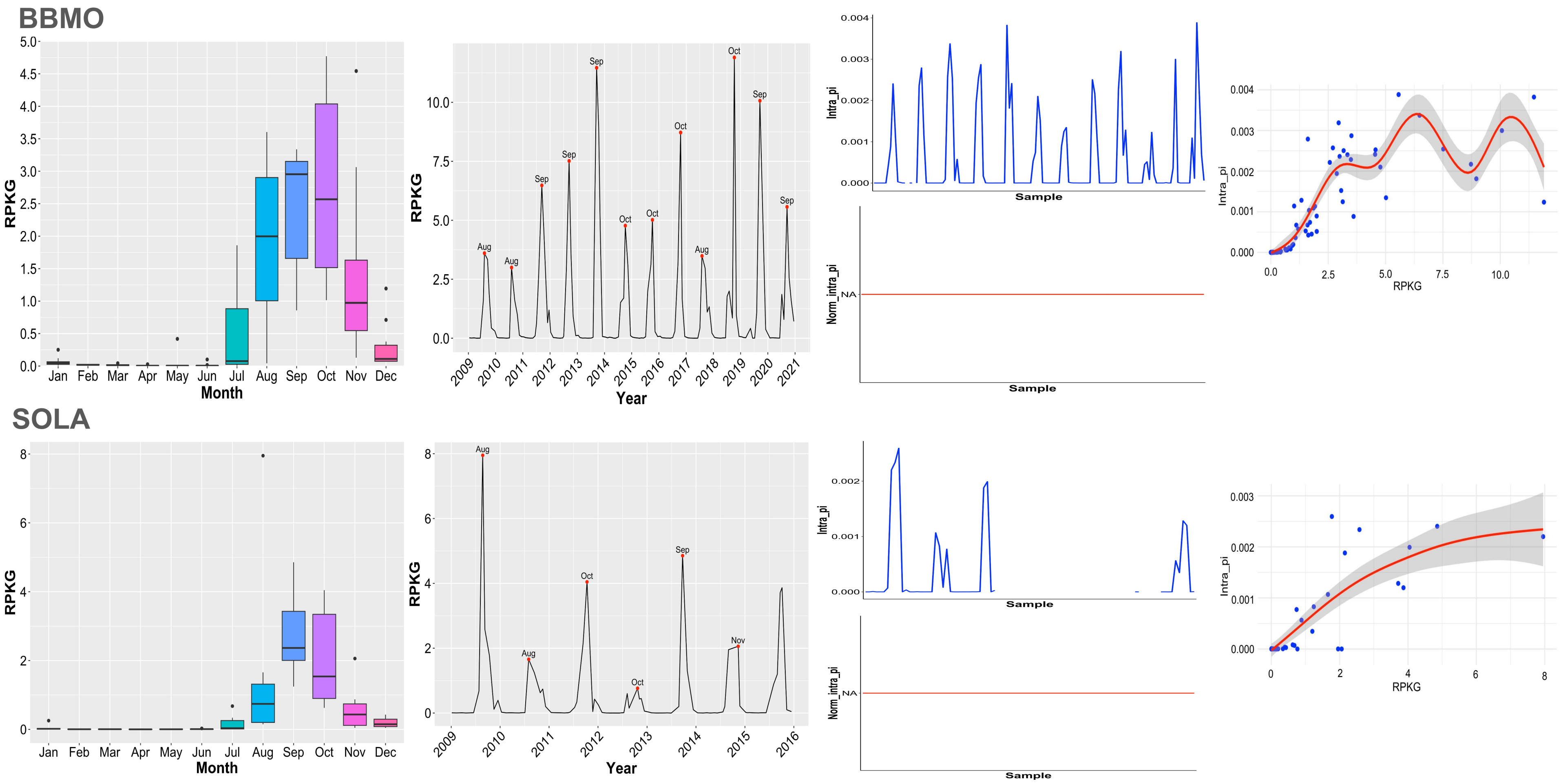

### **Ga0077546 (Obscuribacteraceae) Vampirovibrionia** (BL_pooled_bin.full.1551)

#### BBMO

#### SOLA

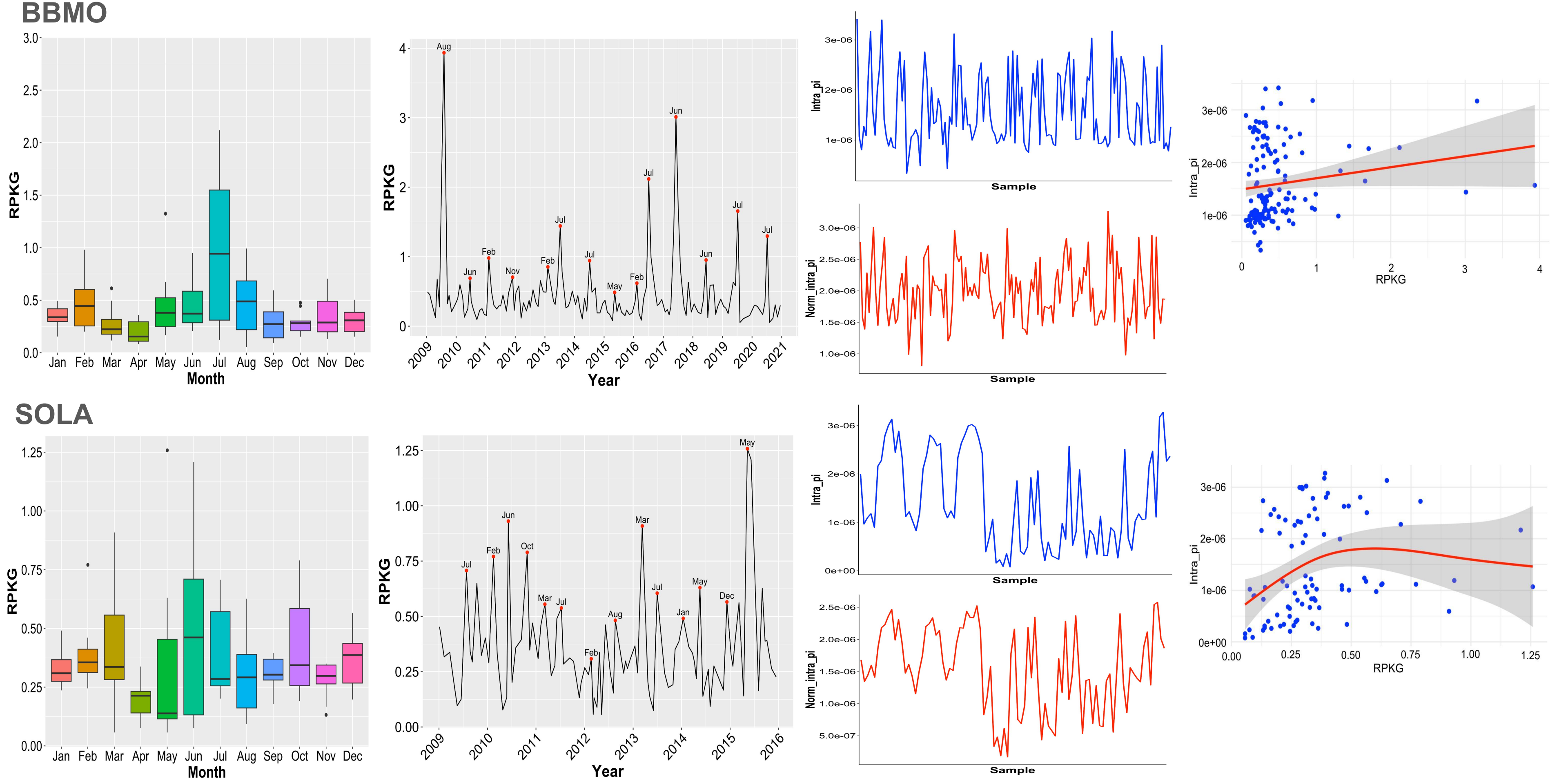
