## Supplementary Figure 8 for "Seasonal population structure and adaptive signatures across long-term marine microbial time series"

**Supplementary Figure 8. Horizontal and vertical coverage of the 30 long-read MAGs across the BBMO and SOLA time series.** Each point represents an individual metagenomic sample. Horizontal coverage (x-axis) indicates the percentage of the MAG covered by mapped reads, whereas vertical coverage (y-axis) indicates the mean sequencing depth across the mapped regions of the MAG. Point shapes indicate sampling location (BBMO or SOLA), and colors indicate seasonal state (Summer–Autumn or Winter–Spring). Each panel corresponds to one MAG, presented in the same order as in Table 1.

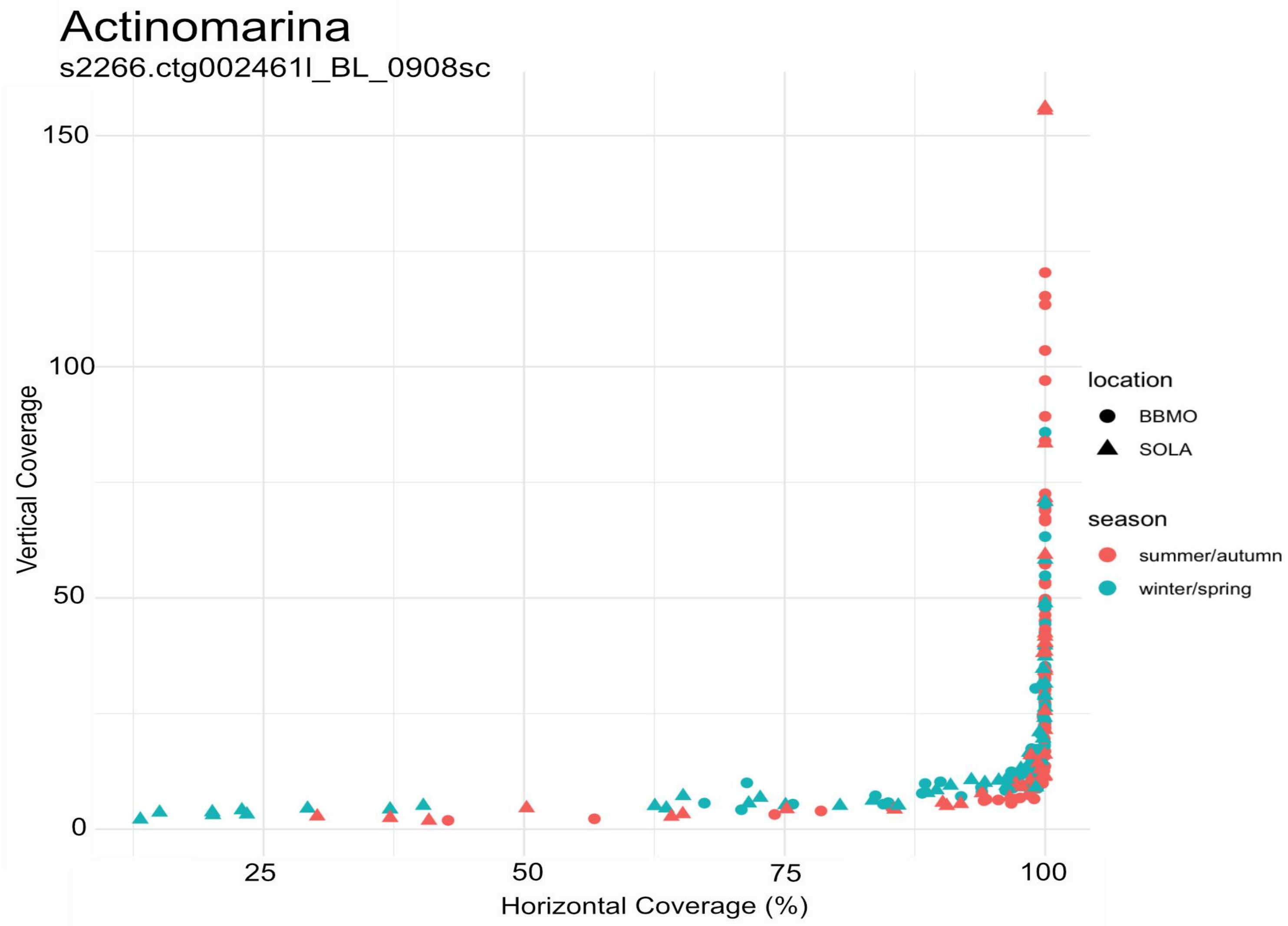
Actinomarina
s2266.ctg002461l_BL_0908sc
Vertical Coverage
150
100
50
0
25
50
75
100
Horizontal Coverage (%)
location
BBMO
SOLA
season
summer/autumn
winter/spring

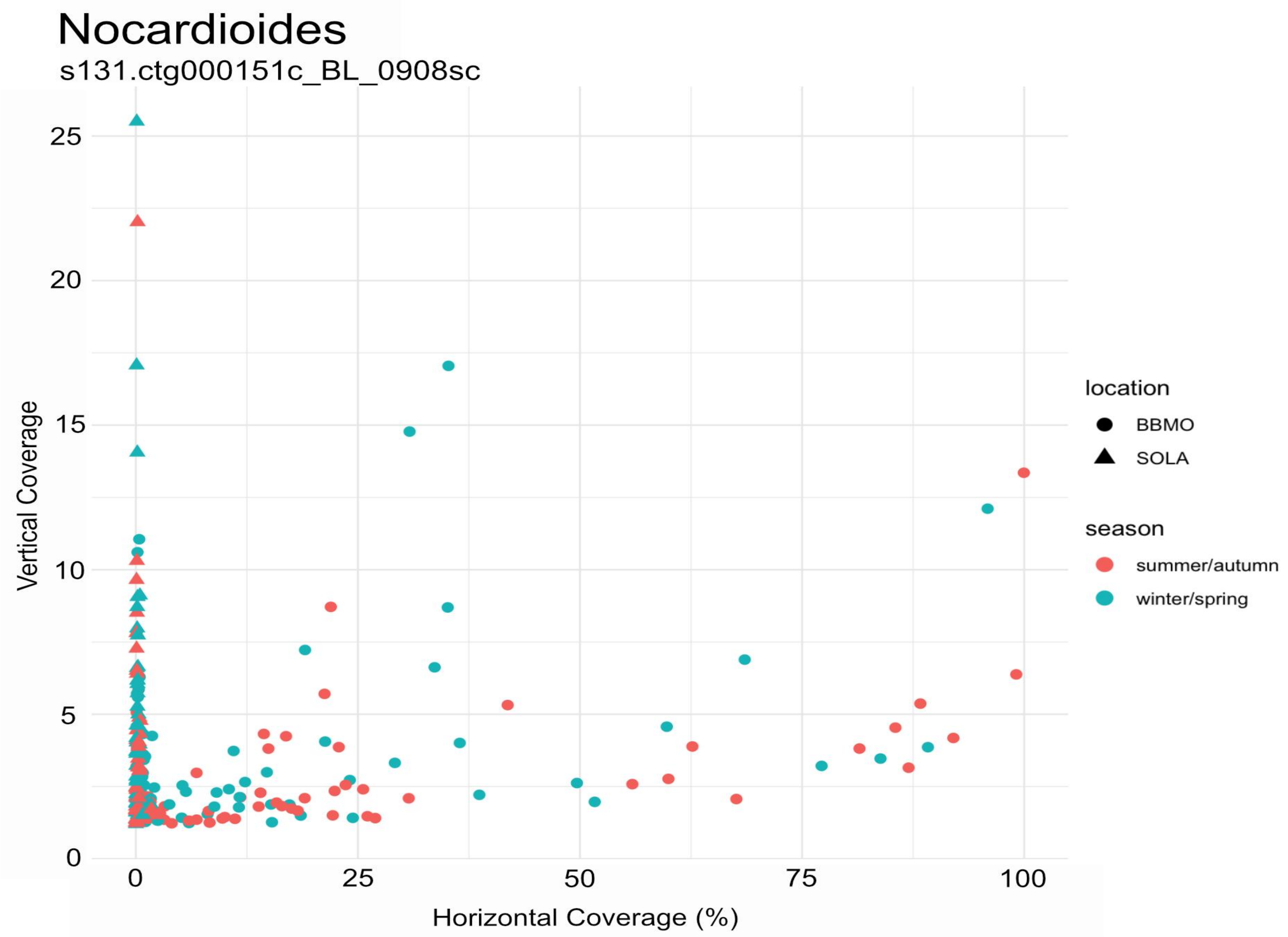
Nocardioides
s131.ctg000151c_BL_0908sc
Vertical Coverage
Horizontal Coverage (%)
0
5
10
15
20
25
0
25
50
75
100
location
BBMO
SOLA
season
summer/autumn
winter/spring

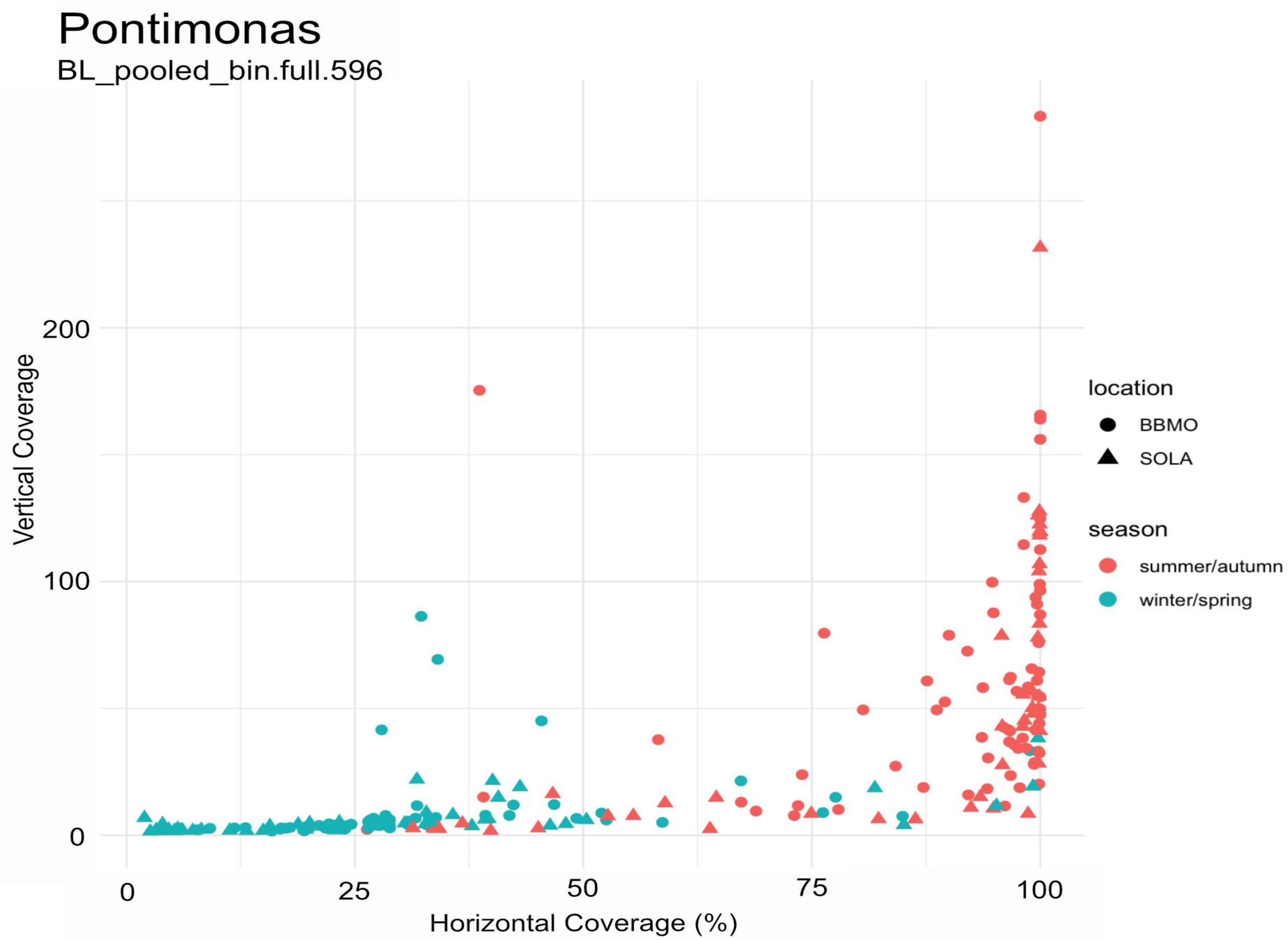
Pontimonas
BL_pooled_bin.full.596
Vertical Coverage
200
100
0
0
25
50
75
100
Horizontal Coverage (%)
location
BBMO
SOLA
season
summer/autumn
winter/spring

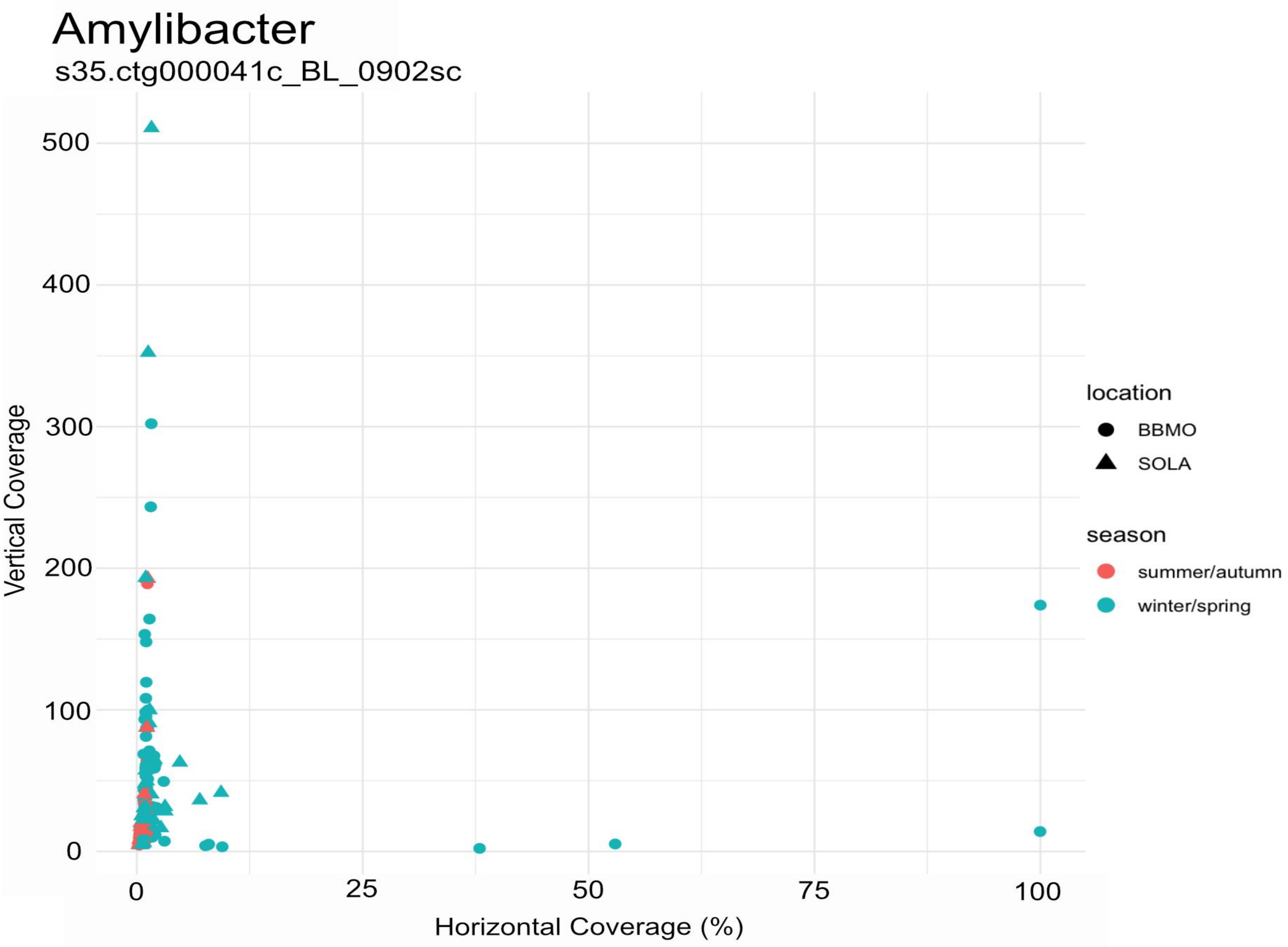
Amylibacter
s35.ctg000041c_BL_0902sc
Vertical Coverage
Horizontal Coverage (%)
0
100
200
300
400
500
0
25
50
75
100
location
BBMO
SOLA
season
summer/autumn
winter/spring

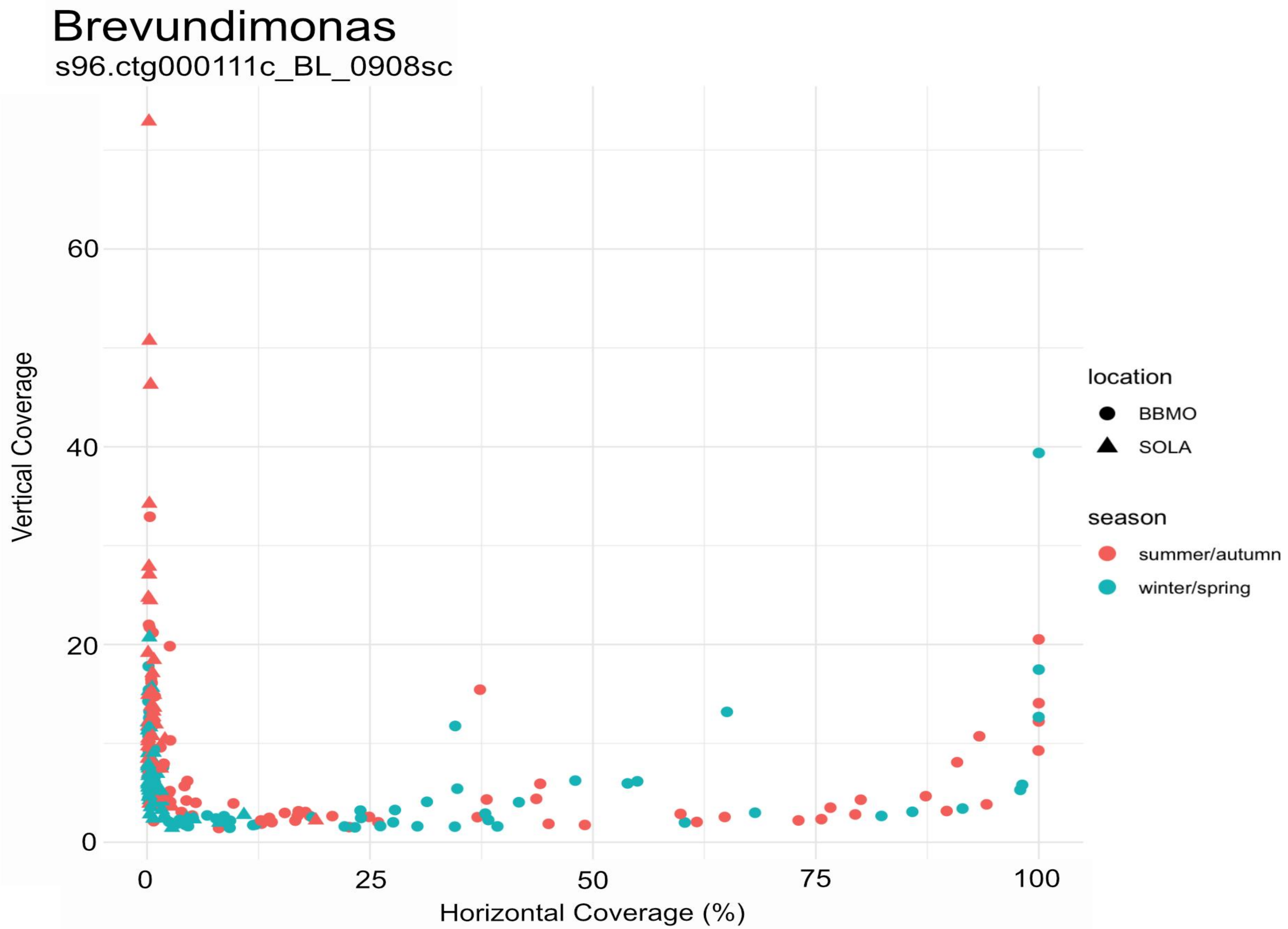
Brevundimonas
s96.ctg000111c_BL_0908sc
Vertical Coverage
Horizontal Coverage (%)
0
20
40
60
0
25
50
75
100
location
BBMO
SOLA
season
summer/autumn
winter/spring

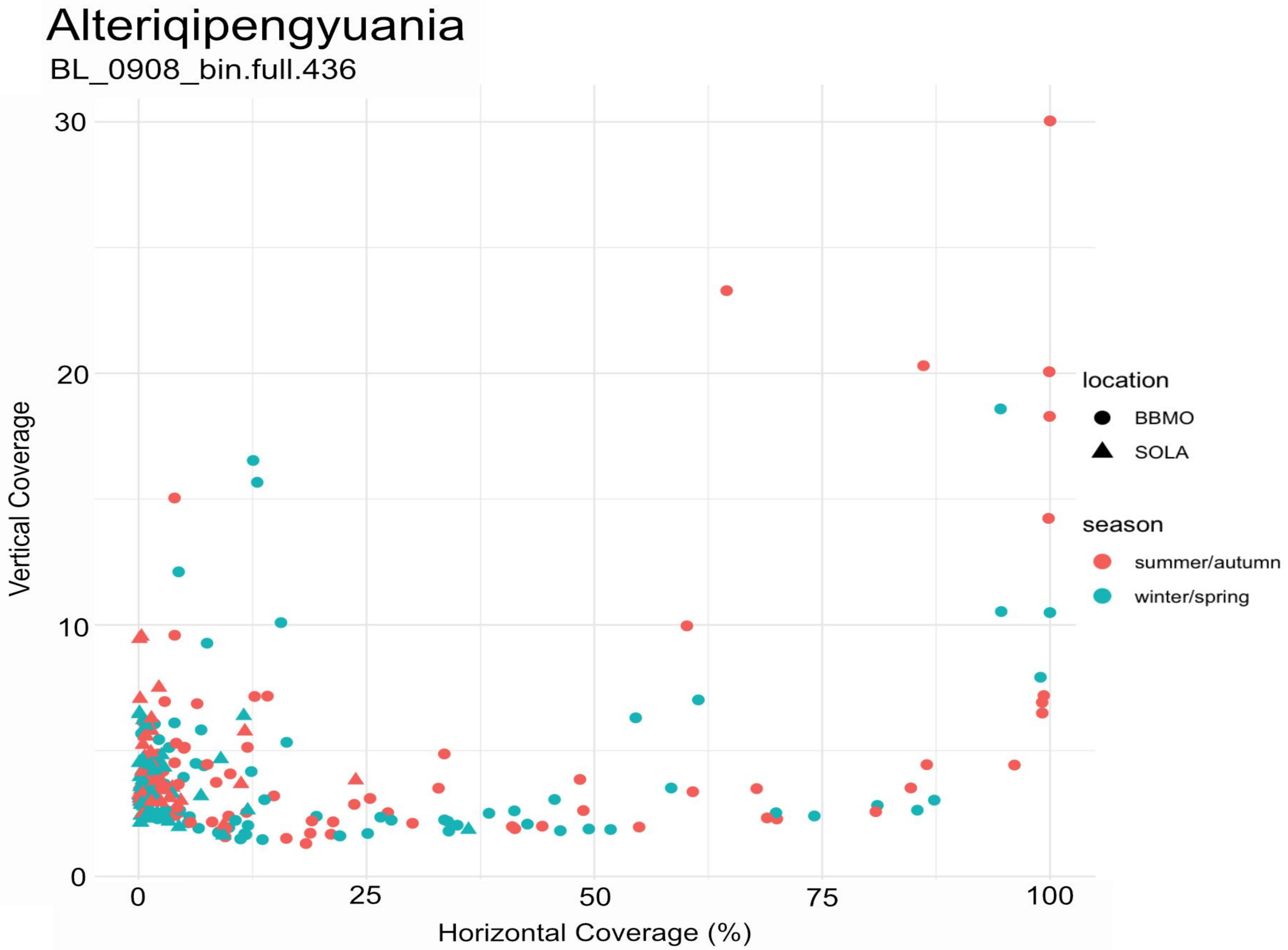

Alteriqipengyuania
BL_0908_bin.full.436
Vertical Coverage
30
20
10
0
0
25
50
75
100
Horizontal Coverage (%)
location
BBMO
SOLA
season
summer/autumn
winter/spring

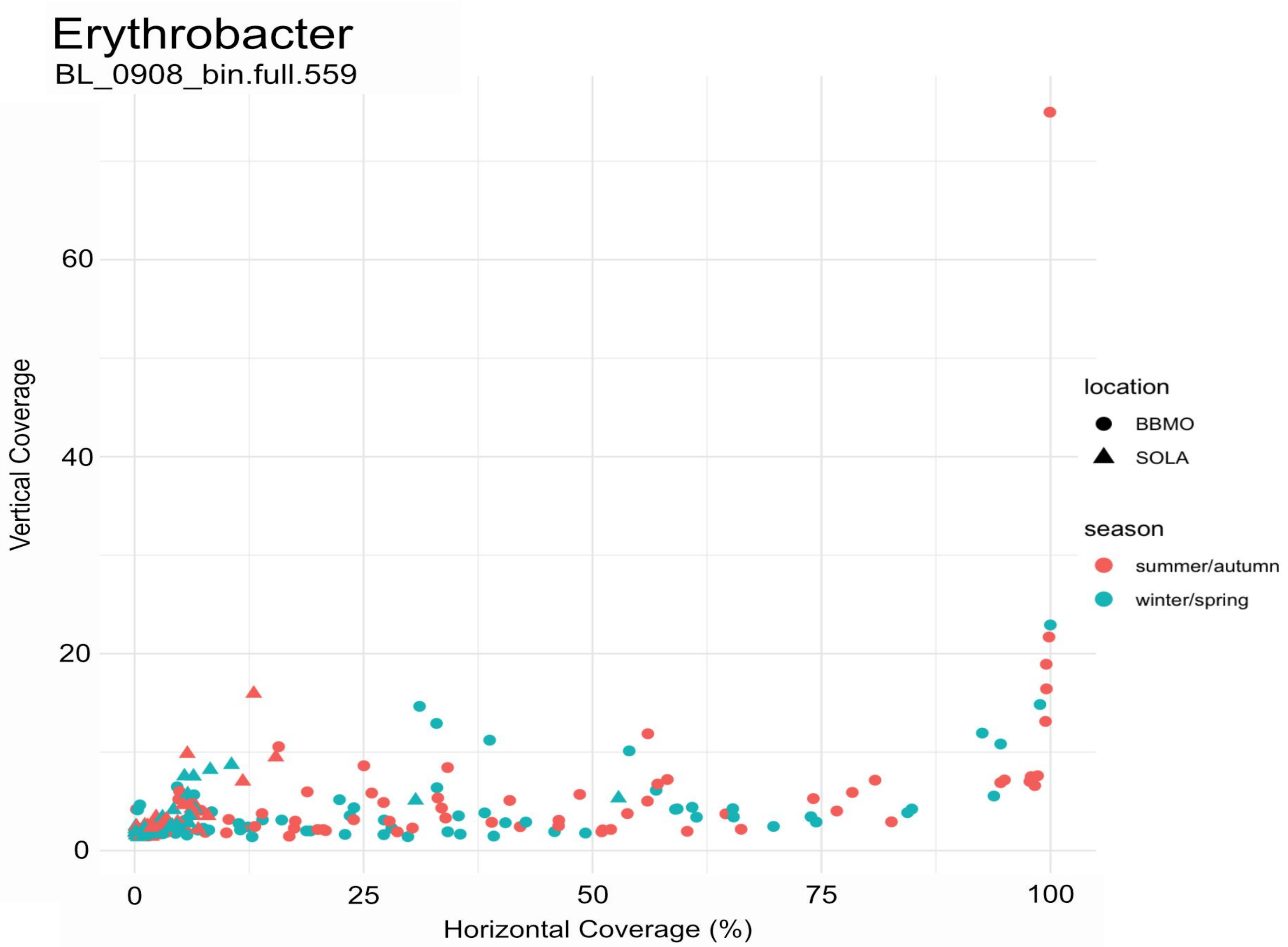

Erythrobacter
BL_0908_bin.full.559
Vertical Coverage
60
40
20
0
0
25
50
75
100
Horizontal Coverage (%)
location
BBMO
SOLA
season
summer/autumn
winter/spring

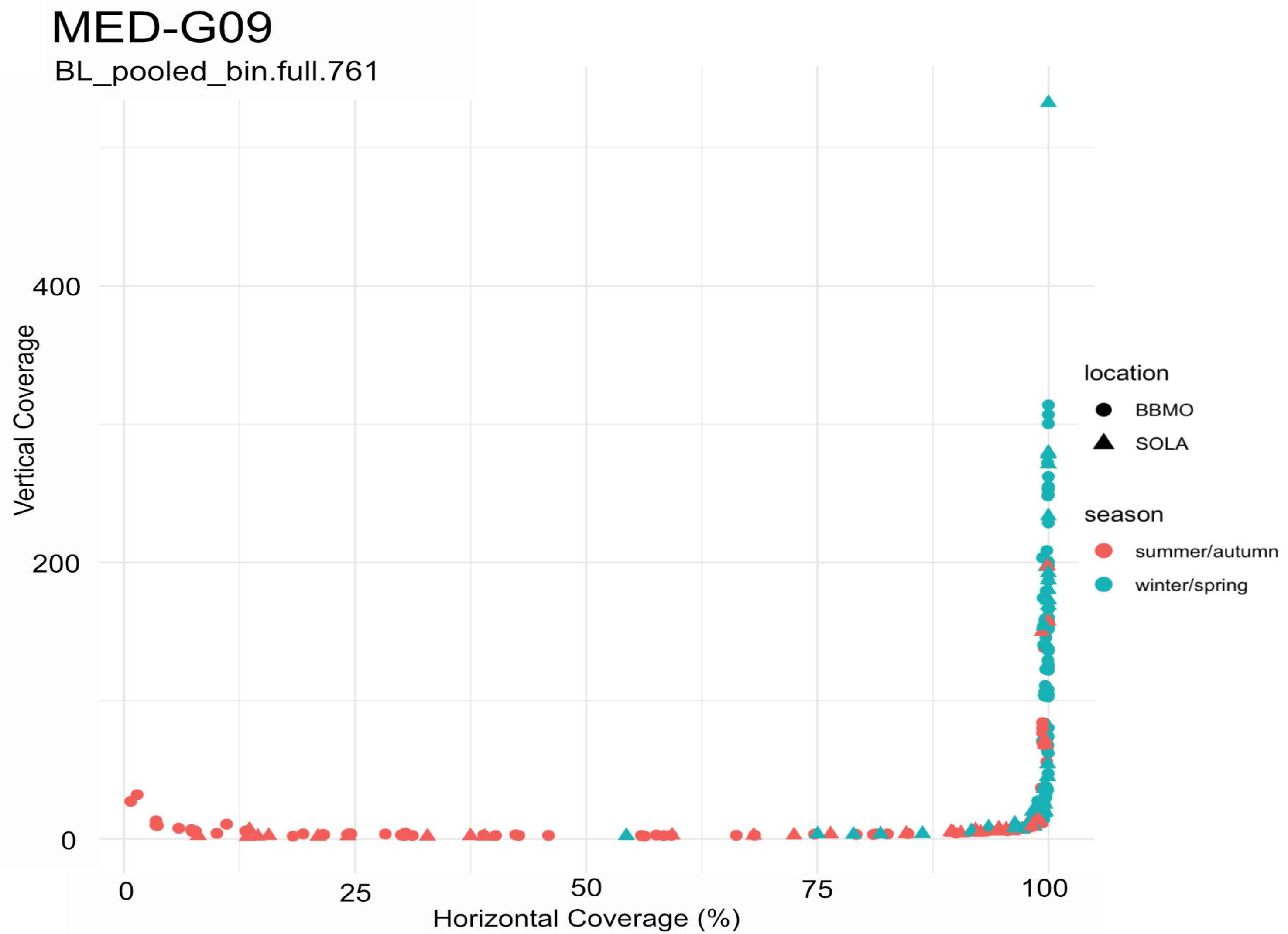
MED-G09
BL_pooled_bin.full.761
Vertical Coverage
400
200
0
0
25
50
75
100
Horizontal Coverage (%)
location
BBMO
SOLA
season
summer/autumn
winter/spring

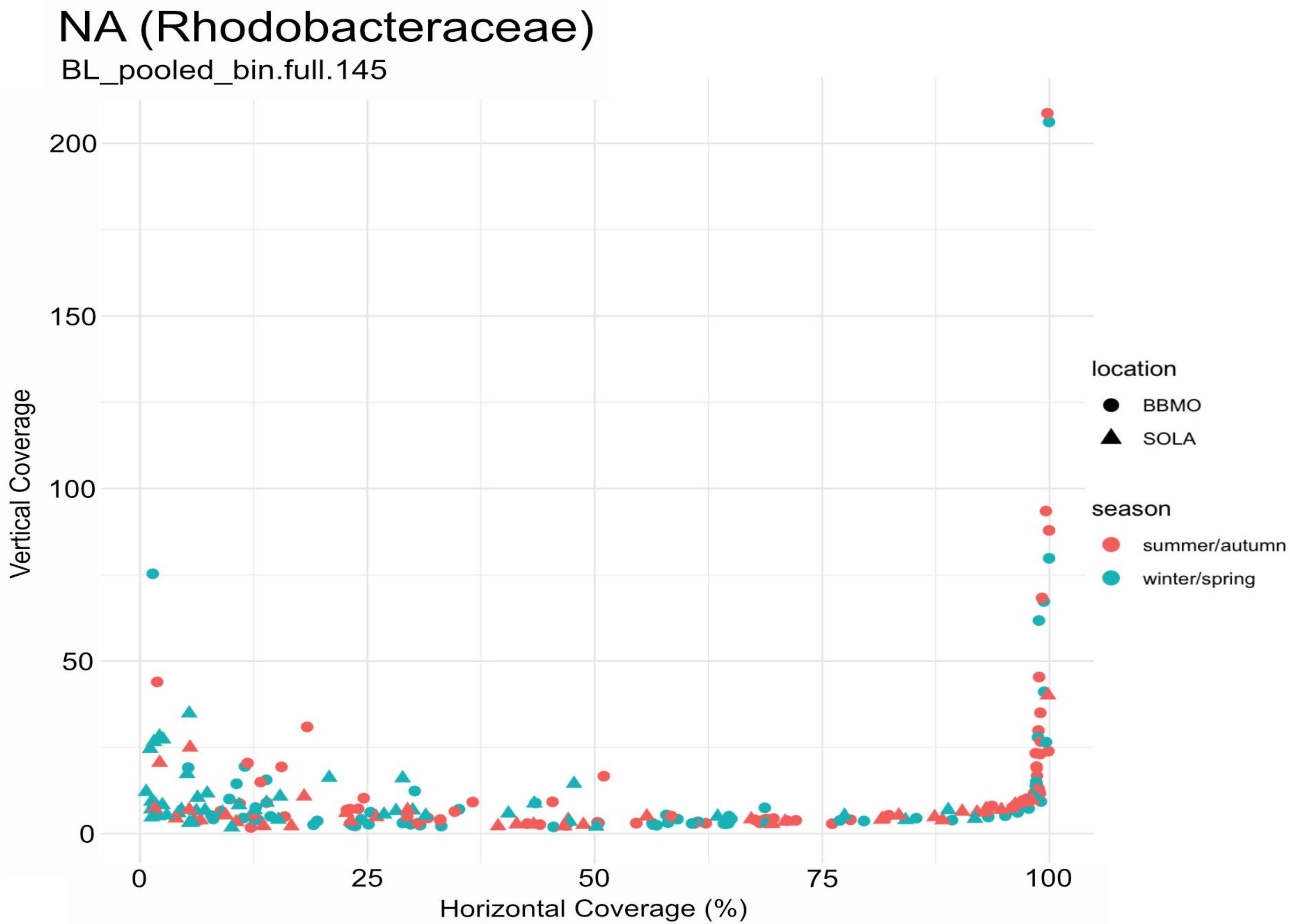
NA (Rhodobacteraceae)
BL_pooled_bin.full.145
Vertical Coverage
200
150
100
50
0
0
25
50
75
100
Horizontal Coverage (%)
location
BBMO
SOLA
season
summer/autumn
winter/spring

### SAR11: HIMB114 (Pelagibacteraceae)

s1032.ctg001132l_BL_0908sc

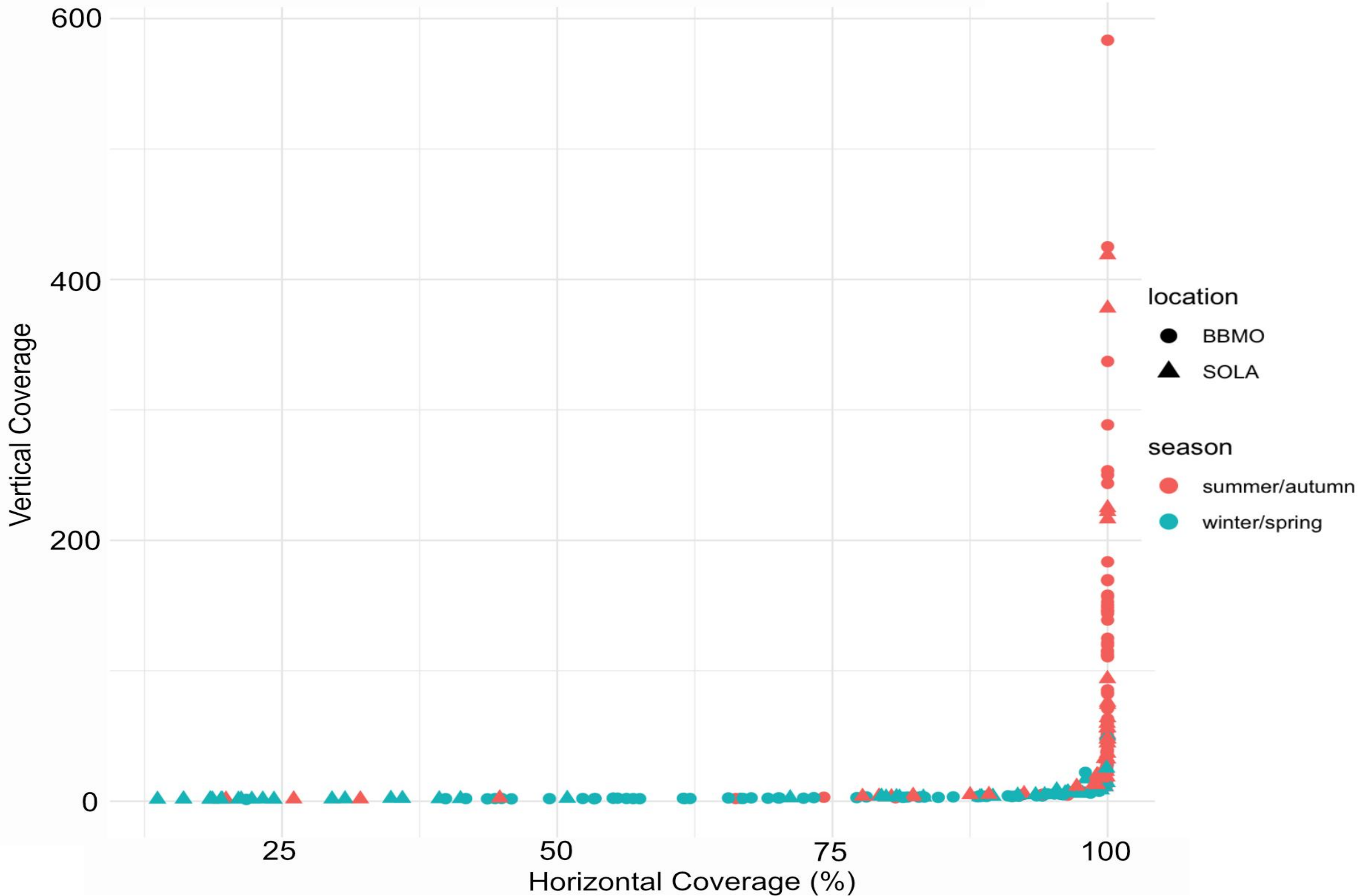

### SAR11: Pelagibacter

s266.ctg000297l_BL_0908sc

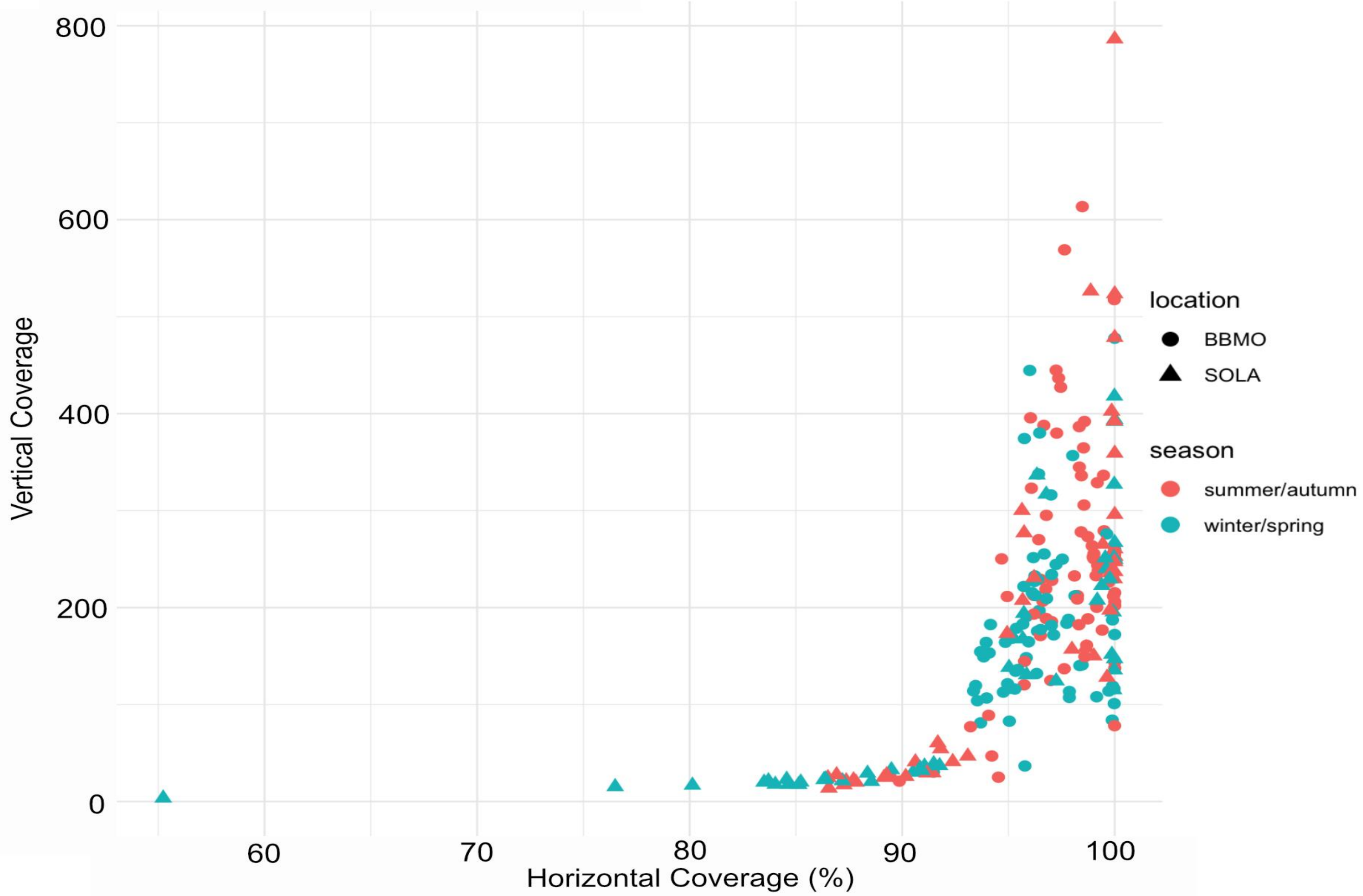

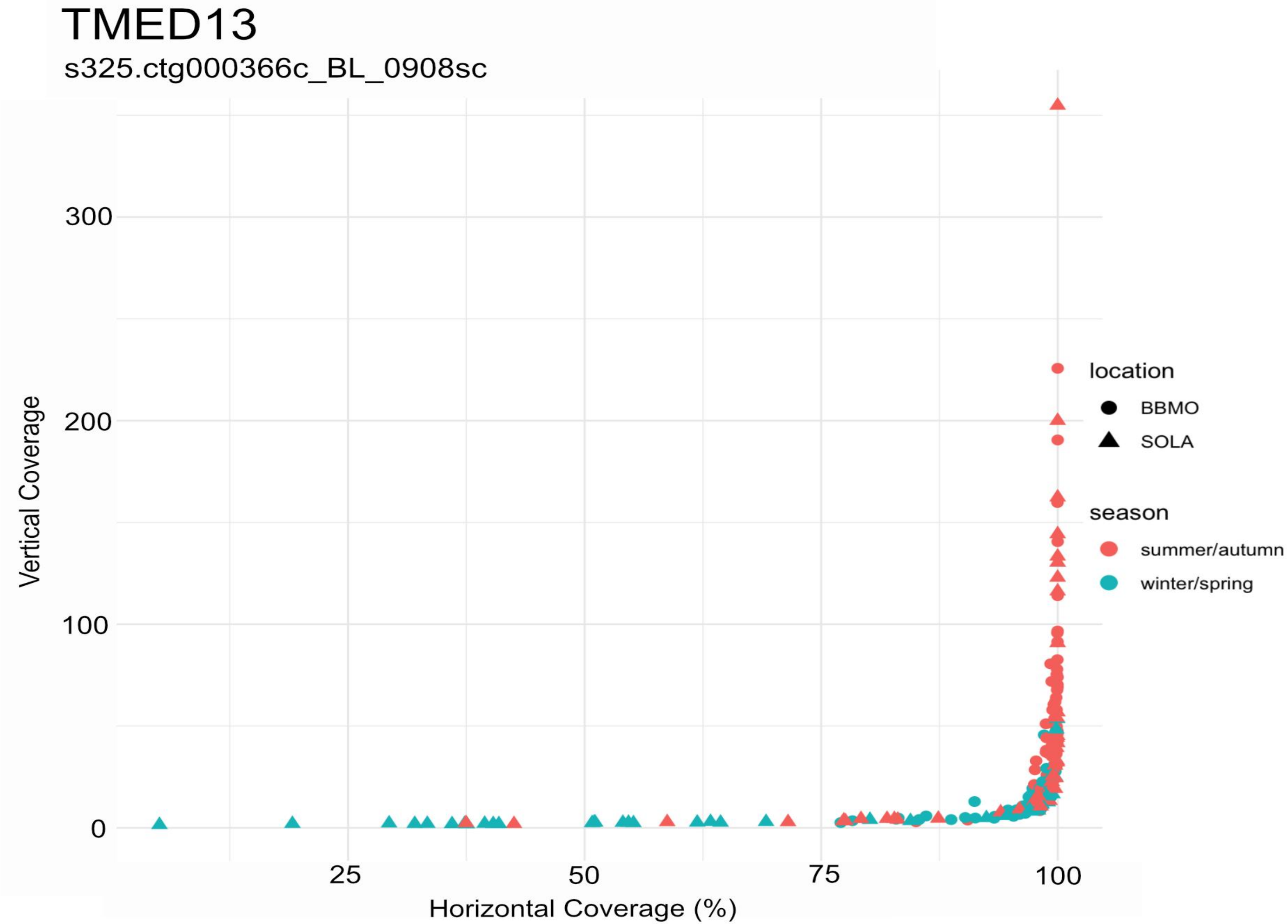
TMED13
s325.ctg000366c_BL_0908sc
Vertical Coverage
Horizontal Coverage (%)
0
100
200
300
25
50
75
100
location
BBMO
SOLA
season
summer/autumn
winter/spring

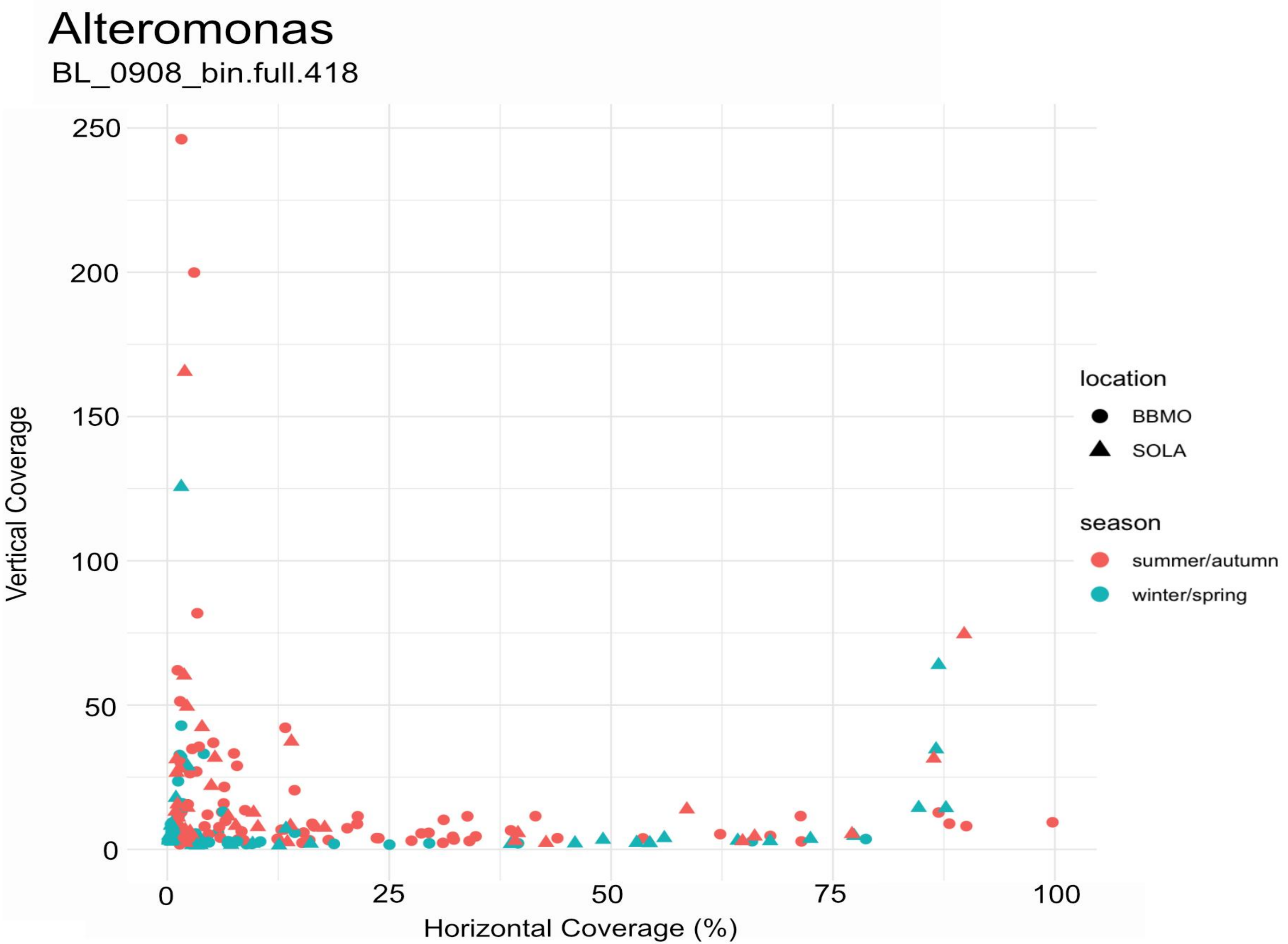

Alteromonas
BL_0908_bin.full.418
Vertical Coverage
250
200
150
100
50
0
0
25
50
75
100
Horizontal Coverage (%)
location
BBMO
SOLA
season
summer/autumn
winter/spring

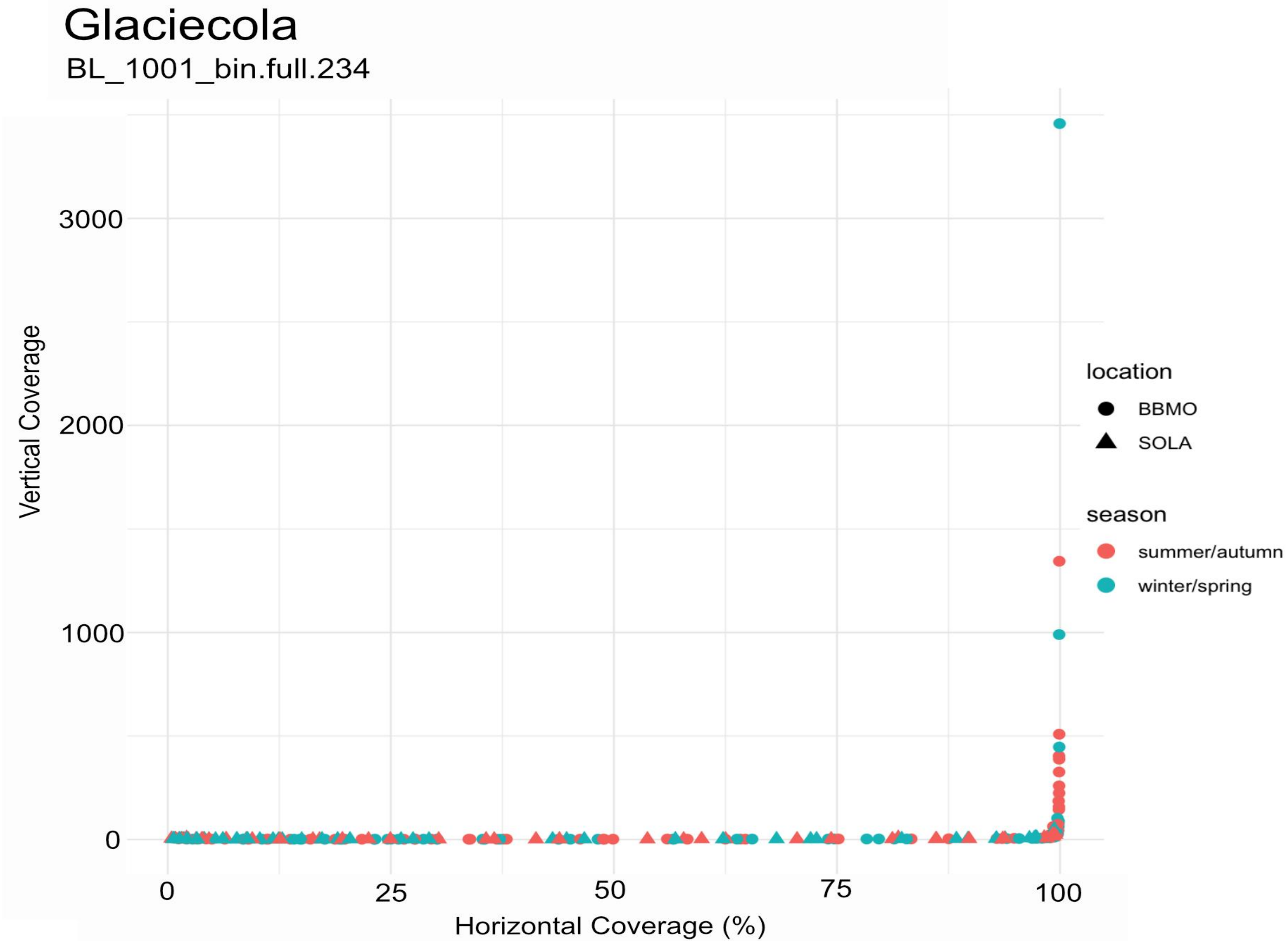

Glaciecola
BL_1001_bin.full.234
Vertical Coverage
3000
2000
1000
0
0
25
50
75
100
Horizontal Coverage (%)
location
BBMO
SOLA
season
summer/autumn
winter/spring

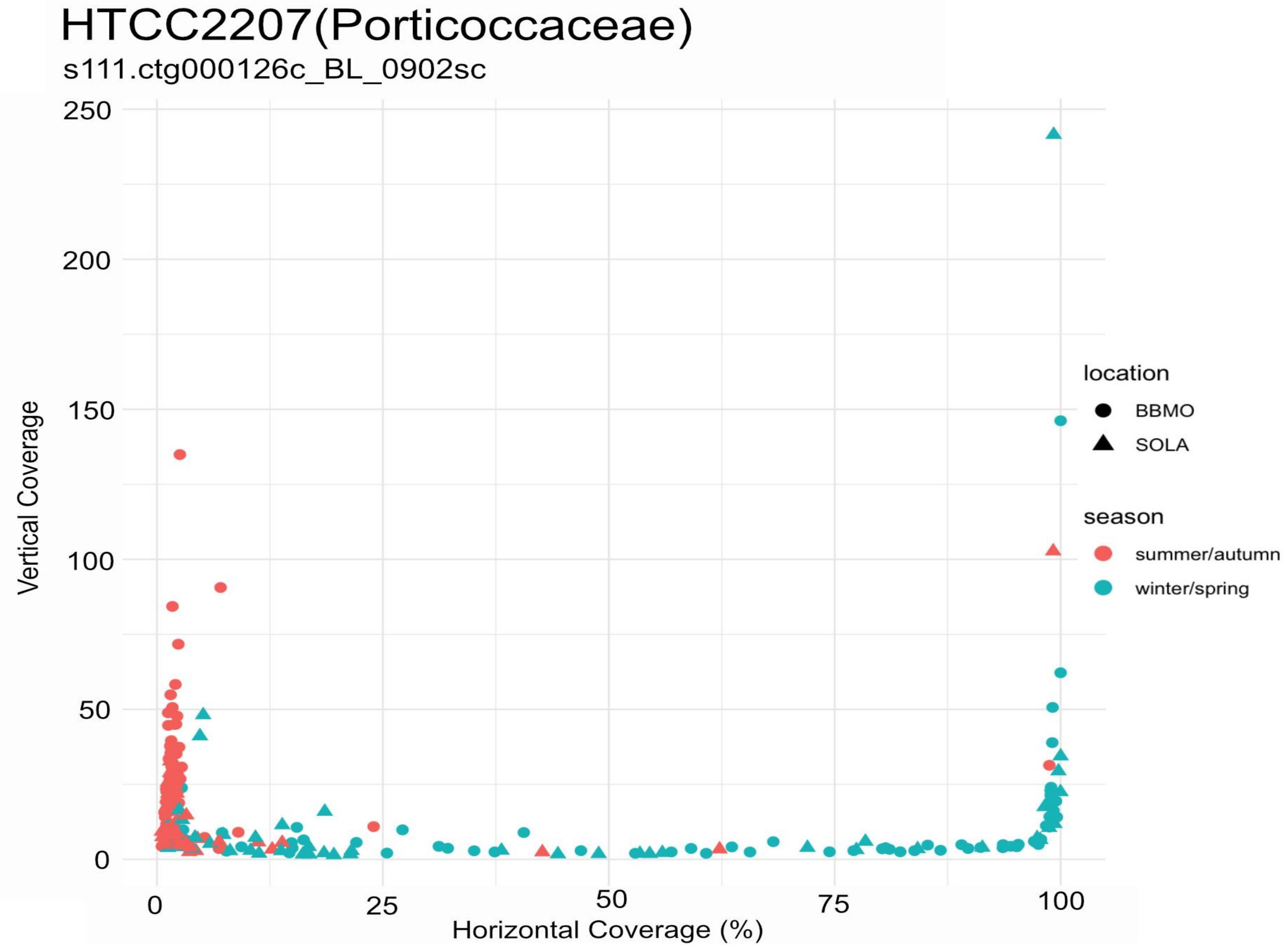
HTCC2207(Porticoccaceae)
s111.ctg000126c_BL_0902sc
Vertical Coverage
Horizontal Coverage (%)
250
200
150
100
50
0
0
25
50
75
100
location
BBMO
SOLA
season
summer/autumn
winter/spring

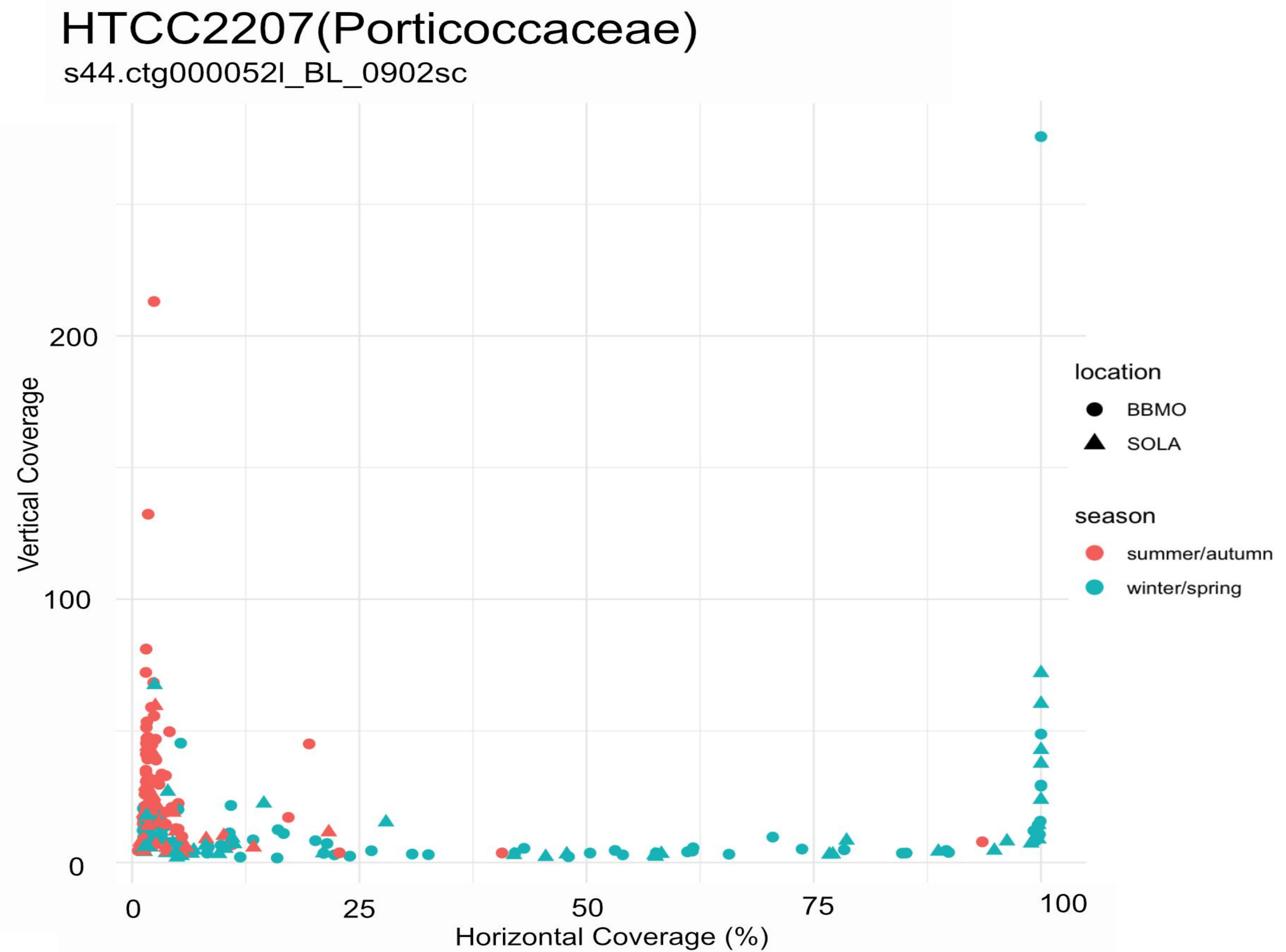
HTCC2207(Porticoccaceae)
s44.ctg000052l_BL_0902sc
Vertical Coverage
0
100
200
Horizontal Coverage (%)
0
25
50
75
100
location
BBMO
SOLA
season
summer/autumn
winter/spring

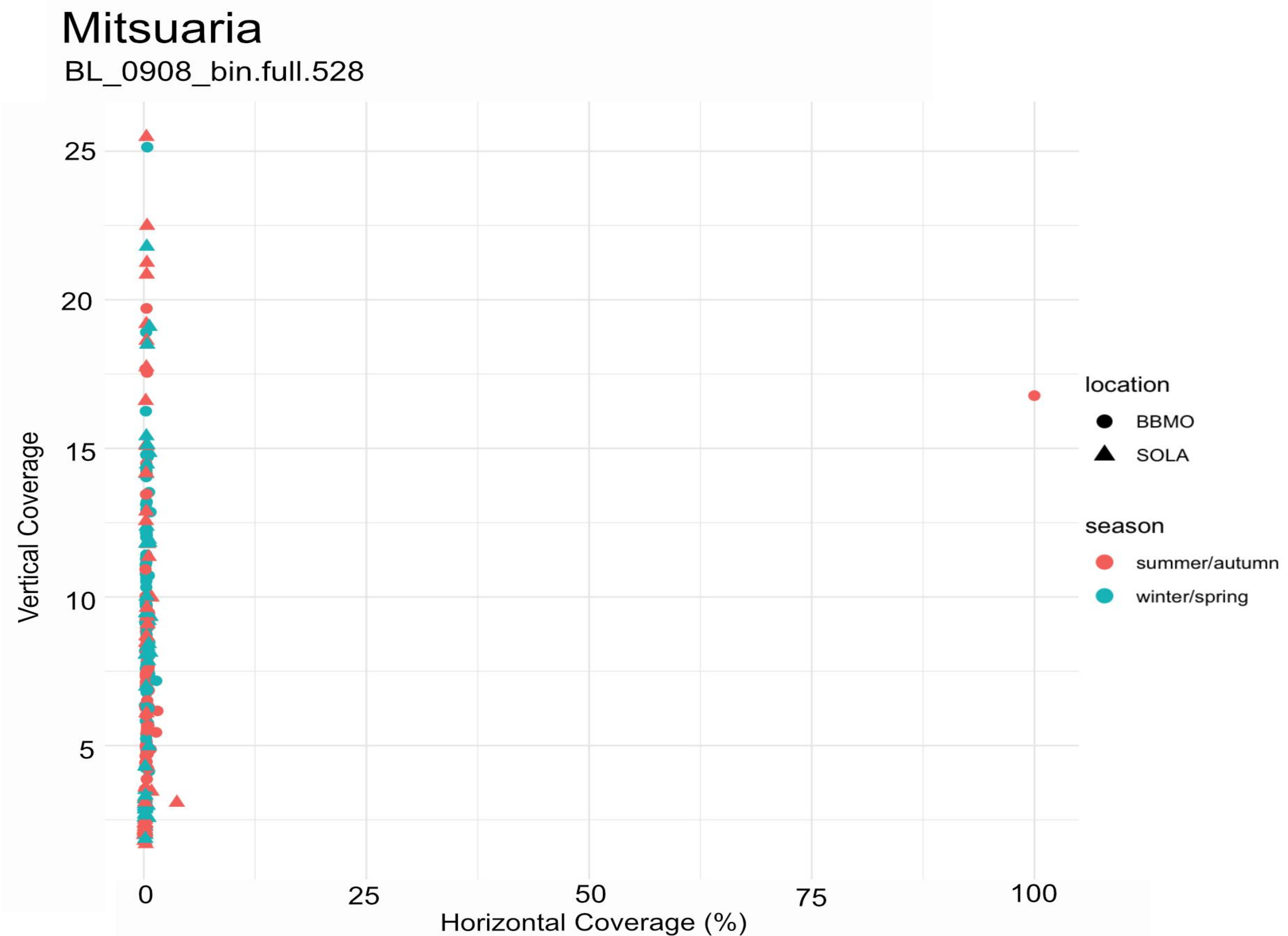
Mitsuaria
BL_0908_bin.full.528
Vertical Coverage
Horizontal Coverage (%)
25
20
15
10
5
0
25
50
75
100
location
BBMO
SOLA
season
summer/autumn
winter/spring

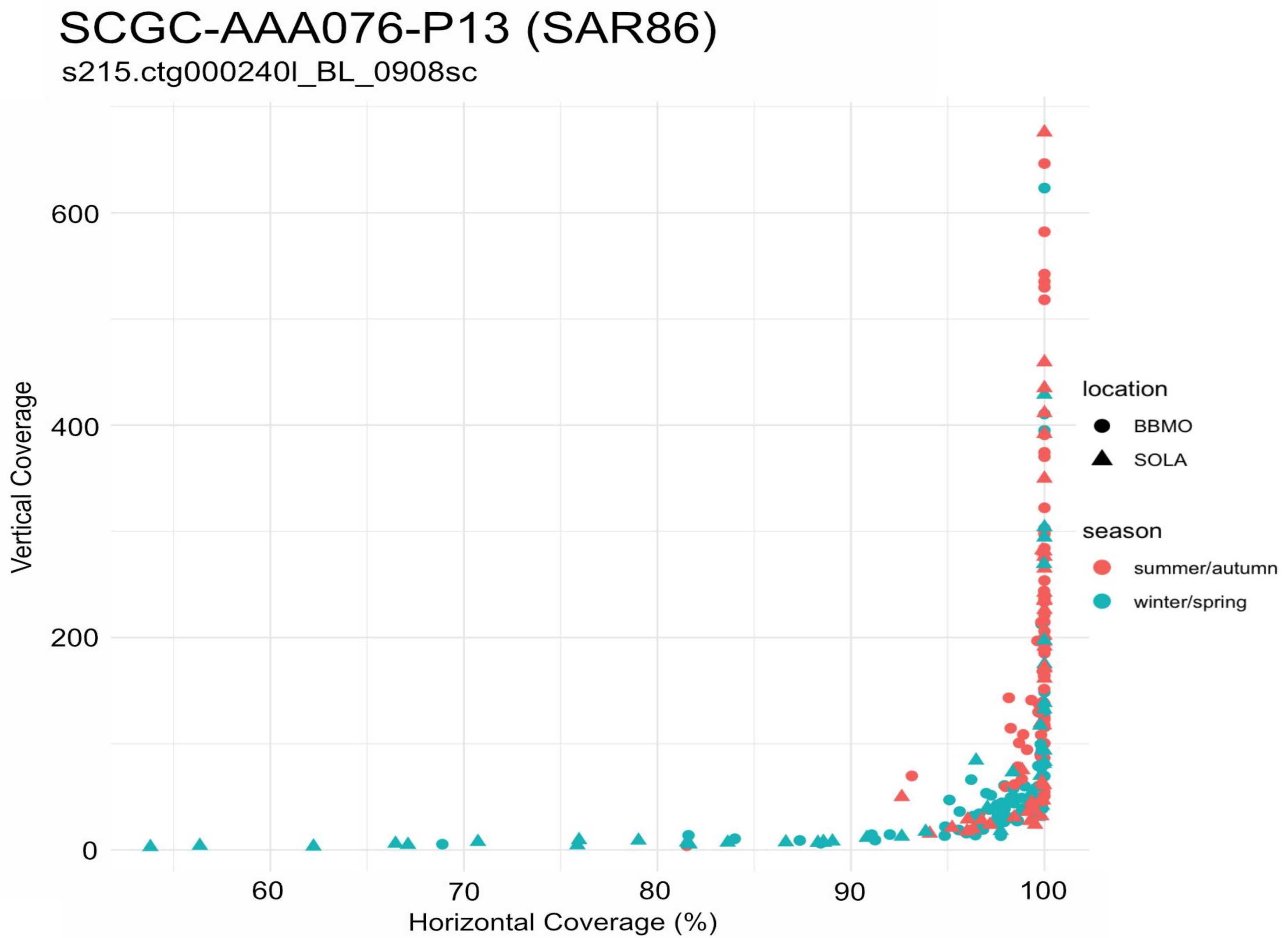

SCGC-AAA076-P13 (SAR86)
s215.ctg000240l_BL_0908sc
Vertical Coverage
600
400
200
0
60
70
80
90
100
Horizontal Coverage (%)
location
BBMO
SOLA
season
summer/autumn
winter/spring

### SCGC-AAA076-P13 (SAR86)

s388.ctg000435l_BL_0908sc

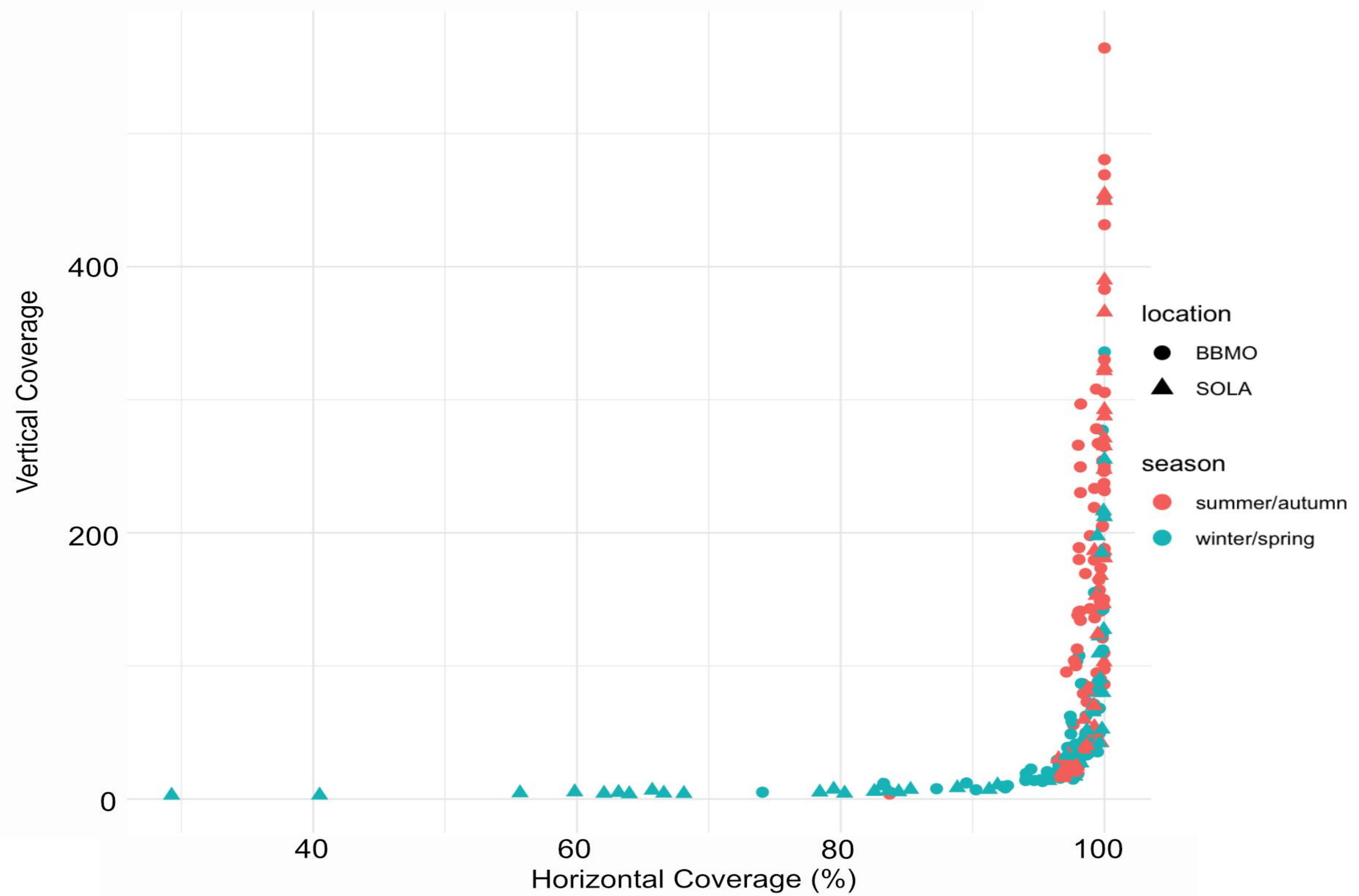

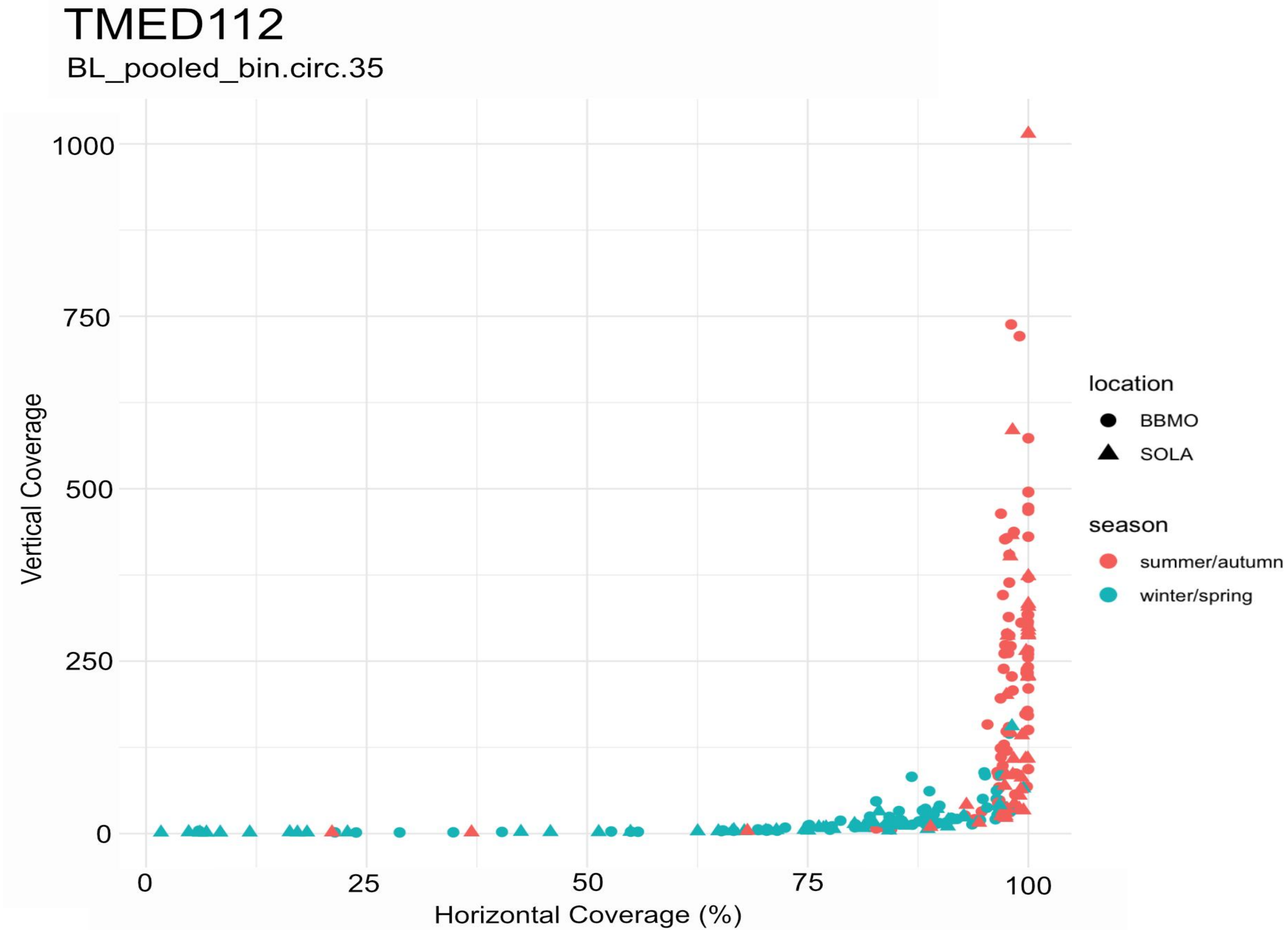

TMED112
BL_pooled_bin.circ.35
Vertical Coverage
1000
750
500
250
0
0
25
50
75
100
Horizontal Coverage (%)
location
BBMO
SOLA
season
summer/autumn
winter/spring

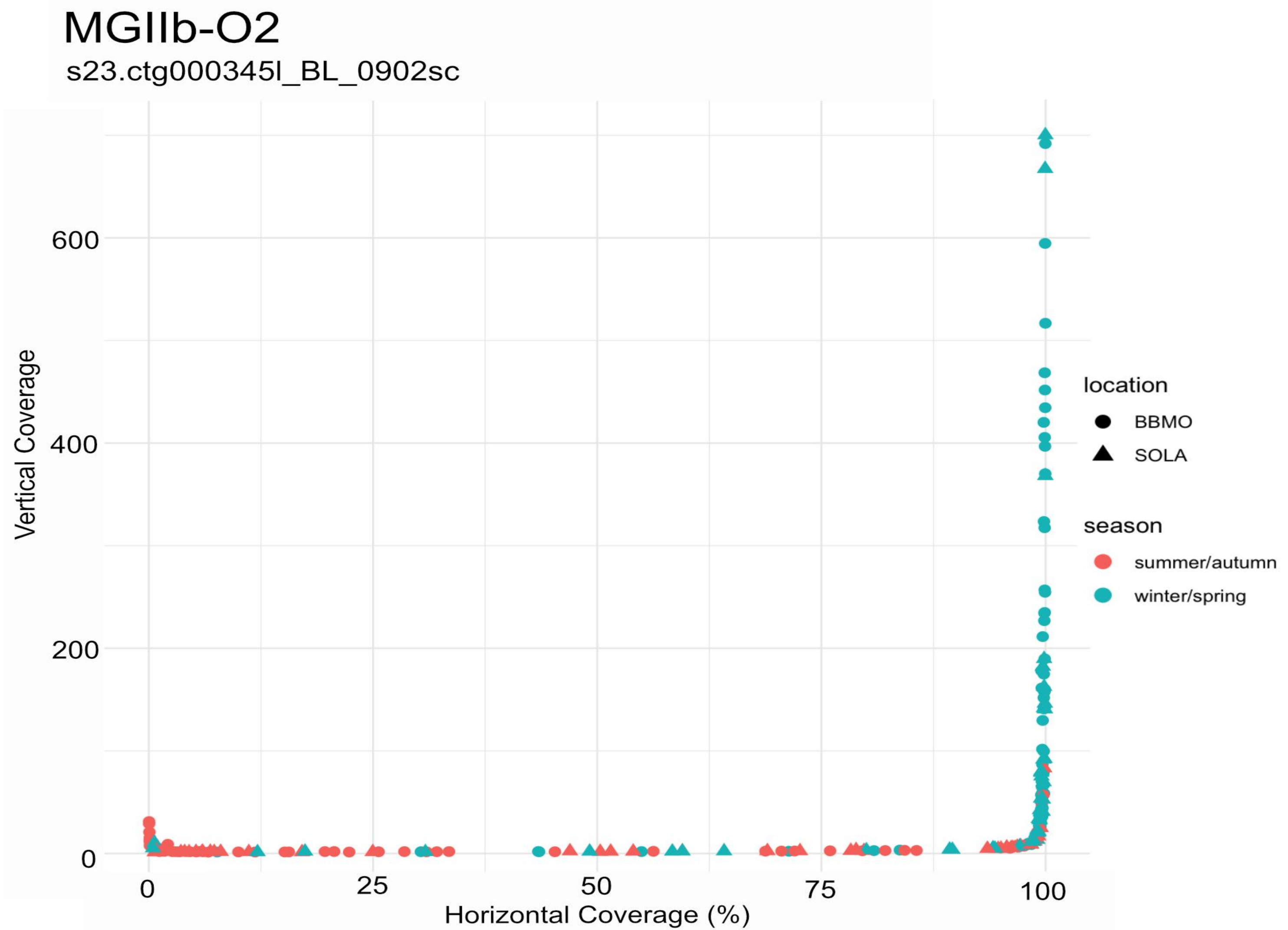

MGIIb-O2
s23.ctg000345l_BL_0902sc
Vertical Coverage
600
400
200
0
0
25
50
75
100
Horizontal Coverage (%)
location
BBMO
SOLA
season
summer/autumn
winter/spring

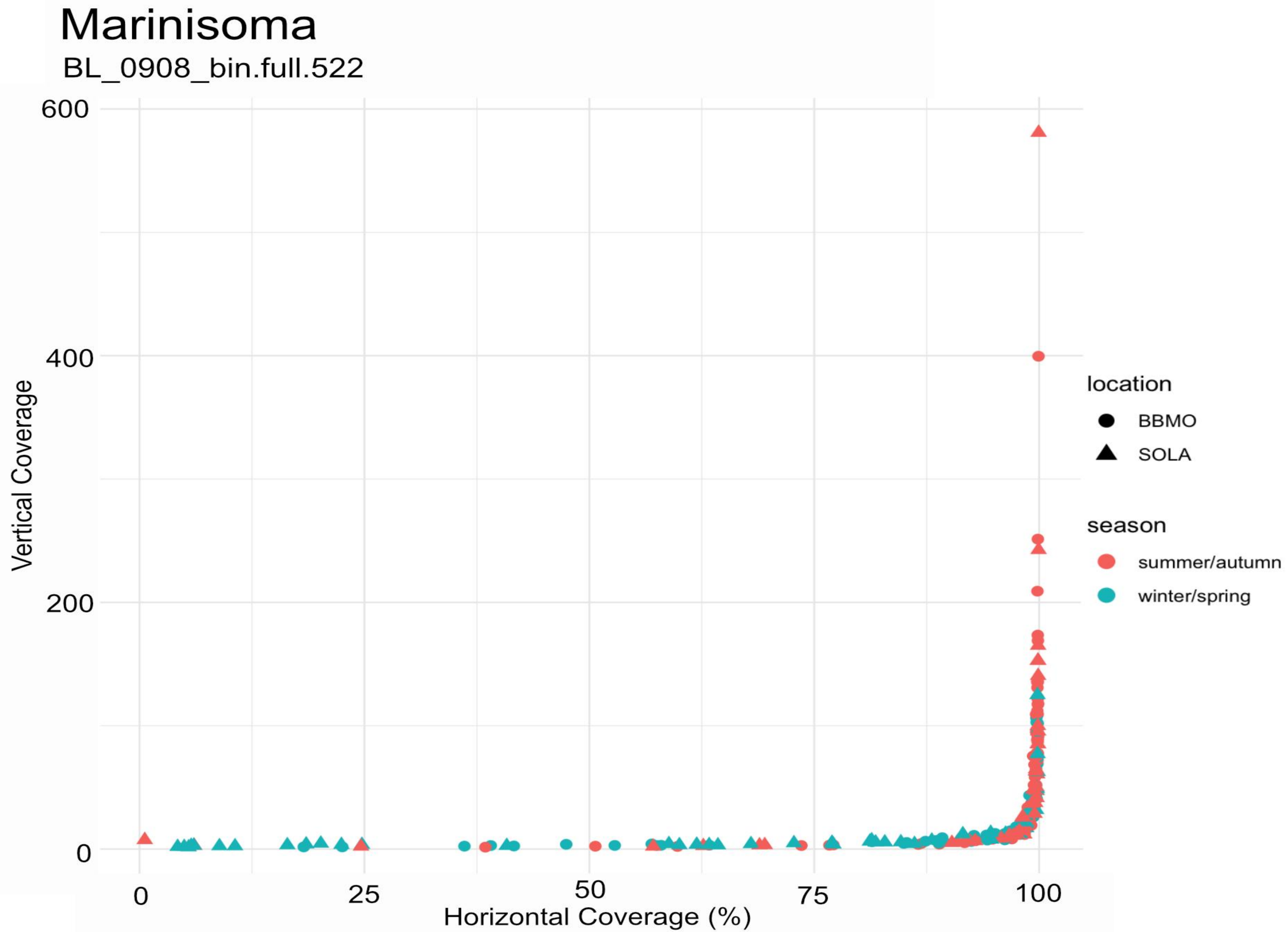
Marinisoma
BL_0908_bin.full.522
Vertical Coverage
Horizontal Coverage (%)
0
200
400
600
0
25
50
75
100
location
BBMO
SOLA
season
summer/autumn
winter/spring

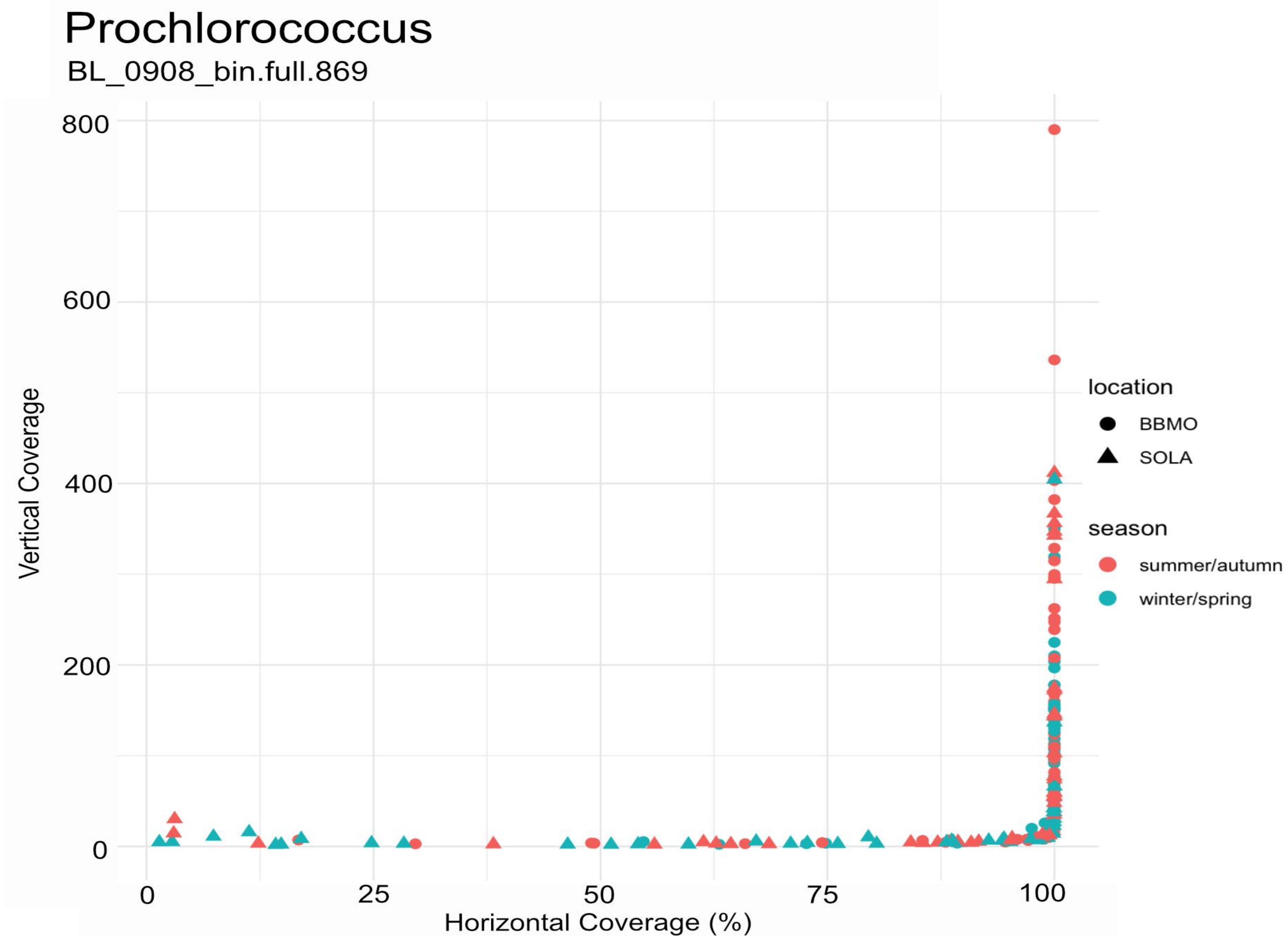

Prochlorococcus
BL_0908_bin.full.869
Vertical Coverage
800
600
400
200
0
0
25
50
75
100
Horizontal Coverage (%)
location
BBMO
SOLA
season
summer/autumn
winter/spring

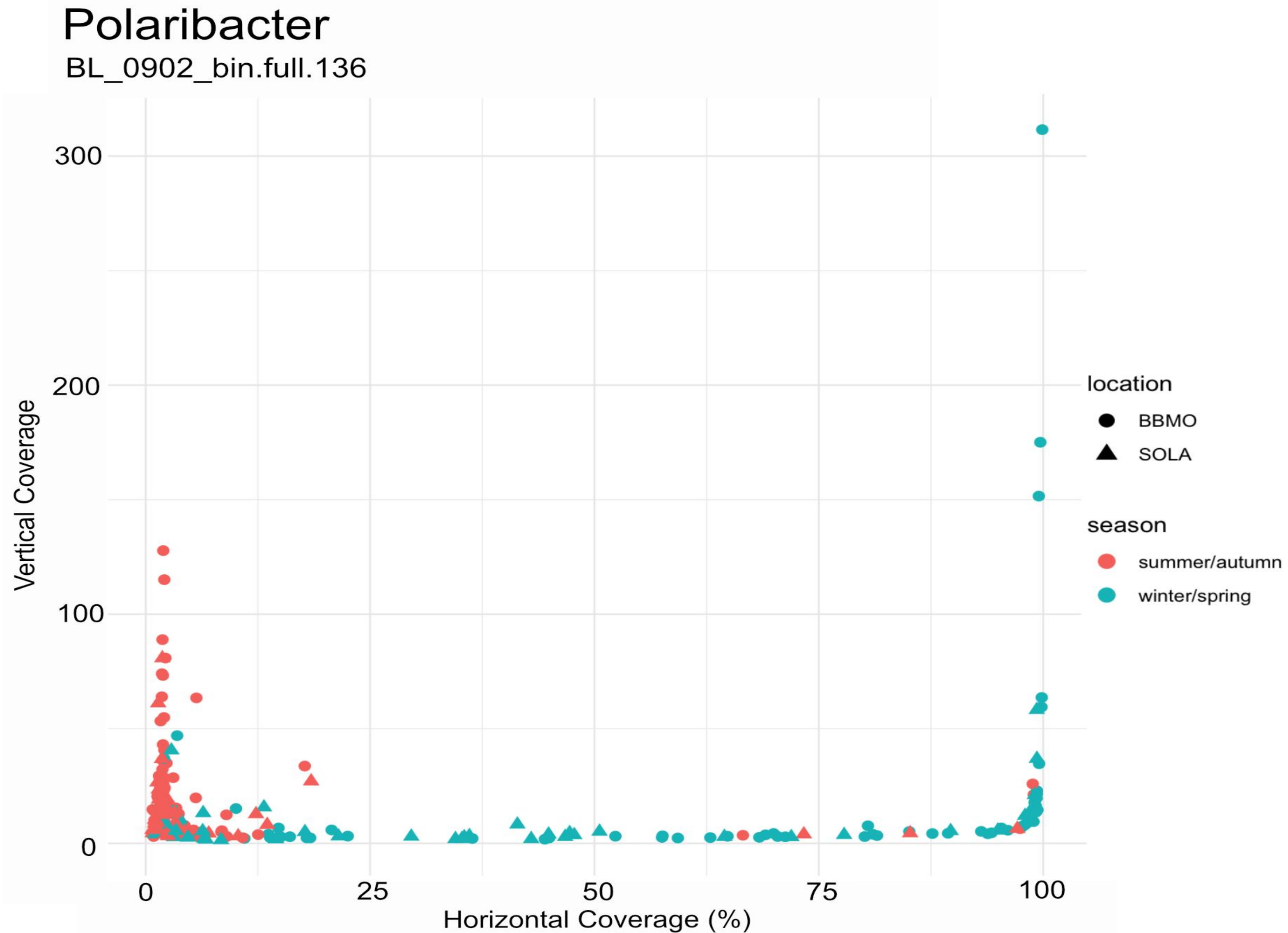

Polaribacter
BL_0902_bin.full.136
Vertical Coverage
Horizontal Coverage (%)
0
100
200
300
0
25
50
75
100
location
BBMO
SOLA
season
summer/autumn
winter/spring

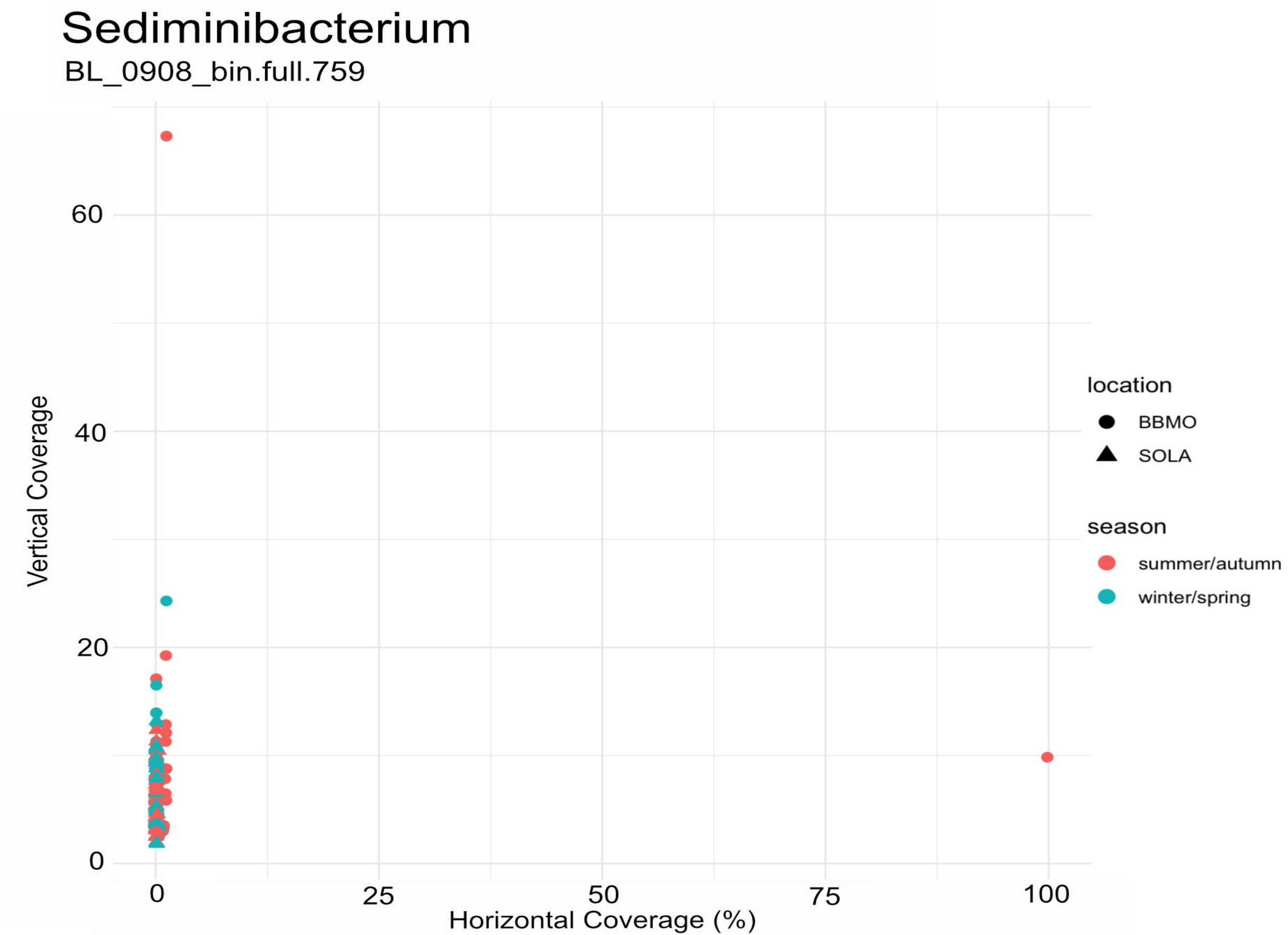

Sediminibacterium
BL_0908_bin.full.759
Vertical Coverage
0
20
40
60
0
25
50
75
100
Horizontal Coverage (%)
location
BBMO
SOLA
season
summer/autumn
winter/spring

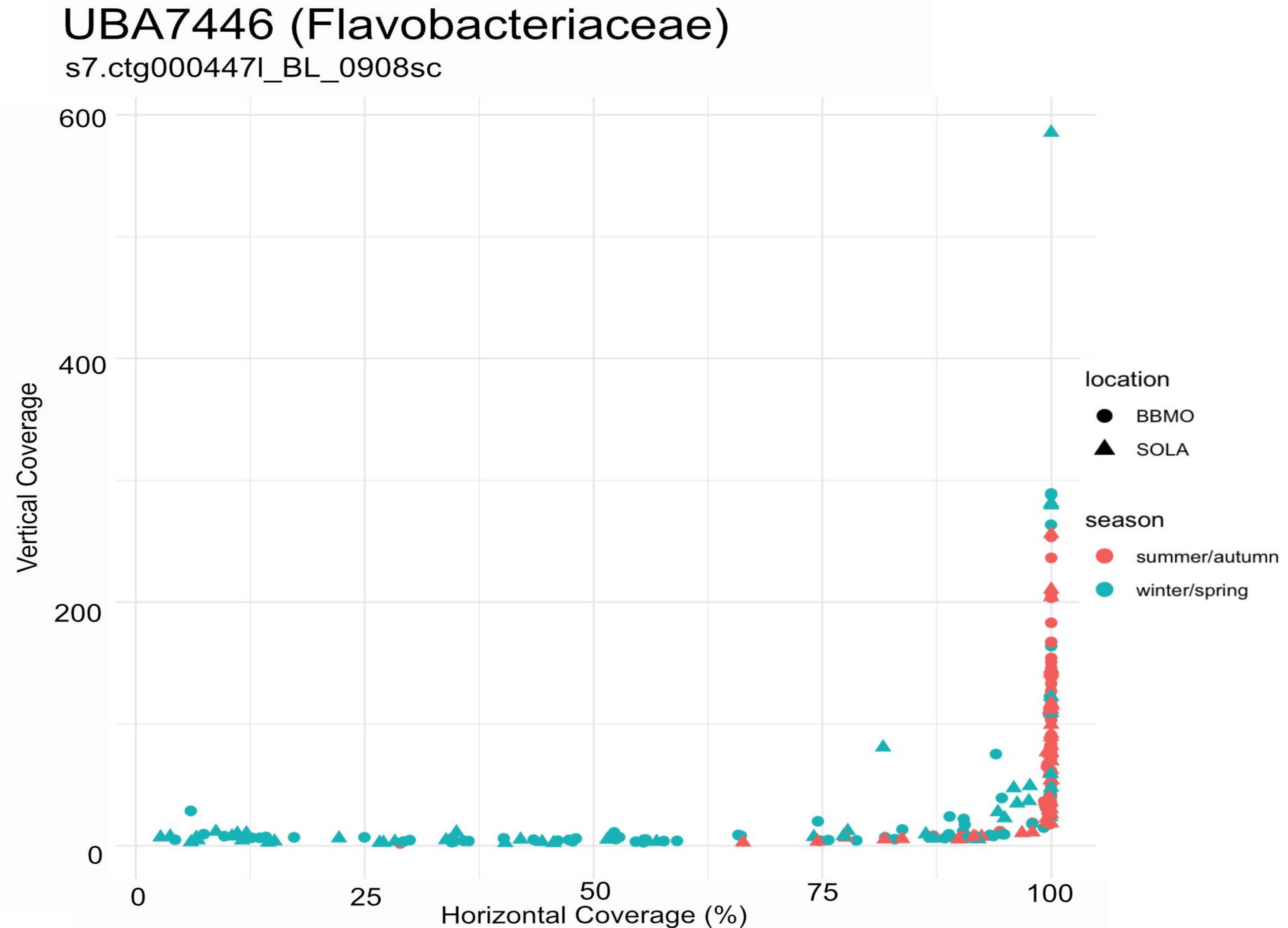
UBA7446 (Flavobacteriaceae)
s7.ctg000447l_BL_0908sc
600
400
200
0
Vertical Coverage
0
25
50
75
100
Horizontal Coverage (%)
location
BBMO
SOLA
season
summer/autumn
winter/spring

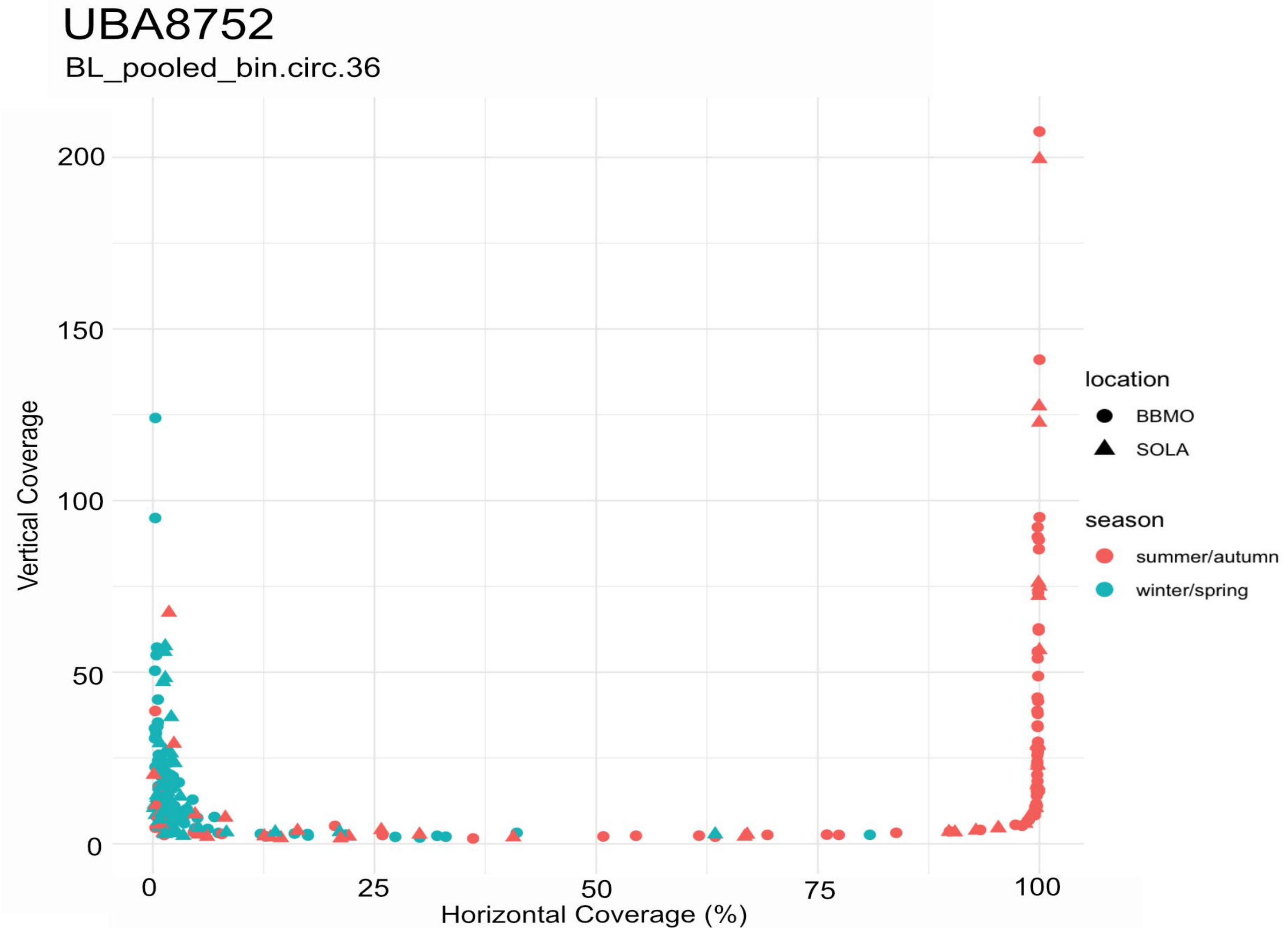
UBA8752
BL_pooled_bin.circ.36
Vertical Coverage
Horizontal Coverage (%)
0
50
100
150
200
0
25
50
75
100
location
BBMO
SOLA
season
summer/autumn
winter/spring

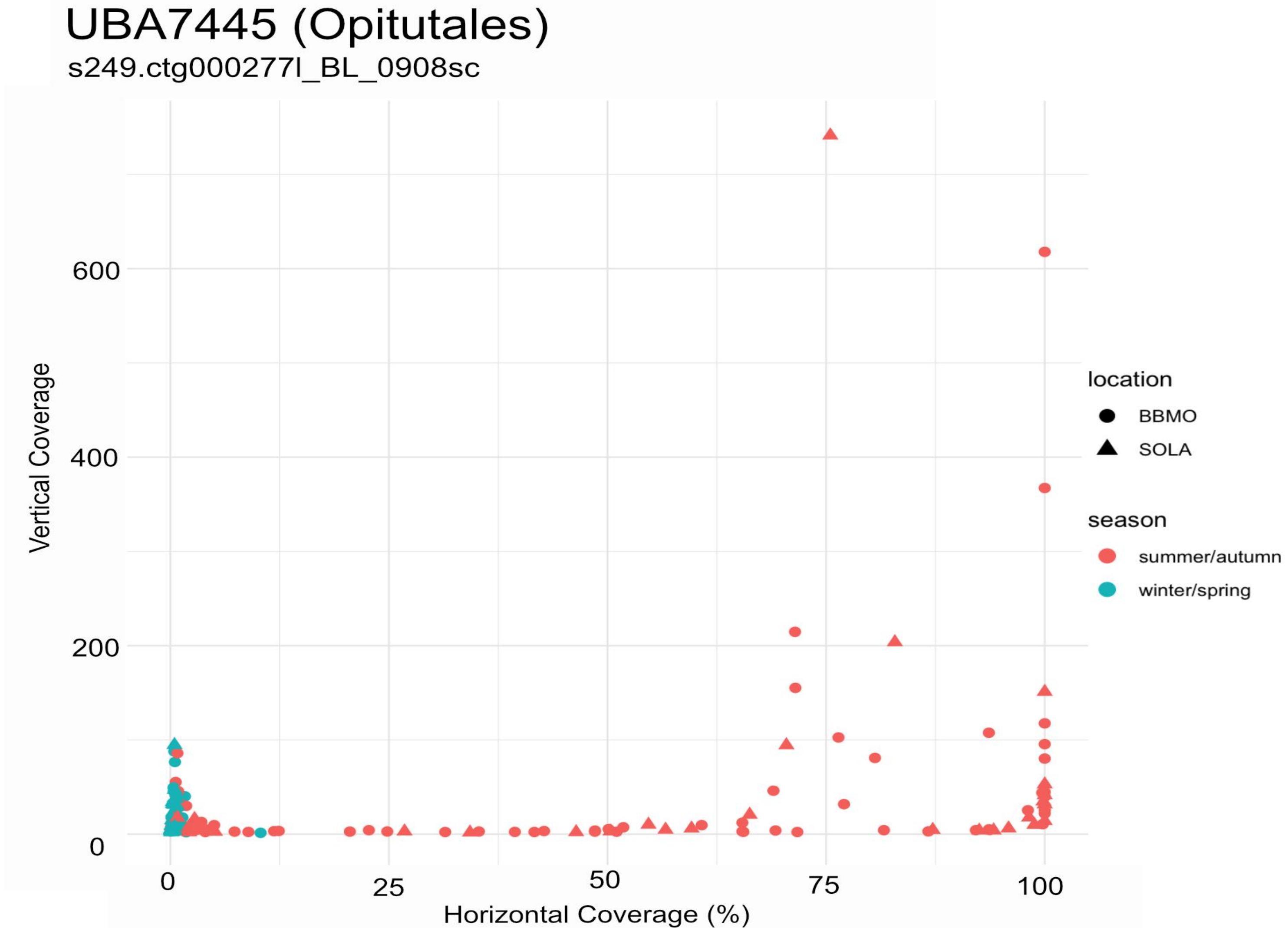
UBA7445 (Opitutales)
s249.ctg000277l_BL_0908sc
Vertical Coverage
600
400
200
0
0
25
50
75
100
Horizontal Coverage (%)
location
BBMO
SOLA
season
summer/autumn
winter/spring

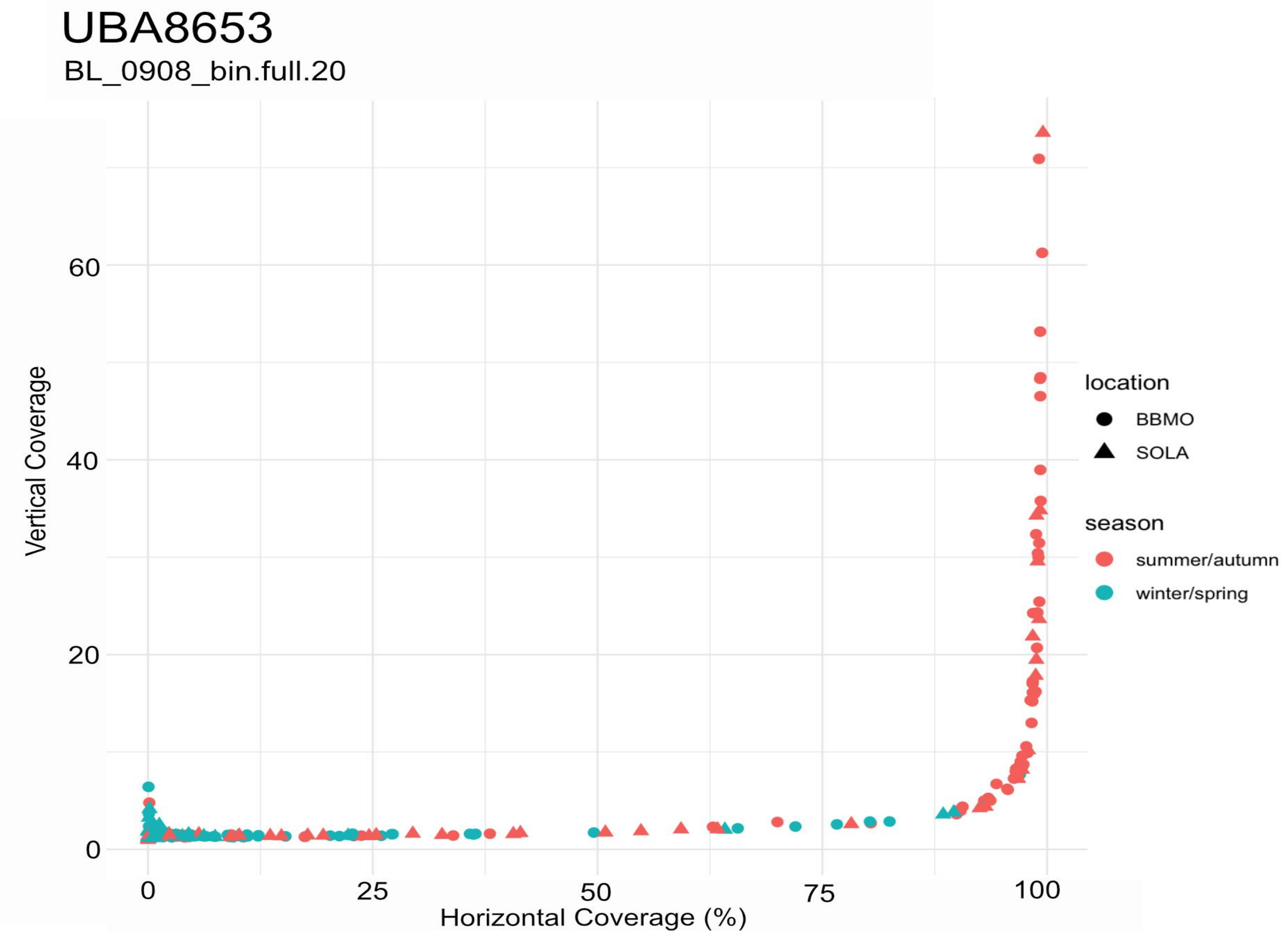
UBA8653
BL_0908_bin.full.20
Vertical Coverage
60
40
20
0
25
50
75
100
Horizontal Coverage (%)
location
BBMO
SOLA
season
summer/autumn
winter/spring

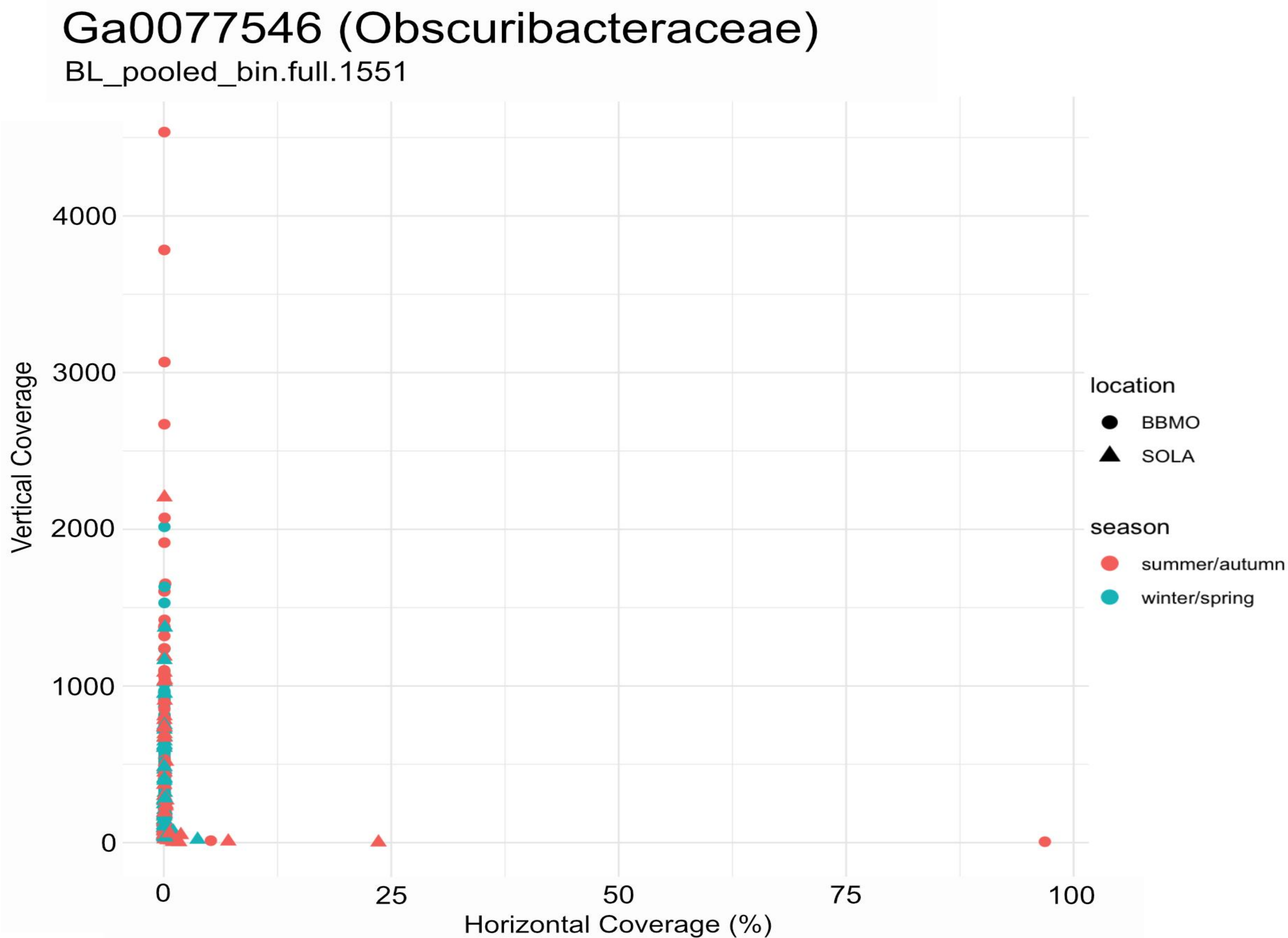
Ga0077546 (Obscuribacteraceae)
BL_pooled_bin.full.1551
Vertical Coverage
4000
3000
2000
1000
0
0
25
50
75
100
Horizontal Coverage (%)
location
BBMO
SOLA
season
summer/autumn
winter/spring
