## Supplementary Figure 9 for "Seasonal population structure and adaptive signatures across long-term marine microbial time series"

**Supplementary Figure 9. Population structure, candidate adaptive-gene pN/pS, and abundance patterns across BBMO and SOLA.** For each MAG, the upper panel shows a UPGMA dendrogram based on pairwise genome-wide Fst among samples meeting the horizontal-coverage criterion described in the Methods. The dashed horizontal line indicates the Fst = 0.15 threshold used to delineate candidate populations, and branch colors distinguish the resulting sample clusters. Bars below the dendrogram indicate the seasonal state of each sample (Winter–Spring, cold waters; Summer–Autumn, warm waters) and sampling location (BBMO or SOLA). The middle panel shows pN/pS values for candidate adaptive genes across the same samples; values >2 were capped at 2 for visualization. White cells indicate zero or unavailable pN/pS values. The lower panel shows MAG abundance (RPKG) across the corresponding samples. Samples with RPKG < 1 were not considered when interpreting population structure. Samples are shown in the order defined by the Fst dendrogram and therefore are not arranged chronologically. MAGs are presented in the same order as in Table 1. Note that *Amylibacter* (s35.ctg000041c_BL_0902sc) is not included, as only a few BBMO samples (metagenomes) pass the threshold of horizontal coverage > 25%, while no sample from SOLA passes this threshold.

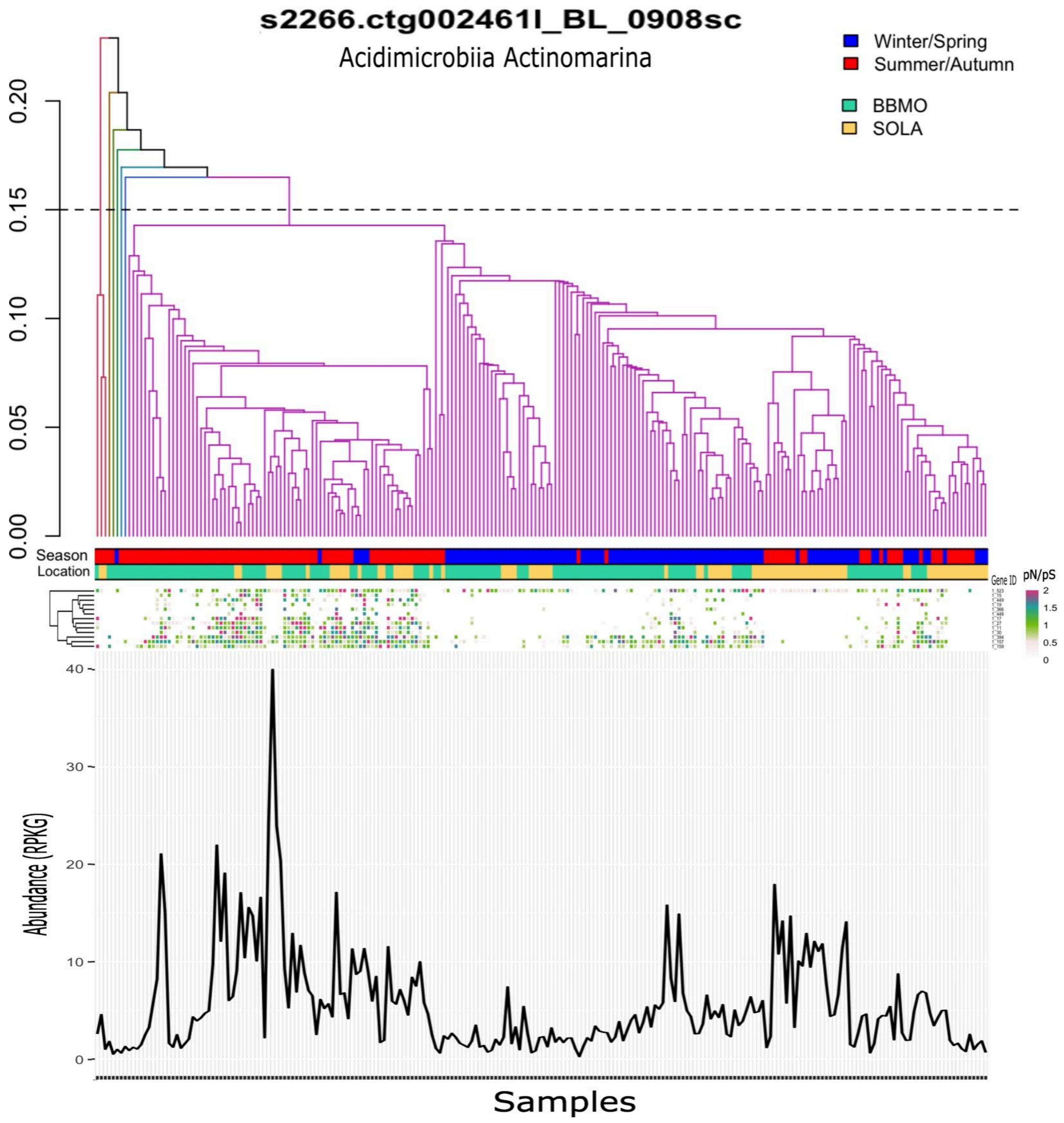
s2266.ctg002461l_BL_0908sc
Acidimicrobiia Actinomarina
Winter/Spring
Summer/Autumn
BBMO
SOLA
0.20
0.15
0.10
0.05
0.00
Season
Location
Gene ID
pN/pS
2
1.5
1
0.5
0
40
30
20
10
0
Abundance (RPKG)
Samples

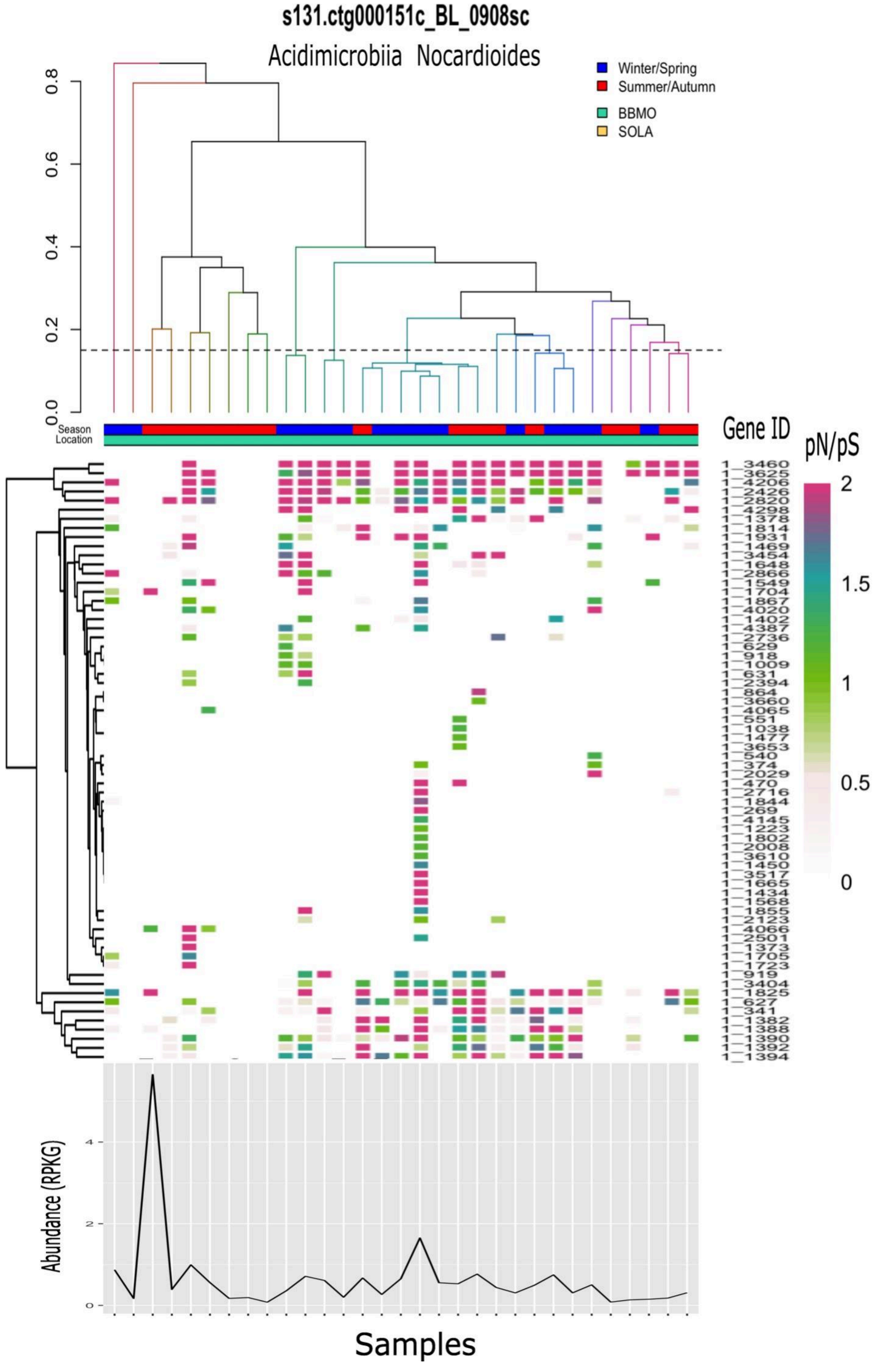

s131.ctg000151c_BL_0908sc
Acidimicrobiia Nocardioides
Winter/Spring
Summer/Autumn
BBMO
SOLA
0.0
0.2
0.4
0.6
0.8
Season
Location
Gene ID
pN/pS
2
1.5
1
0.5
0
1_3460
1_3625
1_4206
1_2426
1_2820
1_4298
1_1378
1_1814
1_1931
1_1469
1_3454
1_1648
1_2866
1_1549
1_1704
1_1867
1_4020
1_1402
1_4387
1_2736
1_629
1_918
1_1009
1_631
1_2394
1_864
1_3660
1_4065
1_551
1_1038
1_1477
1_3653
1_540
1_374
1_2029
1_470
1_2716
1_1844
1_269
1_4145
1_1223
1_1802
1_2008
1_3610
1_1450
1_3517
1_1665
1_1434
1_1568
1_1855
1_2123
1_4066
1_2501
1_1373
1_1705
1_1723
1_919
1_3404
1_1825
1_627
1_341
1_1382
1_1388
1_1390
1_1392
1_1394
Abundance (RPKG)
0
2
4
Samples

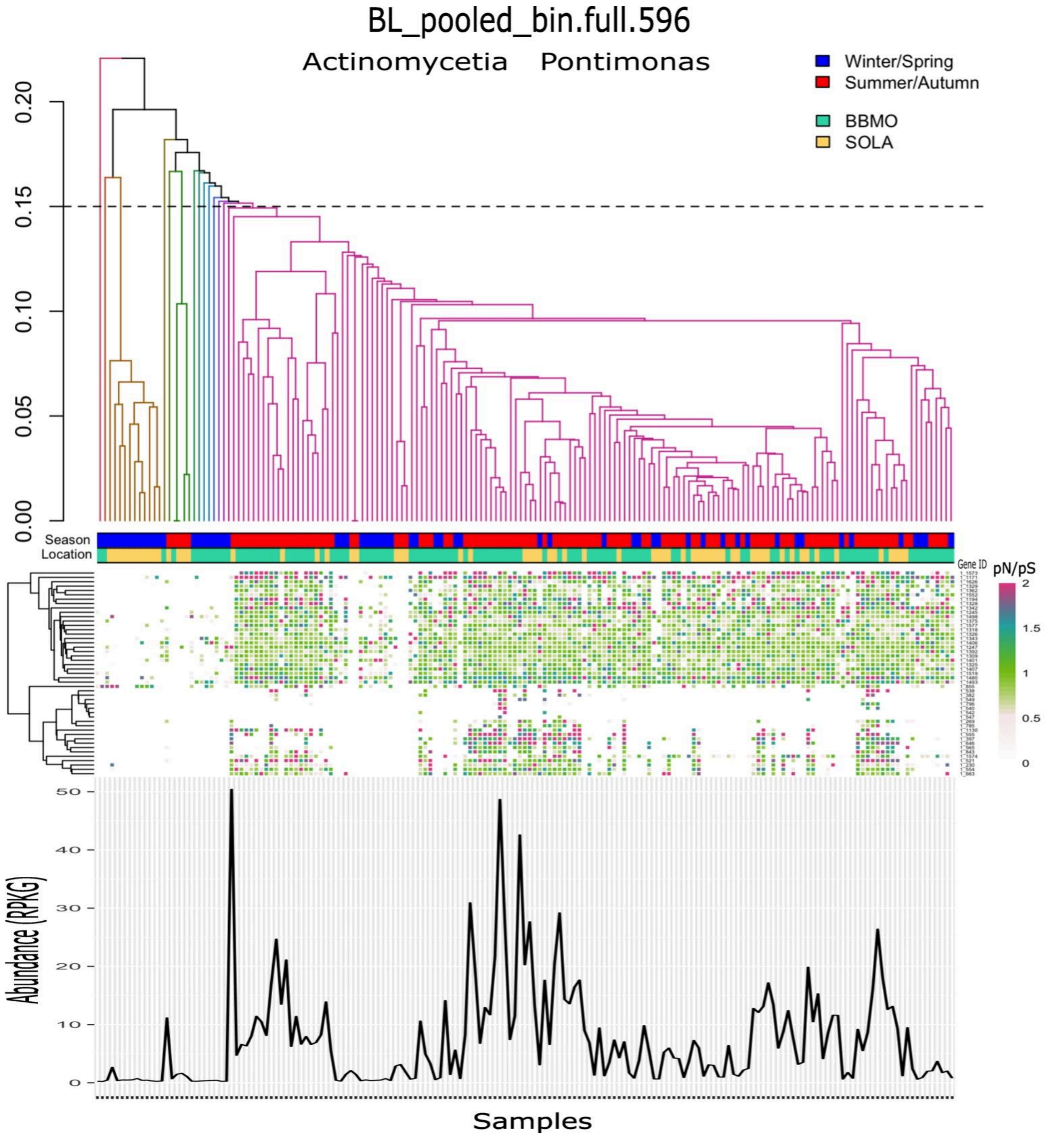

BL_pooled_bin.full.596
Actinomycetia Pontimonas
Winter/Spring
Summer/Autumn
BBMO
SOLA
0.20
0.15
0.10
0.05
0.00
Season
Location
Gene ID
pN/pS
2
1.5
1
0.5
0
50
40
30
20
10
0
Abundance (RPKG)
Samples

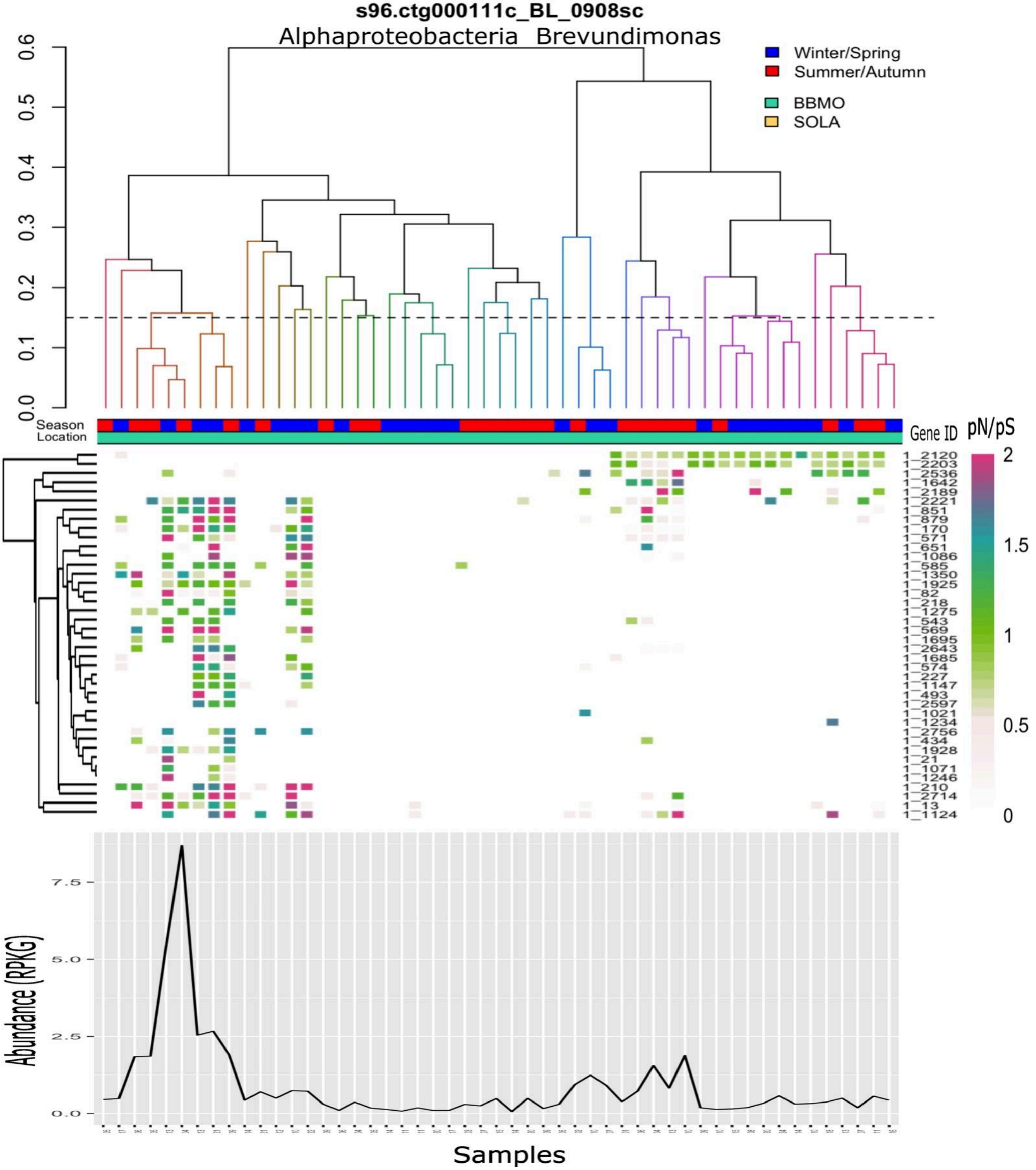
s96.ctg000111c_BL_0908sc
Alphaproteobacteria Brevundimonas
Winter/Spring
Summer/Autumn
BBMO
SOLA
0.0
0.1
0.2
0.3
0.4
0.5
0.6
Season
Location
Gene ID
pN/pS
2
1.5
1
0.5
0
1_2120
1_2203
1_2536
1_1642
1_2189
1_2221
1_851
1_879
1_170
1_571
1_651
1_1086
1_585
1_1350
1_1925
1_82
1_218
1_1275
1_543
1_569
1_1695
1_2643
1_1685
1_574
1_227
1_1147
1_493
1_2597
1_1021
1_1234
1_2756
1_434
1_1928
1_21
1_1071
1_1246
1_210
1_2714
1_13
1_1124
Abundance (RPKG)
0.0
2.5
5.0
7.5
Samples

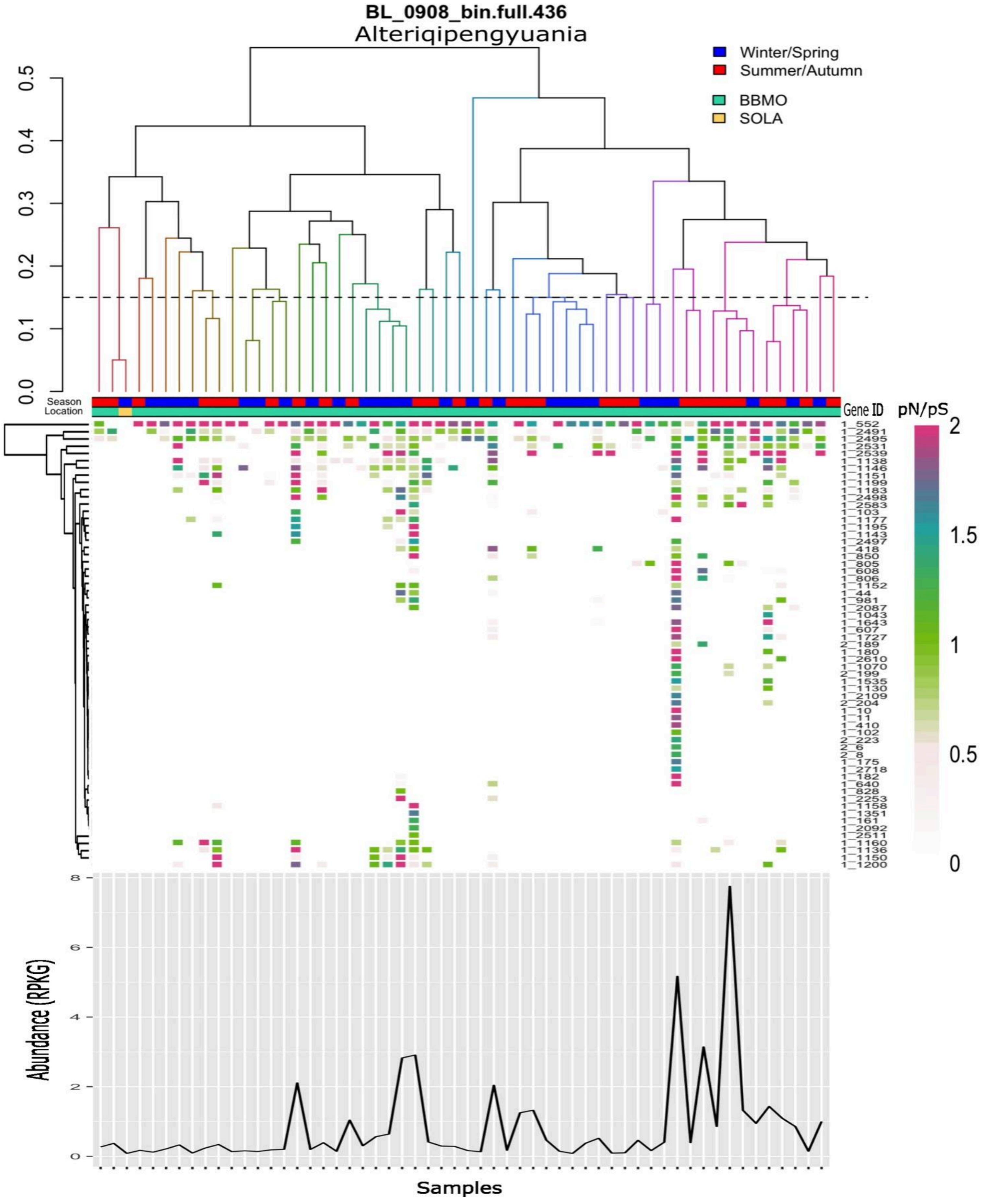
BL_0908_bin.full.436
Alteriqipengyuania
Winter/Spring
Summer/Autumn
BBMO
SOLA
0.0
0.1
0.2
0.3
0.4
0.5
Season
Location
Gene ID
pN/pS
2
1.5
1
0.5
0
1_552
1_2491
1_2495
1_2531
1_2539
1_1138
1_1146
1_1151
1_1199
1_1183
1_2498
1_2583
1_103
1_1177
1_1195
1_1143
1_2497
1_418
1_850
1_805
1_608
1_806
1_1152
1_44
1_981
1_2087
1_1043
1_1643
1_607
1_1727
2_189
1_180
1_2610
1_1070
2_199
1_1535
1_1130
1_2109
2_204
1_10
1_11
1_410
1_102
2_223
2_6
2_8
1_175
1_2718
1_182
1_640
1_828
1_2253
1_1158
1_1351
1_161
1_2092
1_2511
1_1160
1_1136
1_1150
1_1200
Abundance (RPKG)
0
2
4
6
8
Samples

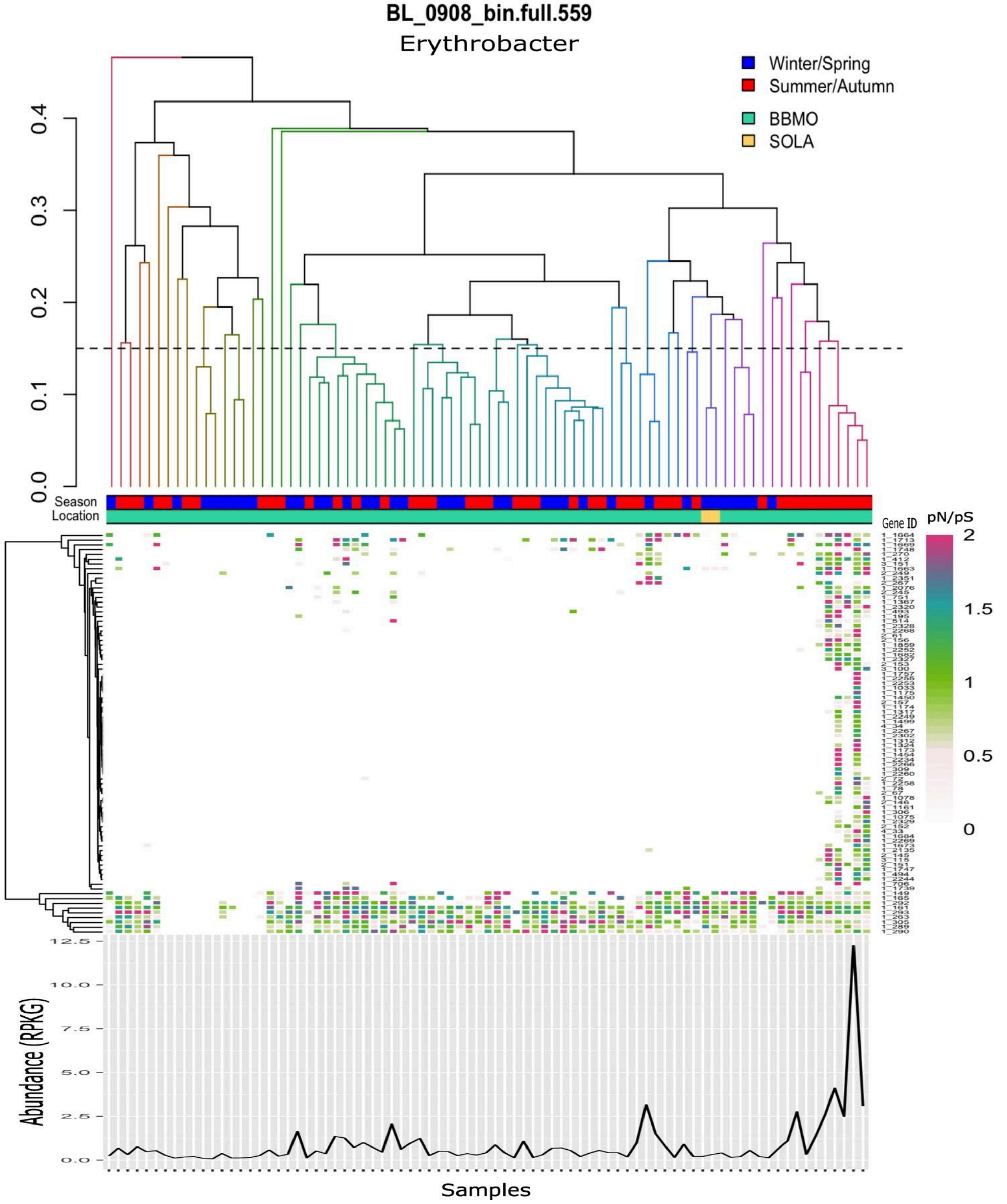

BL_0908_bin.full.559
Erythrobacter
Winter/Spring
Summer/Autumn
BBMO
SOLA
0.0
0.1
0.2
0.3
0.4
Season
Location
Gene ID
pN/pS
2
1.5
1
0.5
0
Abundance (RPKG)
12.5
10.0
7.5
5.0
2.5
0.0
Samples

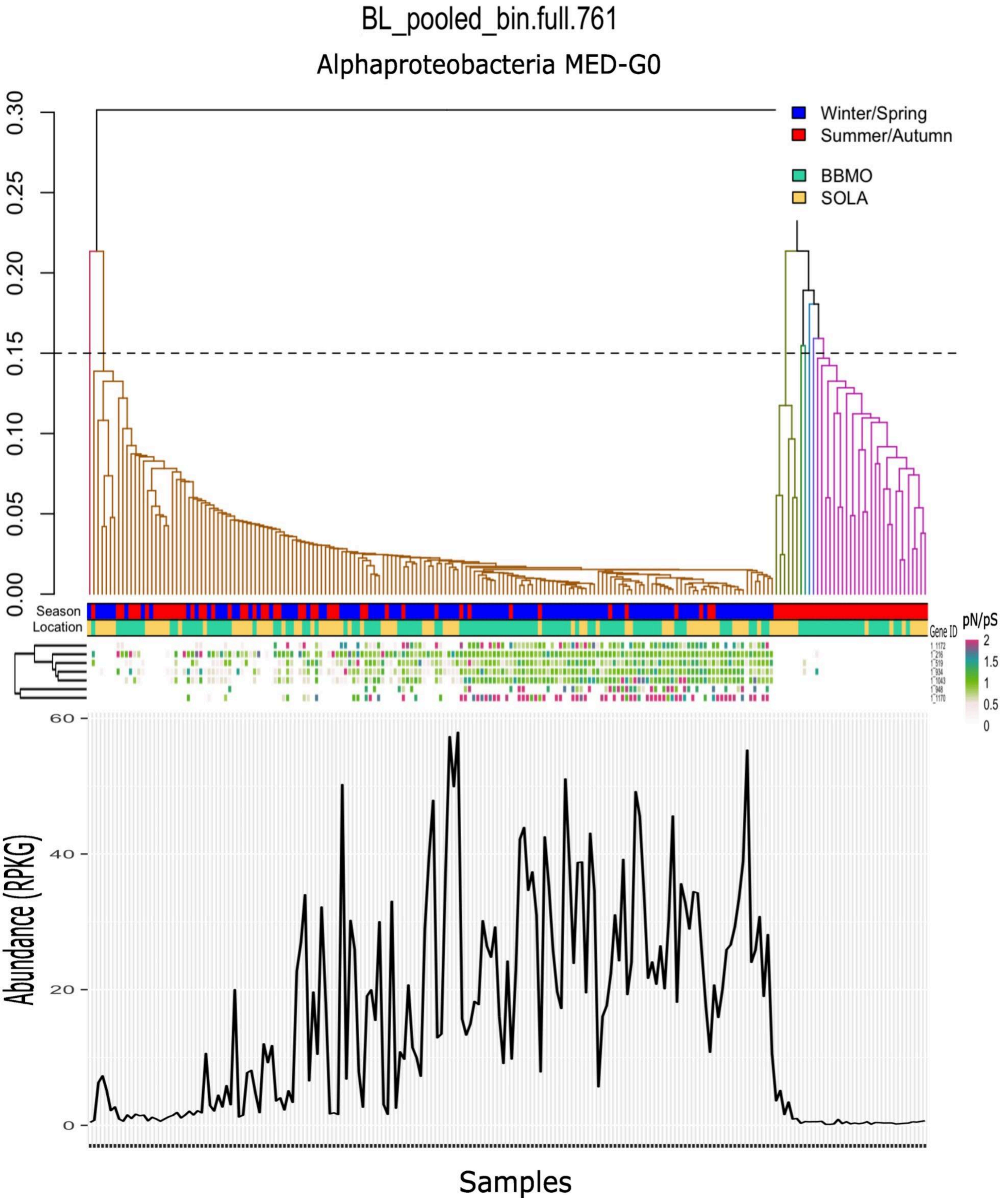

BL_pooled_bin.full.761
Alphaproteobacteria MED-G0
Winter/Spring
Summer/Autumn
BBMO
SOLA
Season
Location
Gene ID
pN/pS
2
1.5
1
0.5
0
0.00
0.05
0.10
0.15
0.20
0.25
0.30
60
40
20
0
Abundance (RPKG)
Samples

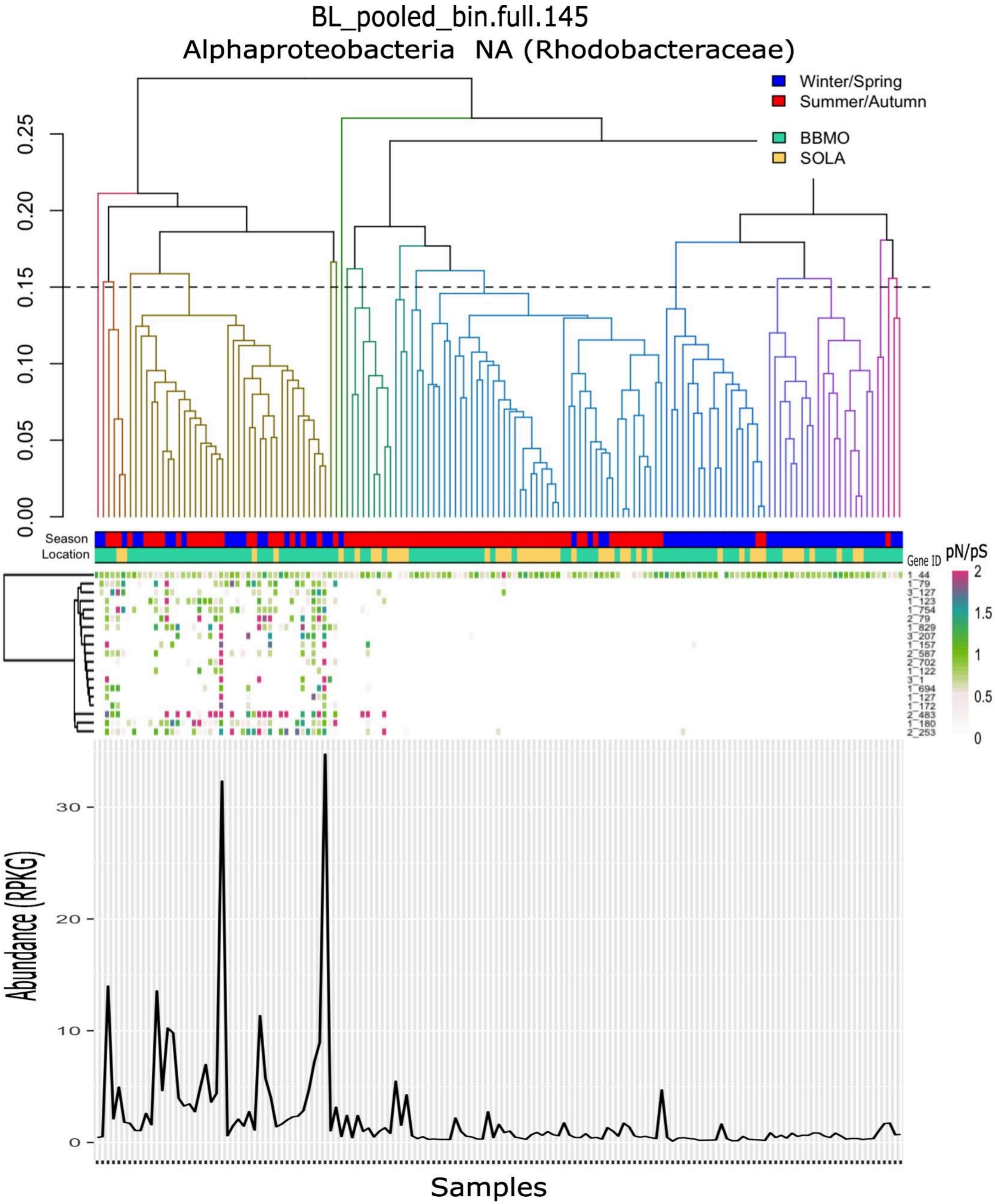

BL_pooled_bin.full.145
Alphaproteobacteria NA (Rhodobacteraceae)
Winter/Spring
Summer/Autumn
BBMO
SOLA
0.00
0.05
0.10
0.15
0.20
0.25
Season
Location
Gene ID
pN/pS
2
1.5
1
0.5
0
1_44
1_79
3_127
1_123
1_754
2_79
1_829
3_207
1_157
2_587
2_702
1_122
3_1
1_694
1_127
1_172
2_483
1_180
2_253
Abundance (RPKG)
0
10
20
30
Samples

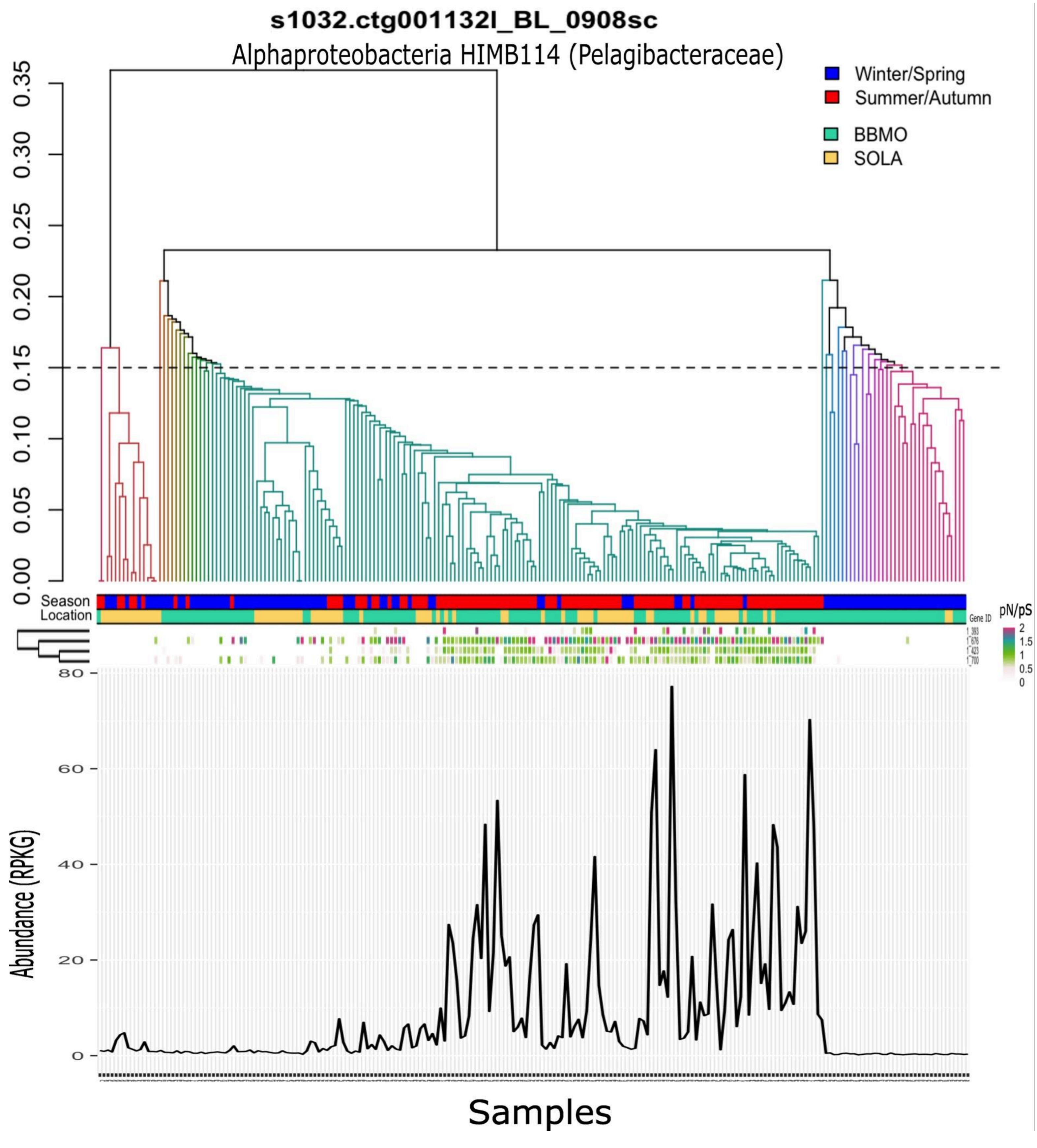

s1032.ctg001132l_BL_0908sc
Alphaproteobacteria HIMB114 (Pelagibacteraceae)
Winter/Spring
Summer/Autumn
BBMO
SOLA
0.00
0.05
0.10
0.15
0.20
0.25
0.30
0.35
Season
Location
Gene ID
pN/pS
2
1.5
1
0.5
0
Abundance (RPKG)
0
20
40
60
80
Samples

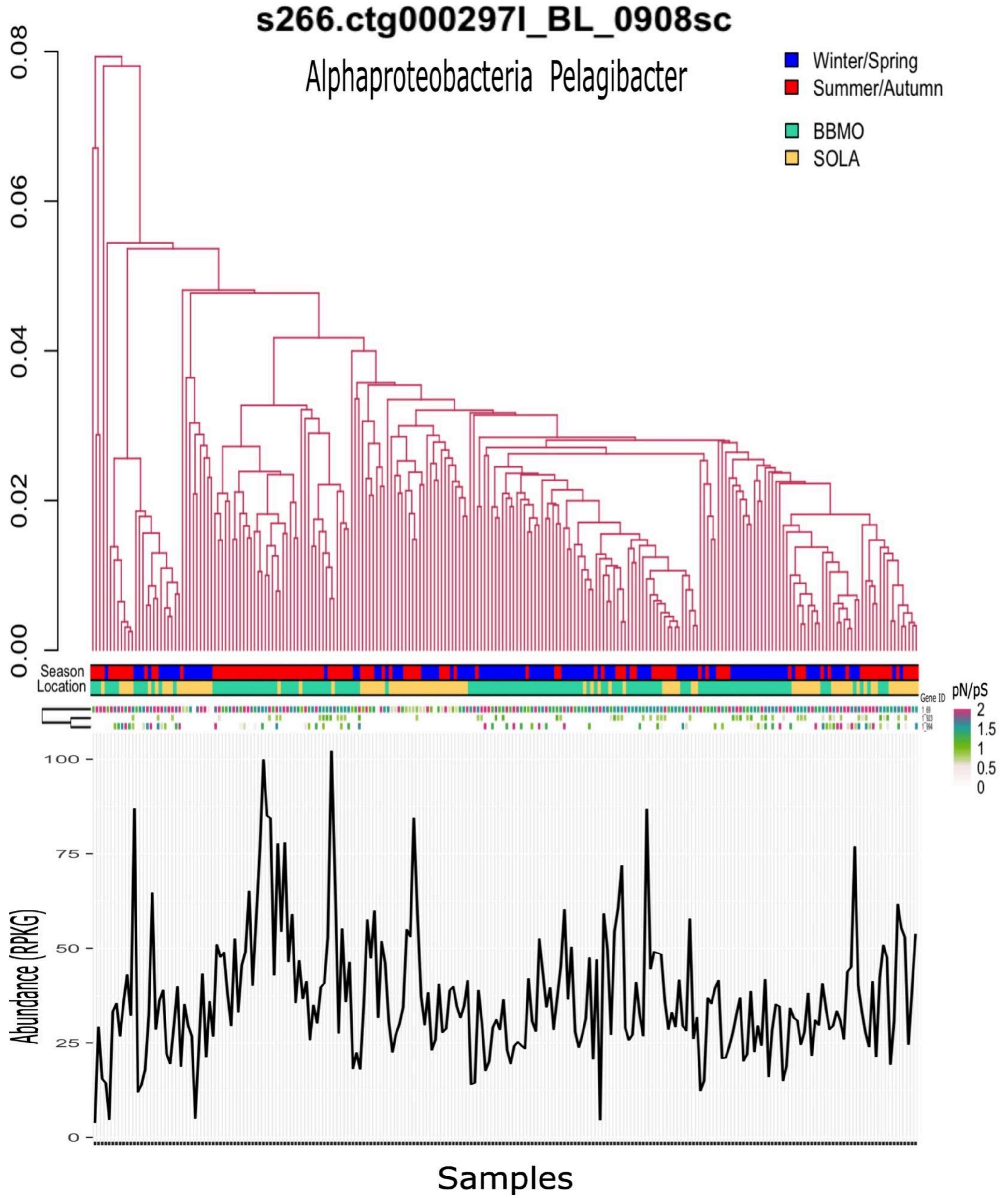

s266.ctg000297l_BL_0908sc
Alphaproteobacteria Pelagibacter
Winter/Spring
Summer/Autumn
BBMO
SOLA
0.08
0.06
0.04
0.02
0.00
Season
Location
Gene ID
pN/pS
2
1.5
1
0.5
0
100
75
50
25
0
Abundance (RPKG)
Samples

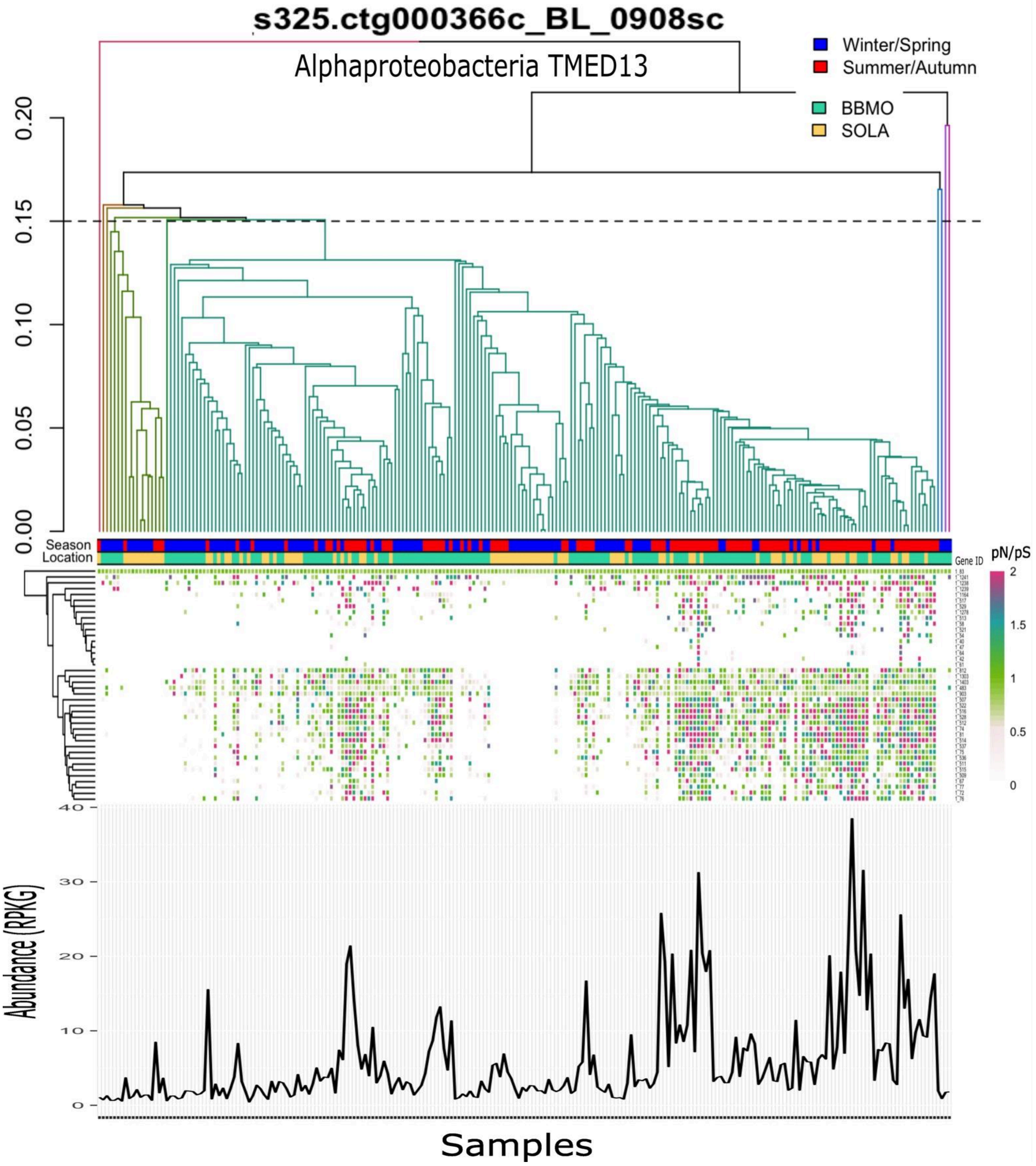
s325.ctg000366c_BL_0908sc
Alphaproteobacteria TMED13
Winter/Spring
Summer/Autumn
BBMO
SOLA
0.00
0.05
0.10
0.15
0.20
Season
Location
Gene ID
pN/pS
2
1.5
1
0.5
0
Abundance (RPKG)
40
30
20
10
0
Samples

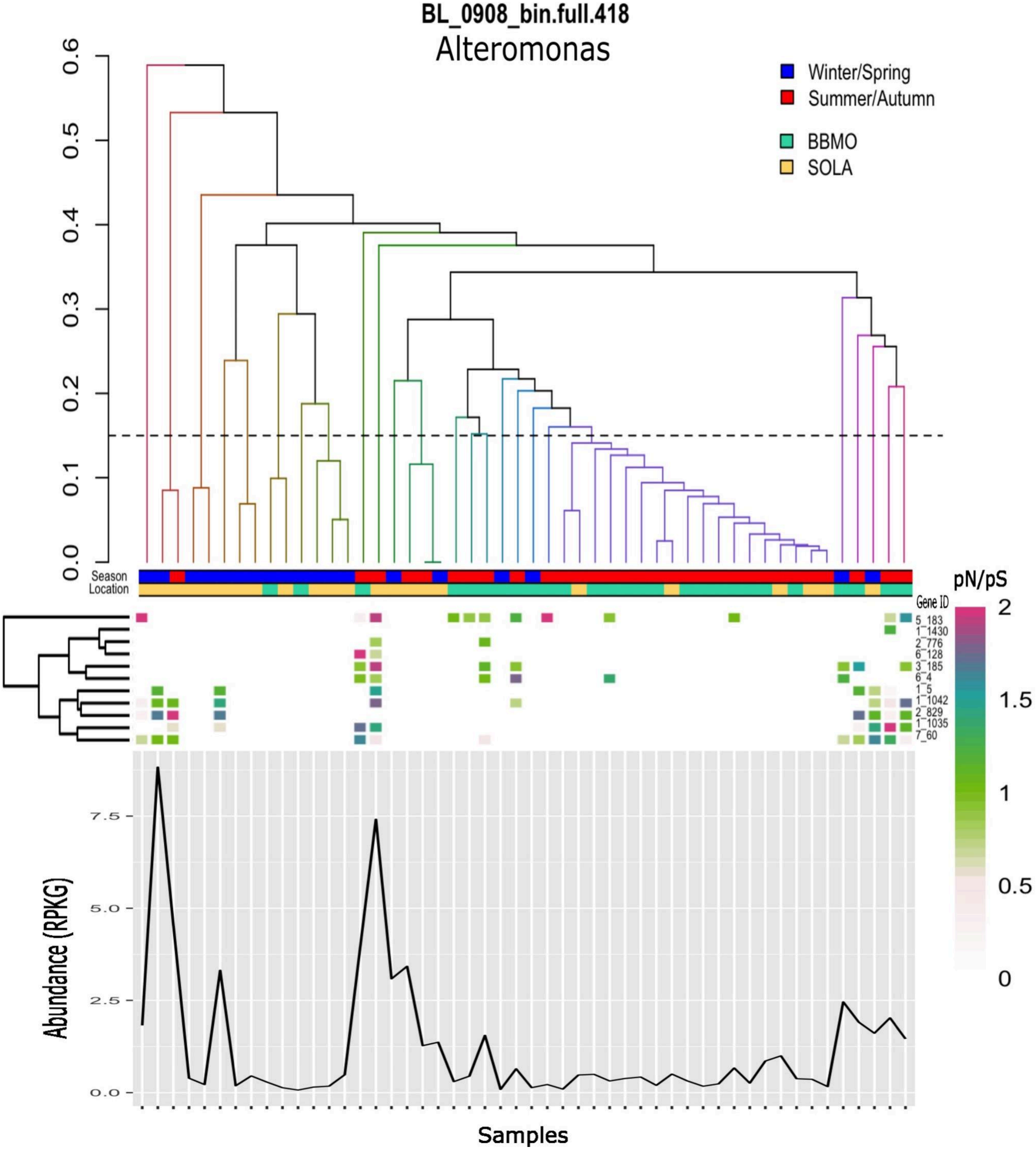
BL_0908_bin.full.418
Alteromonas
Winter/Spring
Summer/Autumn
BBMO
SOLA
0.0
0.1
0.2
0.3
0.4
0.5
0.6
Season
Location
pN/pS
Gene ID
5_183
1_1430
2_776
6_128
3_185
6_4
1_5
1_1042
2_829
1_1035
7_60
2
1.5
1
0.5
0
Abundance (RPKG)
0.0
2.5
5.0
7.5
Samples

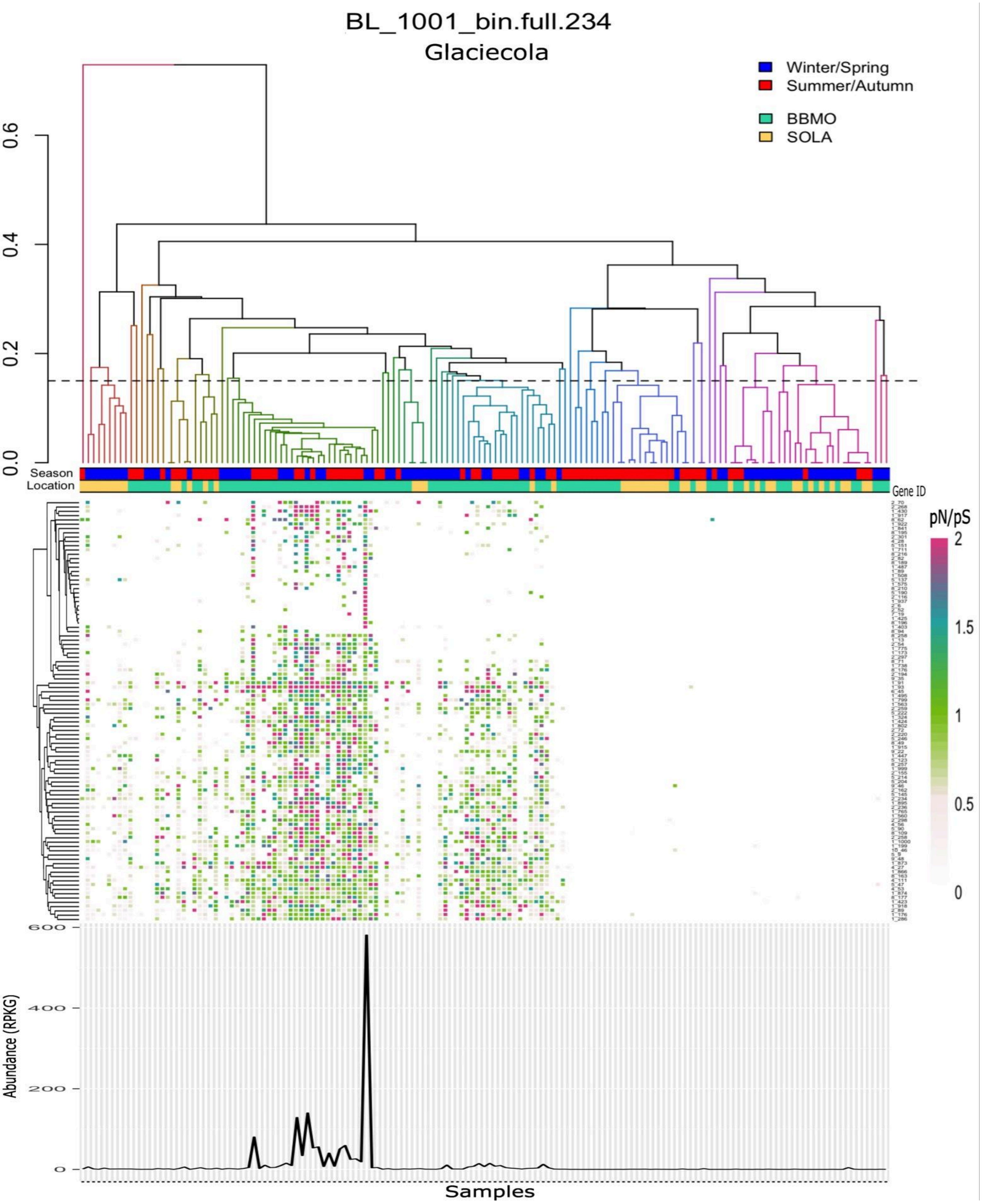
BL_1001_bin.full.234
Glaciecola
Winter/Spring
Summer/Autumn
BBMO
SOLA
0.6
0.4
0.2
0.0
Season
Location
Gene ID
pN/pS
2
1.5
1
0.5
0
600
400
200
0
Abundance (RPKG)
Samples

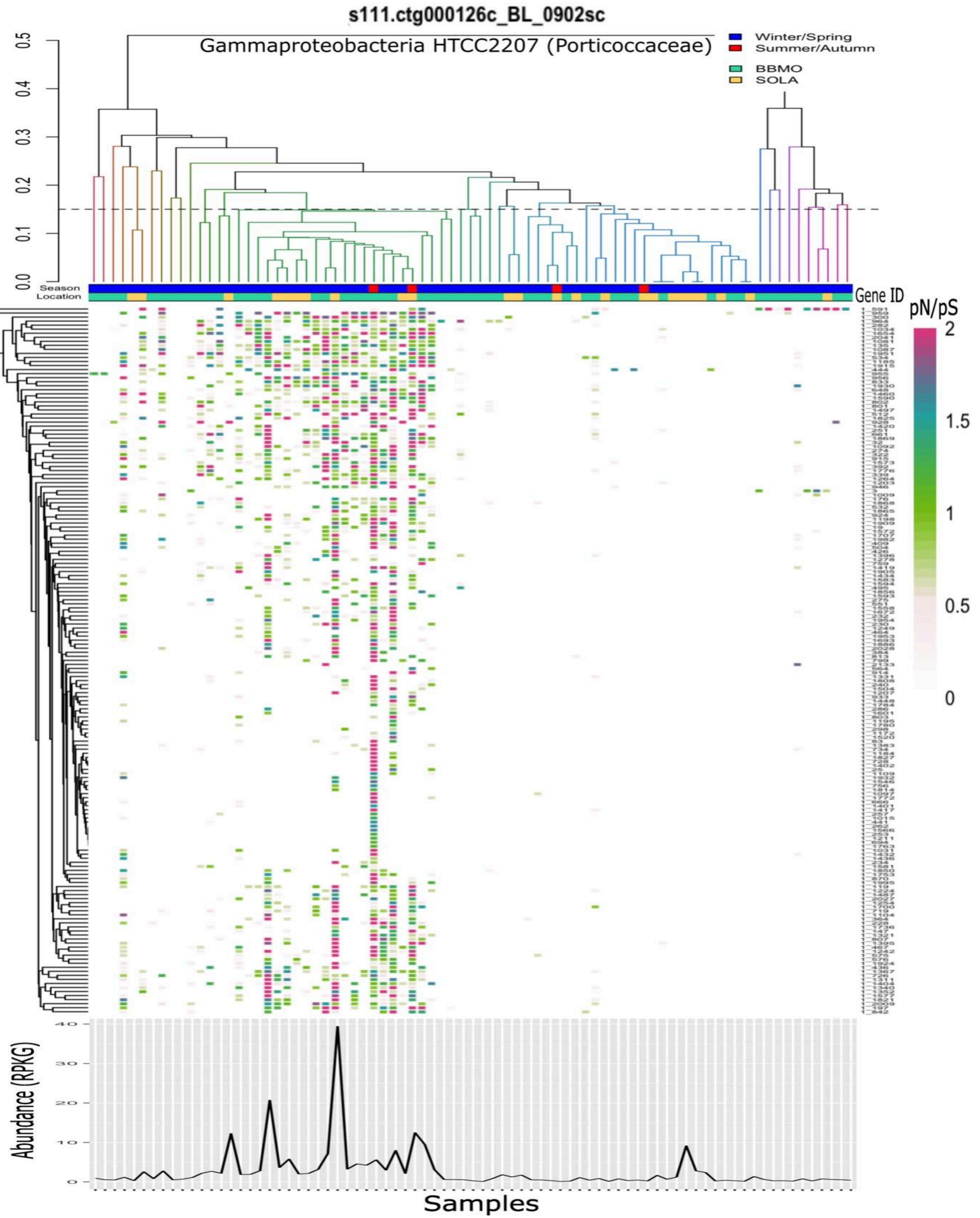
s111.ctg000126c_BL_0902sc
Gammaproteobacteria HTCC2207 (Porticoccaceae)
Winter/Spring
Summer/Autumn
BBMO
SOLA
Season
Location
Gene ID
pN/pS
Abundance (RPKG)
Samples

**s44.ctg000052l_BL_0902sc**

Gammaproteobacteria HTCC2207 (Porticoccaceae)

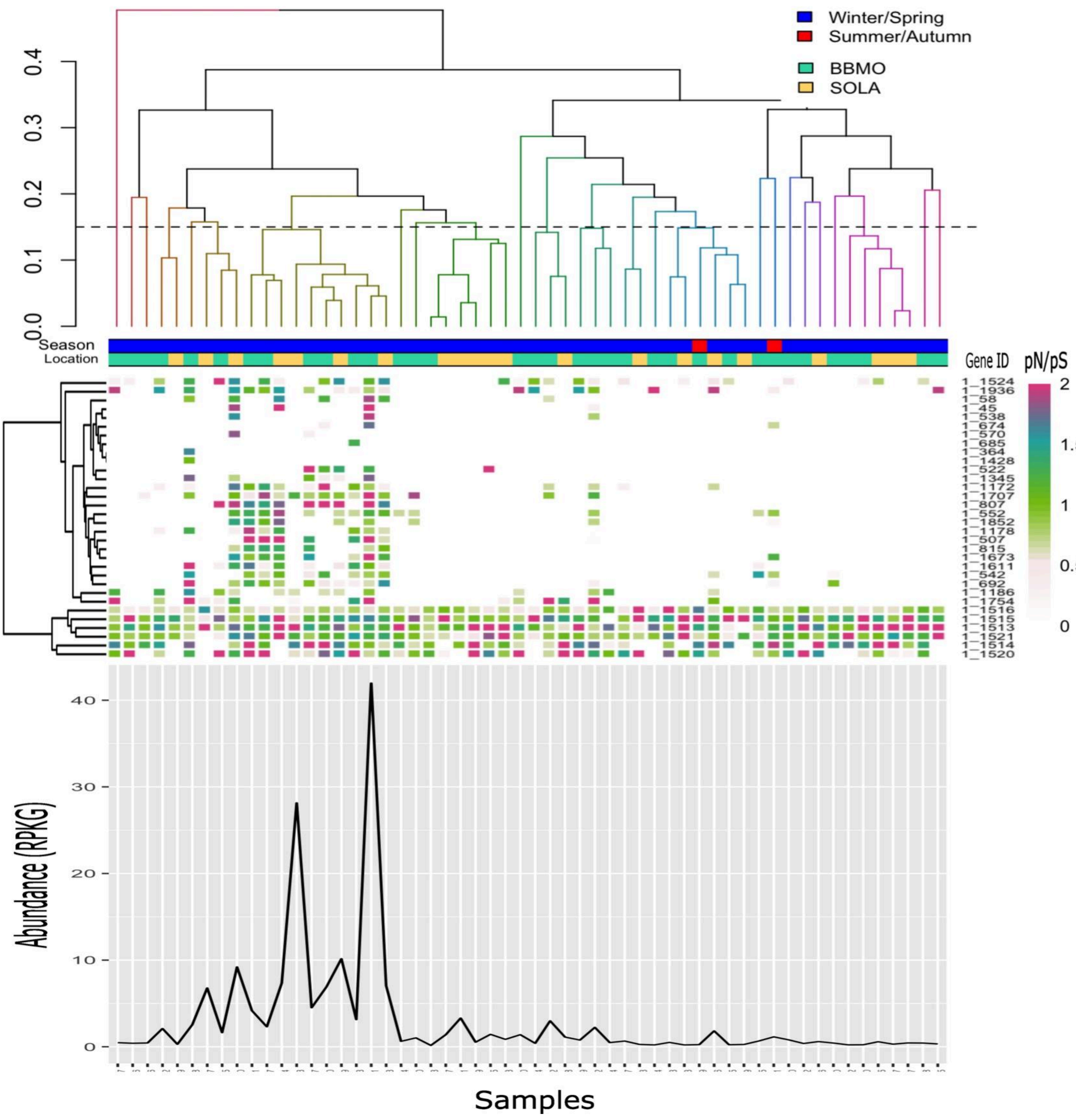

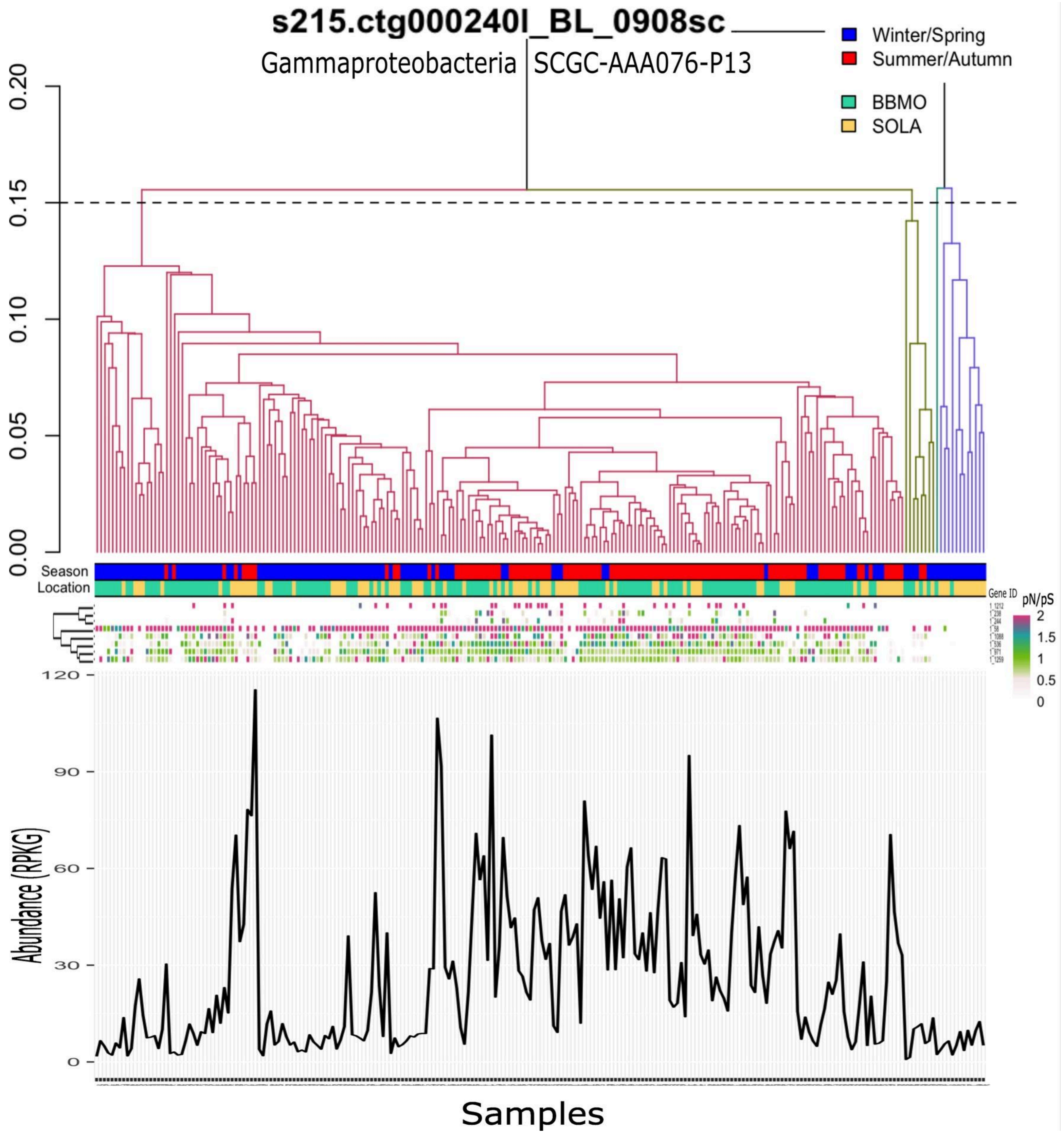

s215.ctg000240l_BL_0908sc
Gammaproteobacteria SCGC-AAA076-P13
Winter/Spring
Summer/Autumn
BBMO
SOLA
0.20
0.15
0.10
0.05
0.00
Season
Location
Gene ID
pN/pS
2
1.5
1
0.5
0
120
90
60
30
0
Abundance (RPKG)
Samples

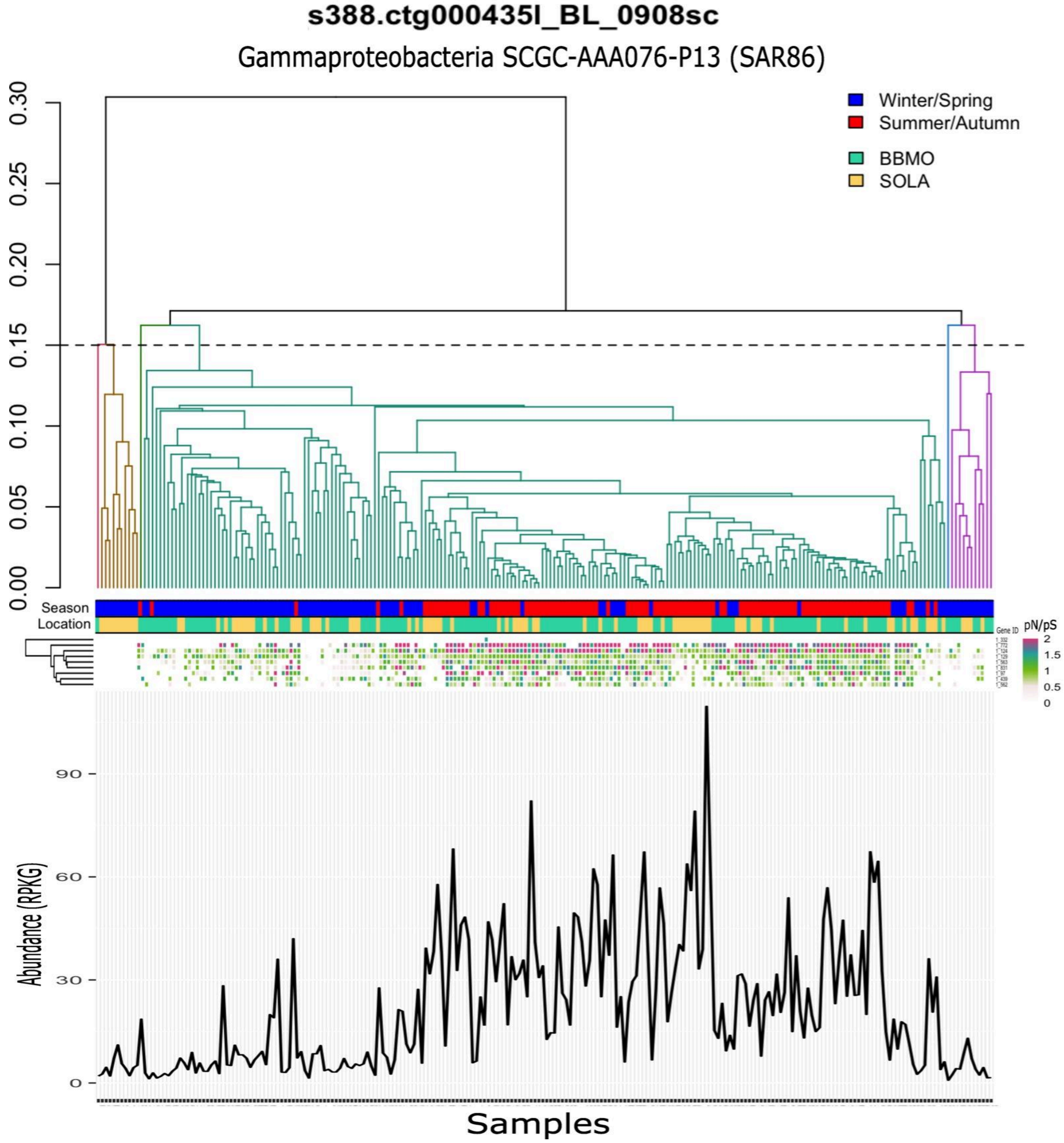
s388.ctg000435l_BL_0908sc
Gammaproteobacteria SCGC-AAA076-P13 (SAR86)
Winter/Spring
Summer/Autumn
BBMO
SOLA
Season
Location
Gene ID
pN/pS
Abundance (RPKG)
Samples

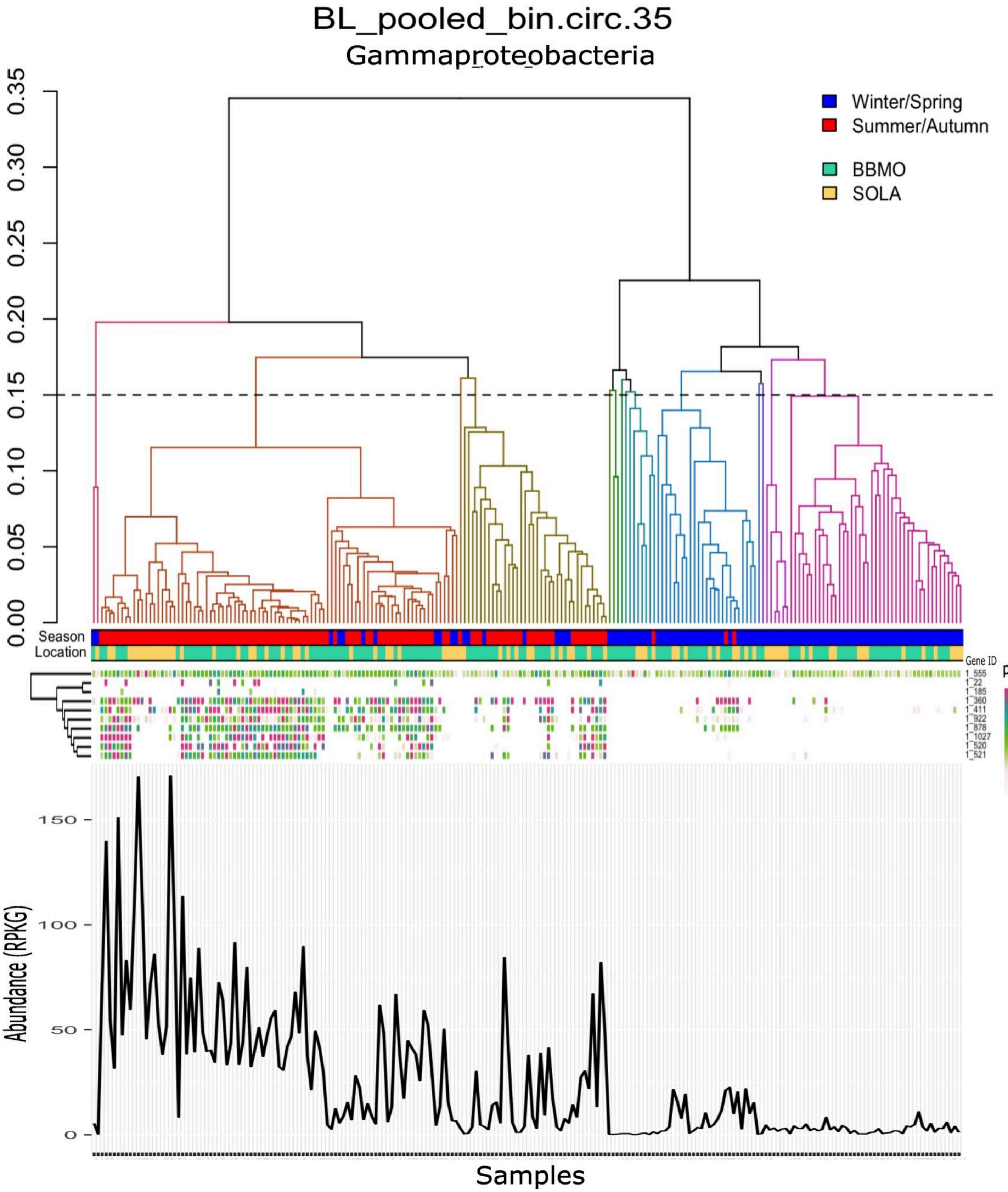

BL_pooled_bin.circ.35
Gammaproteobacteria
Winter/Spring
Summer/Autumn
BBMO
SOLA
0.00
0.05
0.10
0.15
0.20
0.25
0.30
0.35
Season
Location
Gene ID
1_555
1_22
1_185
1_360
1_411
1_922
1_878
1_1027
1_520
1_521
pN/pS
2
1.5
1
0.5
0
Abundance (RPKG)
150
100
50
0
Samples

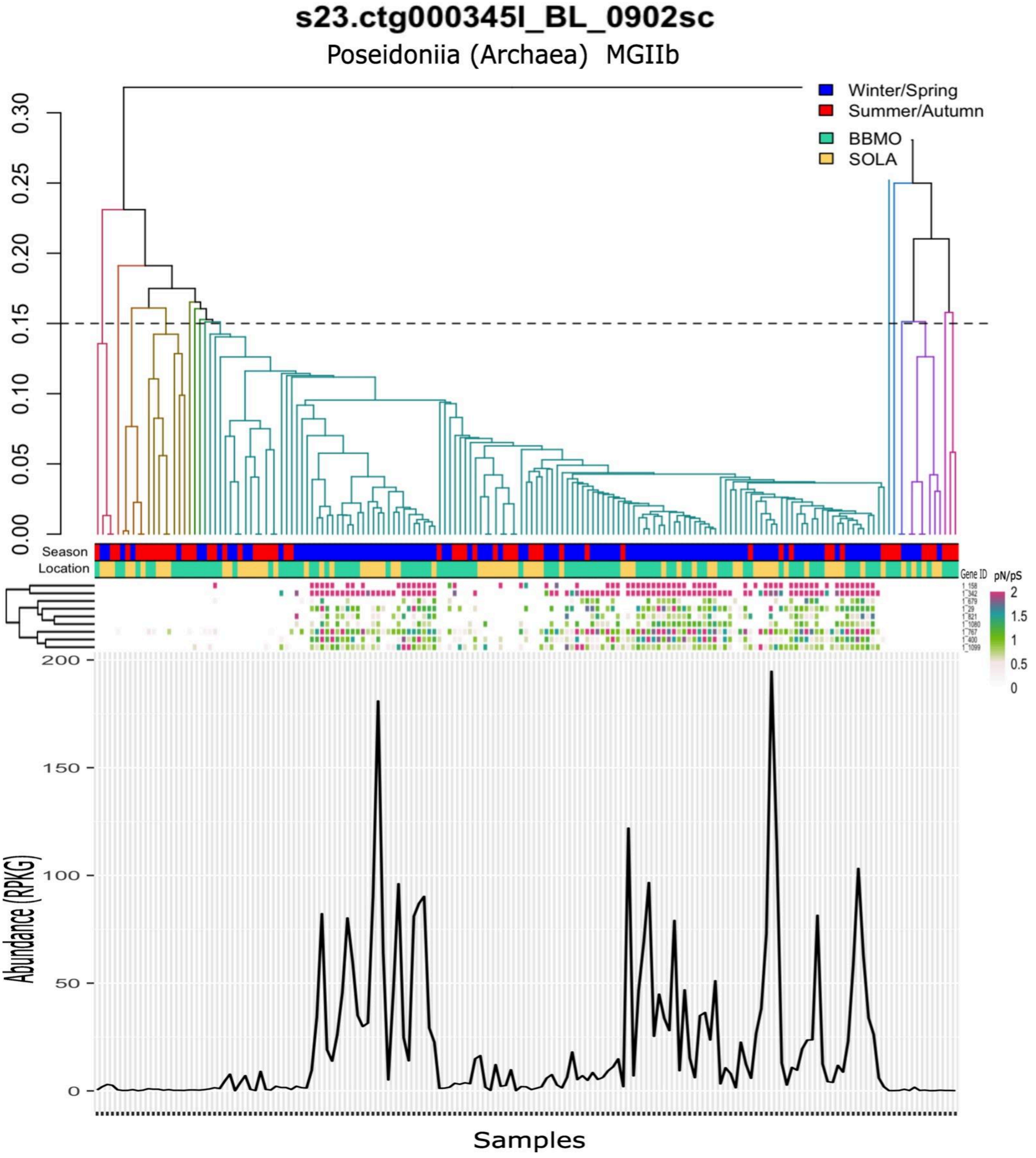
s23.ctg000345l_BL_0902sc
Poseidoniia (Archaea) MGIIb
Winter/Spring
Summer/Autumn
BBMO
SOLA
0.30
0.25
0.20
0.15
0.10
0.05
0.00
Season
Location
Gene ID
pN/pS
2
1.5
1
0.5
0
200
150
100
50
0
Abundance (RPKG)
Samples

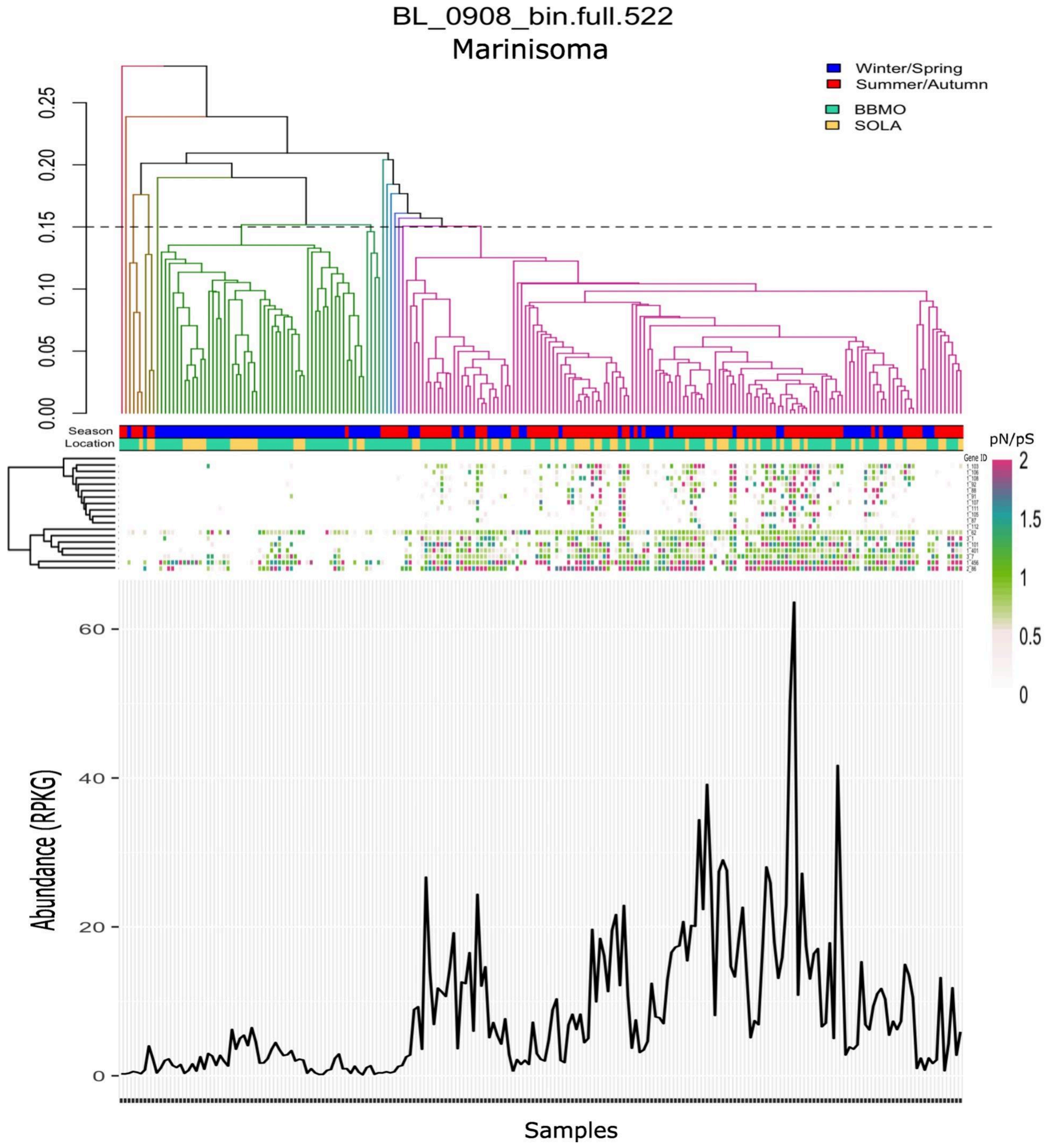

BL_0908_bin.full.522
Marinisoma
Winter/Spring
Summer/Autumn
BBMO
SOLA
0.25
0.20
0.15
0.10
0.05
0.00
Season
Location
pN/pS
Gene ID
2
1.5
1
0.5
0
60
40
20
0
Abundance (RPKG)
Samples

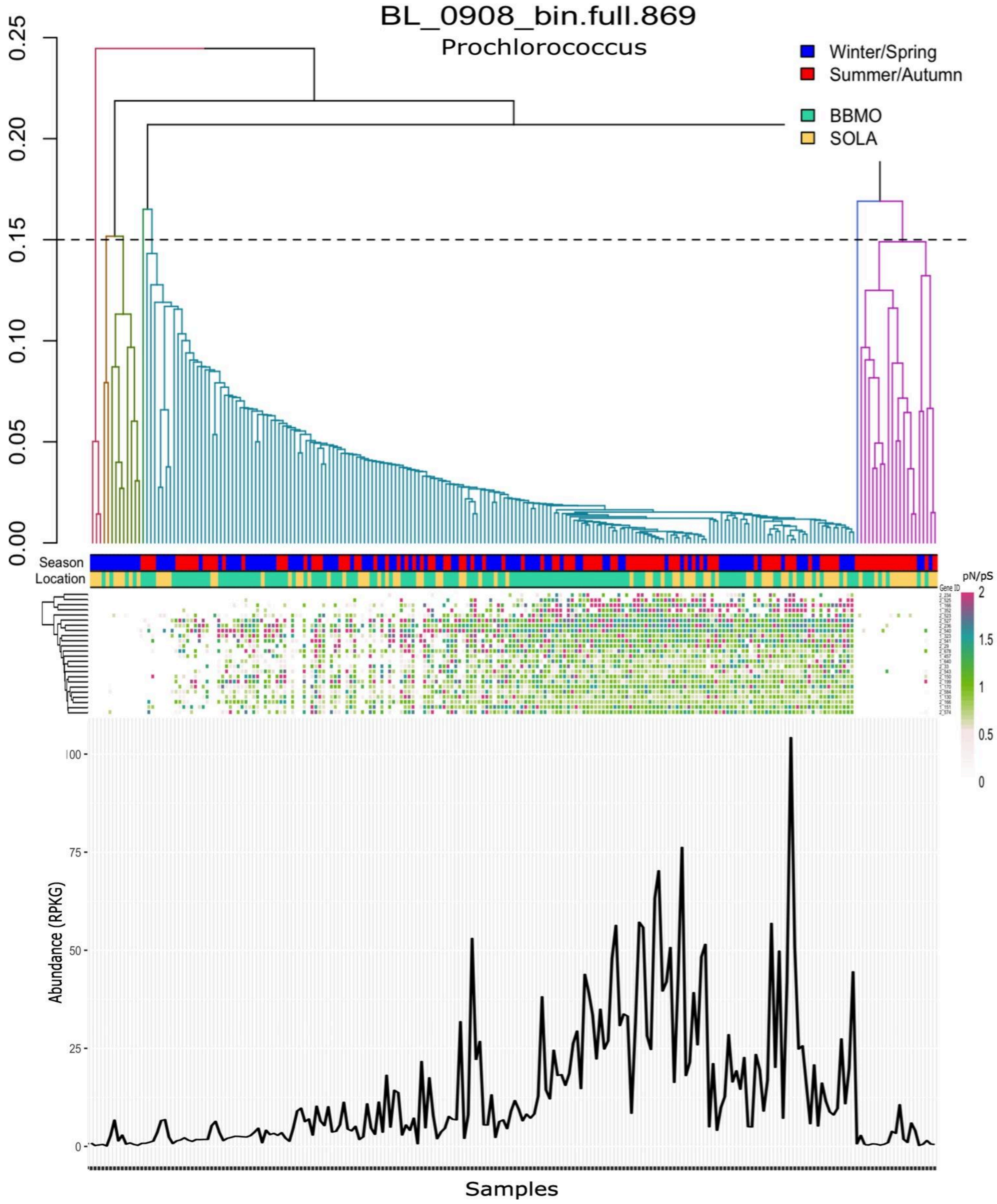

BL_0908_bin.full.869
Prochlorococcus
Winter/Spring
Summer/Autumn
BBMO
SOLA
0.25
0.20
0.15
0.10
0.05
0.00
Season
Location
Gene ID
pN/pS
2
1.5
1
0.5
0
Abundance (RPKG)
100
75
50
25
0
Samples

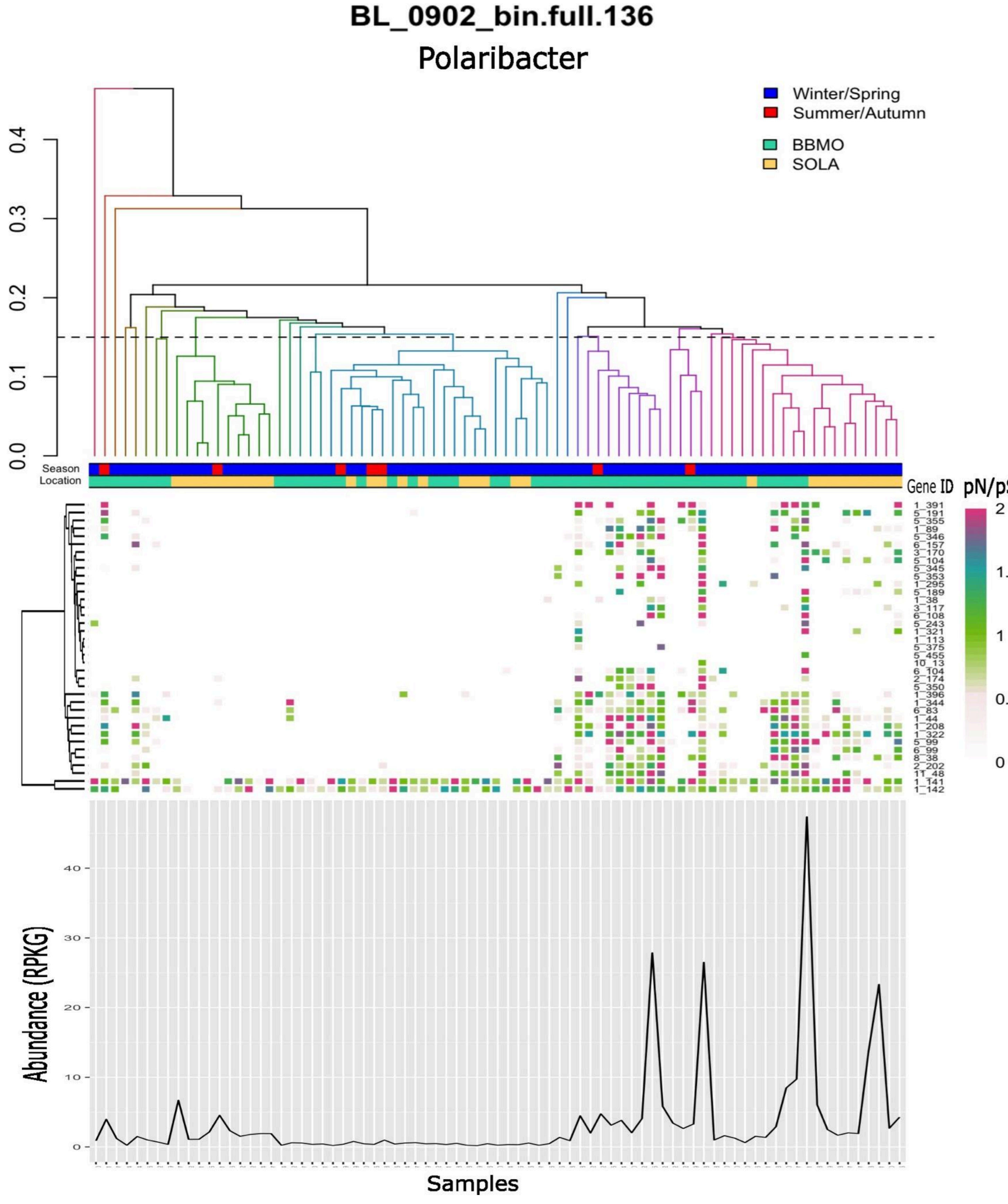

BL_0902_bin.full.136
Polaribacter
Winter/Spring
Summer/Autumn
BBMO
SOLA
0.0
0.1
0.2
0.3
0.4
Season
Location
Gene ID
pN/pS
2
1.5
1
0.5
0
1_391
5_191
5_355
1_89
5_346
6_157
3_170
5_104
5_345
5_353
1_295
5_189
1_38
3_117
6_108
5_243
1_321
1_113
5_375
5_455
10_13
6_104
2_174
5_350
1_396
1_344
6_83
1_44
1_208
1_322
5_99
6_99
8_38
2_202
11_48
1_141
1_142
Abundance (RPKG)
0
10
20
30
40
Samples

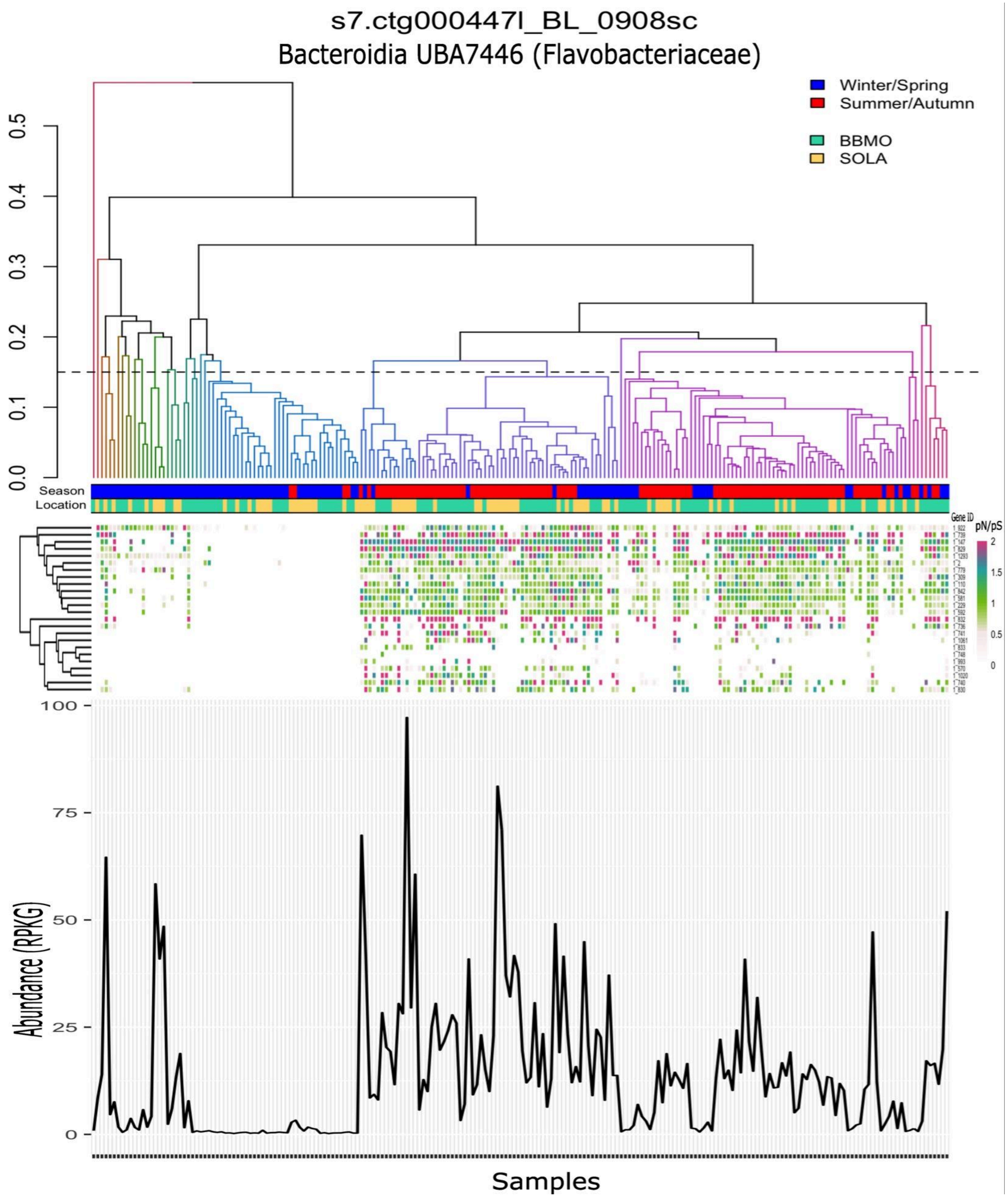
s7.ctg000447l_BL_0908sc
Bacteroidia UBA7446 (Flavobacteriaceae)
Winter/Spring
Summer/Autumn
BBMO
SOLA
0.0
0.1
0.2
0.3
0.4
0.5
Season
Location
Gene ID
pN/pS
2
1.5
1
0.5
0
100
75
50
25
0
Abundance (RPKG)
Samples

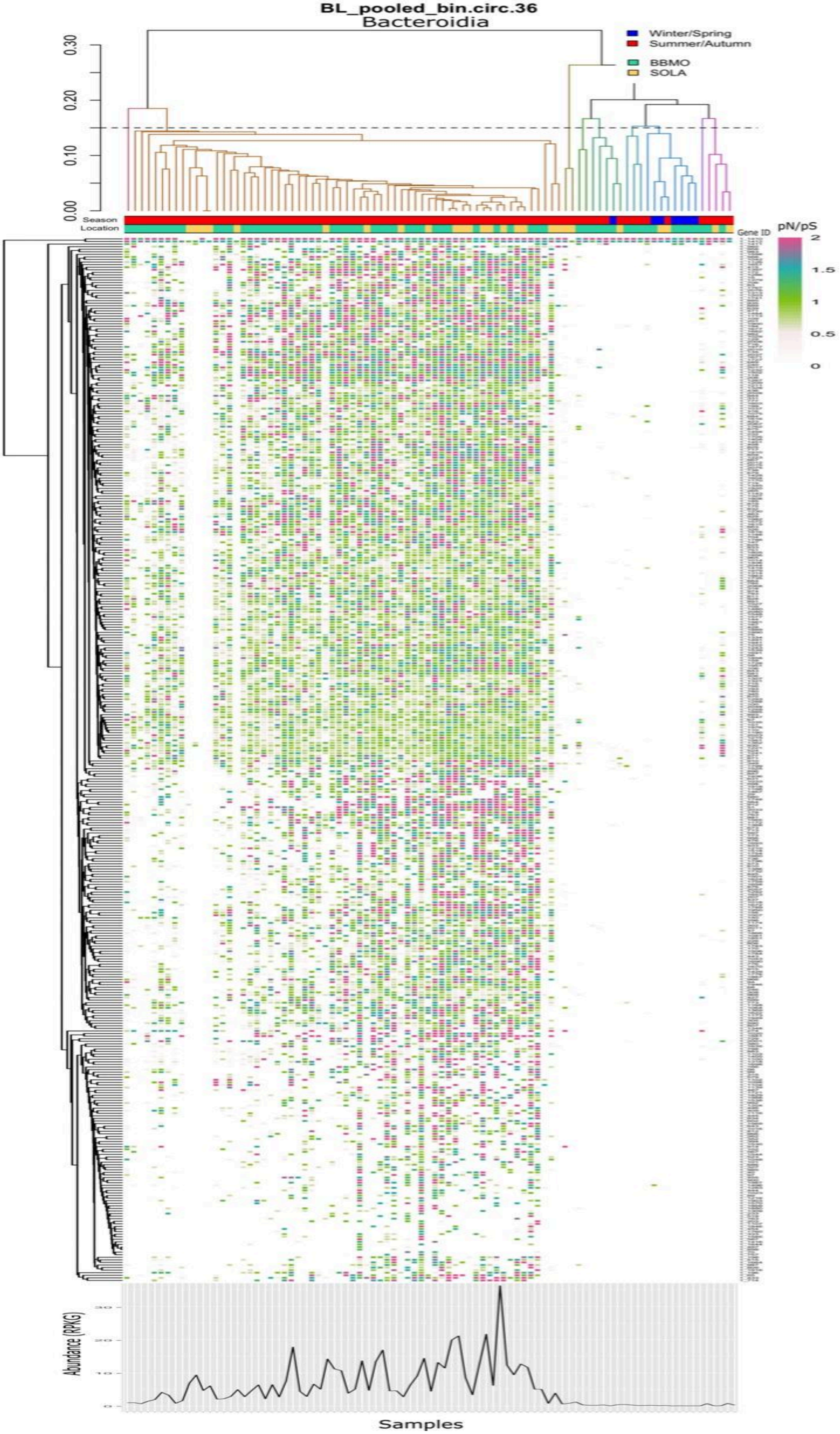
BL_pooled_bin.circ.36
Bacteroidia
Winter/Spring
Summer/Autumn
BBMO
SOLA
0.30
0.20
0.10
0.00
Season
Location
Gene ID
pN/pS
2
1.5
1
0.5
0
Abundance (RPKG)
30
20
10
0
Samples

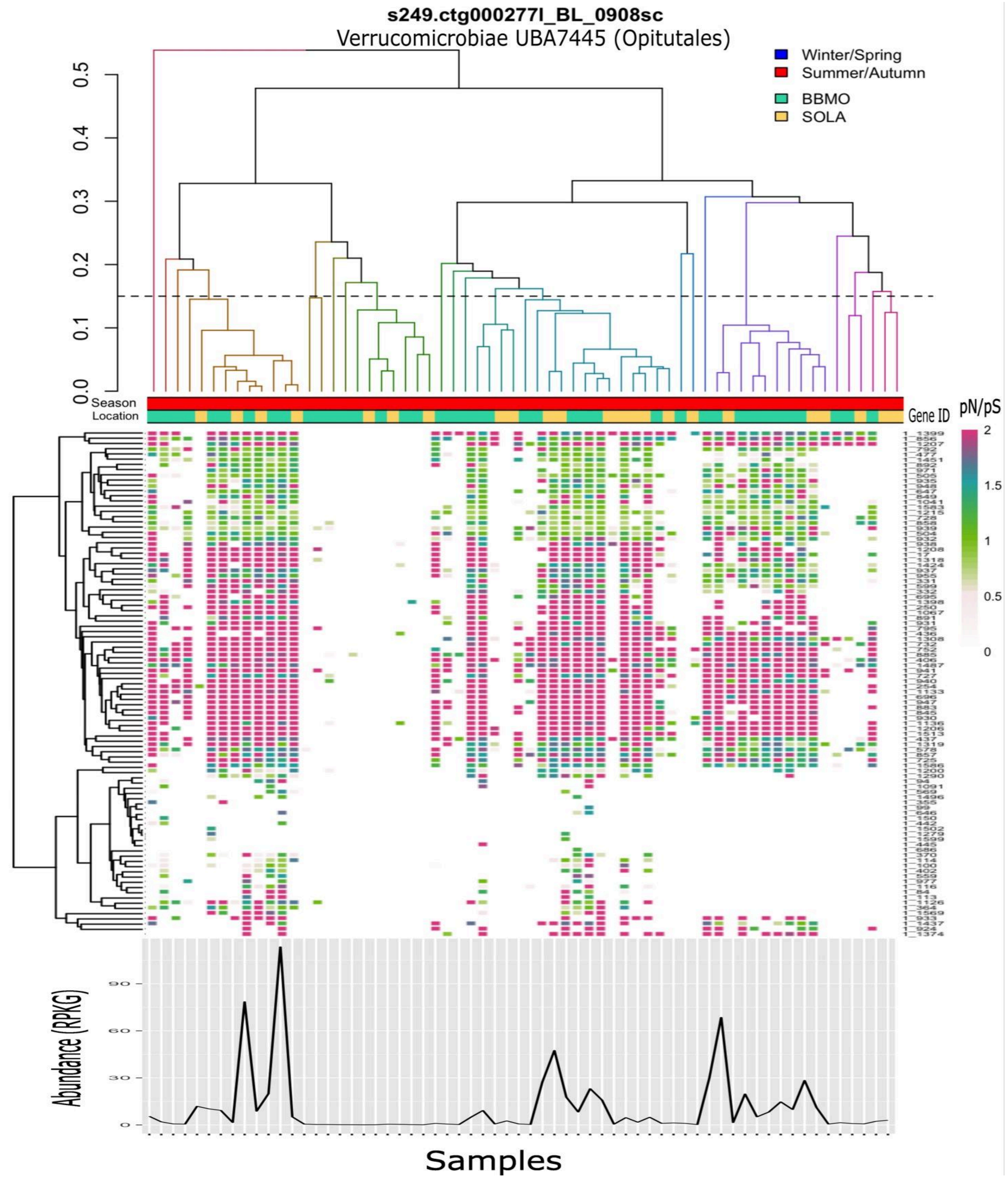

s249.ctg000277l_BL_0908sc
Verrucomicrobiae UBA7445 (Opitutales)
Winter/Spring
Summer/Autumn
BBMO
SOLA
0.5
0.4
0.3
0.2
0.1
0.0
Season
Location
Gene ID
pN/pS
2
1.5
1
0.5
0
Abundance (RPKG)
90
60
30
0
Samples

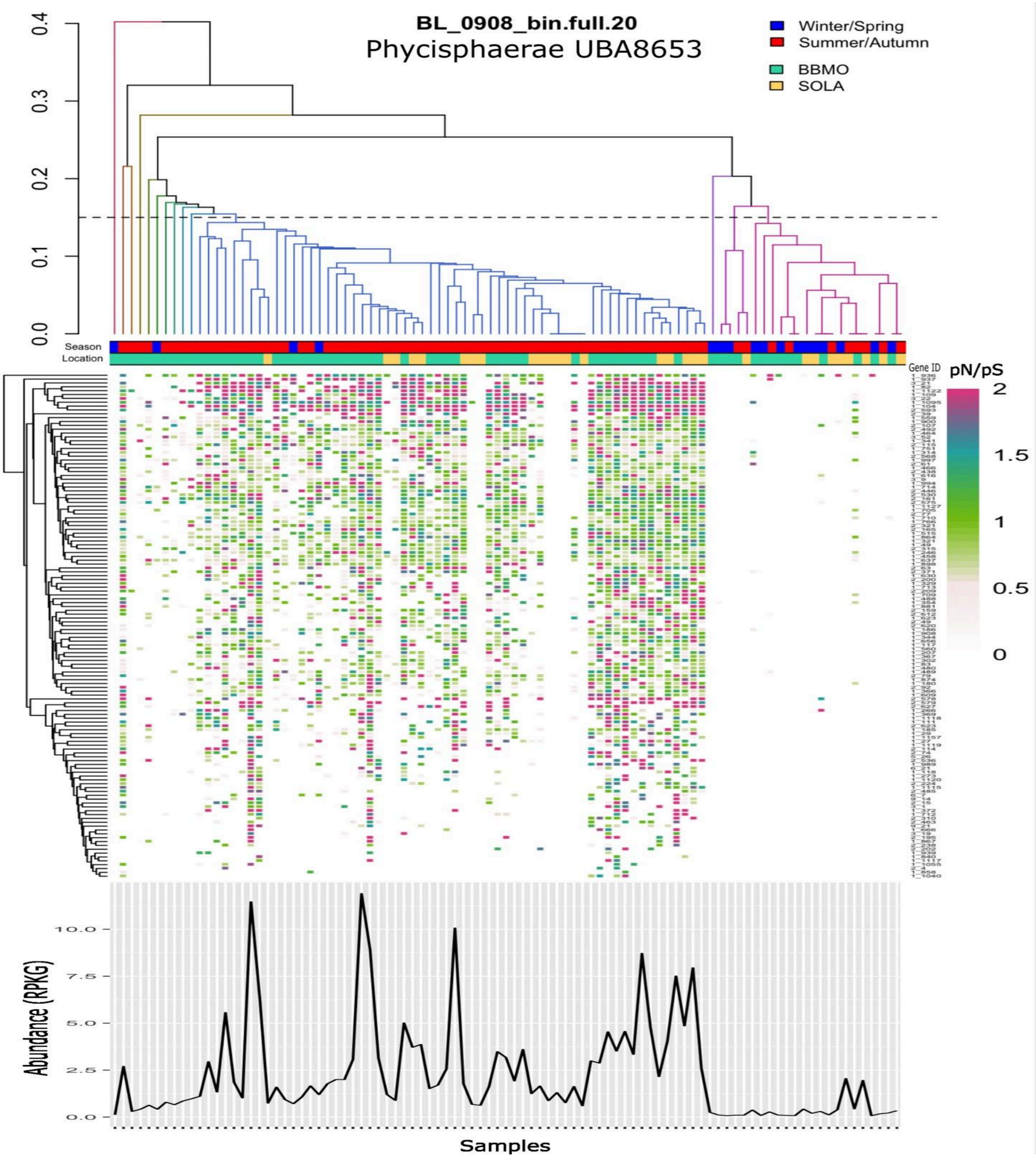
BL_0908_bin.full.20
Phycisphaerae UBA8653
Winter/Spring
Summer/Autumn
BBMO
SOLA
Season
Location
Gene ID
pN/pS
2
1.5
1
0.5
0
0.4
0.3
0.2
0.1
0.0
10.0
7.5
5.0
2.5
0.0
Abundance (RPKG)
Samples
