## Supplementary Figure 10 for "Seasonal population structure and adaptive signatures across long-term marine microbial time series"

**Supplementary Figure 10. Genome-wide population differentiation versus temporal distance for the 27 MAGs.** Pairwise genome-wide Fst values are plotted against the temporal distance (days) between samples for BBMO and SOLA, with each point representing one pair of samples. Only sample pairs for which both samples had >25% horizontal coverage of the corresponding MAG were included. Blue lines show linear regression fits and red dashed lines show LOESS fits; shaded areas indicate 95% confidence intervals around the fitted relationships. The slope, adjusted $R^2$, and p-value of the linear regression are shown in each panel. Analyses are shown only for time series with sufficient data for the corresponding MAG.

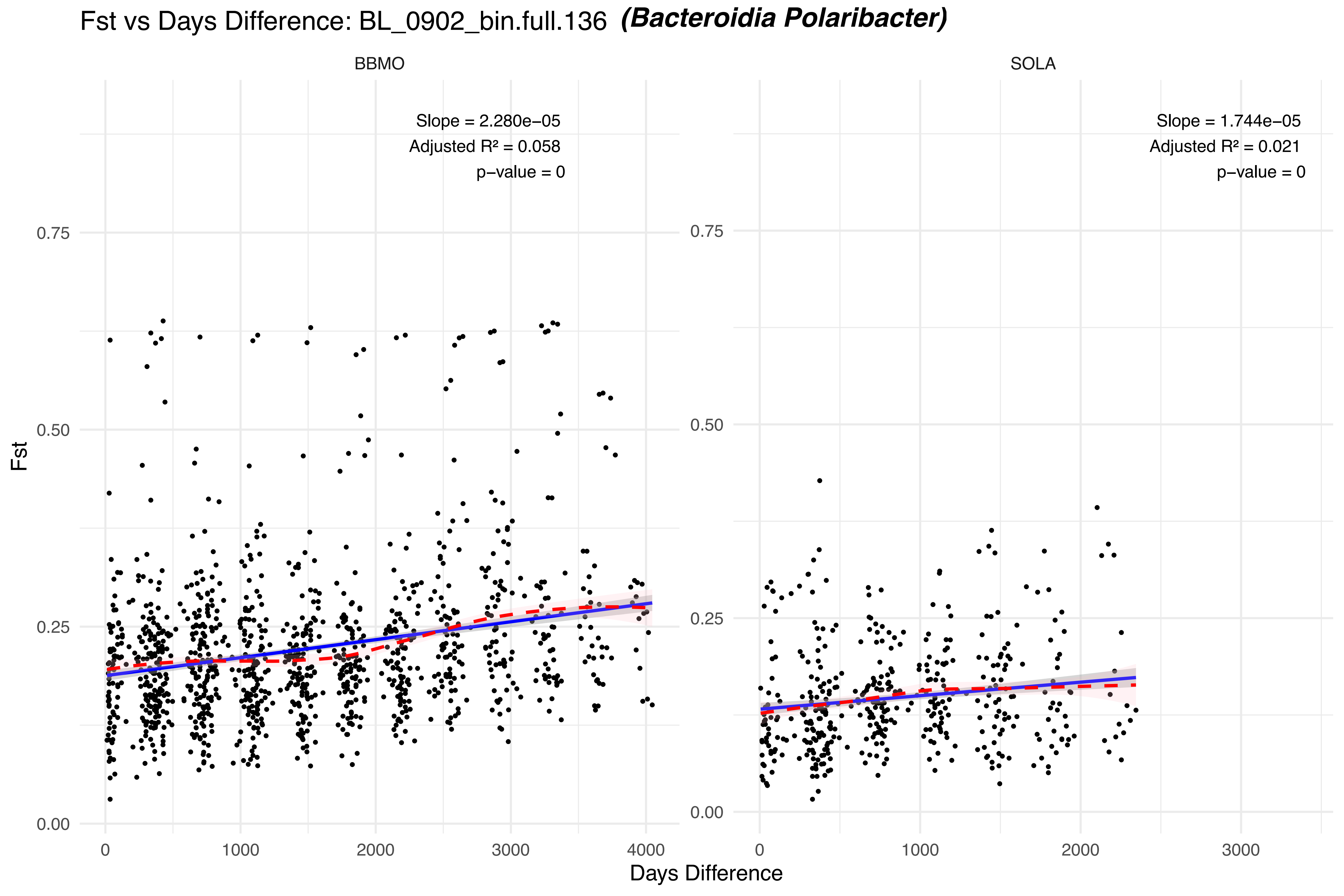

Fst vs Days Difference: BL_0902_bin.full.136 (Bacteroidia Polaribacter)
BBMO
SOLA
Slope = 2.280e−05
Adjusted R² = 0.058
p−value = 0
Slope = 1.744e−05
Adjusted R² = 0.021
p−value = 0
Fst
0.00
0.25
0.50
0.75
0
1000
2000
3000
4000
Days Difference

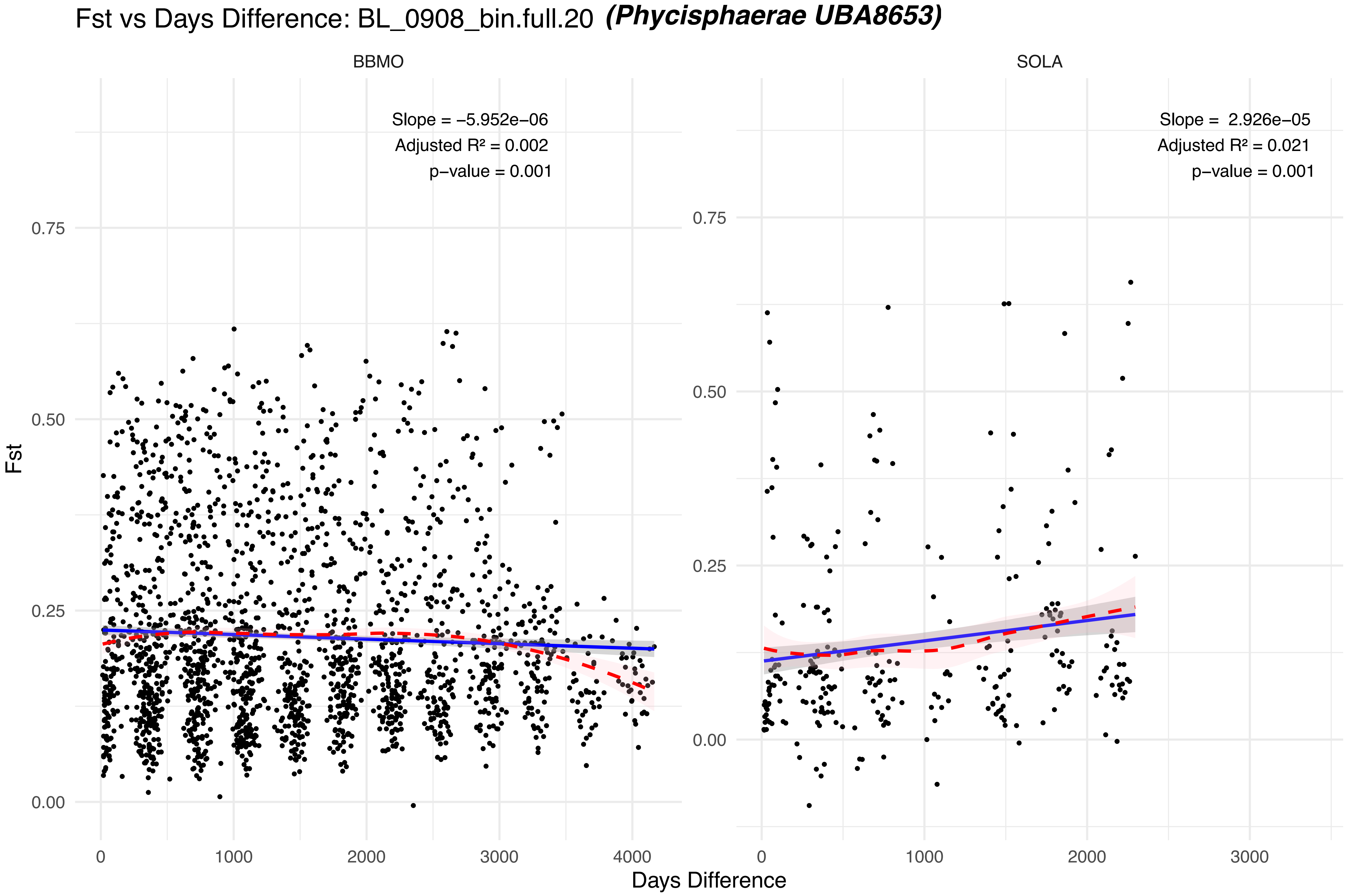

Fst vs Days Difference: BL_0908_bin.full.20 (Phycisphaerae UBA8653)
BBMO
SOLA
Slope = −5.952e−06
Adjusted R² = 0.002
p−value = 0.001
Slope = 2.926e−05
Adjusted R² = 0.021
p−value = 0.001
Fst
Days Difference
0.00
0.25
0.50
0.75
0
1000
2000
3000
4000

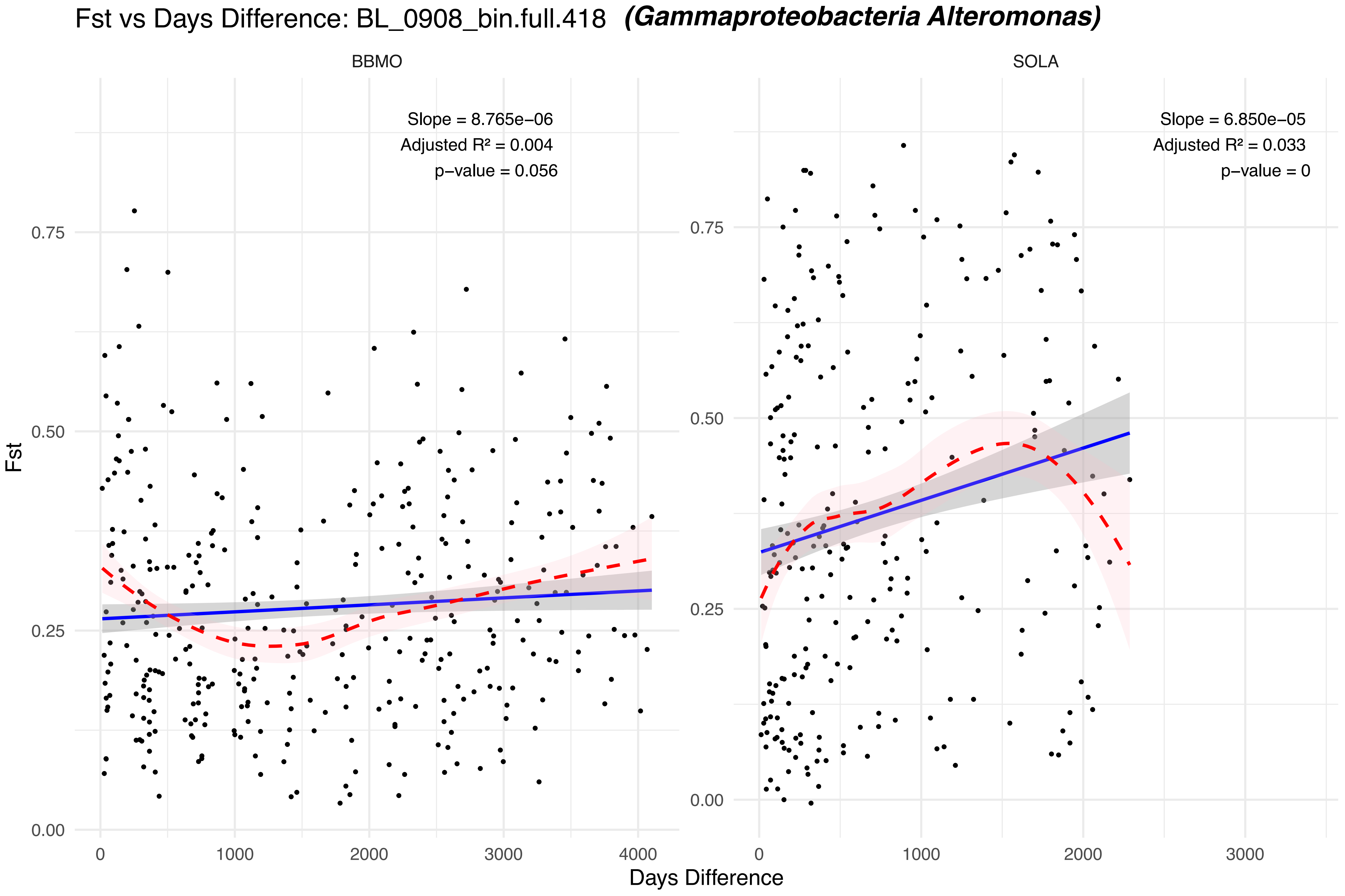

Fst vs Days Difference: BL_0908_bin.full.418 (Gammaproteobacteria Alteromonas)
BBMO
SOLA
Slope = 8.765e−06
Adjusted R² = 0.004
p−value = 0.056
Slope = 6.850e−05
Adjusted R² = 0.033
p−value = 0
Fst
Days Difference
0.00
0.25
0.50
0.75
0
1000
2000
3000
4000

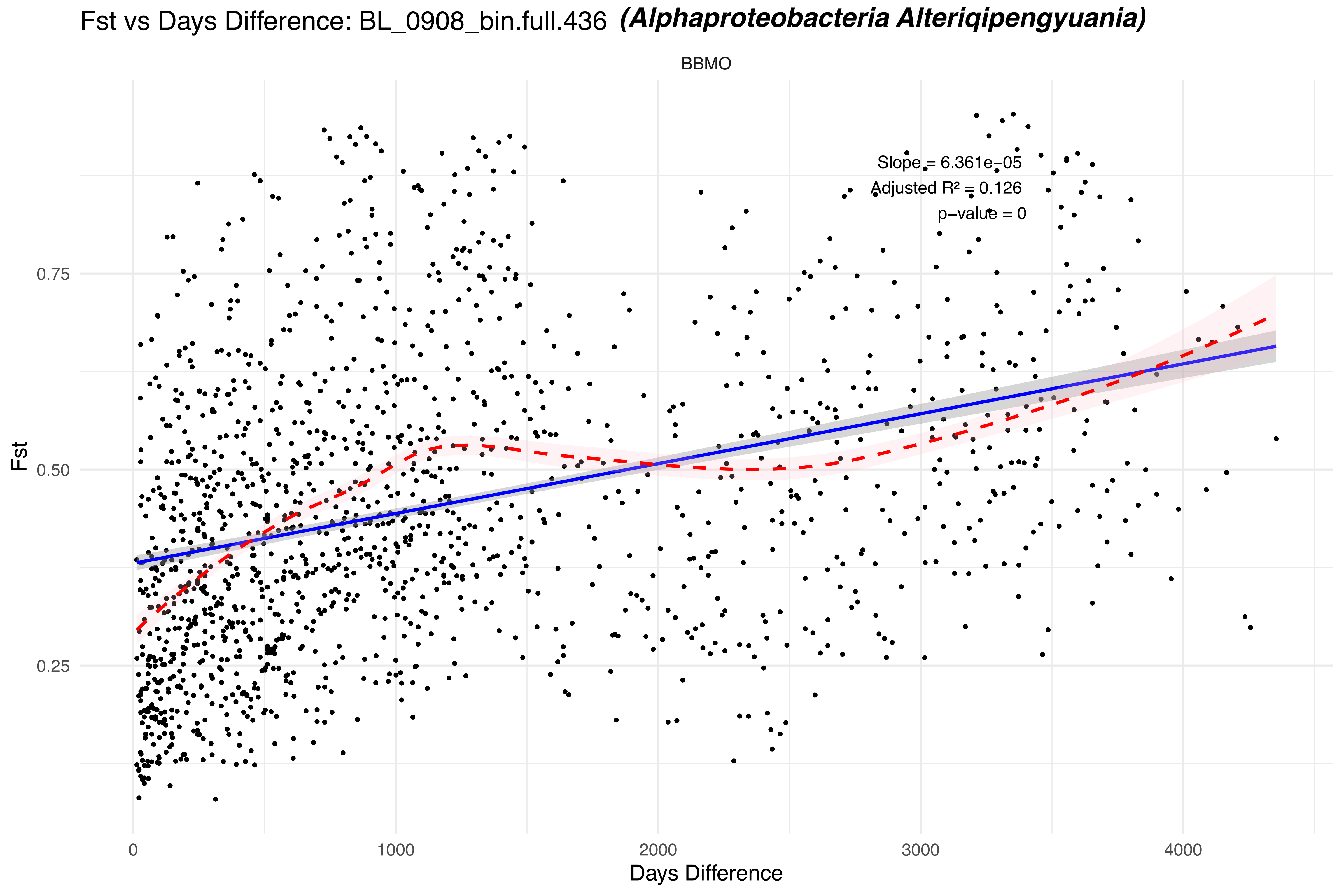

Fst vs Days Difference: BL_0908_bin.full.436 (Alphaproteobacteria Alteriqipengyuania)
BBMO
Slope = 6.361e−05
Adjusted R² = 0.126
p−value = 0
Fst
0.25
0.50
0.75
0
1000
2000
3000
4000
Days Difference

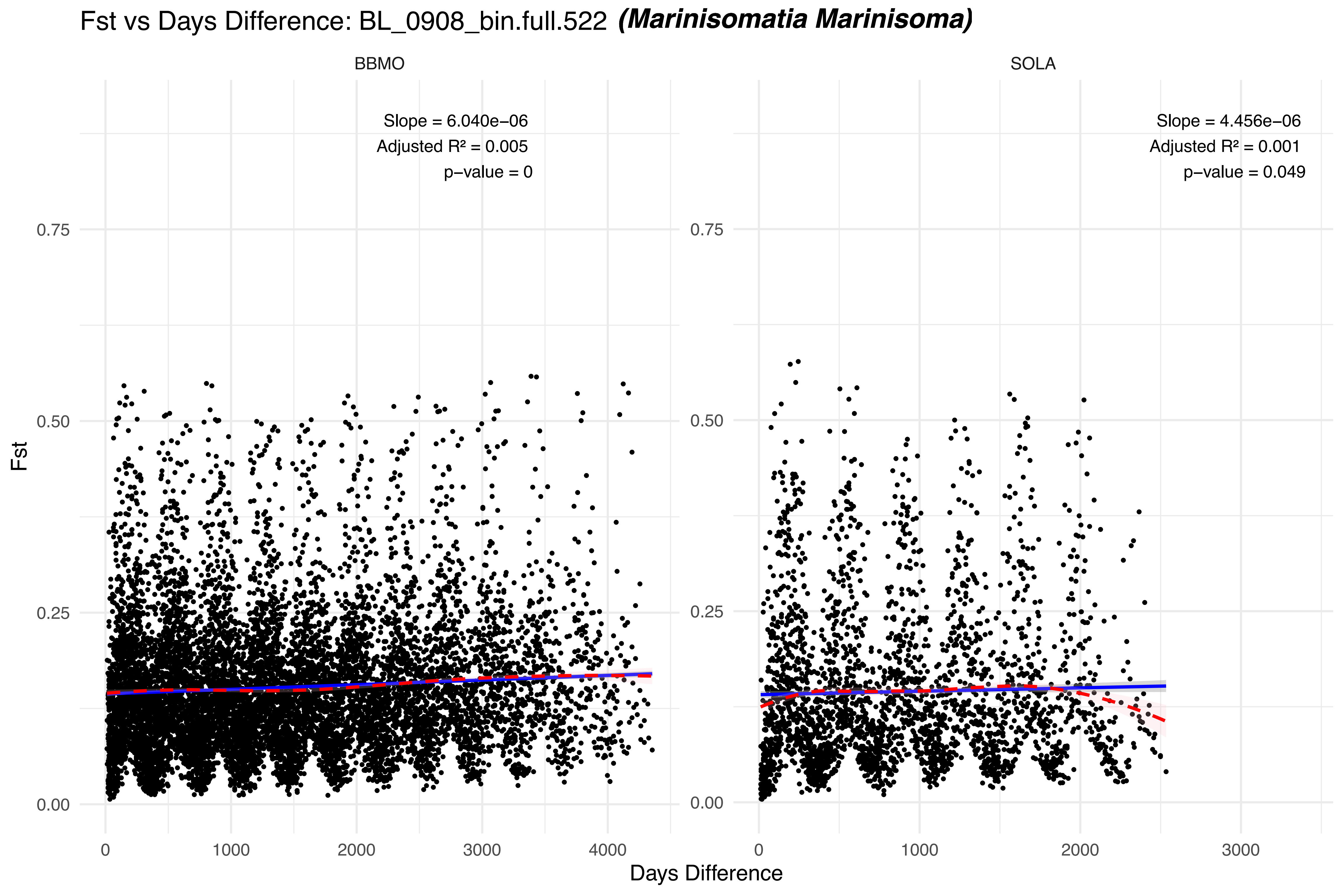

Fst vs Days Difference: BL_0908_bin.full.522 (Marinisomatia Marinisoma)
BBMO
SOLA
Slope = 6.040e−06
Adjusted R² = 0.005
p−value = 0
Slope = 4.456e−06
Adjusted R² = 0.001
p−value = 0.049
Fst
0.00
0.25
0.50
0.75
0
1000
2000
3000
4000
Days Difference

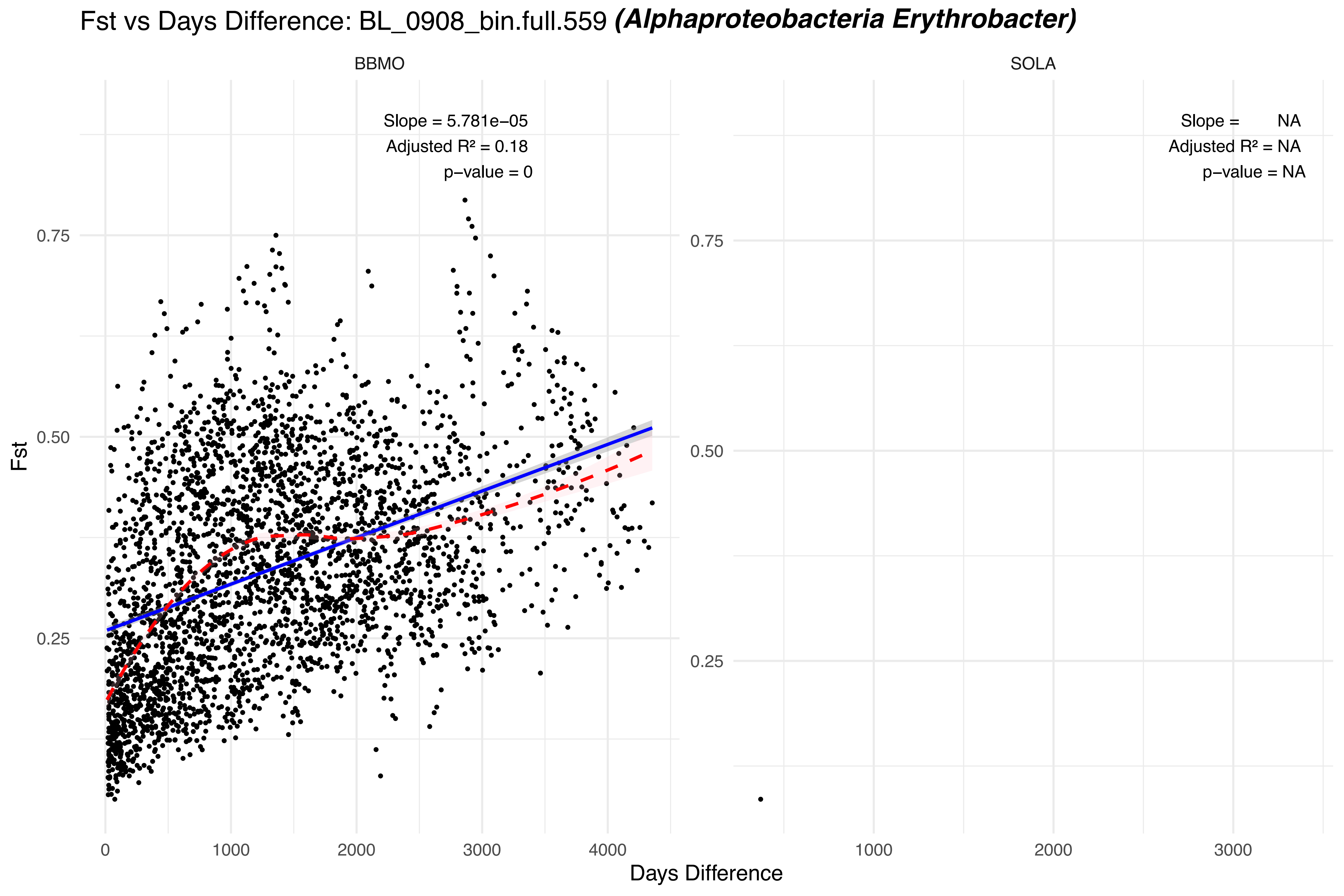

Fst vs Days Difference: BL_0908_bin.full.559 (Alphaproteobacteria Erythrobacter)
BBMO
SOLA
Slope = 5.781e−05
Adjusted R² = 0.18
p−value = 0
Slope = NA
Adjusted R² = NA
p−value = NA
Fst
0.25
0.50
0.75
0
1000
2000
3000
4000
Days Difference

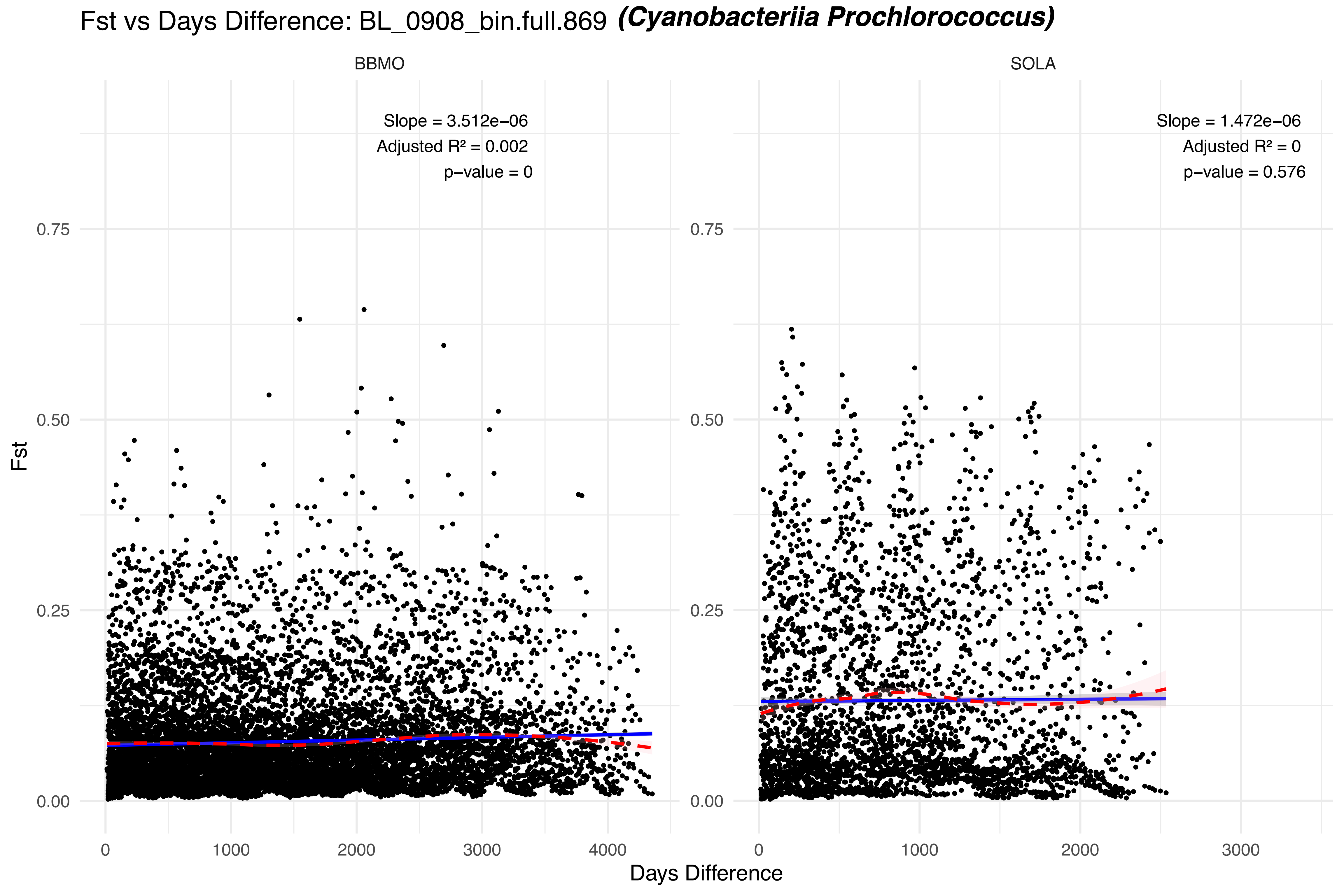

Fst vs Days Difference: BL_0908_bin.full.869 (Cyanobacteriia Prochlorococcus)
BBMO
SOLA
Slope = 3.512e−06
Adjusted R² = 0.002
p−value = 0
Slope = 1.472e−06
Adjusted R² = 0
p−value = 0.576
Fst
0.00
0.25
0.50
0.75
0
1000
2000
3000
4000
Days Difference

### Fst vs Days Difference: BL_1001_bin.full.234 ***(Gammaproteobacteria Glaciecola)***

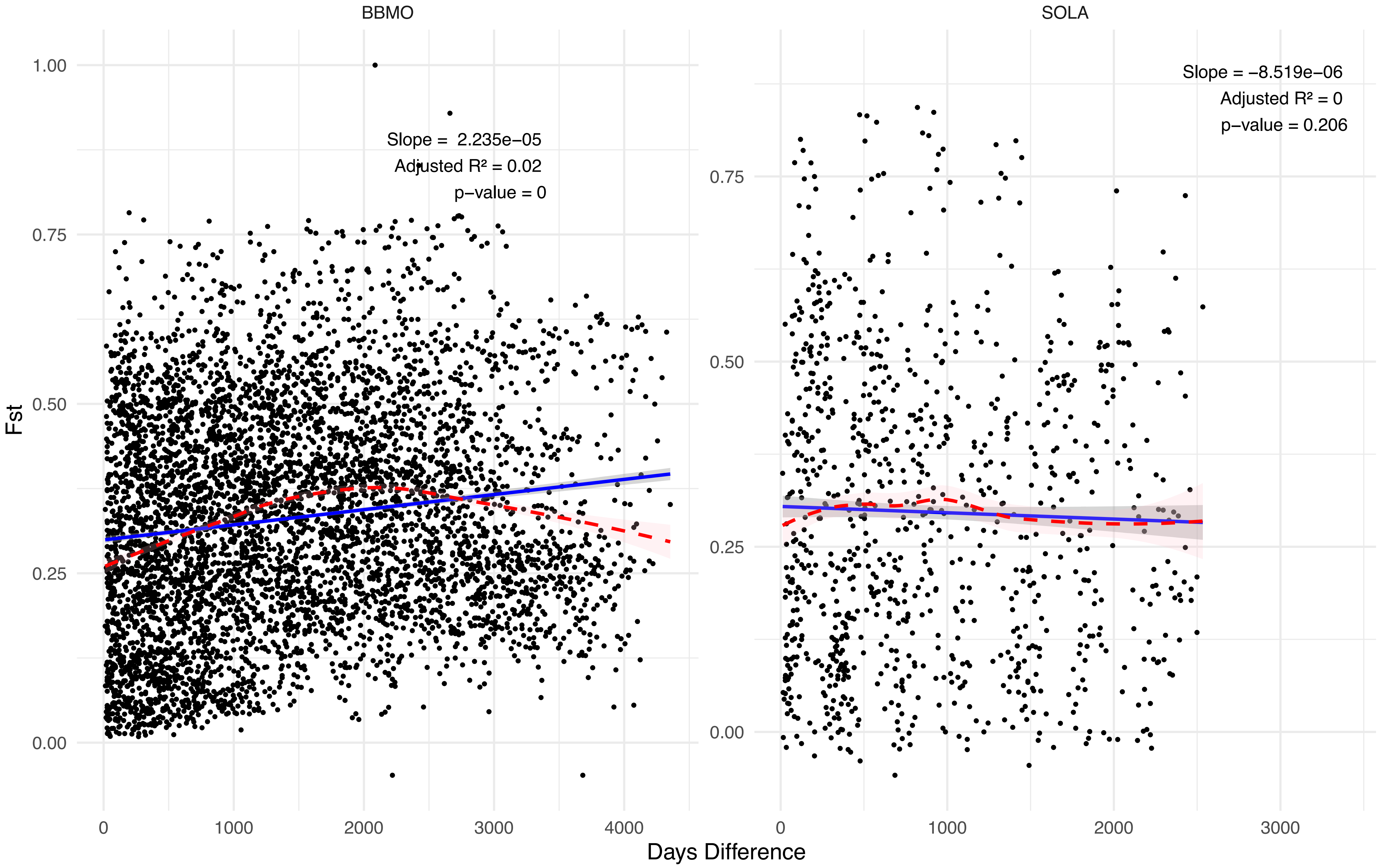

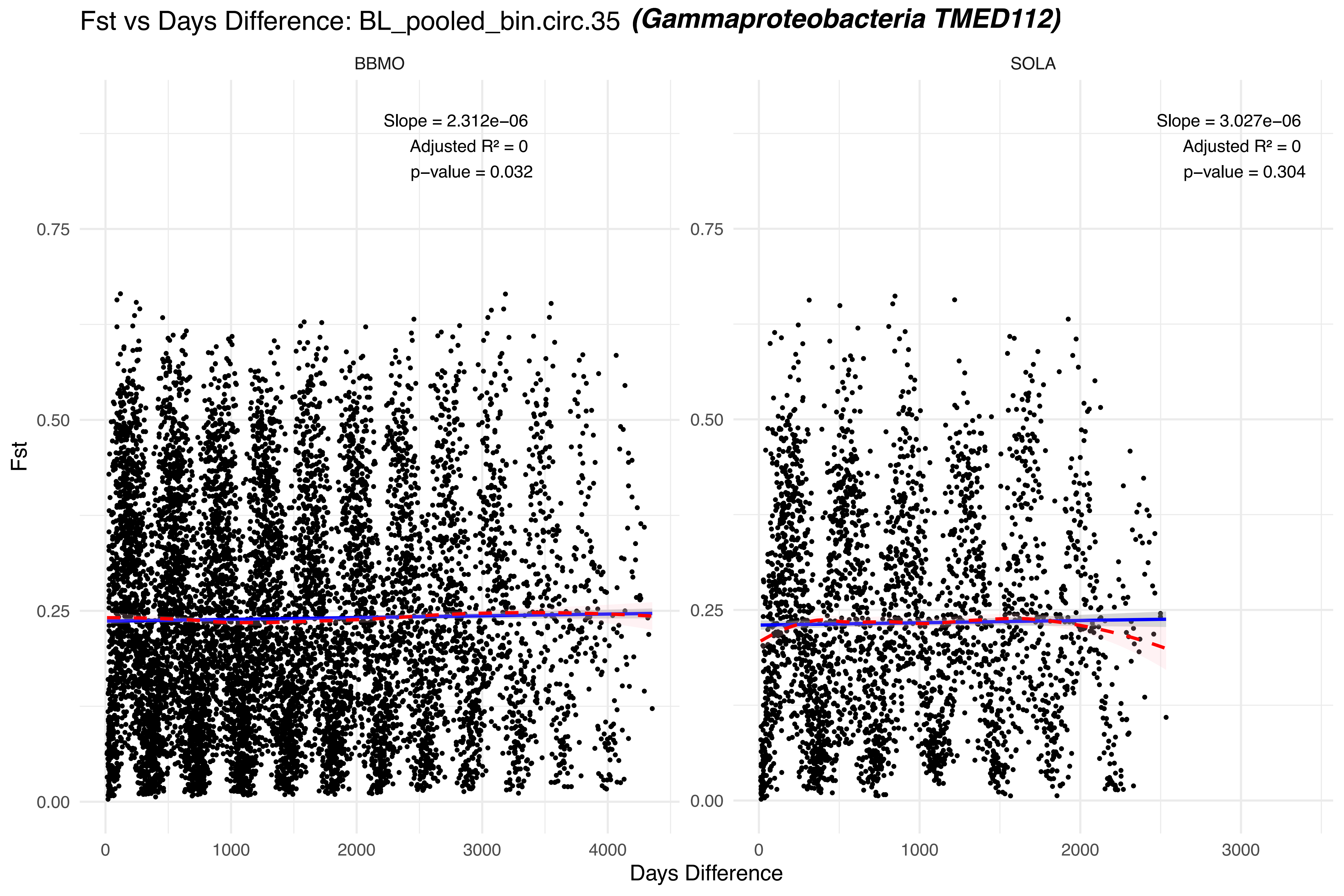

Fst vs Days Difference: BL_pooled_bin.circ.35 (Gammaproteobacteria TMED112)
BBMO
SOLA
Slope = 2.312e−06
Adjusted R² = 0
p−value = 0.032
Slope = 3.027e−06
Adjusted R² = 0
p−value = 0.304
Fst
0.75
0.50
0.25
0.00
0
1000
2000
3000
4000
Days Difference

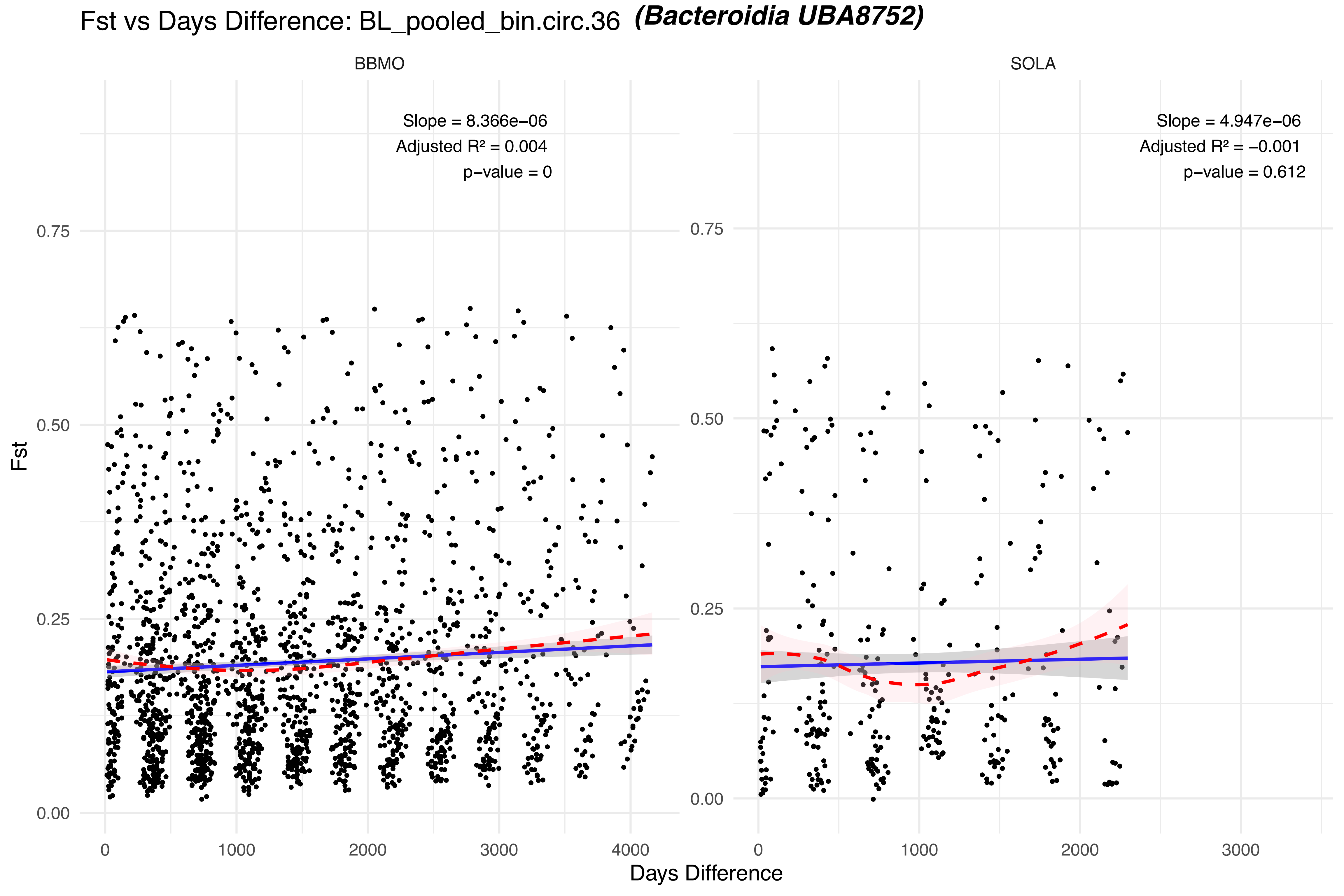

Fst vs Days Difference: BL_pooled_bin.circ.36 (Bacteroidia UBA8752)
BBMO
SOLA
Slope = 8.366e−06
Adjusted R² = 0.004
p−value = 0
Slope = 4.947e−06
Adjusted R² = −0.001
p−value = 0.612
Fst
0.00
0.25
0.50
0.75
0
1000
2000
3000
4000
Days Difference

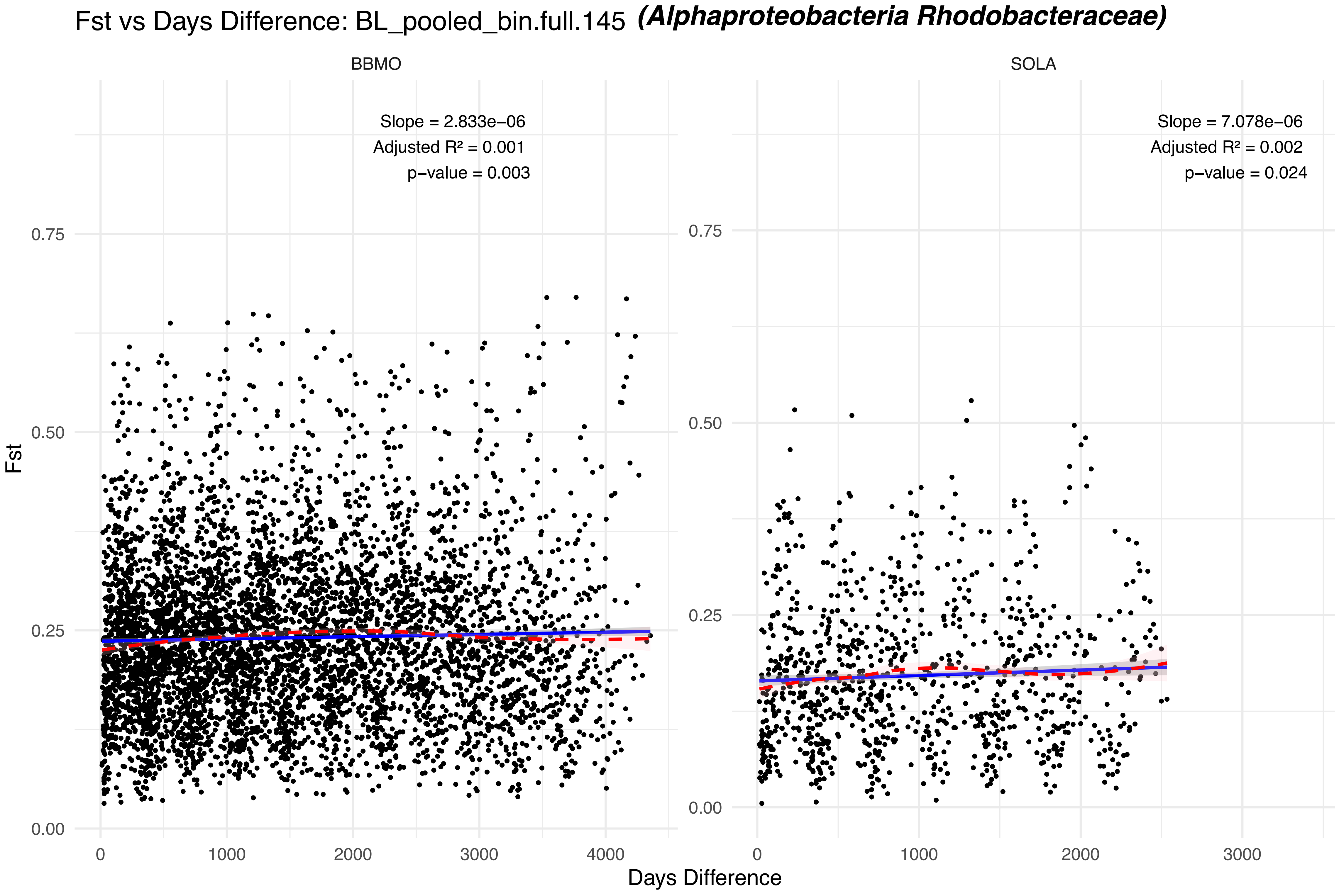

Fst vs Days Difference: BL_pooled_bin.full.145 (Alphaproteobacteria Rhodobacteraceae)
BBMO
SOLA
Slope = 2.833e−06
Adjusted R² = 0.001
p−value = 0.003
Slope = 7.078e−06
Adjusted R² = 0.002
p−value = 0.024
Fst
0.00
0.25
0.50
0.75
Days Difference
0
1000
2000
3000
4000

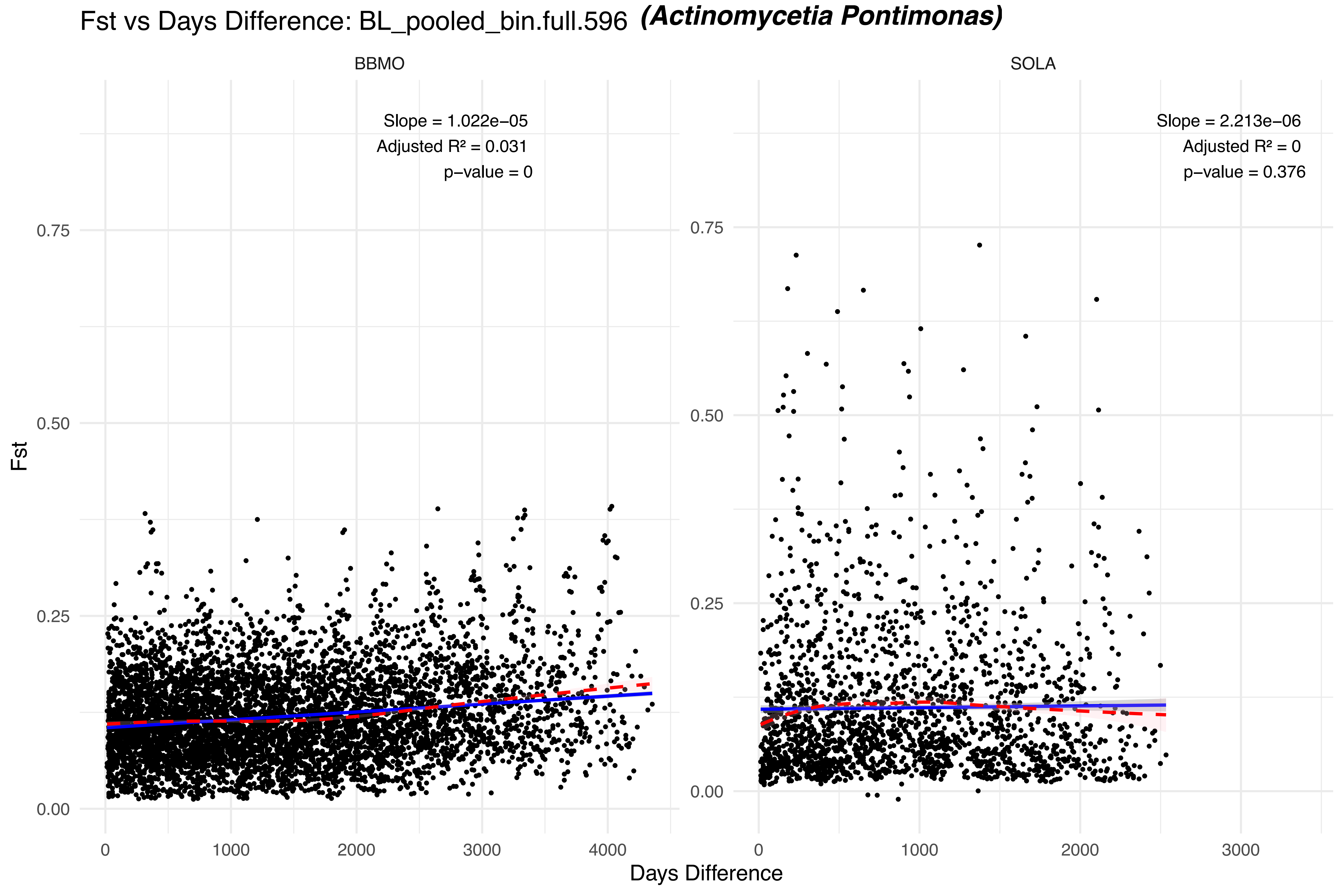

Fst vs Days Difference: BL_pooled_bin.full.596 (Actinomycetia Pontimonas)
BBMO
SOLA
Slope = 1.022e−05
Adjusted R² = 0.031
p−value = 0
Slope = 2.213e−06
Adjusted R² = 0
p−value = 0.376
Fst
0.00
0.25
0.50
0.75
0
1000
2000
3000
4000
Days Difference

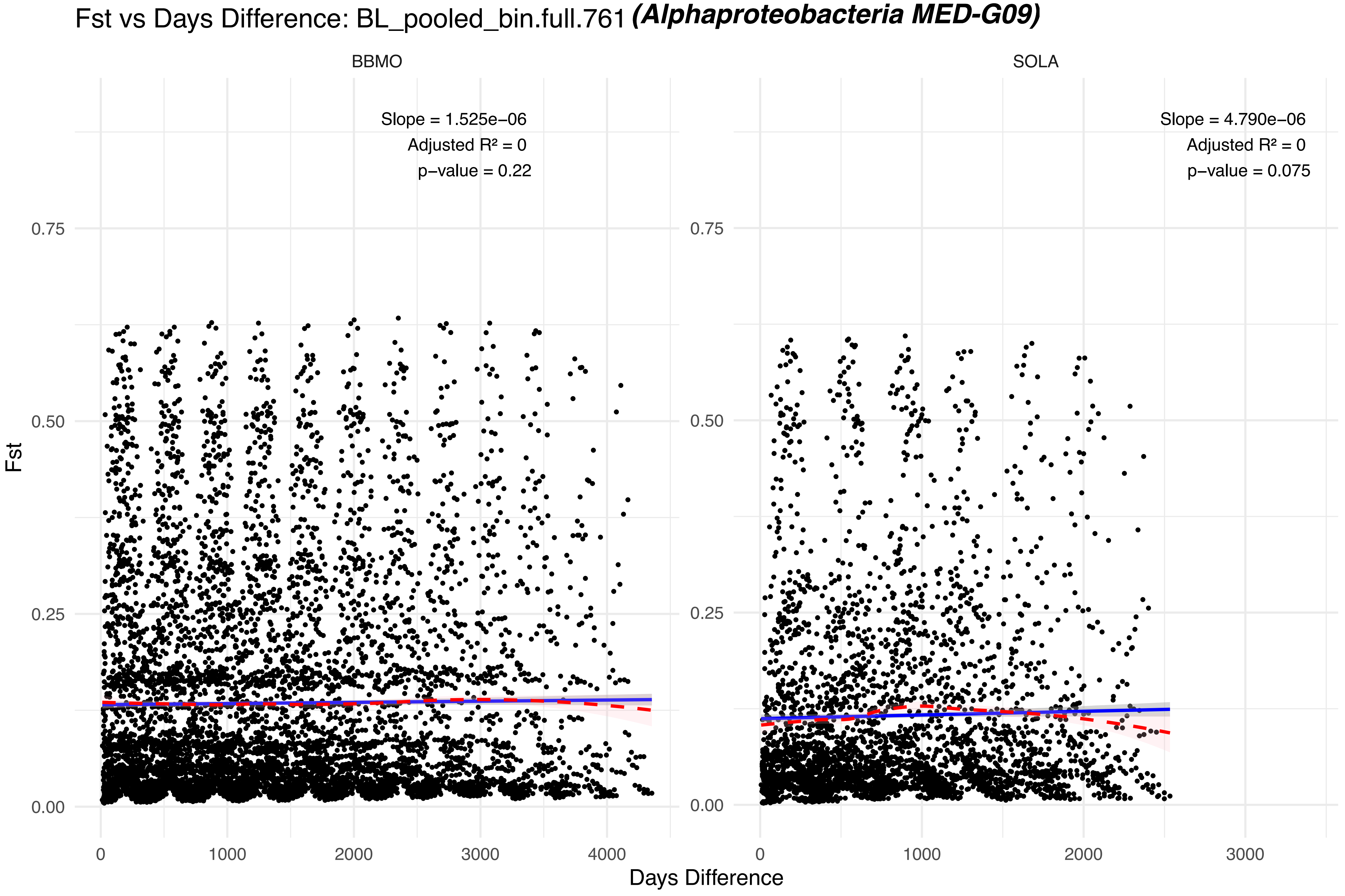
Fst vs Days Difference: BL_pooled_bin.full.761 (Alphaproteobacteria MED-G09)
BBMO
SOLA
Slope = 1.525e−06
Adjusted R² = 0
p−value = 0.22
Slope = 4.790e−06
Adjusted R² = 0
p−value = 0.075
Fst
0.00
0.25
0.50
0.75
0
1000
2000
3000
4000
Days Difference

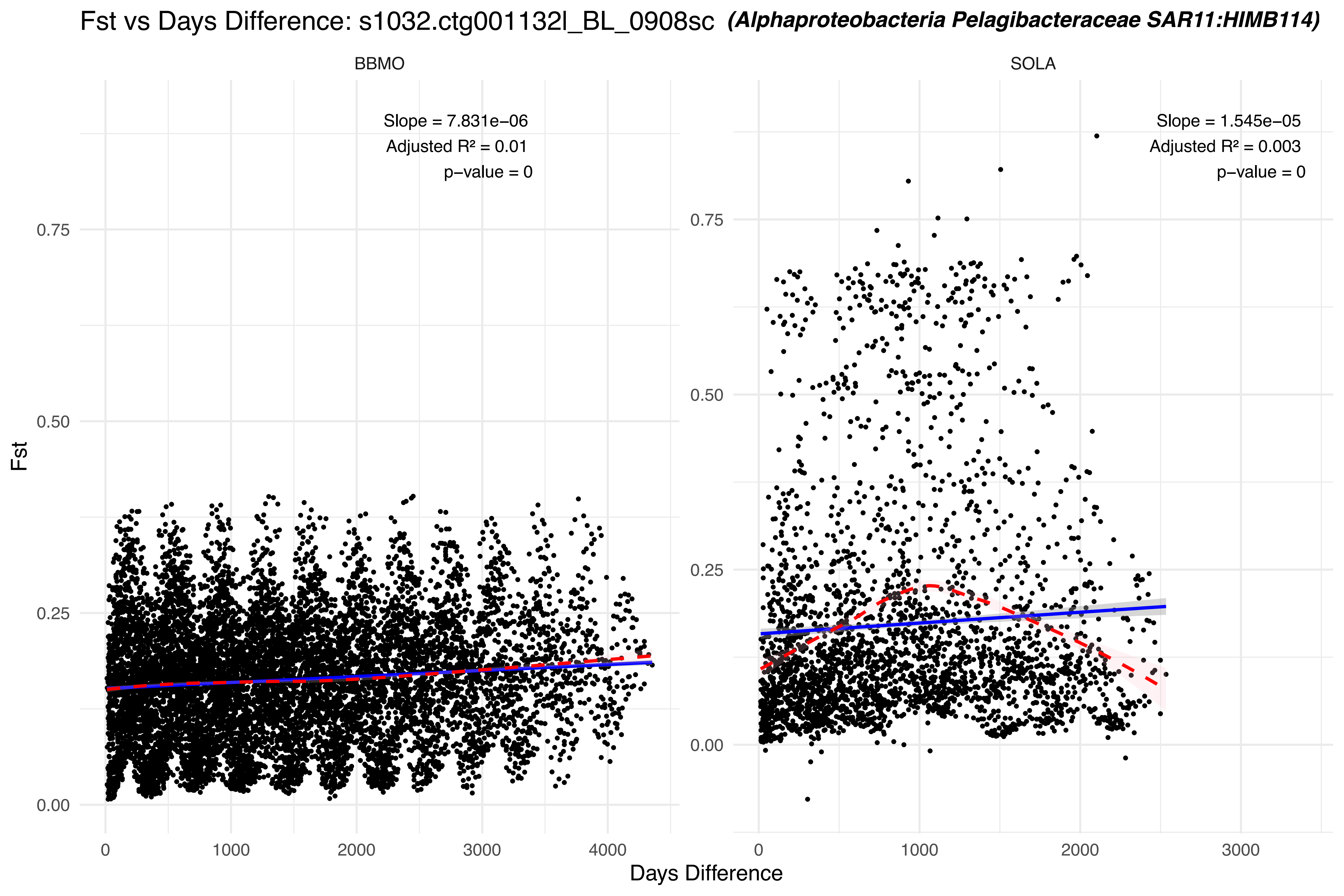

Fst vs Days Difference: s1032.ctg001132l_BL_0908sc (Alphaproteobacteria Pelagibacteraceae SAR11:HIMB114)
BBMO
SOLA
Slope = 7.831e−06
Adjusted R² = 0.01
p−value = 0
Slope = 1.545e−05
Adjusted R² = 0.003
p−value = 0
Fst
0.00
0.25
0.50
0.75
0
1000
2000
3000
4000
Days Difference

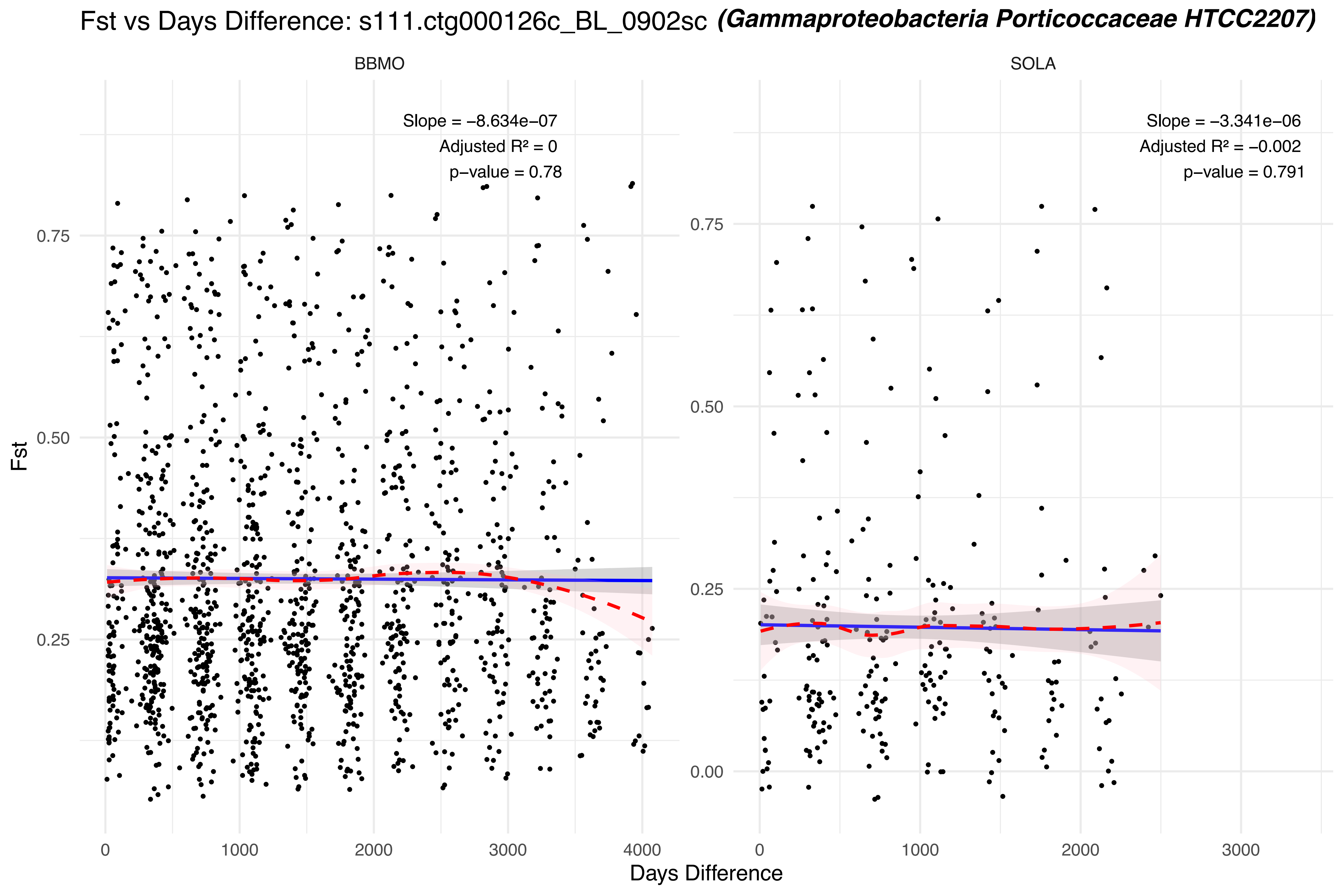
Fst vs Days Difference: s111.ctg000126c_BL_0902sc (Gammaproteobacteria Porticoccaceae HTCC2207)
BBMO
SOLA
Slope = −8.634e−07
Adjusted R² = 0
p−value = 0.78
Slope = −3.341e−06
Adjusted R² = −0.002
p−value = 0.791
Fst
0.75
0.50
0.25
0.00
0
1000
2000
3000
4000
Days Difference

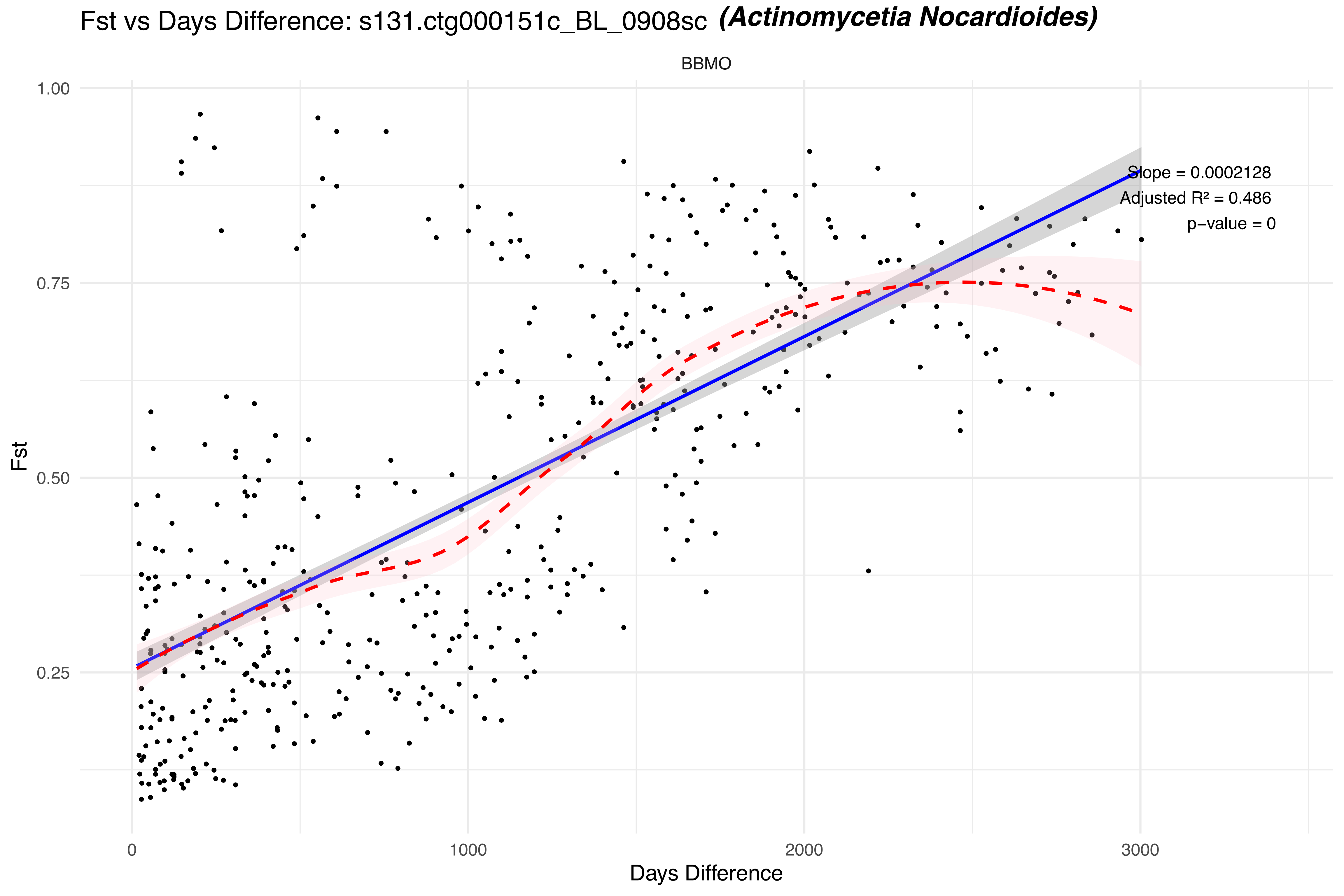

Fst vs Days Difference: s131.ctg000151c_BL_0908sc (Actinomycetia Nocardioides)
BBMO
Slope = 0.0002128
Adjusted R² = 0.486
p−value = 0
Fst
1.00
0.75
0.50
0.25
0
1000
2000
3000
Days Difference

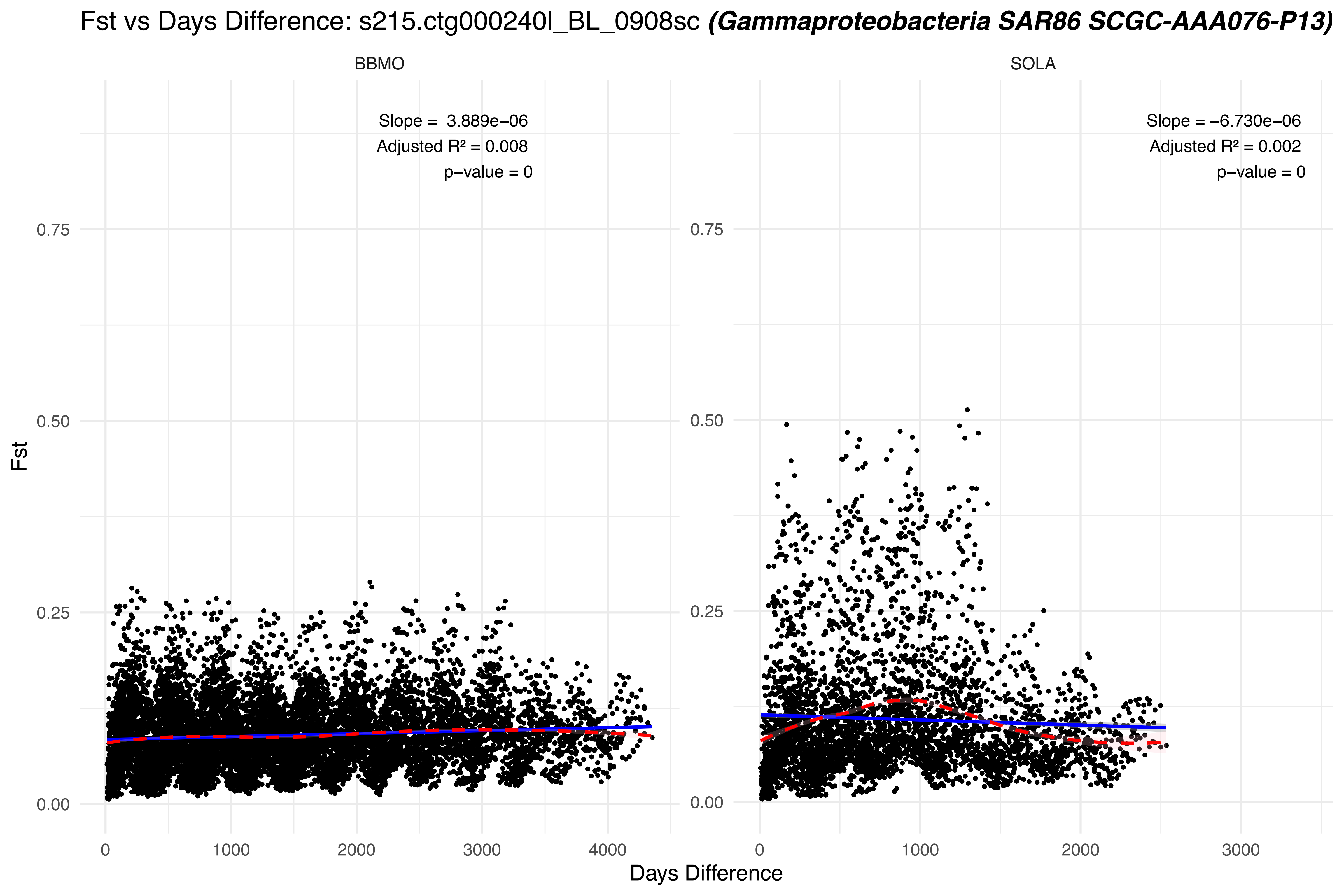

Fst vs Days Difference: s215.ctg000240l_BL_0908sc (Gammaproteobacteria SAR86 SCGC-AAA076-P13)
BBMO
SOLA
Slope = 3.889e−06
Adjusted R² = 0.008
p−value = 0
Slope = −6.730e−06
Adjusted R² = 0.002
p−value = 0
Fst
0.00
0.25
0.50
0.75
0
1000
2000
3000
4000
Days Difference

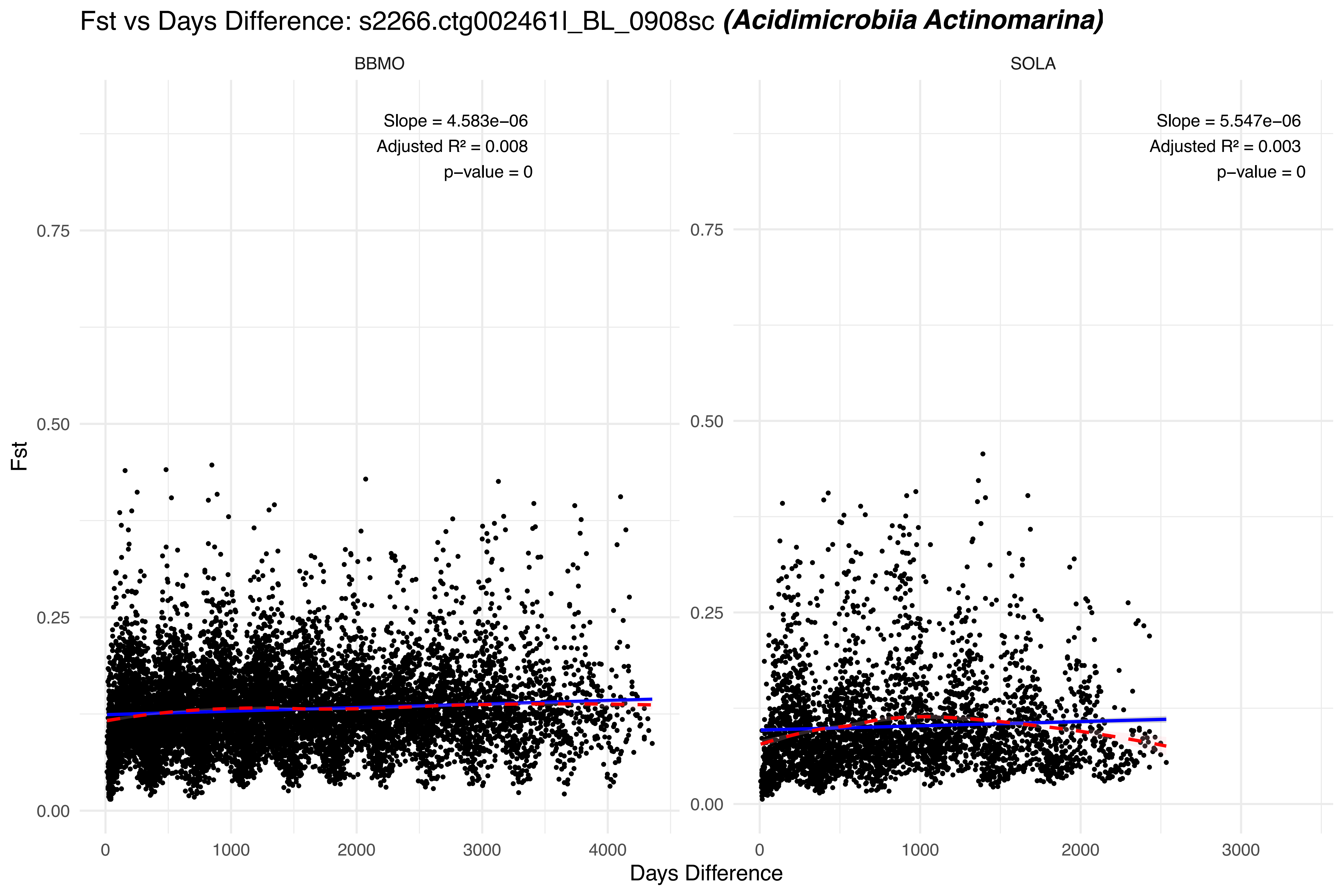
Fst vs Days Difference: s2266.ctg002461l_BL_0908sc *(Acidimicrobiia Actinomarina)*
BBMO
SOLA
Slope = 4.583e−06
Adjusted R² = 0.008
p−value = 0
Slope = 5.547e−06
Adjusted R² = 0.003
p−value = 0
Fst
0.00
0.25
0.50
0.75
0
1000
2000
3000
4000
Days Difference

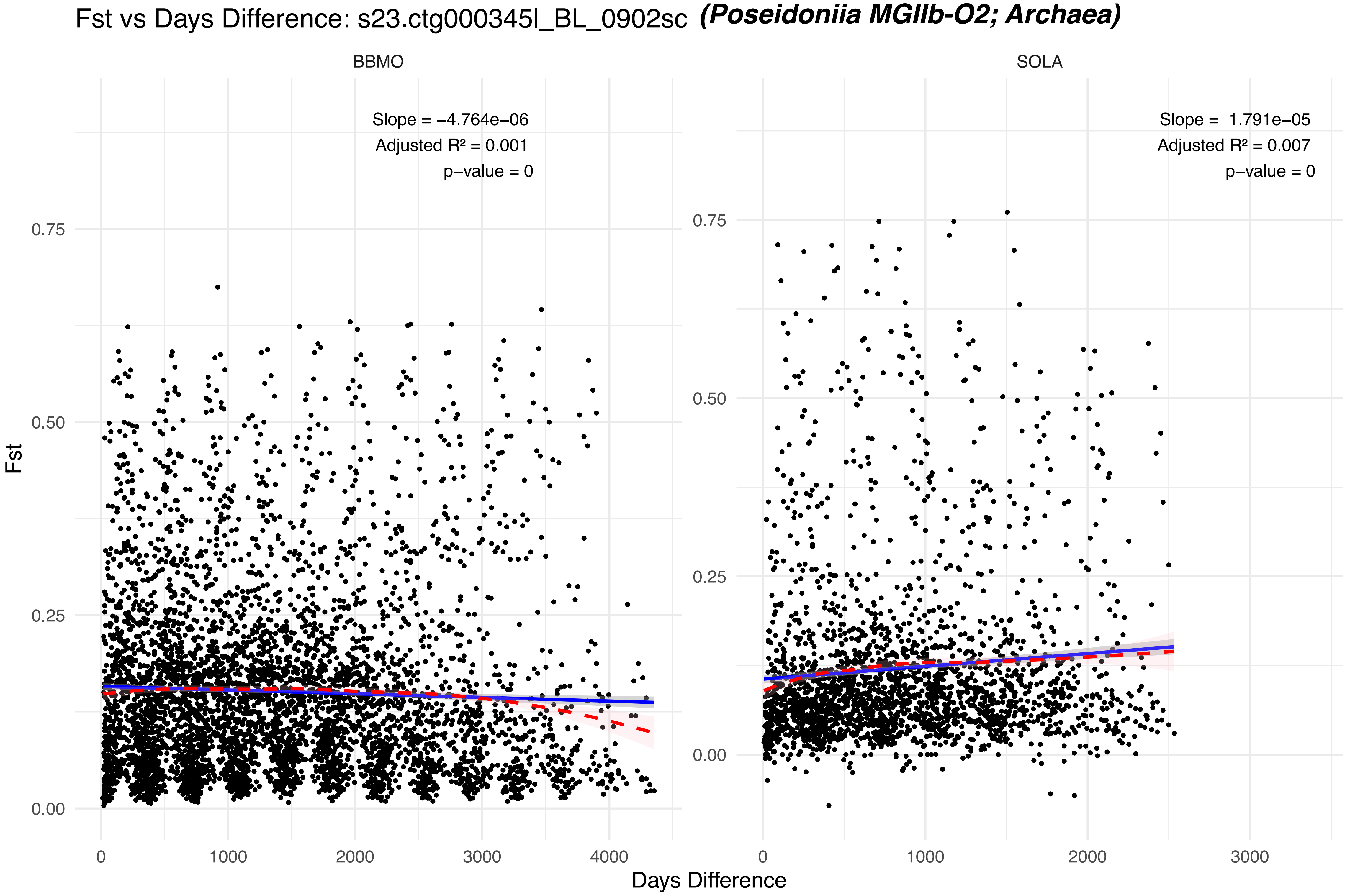

Fst vs Days Difference: s23.ctg000345l_BL_0902sc (Poseidoniia MGIIb-O2; Archaea)
BBMO
SOLA
Slope = −4.764e−06
Adjusted R² = 0.001
p−value = 0
Slope = 1.791e−05
Adjusted R² = 0.007
p−value = 0
Fst
0.00
0.25
0.50
0.75
0
1000
2000
3000
4000
Days Difference

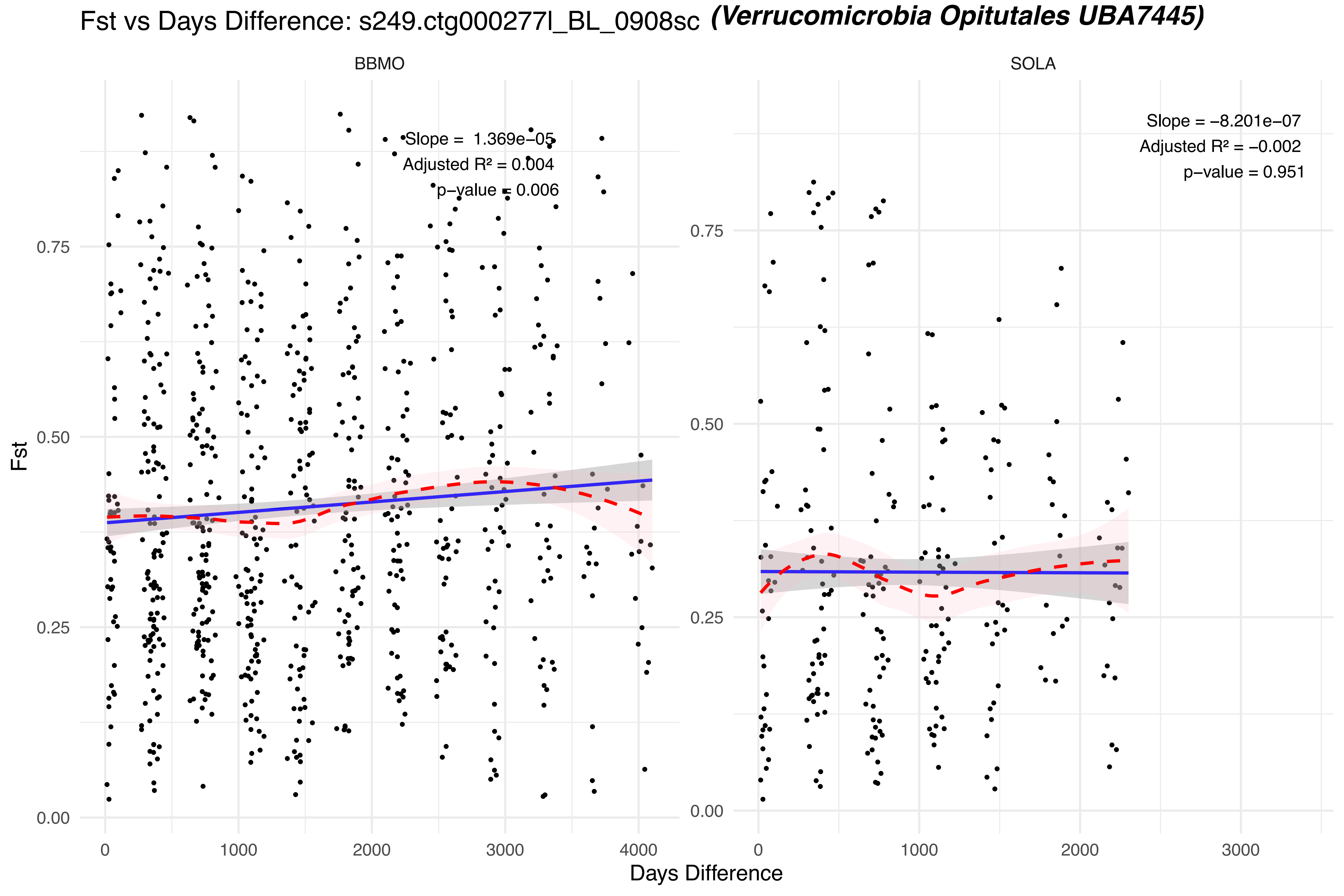

Fst vs Days Difference: s249.ctg000277l_BL_0908sc (Verrucomicrobia Opitutales UBA7445)
BBMO
SOLA
Slope = 1.369e−05
Adjusted R² = 0.004
p−value = 0.006
Slope = −8.201e−07
Adjusted R² = −0.002
p−value = 0.951
Fst
0.00
0.25
0.50
0.75
0
1000
2000
3000
4000
Days Difference

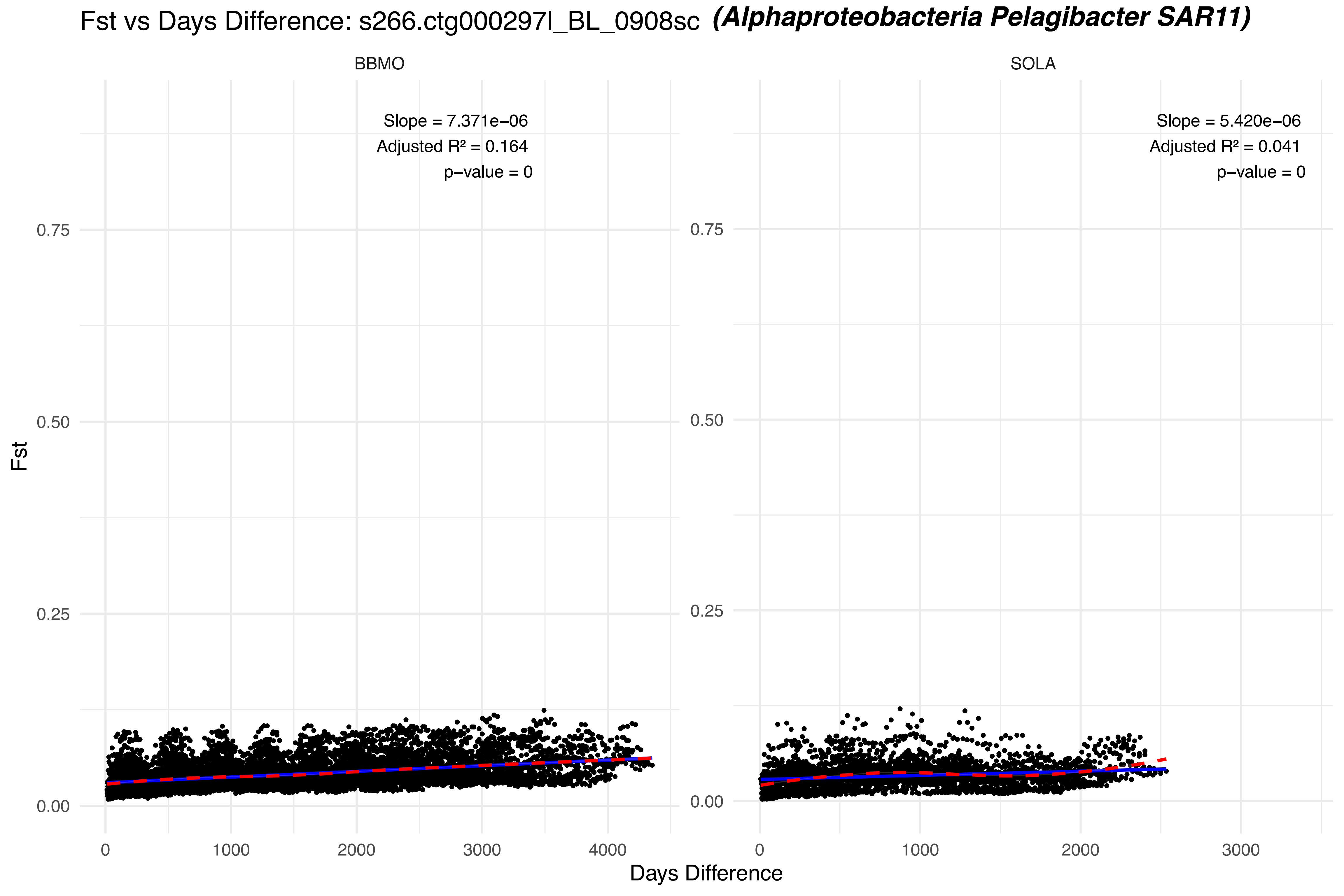
Fst vs Days Difference: s266.ctg000297l_BL_0908sc (Alphaproteobacteria Pelagibacter SAR11)
BBMO
SOLA
Slope = 7.371e−06
Adjusted R² = 0.164
p−value = 0
Slope = 5.420e−06
Adjusted R² = 0.041
p−value = 0
Fst
0.00
0.25
0.50
0.75
0
1000
2000
3000
4000
Days Difference

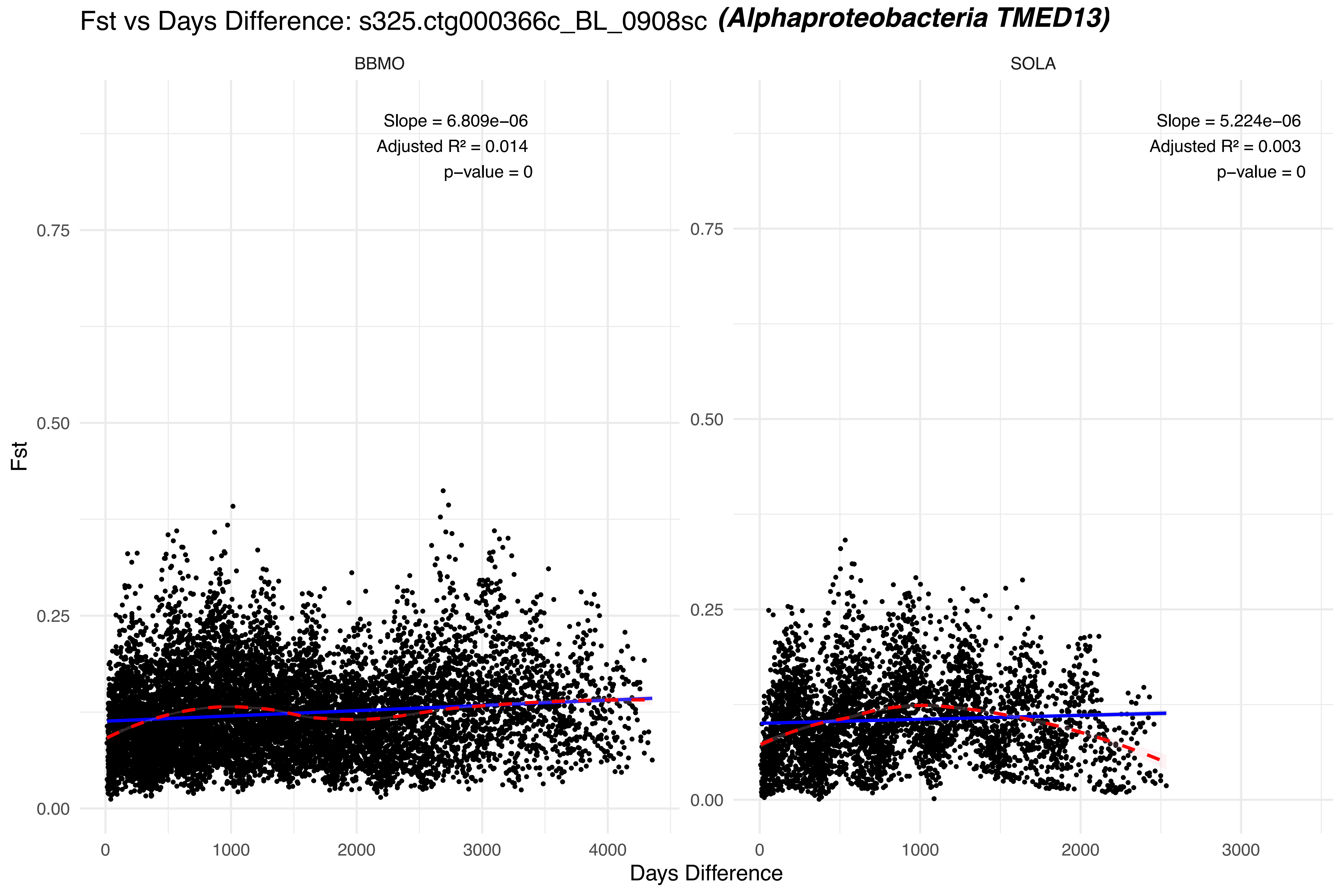

Fst vs Days Difference: s325.ctg000366c_BL_0908sc (Alphaproteobacteria TMED13)
BBMO
SOLA
Slope = 6.809e−06
Adjusted R² = 0.014
p−value = 0
Slope = 5.224e−06
Adjusted R² = 0.003
p−value = 0
Fst
0.00
0.25
0.50
0.75
0
1000
2000
3000
4000
Days Difference

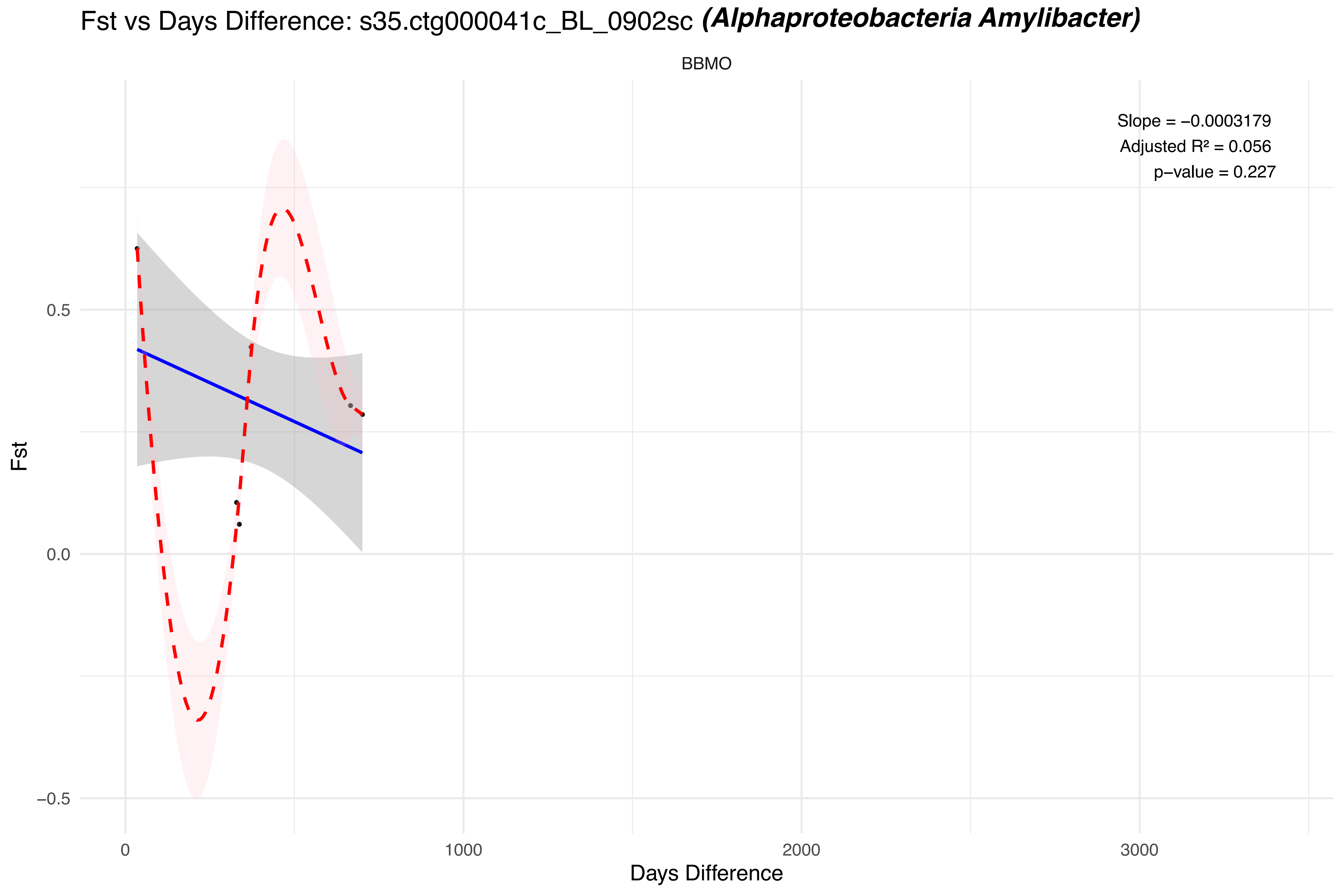
Fst vs Days Difference: s35.ctg000041c_BL_0902sc (Alphaproteobacteria Amylibacter)
BBMO
Slope = −0.0003179
Adjusted R² = 0.056
p−value = 0.227
Fst
0.5
0.0
−0.5
0
1000
2000
3000
Days Difference

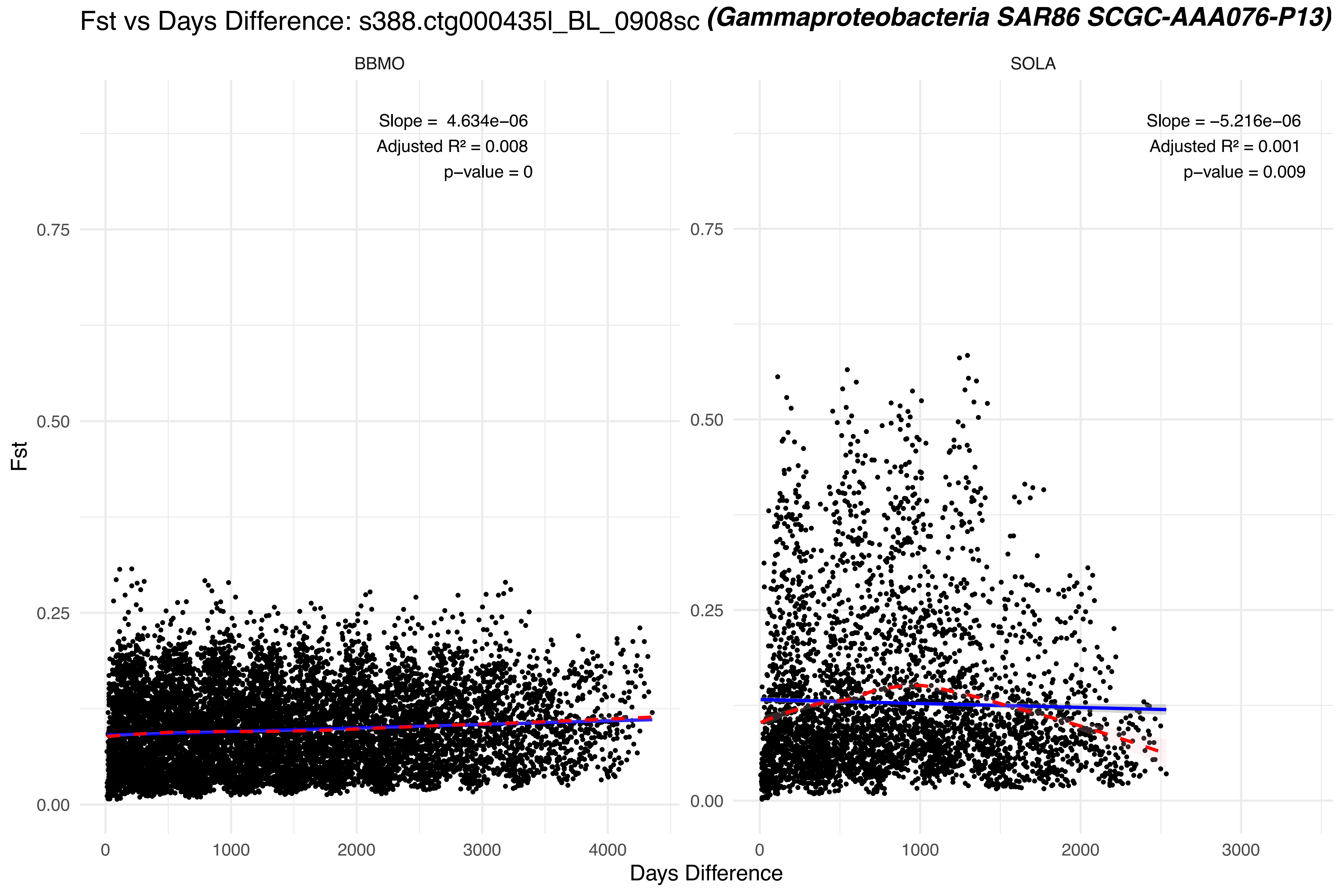

Fst vs Days Difference: s388.ctg000435l_BL_0908sc (Gammaproteobacteria SAR86 SCGC-AAA076-P13)
BBMO
Slope = 4.634e−06
Adjusted R² = 0.008
p−value = 0
SOLA
Slope = −5.216e−06
Adjusted R² = 0.001
p−value = 0.009
Fst
0.00
0.25
0.50
0.75
0
1000
2000
3000
4000
Days Difference

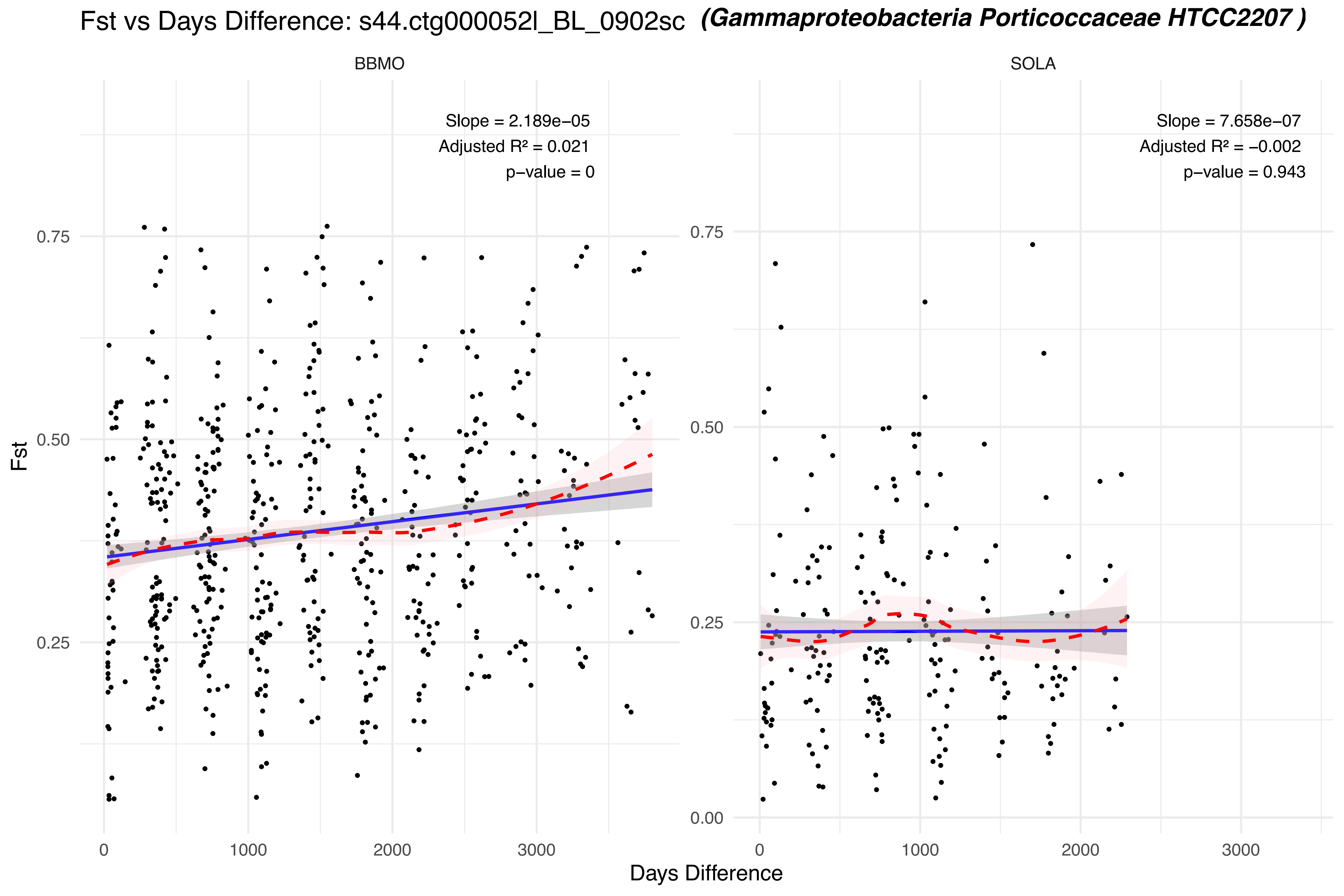

Fst vs Days Difference: s44.ctg000052l_BL_0902sc (Gammaproteobacteria Porticoccaceae HTCC2207 )
BBMO
SOLA
Slope = 2.189e−05
Adjusted R² = 0.021
p−value = 0
Slope = 7.658e−07
Adjusted R² = −0.002
p−value = 0.943
Fst
0.75
0.50
0.25
0.00
0
1000
2000
3000
Days Difference

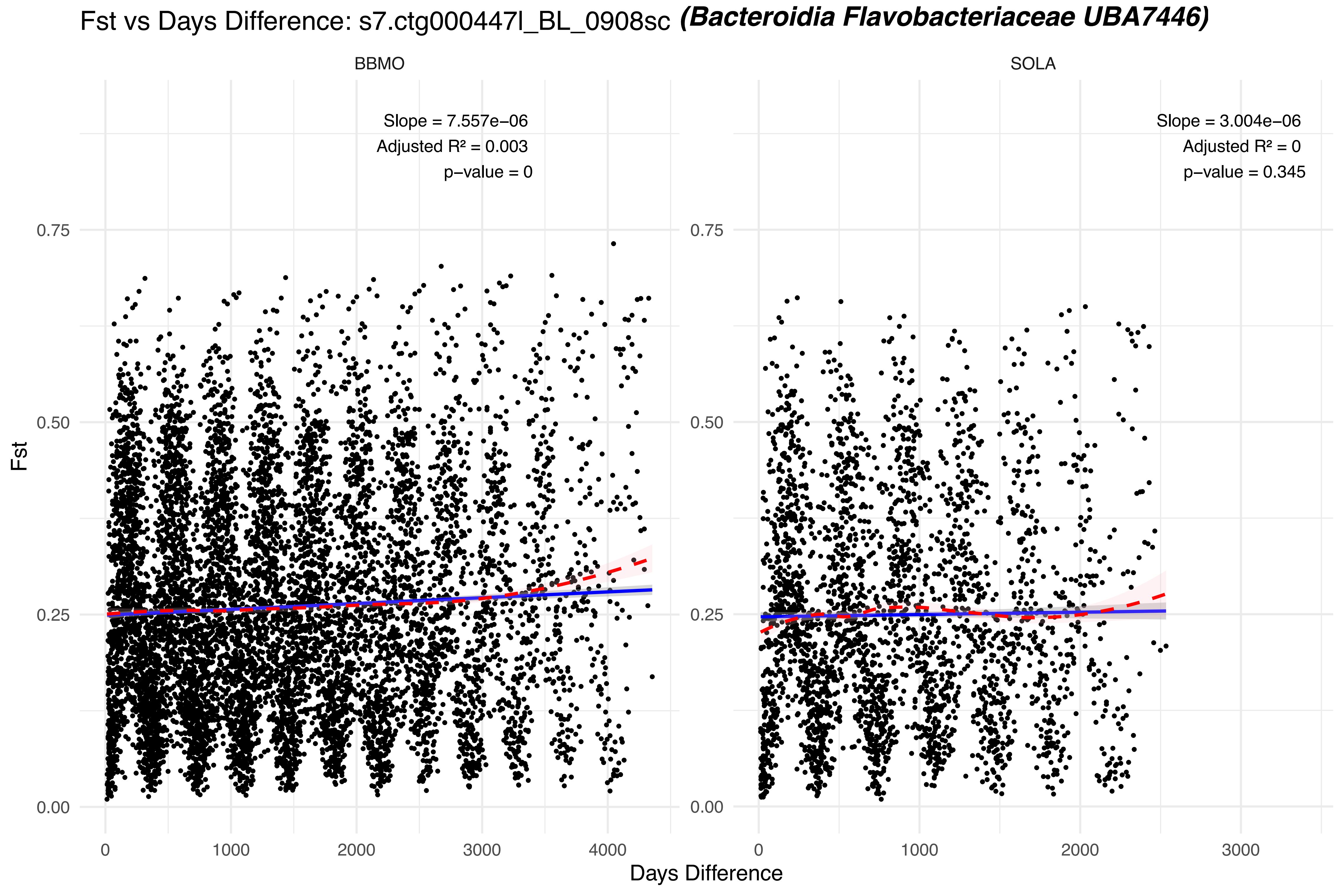

Fst vs Days Difference: s7.ctg000447l_BL_0908sc (Bacteroidia Flavobacteriaceae UBA7446)
BBMO
SOLA
Slope = 7.557e−06
Adjusted R² = 0.003
p−value = 0
Slope = 3.004e−06
Adjusted R² = 0
p−value = 0.345
Fst
0.00
0.25
0.50
0.75
0
1000
2000
3000
4000
Days Difference

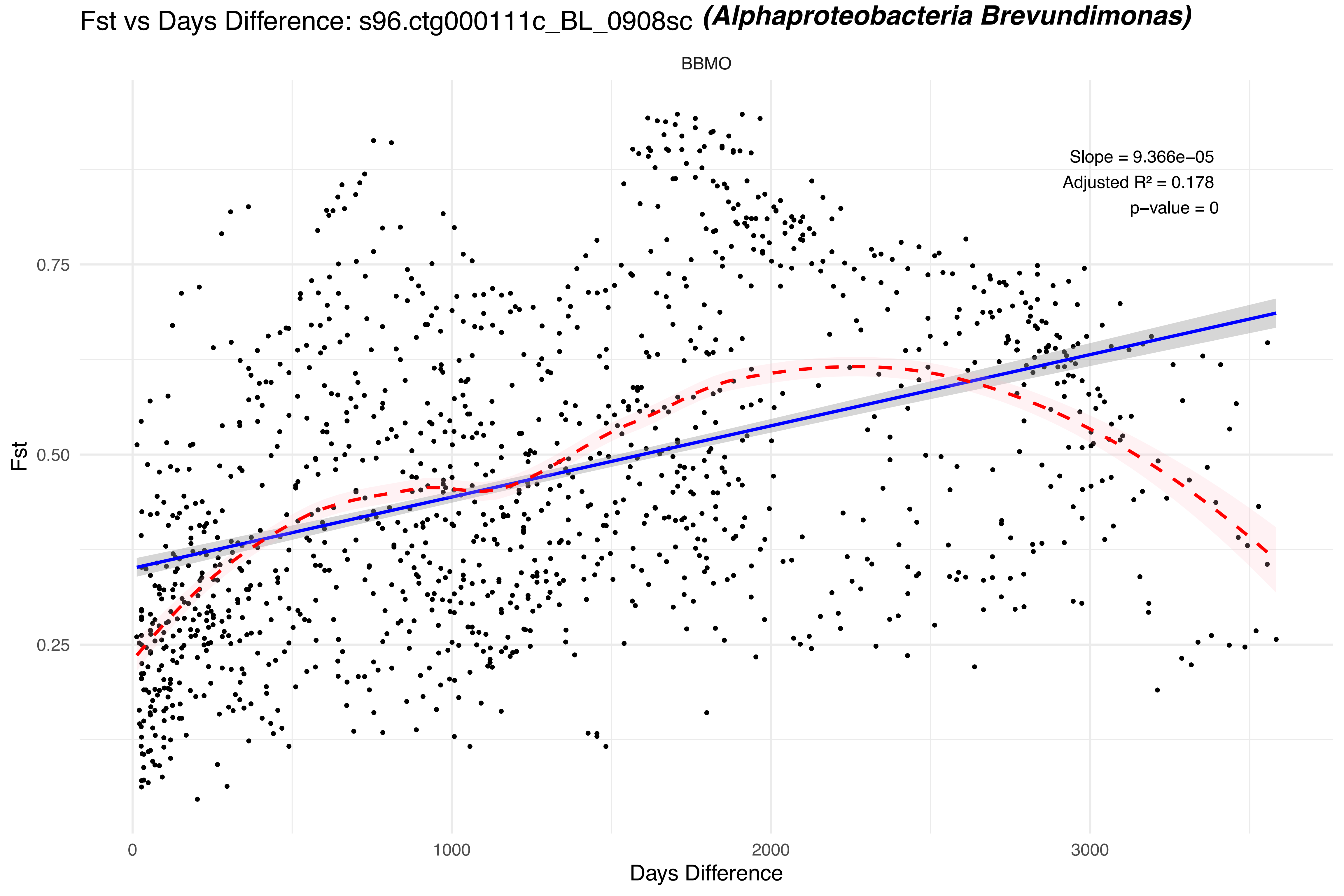
Fst vs Days Difference: s96.ctg000111c_BL_0908sc (Alphaproteobacteria Brevundimonas)
BBMO
Slope = 9.366e−05
Adjusted R² = 0.178
p−value = 0
Fst
0.25
0.50
0.75
0
1000
2000
3000
Days Difference
