## Supplementary Figure 11 for "Seasonal population structure and adaptive signatures across long-term marine microbial time series"

**Supplementary Figure 11. Temporal dynamics of pN/pS for candidate adaptive genes across the BBMO and SOLA time series.** For each MAG, temporal profiles of pN/pS are shown separately for BBMO and SOLA. Each panel represents one candidate adaptive gene, identified by its gene ID (top-left corner), with samples arranged in chronological sampling order. The x-axis indicates successive sampling months (1–140 for BBMO and 1–90 for SOLA), and the y-axis shows pN/pS. Samples with <25% horizontal coverage of the corresponding MAG were excluded. For visualization, pN/pS values >2 were capped at 2, whereas undefined values resulting from pS = 0 were plotted as −1.

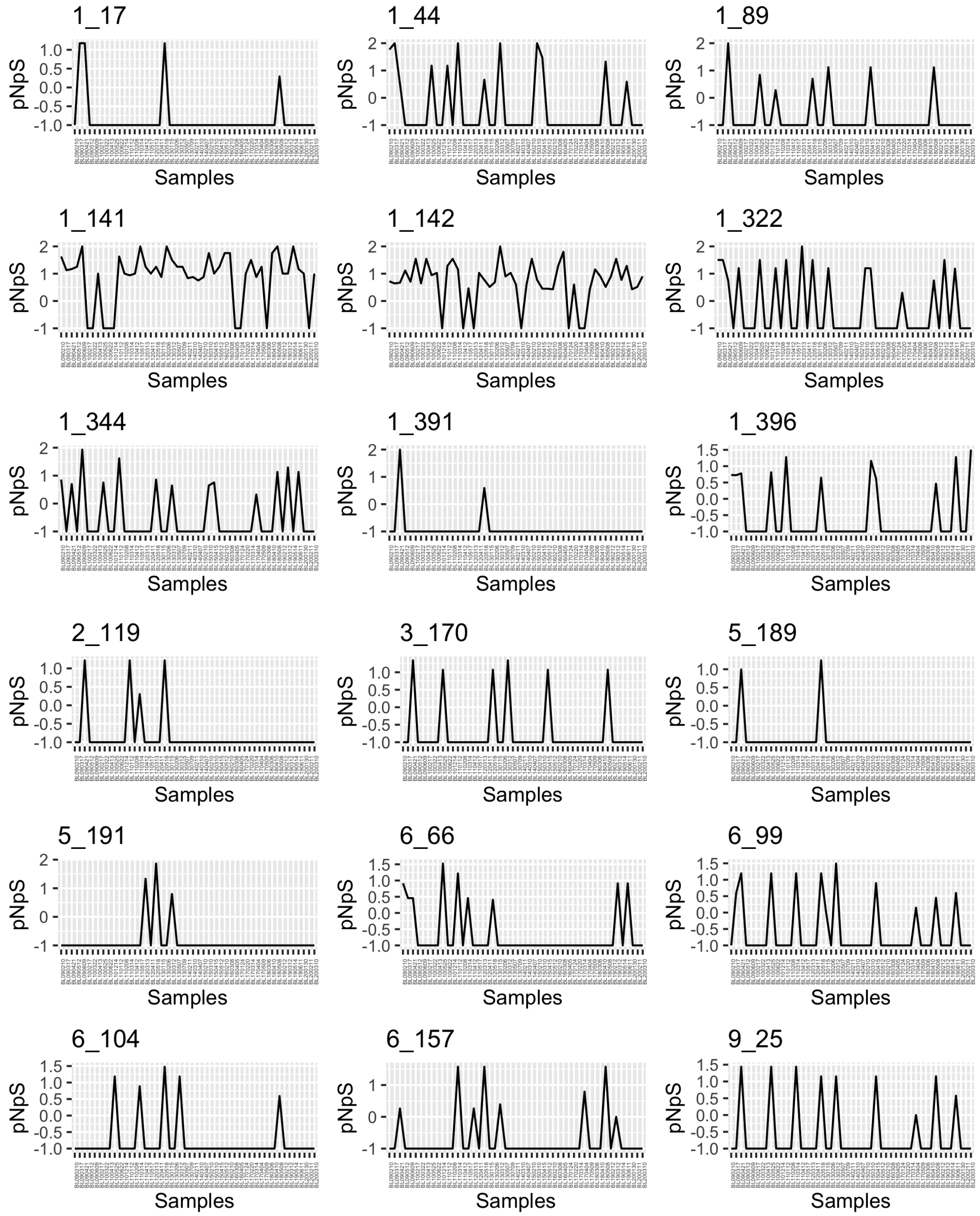


*Polaribacter* MAG (BL_0902_bin.full.136) in BBMO

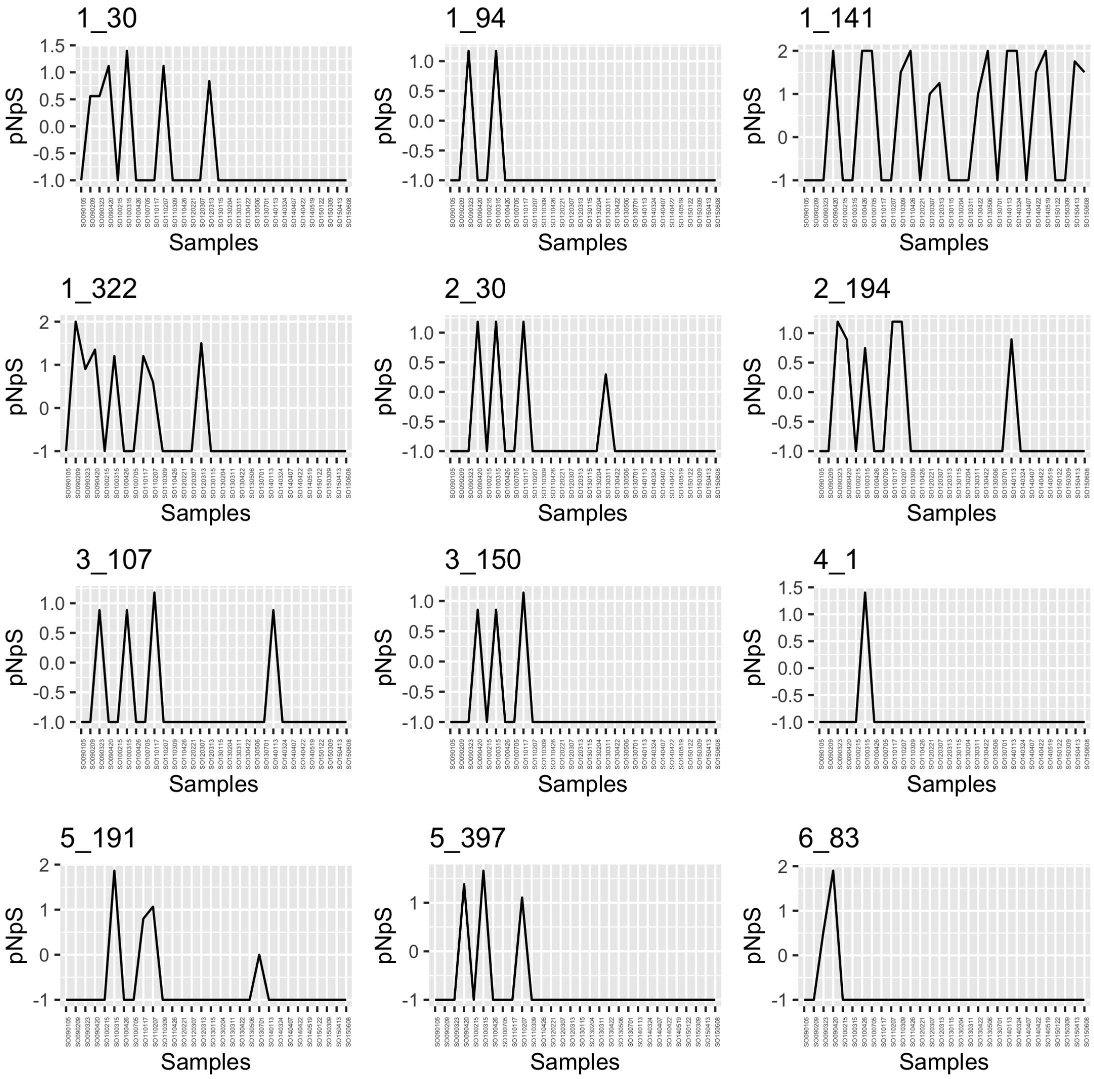


*Polaribacter* MAG (BL_0902_bin.full.136) in SOLA.

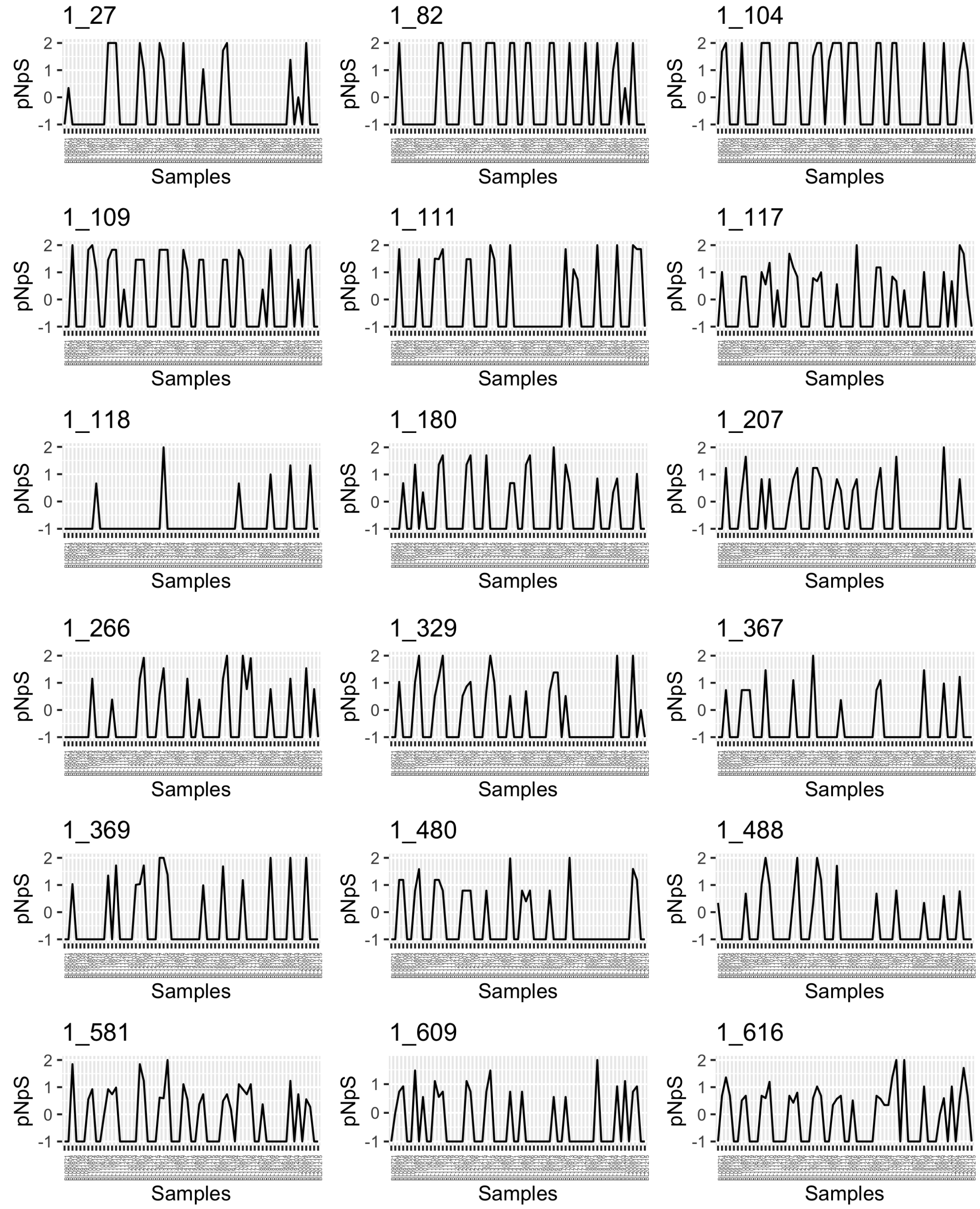


UBA8653 Phycisphaerae MAG (BL_0908_bin.full.20) in BBMO.

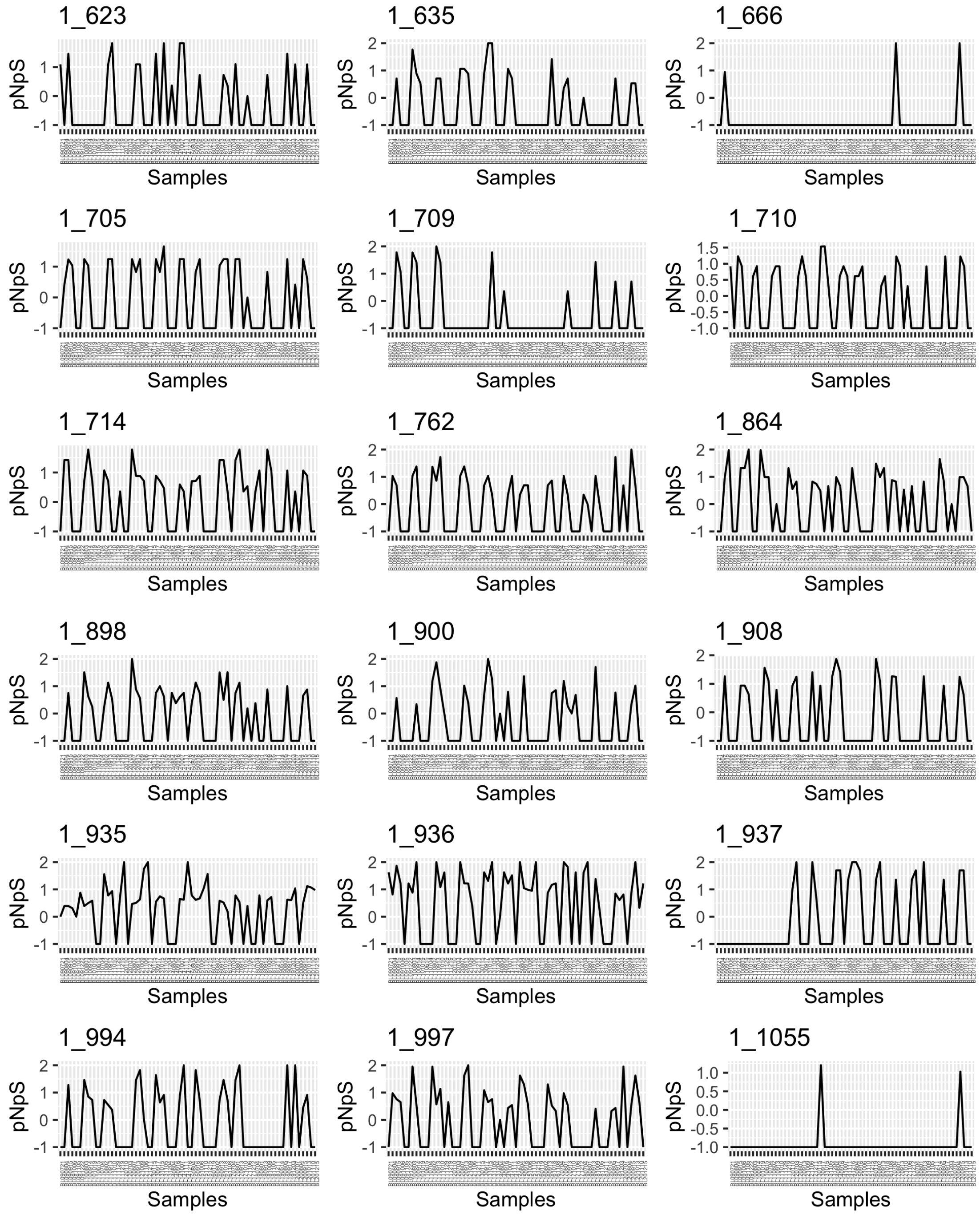


UBA8653 Phycisphaerae MAG (BL_0908_bin.full.20) in BBMO.

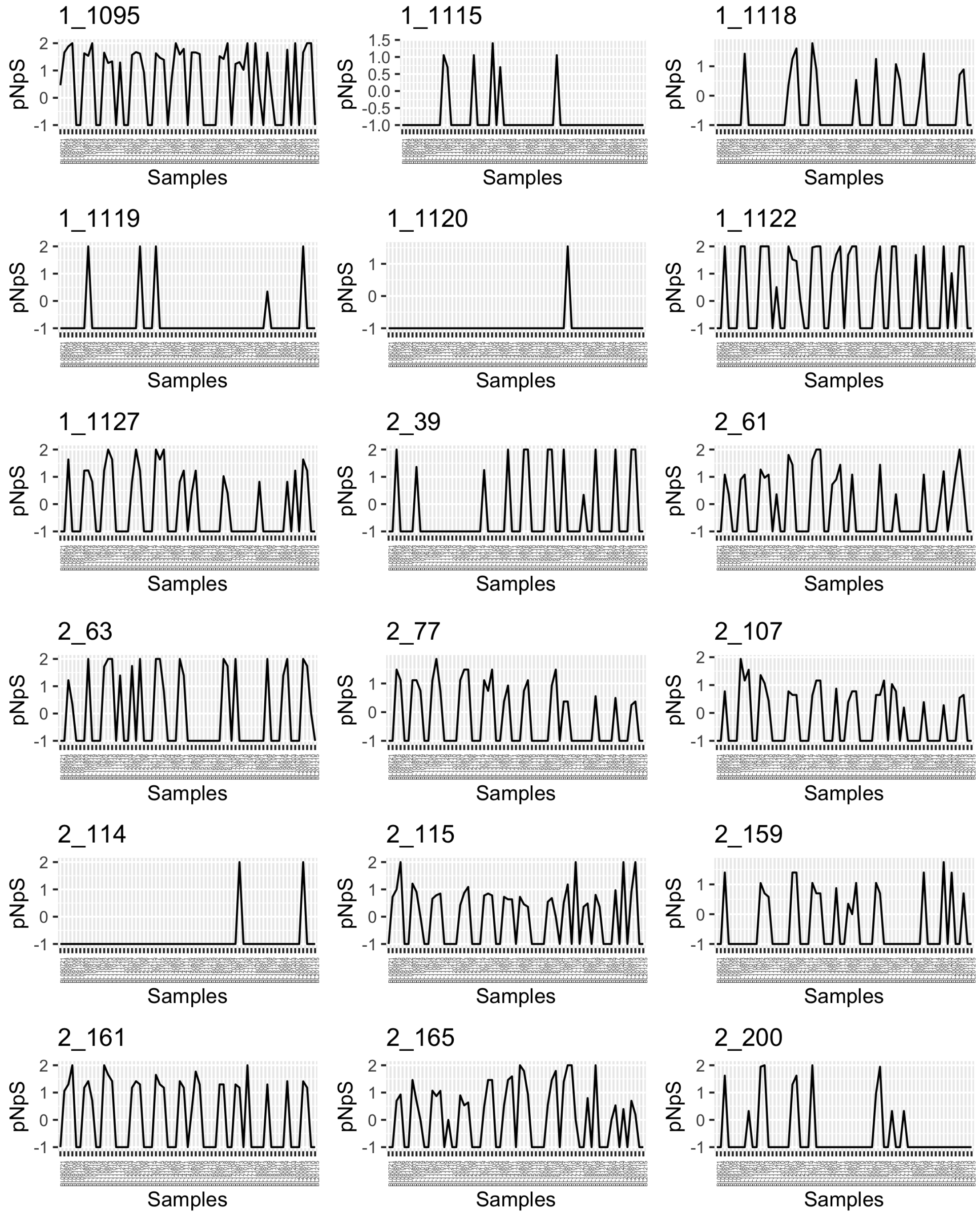


UBA8653 Phycisphaerae MAG (BL_0908_bin.full.20) in BBMO.

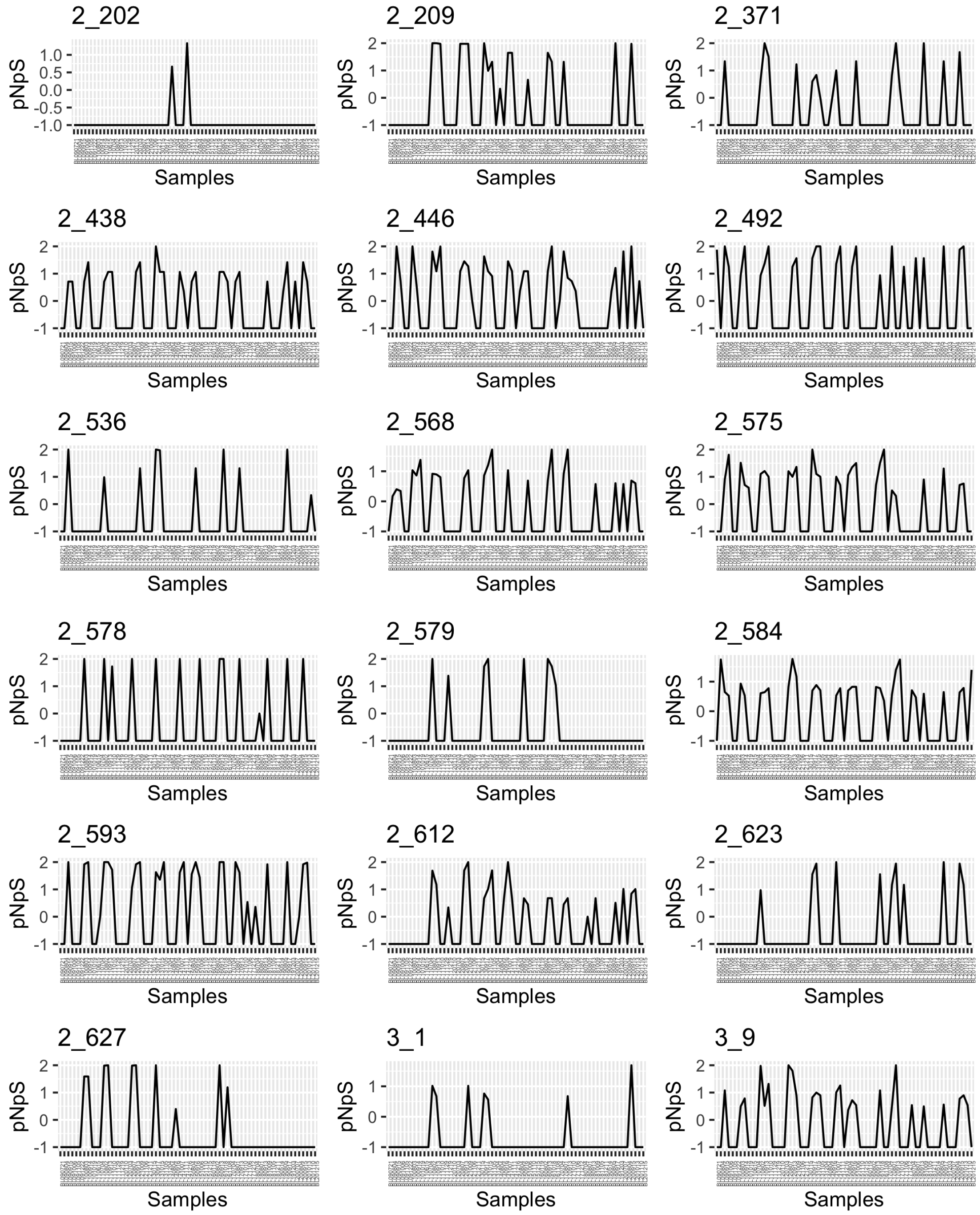


UBA8653 Phycisphaerae MAG (BL_0908_bin.full.20) in BBMO.

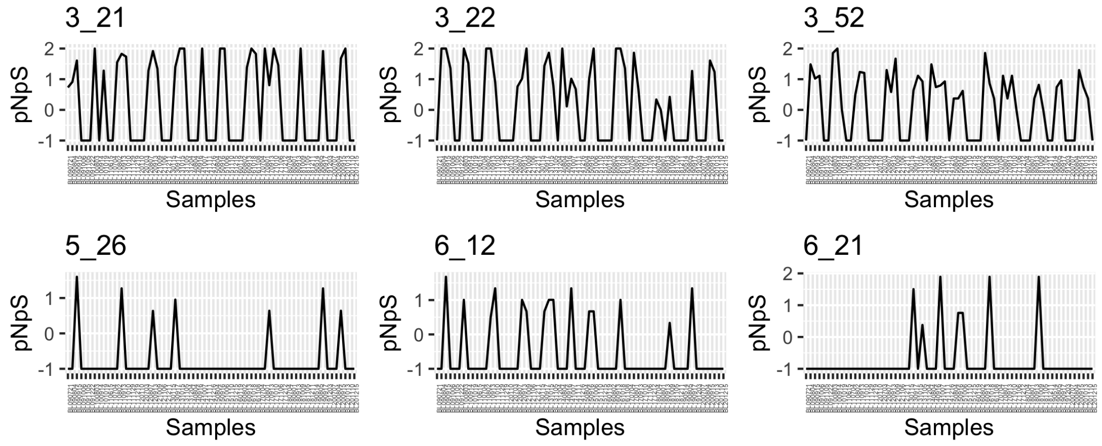


UBA8653 Phycisphaerae MAG (BL_0908_bin.full.20) in BBMO.

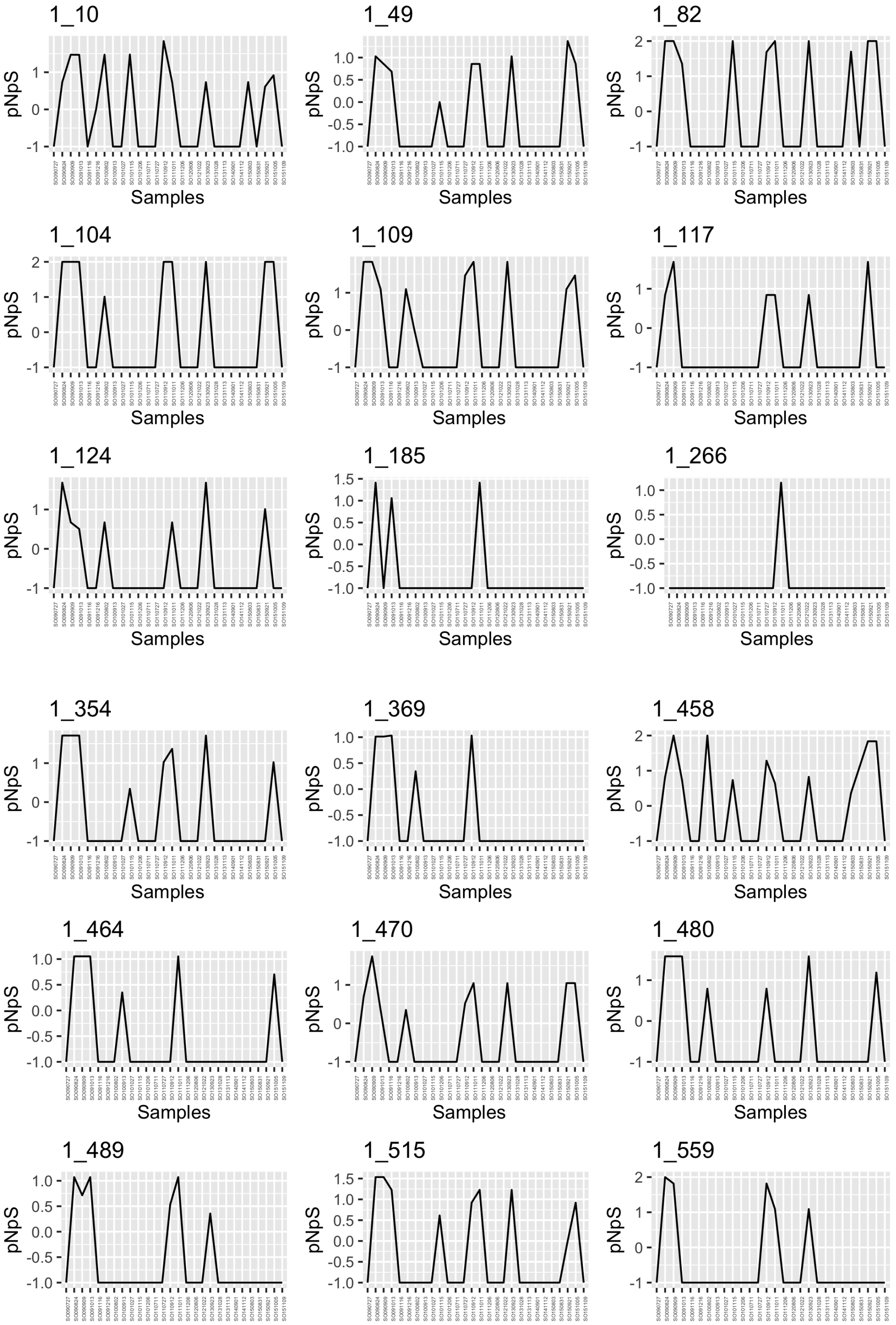


UBA8653 Phycisphaerae MAG (BL_0908_bin.full.20) in SOLA.

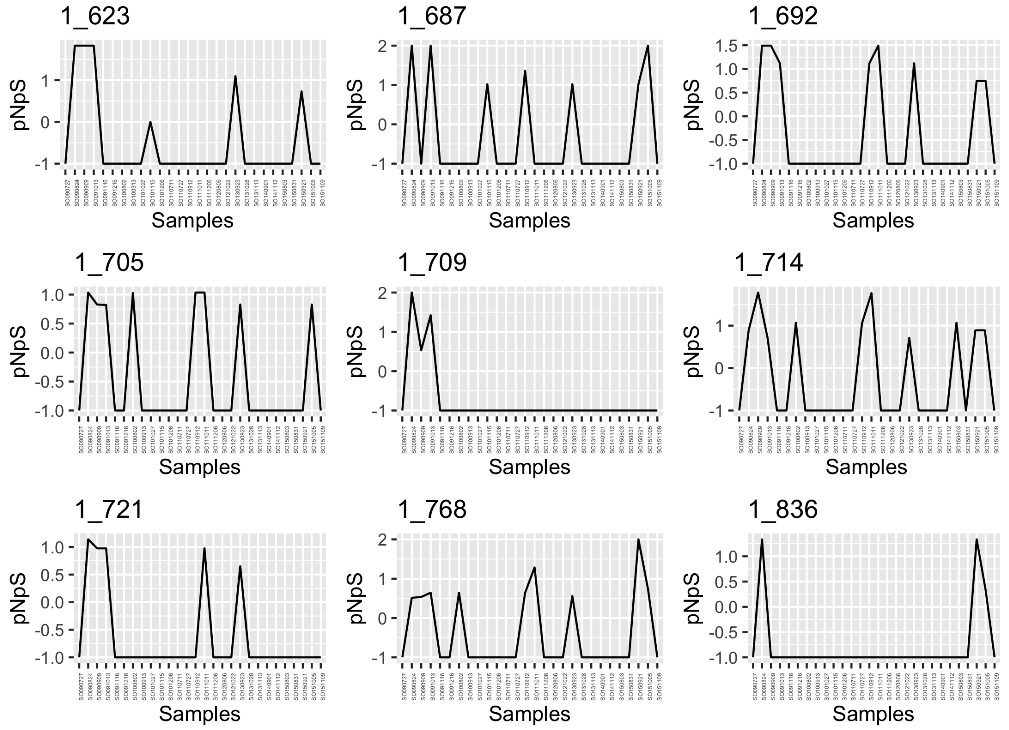


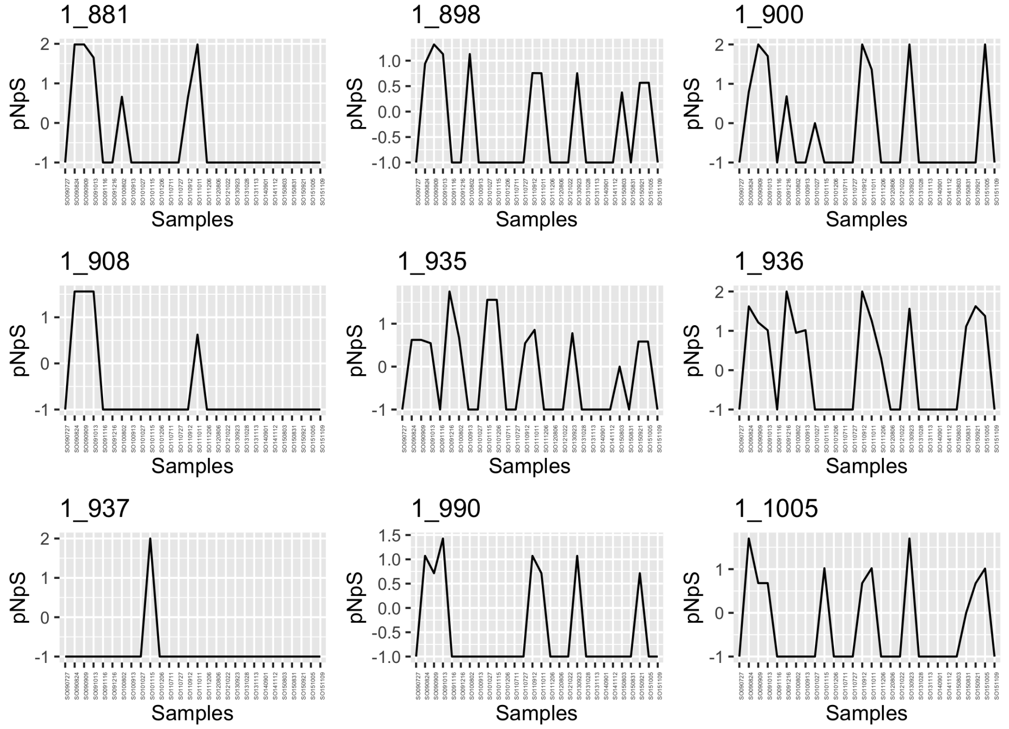


UBA8653 Phycisphaerae MAG (BL_0908_bin.full.20) in SOLA.

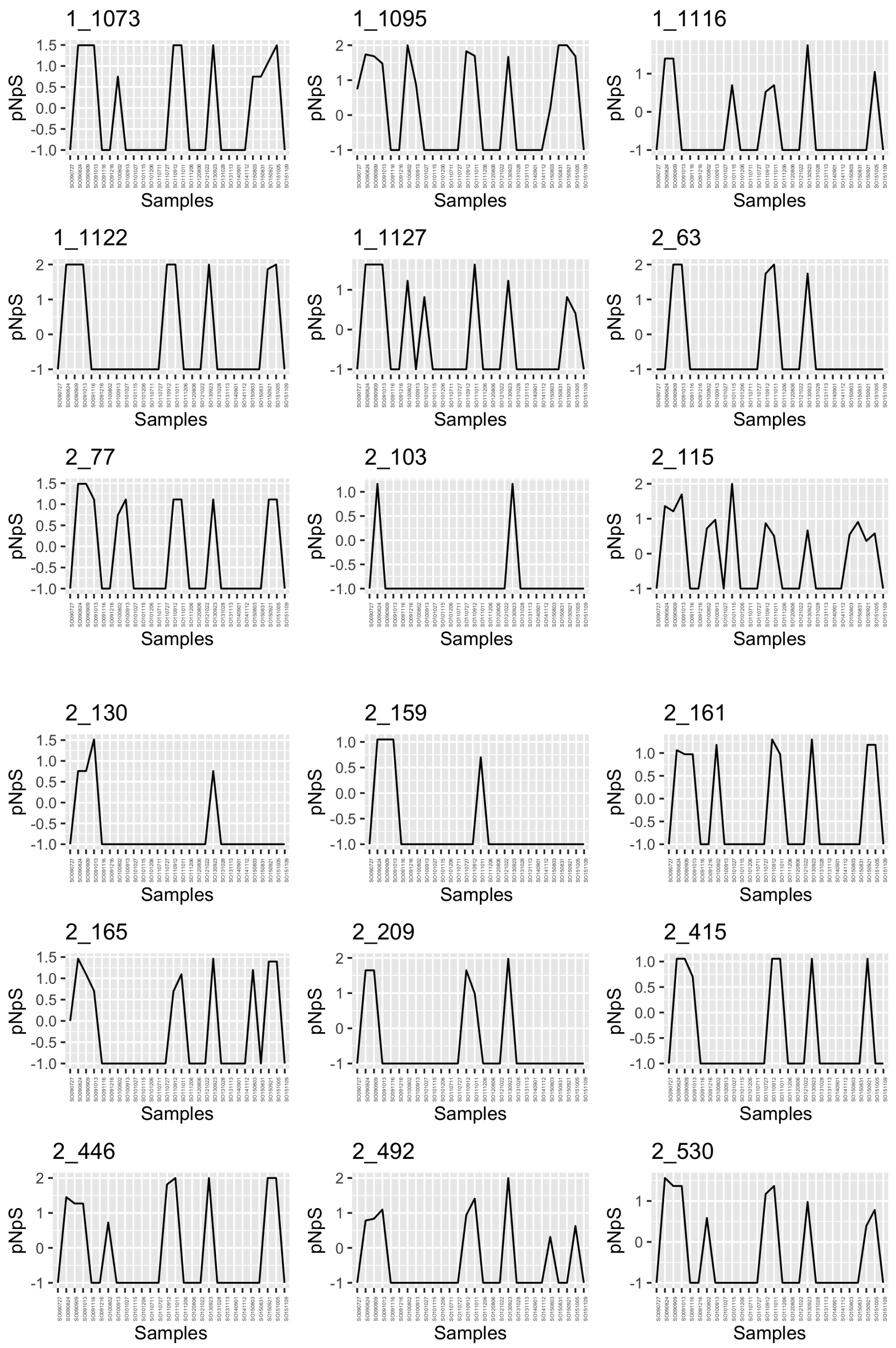


UBA8653 Phycisphaerae MAG (BL_0908_bin.full.20) in SOLA.

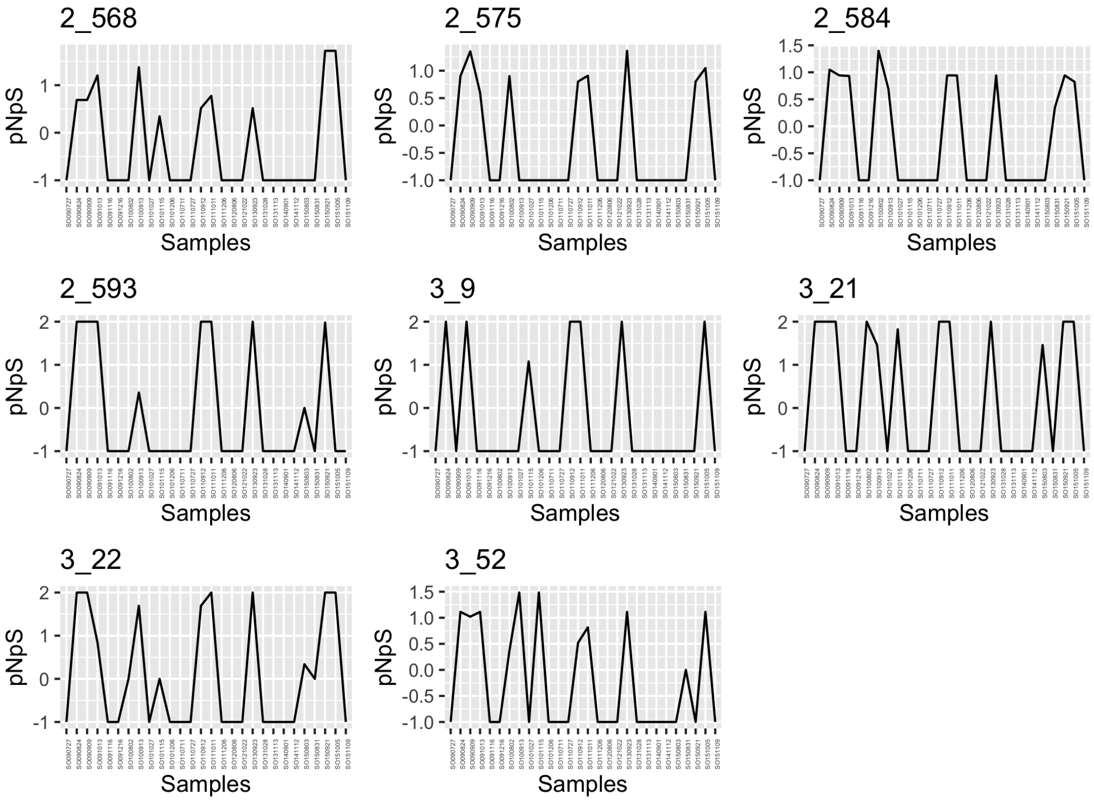

UBA8653 Phycisphaerae MAG (BL_0908_bin.full.20) in SOLA.

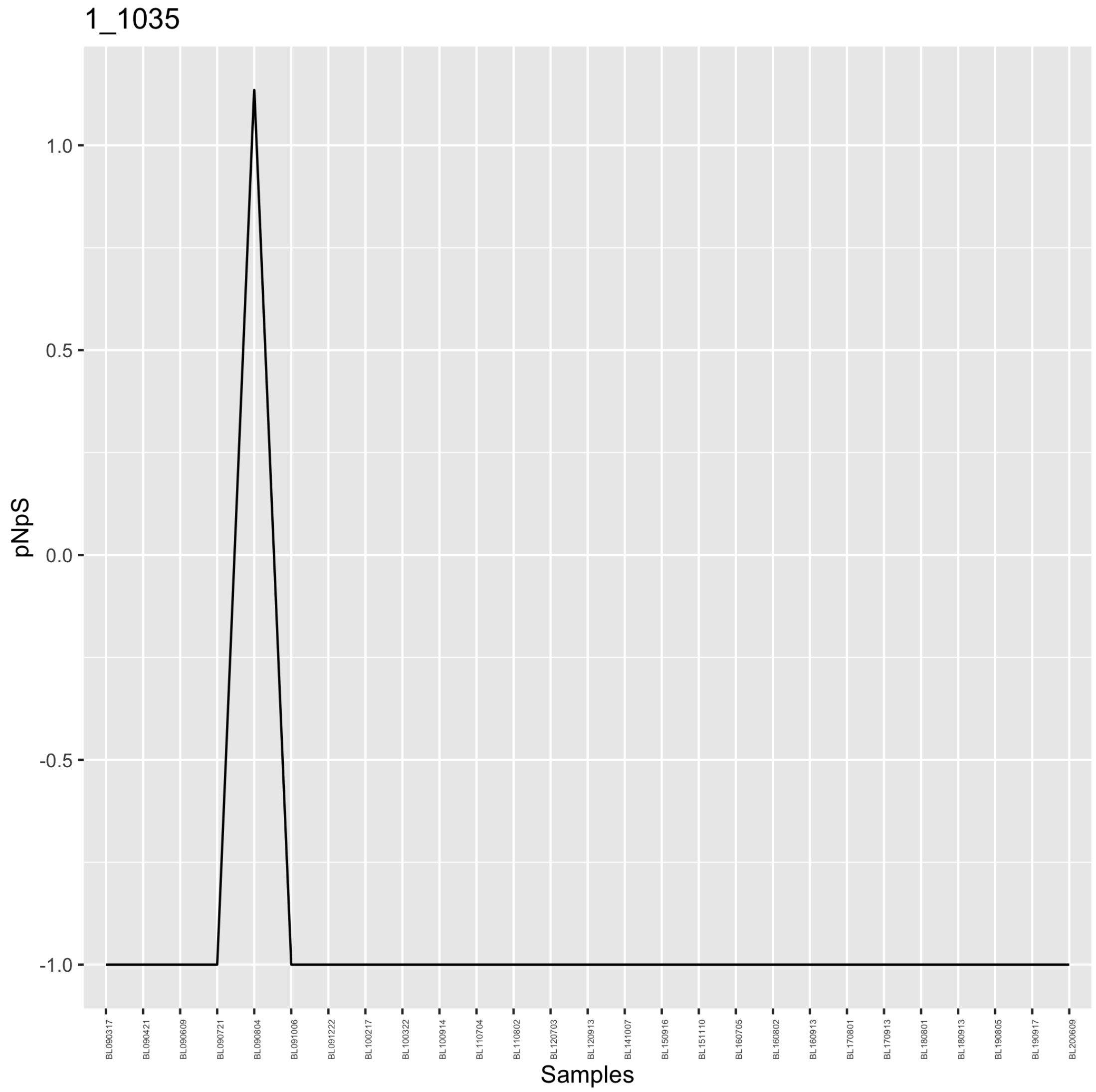


*Alteromonas* MAG (BL_0908_bin.full.418) in BBMO.

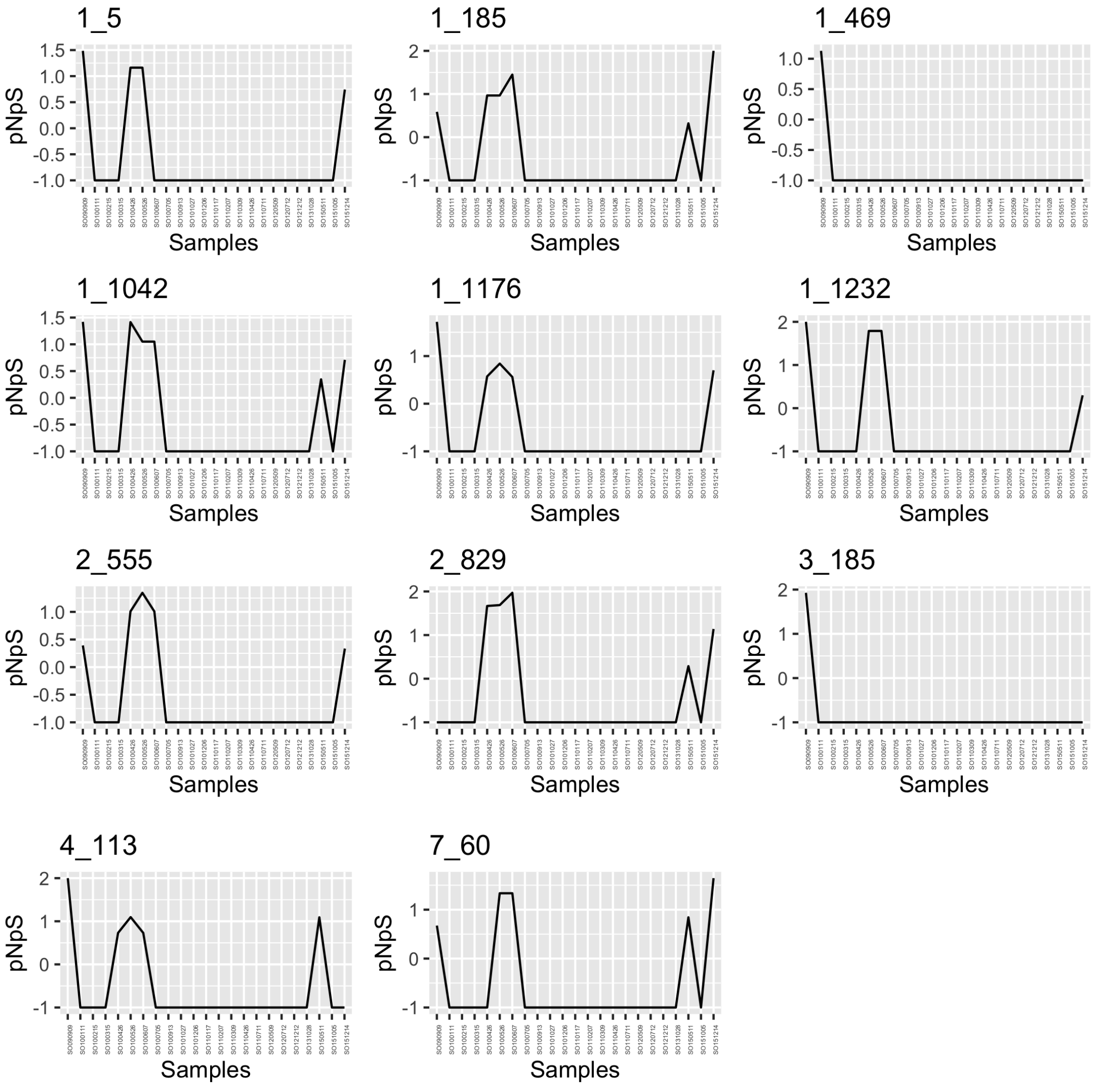


*Alteromonas* MAG (BL_0908_bin.full.418) in SOLA.

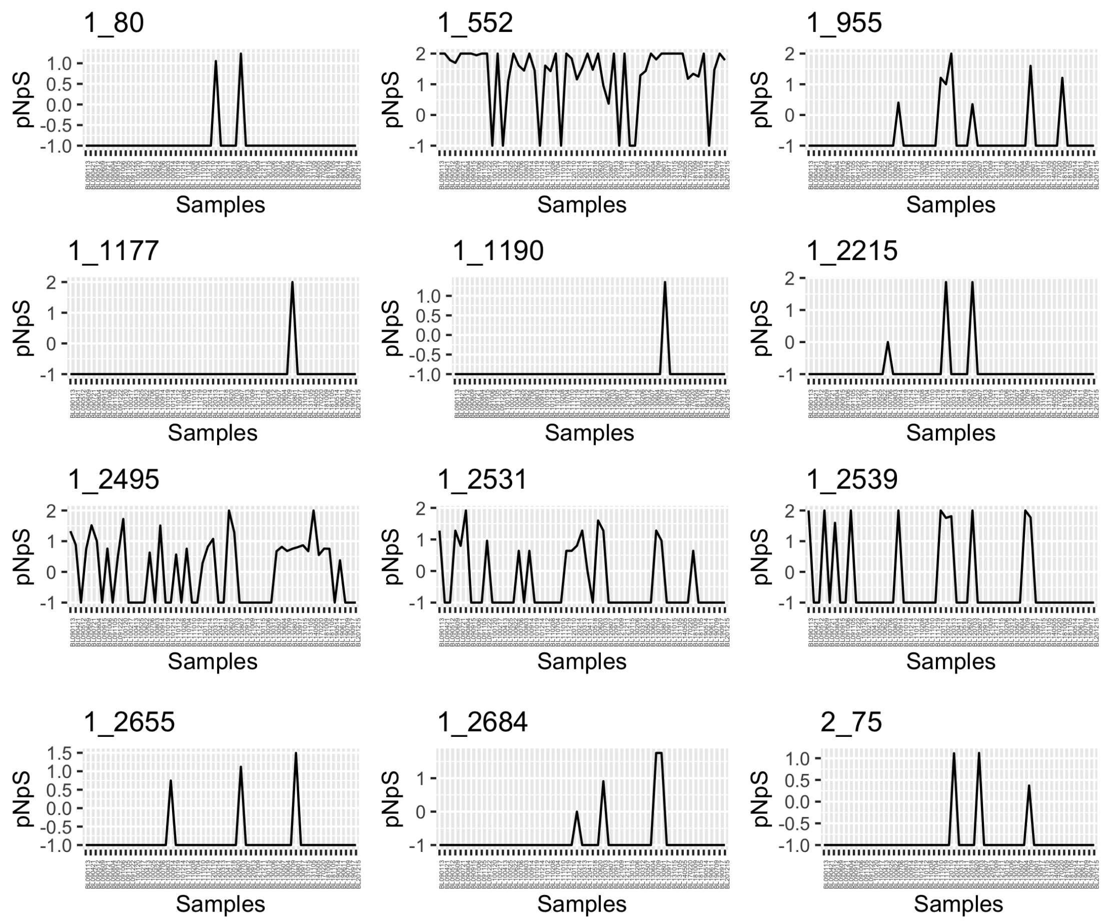


*Alteriqipengyuania* MAG (BL_0908_bin.full.436) in BBMO.

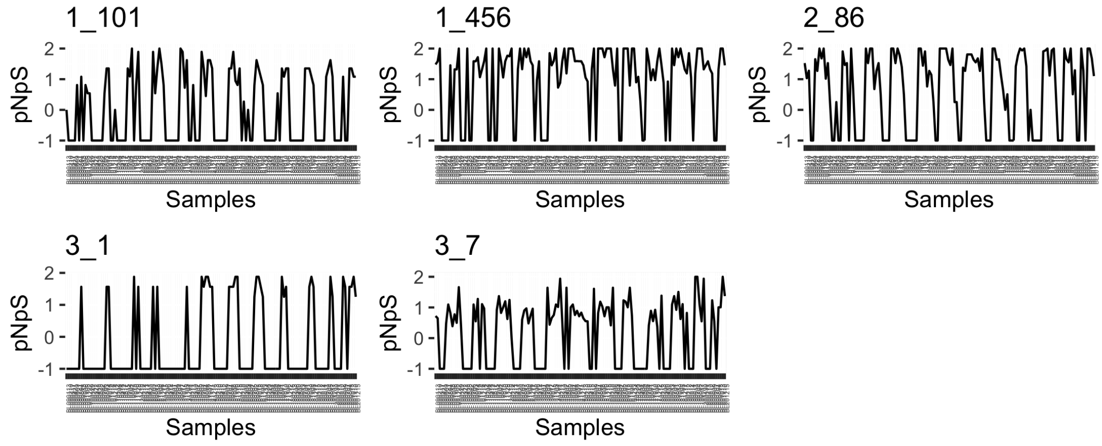


*Marinisoma* MAG (BL_0908_bin.full.522) in BBMO.

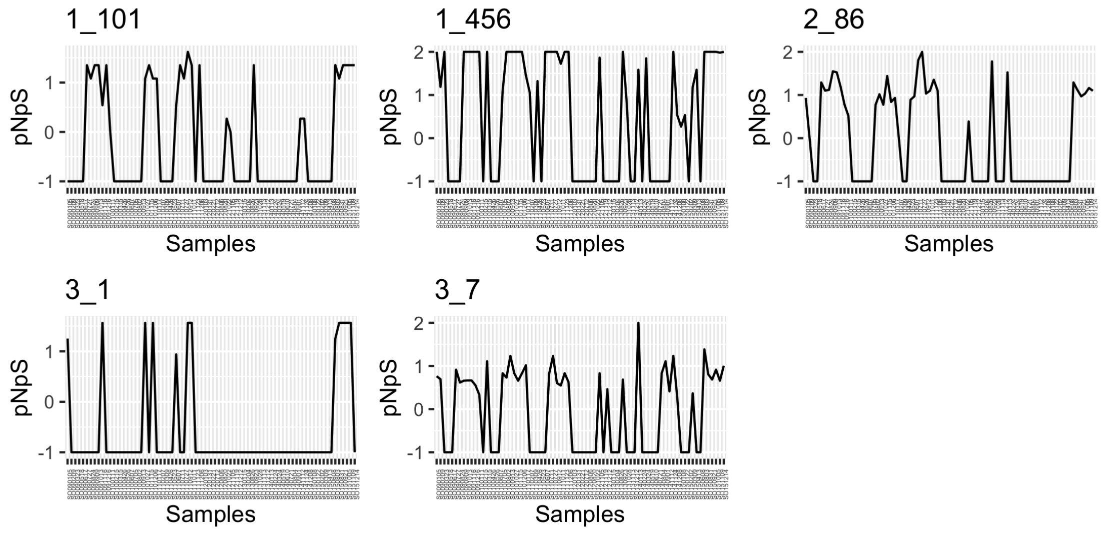


*Marinisoma* MAG (BL_0908_bin.full.522) in SOLA.

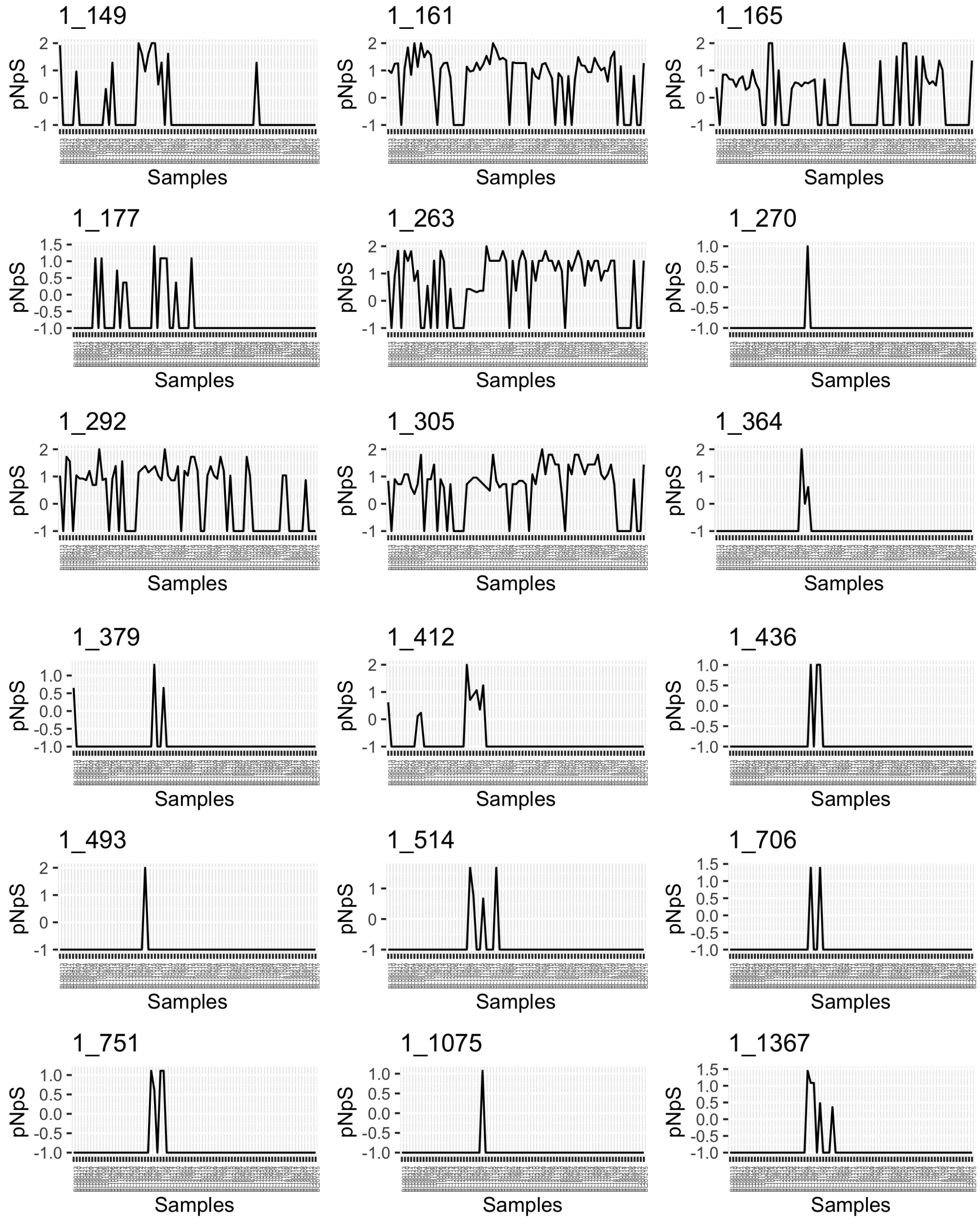


*Erythrobacter* MAG (BL_0908_bin.full.559) in BBMO.

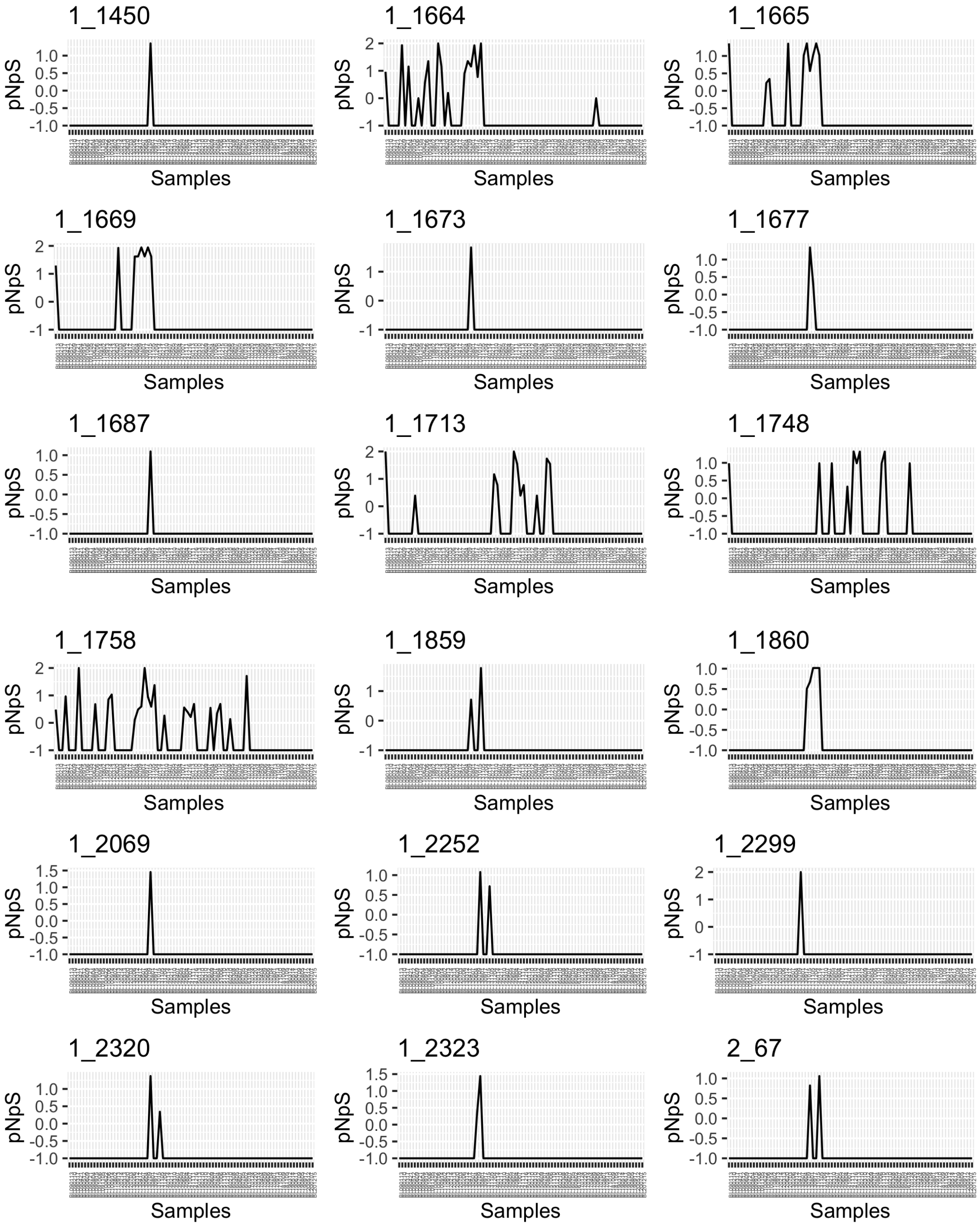


*Erythrobacter* MAG (BL_0908_bin.full.559) in BBMO.

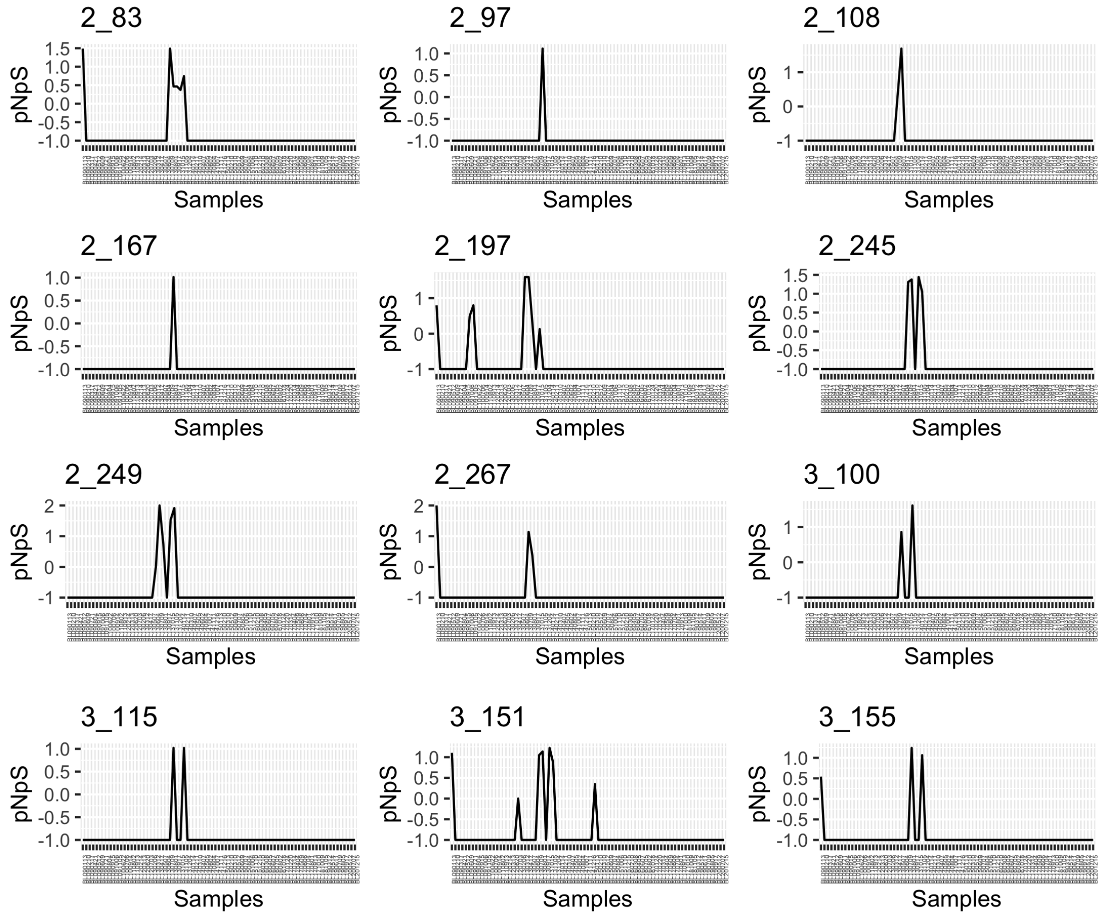


*Erythrobacter* MAG (BL_0908_bin.full.559) in BBMO.

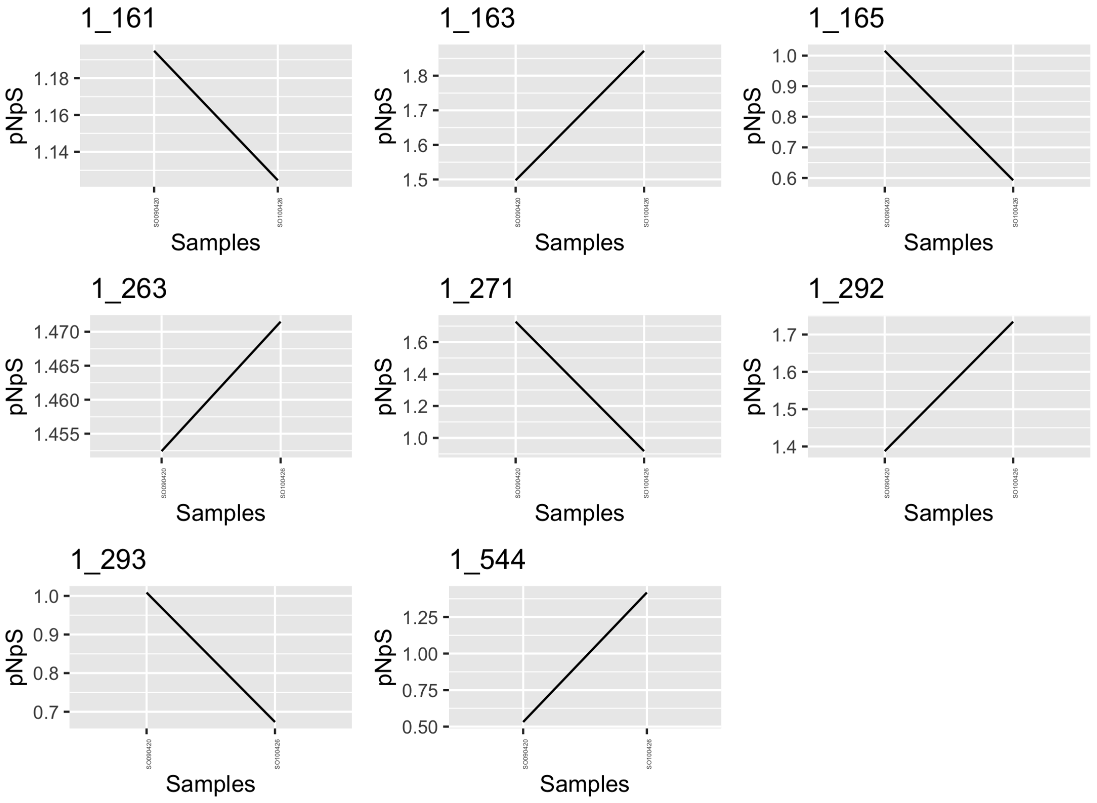


*Erythrobacter* MAG (BL_0908_bin.full.559) in SOLA.

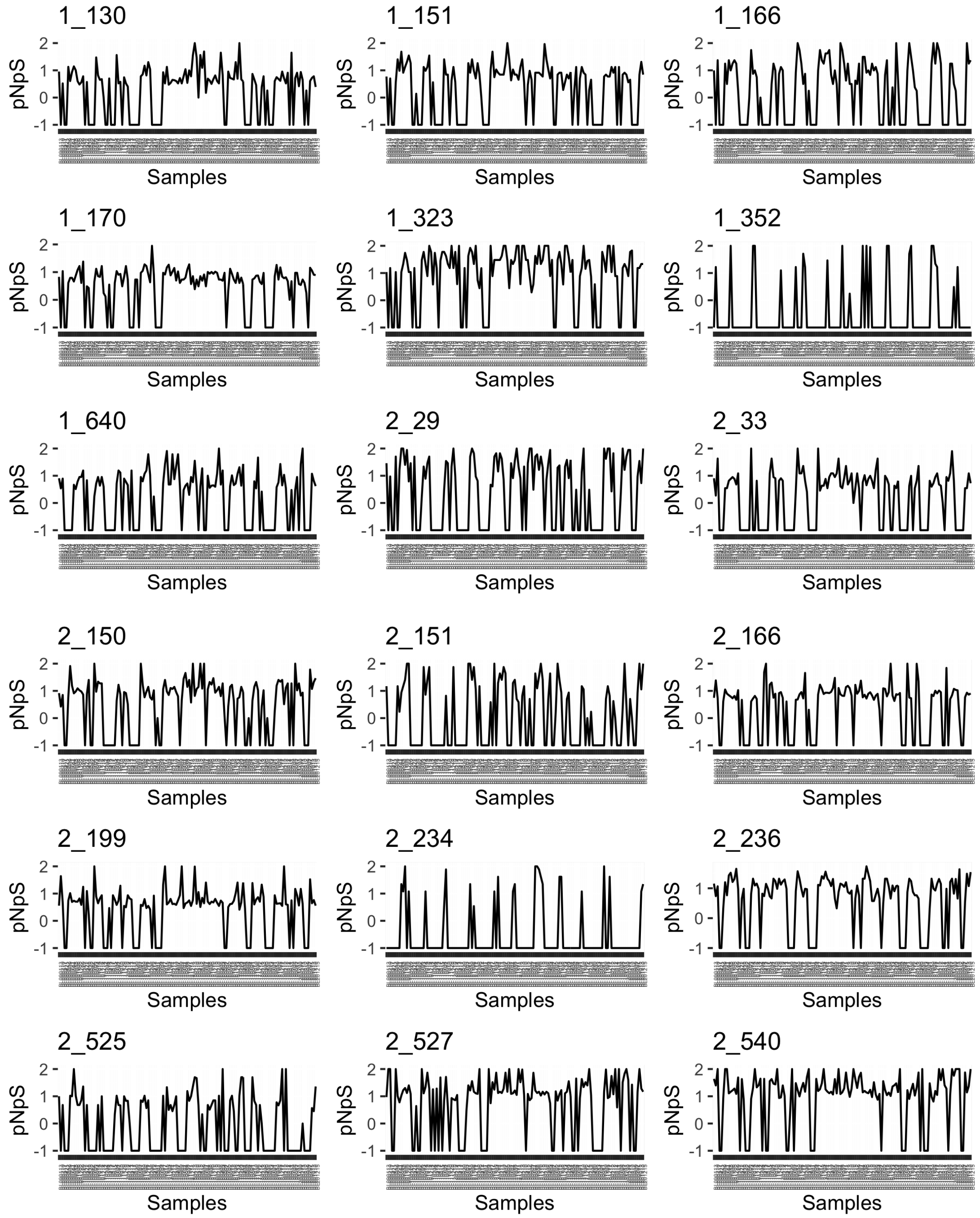


*Prochlorococcus* MAG (BL_0908_bin.full.869) in BBMO.

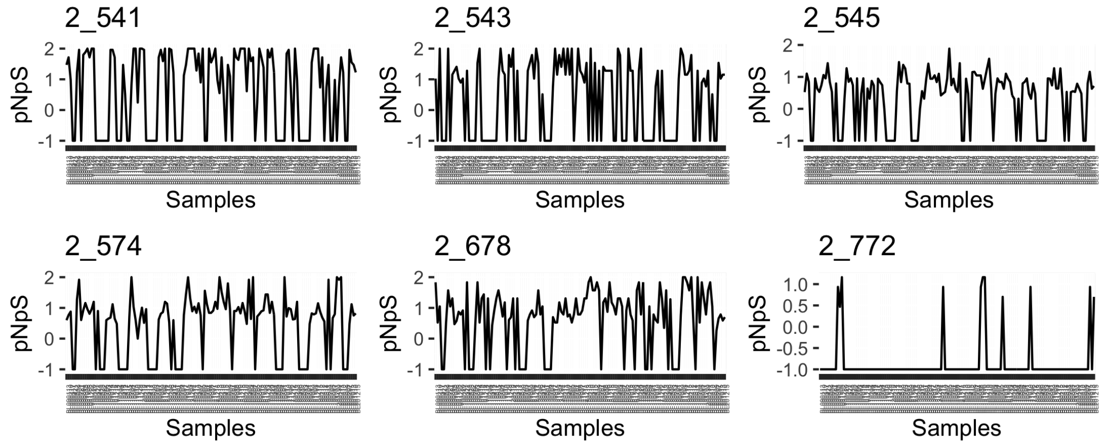


*Prochlorococcus* MAG (BL_0908_bin.full.869) in BBMO.

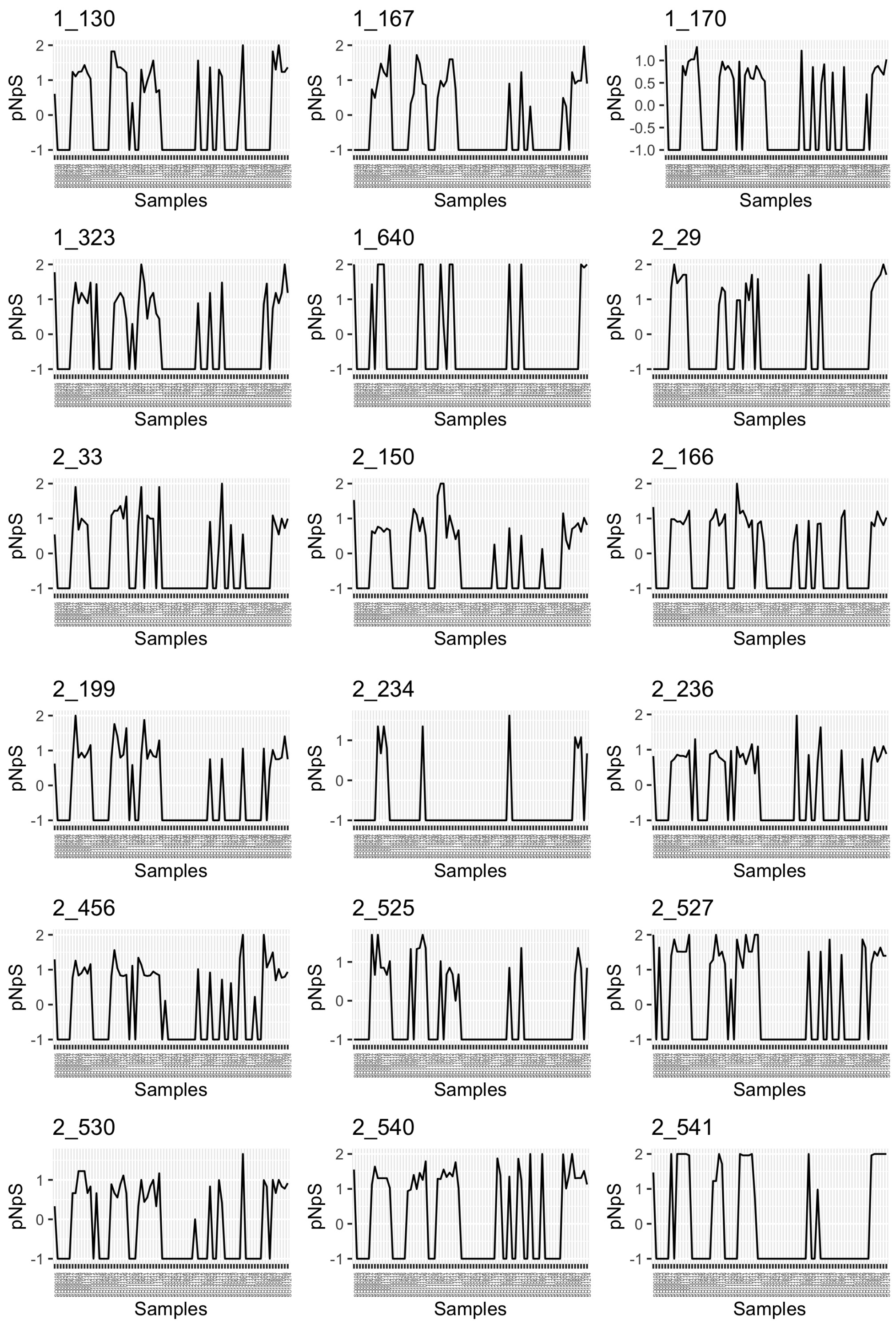


*Prochlorococcus* MAG (BL_0908_bin.full.869) in SOLA.

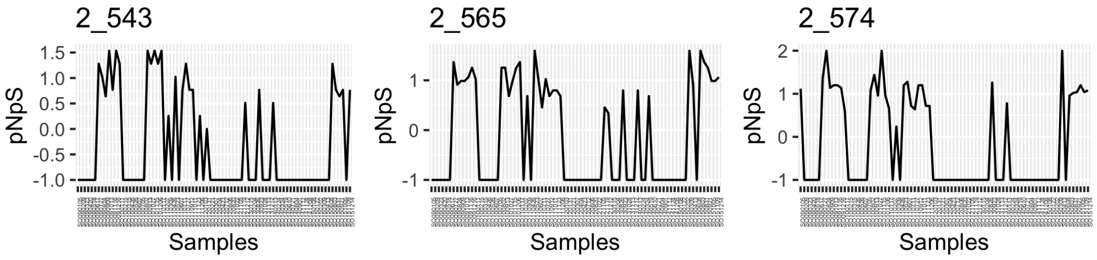


*Prochlorococcus* MAG (BL_0908_bin.full.869) in SOLA.

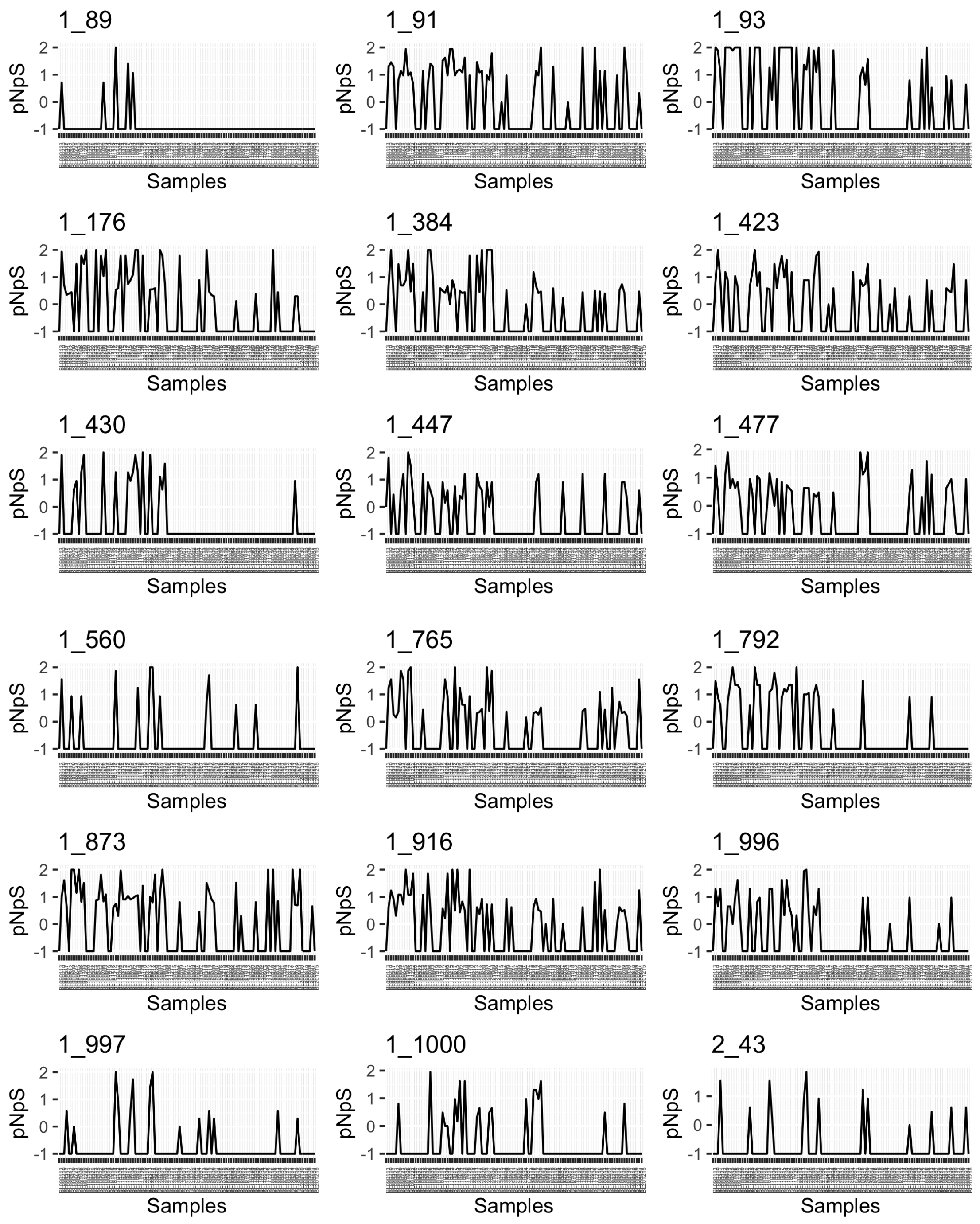


*Glaciecola* MAG (BL_1001_bin.full.234) in BBMO.

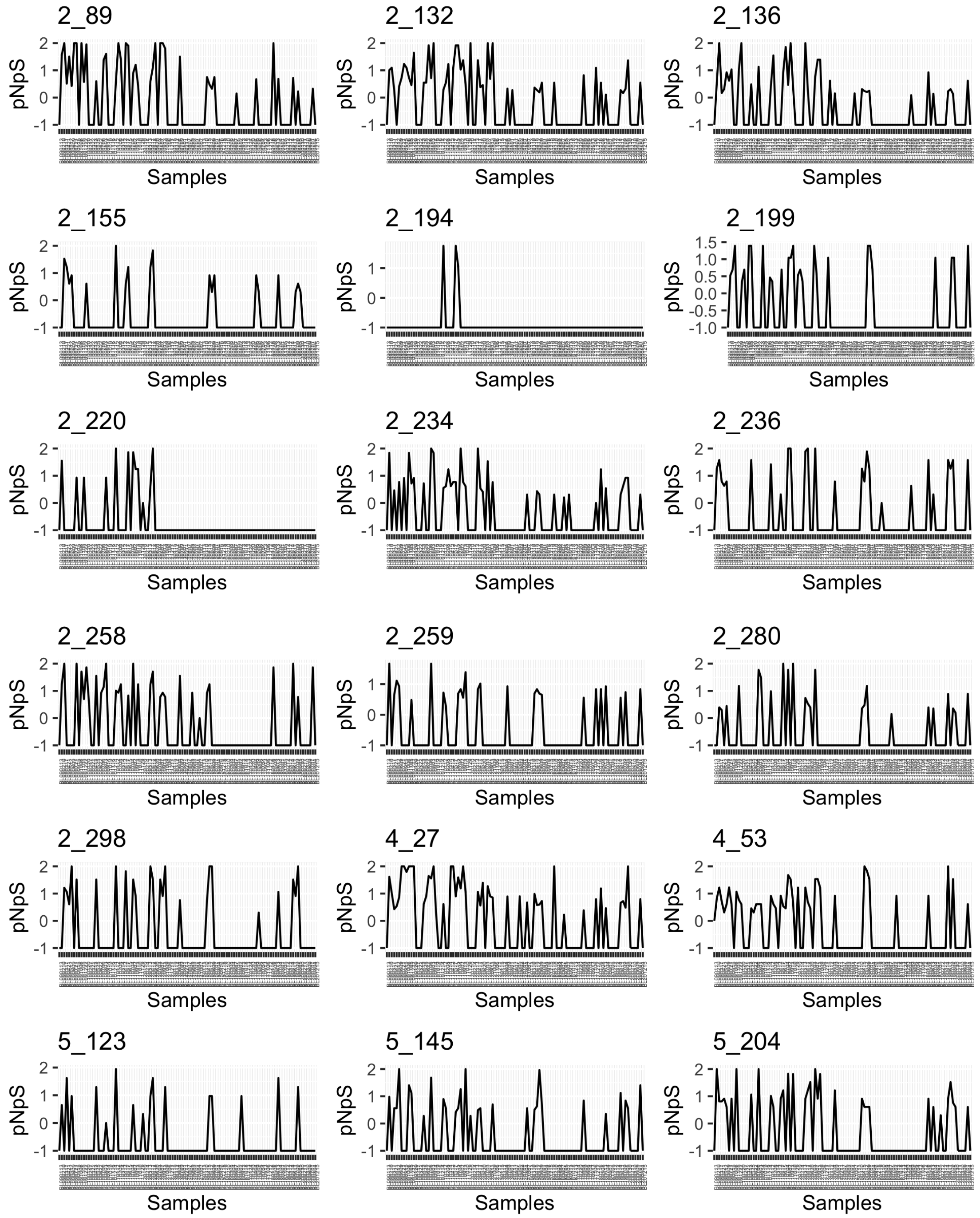


*Glaciecola* MAG (BL_1001_bin.full.234) in BBMO.

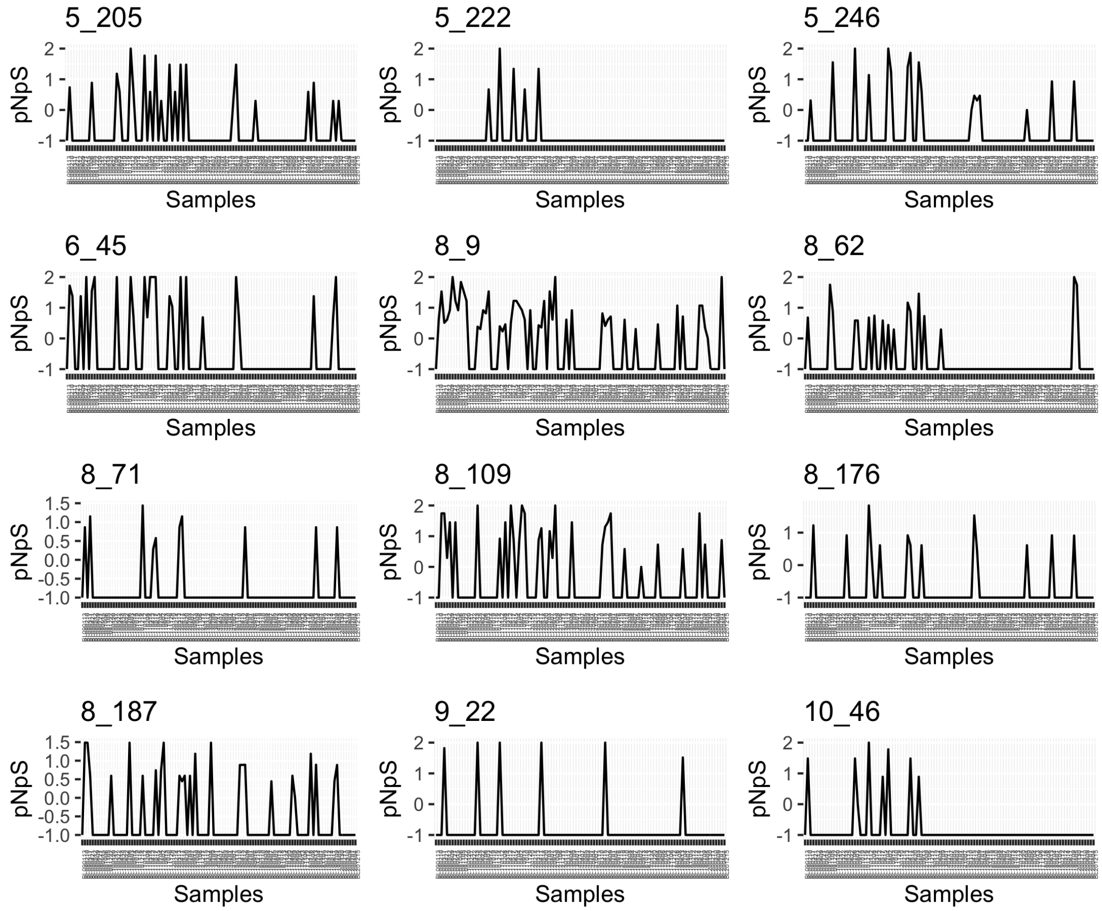


*Glaciecola* MAG (BL_1001_bin.full.234) in BBMO.

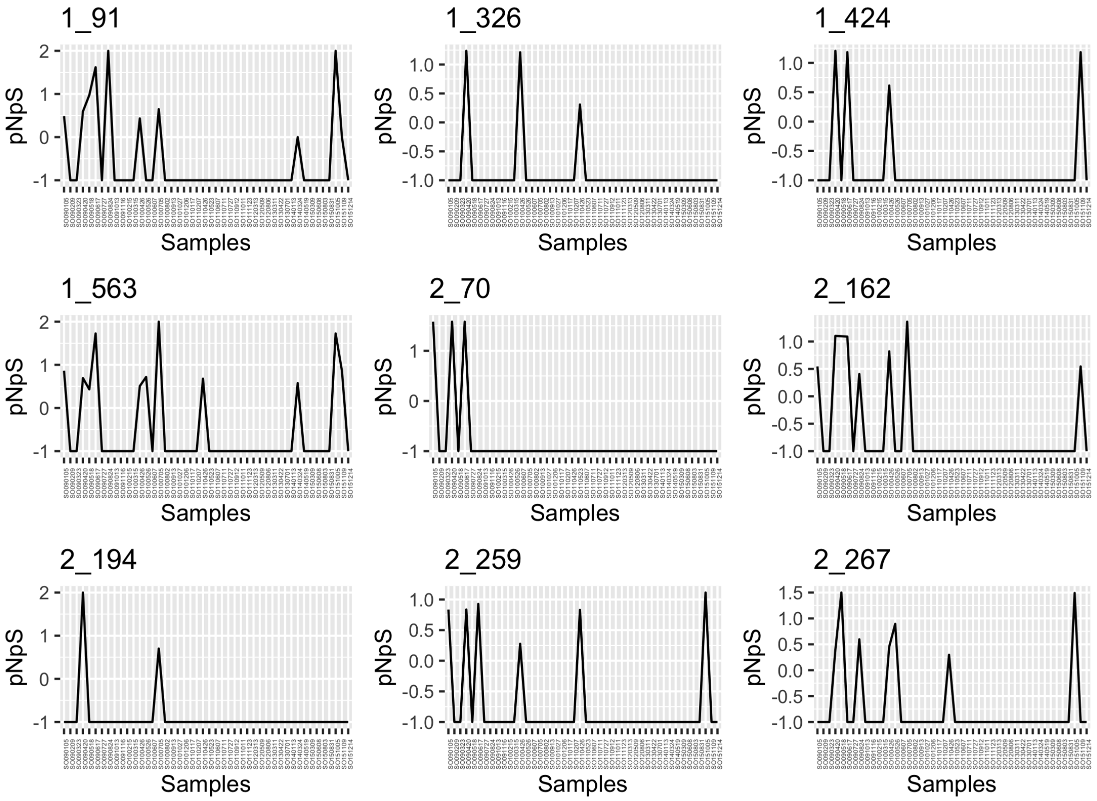


*Glaciecola* MAG (BL_1001_bin.full.234) in SOLA.

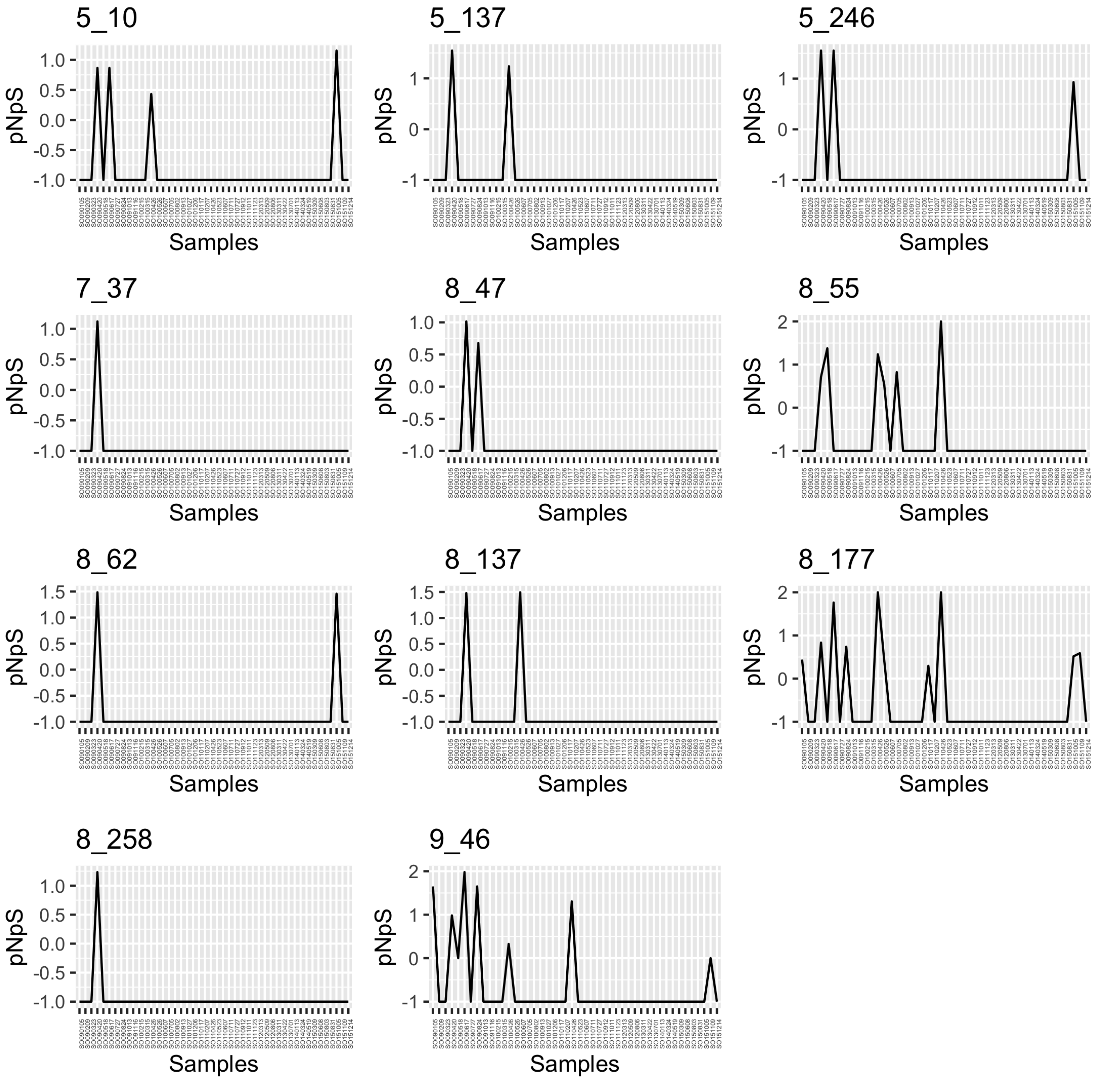


*Glaciecola* MAG (BL_1001_bin.full.234) in SOLA.

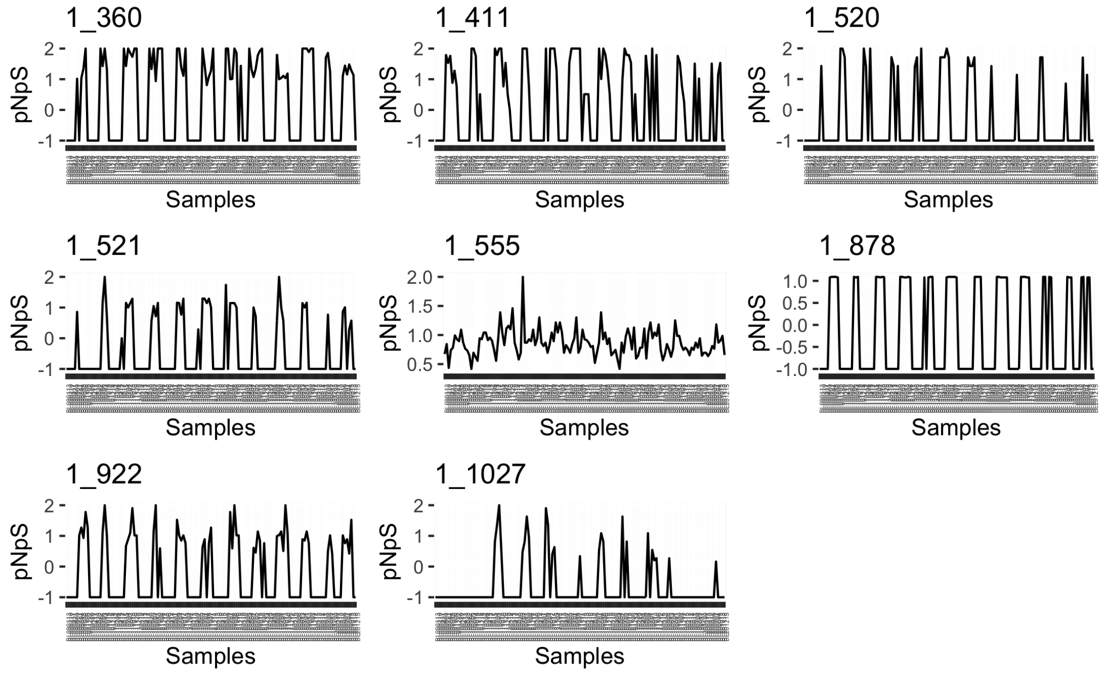


TMED112 Gammaproteobacteria MAG (BL_pooled_bin.circ.35) in BBMO.

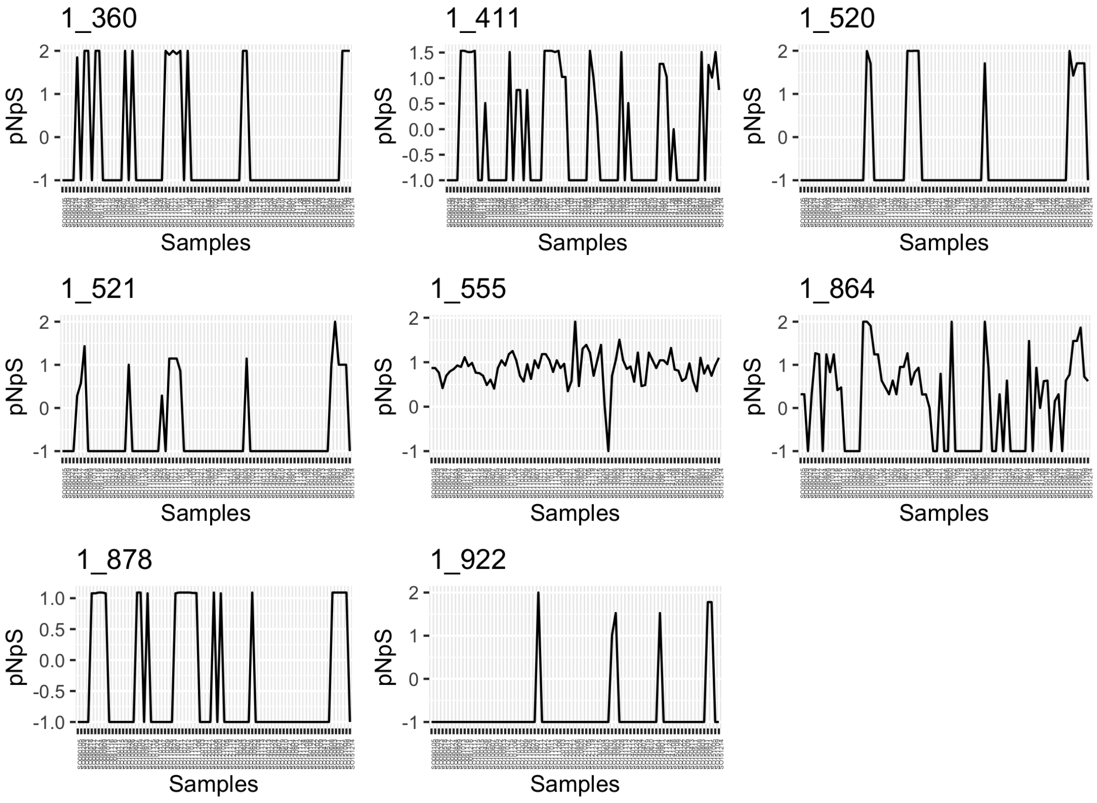


TMED112 Gammaproteobacteria MAG (BL_pooled_bin.circ.35) in SOLA.

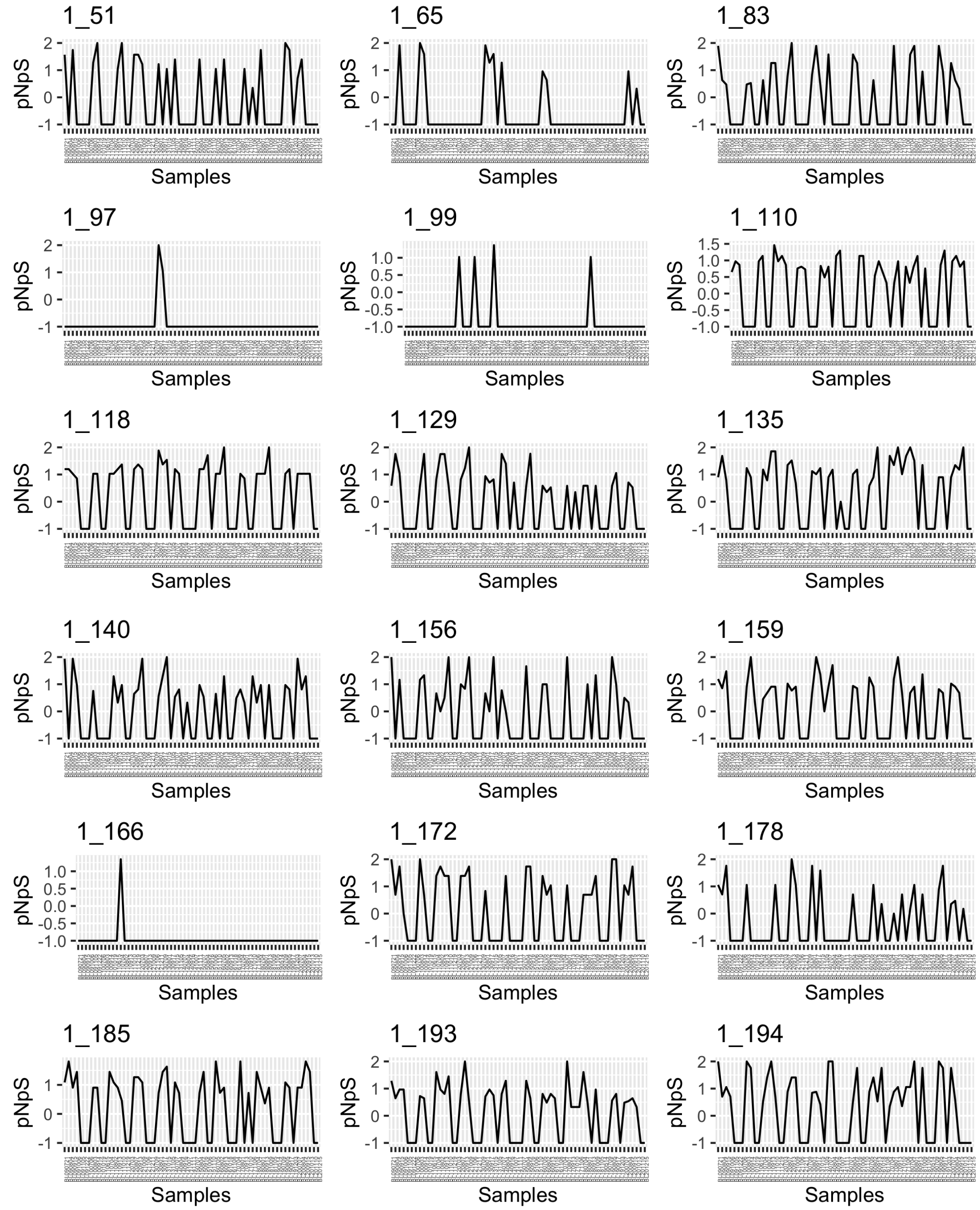


UBA8752 Bacteroidia MAG (BL_pooled_bin.circ.36) in BBMO.

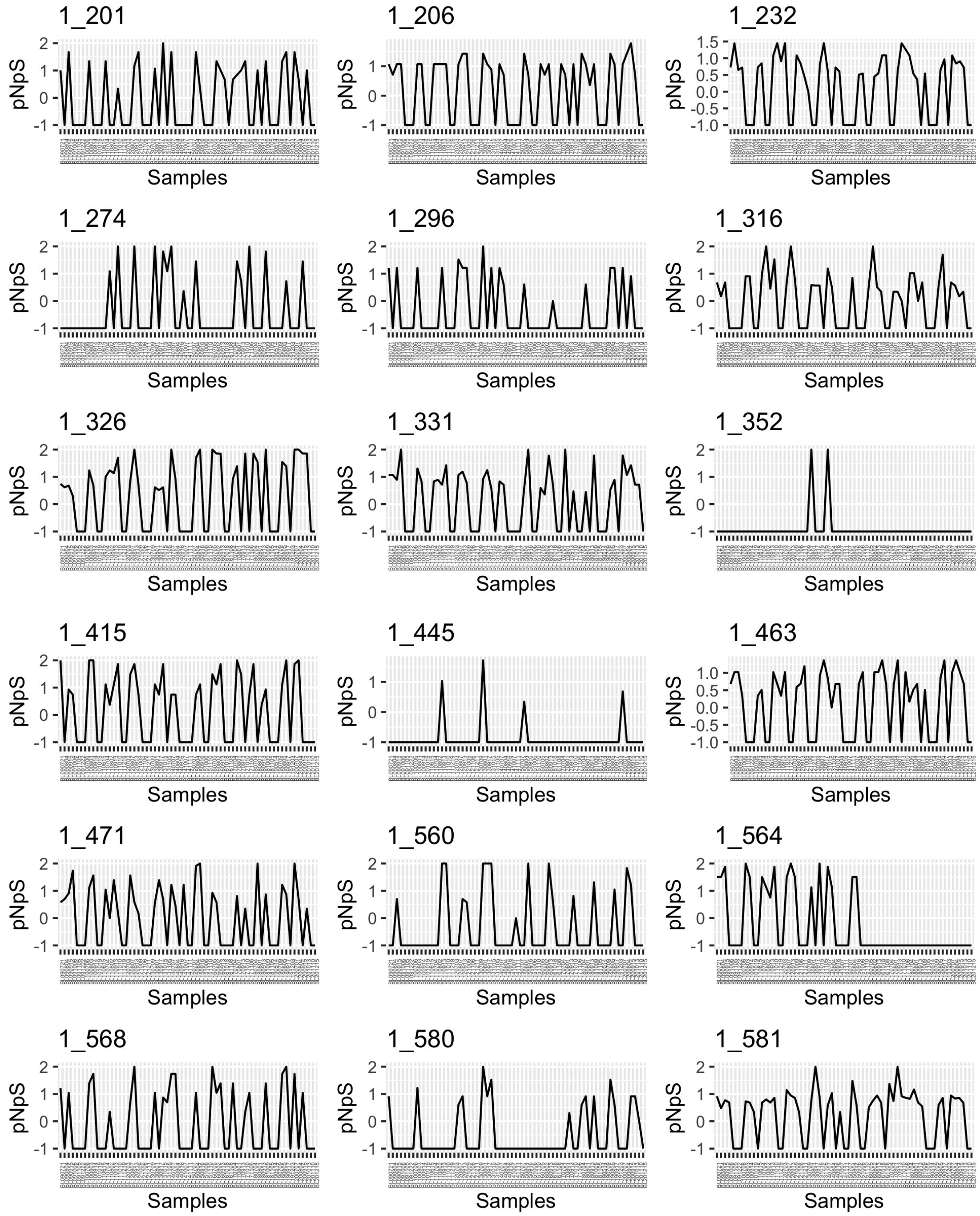


UBA8752 Bacteroidia MAG (BL_pooled_bin.circ.36) in BBMO.

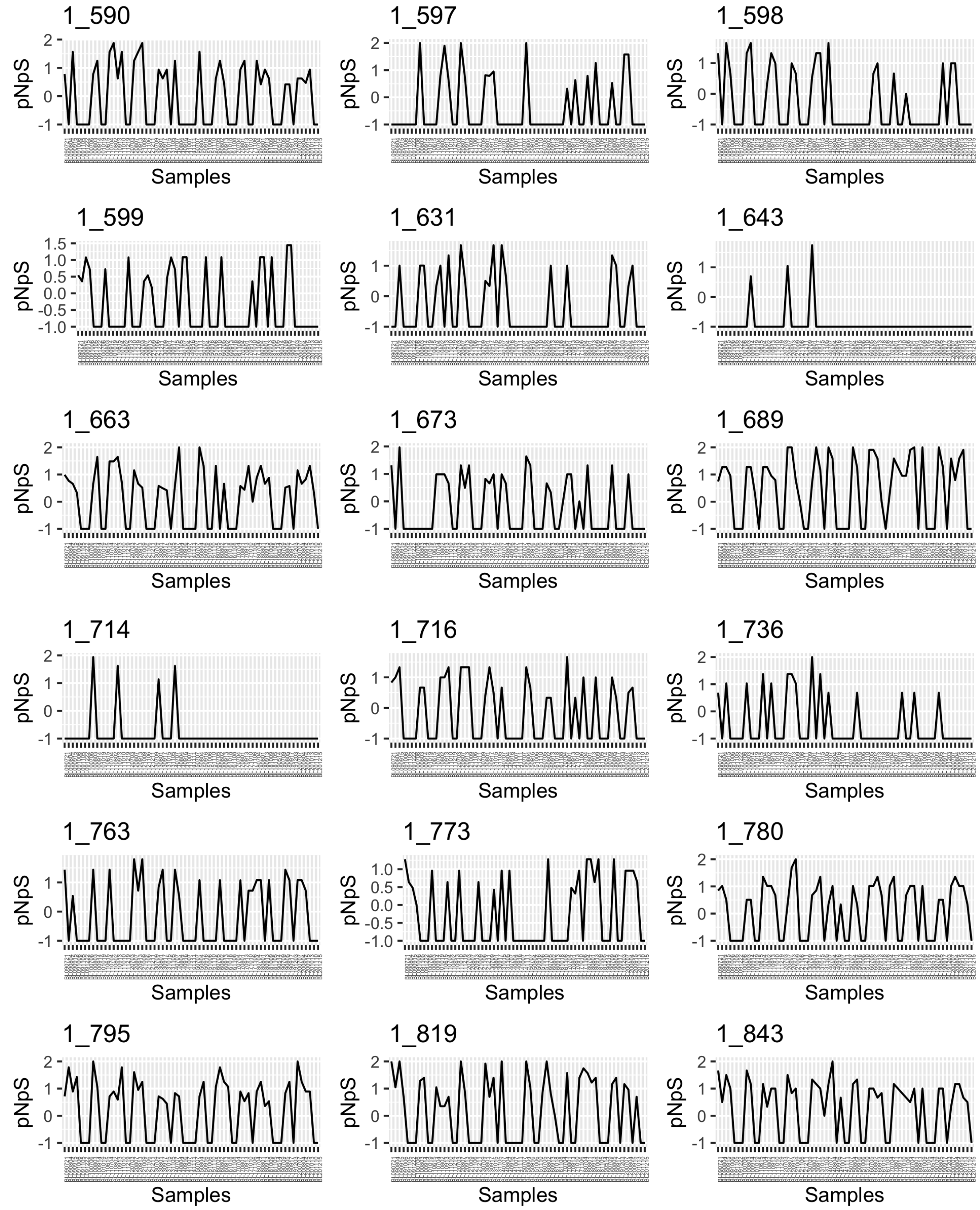


UBA8752 Bacteroidia MAG (BL_pooled_bin.circ.36) in BBMO.

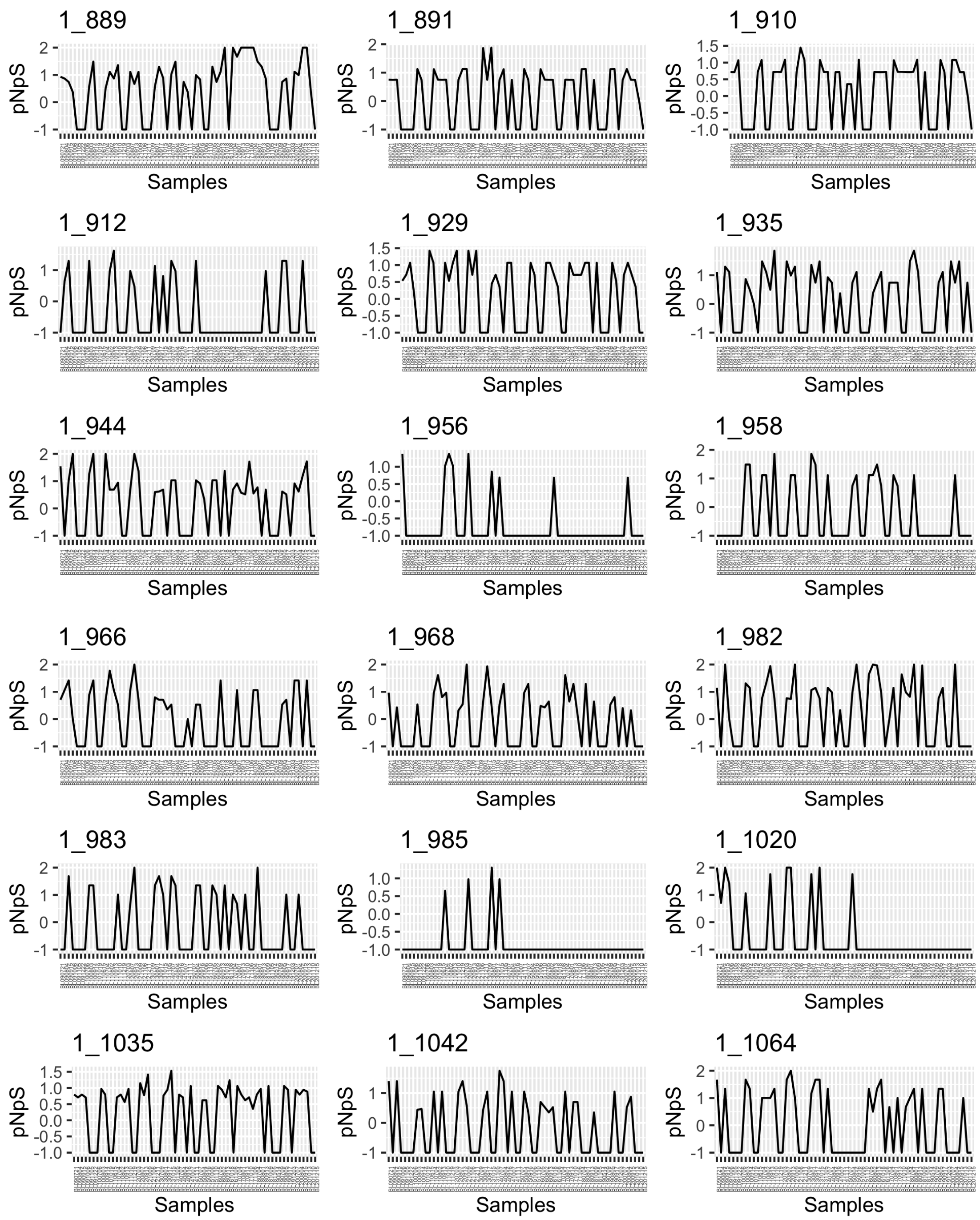


UBA8752 Bacteroidia MAG (BL_pooled_bin.circ.36) in BBMO.

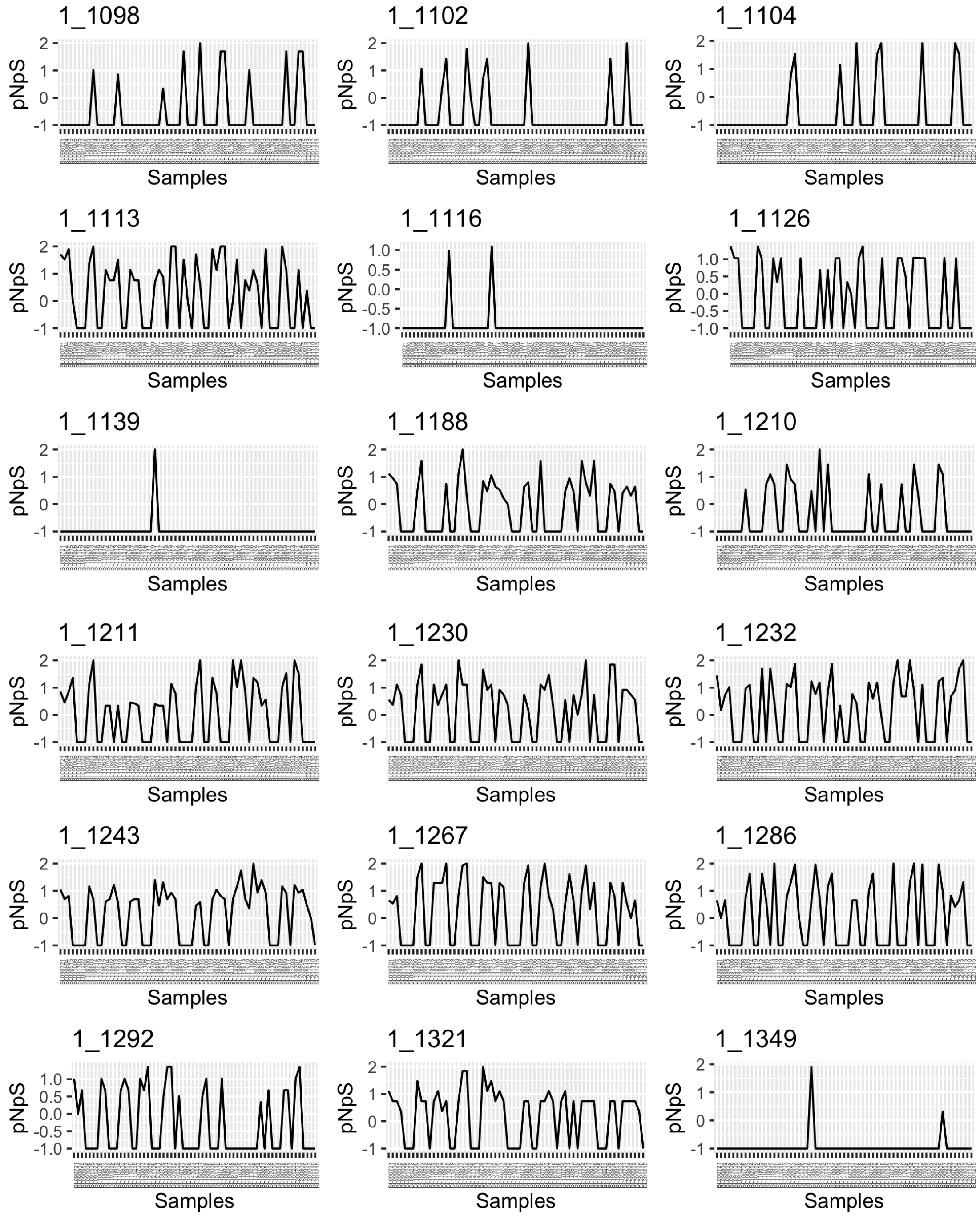


UBA8752 Bacteroidia MAG (BL_pooled_bin.circ.36) in BBMO.

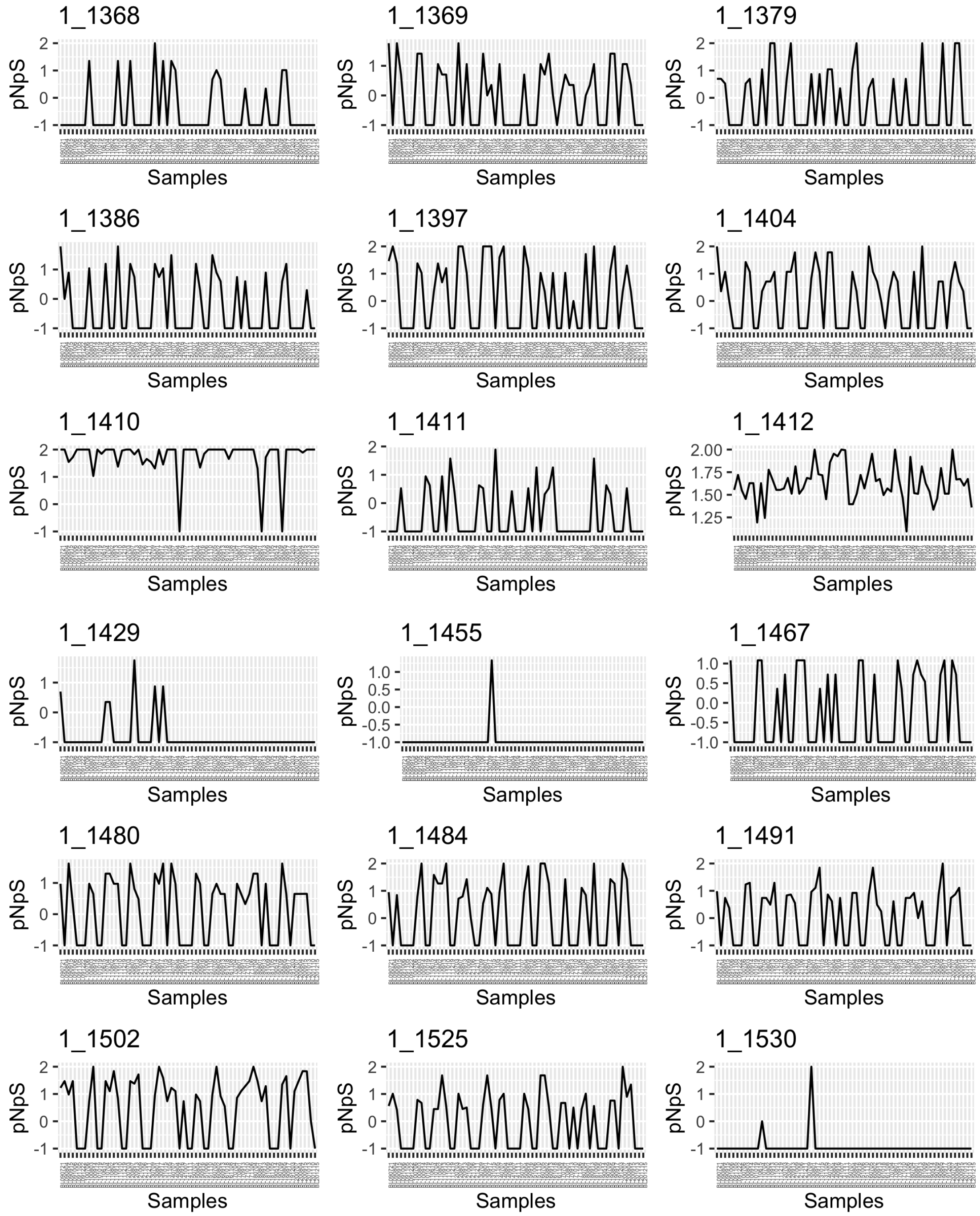


UBA8752 Bacteroidia MAG (BL_pooled_bin.circ.36) in BBMO

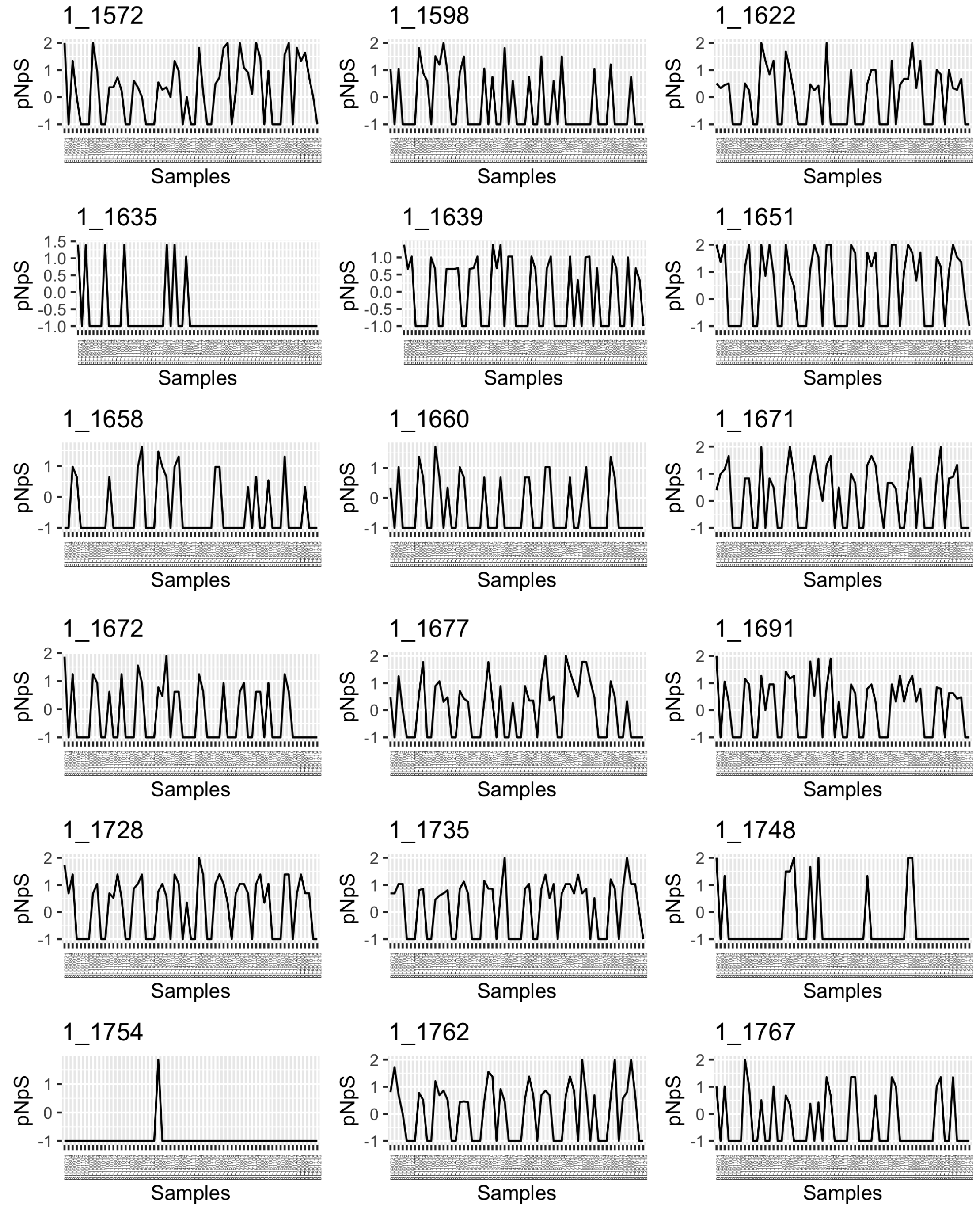


UBA8752 Bacteroidia MAG (BL_pooled_bin.circ.36) in BBMO.

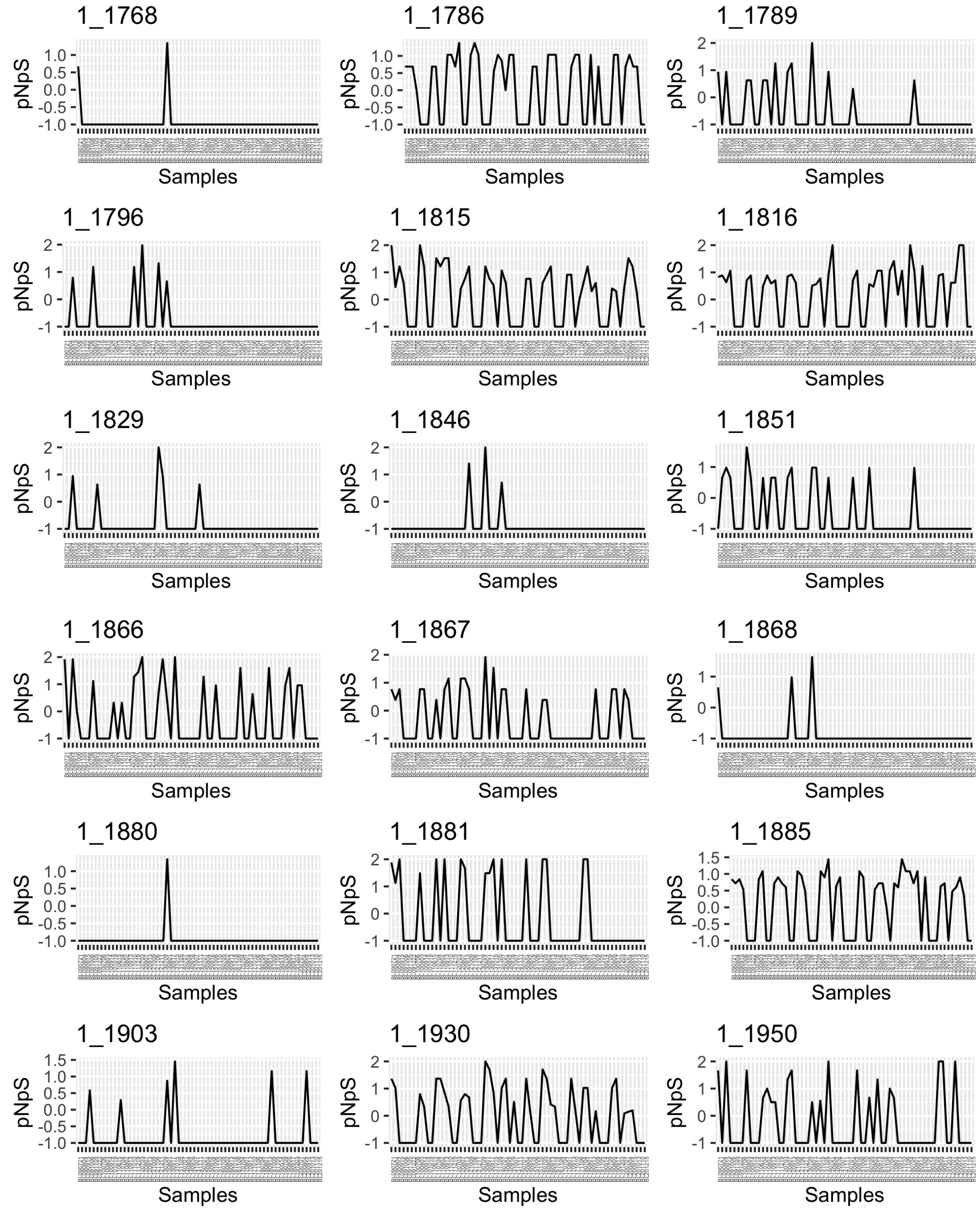


UBA8752 Bacteroidia MAG (BL_pooled_bin.circ.36) in BBMO.

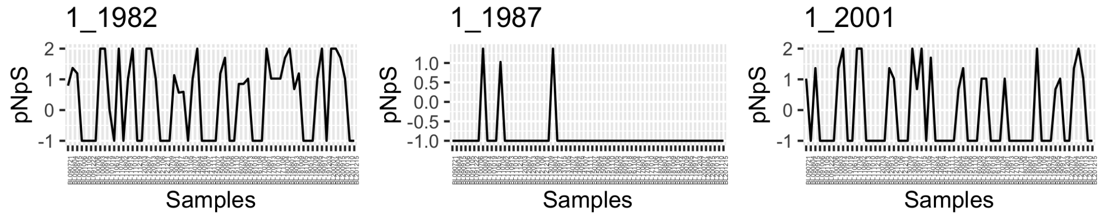


UBA8752 Bacteroidia MAG (BL_pooled_bin.circ.36) in BBMO.

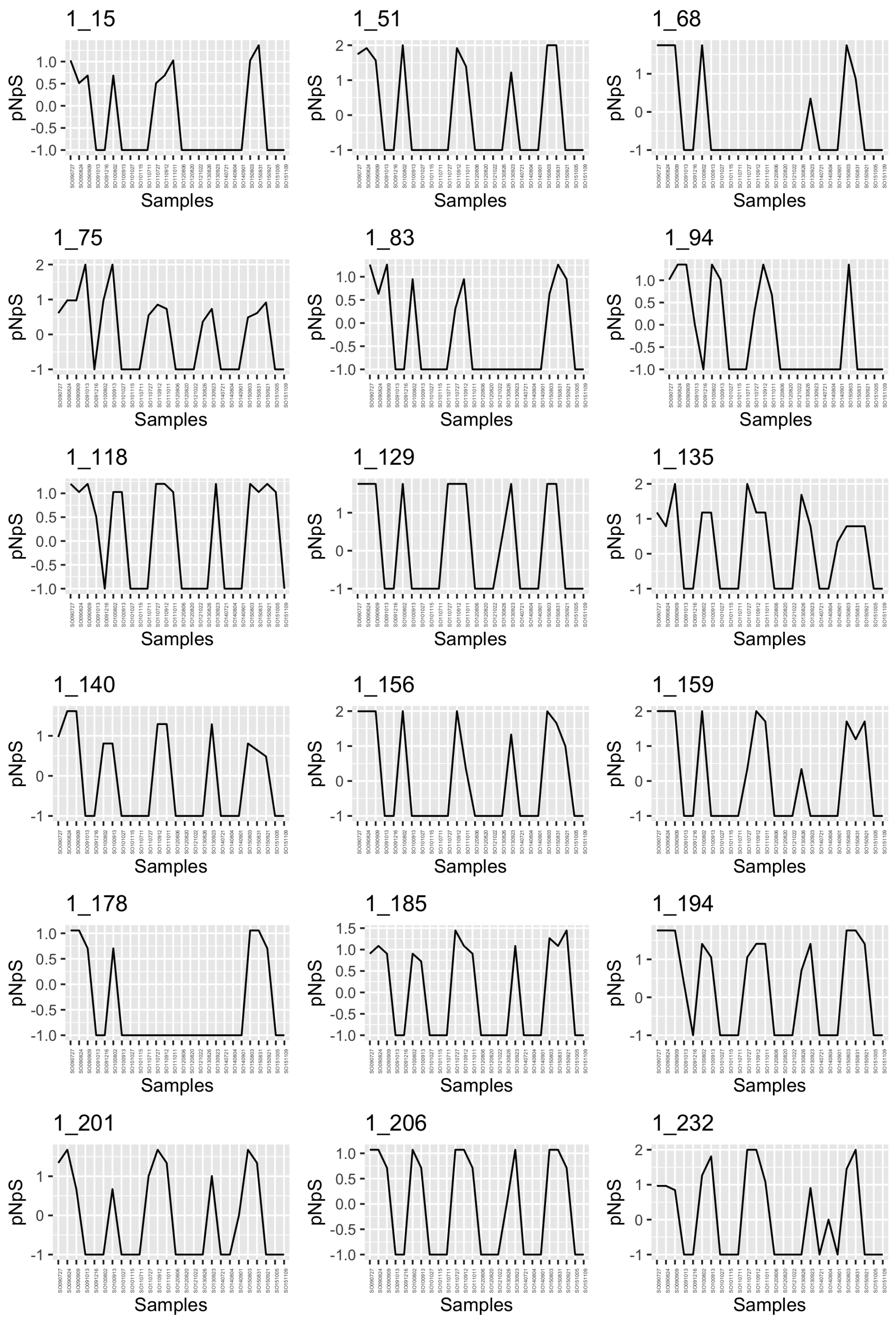


UBA8752 Bacteroidia MAG (BL_pooled_bin.circ.36) in SOLA.

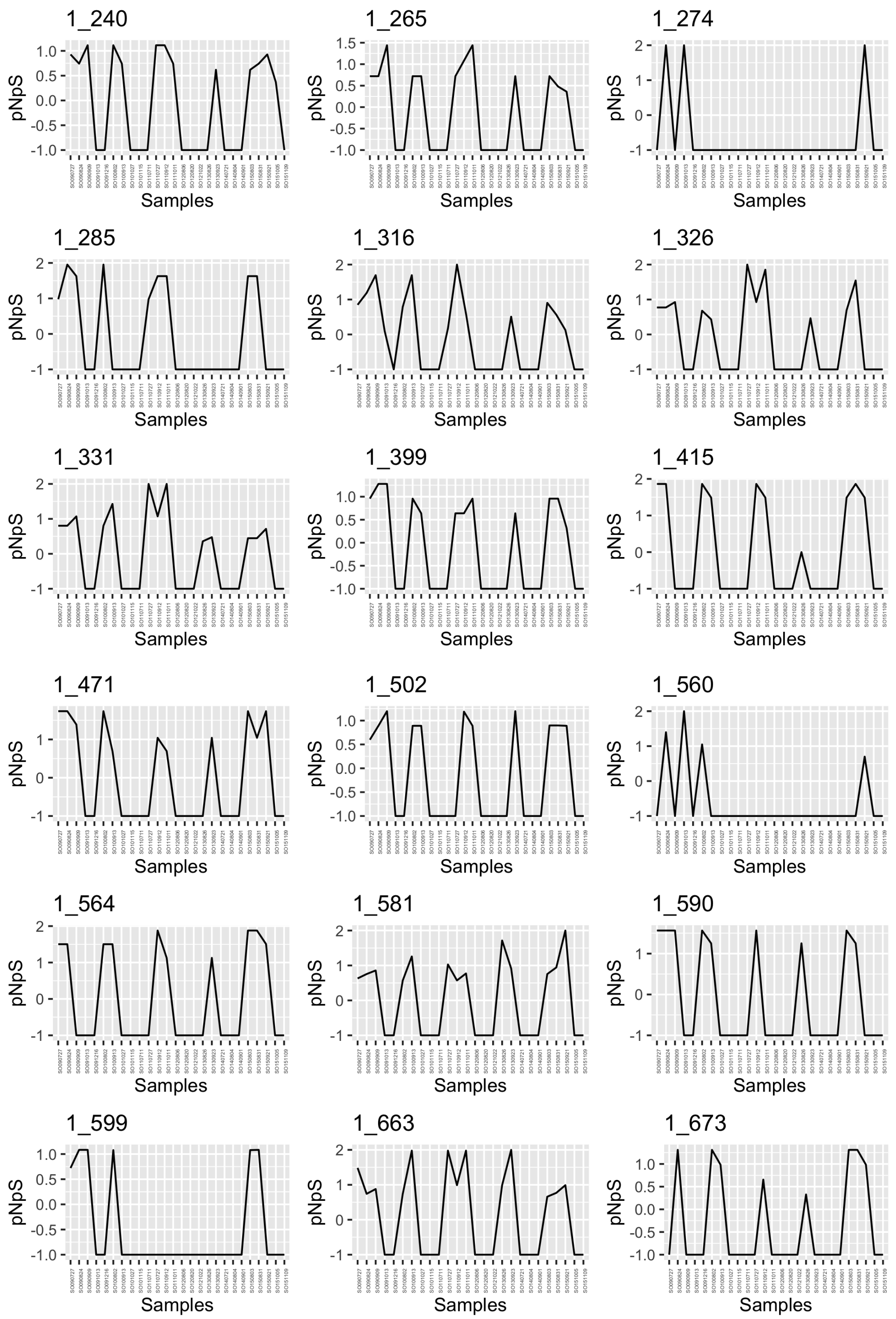


UBA8752 Bacteroidia MAG (BL_pooled_bin.circ.36) in SOLA.

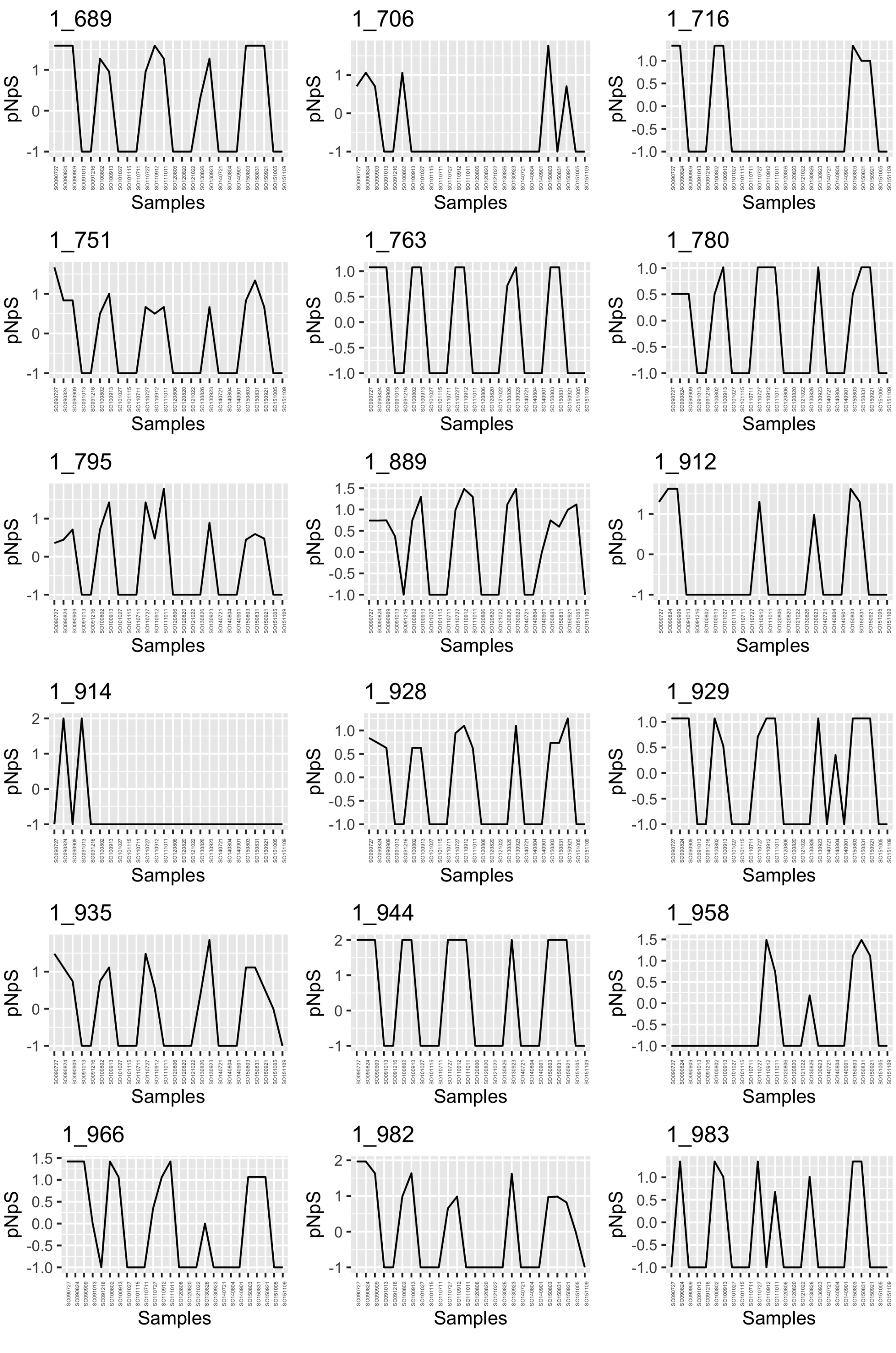


UBA8752 Bacteroidia MAG (BL_pooled_bin.circ.36) in SOLA.

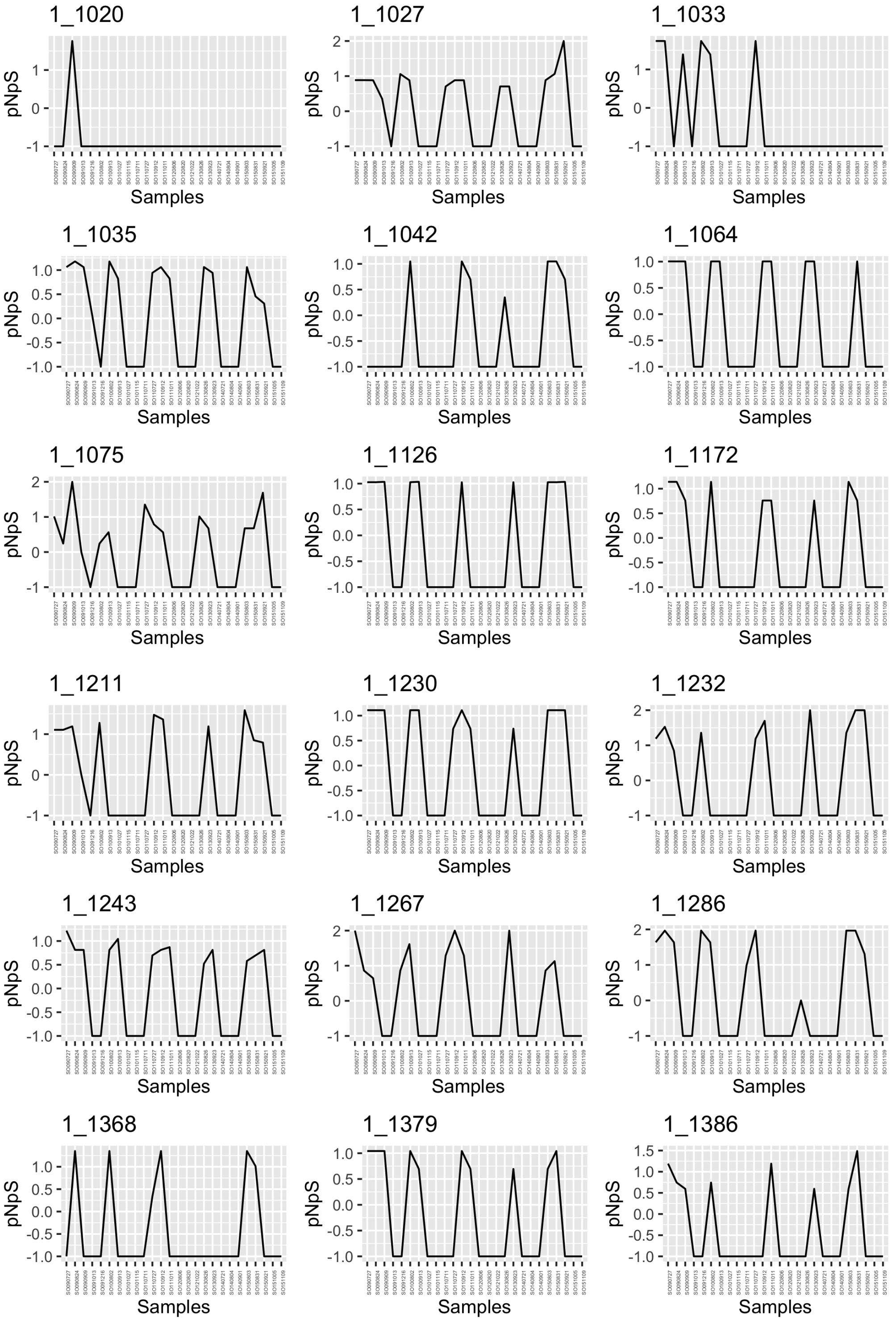


UBA8752 Bacteroidia MAG (BL_pooled_bin.circ.36) in SOLA.

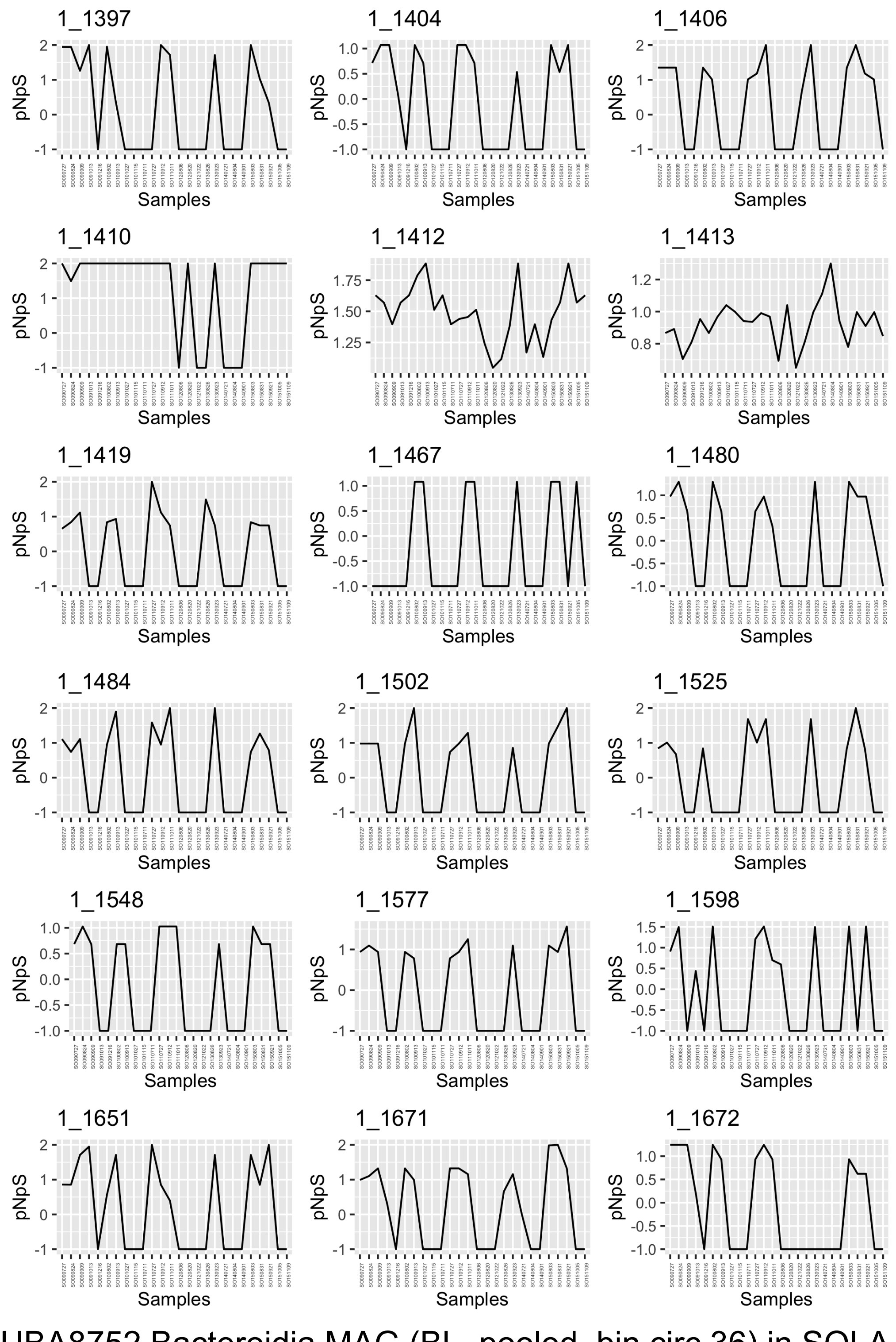


UBA8752 Bacteroidia MAG (BL_pooled_bin.circ.36) in SOLA.

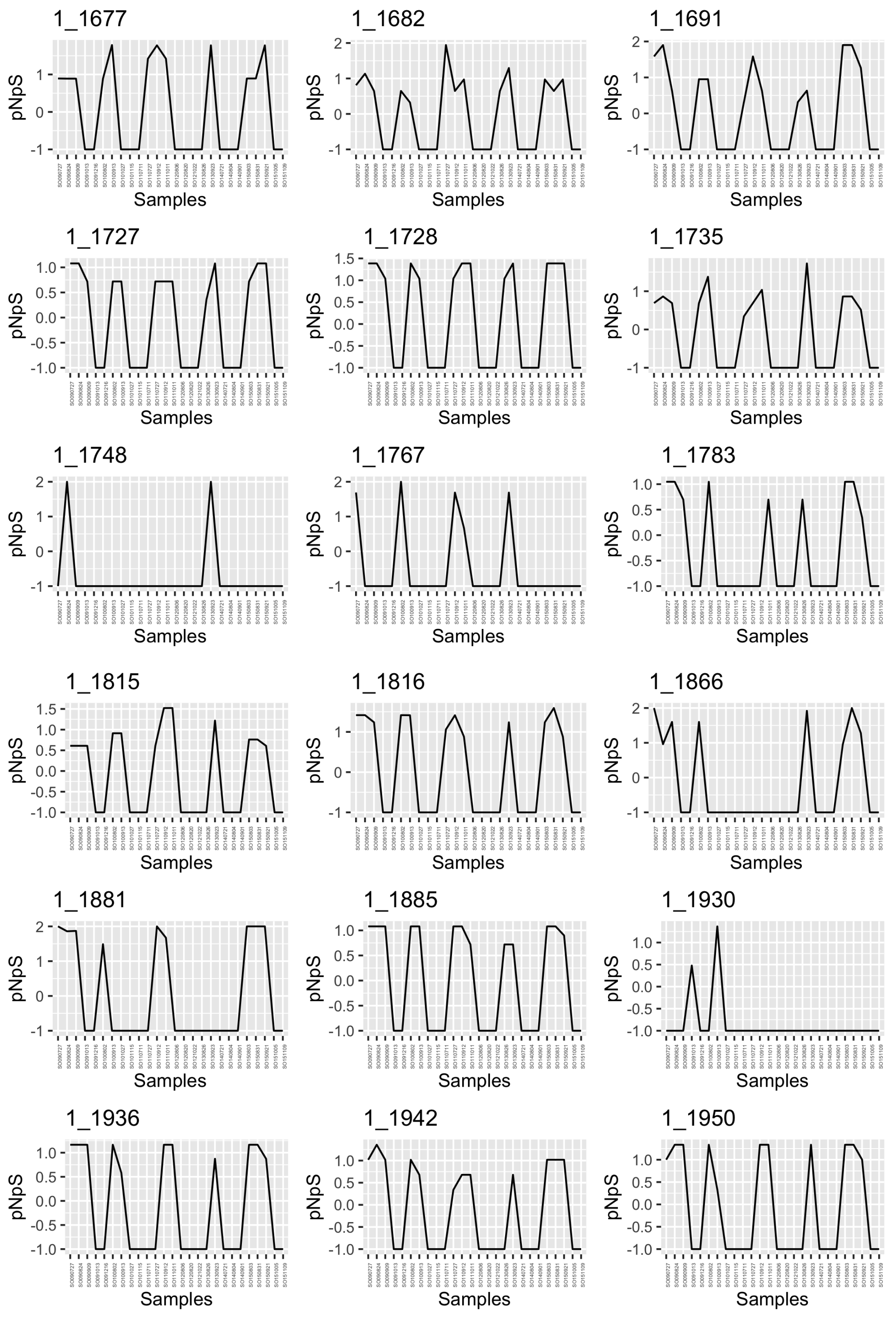


UBA8752 Bacteroidia MAG (BL_pooled_bin.circ.36) in SOLA.

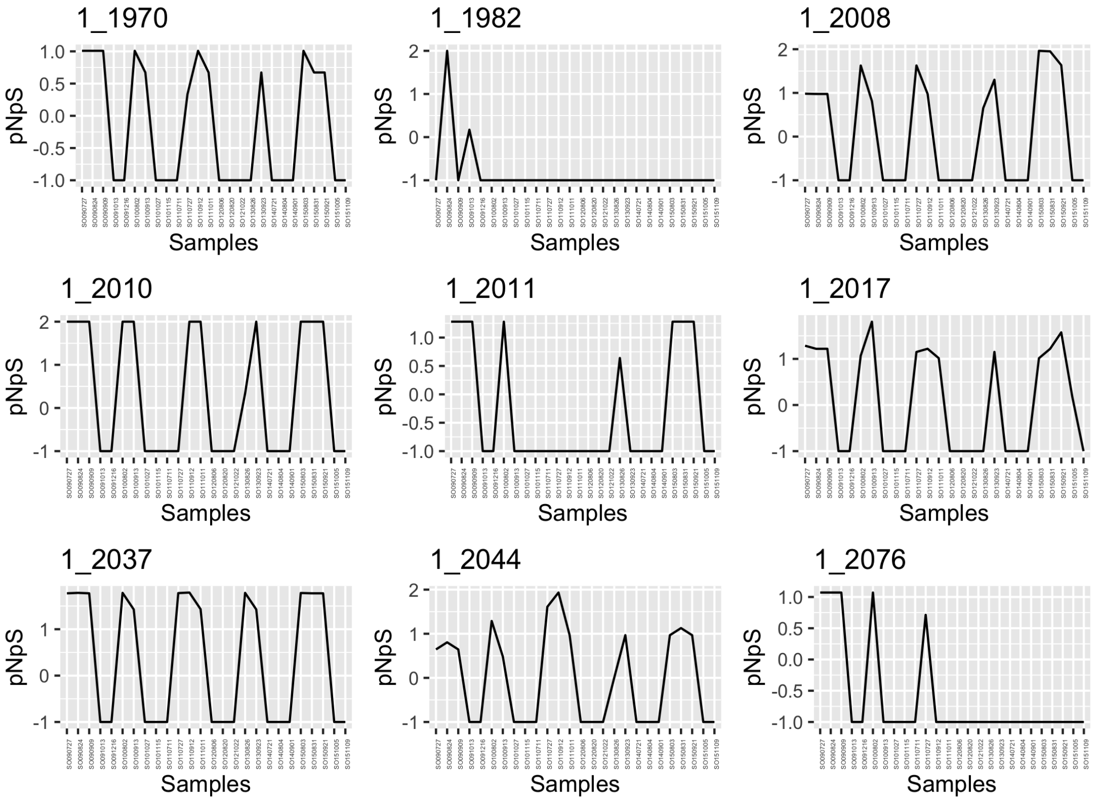


UBA8752 Bacteroidia MAG (BL_pooled_bin.circ.36) in SOLA.

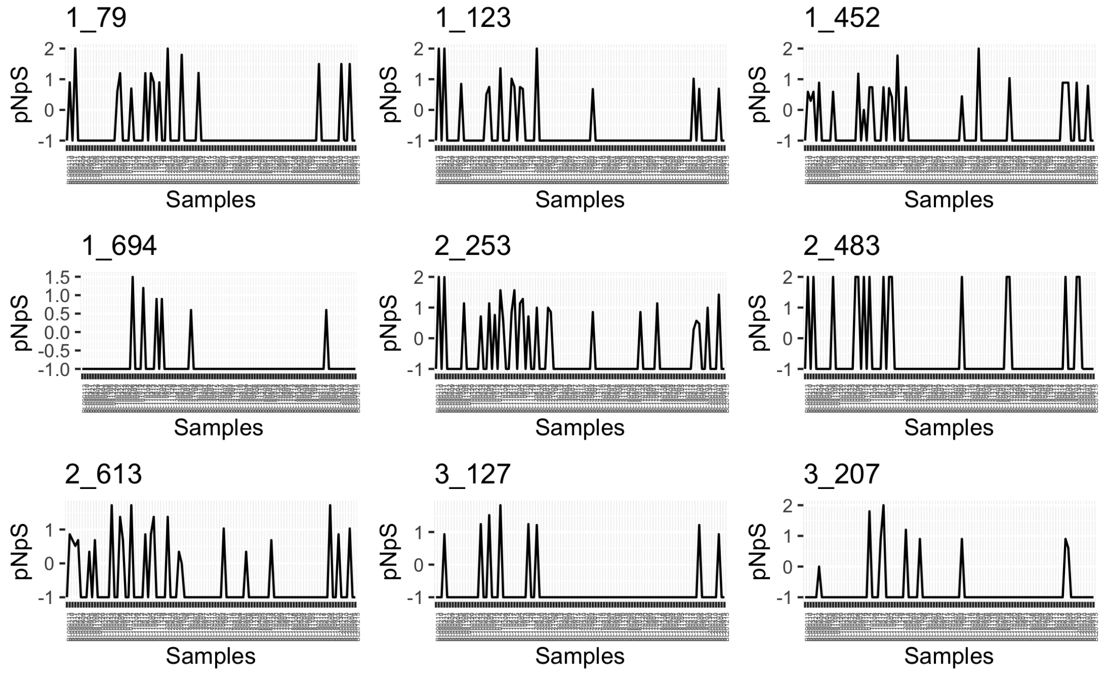


Rhodobacteraceae MAG (BL_pooled_bin.full.145) in BBMO.

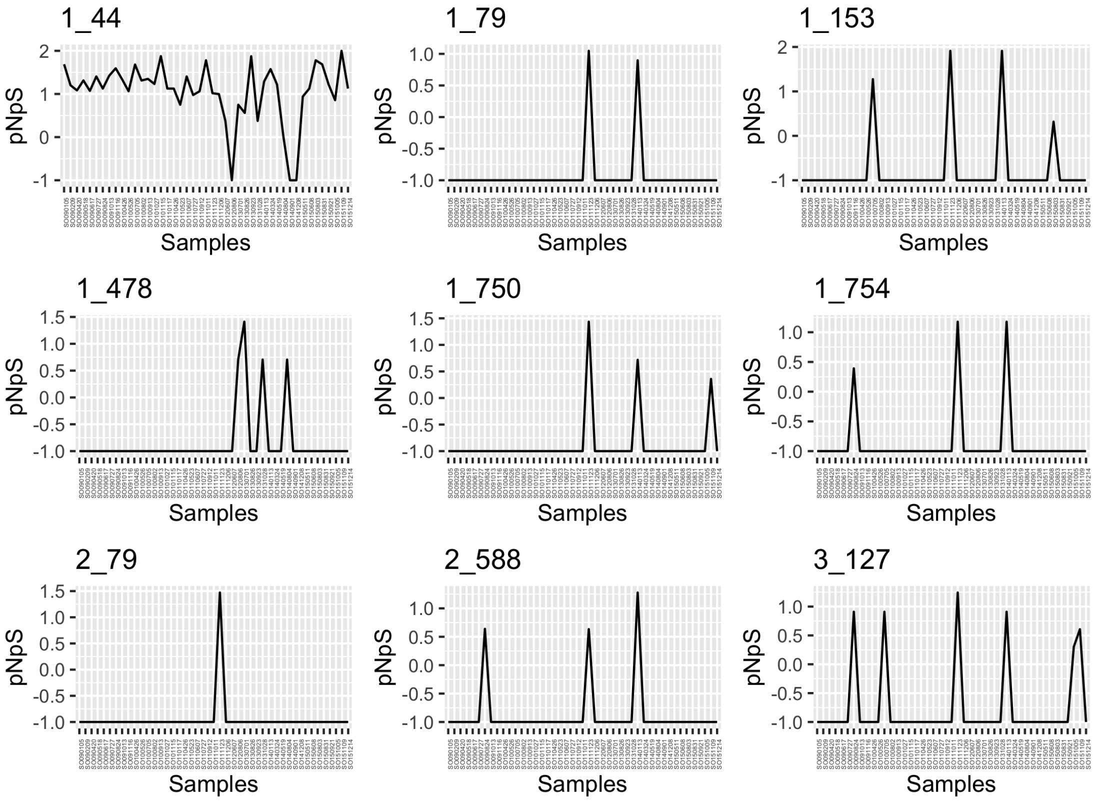


Rhodobacteraceae MAG (BL_pooled_bin.full.145) in SOLA.

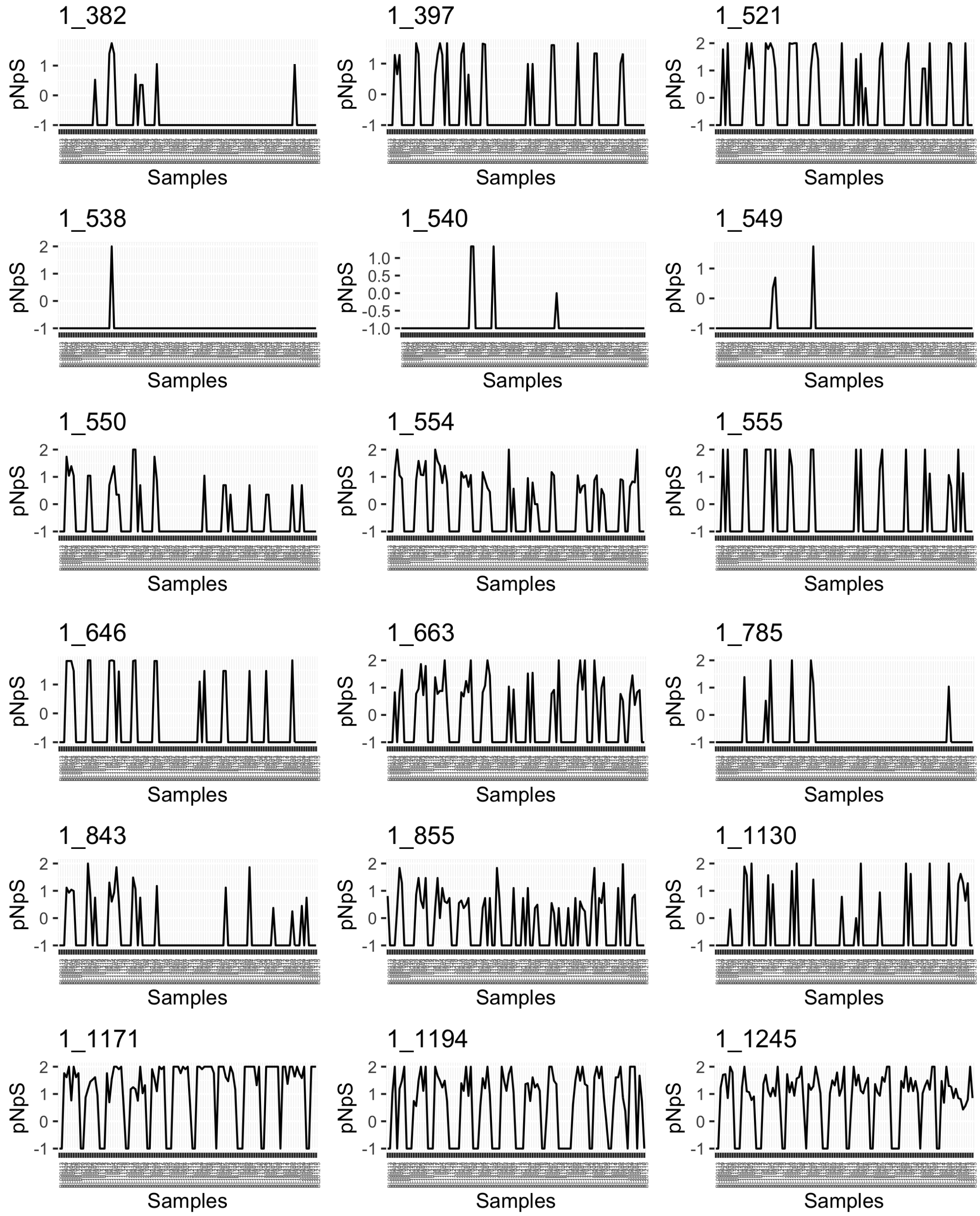


*Pontimonas* MAG (BL_pooled_bin.full.596) in BBMO.

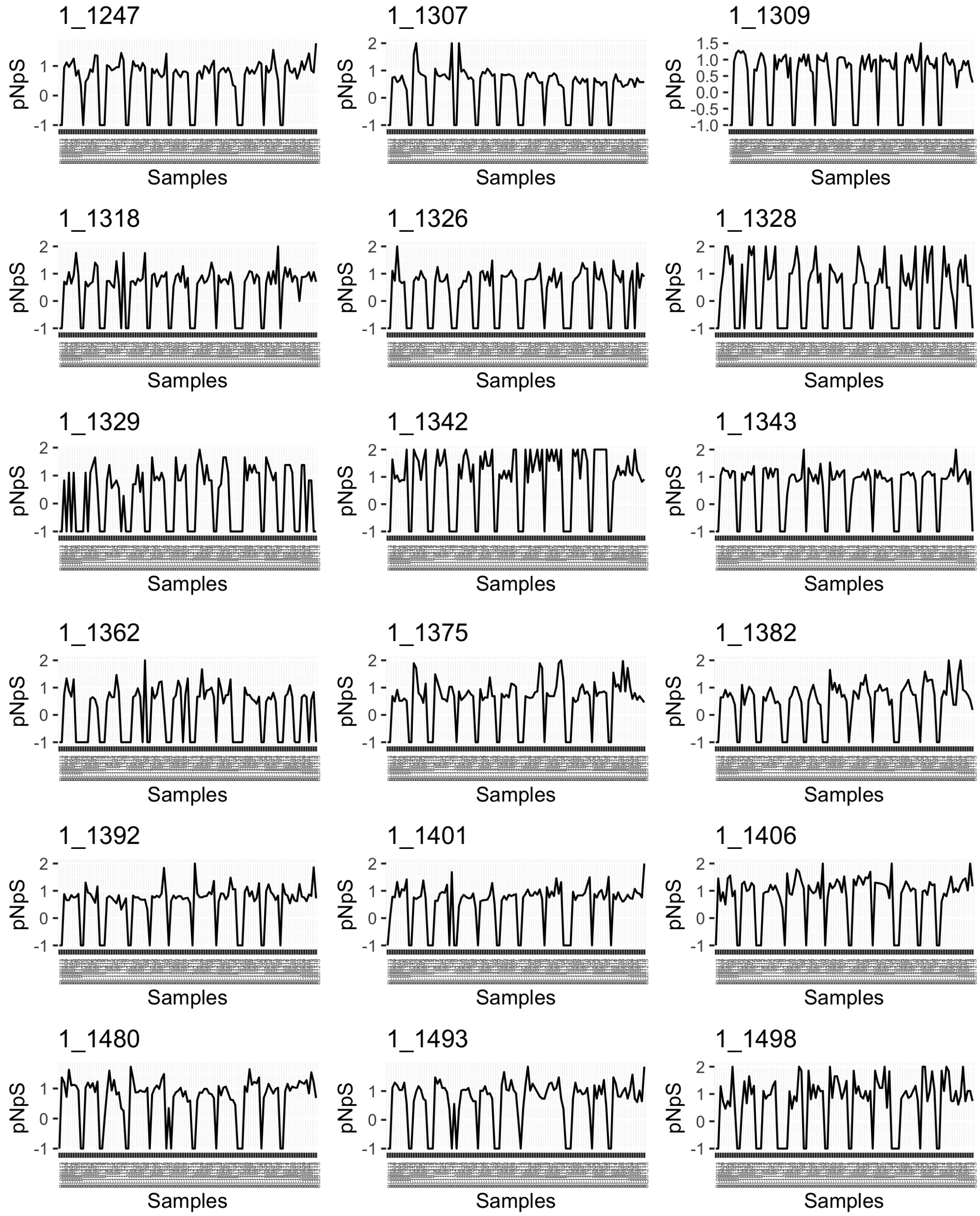


*Pontimonas* MAG (BL_pooled_bin.full.596) in BBMO.

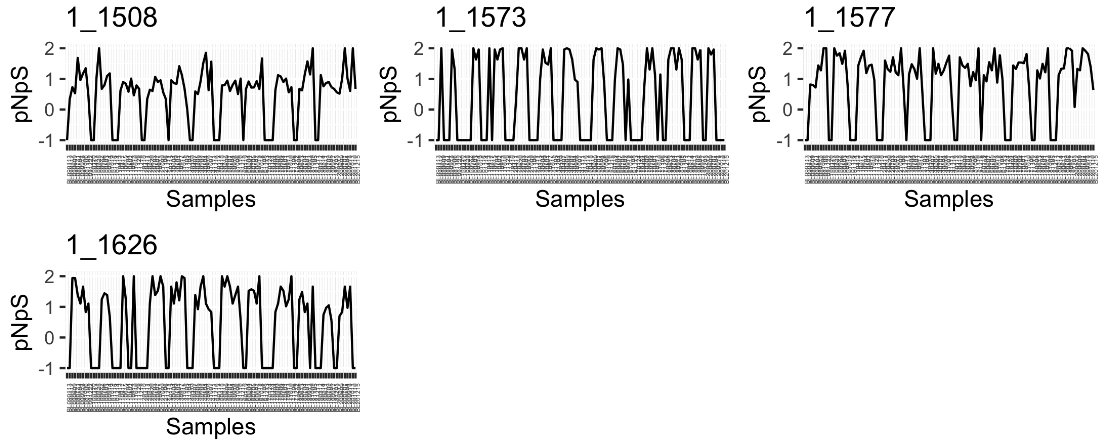


*Pontimonas* MAG (BL_pooled_bin.full.596) in BBMO.

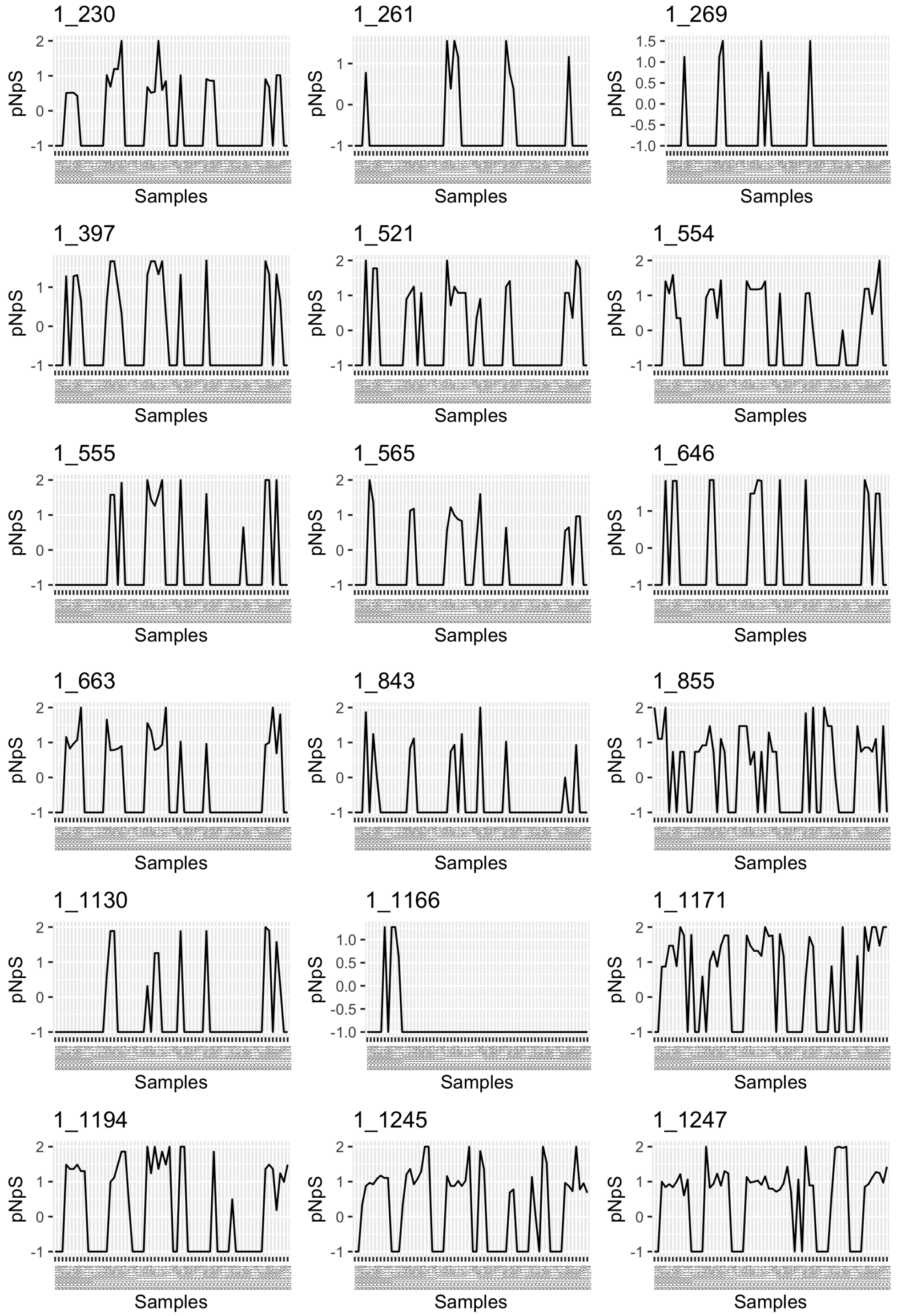


*Pontimonas* MAG (BL_pooled_bin.full.596) in SOLA.

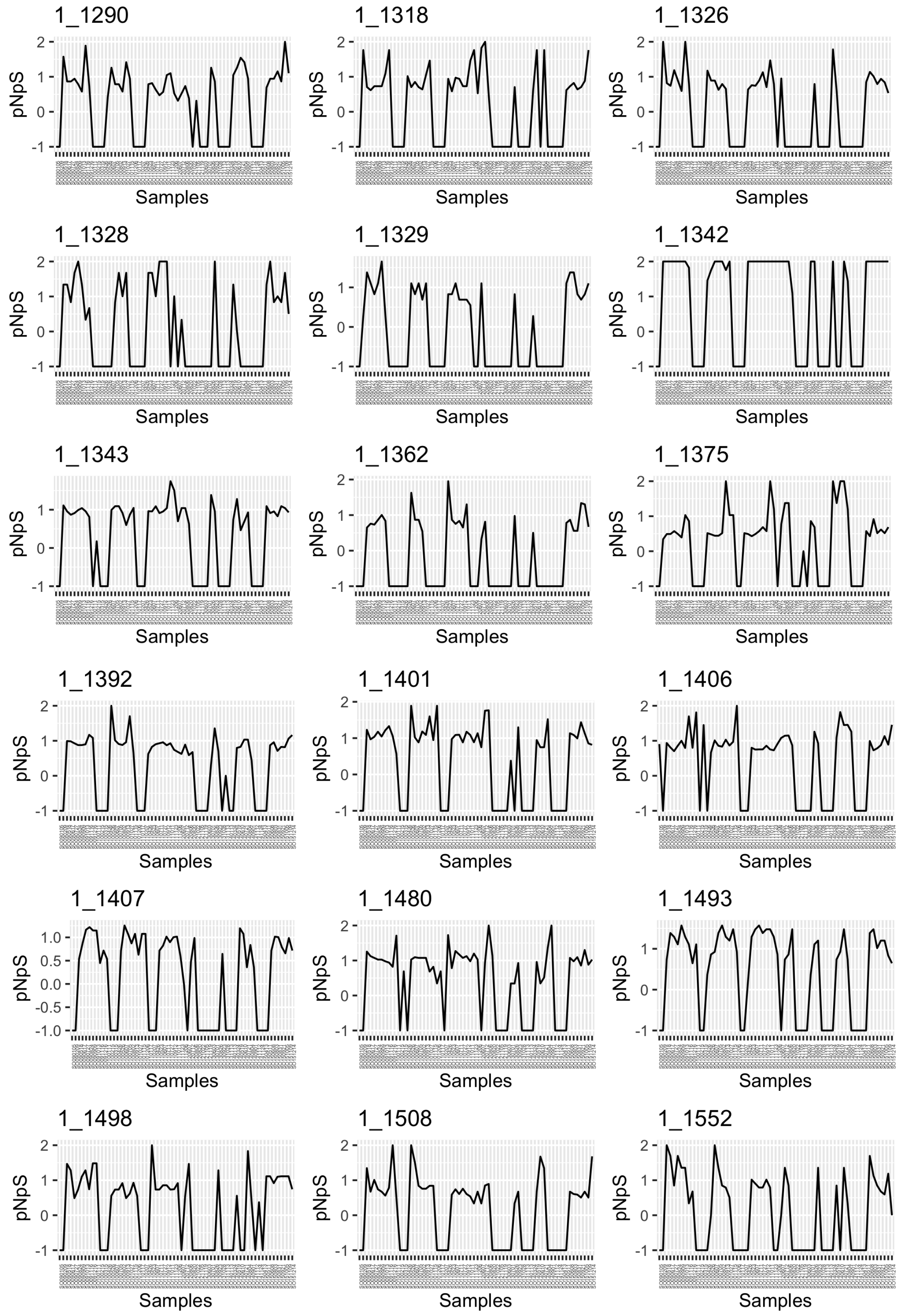


*Pontimonas* MAG (BL_pooled_bin.full.596) in SOLA.

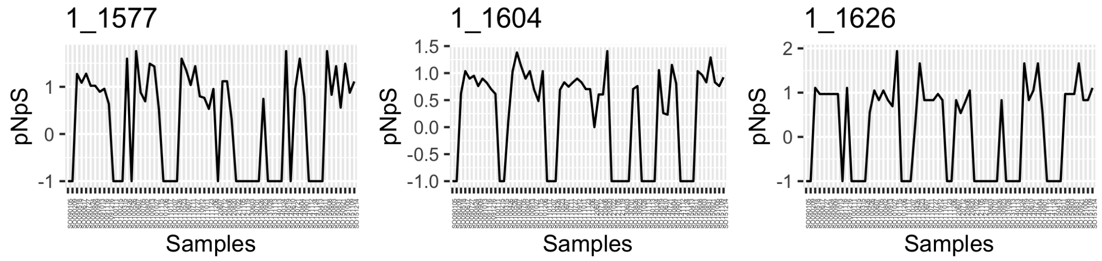


*Pontimonas* MAG (BL_pooled_bin.full.596) in SOLA.

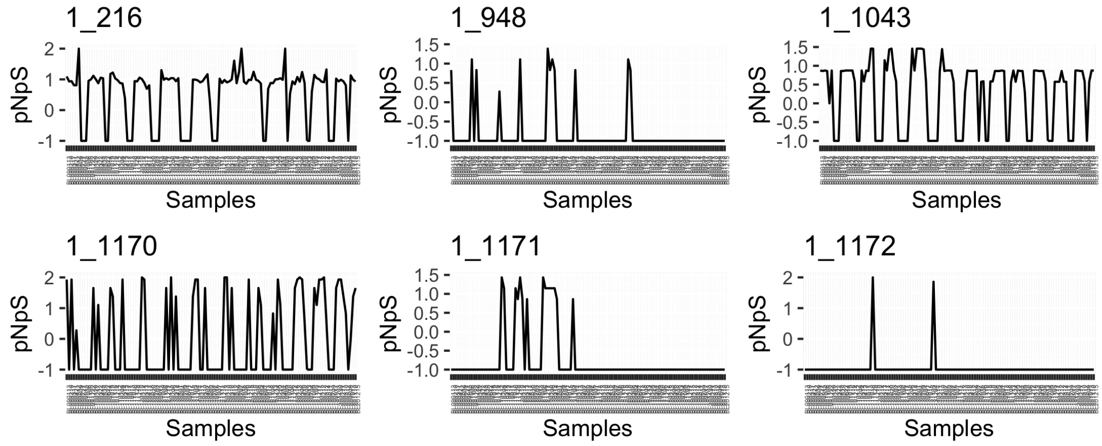


MED-G09 MAG (BL_pooled_bin.full.761) in BBMO.

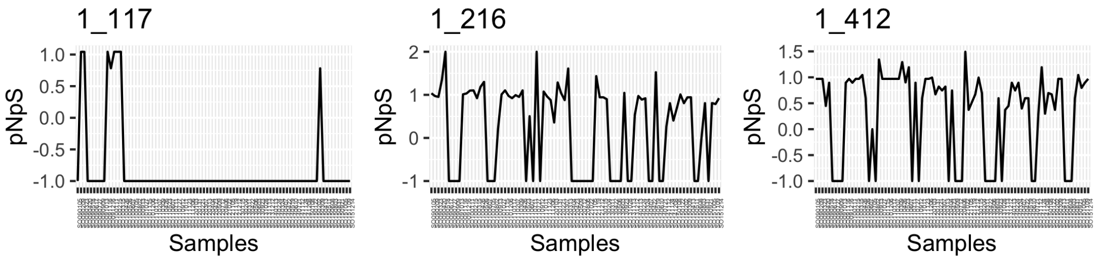


MED-G09 MAG (BL_pooled_bin.full.761) in SOLA.

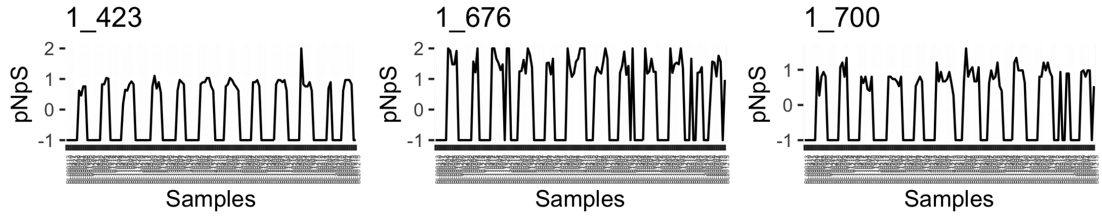


HIMB114 (SAR11) MAG (s1032.ctg001132l_BL_0908sc) in BBMO.

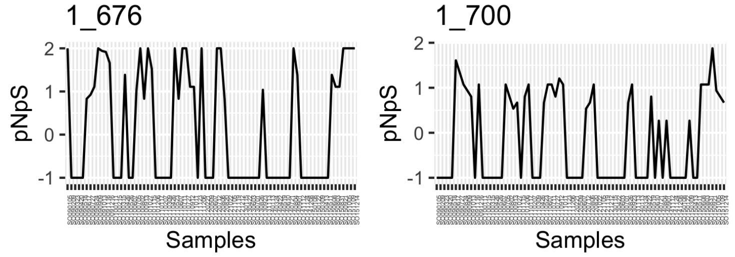


HIMB114 (SAR11) MAG (s1032.ctg001132l_BL_0908sc) in SOLA.

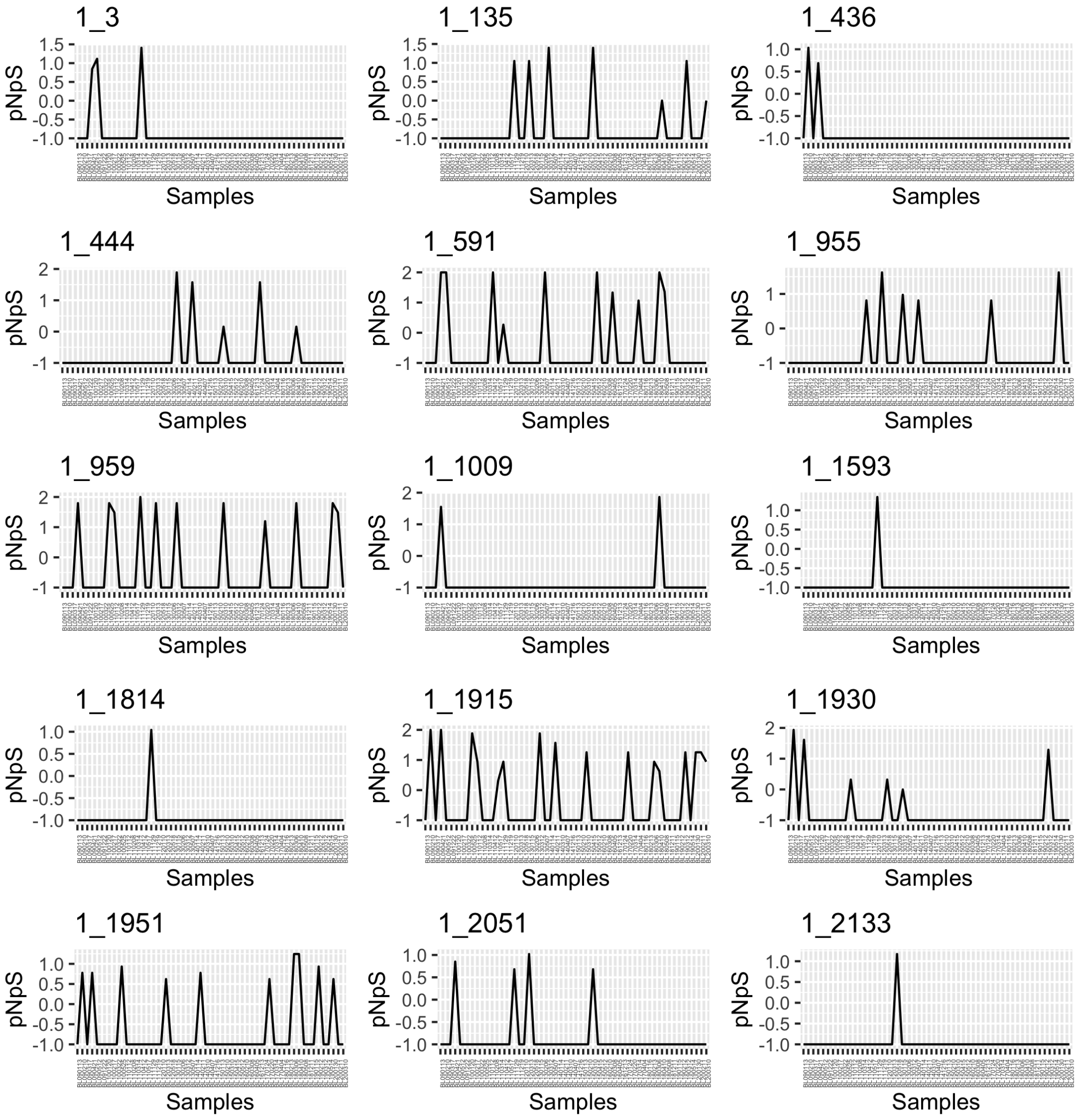


HTCC2207 (Porticoccaceae) MAG (s111.ctg000126c_BL_0902sc) in BBMO.

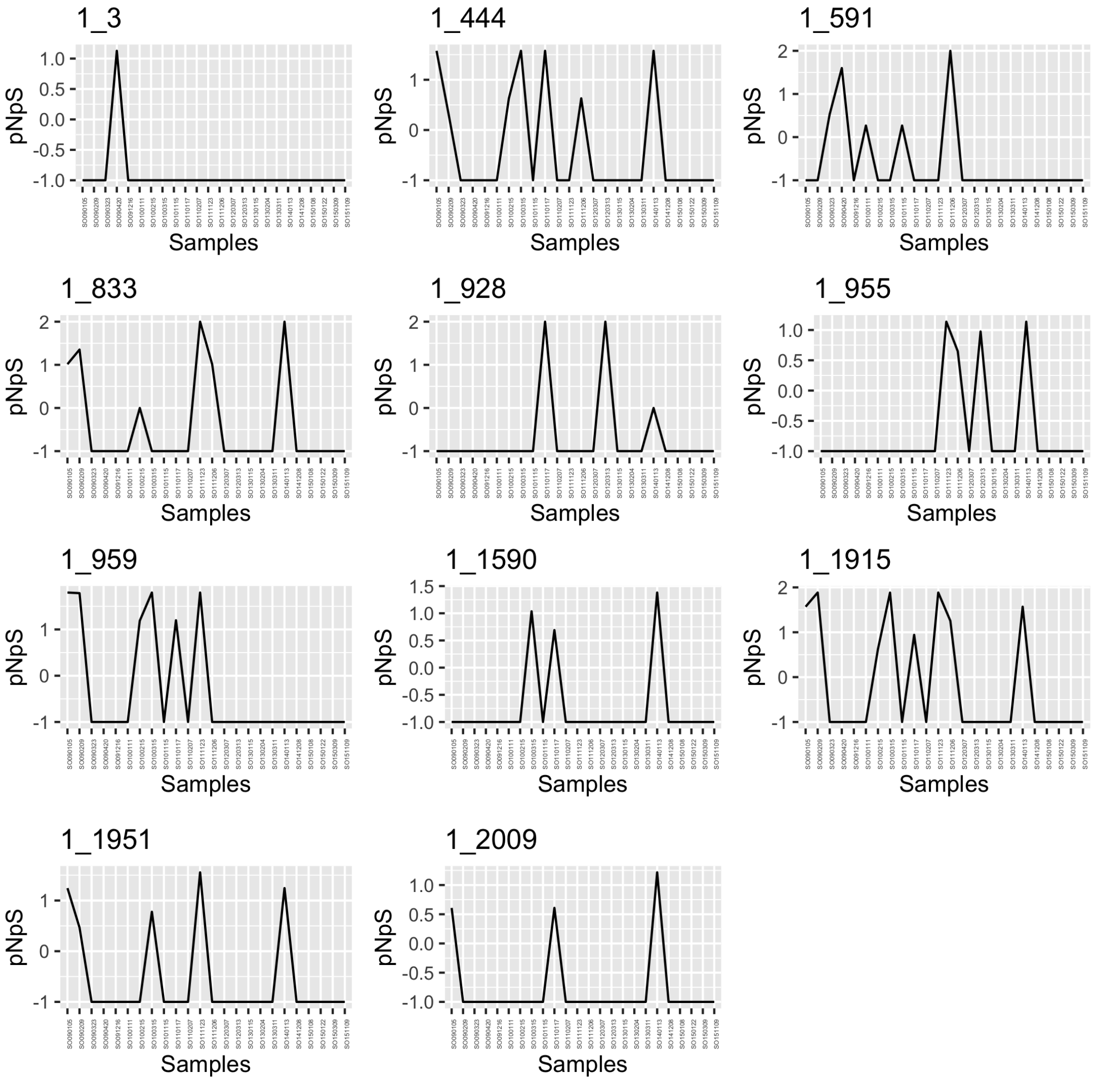


HTCC2207 (Porticoccaceae) MAG (s111.ctg000126c_BL_0902sc) in SOLA.

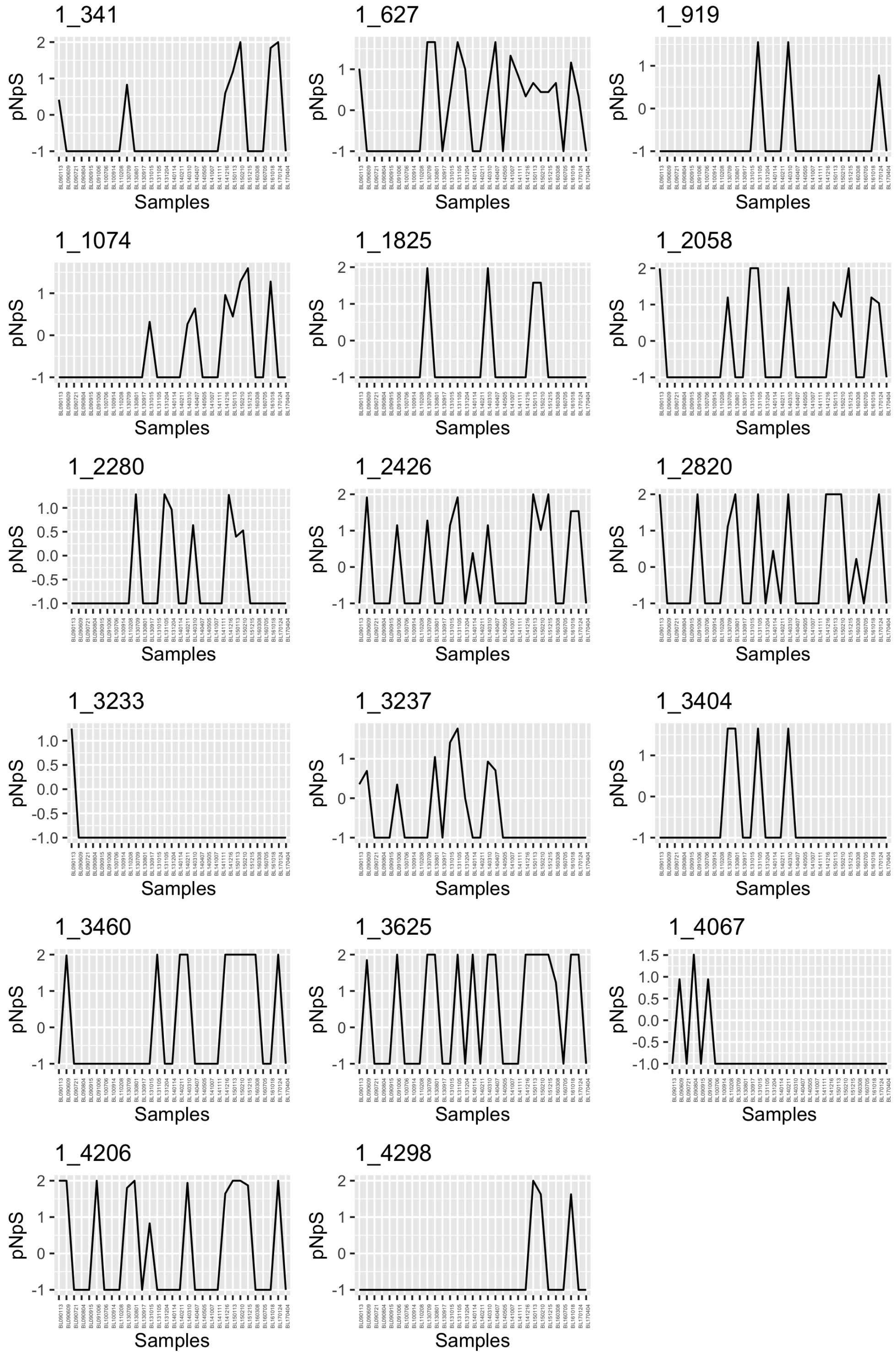


*Nocardioides* MAG (s131.ctg000151c_BL_0908sc) in BBMO.

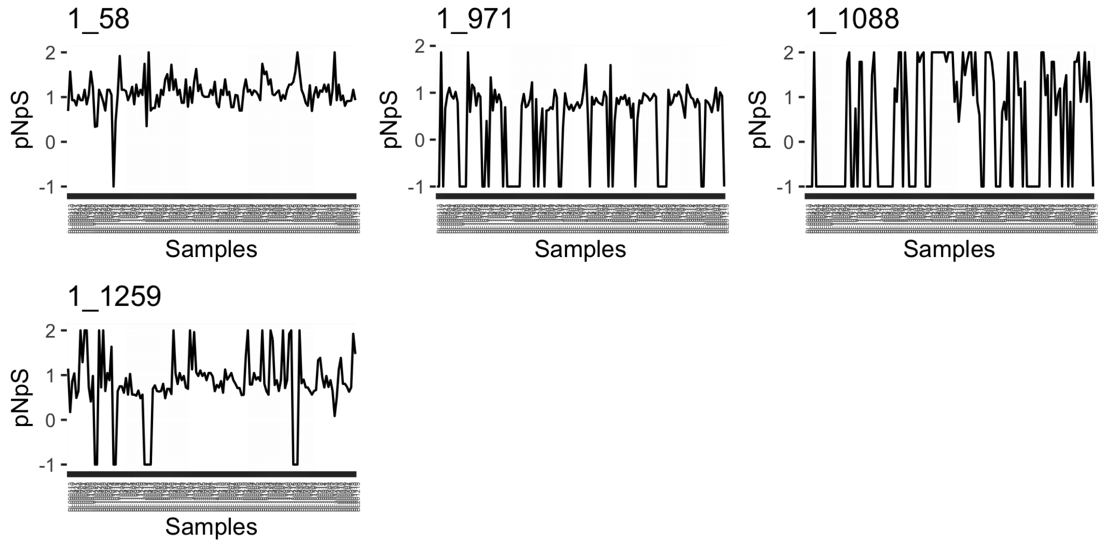


SCGC-AAA076-P13 (SAR86) MAG (s215.ctg000240l_BL_0908sc) in BBMO.

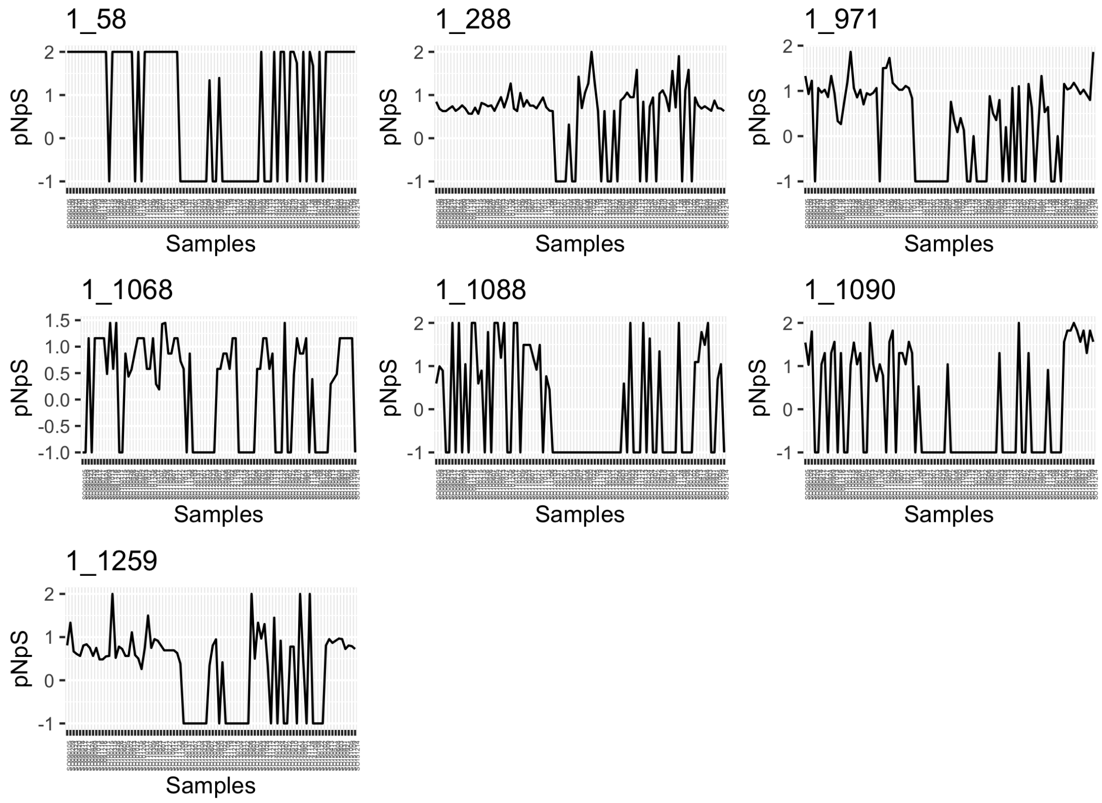


SCGC-AAA076-P13 (SAR86) MAG (s215.ctg000240l_BL_0908sc) in SOLA.

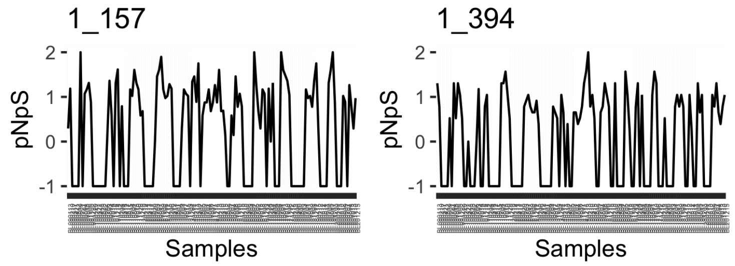


*Actinomarina* MAG (s2266.ctg002461l_BL_0908sc) in BBMO.

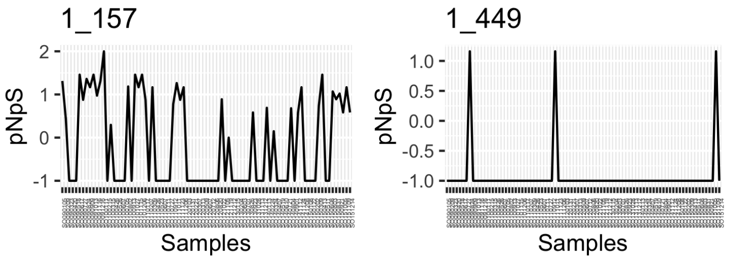


*Actinomarina* MAG (s2266.ctg002461l_BL_0908sc) in SOLA.

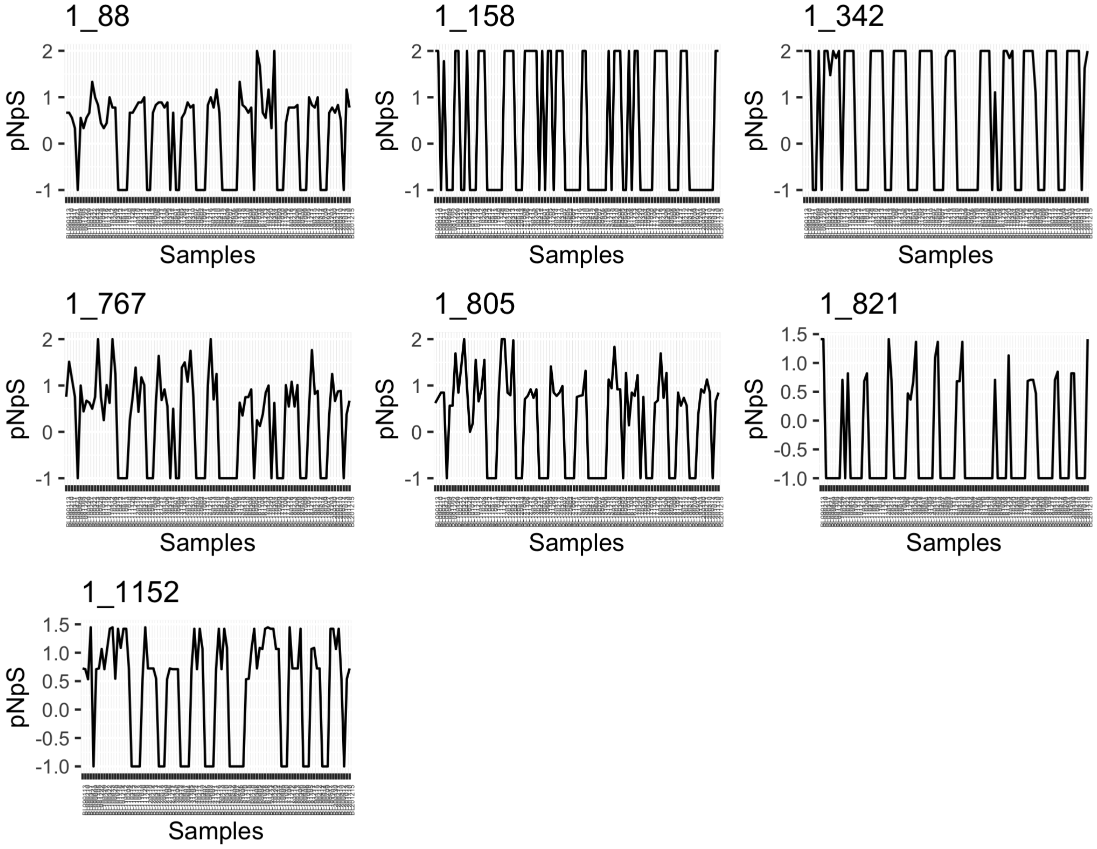


MGIIb-O2 (Archaea) MAG (s23.ctg000345l_BL_0902sc) in BBMO.

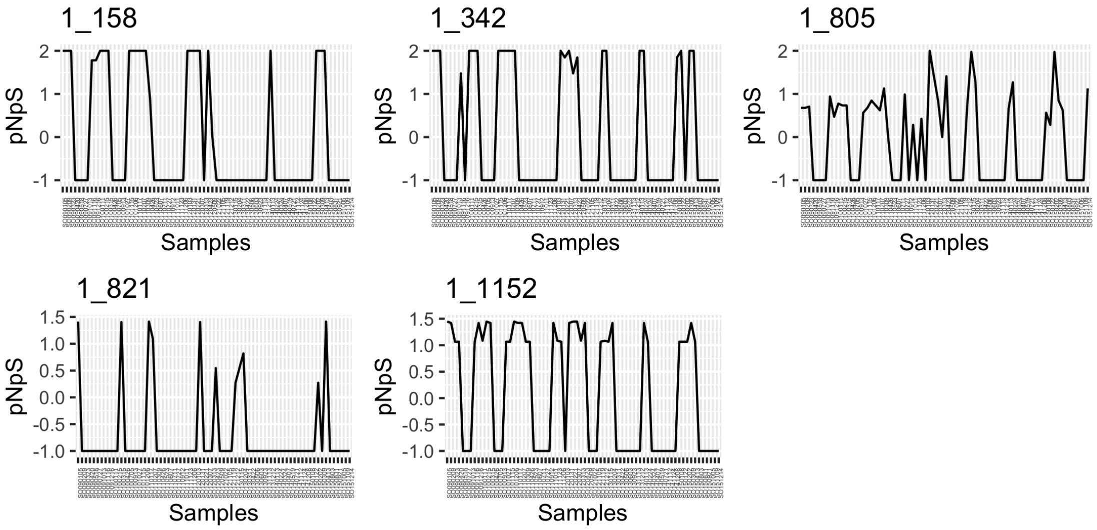


MGIIb-O2 (Archaea) MAG (s23.ctg000345l_BL_0902sc) in SOLA.

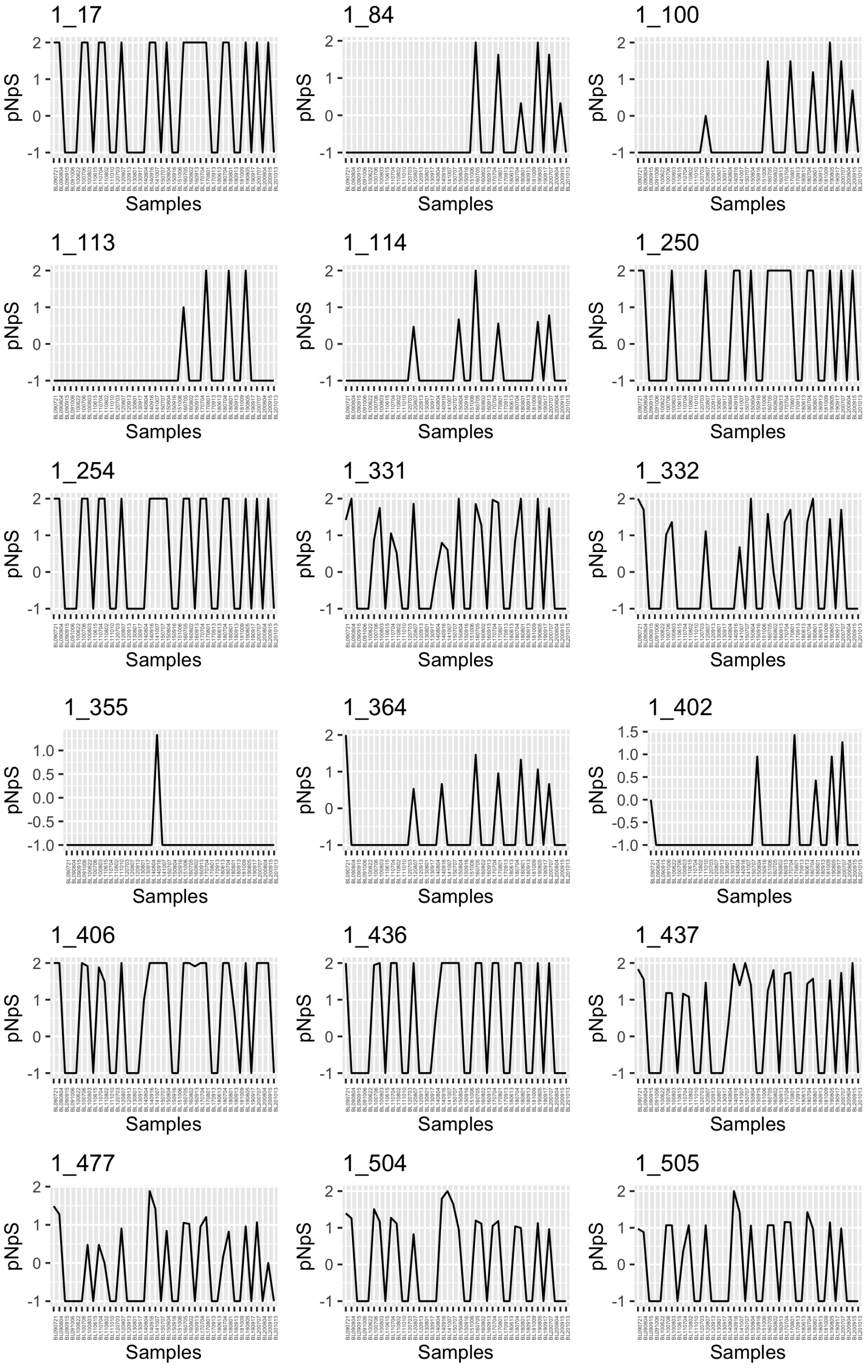


UBA7445 (Opitutales) MAG (s249.ctg000277I_BL_0908sc) in BBMO.

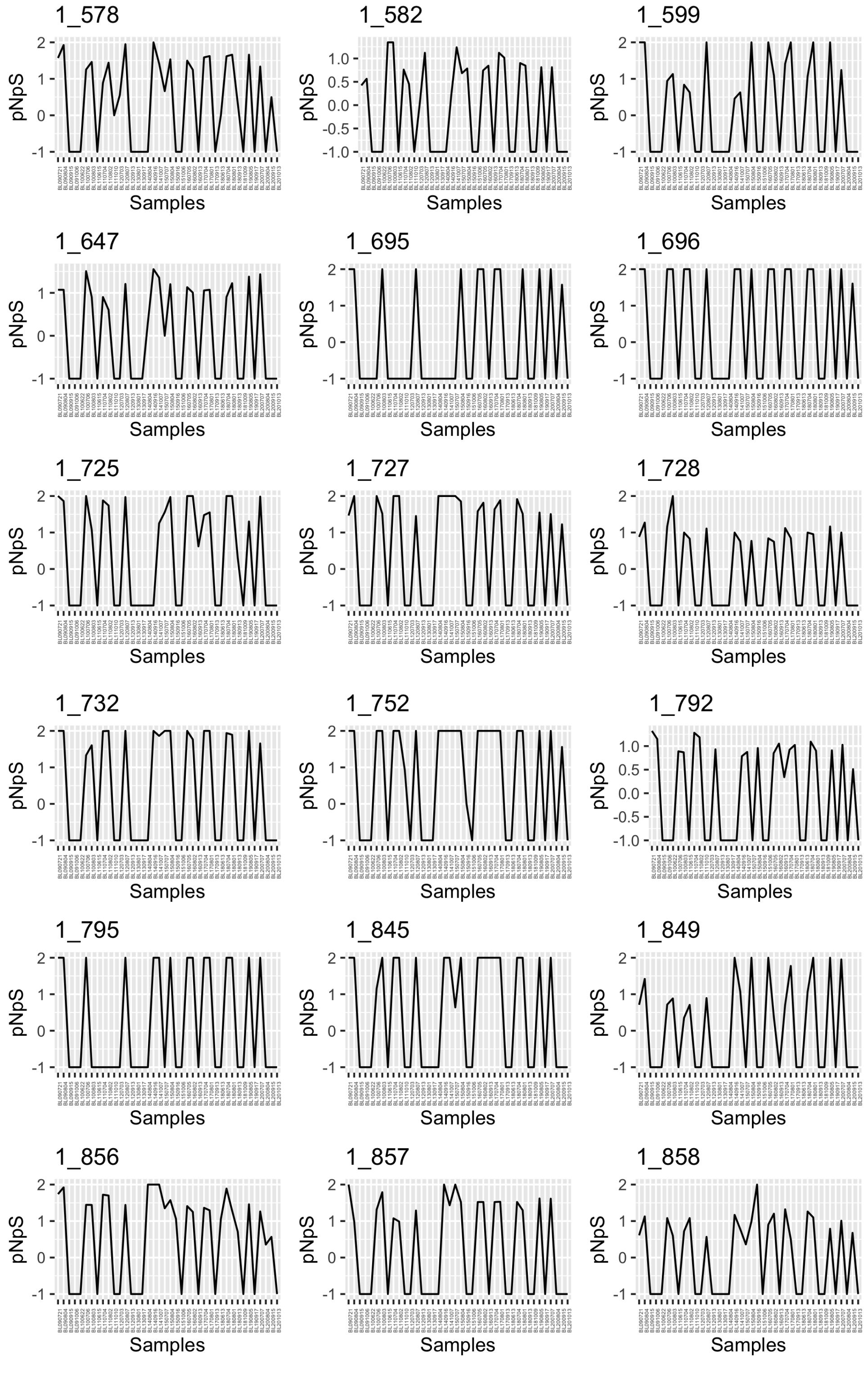


UBA7445 (Opitutales) MAG (s249.ctg000277l_BL_0908sc) in BBMO.

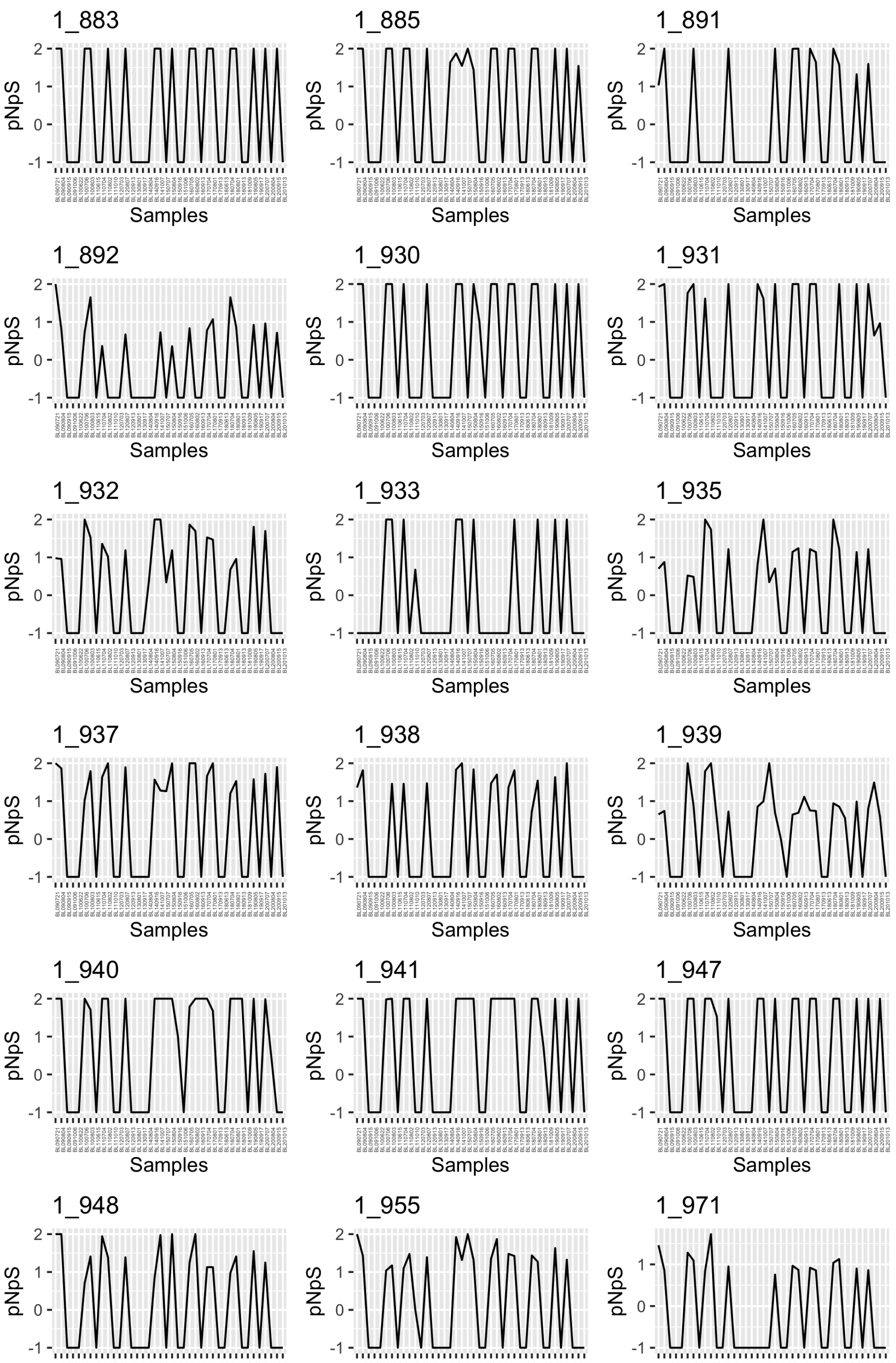


UBA7445 (Opitutales) MAG (s249.ctg000277I_BL_0908sc) in BBMO.

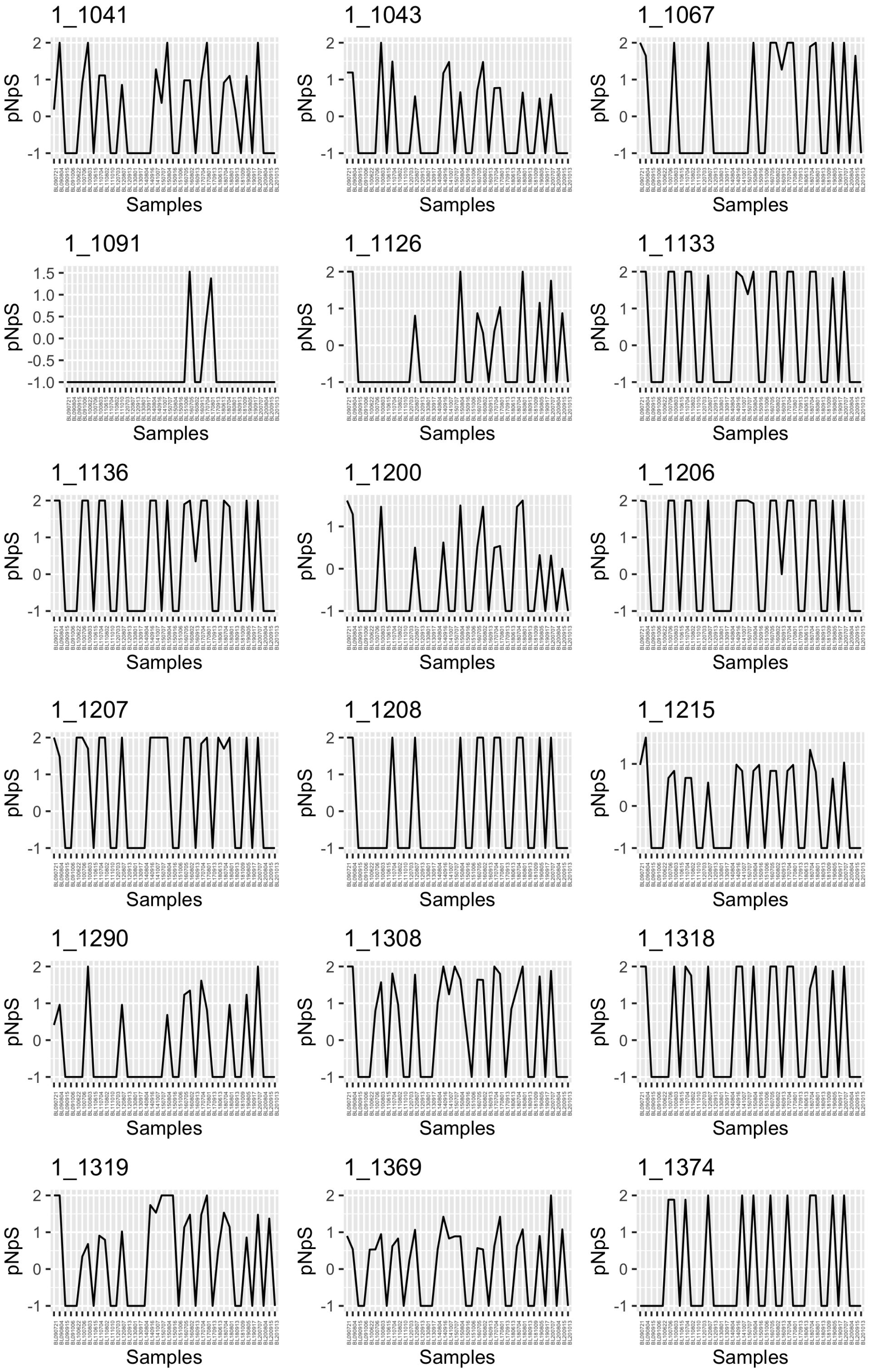


UBA7445 (Opitutales) MAG (s249.ctg000277I_BL_0908sc) in BBMO.

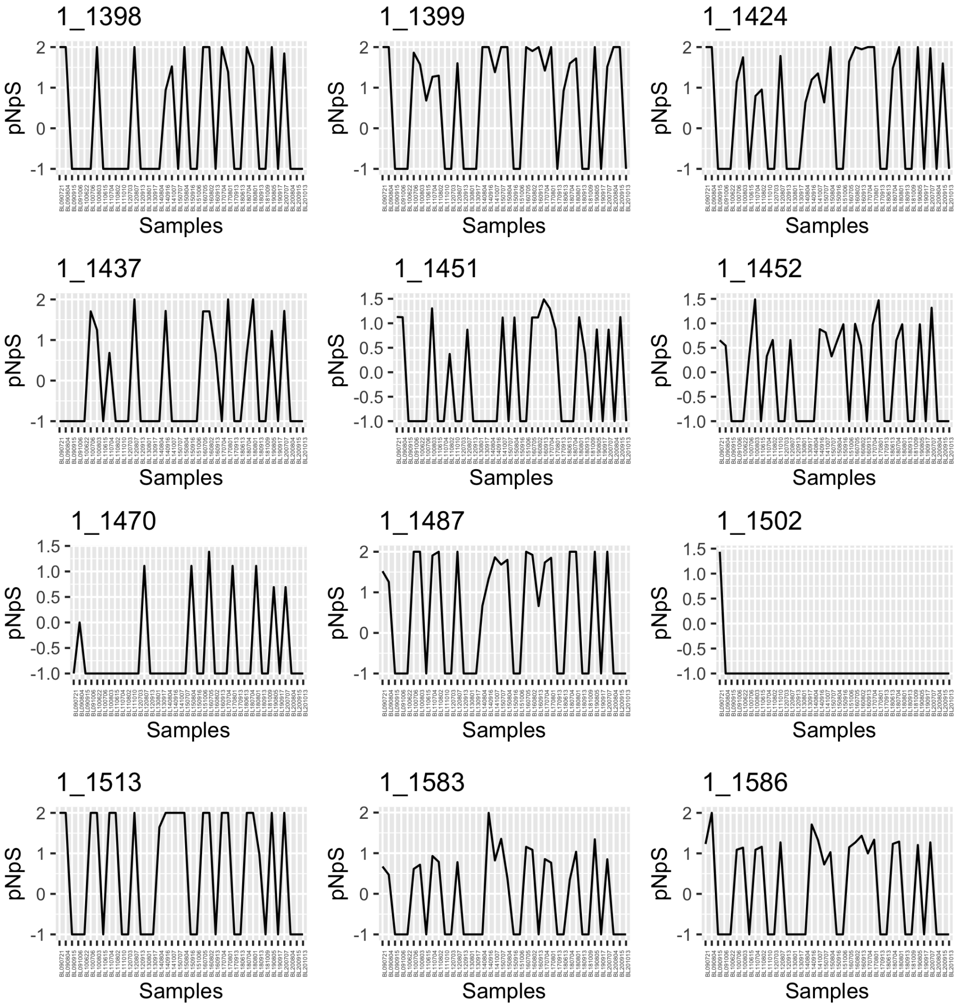


UBA7445 (Opitutales) MAG (s249.ctg000277I_BL_0908sc) in BBMO.

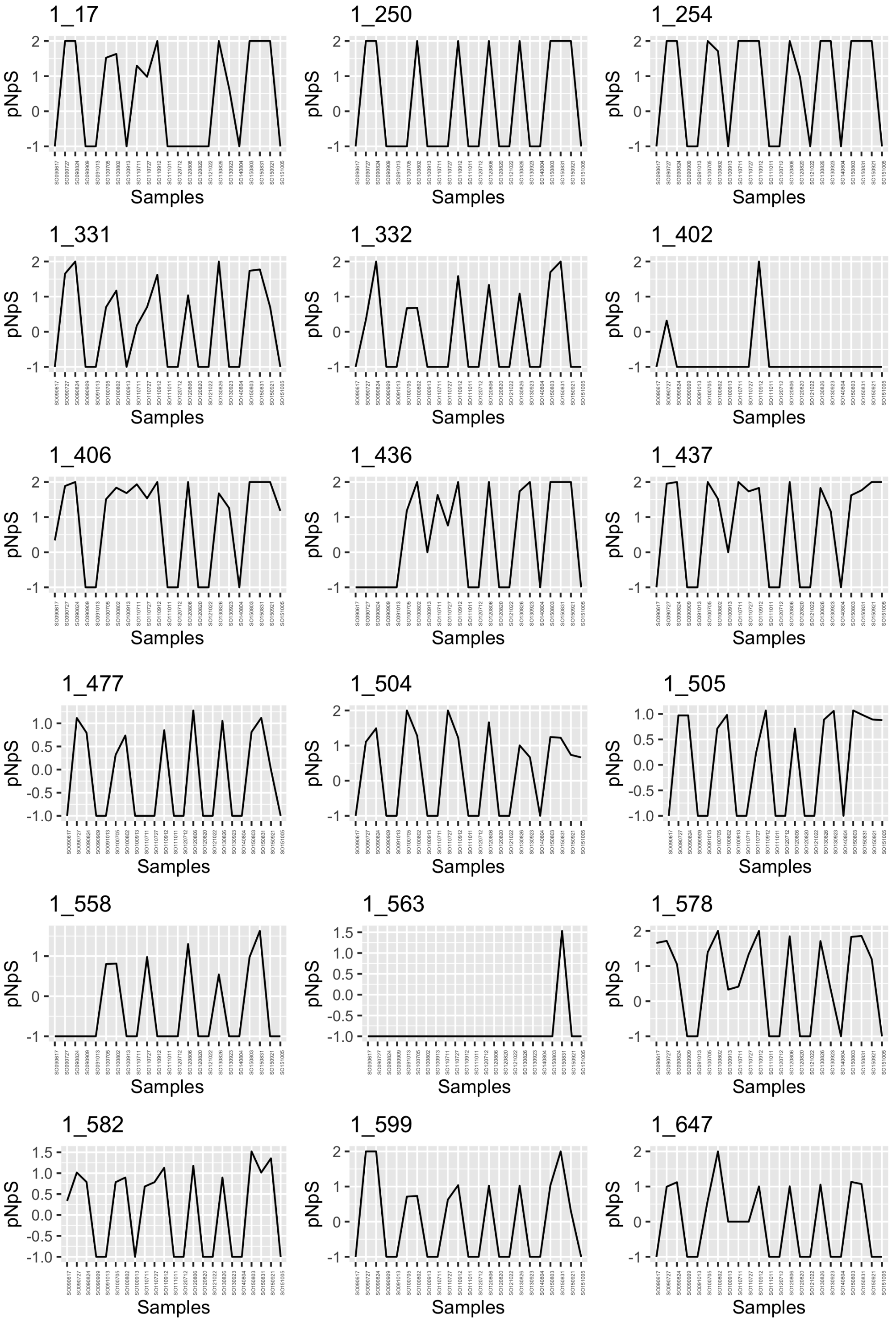


UBA7445 (Opitutales) MAG (s249.ctg000277l_BL_0908sc) in SOLA.

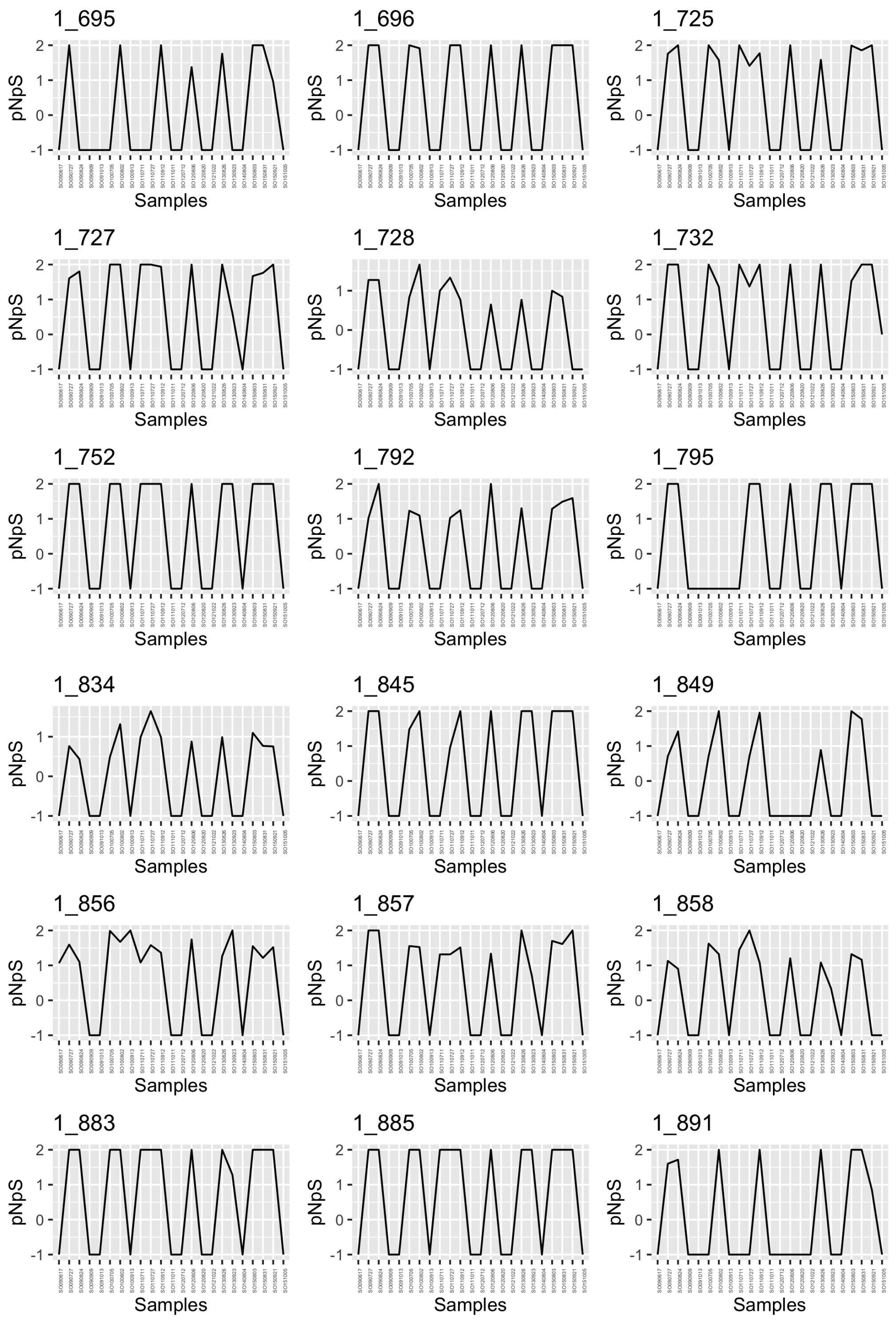


UBA7445 (Opitutales) MAG (s249.ctg000277I_BL_0908sc) in SOLA.

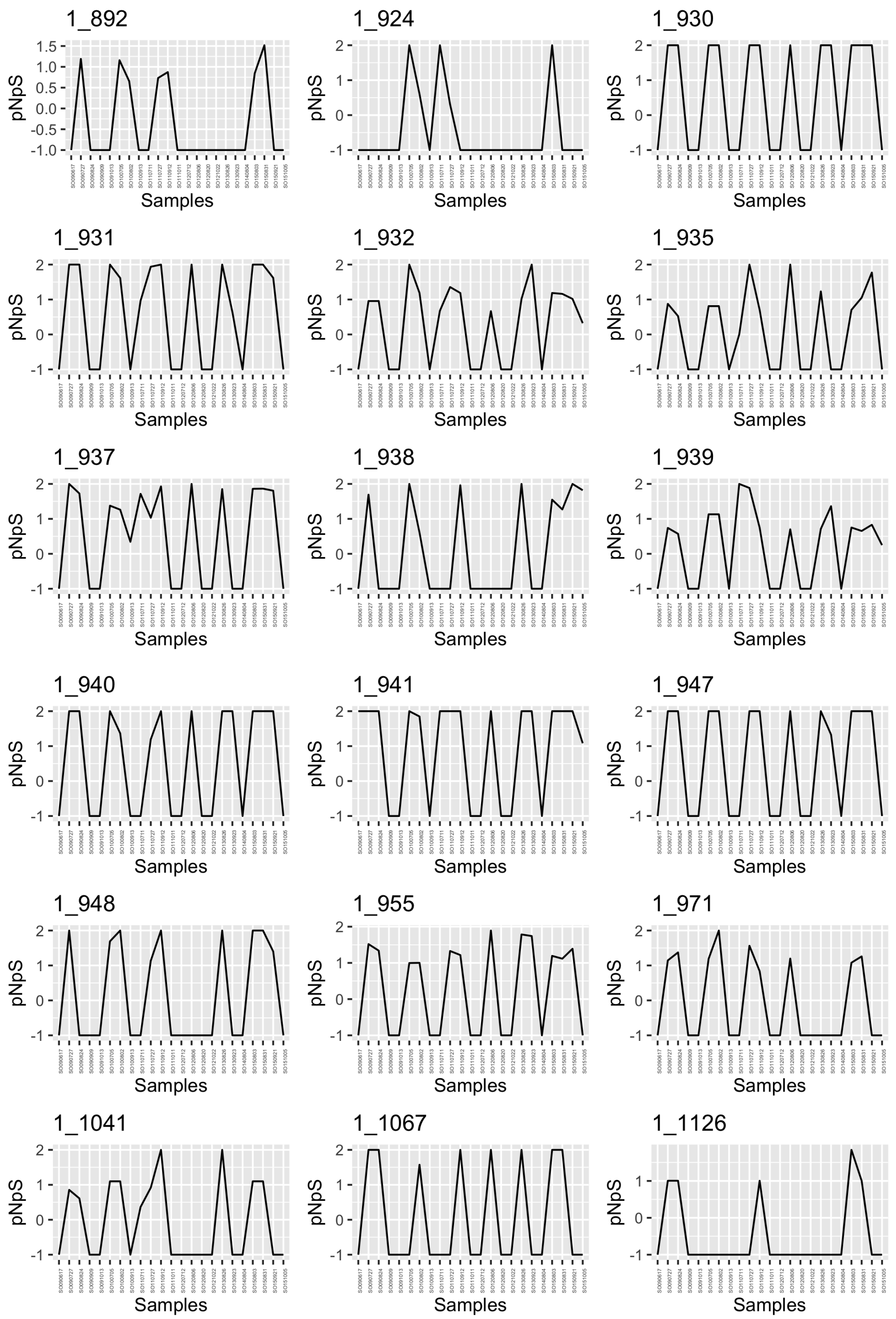


UBA7445 (Opitutales) MAG (s249.ctg000277l_BL_0908sc) in SOLA.

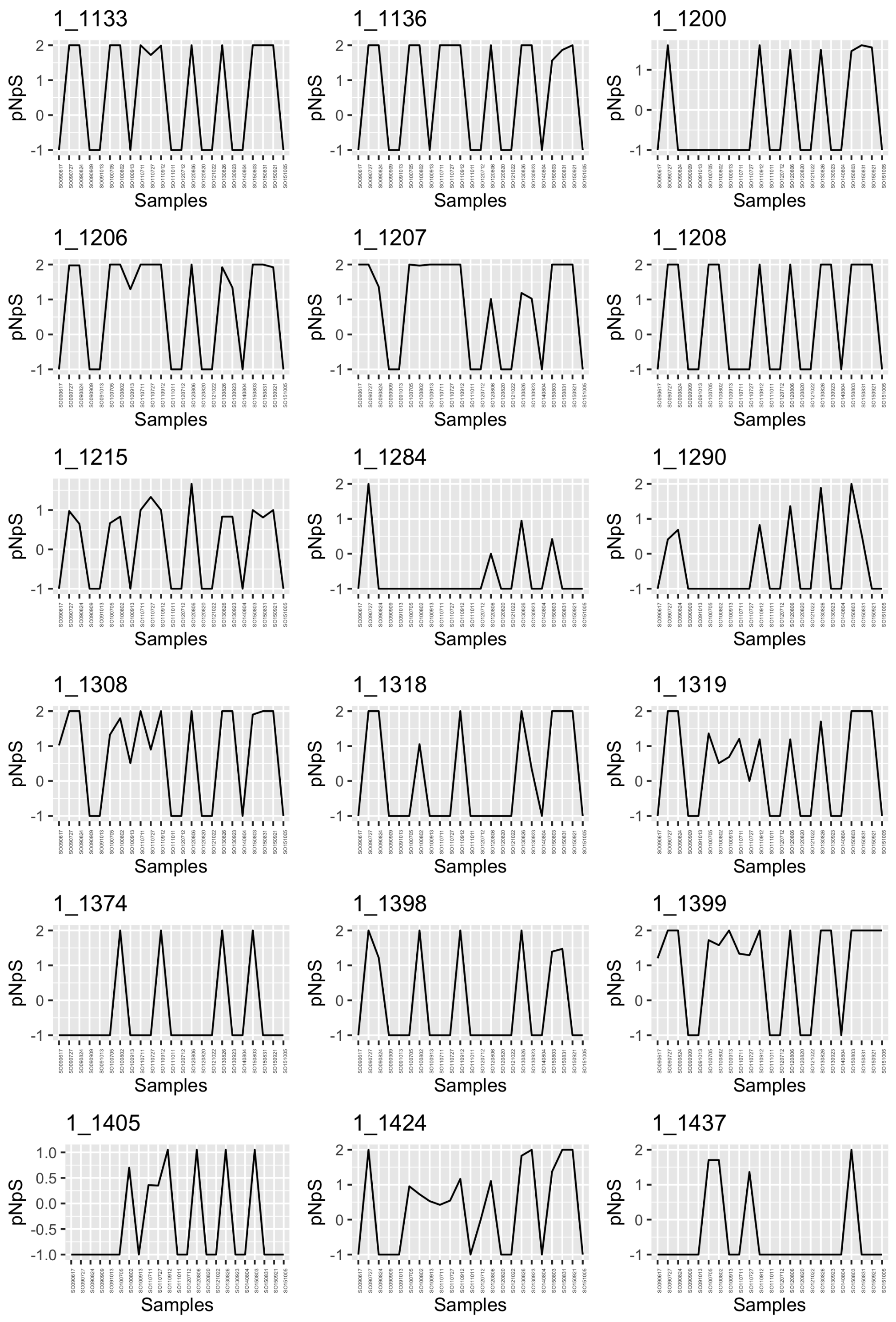


UBA7445 (Opitutales) MAG (s249.ctg000277l_BL_0908sc) in SOLA.

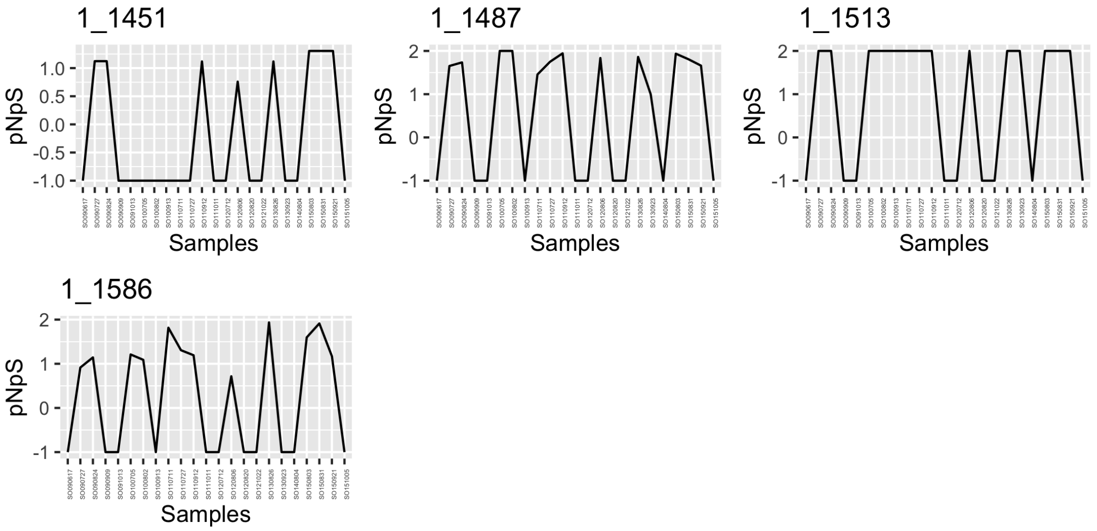


UBA7445 (Opitutales) MAG (s249.ctg000277I_BL_0908sc) in SOLA.

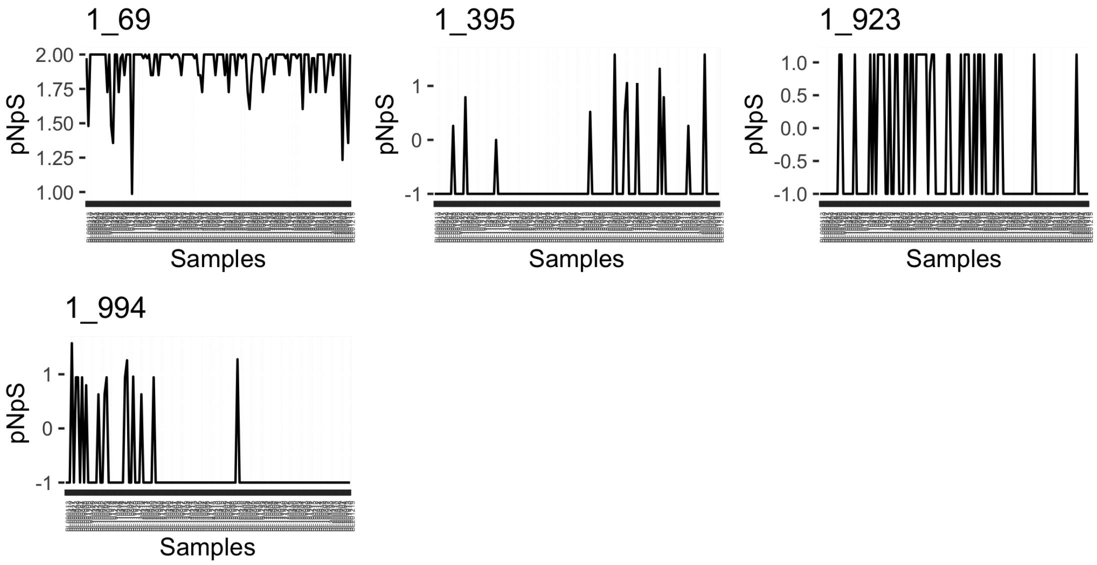


*Pelagibacter* MAG (s266.ctg000297l_BL_0908sc) in BBMO.

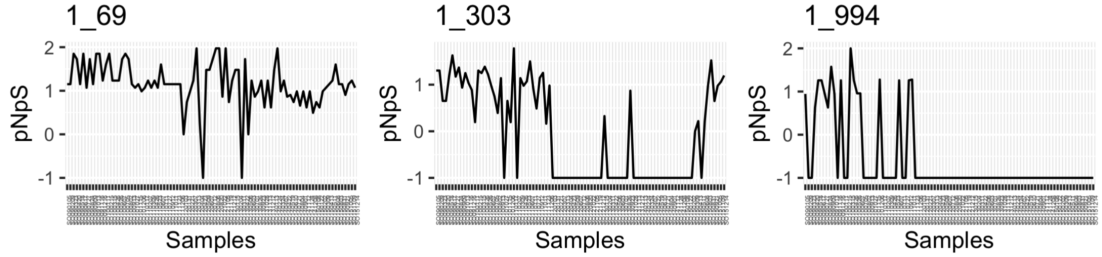


*Pelagibacter* MAG (s266.ctg000297l_BL_0908sc) in SOLA

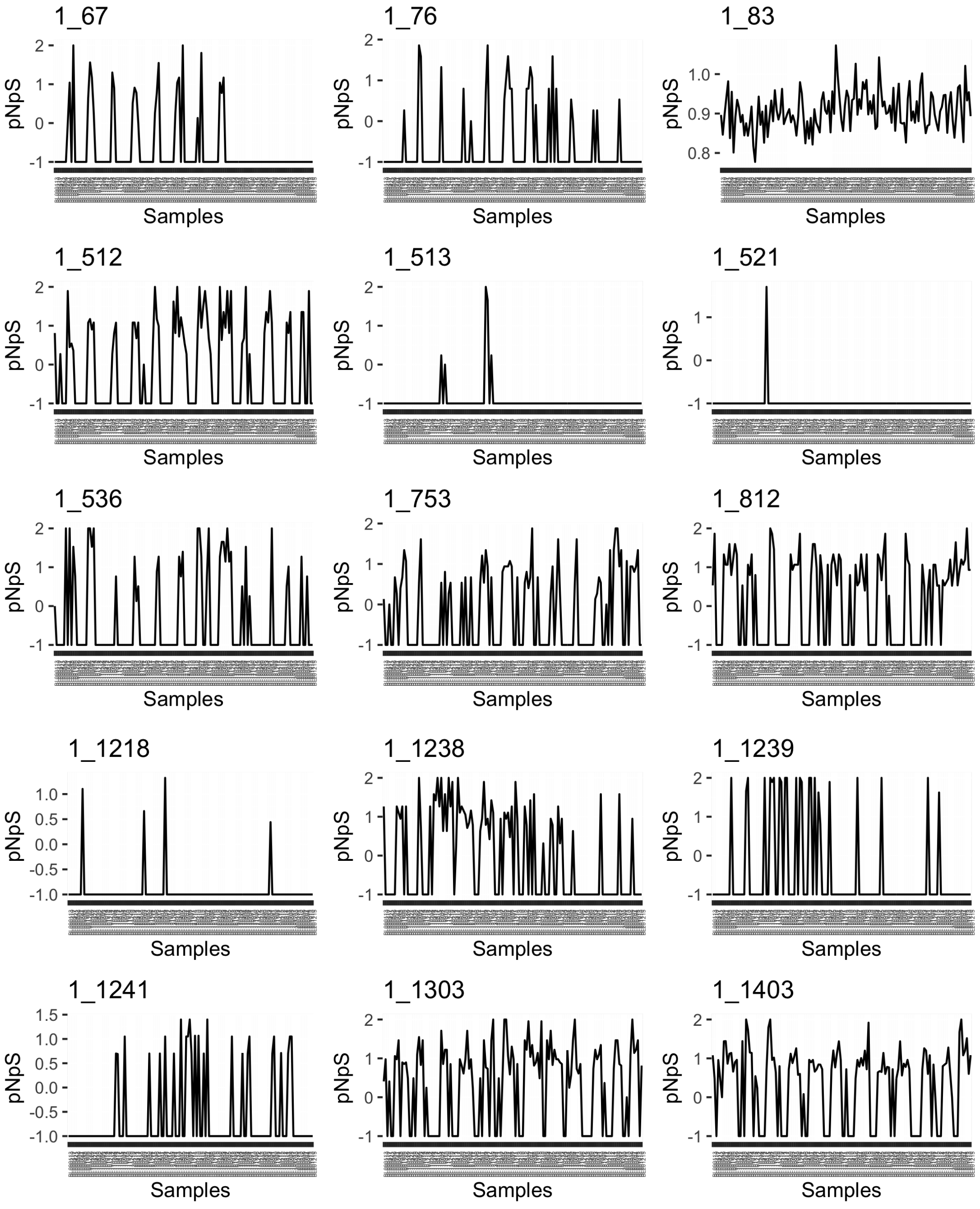


TMED13 MAG (s325.ctg000366c_BL_0908sc) in BBMO.

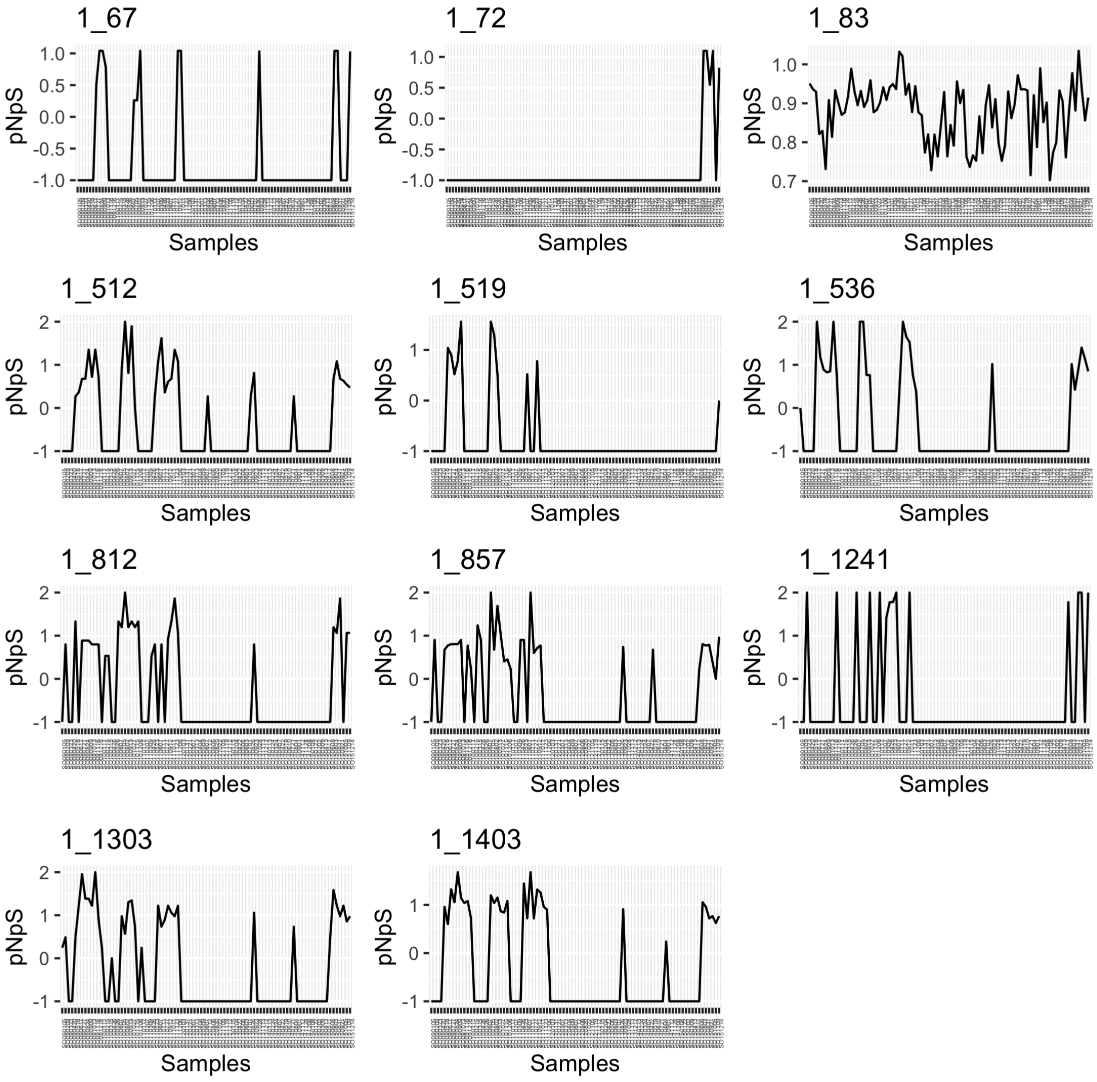


TMED13 MAG (s325.ctg000366c_BL_0908sc) in SOLA.

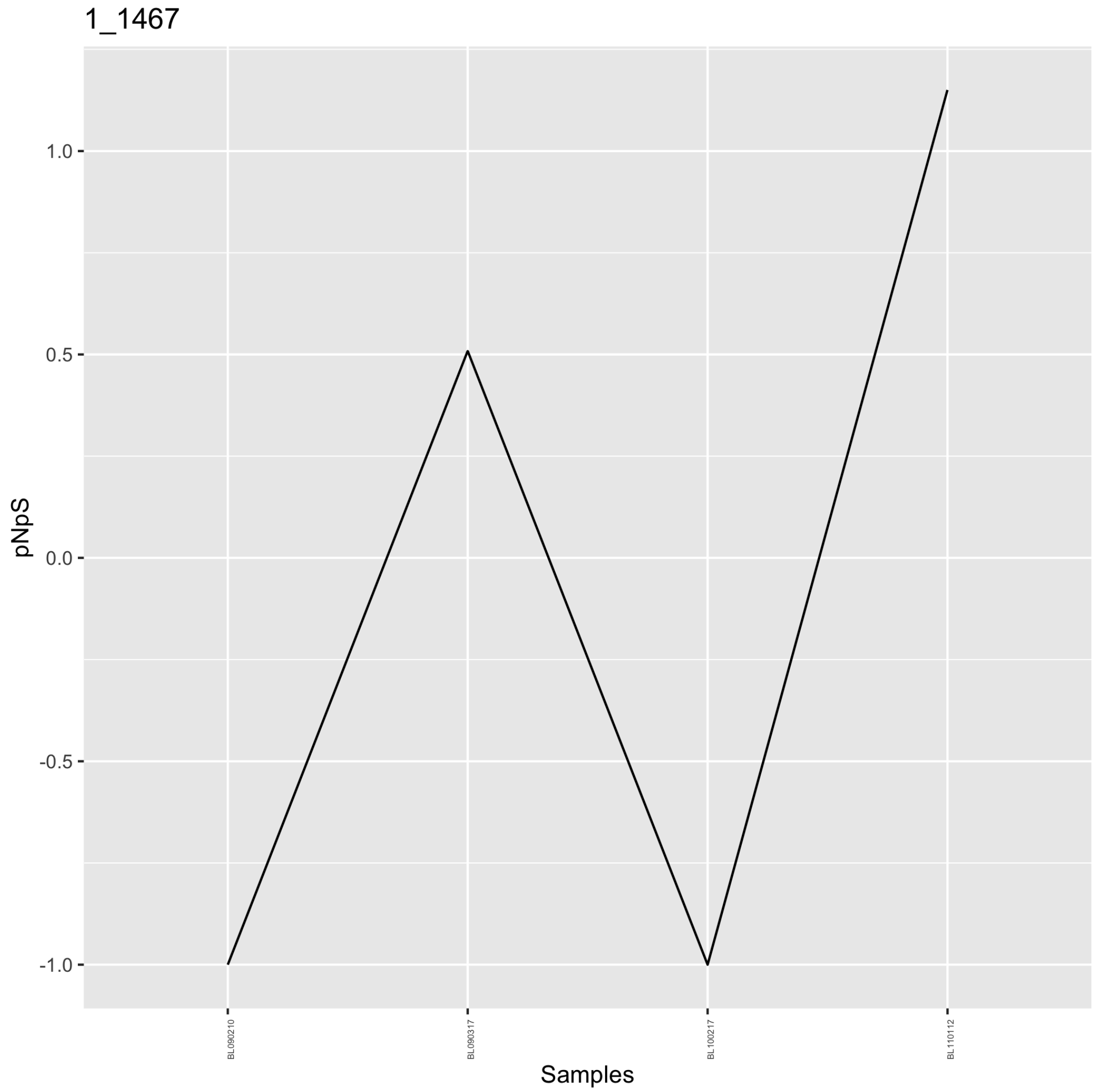


*Amylibacter* MAG (s35.ctg000041c_BL_0902sc) in BBMO.

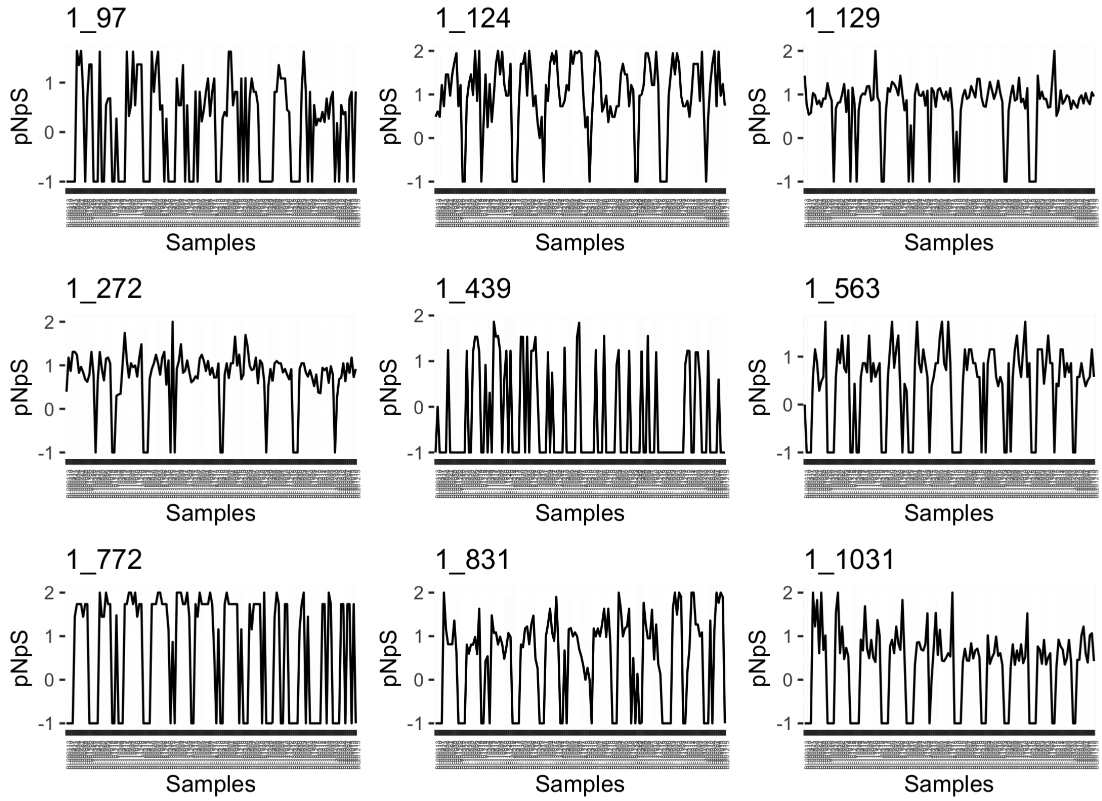


SCGC-AAA076-P13 (SAR86) MAG (s388.ctg000435l_BL_0908sc) in BBMO.

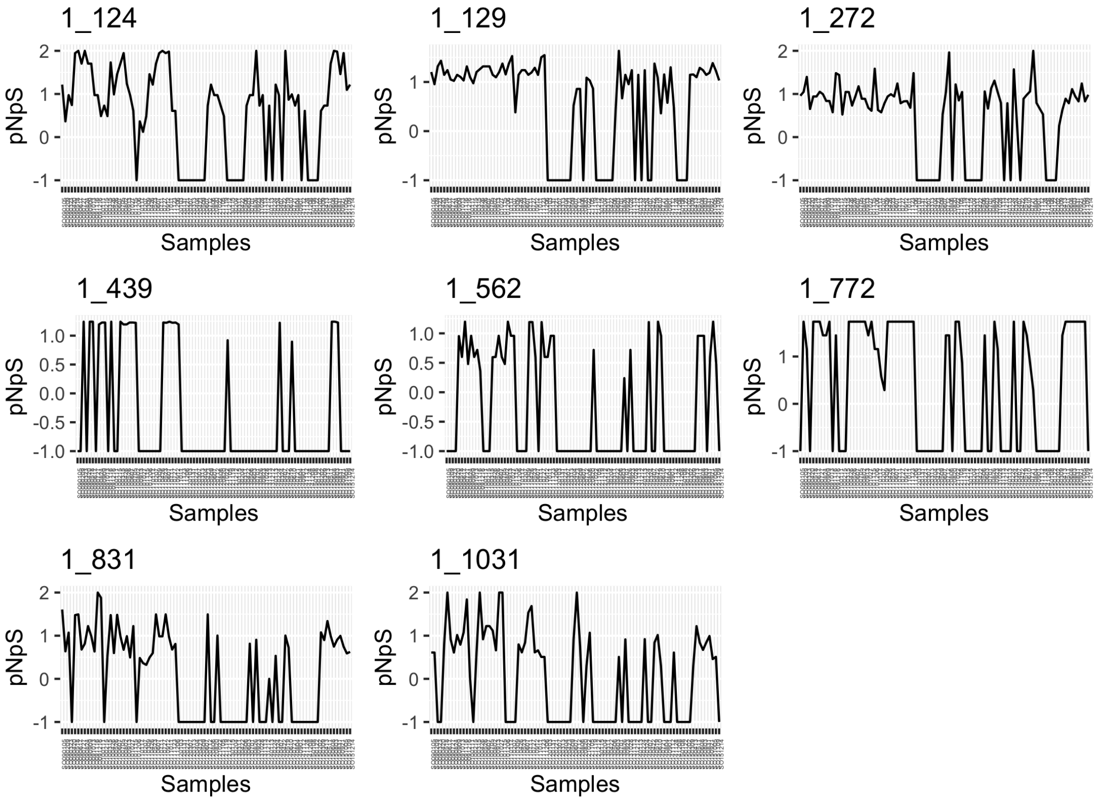


SCGC-AAA076-P13 (SAR86) MAG (s388.ctg000435l_BL_0908sc) in SOLA.

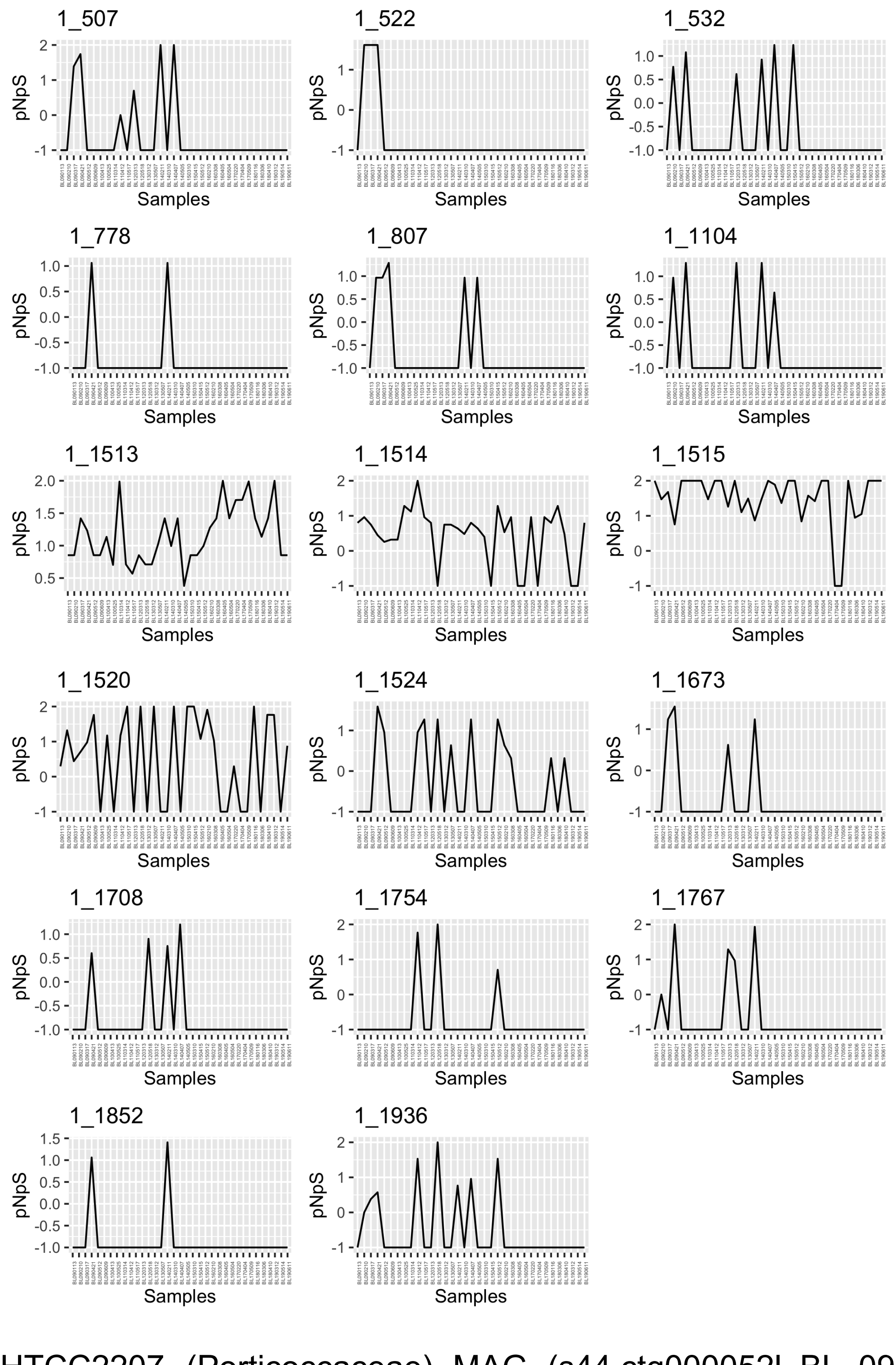


HTCC2207 (Porticoccaceae) MAG (s44.ctg000052l_BL_0902sc) in BBMO.

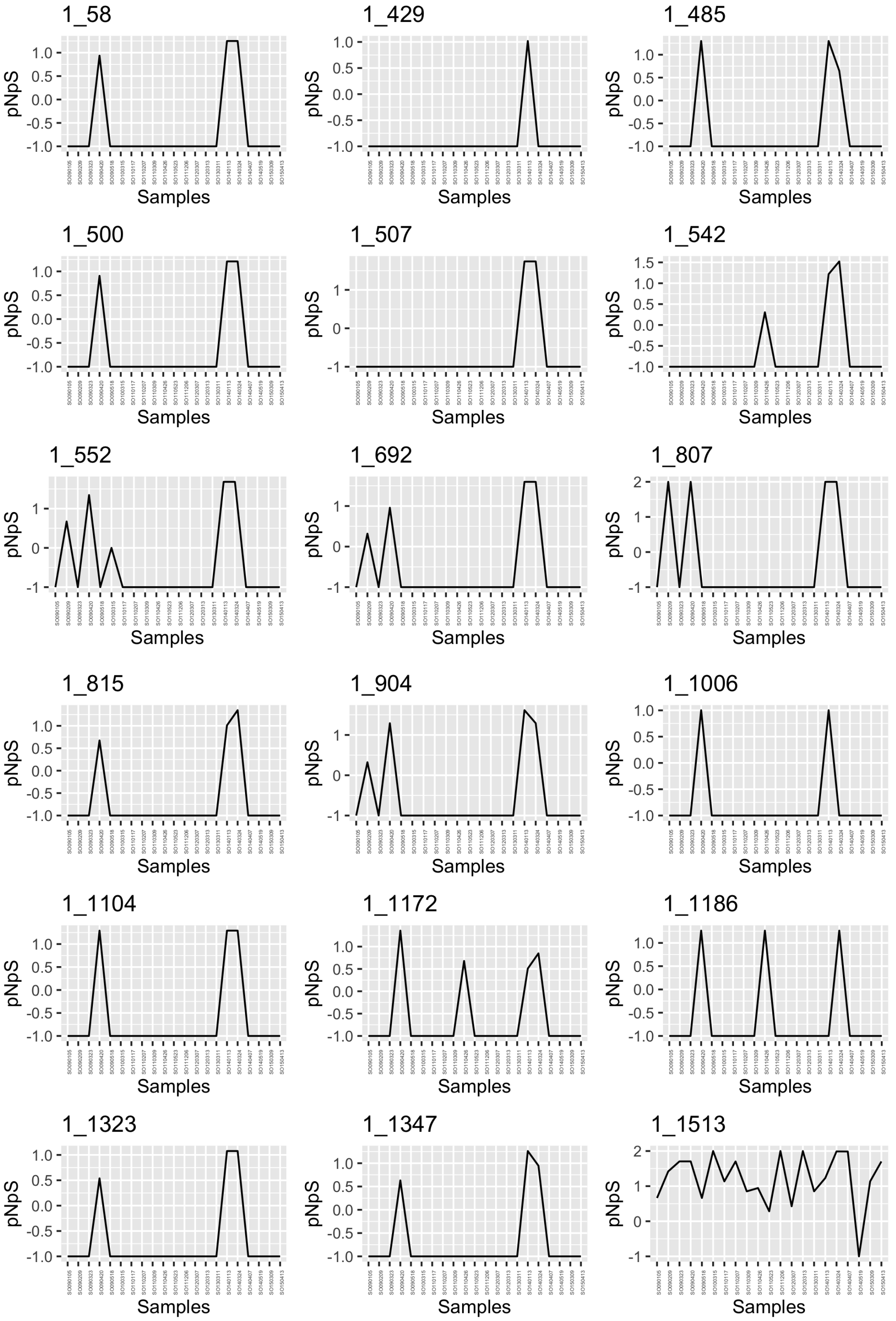


HTCC2207 (Porticoccaceae) MAG (s44.ctg000052l_BL_0902sc) in SOLA.

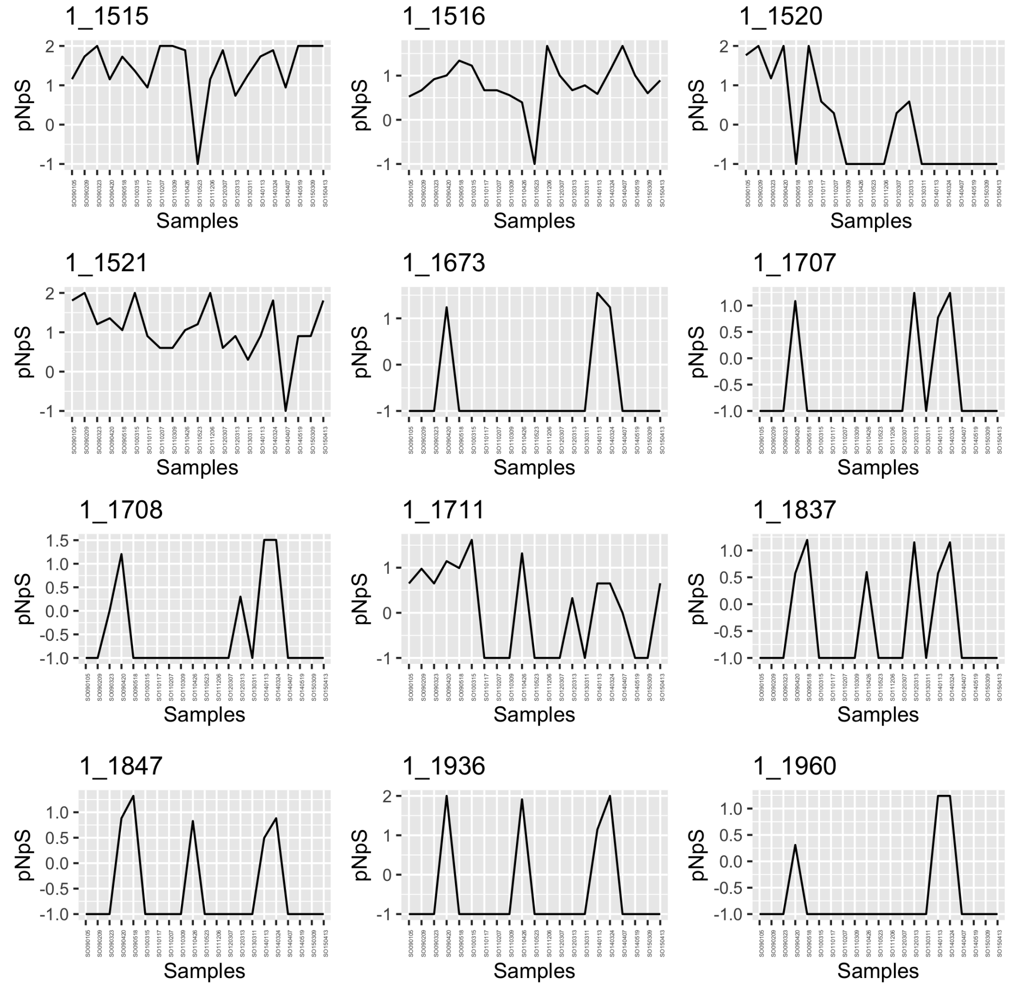


HTCC2207 (Porticoccaceae) MAG (s44.ctg000052l_BL_0902sc) in SOLA.

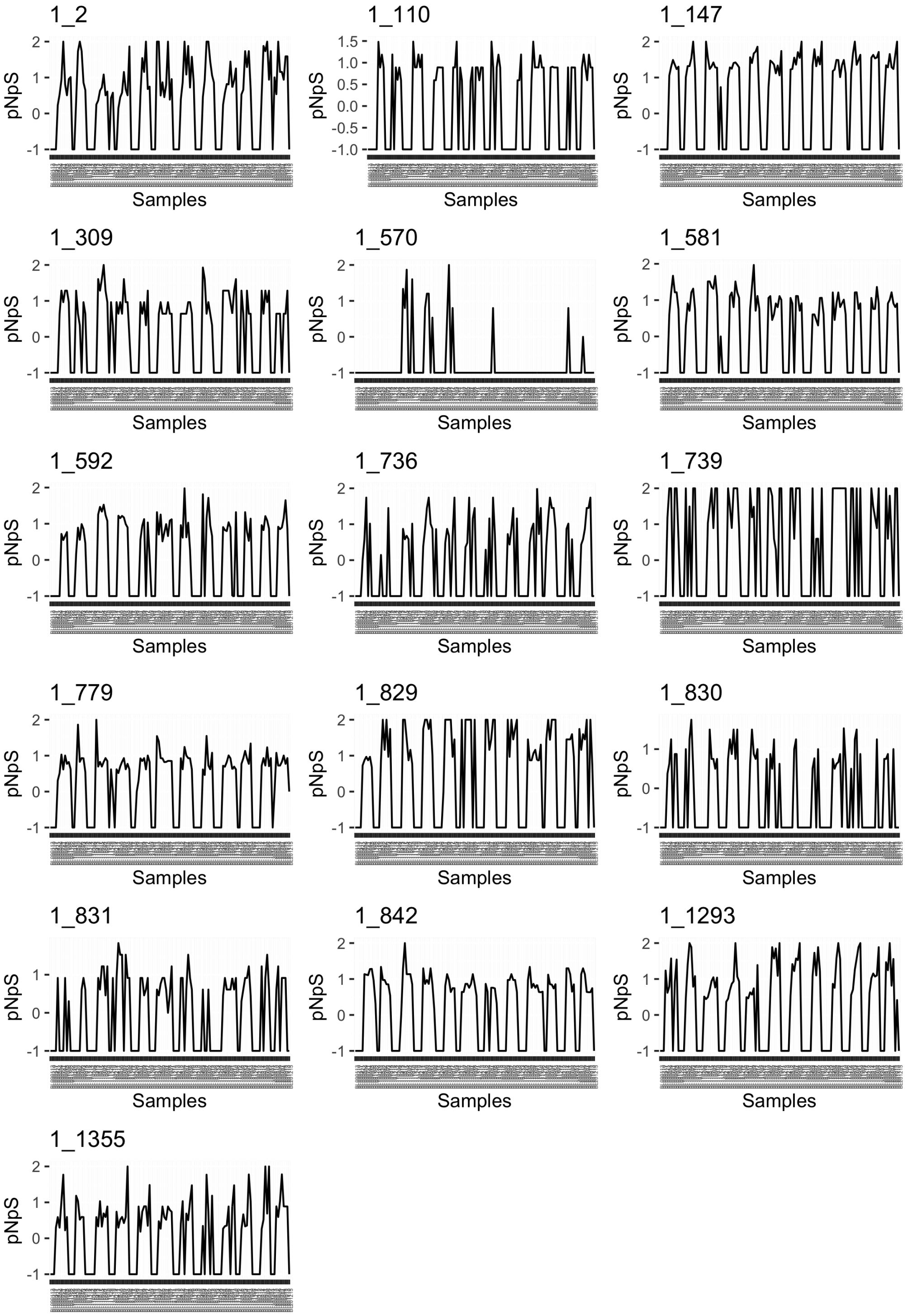


UBA7446 (Flavobacteriaceae) MAG (s7.ctg000447l_BL_0908sc) in BBMO.

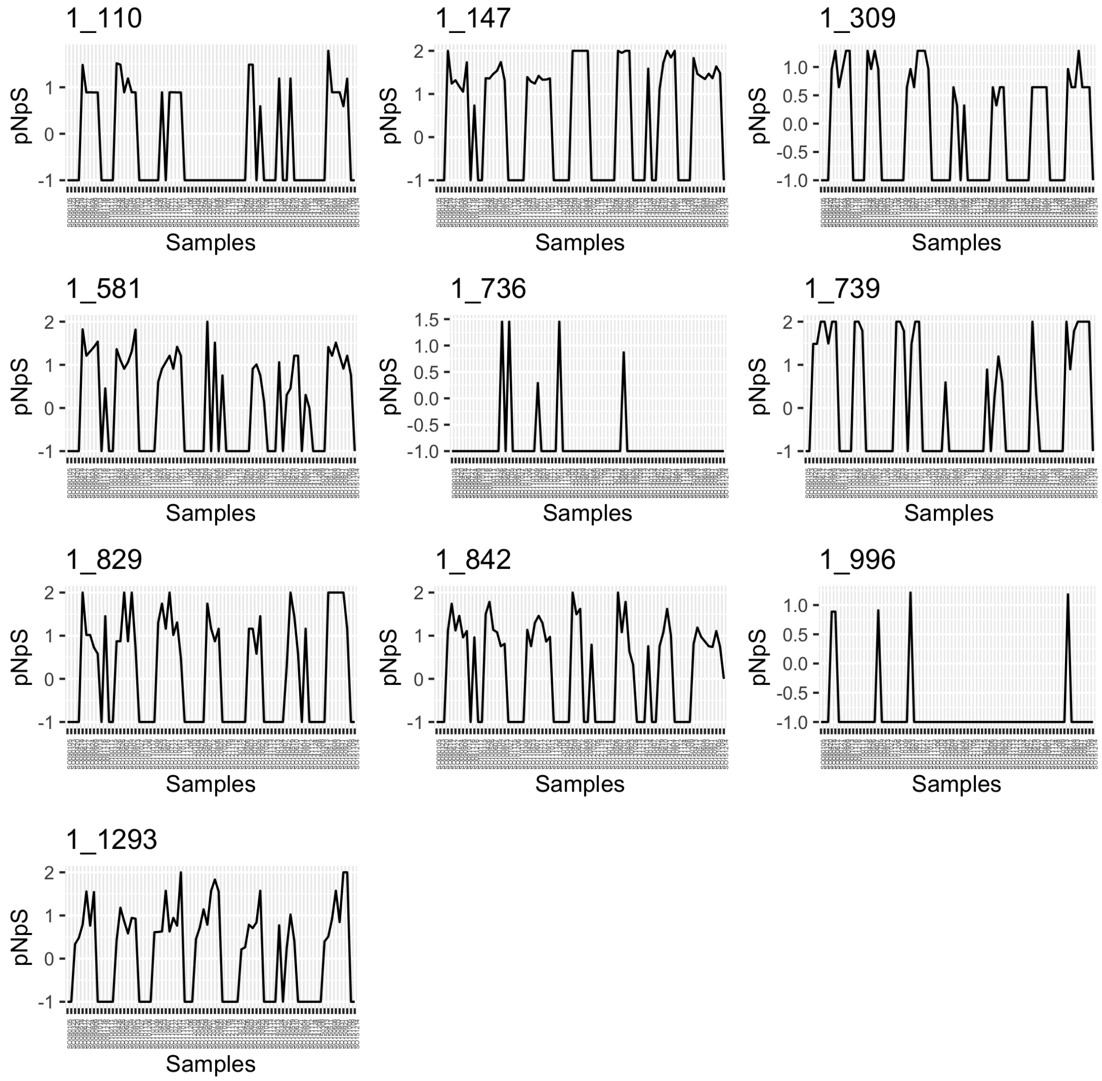


UBA7446 (Flavobacteriaceae) MAG (s7.ctg000447l_BL_0908sc) in SOLA.

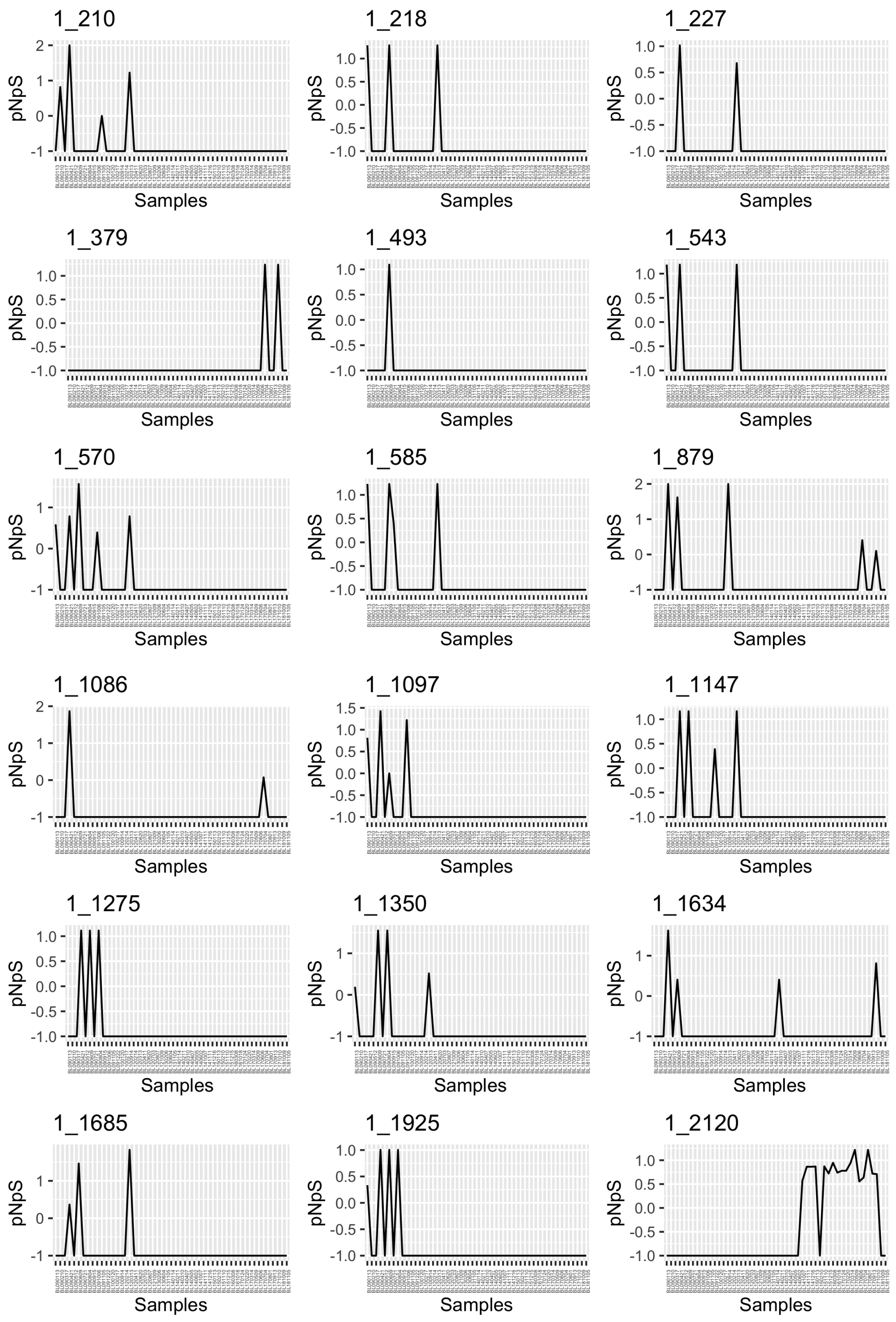


*Brevundimonas* MAG (s96.ctg000111c_BL_0908sc) in BBMO.

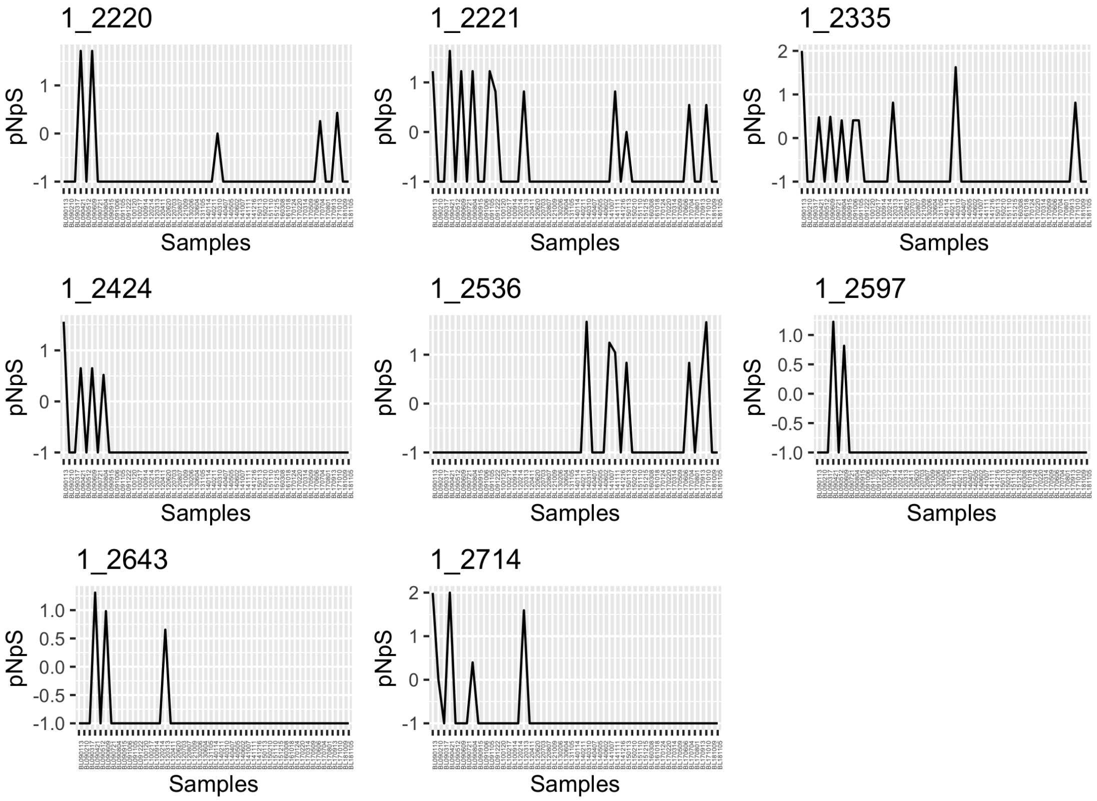


*Brevundimonas* MAG (s96.ctg000111c_BL_0908sc) in BBMO.
